# CellDART: Cell type inference by domain adaptation of single-cell and spatial transcriptomic data

**DOI:** 10.1101/2021.04.26.441459

**Authors:** Sungwoo Bae, Kwon Joong Na, Jaemoon Koh, Dong Soo Lee, Hongyoon Choi, Young Tae Kim

**Affiliations:** Department of Molecular Medicine and Biopharmaceutical Sciences, Graduate School of Convergence Science and Technology, Seoul National University, Seoul, Republic of Korea; Department of Nuclear Medicine, Seoul National University Hospital, Seoul, Republic of Korea; Department of Thoracic and Cardiovascular Surgery, Seoul National University Hospital, Seoul, Republic of Korea; Seoul National University Cancer Research Institute, Seoul National University College of Medicine, Seoul, Republic of Korea; Department of Pathology, Seoul National University Hospital, Seoul, Republic of Korea; Department of Nuclear Medicine, Seoul University College of Medicine, Republic of Korea

**Author notes:** Correspondence Hongyoon Choi, MD., Ph.D. Department of Nuclear Medicine, Seoul National University Hospital 101, Daehak-ro, Jongno-gu, Seoul 03080, Republic of Korea, Young Tae Kim, MD., Ph.D. Department of Thoracic and Cardiovascular Surgery, Seoul National University Hospital 101 Daehak-ro, Jongno-gu, Seoul, Republic of Korea, 03080. These authors contributed equally to this work (Sungwoo Bae, Kwon Joong Na).

**Keywords:** spatially resolved transcriptomics, single-cell genomics, domain adaptation, deep learning, cell label transfer

## Abstract

Deciphering the cellular composition in genome-wide spatially resolved transcriptomic data is a critical task to clarify the spatial context of cells in a tissue. In this study, we developed a method, CellDART, which estimates the spatial distribution of cells defined by single-cell level data using domain adaptation of neural networks and applied it to the spatial mapping of human lung tissue. The neural network that predicts the cell proportion in a pseudospot, a virtual mixture of cells from single-cell data, is translated to decompose the cell types in each spatial barcoded region. First, CellDART was applied to mouse brain and human dorsolateral prefrontal cortex tissue to identify cell types with a layer-specific spatial distribution. Overall, the suggested approach was competent to the other computational methods in predicting the spatial localization of excitatory neurons. Besides, CellDART was capable of decomposing cellular proportion in mouse hippocampus Slide-seq data. Furthermore, CellDART elucidated the cell type predominance defined by the human lung cell atlas across the lung tissue compartments and it corresponded to the known prevalent cell types. CellDART is expected to help to elucidate the spatial heterogeneity of cells and their close interactions in various tissues.

## INTRODUCTION

Rapid progress in spatially resolved transcriptomics helped to comprehensively characterize the spatial interaction of cells in a tissue (1, 2). Breakthrough technologies enabled capturing genome-wide spatial gene expression at a resolution of several cells (3) to the single-cell (4–6) and even subcellular levels (7). These methods have been used in various disease models to decipher spatial maps of genes of interest and culprit cells (8–12). Furthermore, emerging computational approaches facilitated the spatiotemporal tracking of specific cells and elucidated cell-to-cell interactions by preserving the spatial context (12–14). However, there is an inherent limitation in the spatial transcriptomic analysis that each spot or bead covers more than one cell in most cases. Even with a high-resolution technique, a small portion of several cells can be contained in the same spatial barcoded region. In addition, a tissue with a high level of heterogeneity, such as cancer, consists of a variety of cells in each small domain of the tissue (15). Thus, the identification of different cell types in each spot is a crucial task to understand the spatial context of pathophysiology using a spatially resolved transcriptome.

In this regard, recent computational tools have focused on integrating different types of transcriptomic data, particularly spatially resolved transcriptomic and single-cell RNA-sequencing (scRNA-seq) data (14,16–24). These tools have utilized the cell type signatures or variable genes defined by scRNA-seq and transferred the cell labels into spatial transcriptomic data. The majority of the approaches applied a statistical model or a matrix decomposition to infer the cell fraction in each spot (18,20–23). Meanwhile, calculating the proportion of cell types defined by scRNA-seq data from spots of spatially resolved transcriptomic data can be considered a domain adaptation task (25, 26). A model that predicts cell fractions from the gene expression profile of a group of cells can be transferred to predict the spatial cell-type distribution.

In this paper, we suggest a method, CellDART, that implements modified adversarial discriminative domain adaptation (ADDA) (27) to infer the cell fraction in spatial transcriptomic data. The randomly selected cells from scRNA-seq data constitute a pseudospot in which the fraction of cells is known. The neural network model that extracts the cell fraction from the gene expression of a pseudospot is adapted to a different domain where spatial transcriptomic data are present. Consequently, the joint analysis of spatial and single-cell transcriptomic data elucidates the spatial cell composition and unveils the spatial heterogeneity of the cells. We utilized the proposed method to provide a resource for spatial mapping of the human lung cell atlas using the spatially resolved transcriptome of human lung tissue.

## MATERIALS AND METHODS

### Human brain cortex data

A publicly available Visium spatial transcriptomics dataset obtained from the DLPFC of postmortem neurotypical humans was downloaded from the data repository provided by the paper (28) (Two datasets in this study included ’tissue 151673’ with 3639 spots and 33538 genes and ‘tissue 151509’ with 4789 spots and 33538 genes). The count matrix for the tissue and brain layer information for each spot (cortical layers 1-6 and white matter) was added, and spots with no layer information were excluded from further analysis. Single-nucleus transcriptomic data acquired from the DLPFC of a healthy human control group (n=17) were utilized for the joint analysis (29). The count matrix for 35212 cells and 30062 genes and the cell type annotation were included in the analysis.

### Mouse brain data

Visium spatial transcriptomics dataset for the mouse brain was downloaded from the 10X Genomics Data Repository. The ‘Mouse Brain Serial Section (Sagittal-Anterior)’ slide, which contains 2695 spots and 32285 genes, was utilized for the CellDART analysis. For the joint analysis, scRNA-seq data obtained from the mouse primary visual cortex and anterior lateral motor cortex were selected. The count matrix was comprised of 23178 cells and 45768 genes. The layer-specific excitatory neuron types [L2/3 IT (intratelencephalic), L4, L5 IT, L5 PT (pyramidal tract), L6b, L6 CT (corticothalamic) and L6 IT] were determined based on the markers discovered in a previous study (30).

### Mouse hippocampus Slide-seq data

Slide-seq data for mouse hippocampus was obtained from Single Cell Portal repository offered by the paper (6). The spatial data contains count matrix for 23,264 genes across 53,173 beads. To infer the cellular composition in the Slide-seq data, scRNA-seq data of mouse hippocampus with 52,846 cells and 27,953 genes was utilized for the integration. The cell labels were determined based on the cell sub-clustering results suggested by the paper (31).

### Normal human lung data

Two normal lung samples were acquired from lung specimens from one patient who underwent surgical resection for lung cancer. We acquired samples and embedded them in optimal cutting temperature (OCT) compound in the operating room and stored them at -80°C until cryosectioning. For cryosectioning, samples were equilibrated to -20°C with a cryotome (Thermo Scientific, USA). Sections were imaged and processed for spatially resolved gene expression using the Visium Spatial Transcriptomic kit (10X Genomics, USA). The protocol of this study was reviewed and approved by the institutional review board of Seoul National University (Application number: H-2009-081-1158). ’Lung 1’ consists of 1591 spots and 36601 genes, and ’Lung 2’ consists of 2683 spots and 36601 genes. Single-cell data from the normal lung tissue of three subjects were downloaded and utilized for the integrated analysis (32). The count matrix for 65662 cells and 26485 genes and the cell labels were included in the downstream analysis.

### Preprocessing spatial and single-cell datasets

All of the preprocessing steps were performed with Python (version 3.7) with the Scanpy toolkit (version 1.5.1) (33). The count matrices for both spatial and single-cell datasets were normalized with the ’scanpy.pp.normalize_total’ function such that gene expression was comparable between spots or cells. For the single-cell data, the counts were log-transformed (scanpy.pp.log1p) followed by scaling (scanpy.pp.scale) and dimensionality reduction by principal component analysis (scanpy.tl.pca). Finally, the cells were represented with a t-distributed stochastic neighborhood embedding (t-SNE) plot (scanpy.tl.tsne and scanpy.pl.tsne) and were named based on the annotation data from publicly available datasets.

Meanwhile, the top *l* highly expressed marker genes for each cell cluster from brain and lung samples were extracted with the Wilcoxon rank-sum test (’scanpy.tl.rank_genes_groups’) based on the log-normalized count. Multiple comparison correction with the Benjamini-Hochberg method was applied, and genes were ranked by the corrected p-values. All of the cell type markers were pooled to form cell signature genes (**Supplementary Figure S1A**). The intersection between the cell signature genes and all provided genes from the spatial data was obtained. The downstream analysis was performed only with these intersecting genes.

For the next step, *k* cells were randomly selected from the mouse or human brain and lung single-cell datasets. Random weights were given to each cell to mimic the cases in which only the portion of the cells are contained in a spatial spot (**Supplementary Figure S1A**). The virtual mixture of the cells was defined as a ’pseudospot’. A total of *n* pseudospots were generated, and the composite gene expression values were calculated for each pseudospot.

Then, the log-normalized count matrices for single-cell, pseudospot, and real spot data were scaled such that the value lies between 0 and 1 in each cell or spot.

### CellDART: Cell type inference with domain adaptation

The modified ADDA algorithm (27) was applied to develop a model to predict cell type proportions for each spot (**Figure 1A and Supplementary Figure S1B**). The training of neural networks was implemented based on Keras (version 2.3.1), TensorFlow (version 1.14.0) and scikit-learn (version 0.24.1) packages. First, a feature embedder that computes 64-dimensional embedding features from the gene expression data of either real spatial spots or pseudospots was defined. The feature embedder was comprised of two fully connected layers, each of which underwent batch normalization and activation by the ELU function. The outputs of the first layer and second layer have 1024 and 64 dimensions, respectively. Source and domain classifiers were defined such that they could predict the cell fraction in each spot and discriminate pseudospots from spots, respectively. The domain classifier consisted of two fully connected layers. The first layer with 32-dimensional output was connected to the embedded features. After batch normalization, ELU activation, and dropout, another layer to discriminate real spots from pseudospots was applied. The source classifier is directly connected to the embedded features of the feature extractor as a one-layer model connected to the feature embedder. Therefore, the feature extractor attached to either of the classifiers was named a source or domain classification model. The source and domain classification model shared the feature extractor.

**Figure 1.**
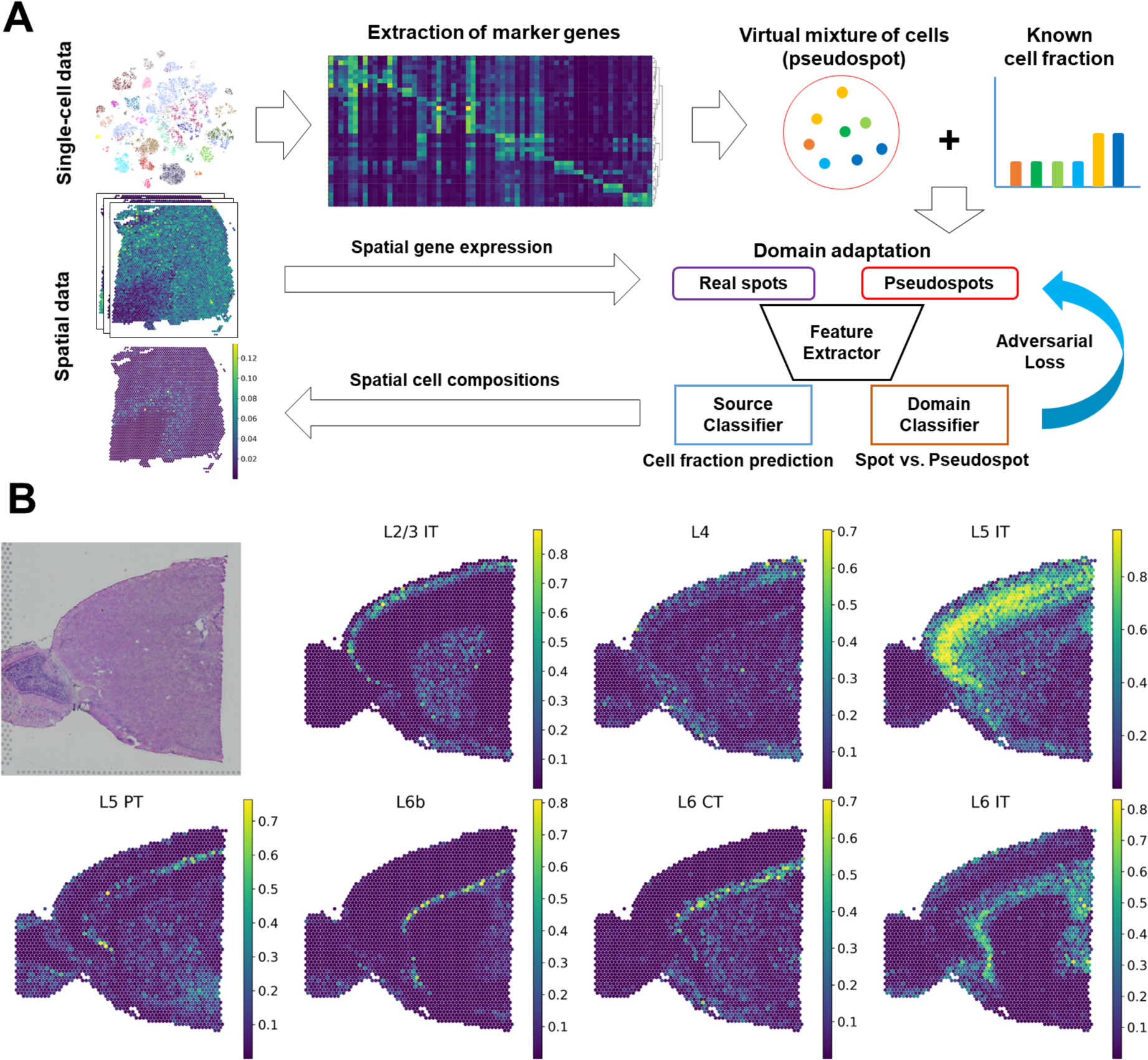
CellDART analysis in human and mouse brain tissues. **(A)** Schematic diagram for CellDART analysis. The human dorsolateral prefrontal cortex (DLPFC) dataset was preprocessed, and the marker genes for each cell cluster were extracted. The shared genes between the pooled cluster markers and the spatial transcriptomic data were selected for the downstream analysis. Then, 20000 pseudospots were generated by randomly selecting 8 cells from the single-cell data and giving them random weights. A feature extractor with a source and domain classifier was trained to estimate the cell fraction from the pseudospot and distinguish the pseudospots from the real spatial spots. First, weights of the neural network were updated except for the domain classifier. Next, the data label for the spot and pseudospot was inverted, and only the domain classifier was updated. Finally, the trained CellDART model was applied to spatial transcriptomics data to estimate the cell proportion in each spot. **(B)** Spatial mapping of 7 layer-specific mouse excitatory neurons predicted by CellDART. The figure in the top left corner shows the mouse brain tissue slide. Colormaps present the maximum and minimum values for the corresponding cell fraction.

The loss function of the source classifier that predicts cell type proportions was defined by Kullback-Leibler divergence (KLD). KLD is decreased when the distribution of predicted and real cell type proportions is similar. For the initialization of weights of feature embedder and source classifier, initial training to predict cell type proportions for pseudospots was performed. The optimization process can be summarized by these formulae:

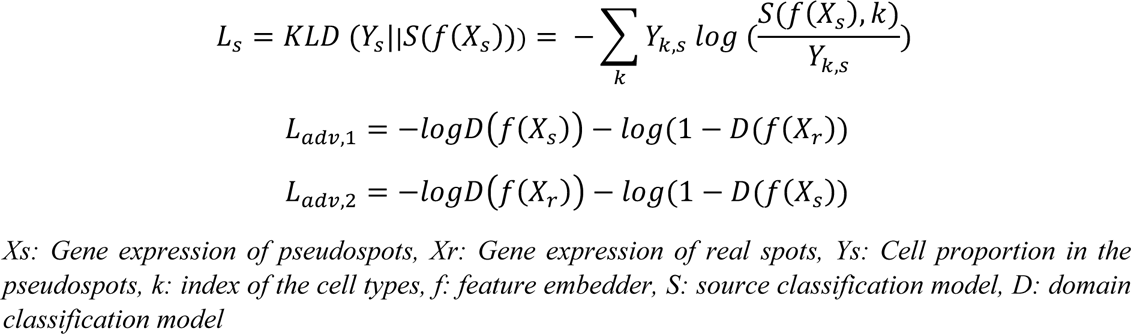

First, the model was trained to minimize *Ls* as a pre-training process. As an adversarial loss, it is typical to train the model with the standard loss function with inverted labels as in the above formula. Thus, two optimization processes were applied (**Supplementary Figure S1B**). The networks were optimized to minimize *Ls* and *Ladv,1* with fixed weights of the domain classifier. Then, the domain classifier was trained to minimize *Ladv,2* with fixed weights of the feature embedder, *f* and the source classifier, *S*. These two processes were repeated with a training parameter of the number of iterations.

Finally, the trained model, CellDART, predicted the cell fraction in each spot from the spatial data, and the results for each cell type were spatially mapped to the tissue by the ’scanpy.pl.spatial’ function. Additionally, the distribution of cell type compositions across the brain layer was represented with the ‘scanpy.pl.stacked_violin’ function.

### Optimal parameter selection

To search optimal parameters for CellDART, the performance of the model was evaluated in different parameter settings with DLPFC dataset as a reference. The DLPFC spatial data contain the brain layer information (layer 1 to layer 6, white matter, and unknown), and the single-cell data have ten layer-specific excitatory neuron clusters (*Ex_1_L5_6*, *Ex_2_L5*, *Ex_3_L4_5*, *Ex_4_L6*, *Ex_5_L5*, *Ex_6_L4_6*, *Ex_7_L4_6*, *Ex_8_L5_6*, *Ex_9_L5_6*, and *Ex_10_L2_4*). The layer specificity of excitatory neurons was determined by the cell types identified from the single-nucleus RNA-seq data (29) with the layer markers suggested by several studies (34–36). Receiver operating characteristic (ROC) analysis was performed to determine whether the spatial cell fraction of excitatory neuron clusters could differentiate the specific cortical layer.

The performance of CellDART was tested by modulating the number of markers in each cell cluster from 5 to 80 (5, 10, 20, 40, 80) which corresponds to the total number of markers from 109 to 1157 (109, 203, 364, 642, 1157). It showed that setting total number of markers between 300 and 400 leads to overall high AUC value (**Supplementary Figure S2A**). Also, the CellDART was implemented for different number of pseudospots ranging from 800 to 80000 (800, 4000, 20000, 40000, 80000). The performance was optimal when the number was larger than 20000 which was approximately five to ten times the number of real spots in one spatial dataset (**Supplementary Figure S2A**). The time consumed for the training gradually increased as the number of markers and pseudospots increased (**Supplementary Figure S2B**). The number of cells in a pseudospot was fixed (brain Visium: *k* = 8, brain Slide-seq: *k*=2, and lung: *k*=10) and other parameters were tuned to obtain an optimal performance.

Finally, the number of marker genes for each cell cluster (*l*) was set to 20 and 10 for brain and lung tissues, respectively. The number of pseudospots (*n*) was set to 20000 and 500000 for Visium and Slide-seq data. The iteration number was 3000, the minibatch size was 512, and the learning rate for the training domain classifier was 0.005. The loss weights between the source and domain classifiers were 1:0.6 in brain tissues and 1:1 in lung tissues.

### Comparison to other tools

For the DLPFC datasets, the performance of CellDART was compared with five other computational tools: Scanorama (16), Cell2location (20), RCTD (21), SPOTlight (23), and Seurat (version 3) (17). Briefly, Scanorama and Seurat aligns the single-cell and spatial datasets based on the mutual nearest neighbors between the two. Meanwhile, Cell2location assumes that the count matrix from spatial data follows a negative binomial distribution and can be decomposed into a linear combination of cell type signatures. RCTD postulates that counts in spatial spots follow a Poisson distribution and applies the maximum likelihood method to estimate cell proportions. Finally, SPOTlight utilizes non-negative matrix factorization regression to obtain cell type specific gene signatures and applies it to infer cellular composition of spots. These five toolkits were applied, and cell density from Cell2location and cell fraction from Scanorama, RCTD, SPOTlight, and Seurat were spatially mapped to the tissue. For Cell2location analysis, we set the total number of cells, cell types, and groups of cell types in each spatial spot as 8, 9, and 5, respectively. Next, for RCTD, we selected ‘full mode’, which does not restrict the number of cells in each spot. Finally, for SPOTlight, we randomly selected 100 cells per each cell cluster and extracted top 3000 highly variable genes for the downstream analysis. The ROC analysis was performed in the DLPFC dataset for discriminating spatial localizing patterns of layer-specific excitatory neurons. AUC values were compared between CellDART, Scanorama, Cell2location, RCTD, SPOTlight, and Seurat. The cell fraction and layer information of each spot were bootstrapped, and 1000 null distribution samples for the area under the curve (AUC) value were calculated. The statistical analysis and visualization were implemented with scikit-learn and matplotlib (version 3.3.4).

### Spatial mapping of lung cells to normal lung tissue data: investigation of the spatial heterogeneity of the cells

The boundary of the tissue structures in two lung samples (lung 1 and lung 2) was delineated by a pathologist on H&E staining images, and the spots were classified into 6 domains: alveolar space, bronchial epithelium, fibrous stroma, immune cluster, terminal bronchiole, and vessels. The uncertain region of the tissue was named ‘unknown’ and more specifically ‘unknown stroma’ if the corresponding region was stromal tissue. After transferring the single-cell cluster labels to the spatial data, the minimum and maximum cell fraction values across all spots were scaled to 0 and 1, respectively. The cell types were divided into six categories based on where the cells were commonly found (32) (‘airway epithelium’, ‘alveoli epithelium’, ‘endothelial’, ‘muscle stromal’, and ‘other stromal’). The average scaled cell fraction in each tissue domain according to cell types was visualized with a seaborn clustermap function (version 0.11.1), and the cell type categories were color-coded and presented on top. For the next step, the cell types showing highly different cell fractions across the histological domains were selected, and their spatial composition was mapped to the tissue. Cell type selection was performed with the Wilcoxon rank-sum test, and Benjamini-Hochberg corrected p-values were computed. The cell types in each tissue domain were ranked based on a ratio of the average scaled cell fraction in a specific tissue domain to the rest of the domains. The cell types with an average scaled fraction below 0.2 were excluded from further analysis. An adjusted p-value below 0.05 was considered significant.

## RESULTS

### Decomposition of spatial cell distribution with CellDART in human and mouse brain data

The performance of CellDART was assessed in publicly available single-nucleus (also considered single-cell data) and spatial transcriptomic autopsy samples of the human dorsolateral prefrontal cortex (DLPFC), each of which was obtained from two different subject groups with no neurological disorders. Additionally, single-cell and spatial datasets acquired from the mouse brain were utilized. First, both single-cell datasets were preprocessed, and the cells were named after the annotation data provided by the original papers (29, 30). The 33 and 29 annotated cell clusters from the human and mouse brains were visualized by t-distributed stochastic neighbor embedding (t-SNE) plots (**Supplementary Figure S3A, B**), and marker genes for each cluster were extracted (**Supplementary Figure S3C, D and Tables S1, S2**). The cell clusters showed distinct gene expression patterns represented by cell type-specific marker genes.

A specific number of cells (*k* = 8) were randomly sampled from the single-cell data with random weights to generate pseudospots (number of pseudospots = 20000). Then, composite gene expression values were computed based on marker genes (**Figure 1A**). A neural network was trained to accurately decompose the pseudospots, and another network, the domain classifier, was trained to discriminate spots of real spatially resolved transcriptomes from pseudospots. During the training process, the weights of neural networks were updated to predict cell fractions and fool the domain classifier to avoid discriminating spots and pseudospots (**Figure 1A**). As a result, the neural network, source classifier, was trained to estimate cell fractions in both the pseudospots and the real spatial spots as an adversarial domain adaptation process.

The layer-specific excitatory neuron fraction in each spot was predicted by CellDART and spatially mapped to the tissue. In the case of mouse brain tissue, 7 excitatory neurons showed spatially restricted patterns in a specific cortical layer (**Figure 1B**). In addition, the 10 excitatory neuron clusters in the human brain presented layer-specific distribution patterns across the six cortical layers (layers 1 to 6) (**Figure 2A and Supplementary Figure S4A**). Additionally, the spatial density of human non-neuronal cells was estimated with a neural network (**Supplementary Figure S4B**). Astrocytes were mainly located in layer 1 and layer 6, while oligodendrocyte cluster 3 (*Oligos_3*), which showed an approximately 10 to 60 times higher cell fraction than the other two clusters (*Oligos_1* and *Oligos_2*), was predominantly localized in white matter. Endothelial cells, microglia, and macrophages were spatially distributed across the six cortical layers with low cell proportions.

**Figure 2.**
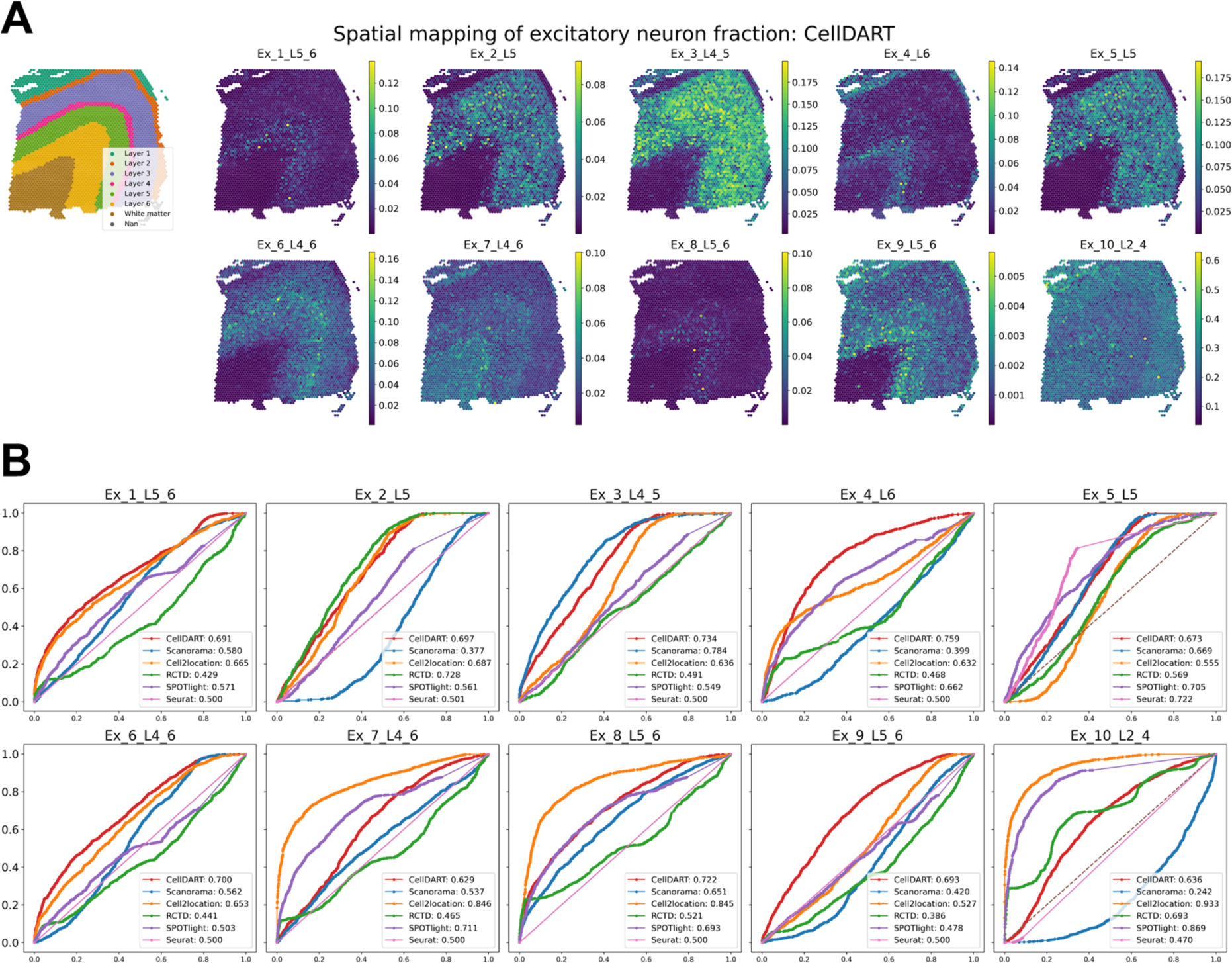
Implementation of CellDART in a human dorsolateral prefrontal cortex dataset. **(A)** Spatial mapping of 10 layer-specific excitatory neurons predicted by CellDART. The figure in the top left corner shows the layer annotation for each spatial spot. The layer consists of cortical layers 1 to 6 and white matter. ‘Nan’ represents the spot without the layer information. Colormaps present the maximum and minimum values for the corresponding cell fraction. **(B)** Receiver operating characteristic (ROC) analysis for predicting the layer-specific distribution of excitatory neurons. The computational tools CellDART, Scanorama, Cell2location, and RCTD, which estimate cell types in the spatial spots, were compared by means of the area under the curve (AUC). The ROC curves for CellDART, Scanorama, Cell2location, RCTD, SPOTlight, and Seurat (version 3) are color-coded, and AUC values are presented in the lower right corner of each plot.

Meanwhile, the impact of domain adaptation on improvement of model performance was assessed. The feature embedder with source classifier were trained only with pseudospots and domain adaptation was not applied. The model without domain adaptation was named as ‘NN_wo_da’. The cell fraction predicted by ‘NN_wo_da’ presented spatially localizing patterns in *Ex_2_L5* and *Ex_5_L5*; however, other cell types showed uneven spatial distributions (**Supplementary Figure S5A**). ROC analysis was implemented and the accuracy of CellDART in spatially localizing layer-specific excitatory neuron fraction was compared with that of NN_wo_da (**Supplementary Figure S5B**). In general, the performance of NN_wo_da was inferior to CellDART, which proves the necessity of domain adaptation to obtain an optimal result.

### Comparison of CellDART with other integration tools in human brain tissue

The capability of CellDART to accurately assign cell types in spatial spots was compared with that of five computational tools: Scanorama, Cell2location, RCTD, SPOTlight, and Seurat version 3. The five methods were employed to decipher the spatial distribution of excitatory neurons and non-neuronal cells in the DLPFC 151673 dataset.

Scanorama showed a few excitatory neurons of cortical layer-specific distribution patterns, whereas *Ex_2_L5*, *Ex_4_L6*, *Ex_9_L5_6*, and *Ex_10_L2_4* excitatory neurons were distributed differently from the known cortical distribution (**Supplementary Figure S6A, B**). Astrocytes and oligodendrocytes did not show consistent cell distribution patterns across the cell subtypes. Endothelial cells, microglia, and macrophages were predominantly localized in layer 1, layer 6, and the white matter according to the Scanorama analysis (**Supplementary Figure S6C**).

In the case of Cell2location, excitatory neurons showed layer-specific distribution patterns except for *Ex_5_L5* and *Ex_9_L5_6* where the distribution was uneven and not restricted to layer 5 and layer 5-6, respectively. Overall, non-neuronal cells showed layer-specific localization pattern; however *Astros_2* and *Astros_3* exhibited heterogeneous patterns in the same layer (**Supplementary Figure S7**).

Next, for RCTD, a few excitatory neurons (*Ex_2_L5* and *Ex_10_L2_4*) exhibited a high cell fraction in the corresponding cortical layer of a known layer specificity; however, other excitatory neurons presented heterogeneous patterns of distribution (**Supplementary Figure S8A, B**). Additionally, in the non-neuronal cells, the spatial distribution was relatively uneven and not layer-specific except for three oligodendrocyte cell clusters (**Supplementary Figure S8C**).

SPOTlight exhibited spatially restricted patterns of distribution for some excitatory neurons and non-neuronal cells. However, *Ex_1_L5_6*, *Ex_2_L5*, *Ex_3_L4_5*, *Ex_6_L4_6, Ex_9_L5_6*, *Astros_2*, *Astros_3*, and *Oligos_3* were not layer-specific or did not show even distribution in the same layer (**Supplementary Figure S9**).

Finally, Seurat version 3 was successful in discriminating layer-specific distribution patterns for few cell types (*Ex_5_L5*, *Oligos_1*, and Oligos_3). In other cell types, spatial distribution could not be obtained or did not exhibit layer-specific localization patterns (**Supplementary Figure S10**).

Receiver operating characteristic (ROC) curve analysis was implemented to compare the performance of the four different tools in predicting the layer-specific distribution of excitatory neurons (**Figure 2B**). The spatial spots in the DFPLC data were classified into a specific layer, layer 1 to layer 6, white matter, or unknown, with manual annotation data (28) based on the tissue morphology and marker genes (34). The ROC curves for 10 excitatory neurons revealed that CellDART has overall good prediction accuracy, with an area under the curve (AUC) ranging from 0.629 in *Ex_7_L4-6* to 0.759 in *Ex_4_L6*. On the other hand, Scanorama and RCTD exhibited relatively low discriminative accuracy in several cell types with AUCs below 0.500 and comparable AUCs with CellDART for a few cell types. In addition, Cell2location showed several cell types with AUCs below 0.500 and a maximum AUC of 0.591. The confidence interval of the AUC was generated by bootstrapping, and the results were compared among the six methods (**Supplementary Figure S11A**). In general, CellDART showed superior performance in predicting layer-specific localization patterns across all excitatory neurons (**Supplementary Figure S11B**).

The performance of CellDART was further validated in another DLPFC 151509 spatial data. Although *Ex_6_L4_6* did not show layer-specific distribution pattern (AUC: 0.433), the CellDART was overall competent in predicting spatial fraction in other excitatory neuron types (AUC: 0.566 – 0.894) (**Supplementary Figure S12A, B**). Other computational tools showed poor accuracy in more than two cell types with AUC same or below 0.500 (**Supplementary Figure 12B**). Furthermore, non-neuronal cells showed similar localizing patterns compared with DLPFC 151673 dataset. Three astrocyte cell types were mainly located in layer 1 or layer 6 while oligodendrocytes mainly in white matter (**Supplementary Figure 12C**). In summary, CellDART exhibited consistently good performance in predicting layer-specific distribution patterns of cells in brain.

### Application of CellDART in Slide-seq data

The capability of CellDART was evaluated in Slide-seq data to decompose cellular proportion in each spatial bead. CellDART could precisely localize the cells, especially for CA1, CA2, CA3, dentate principal cells, and entorhinal cortex, which are known to be restricted in specific anatomical locations (**Supplementary Figure S13**) (37). Also, astrocytes are found to have high density in stratum oriens and stratum radiatum and oligodendrocytes in corpus callosum as previously reported (38–40). In short, CellDART can be applied to spatial transcriptome with different spatial resolution.

### Discovery of spatial heterogeneity of human lung tissue with CellDART

CellDART was further applied to normal lung spatial transcriptomic data. Human lung tissue was obtained from one patient who underwent lobectomy for surgical resection of lung cancer. The two normal lung tissues were dissected far from the tumor and pathologically confirmed to have no tumor cells (**Supplementary Figure S14**). The demographic features of the patient are summarized in **Supplementary Table S3**. The publicly available human lung cell atlas data were used for spatial mapping of lung cell types using CellDART. They consisted of scRNA-seq from three human normal lung tissues (32). The single-cell data were embedded in low-dimensional space by a t-SNE plot, and 57 cell clusters showed discrete gene expression patterns (**Supplementary Figure S15A**). The marker genes selected in each cell cluster were pooled and utilized in the downstream analysis (**Supplementary Figure S15B and Supplementary Table S4**). After the generation of pseudospots, CellDART was trained to assign the proportion of cells in the real spatial spots. The tissue slides from two spatial datasets (lung 1 and lung 2) were manually segmented, and each spot was classified into 7 categories: alveolar space, bronchial epithelium, fibrous stroma, immune cluster, terminal bronchiole, vessels, and unknown region (**Figure 3A, B**). In addition, the cell types were classified into five categories based on a previous study (32). An average scaled cell proportion of spots in the same tissue domain was calculated, and the values were expressed with heatmaps.

**Figure 3.**
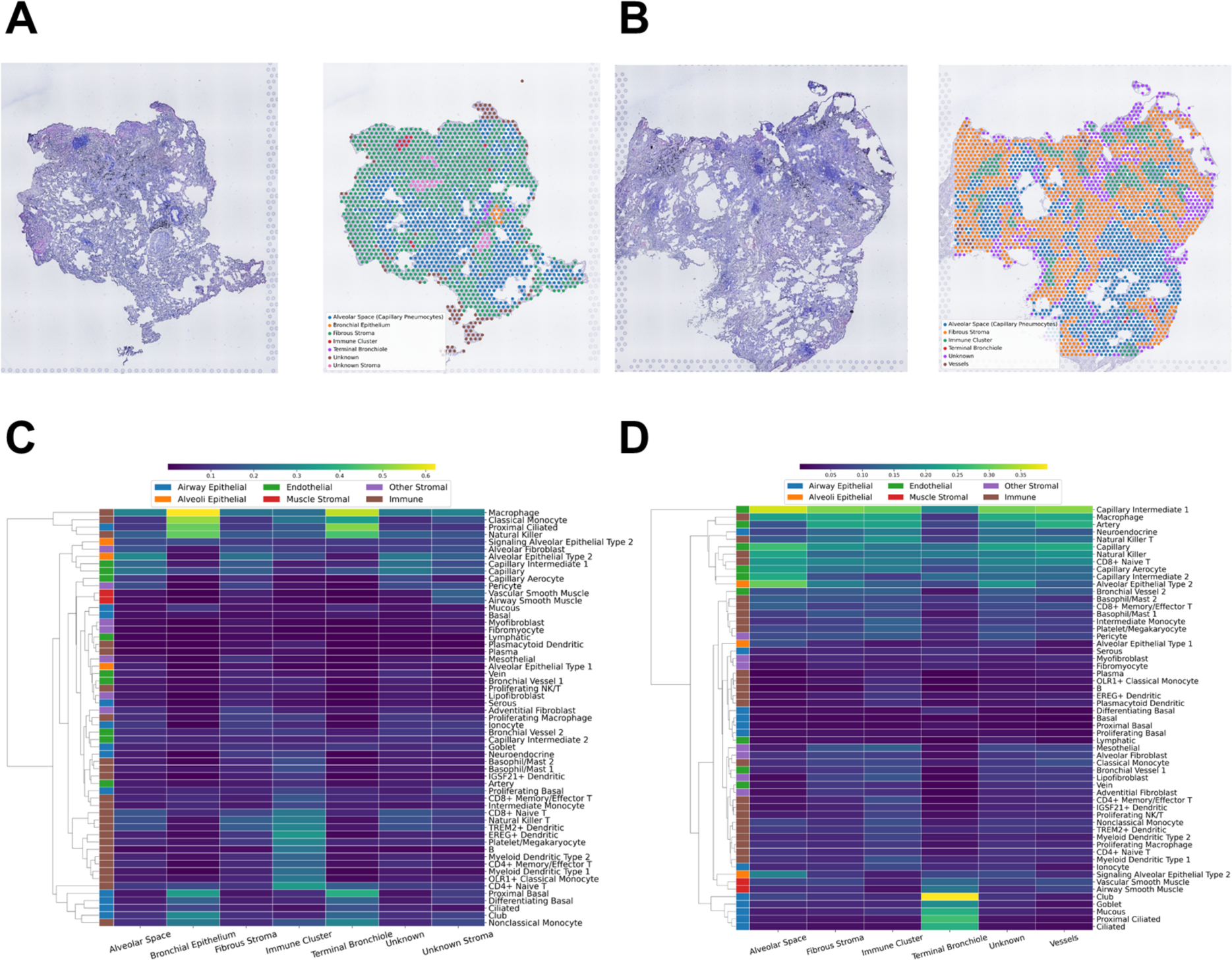
Application of CellDART in human lung data to decipher the tissue microenvironment. **(A-B)** Segmentation of the histological structures in normal lung tissue (A) 1 and (B) 2. The tissue was divided into six domains: alveolar space (capillary pneumocyte), bronchial epithelium, fibrous stroma, immune cluster, terminal bronchiole, and vessels. Uncertain stromal tissue was defined as ‘unknown stroma’, and the other unspecified areas were defined as ‘unknown’. **(C-D)** Heatmaps for the average scaled cell fraction in each histological domain of (C) lung 1 and (D) 2 tissues. The cell types were classified into 6 categories (‘alveolar epithelial’, ‘alveoli epithelial’, ‘endothelial’, ‘muscle stromal’, ‘other stromal’, and ‘immune’) based on the original paper of the human lung cell atlas (32) and color-coded on top of the heatmaps. Additionally, hierarchical clustering was performed based on cell fraction profiles across the tissue compartment to visualize the similarity between the cell types.

In both the lung 1 and lung 2 datasets, each cell type showed different distribution patterns across the segmented tissue domains (**Figure 3C, D**). Among the ‘airway epithelial’ cells (blue color on the left side of the heatmaps), proximal ciliated, ciliated, mucous, and club were mainly localized in bronchial epithelium or terminal bronchiole. ‘Alveoli epithelial’ cells (orange color) were localized in the alveolar space of lung 2 data. ‘Muscle stromal’ cells (red color) were mainly distributed in unknown stroma in the lung 1 data and terminal bronchioles or vessels in the lung 2 data. ‘Immune’ cells (brown color), particularly B-cells, monocytes, and dendritic cells, were predominantly located in the immune cluster tissue domain. Finally, ‘endothelial’ and ‘other stromal’ cells did not present spatially localized patterns of distribution.

For the next step, cell types that showed highly different cell fractions across the tissue domains were selected (**Supplementary Table S5**). The top 7 cell types were ranked by the ratio of the scaled cell fraction in the specific tissue domain compared to the other domains and were mapped to the tissue (**Figure 4A, B**). The cell types with an average scaled cell fraction in the domain below 0.2 were excluded. For lung 1 tissue, ‘proximal basal’ and ‘proximal ciliated’ cells, which were previously described as ‘airway epithelial’ cells (blue color), were predominantly distributed in the bronchial epithelium or terminal bronchiole tissue domain (**Figure 4A**). Most ‘immune’ cell types (brown color) were localized in the immune cluster domain except for ‘classical monocytes’, which are commonly found in bronchial epithelium. In lung 2 tissue, ‘ciliated’, ‘proximal ciliated’, ‘club’, and ‘mucous’ cells, which are included in ‘airway epithelial’ (blue color) cells, were mainly located in the terminal bronchiole domain (**Figure 4B**). ‘Capillary intermediate 2’ included in the ‘endothelial’ cell type (green color) was localized in the alveolar space domain, while another endothelial cell type, ‘artery’, was mainly located in the fibrous stroma and vessels. Additionally, ‘alveolar epithelial type 2’ in the ‘alveoli epithelial’ (orange color) was predominantly distributed in the alveolar space domain. In summary, CellDART could precisely localize the spatial distribution of heterogeneous cell types in normal lung tissue.

**Figure 4.**
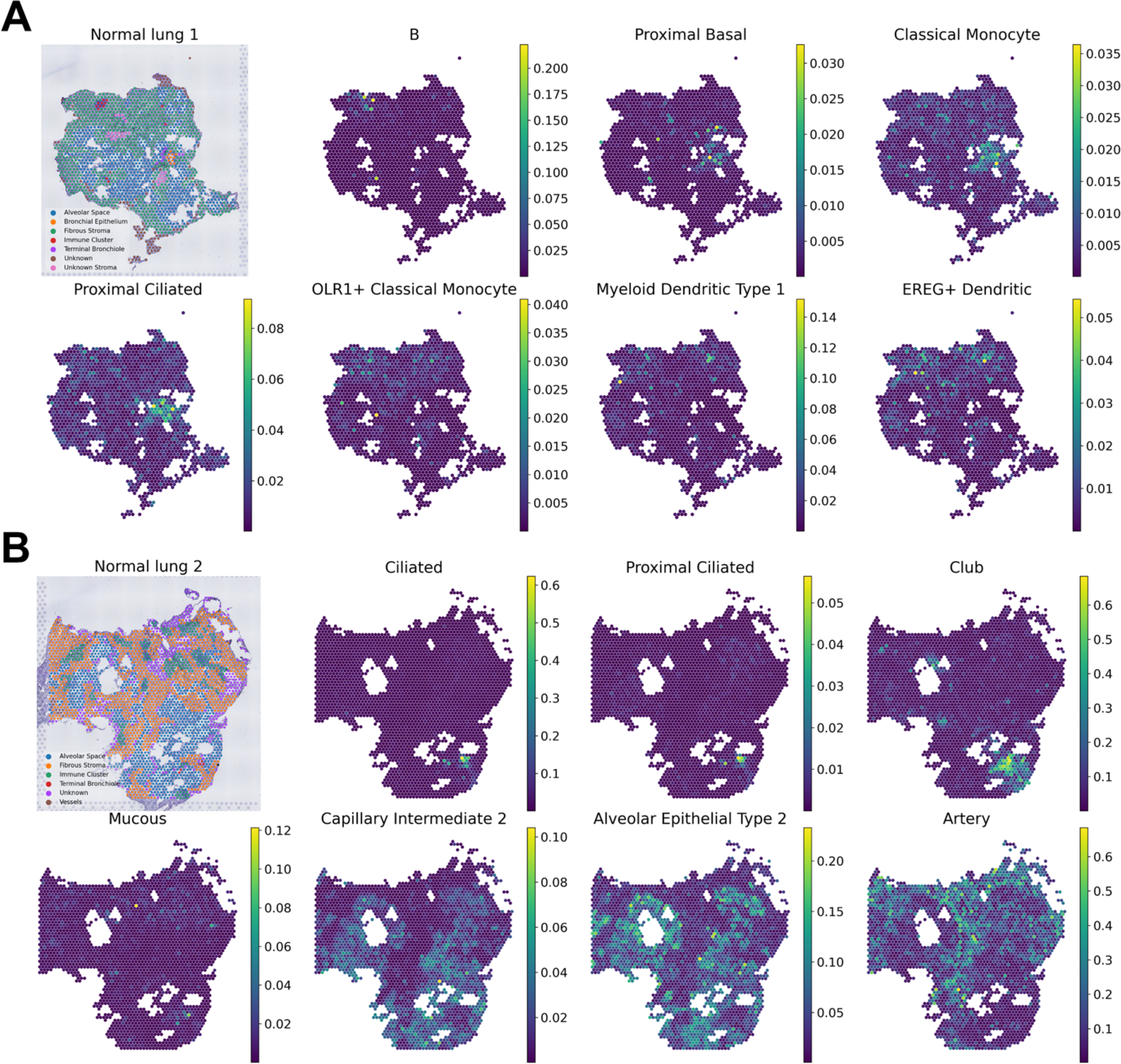
Spatial compositions of tissue compartment-specific cell types in the human lung. **(A-B)** Spatial mapping of the lung cell fraction for the top 7 cell types predominant in a specific tissue compartment of the (A) lung 1 and (B) 2 datasets. The cell types were ranked based on the average scaled cell fraction in a specific tissue domain compared to the other domains. The figure in the top left corner shows the classification of the tissue domain for each spatial spot. Colormaps present the maximum and minimum values for the corresponding cell fraction.

## DISCUSSION

CellDART, which adapts the domain of single-cell and spatial transcriptomic data, could be flexibly applied to brain and lung tissues to decompose the spatial distribution of various cell types. The suggested approach was capable of accurately predicting the layer-specific localization of excitatory neurons and non-neuronal cells in the brain. Additionally, CellDART was competent to the other six computational tools, Scanorama, Cell2location, RCTD, SPOTlight, and Seurat in spatially localizing multiple excitatory neuron subtypes. Besides, CellDART could be applied to Slide-seq data to estimate spatial cell compositions. Finally, our domain adaptation method deciphered the spatial distribution patterns of various lung cells across the different tissue compartments. Since tissues consist of variable cells, spatial mapping of various cell types is crucial to understand functions and pathophysiology. As examples of brain and lung tissues in our results, deciphering layer-specific cell types and tissue compartments associated with specific cell types could be a resource to understand the underlying biology. Furthermore, although a simple approach using a priori cell type markers of specific cells could be used to understand brief patterns of cell types, tissues with complex and heterogeneous cell types such as lung require spatial mapping of precisely defined cell types based on single-cell level studies.

CellDART can be adopted for spatial transcriptomic data to portray the cellular landscape of the tissue by preserving the spatial context. In the brain, six cortical layers and the white matter contain different cell types, and the heterogeneous cells in each layer shape distinct functional characteristics (41). Therefore, it is crucial to precisely decompose brain cell types in spatial transcriptomic data to comprehensively analyze the spatial crosstalk among cells. In the mouse brain tissue, layer-specific excitatory neurons revealed localized patterns of distribution across the cortical layers (**Figure 1B**). Also, when applied to Slide-seq data, CellDART was successful in mapping various hippocampal cell types into anatomically relevant position. Additionally, in the validation study with human DLPFC tissue, not only layer-specific excitatory neurons but also glial cells such as astrocytes and oligodendrocytes showed spatially restricted patterns (**Figure 2 and Supplementary Figure S4B**). Three astrocyte subtypes, Astros_1, Astros_2, and Astros_3, were predominantly located in L1 and L6. It has been reported that astrocytes form cortical layer-specific morphological and gene expression features (41–43); however, the abundance of excitatory neurons in the mid cortical layers may have masked the presence of diverse astrocyte populations. Meanwhile, one of the oligodendrocyte subtypes, Oligos_3, which presented a higher absolute cell fraction than the other subtypes, was localized in the white matter. This finding is in line with a previous study showing that oligodendrocytes are highly restricted in white matter compared to gray matter (39). On the other hand, the spatial distribution of glial cells did not match the known localization patterns and presented heterogeneous distribution in Scanorama and Cell2location, respectively (**Supplementary Figures S6, 7**). In the case of RCTD, SPOTlight, and Seurat, oligodendrocytes showed strong localization patterns in white matter; however, other glial cells did not present a layer-specific distribution (**Supplementary Figure S8-10**).

Our suggested method, CellDART, was further applied to human lung tissues where a mixture of various cells was present across the tissue compartments (32). The cell types that exhibited a high proportion in a specific tissue domain corresponded with a previous paper presenting the cell type predominance in the lung compartments (**Figures 3, 4**). In addition, when the top 7 highly localized cell types in each tissue compartment were listed, the selected cell types were shared between the lung 1 and lung 2 tissues (**Supplementary Table S5**). More specifically, for the alveolar space tissue domain, three alveolar epithelial cells (‘alveolar epithelial type 1, 2,’ and ‘signaling alveolar epithelial type 2’) and two capillary cells (‘capillary intermediate 2 and ‘capillary aerocyte’) overlapped in both tissues. The cell types were also shared in the fibrous stroma (‘fibromyocyte’, ‘mesothelial’, and ‘myofibroblast’), immune cluster (‘B’, ‘OLR1+ classical monocyte’, ‘EREG+ dendritic’, and ‘plasmacytoid dendritic’), and terminal bronchiole (‘proximal basal’, ’proximal ciliated’, ‘differentiating basal’, and ‘ciliated’) tissue domains. In short, CellDART can accurately assign prevalent cell types in the tissue compartments and is reproducible across replicates of the tissue. Considering the heterogeneous cell types in the lung, our resource of spatially resolved cell types derived from human lung tissue data provides the spatial distribution of cell types and may be used as controls to analyze pathologic patterns of various lung diseases.

There are several important issues to consider before applying CellDART to transfer cell labels. First, the density of cells may vary in the different regions of the tissue. In our method, the sampled number of cells in a pseudospot is fixed during the training; however, the domain adaptation process aligns the pseudospot to the spatial data, and the impact of spatial cell density variance on the result may be attenuated. In addition, the small population of cells in the spatial data may be neglected during the prediction of the cell proportion. The proportion of those cell types can be masked due to other predominant cell types in the spatial spots. In that case, CellDART can be implemented for the corresponding subpopulation of cells by extracting the marker genes for the subclusters. Lastly, CellDART adopts shared feature embedder between pseudospots and real spots in the adversarial domain adaptation. There have been several deep learning-based domain adaptation approaches that have differences in feature embedders as well as training methods (44). Notably, our approach used the shared feature embedder considering the similar features representing cell types between pseudospots and real spots. Though CellDART showed competent performance with stable training results, there could be a room for improvement by finding optimized domain adaptation methods among various recently developed approaches.

In conclusion, CellDART is capable of estimating the spatial cell compositions in complex tissues with high levels of heterogeneity by aligning the domain of single-cell and spatial transcriptomics data. The suggested method may help elucidate the spatial interaction of various cells in close proximity and track the cell-level transcriptomic changes while preserving the spatial context.

## SUPPLEMENTARY DATA

Supplementary Data are available at NAR online.

## ACKNOWLEDGEMENTS

*Author’s contributions:* H.C. and Y.T.K. conceived the study. S.B., K.J.N., and H.C. designed the experiments. K.J.N. and Y.T.K. mainly contributed to acquire spatially resolved transcriptomic data. H.C. initially developed a new algorithm and S.B. modified and optimized the analytic method. H.C., D.S.L. and Y.T.K. devised the data analysis. S.B. mainly conducted the data analysis. J.K. conducted histologic image analysis. All authors contributed to the interpretation of the data and wrote the paper.

