## Supplementary Figure for "CellDART: Cell type inference by domain adaptation of single-cell and spatial transcriptomic data"

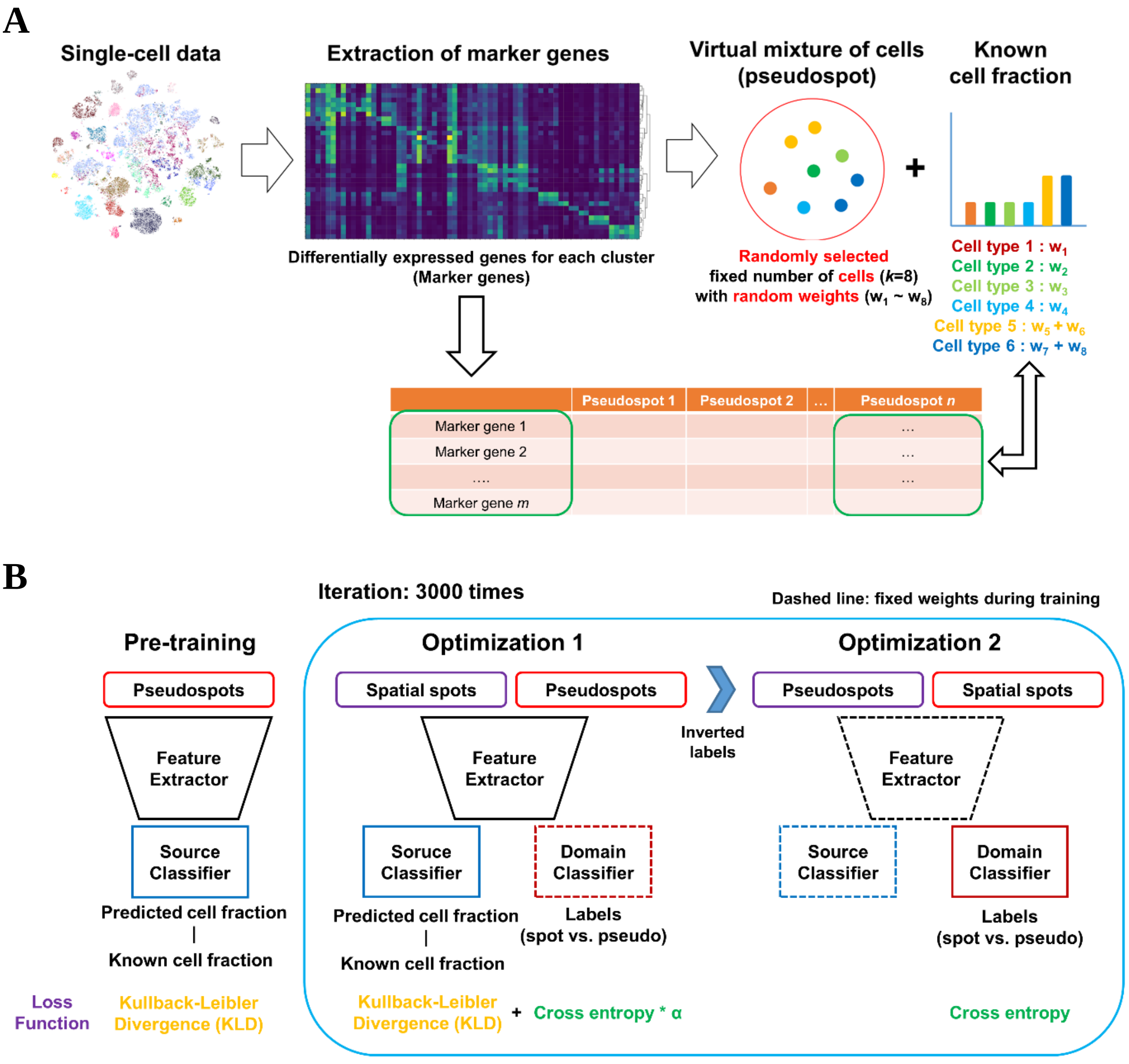


**Figure S1: Detailed workflow of CellDART analysis.**

**(A)** The top *l* (*l* = 20 for brain and *l* = 10 for lung) differentially expressed genes for each single-cell cluster were pooled to form a total of *m* marker genes. The fixed number of cells (*k* = 8 for brain Visium, *k* = 2 for brain Slide-seq, and *k* = 10 for lung) were randomly sampled from the single-cell data and random weights ($w_1$, $w_2$, ⋯, $w_k$) were given to each cell. The consequent mixture of the cells was called as a 'pseudospot'. The weighted sums were calculated for each marker gene expression in cells comprising a pseudospot. Also, the cell fraction was calculated by adding the weights given to each cell-type. The process was repeated to generate a total of *n* pseudospots.

**(B)** The training process was composed of two parts. In the pre-training, a feature extractor with a source classifier was trained to predict cell fraction. The neural network was trained such that loss function ($L_s$), Kullback-Leibler divergence (KLD), between predicted and known cell fraction is minimum. For the first optimization process, the feature extractor with the source and the domain classifier was jointly trained to minimize the loss represented as weighted sum of KLD ($L_s$) and cross entropy ($L_{adv,1}$). The weights of the domain classifier were fixed during the optimization. In the second optimization process, only the domain classifier was updated with the inverted label for spots and pseudospots ($L_{adv,2}$). The two optimization processes were iterated for 3000 times during the training.

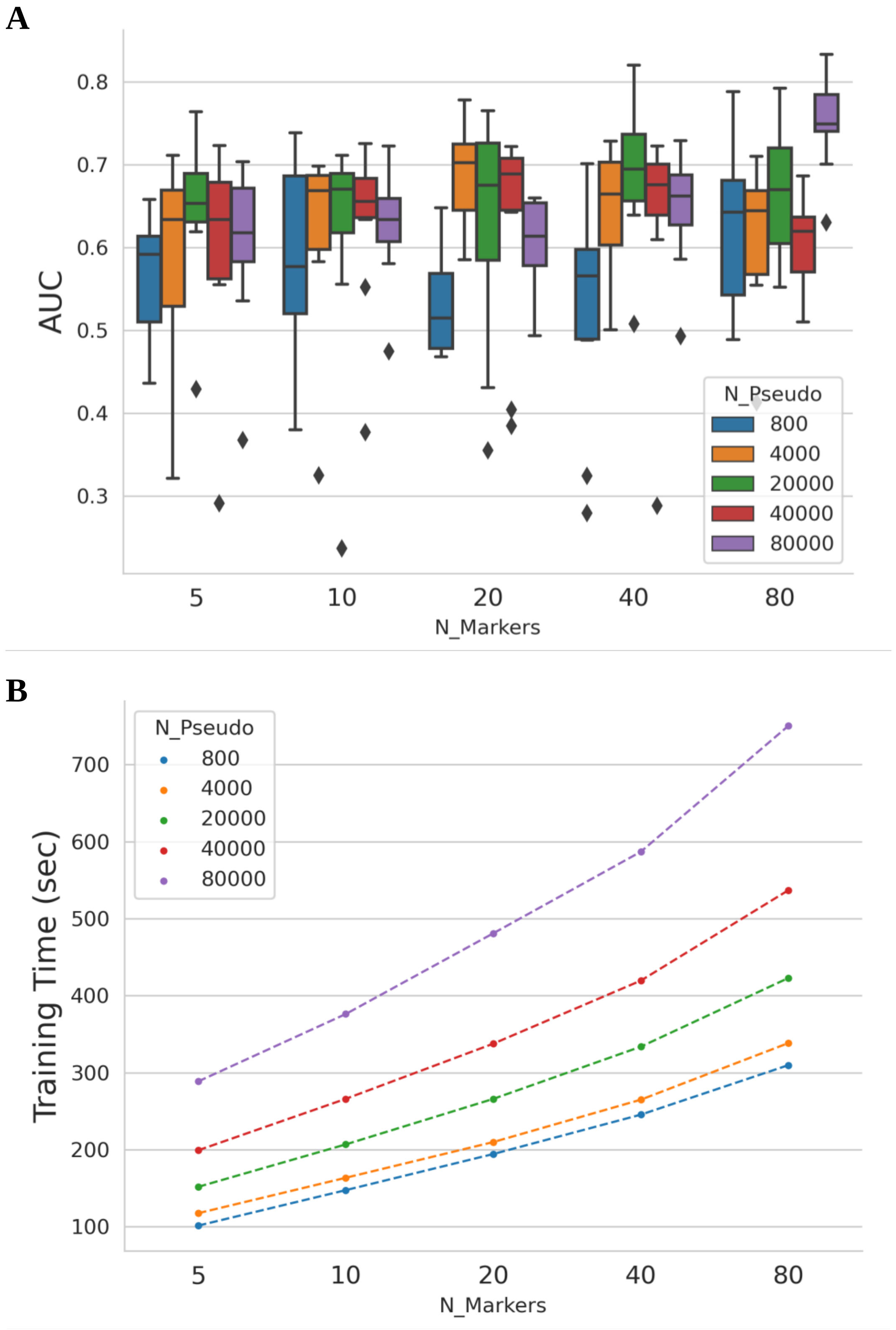

A
AUC
0.8
0.7
0.6
0.5
0.4
0.3
5
10
20
40
80
N_Markers
N_Pseudo
800
4000
20000
40000
80000
B
Training Time (sec)
700
600
500
400
300
200
100
5
10
20
40
80
N_Markers
N_Pseudo
800
4000
20000
40000
80000

**Figure S2: Evaluation of CellDART performance according to parameters**

**(A)** CellDART models with different parameters (number of marker genes in each cell cluster and number of pseudospots) were tested by means of prediction accuracy of layer-specific excitatory neuron distribution. Area under the curve (AUC) values were calculated for CellDART models with different number of marker genes (N_markers) and pseudospots (N_pseudo). The results across the ten excitatory neuron clusters were pooled and visualized in each boxplot.

**(B)** Training time for CellDART models with different number of marker genes (N_markers) and pseudospots (N_pseudo) were visualized with a line plot. The models were trained on one GPU (GeForce RTX 3090) device.

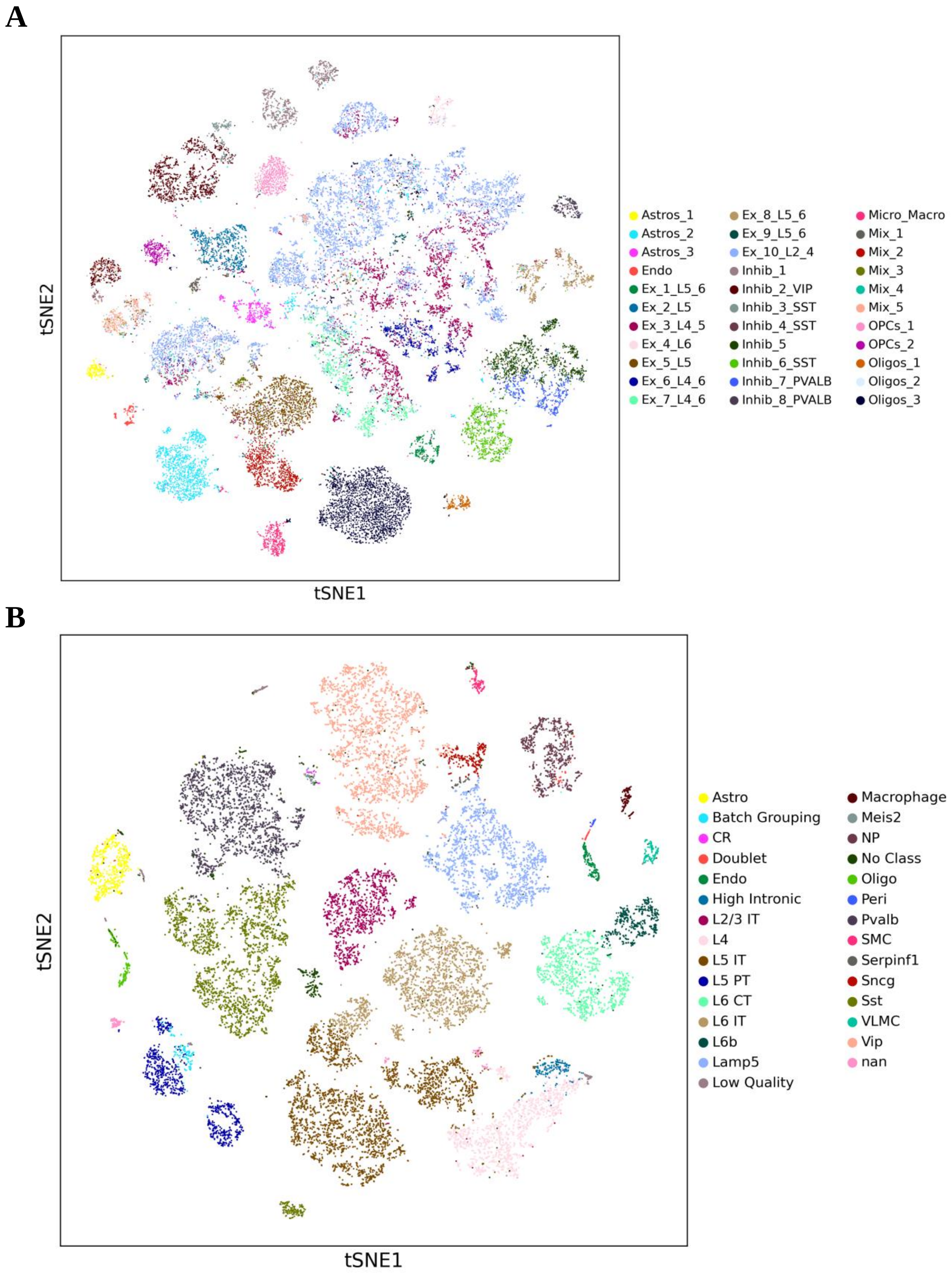

A
tSNE2
tSNE1
Astros_1
Astros_2
Astros_3
Endo
Ex_1_L5_6
Ex_2_L5
Ex_3_L4_5
Ex_4_L6
Ex_5_L5
Ex_6_L4_6
Ex_7_L4_6
Ex_8_L5_6
Ex_9_L5_6
Ex_10_L2_4
Inhib_1
Inhib_2_VIP
Inhib_3_SST
Inhib_4_SST
Inhib_5
Inhib_6_SST
Inhib_7_PVALB
Inhib_8_PVALB
Micro_Macro
Mix_1
Mix_2
Mix_3
Mix_4
Mix_5
OPCs_1
OPCs_2
Oligos_1
Oligos_2
Oligos_3
B
tSNE2
tSNE1
Astro
Batch Grouping
CR
Doublet
Endo
High Intronic
L2/3 IT
L4
L5 IT
L5 PT
L6 CT
L6 IT
L6b
Lamp5
Low Quality
Macrophage
Meis2
NP
No Class
Oligo
Peri
Pvalb
SMC
Serpinf1
Sncg
Sst
VLMC
Vip
nan

**C**

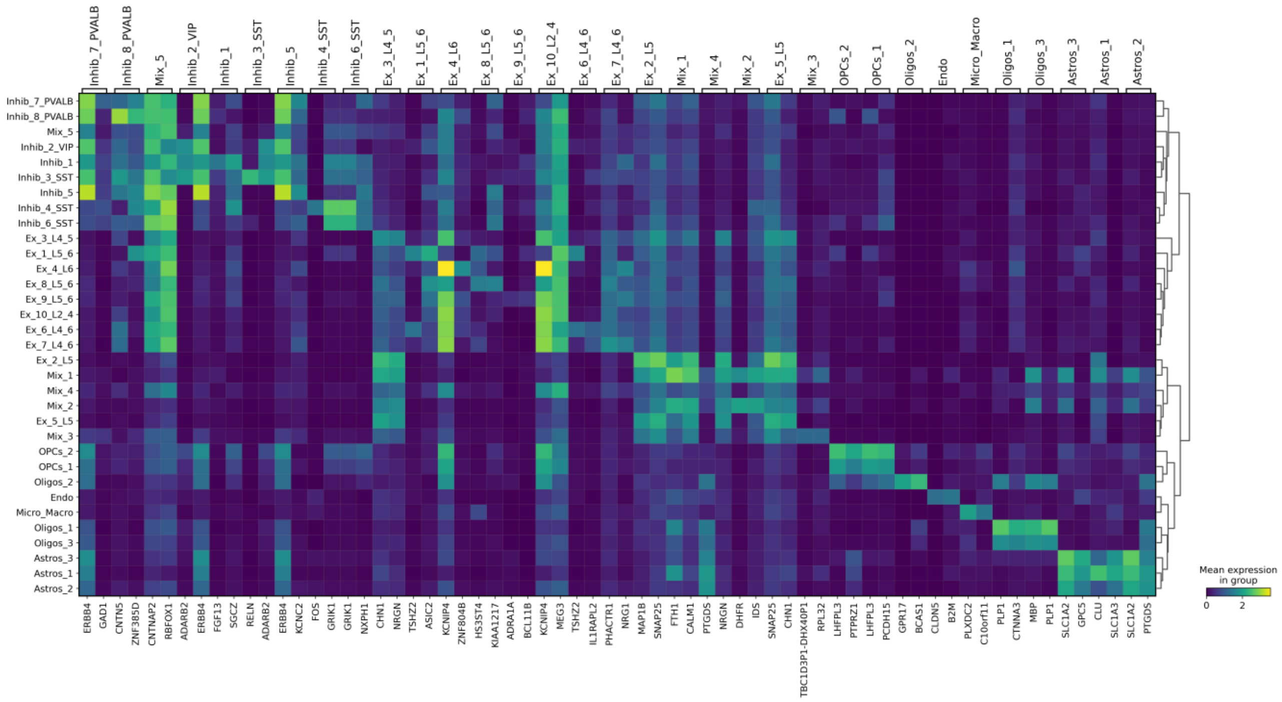
Inhib_7_PVALB
Inhib_8_PVALB
Mix_5
Inhib_2_VIP
Inhib_1
Inhib_3_SST
Inhib_5
Inhib_4_SST
Inhib_6_SST
Ex_3_L4_5
Ex_1_L5_6
Ex_4_L6
Ex_8_L5_6
Ex_9_L5_6
Ex_10_L2_4
Ex_6_L4_6
Ex_7_L4_6
Ex_2_L5
Mix_1
Mix_4
Mix_2
Ex_5_L5
Mix_3
OPCs_2
OPCs_1
Oligos_2
Endo
Micro_Macro
Oligos_1
Oligos_3
Astros_3
Astros_1
Astros_2
ERBB4 GAD1 CNTN5 ZNF385D CNTNAP2 RBFOX1 ADARB2 ERBB4 FGF13 SGCZ RELN ADARB2 ERBB4 KCNC2 FOS GRIK1 GRIK1 NXPH1 CHN1 NRGN TSHZ2 ASIC2 KCNIP4 ZNF804B HS3ST4 KIAA1217 ADRA1A BCL11B KCNIP4 MEG3 TSHZ2 IL1RAPL2 PHACTR1 NRG1 MAP1B SNAP25 FTH1 CALM1 PTGDS NRGN DHFR IDS SNAP25 CHN1 TBC1D3P1-DHX40P1 RPL32 LHFPL3 PTPRZ1 LHFPL3 PCDH15 GPR17 BCAS1 CLDN5 B2M PLXDC2 C10orf11 PLP1 CTNNA3 MBP PLP1 SLC1A2 GPC5 CLU SLC1A3 SLC1A2 PTGDS
Mean expression in group
0
2

**D**

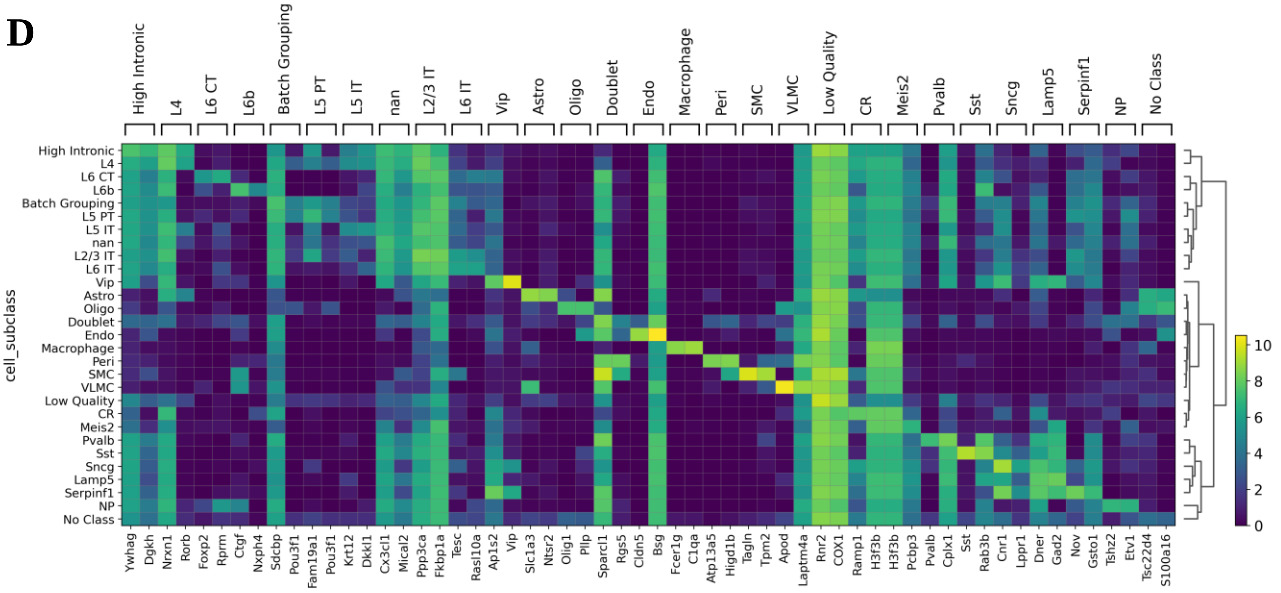
High Intronic
L4
L6 CT
L6b
Batch Grouping
L5 PT
L5 IT
nan
L2/3 IT
L6 IT
Vip
Astro
Oligo
Doublet
Endo
Macrophage
Peri
SMC
VLMC
Low Quality
CR
Meis2
Pvalb
Sst
Sncg
Lamp5
Serpinf1
NP
No Class
cell_subclass
Ywhag Dgkh Nrxn1 Rorb Foxp2 Rprm Ctgf Nxph4 Sdcbp Pou3f1 Fam19a1 Pou3f1 Krt12 Dkkl1 Cx3cl1 Mical2 Ppp3ca Fkbp1a Tesc Rasl10a Ap1s2 Vip Slc1a3 Ntsr2 Olig1 Plp Sparcl1 Rgs5 Cldn5 Bsg Fcer1g C1qa Atp13a5 Higd1b Tagln Tpm2 Apod Laptm4a Rnr2 COX1 Ramp1 H3f3b H3f3b Pcbp3 Pvalb Cplx1 Sst Rab3b Cnr1 Lppr1 Dner Gad2 Nov Gsto1 Tshz2 Etv1 Tsc22d4 S100a16
0
2
4
6
8
10

**Figure S3: Preprocessing mouse brain and human dorsolateral prefrontal cortex (DLPFC) single-cell data**

**(A-B)** t-SNE map for gene expression in (A) human DLPFC and (B) mouse brain single-cell data. The identity of the 33 and 29 clusters are color-coded and presented on the right side of the plot.

**(C-D)** Heatmaps for average log-normalized gene expression in the top 2 marker genes of each cell cluster in (C) human DLPFC and (D) mouse brain tissues. Hierarchical clustering was performed for the 33 and 29 cell types based on expression profiles of the presented genes.

**A**

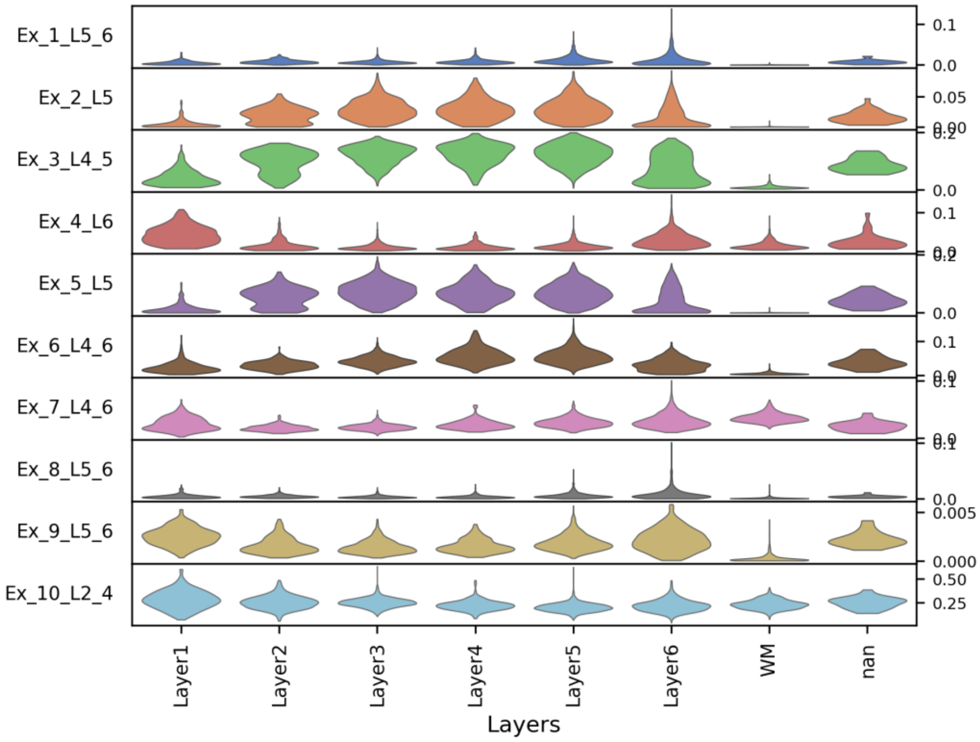

Ex_1_L5_6
Ex_2_L5
Ex_3_L4_5
Ex_4_L6
Ex_5_L5
Ex_6_L4_6
Ex_7_L4_6
Ex_8_L5_6
Ex_9_L5_6
Ex_10_L2_4
Layer1
Layer2
Layer3
Layer4
Layer5
Layer6
WM
nan
Layers


**B**

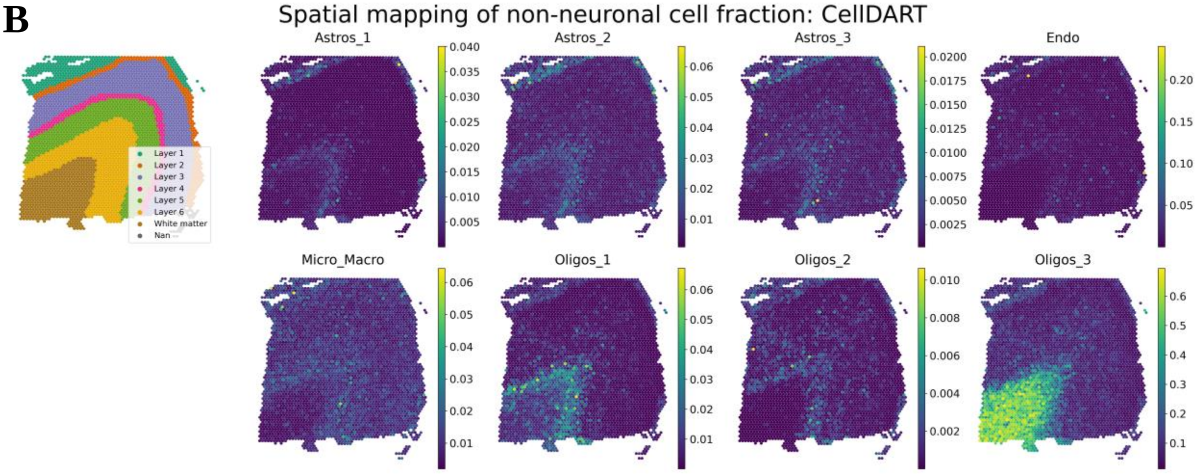

Spatial mapping of non-neuronal cell fraction: CellDART
Layer 1
Layer 2
Layer 3
Layer 4
Layer 5
Layer 6
White matter
Nan
Astros_1
Astros_2
Astros_3
Endo
Micro_Macro
Oligos_1
Oligos_2
Oligos_3

**Figure S4: Discovery of spatial cell localization patterns by CellDART**

**(A)** A stacked violin plot for the layer-specific excitatory neuron fraction predicted by CellDART. The distribution of a specific excitatory neuron fraction in the spots corresponding to one of the cortical layers (from layer 1 to 6), white matter (WM), or non-specified spots (nan) was presented in each row.

**(B)** Spatial mapping of glial cells [astrocytes (Astros) and oligodendrocytes (Oligos)], endothelial cells (Endo), and microglia or macrophages (Micro_Macro) fraction predicted by CellDART. The figure in the top left corner shows the layer annotation for each spatial spot. The layer consists of cortical layer 1 to 6 and white matter. 'Nan' represents the spot without the layer information. Colormaps present the maximum and minimum values for the corresponding cell fraction.

**A**

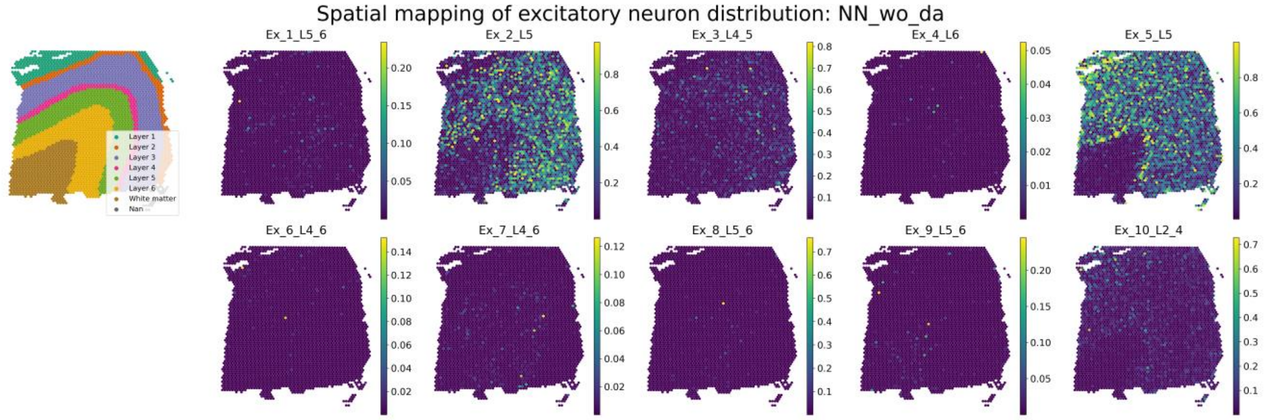

Spatial mapping of excitatory neuron distribution: NN_wo_da
Layer 1
Layer 2
Layer 3
Layer 4
Layer 5
Layer 6
White matter
Ex_1_L5_6
Ex_2_L5
Ex_3_L4_5
Ex_4_L6
Ex_5_L5
Ex_6_L4_6
Ex_7_L4_6
Ex_8_L5_6
Ex_9_L5_6
Ex_10_L2_4


**B**

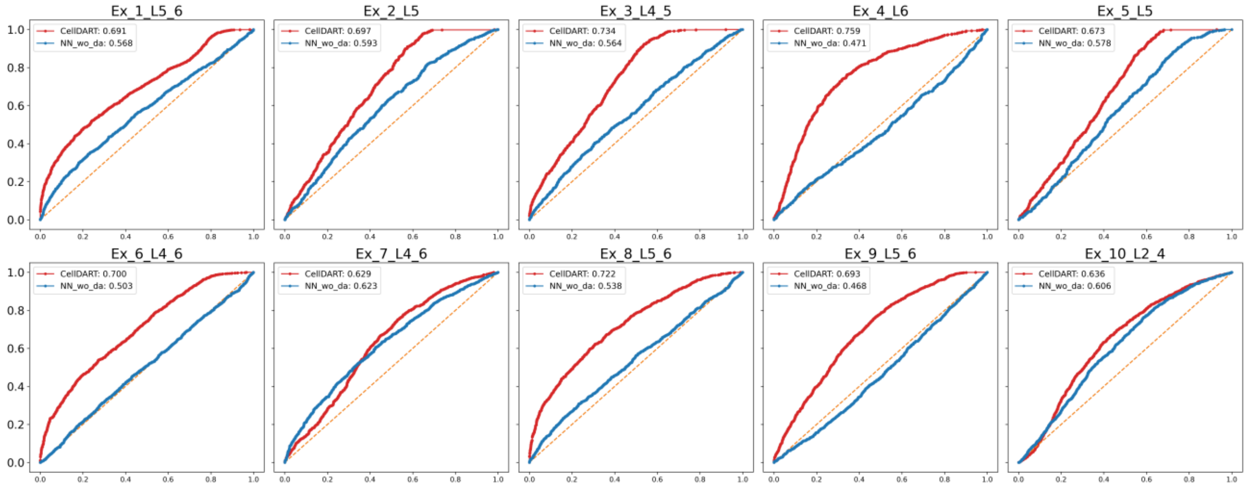

Ex_1_L5_6
CellDART: 0.691
NN_wo_da: 0.568
Ex_2_L5
CellDART: 0.697
NN_wo_da: 0.593
Ex_3_L4_5
CellDART: 0.734
NN_wo_da: 0.564
Ex_4_L6
CellDART: 0.759
NN_wo_da: 0.471
Ex_5_L5
CellDART: 0.673
NN_wo_da: 0.578
Ex_6_L4_6
CellDART: 0.700
NN_wo_da: 0.503
Ex_7_L4_6
CellDART: 0.629
NN_wo_da: 0.623
Ex_8_L5_6
CellDART: 0.722
NN_wo_da: 0.538
Ex_9_L5_6
CellDART: 0.693
NN_wo_da: 0.468
Ex_10_L2_4
CellDART: 0.636
NN_wo_da: 0.606

**Figure S5: Comparison of CellDART with neural network without domain adaptation**

**(A)** Spatial mapping of 10 layer-specific excitatory neuron fraction predicted by neural network model without domain adaptation (NN_wo_da). The figure in the top left corner shows the layer annotation for each spatial spot. The layer consists of cortical layer 1 to 6 and white matter. 'Nan' represents the spot without the layer information. Colormaps present the maximum and minimum values for the corresponding cell fraction.

**(B)** Receiver operating characteristic (ROC) analysis for predicting the layer-specific distribution of excitatory neurons. The CellDART and NN_wo_da were compared by means of the area under the curve (AUC). The ROC curves for CellDART and NN_wo_da are color-coded, and AUC values are presented in the lower right corner of each plot.

**A**

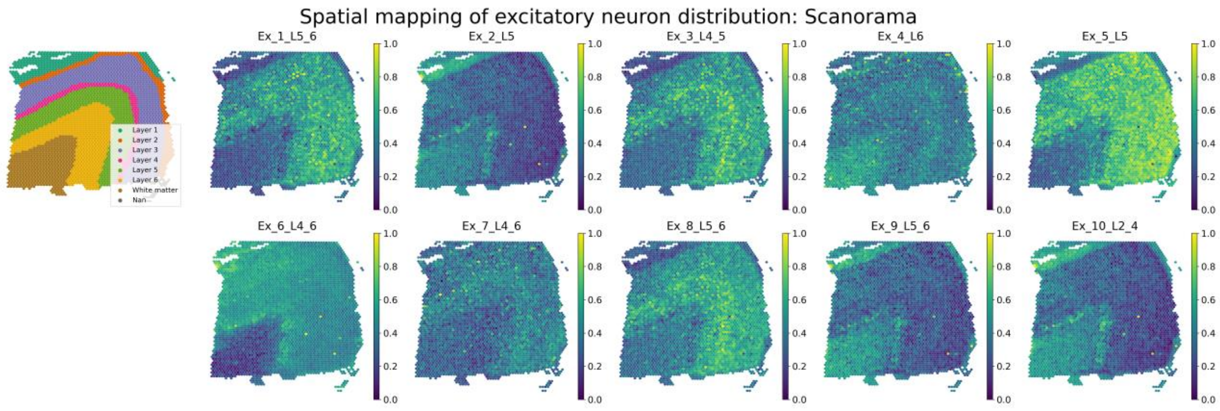
Spatial mapping of excitatory neuron distribution: Scanorama
Ex_1_L5_6
Ex_2_L5
Ex_3_L4_5
Ex_4_L6
Ex_5_L5
Ex_6_L4_6
Ex_7_L4_6
Ex_8_L5_6
Ex_9_L5_6
Ex_10_L2_4
Layer 1
Layer 2
Layer 3
Layer 4
Layer 5
Layer 6
White matter
Nan
0.0
0.2
0.4
0.6
0.8
1.0

**B**

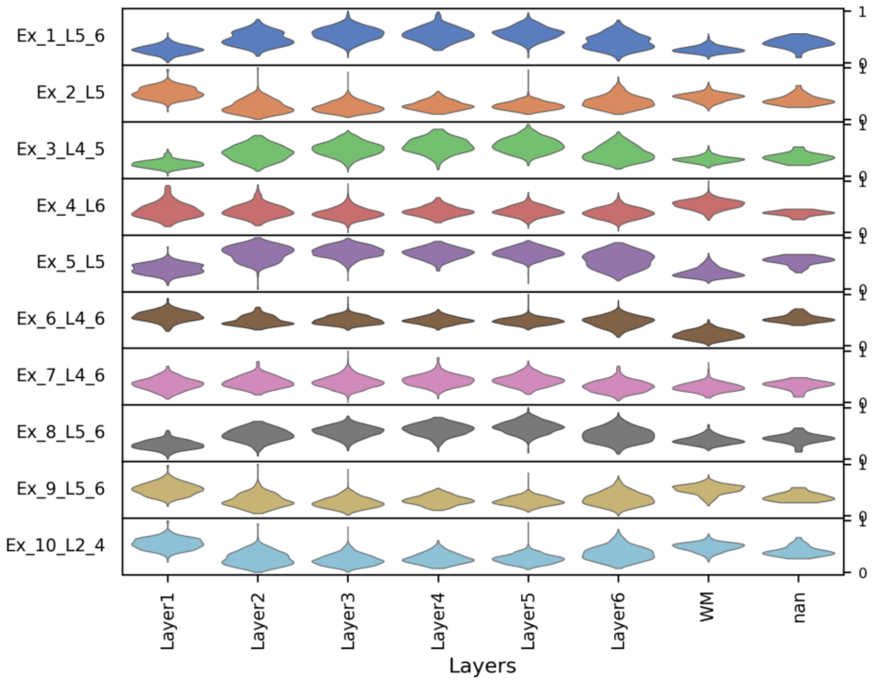
Ex_1_L5_6
Ex_2_L5
Ex_3_L4_5
Ex_4_L6
Ex_5_L5
Ex_6_L4_6
Ex_7_L4_6
Ex_8_L5_6
Ex_9_L5_6
Ex_10_L2_4
0
1
Layer1
Layer2
Layer3
Layer4
Layer5
Layer6
WM
nan
Layers

**C**

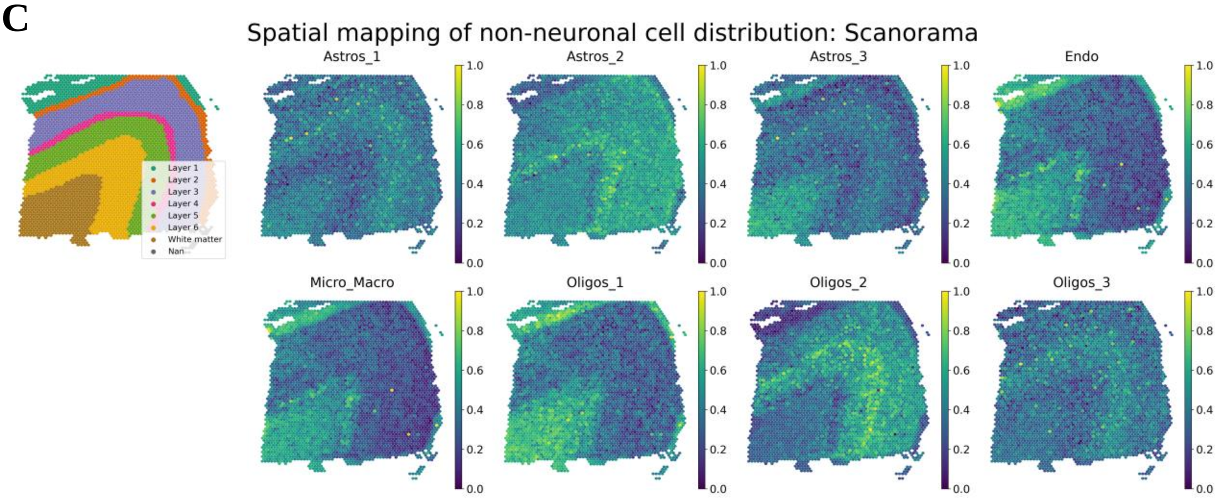
Spatial mapping of non-neuronal cell distribution: Scanorama
Astros_1
Astros_2
Astros_3
Endo
Micro_Macro
Oligos_1
Oligos_2
Oligos_3
Layer 1
Layer 2
Layer 3
Layer 4
Layer 5
Layer 6
White matter
Nan
0.0
0.2
0.4
0.6
0.8
1.0

**Figure S6: Estimation of spatial cell proportion by Scanorama**

**(A)** Spatial mapping of 10 layer-specific excitatory neuron fraction predicted by Scanorama. The figure in the top left corner shows the layer annotation for each spatial spot. The layer consists of cortical layer 1 to 6 and white matter. 'Nan' represents the spot without the layer information. Colormaps present the maximum and minimum values for the corresponding cell fraction.

**(B)** A stacked violin plot for the layer-specific excitatory neuron fraction predicted by Scanorama. The distribution of a specific excitatory neuron fraction in the spots corresponding to one of the cortical layers (from layer 1 to 6), white matter (WM), or non-specified spots (nan) was presented in each row.

**(C)** Spatial mapping of non-neuronal cells (Astros, Oligos, Endo, and Micro_Macro) fraction predicted by Scanorama. Colormaps present the maximum and minimum values for the corresponding cell fraction.

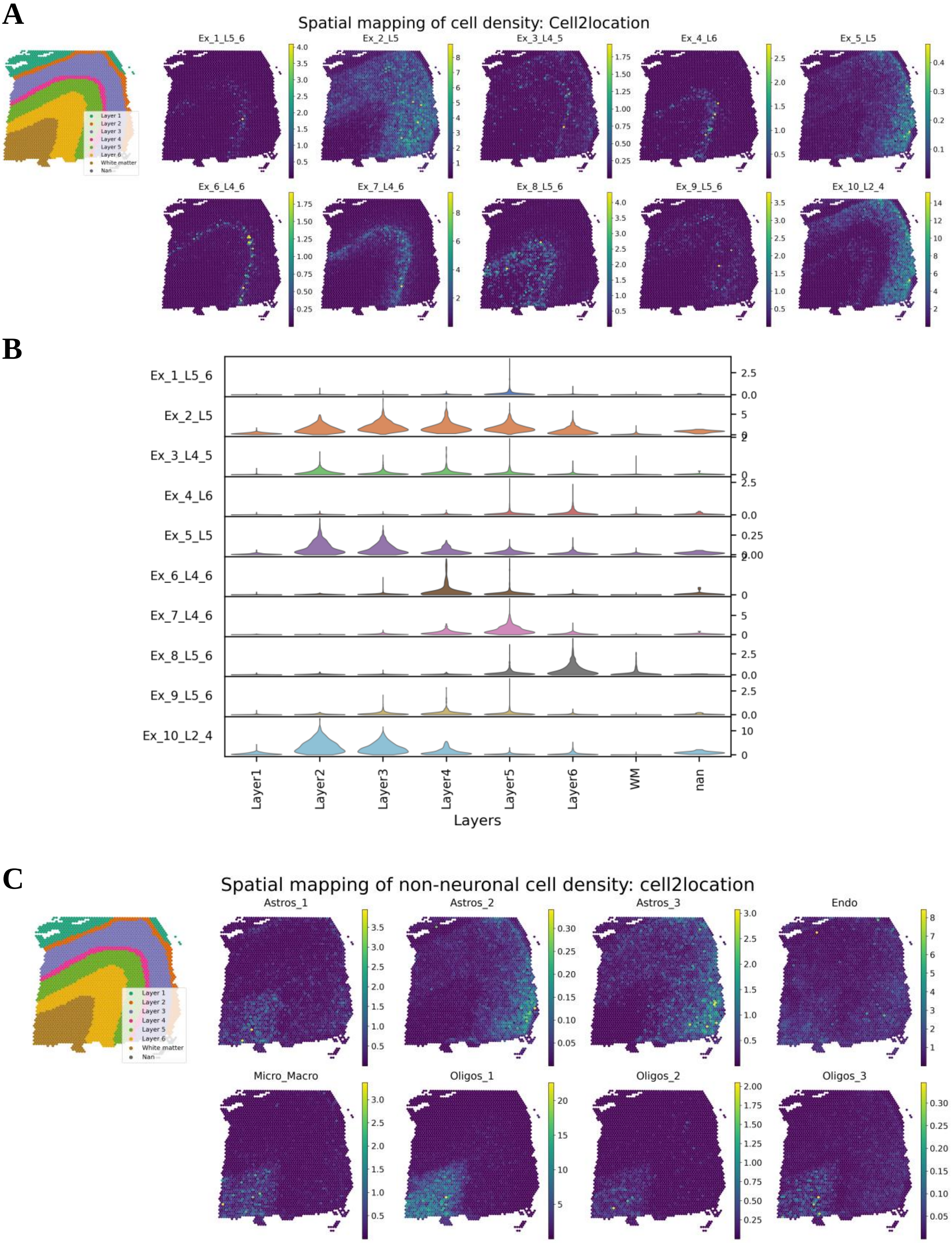

A
Spatial mapping of cell density: Cell2location
Ex_1_L5_6
Ex_2_L5
Ex_3_L4_5
Ex_4_L6
Ex_5_L5
Ex_6_L4_6
Ex_7_L4_6
Ex_8_L5_6
Ex_9_L5_6
Ex_10_L2_4
Layer 1
Layer 2
Layer 3
Layer 4
Layer 5
Layer 6
White matter
Nan
B
Layer1
Layer2
Layer3
Layer4
Layer5
Layer6
WM
nan
Layers
C
Spatial mapping of non-neuronal cell density: cell2location
Astros_1
Astros_2
Astros_3
Endo
Micro_Macro
Oligos_1
Oligos_2
Oligos_3

**Figure S7: Prediction of spatial cell fraction by Cell2location**

**(A)** Spatial mapping of 10 layer-specific excitatory neuron fraction predicted by Cell2location. The figure in the top left corner shows the layer annotation for each spatial spot. The layer consists of cortical layer 1 to 6 and white matter. ‘Nan’ represents the spot without the layer information. Colormaps present the maximum and minimum values for the corresponding cell fraction.

**(B)** A stacked violin plot for the layer-specific excitatory neuron fraction predicted by Cell2location. The distribution of a specific excitatory neuron fraction in the spots corresponding to one of the cortical layers (from layer 1 to 6), white matter (WM), or non-specified spots (nan) was presented in each row.

**(C)** Spatial mapping of non-neuronal cells (Astros, Oligos, Endo, and Micro_Macro) fraction predicted by Cell2location. Colormaps present the maximum and minimum values for the corresponding cell fraction.

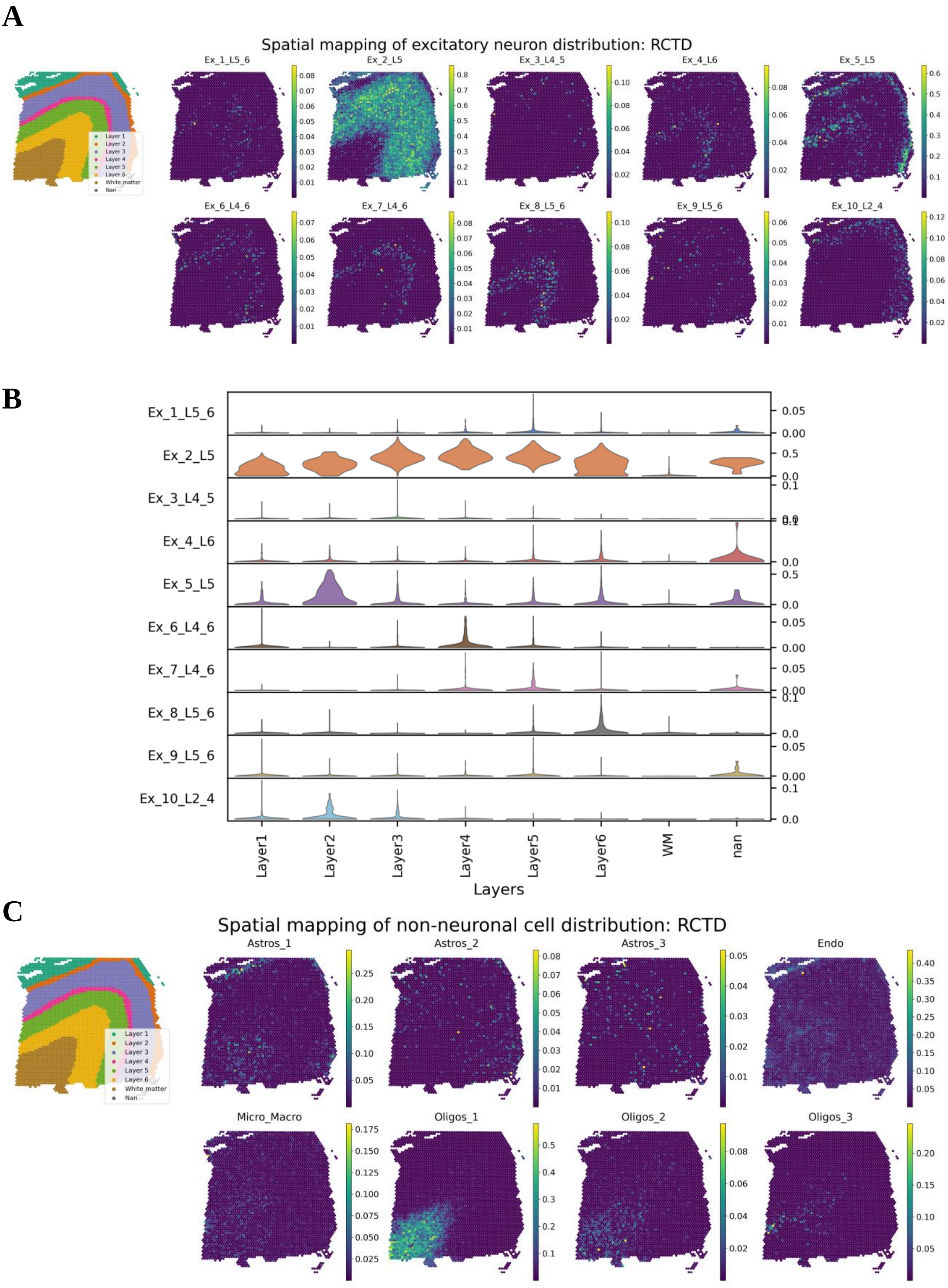
A
Spatial mapping of excitatory neuron distribution: RCTD
Layer 1
Layer 2
Layer 3
Layer 4
Layer 5
Layer 6
White matter
Nan
Ex_1_L5_6
Ex_2_L5
Ex_3_L4_5
Ex_4_L6
Ex_5_L5
Ex_6_L4_6
Ex_7_L4_6
Ex_8_L5_6
Ex_9_L5_6
Ex_10_L2_4
B
Ex_1_L5_6
Ex_2_L5
Ex_3_L4_5
Ex_4_L6
Ex_5_L5
Ex_6_L4_6
Ex_7_L4_6
Ex_8_L5_6
Ex_9_L5_6
Ex_10_L2_4
Layer1
Layer2
Layer3
Layer4
Layer5
Layer6
WM
nan
Layers
C
Spatial mapping of non-neuronal cell distribution: RCTD
Layer 1
Layer 2
Layer 3
Layer 4
Layer 5
Layer 6
White matter
Nan
Astros_1
Astros_2
Astros_3
Endo
Micro_Macro
Oligos_1
Oligos_2
Oligos_3

**Figure S8: Estimation of spatial cell fraction by RCTD**

**(A)** Spatial mapping of 10 layer-specific excitatory neuron fraction predicted by RCTD. The figure in the top left corner shows the layer annotation for each spatial spot. The layer consists of cortical layer 1 to 6 and white matter. 'Nan' represents the spot without the layer information. Colormaps present the maximum and minimum values for the corresponding cell fraction.

**(B)** A stacked violin plot for the layer-specific excitatory neuron fraction predicted by RCTD. The distribution of a specific excitatory neuron fraction in the spots corresponding to one of the cortical layers (from layer 1 to 6), white matter (WM), or non-specified spots (nan) was presented in each row.

**(C)** Spatial mapping of non-neuronal cells (Astros, Oligos, Endo, and Micro_Macro) fraction estimated by RCTD. Colormaps present the maximum and minimum values for the corresponding cell fraction.

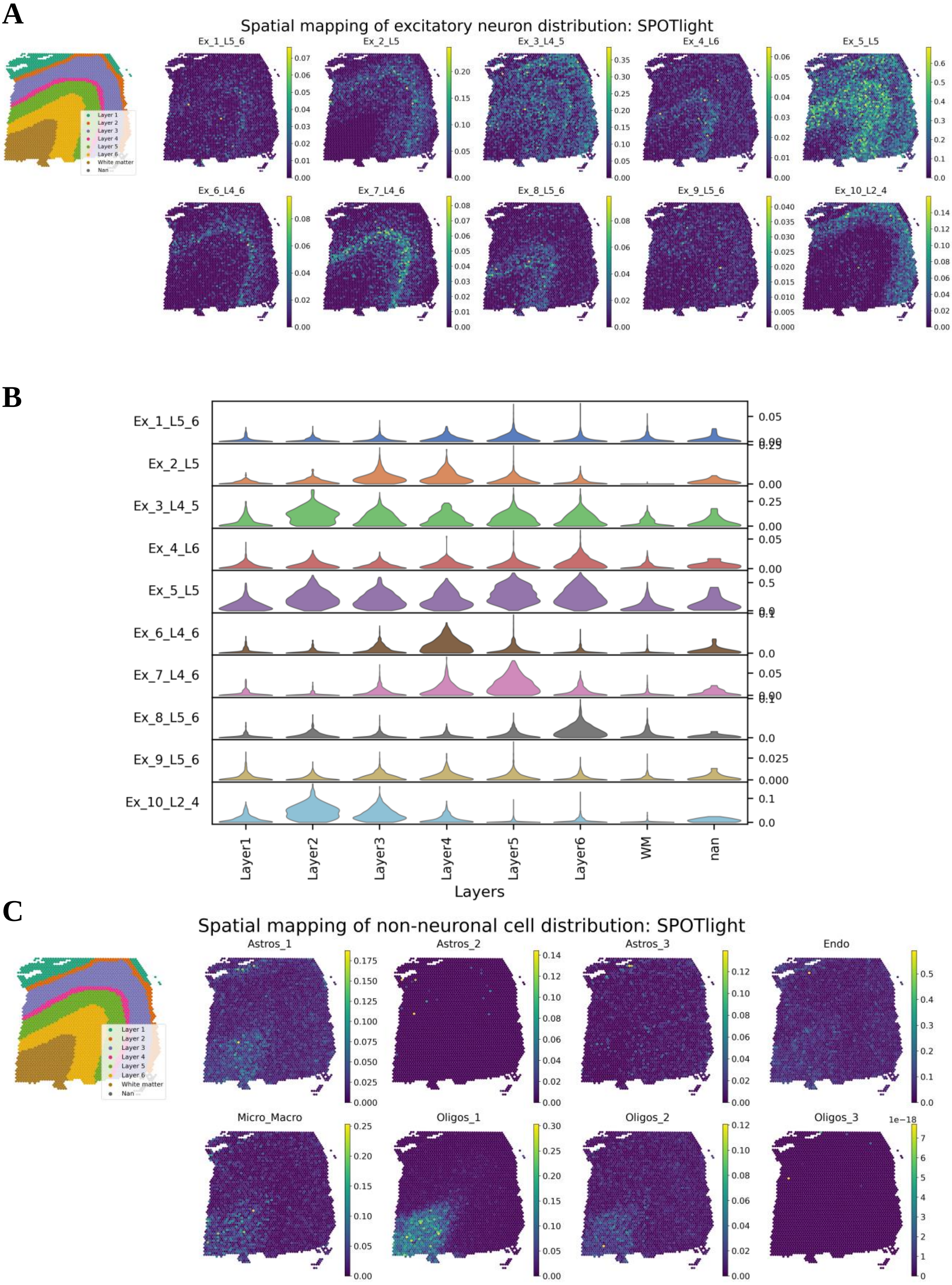
A
Spatial mapping of excitatory neuron distribution: SPOTlight
Layer 1
Layer 2
Layer 3
Layer 4
Layer 5
Layer 6
White matter
Nan
Ex_1_L5_6
Ex_2_L5
Ex_3_L4_5
Ex_4_L6
Ex_5_L5
Ex_6_L4_6
Ex_7_L4_6
Ex_8_L5_6
Ex_9_L5_6
Ex_10_L2_4
B
Ex_1_L5_6
Ex_2_L5
Ex_3_L4_5
Ex_4_L6
Ex_5_L5
Ex_6_L4_6
Ex_7_L4_6
Ex_8_L5_6
Ex_9_L5_6
Ex_10_L2_4
Layer1
Layer2
Layer3
Layer4
Layer5
Layer6
WM
nan
Layers
C
Spatial mapping of non-neuronal cell distribution: SPOTlight
Layer 1
Layer 2
Layer 3
Layer 4
Layer 5
Layer 6
White matter
Nan
Astros_1
Astros_2
Astros_3
Endo
Micro_Macro
Oligos_1
Oligos_2
Oligos_3
1e−18

**Figure S9: Prediction of spatial cell proportion by SPOTlight**

**(A)** Spatial mapping of 10 layer-specific excitatory neuron fraction predicted by SPOTlight. The figure in the top left corner shows the layer annotation for each spatial spot. The layer consists of cortical layer 1 to 6 and white matter. ‘Nan’ represents the spot without the layer information. Colormaps present the maximum and minimum values for the corresponding cell fraction.

**(B)** A stacked violin plot for the layer-specific excitatory neuron fraction predicted by SPOTlight. The distribution of a specific excitatory neuron fraction in the spots corresponding to one of the cortical layers (from layer 1 to 6), white matter (WM), or non-specified spots (nan) was presented in each row.

**(C)** Spatial mapping of non-neuronal cells (Astros, Oligos, Endo, and Micro_Macro) fraction estimated by SPOTlight. Colormaps present the maximum and minimum values for the corresponding cell fraction.

**A**

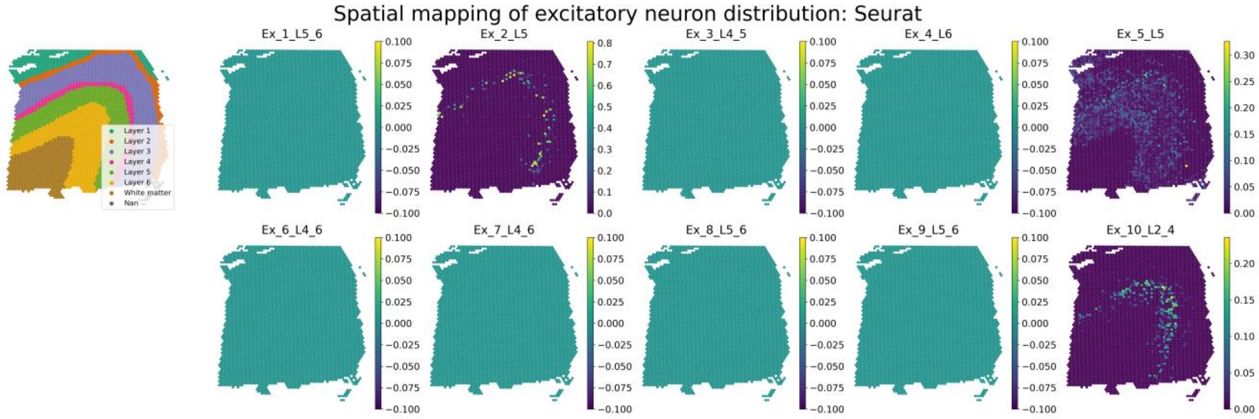

Spatial mapping of excitatory neuron distribution: Seurat
Layer 1
Layer 2
Layer 3
Layer 4
Layer 5
Layer 6
White matter
Nan
Ex_1_L5_6
Ex_2_L5
Ex_3_L4_5
Ex_4_L6
Ex_5_L5
Ex_6_L4_6
Ex_7_L4_6
Ex_8_L5_6
Ex_9_L5_6
Ex_10_L2_4


**B**

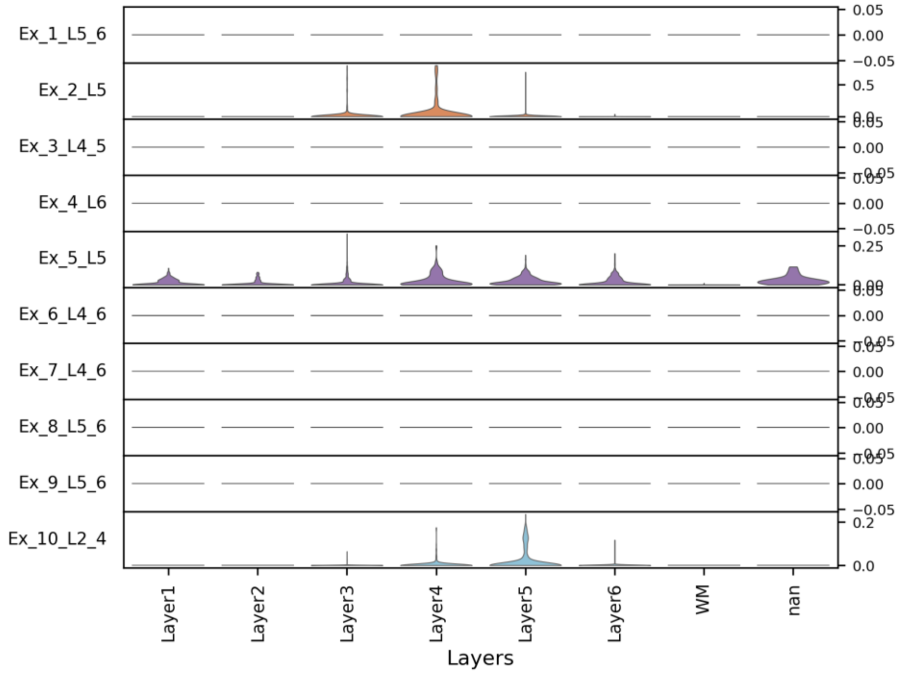

Ex_1_L5_6
Ex_2_L5
Ex_3_L4_5
Ex_4_L6
Ex_5_L5
Ex_6_L4_6
Ex_7_L4_6
Ex_8_L5_6
Ex_9_L5_6
Ex_10_L2_4
Layer1
Layer2
Layer3
Layer4
Layer5
Layer6
WM
nan
Layers


**C**

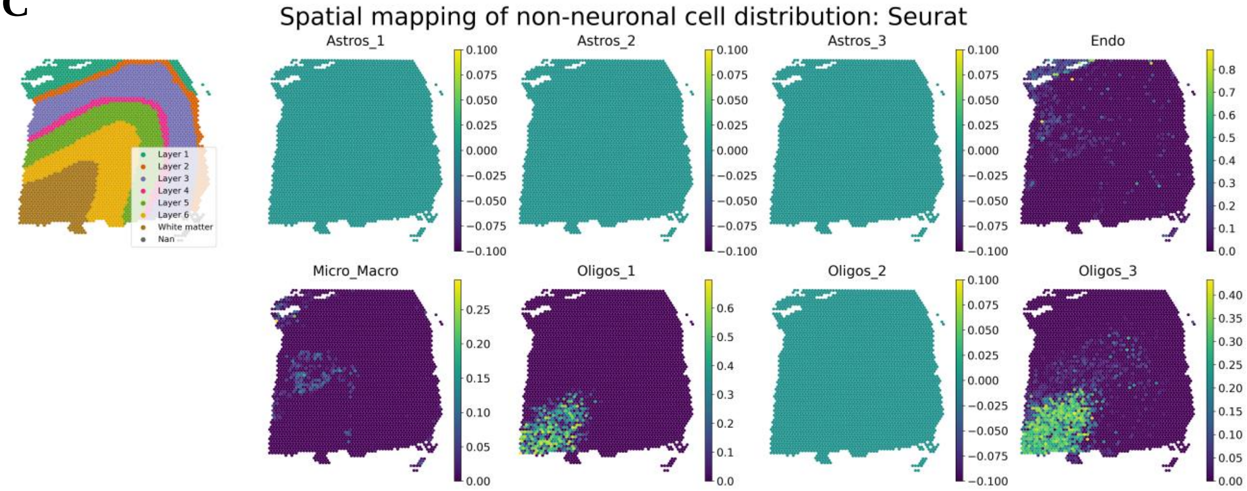

Spatial mapping of non-neuronal cell distribution: Seurat
Layer 1
Layer 2
Layer 3
Layer 4
Layer 5
Layer 6
White matter
Nan
Astros_1
Astros_2
Astros_3
Endo
Micro_Macro
Oligos_1
Oligos_2
Oligos_3

**Figure S10: Prediction of spatial cell proportion by Seurat**

**(A)** Spatial mapping of 10 layer-specific excitatory neuron fraction predicted by Seurat. The figure in the top left corner shows the layer annotation for each spatial spot. The layer consists of cortical layer 1 to 6 and white matter. 'Nan' represents the spot without the layer information. Colormaps present the maximum and minimum values for the corresponding cell fraction.

**(B)** A stacked violin plot for the layer-specific excitatory neuron fraction predicted by Seurat. The distribution of a specific excitatory neuron fraction in the spots corresponding to one of the cortical layers (from layer 1 to 6), white matter (WM), or non-specified spots (nan) was presented in each row.

**(C)** Spatial mapping of non-neuronal cells (Astros, Oligos, Endo, and Micro_Macro) fraction estimated by Seurat. Colormaps present the maximum and minimum values for the corresponding cell fraction.

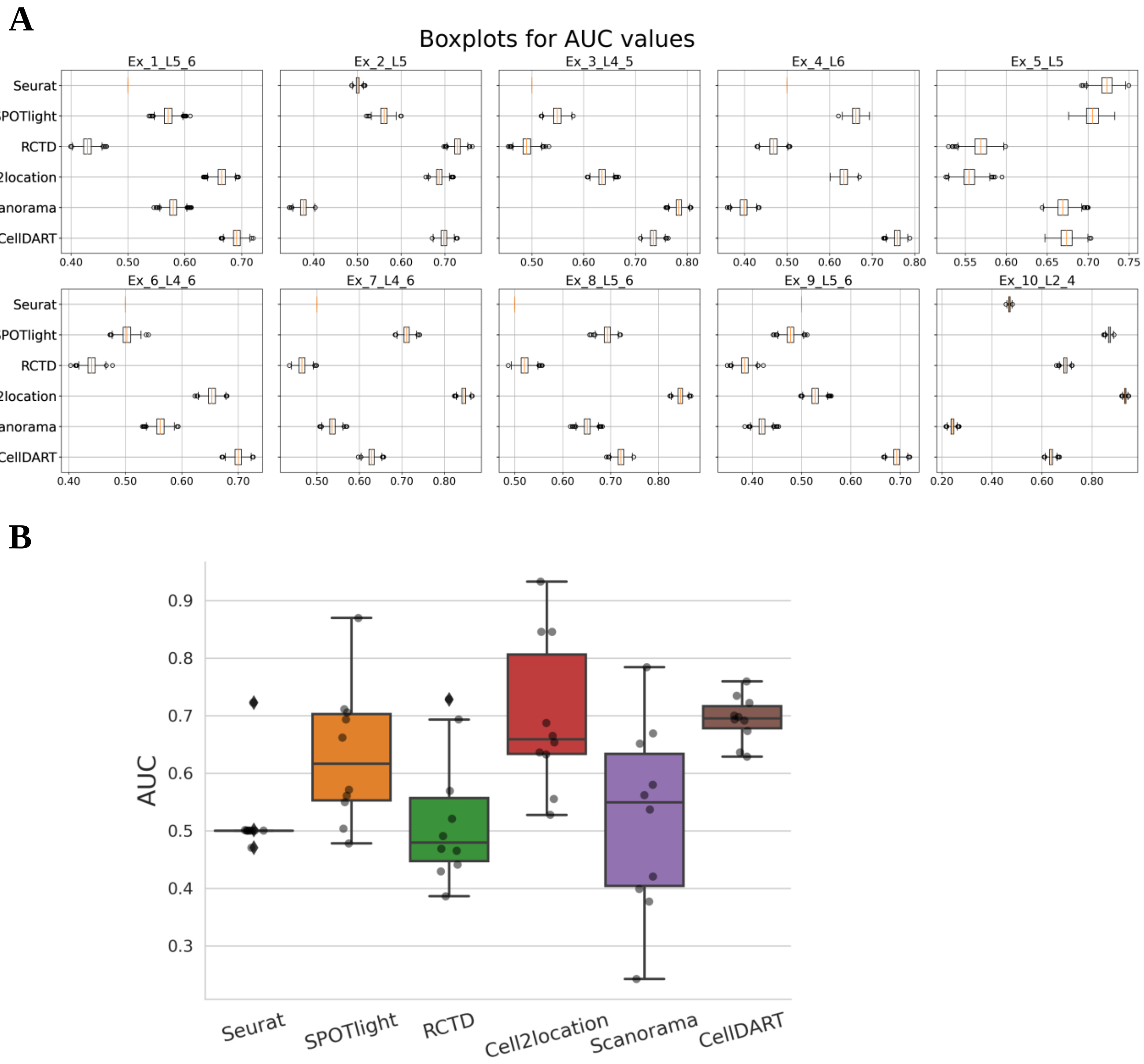

A
Boxplots for AUC values
Ex_1_L5_6
Ex_2_L5
Ex_3_L4_5
Ex_4_L6
Ex_5_L5
Ex_6_L4_6
Ex_7_L4_6
Ex_8_L5_6
Ex_9_L5_6
Ex_10_L2_4
Seurat
SPOTlight
RCTD
Cell2location
Scanorama
CellDART
B
AUC
Seurat
SPOTlight
RCTD
Cell2location
Scanorama
CellDART

**Figure S11: Comparison of AUC values for predicting layer-specific distribution of excitatory neurons in DLPFC 151673 dataset.**

**(A)** The cell fraction values and layer labels of spatial spots for 10 excitatory neurons were bootstrapped and the null distribution for the AUC values were obtained. The results derived from six different tools, CellDART, Scanorma, Cell2location, RCTD, SPOTlight, and Seurat were visualized with boxplots.

**(B)** The AUC values for the 10 excitatory neurons were collected and their distribution across six computational tools was visualized with a boxplot. Grey points in the boxplot represent the individual AUC values for each excitatory neuron subtype.

**A**

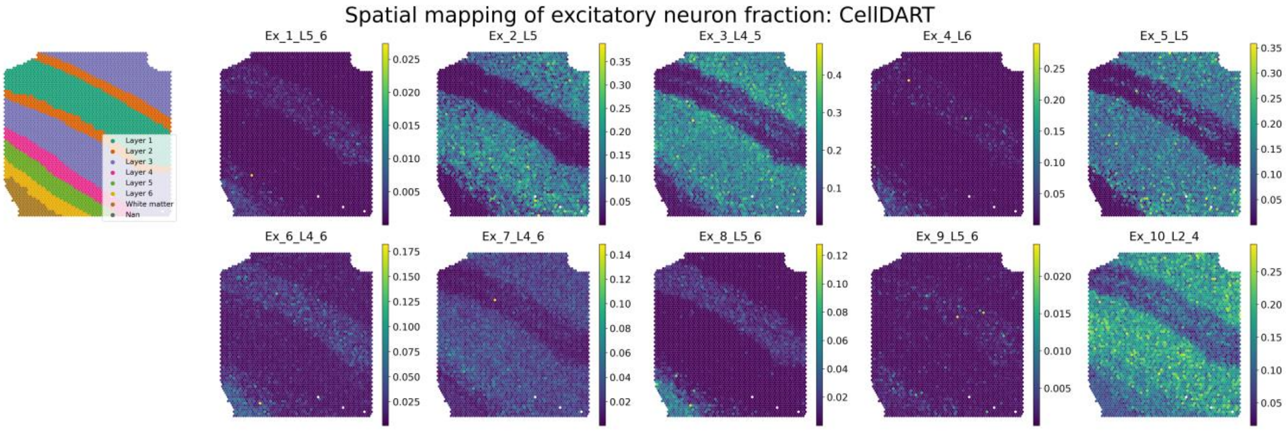
Spatial mapping of excitatory neuron fraction: CellDART
Ex_1_L5_6
Ex_2_L5
Ex_3_L4_5
Ex_4_L6
Ex_5_L5
Ex_6_L4_6
Ex_7_L4_6
Ex_8_L5_6
Ex_9_L5_6
Ex_10_L2_4
Layer 1
Layer 2
Layer 3
Layer 4
Layer 5
Layer 6
White matter
Nan

**B**

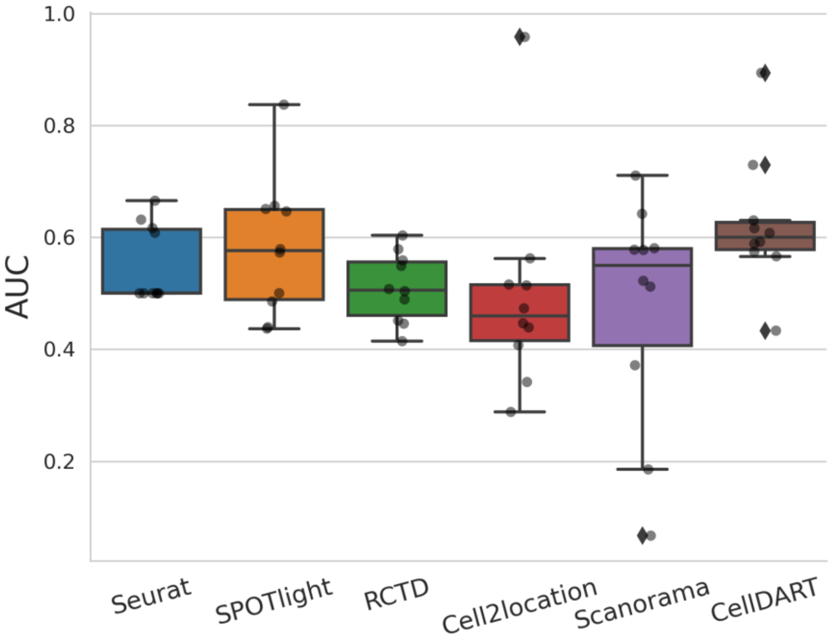
AUC
1.0
0.8
0.6
0.4
0.2
Seurat
SPOTlight
RCTD
Cell2location
Scanorama
CellDART

**C**

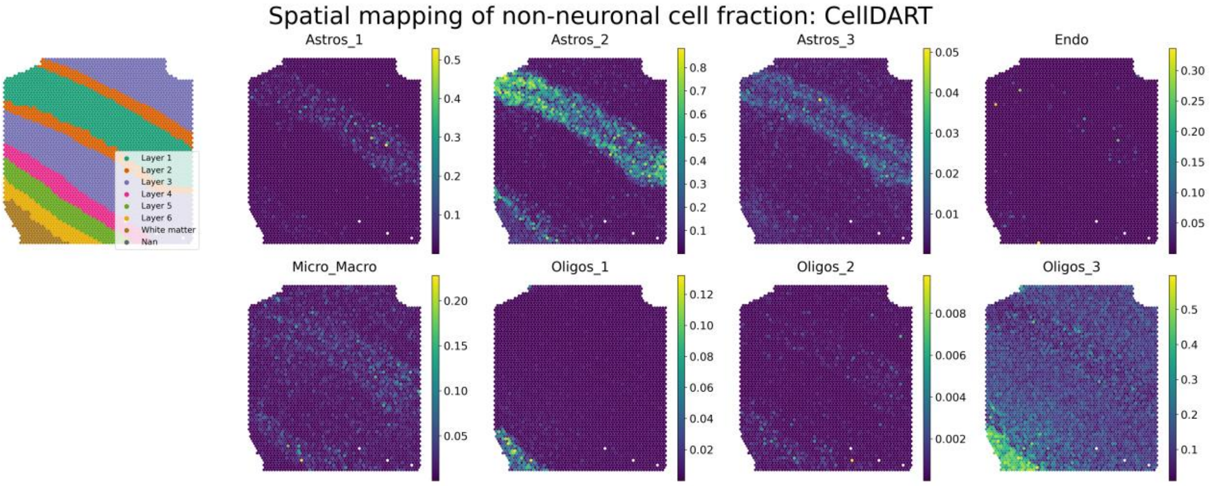
Spatial mapping of non-neuronal cell fraction: CellDART
Astros_1
Astros_2
Astros_3
Endo
Micro_Macro
Oligos_1
Oligos_2
Oligos_3
Layer 1
Layer 2
Layer 3
Layer 4
Layer 5
Layer 6
White matter
Nan

**Figure S12: Validation of CellDART in DLPFC 151509 dataset.**

(A) Spatial mapping of 10 layer-specific excitatory neuron fraction predicted by CellDART. The figure in the top left corner shows the layer annotation for each spatial spot. The layer consists of cortical layer 1 to 6 and white matter. ‘Nan’ represents the spot without the layer information. Colormaps present the maximum and minimum values for the corresponding cell fraction.

(B) Comparison of AUC values for predicting layer-specific distribution of excitatory neurons across six computational tools: CellDART, Scanorama, Cell2location, RCTD, SPOTlight and Seurat. The AUC values for the 10 excitatory neurons were collected and their distribution in each toolkits was visualized with a boxplot. Grey points in the boxplot represent the individual AUC values for each excitatory neuron subtype.

(C) Spatial mapping of non-neuronal cells (Astros, Oligos, Endo, and Micro_Macro) fraction predicted by CellDART. Colormaps present the maximum and minimum values for the corresponding cell fraction.

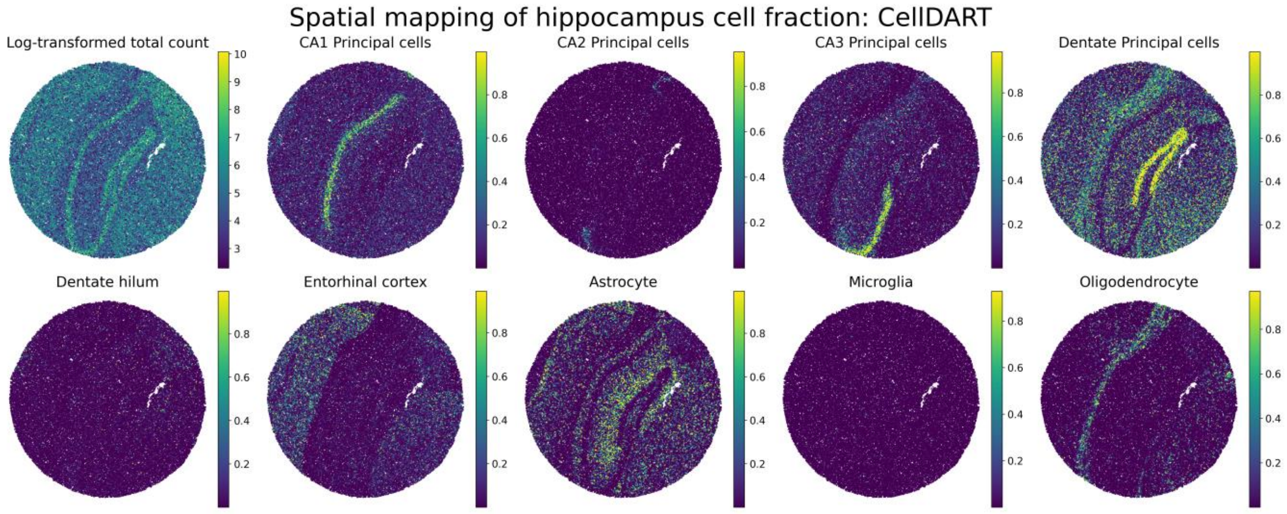


**Figure S13: Application of CellDART in mouse hippocampus Slide-seq data.**

Spatial mapping of 9 hippocampal cell fraction on the Slide-seq data. The figure in the top left corner shows the log-transformed total count in each bead. Colormaps present the maximum and minimum values for the corresponding log-transformed count or cell fraction.

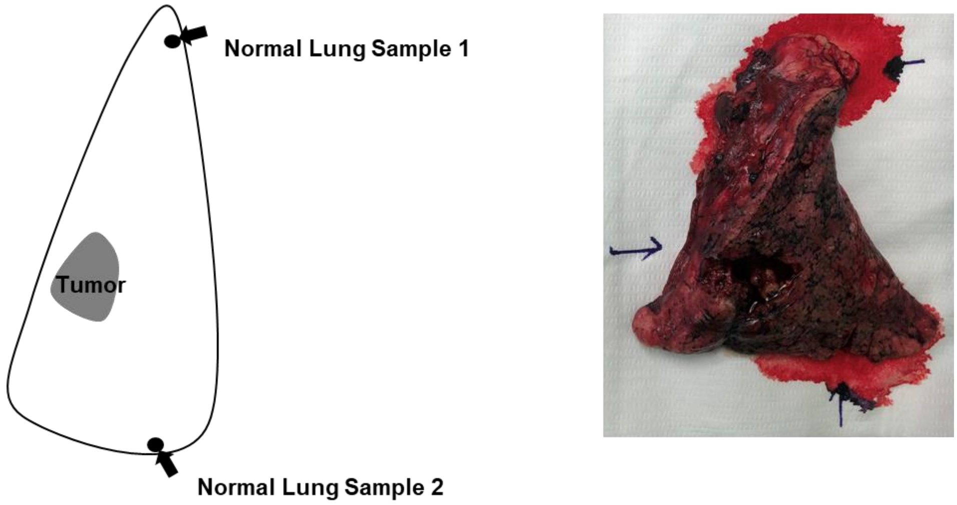


**Figure S14: Gross specimen of human lung tissue 1 and 2 data**

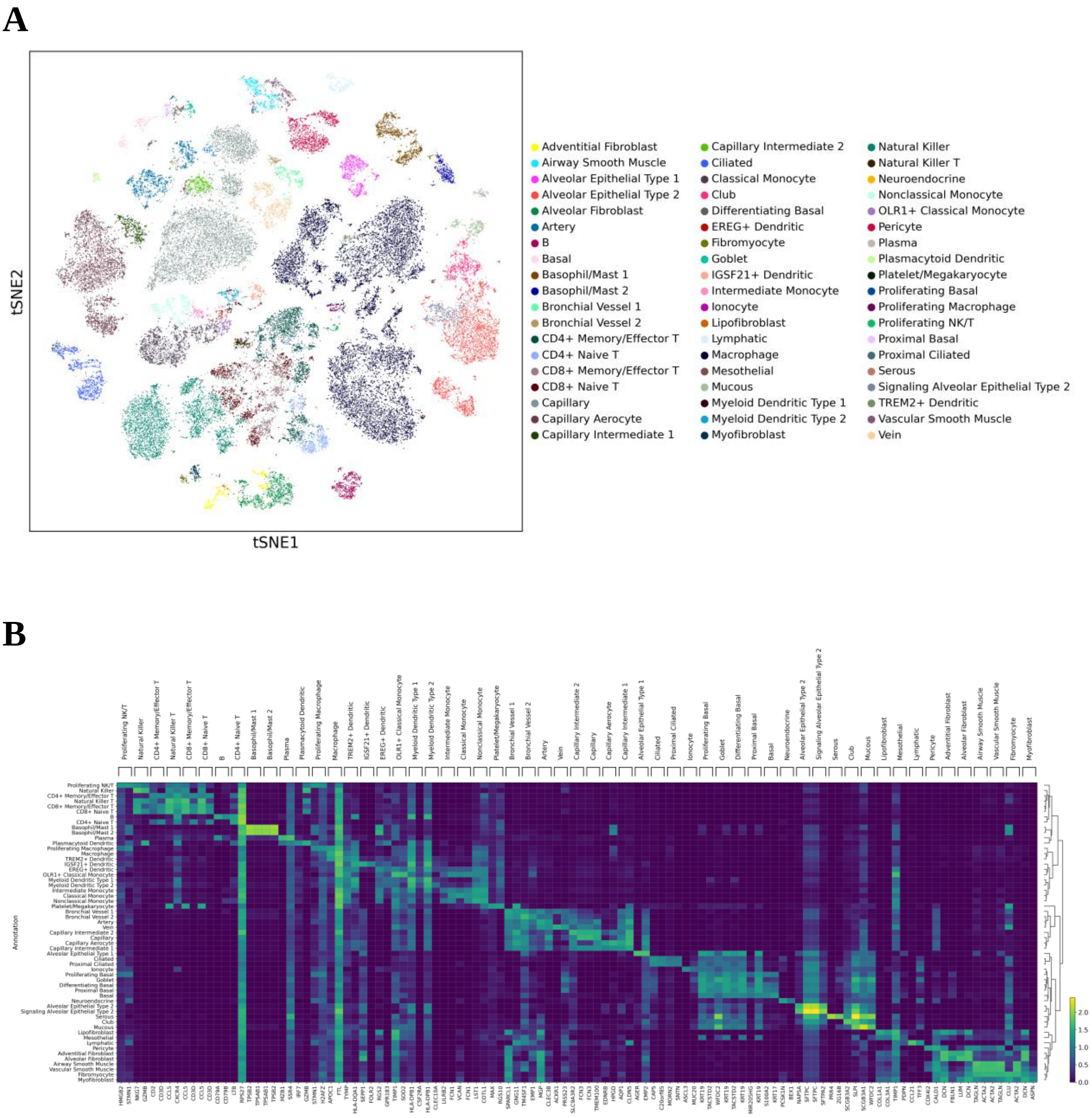


**Figure S15: Preprocessing human lung single-cell data**

**(A)** t-SNE map for gene expression in lung single-cell data. The identity of the 57 clusters are color-coded and presented on the right side of the plot.

**(B)** A heatmap for average log-normalized gene expression in the top 2 marker genes of each cell cluster. Hierarchical clustering was performed for the 57 cell types based on expression profiles of the presented genes.
