## Supplementary Table 1 for "CellDART: Cell type inference by domain adaptation of single-cell and spatial transcriptomic data"

| **Table 1** | The top 100 marker genes for each cell cluster in human normal dorsolateral prefrontal cortex single-cell dataset. The genes were ranked based on the Benjamini-Hochberg adjusted p-values. The name of the cells are based on the metadata provided by the paper (Maynard, K.R. et al. Transcriptome-scale spatial gene expression in the human dorsolateral prefrontal cortex. *Nat Neurosci* **24**, 425-436 (2021)). | | | |
|---|---|---|---|---|
| Number | Genes | Adjusted p-value | Log fold change | Cluster |
| 0 | CLU | 5.91E-134 | 4.279877 | Astros_1 |
| 1 | SLC1A3 | 6.88E-128 | 5.293424 | Astros_1 |
| 2 | APOE | 3.46E-121 | 5.169107 | Astros_1 |
| 3 | GPC5 | 7.11E-120 | 4.2819195 | Astros_1 |
| 4 | GLUL | 2.51E-116 | 4.4119196 | Astros_1 |
| 5 | ANGPTL4 | 5.73E-116 | 7.781683 | Astros_1 |
| 6 | DTNA | 1.01E-115 | 3.1554394 | Astros_1 |
| 7 | AQP4 | 1.55E-115 | 6.1355166 | Astros_1 |
| 8 | SLC1A2 | 1.40E-111 | 4.5177894 | Astros_1 |
| 9 | GJA1 | 9.69E-108 | 5.9901986 | Astros_1 |
| 10 | CST3 | 1.44E-105 | 4.1773524 | Astros_1 |
| 11 | MACF1 | 1.46E-104 | 2.6622176 | Astros_1 |
| 12 | PTGDS | 7.97E-104 | 3.6867716 | Astros_1 |
| 13 | ATP1B2 | 7.11E-103 | 4.5717854 | Astros_1 |
| 14 | GFAP | 4.47E-102 | 4.9359474 | Astros_1 |
| 15 | LRP4 | 1.56E-100 | 2.5164092 | Astros_1 |
| 16 | FTL | 2.83E-98 | 3.0015345 | Astros_1 |
| 17 | MT2A | 1.07E-94 | 4.976446 | Astros_1 |
| 18 | DST | 4.77E-93 | 2.4389877 | Astros_1 |
| 19 | ATP1A2 | 7.18E-92 | 4.724029 | Astros_1 |
| 20 | NPAS3 | 2.81E-90 | 2.7396767 | Astros_1 |
| 21 | F3 | 1.05E-89 | 5.5254164 | Astros_1 |
| 22 | ADGRV1 | 5.79E-88 | 3.8552217 | Astros_1 |
| 23 | RYR3 | 5.89E-88 | 3.5340269 | Astros_1 |
| 24 | MGST1 | 1.11E-87 | 3.279118 | Astros_1 |
| 25 | GPM6A | 1.27E-85 | 1.9346691 | Astros_1 |
| 26 | PITPNC1 | 2.53E-85 | 3.1302824 | Astros_1 |
| 27 | CPE | 1.69E-84 | 2.8412752 | Astros_1 |
| 28 | CADM1 | 5.55E-78 | 2.3973832 | Astros_1 |
| 29 | NEAT1 | 6.02E-78 | 3.1995096 | Astros_1 |
| 30 | ITM2C | 1.10E-75 | 3.0907598 | Astros_1 |
| 31 | HEPN1 | 2.02E-75 | 5.4121146 | Astros_1 |
| 32 | NFIA | 6.26E-75 | 2.9112587 | Astros_1 |
| 33 | AHCYL1 | 7.97E-75 | 3.254617 | Astros_1 |
| 34 | PTN | 3.09E-73 | 4.0195546 | Astros_1 |
| 35 | AGT | 1.49E-71 | 5.453191 | Astros_1 |
| 36 | NTM | 6.19E-70 | 2.0314386 | Astros_1 |
| 37 | ZBTB20 | 1.63E-67 | 2.7558603 | Astros_1 |
| 38 | TUBB2B | 3.01E-65 | 4.3452806 | Astros_1 |
| 39 | GPRC5B | 3.50E-64 | 3.6493795 | Astros_1 |
| 40 | FAM189A2 | 8.64E-64 | 6.24409 | Astros_1 |
| 41 | RNF219-AS1 | 8.12E-63 | 3.8787997 | Astros_1 |
| 42 | TPD52L1 | 2.59E-59 | 4.342192 | Astros_1 |

| | | | | |
|---|---|---|---|---|
| 43 | FTH1 | 9.54E-59 | 2.149782 | Astros_1 |
| 44 | BMPR1B | 9.96E-59 | 4.2634716 | Astros_1 |
| 45 | BCL6 | 7.32E-58 | 4.494747 | Astros_1 |
| 46 | SERPINE2 | 7.68E-58 | 3.029796 | Astros_1 |
| 47 | PARD3 | 1.25E-57 | 3.3649728 | Astros_1 |
| 48 | GPR37L1 | 1.42E-57 | 4.4299803 | Astros_1 |
| 49 | ITM2B | 9.99E-56 | 2.061388 | Astros_1 |
| 50 | ASPH | 2.44E-55 | 2.5775685 | Astros_1 |
| 51 | MT1E | 1.28E-54 | 5.1699896 | Astros_1 |
| 52 | LSAMP | 1.41E-54 | 1.3553015 | Astros_1 |
| 53 | HILPDA | 3.24E-54 | 6.8582935 | Astros_1 |
| 54 | VEGFA | 8.23E-54 | 4.542344 | Astros_1 |
| 55 | FBXL7 | 2.26E-53 | 2.9255433 | Astros_1 |
| 56 | SLC14A1 | 1.40E-52 | 5.8348603 | Astros_1 |
| 57 | RP11-436D23.1 | 3.22E-51 | 2.4200823 | Astros_1 |
| 58 | CDH20 | 3.64E-51 | 2.661026 | Astros_1 |
| 59 | RFX4 | 7.65E-51 | 3.8435752 | Astros_1 |
| 60 | SLC4A4 | 8.33E-51 | 3.1377878 | Astros_1 |
| 61 | SPARCL1 | 1.39E-49 | 1.829308 | Astros_1 |
| 62 | TTYH1 | 3.14E-49 | 2.9115121 | Astros_1 |
| 63 | LRRC3B | 1.51E-48 | 3.969512 | Astros_1 |
| 64 | SORBS1 | 1.68E-48 | 1.9018283 | Astros_1 |
| 65 | PREX2 | 2.67E-48 | 3.434686 | Astros_1 |
| 66 | NTRK2 | 4.17E-48 | 1.4709163 | Astros_1 |
| 67 | SDC4 | 8.38E-48 | 4.9494185 | Astros_1 |
| 68 | WWOX | 1.80E-47 | 1.8537966 | Astros_1 |
| 69 | PTPRZ1 | 3.37E-47 | 2.631083 | Astros_1 |
| 70 | COL5A3 | 3.39E-47 | 3.000803 | Astros_1 |
| 71 | RGS20 | 5.39E-47 | 4.9334464 | Astros_1 |
| 72 | GLIS3 | 5.80E-47 | 3.238788 | Astros_1 |
| 73 | ACSS1 | 6.42E-47 | 3.609841 | Astros_1 |
| 74 | CHPT1 | 2.44E-46 | 2.5830915 | Astros_1 |
| 75 | TNIK | 6.37E-46 | 1.916651 | Astros_1 |
| 76 | DLC1 | 9.13E-46 | 2.668396 | Astros_1 |
| 77 | RP11-472M19.2 | 1.94E-45 | 4.2909155 | Astros_1 |
| 78 | HEPACAM | 3.85E-45 | 3.8042614 | Astros_1 |
| 79 | HPSE2 | 8.07E-45 | 3.4611673 | Astros_1 |
| 80 | SLCO1C1 | 4.13E-44 | 4.7457213 | Astros_1 |
| 81 | FGFR3 | 4.99E-44 | 3.1213646 | Astros_1 |
| 82 | CHI3L1 | 4.99E-44 | 6.331985 | Astros_1 |
| 83 | METTL7A | 1.67E-43 | 4.4203763 | Astros_1 |
| 84 | RORA | 1.89E-43 | 1.608593 | Astros_1 |
| 85 | EMX2 | 4.18E-43 | 4.544139 | Astros_1 |
| 86 | IGFBP7 | 4.90E-43 | 5.112382 | Astros_1 |
| 87 | DOCK4 | 4.01E-42 | 1.688624 | Astros_1 |
| 88 | BSG | 7.86E-40 | 2.2166276 | Astros_1 |
| 89 | NDRG2 | 1.44E-39 | 2.2876375 | Astros_1 |
| 90 | PCDH9-AS2 | 1.64E-39 | 1.895119 | Astros_1 |
| 91 | GRIN2C | 4.23E-39 | 4.306367 | Astros_1 |
| 92 | PARD3B | 4.24E-39 | 3.3597906 | Astros_1 |
| 93 | PSAP | 7.83E-39 | 1.6060593 | Astros_1 |
| 94 | LRIG1 | 9.86E-39 | 4.30274 | Astros_1 |
| 95 | PLPP3 | 9.99E-39 | 3.8779511 | Astros_1 |

| | | | | |
|---|---|---|---|---|
| 96 | LGI4 | 2.55E-38 | 2.7712145 | Astros_1 |
| 97 | TSC22D4 | 4.77E-38 | 3.4468675 | Astros_1 |
| 98 | RALGDS | 7.70E-37 | 2.6438828 | Astros_1 |
| 99 | NKAIN3 | 9.51E-37 | 2.6188796 | Astros_1 |
| 0 | SLC1A2 | 0 | 4.472158 | Astros_2 |
| 1 | PTGDS | 0 | 3.8649628 | Astros_2 |
| 2 | SLC1A3 | 0 | 4.4820786 | Astros_2 |
| 3 | GPC5 | 0 | 3.8927445 | Astros_2 |
| 4 | COL5A3 | 0 | 4.596989 | Astros_2 |
| 5 | FGFR3 | 0 | 5.1734343 | Astros_2 |
| 6 | NDRG2 | 0 | 3.8730354 | Astros_2 |
| 7 | ADGRV1 | 0 | 4.222826 | Astros_2 |
| 8 | GPM6A | 0 | 1.5394098 | Astros_2 |
| 9 | NPAS3 | 0 | 2.246582 | Astros_2 |
| 10 | LSAMP | 2.39E-305 | 1.2529488 | Astros_2 |
| 11 | GLUL | 3.16E-288 | 3.3410966 | Astros_2 |
| 12 | NTM | 3.08E-273 | 1.8763034 | Astros_2 |
| 13 | CTNNA2 | 3.71E-270 | 1.5621982 | Astros_2 |
| 14 | RNF219-AS1 | 5.42E-175 | 4.266779 | Astros_2 |
| 15 | CST3 | 4.83E-167 | 2.8812706 | Astros_2 |
| 16 | ACSS1 | 7.72E-163 | 4.416618 | Astros_2 |
| 17 | PITPNC1 | 6.72E-152 | 2.5198433 | Astros_2 |
| 18 | ZBTB20 | 2.39E-140 | 2.1245315 | Astros_2 |
| 19 | APOE | 1.50E-139 | 3.2922711 | Astros_2 |
| 20 | RORA | 3.64E-139 | 1.5149206 | Astros_2 |
| 21 | C1orf61 | 1.51E-138 | 2.7967994 | Astros_2 |
| 22 | DTNA | 3.66E-131 | 1.7599413 | Astros_2 |
| 23 | ATP1A2 | 5.44E-122 | 3.7661688 | Astros_2 |
| 24 | NKAIN3 | 5.31E-121 | 2.9530652 | Astros_2 |
| 25 | NRXN1 | 2.56E-109 | 0.7922961 | Astros_2 |
| 26 | NEAT1 | 1.37E-103 | 2.140488 | Astros_2 |
| 27 | SLC25A18 | 4.99E-102 | 4.4320164 | Astros_2 |
| 28 | HPSE2 | 1.59E-100 | 3.2902963 | Astros_2 |
| 29 | NFIA | 1.04E-87 | 1.952651 | Astros_2 |
| 30 | MSI2 | 2.36E-83 | 2.4231212 | Astros_2 |
| 31 | CDH20 | 1.61E-81 | 2.2164755 | Astros_2 |
| 32 | CADM1 | 4.94E-81 | 1.4210589 | Astros_2 |
| 33 | PLXNB1 | 9.33E-80 | 3.5515726 | Astros_2 |
| 34 | SFXN5 | 2.48E-77 | 2.5670898 | Astros_2 |
| 35 | PREX2 | 1.64E-74 | 2.876665 | Astros_2 |
| 36 | RAPGEF3 | 1.24E-73 | 3.5415957 | Astros_2 |
| 37 | LCNL1 | 5.82E-73 | 3.299153 | Astros_2 |
| 38 | FBXL7 | 1.16E-72 | 2.2356074 | Astros_2 |
| 39 | SLC4A4 | 1.79E-71 | 2.7705328 | Astros_2 |
| 40 | HIF3A | 4.76E-70 | 4.075128 | Astros_2 |
| 41 | MACF1 | 2.17E-67 | 1.4218498 | Astros_2 |
| 42 | TTYH1 | 2.35E-66 | 2.4542193 | Astros_2 |
| 43 | DLC1 | 4.47E-66 | 2.0705633 | Astros_2 |
| 44 | GJA1 | 9.52E-65 | 3.7225735 | Astros_2 |
| 45 | MIR99AHG | 1.27E-64 | 1.1036259 | Astros_2 |
| 46 | F3 | 7.13E-62 | 4.095997 | Astros_2 |
| 47 | ZNF98 | 4.69E-59 | 3.9799843 | Astros_2 |
| 48 | ZNRF3 | 2.55E-58 | 2.8828943 | Astros_2 |

| | | | | |
|---|---|---|---|---|
| 49 | TNIK | 3.88E-58 | 1.6456398 | Astros_2 |
| 50 | ATP1B2 | 3.56E-57 | 2.6730492 | Astros_2 |
| 51 | CTNND2 | 5.09E-57 | 0.71431553 | Astros_2 |
| 52 | RYR3 | 8.11E-56 | 2.3028996 | Astros_2 |
| 53 | RP11-436D23.1 | 1.24E-53 | 1.8221276 | Astros_2 |
| 54 | EMX2 | 2.48E-51 | 4.4784904 | Astros_2 |
| 55 | NHSL1 | 1.63E-50 | 3.6184068 | Astros_2 |
| 56 | MICALL2 | 1.06E-47 | 2.619722 | Astros_2 |
| 57 | GRIN2C | 1.14E-46 | 4.1747503 | Astros_2 |
| 58 | CLU | 3.50E-46 | 1.3641616 | Astros_2 |
| 59 | MT2A | 5.27E-46 | 2.743265 | Astros_2 |
| 60 | MLC1 | 1.69E-43 | 3.98654 | Astros_2 |
| 61 | GPR37L1 | 1.28E-42 | 3.4485781 | Astros_2 |
| 62 | SDC4 | 3.85E-41 | 4.332987 | Astros_2 |
| 63 | NCAN | 8.04E-40 | 2.4176795 | Astros_2 |
| 64 | PAMR1 | 1.34E-38 | 3.4683776 | Astros_2 |
| 65 | QKI | 6.23E-38 | 1.0377972 | Astros_2 |
| 66 | MTSS1L | 8.04E-38 | 2.2762964 | Astros_2 |
| 67 | CABLES1 | 1.15E-36 | 2.8757534 | Astros_2 |
| 68 | BMPR1B | 1.35E-36 | 3.028023 | Astros_2 |
| 69 | GRAMD3 | 4.15E-36 | 2.8093047 | Astros_2 |
| 70 | HEPACAM | 1.02E-35 | 2.906935 | Astros_2 |
| 71 | LRP4 | 3.05E-35 | 0.748112 | Astros_2 |
| 72 | ITGA7 | 4.79E-34 | 3.1187382 | Astros_2 |
| 73 | ERBB4 | 7.12E-34 | 0.7865683 | Astros_2 |
| 74 | PLPP3 | 8.40E-33 | 2.9507651 | Astros_2 |
| 75 | SLC7A11 | 2.20E-32 | 3.5559185 | Astros_2 |
| 76 | TPD52L1 | 2.78E-31 | 2.9433289 | Astros_2 |
| 77 | SLCO1C1 | 3.70E-31 | 3.9444609 | Astros_2 |
| 78 | PTPRZ1 | 7.70E-31 | 1.6170688 | Astros_2 |
| 79 | PARD3 | 1.15E-30 | 2.1430085 | Astros_2 |
| 80 | PARD3B | 2.23E-30 | 2.4681518 | Astros_2 |
| 81 | LINC00982 | 6.38E-30 | 4.532776 | Astros_2 |
| 82 | ADGRG1 | 1.10E-29 | 2.9346845 | Astros_2 |
| 83 | CKB | 1.36E-29 | 0.91072357 | Astros_2 |
| 84 | MMD2 | 5.64E-29 | 3.286013 | Astros_2 |
| 85 | WIF1 | 4.19E-28 | 3.1657596 | Astros_2 |
| 86 | MAML2 | 1.13E-27 | 1.7712533 | Astros_2 |
| 87 | PTN | 5.18E-27 | 2.081871 | Astros_2 |
| 88 | HTRA1 | 5.92E-27 | 2.2716162 | Astros_2 |
| 89 | WWOX | 6.04E-27 | 1.1102937 | Astros_2 |
| 90 | GFAP | 1.28E-26 | 2.564529 | Astros_2 |
| 91 | COL16A1 | 2.75E-26 | 2.0046327 | Astros_2 |
| 92 | SOX9 | 1.11E-25 | 3.2732472 | Astros_2 |
| 93 | STON2 | 1.35E-25 | 3.532648 | Astros_2 |
| 94 | LINC00499 | 4.35E-25 | 4.1594195 | Astros_2 |
| 95 | ALDH1L1 | 6.90E-24 | 3.8969295 | Astros_2 |
| 96 | CIRBP | 7.46E-24 | 1.1625935 | Astros_2 |
| 97 | ACACB | 1.14E-23 | 3.3935366 | Astros_2 |
| 98 | PSD2 | 1.18E-23 | 3.076245 | Astros_2 |
| 99 | NTRK2 | 1.18E-23 | 0.6028643 | Astros_2 |
| 0 | SLC1A2 | 9.25E-286 | 5.0210843 | Astros_3 |
| 1 | GPC5 | 1.58E-280 | 4.811595 | Astros_3 |

| | | | | |
|---|---|---|---|---|
| 2 | ADGRV1 | 6.17E-256 | 4.971284 | Astros_3 |
| 3 | SLC1A3 | 7.26E-256 | 4.9616747 | Astros_3 |
| 4 | NPAS3 | 1.11E-221 | 3.0532203 | Astros_3 |
| 5 | GPM6A | 2.43E-214 | 2.2579572 | Astros_3 |
| 6 | RNF219-AS1 | 4.19E-210 | 5.122293 | Astros_3 |
| 7 | DTNA | 2.61E-203 | 2.6665585 | Astros_3 |
| 8 | LSAMP | 8.32E-201 | 1.8681517 | Astros_3 |
| 9 | GLUL | 4.21E-196 | 3.7721992 | Astros_3 |
| 10 | ZBTB20 | 4.44E-193 | 3.3122256 | Astros_3 |
| 11 | NTM | 5.65E-193 | 2.3704078 | Astros_3 |
| 12 | COL5A3 | 3.82E-188 | 3.9337196 | Astros_3 |
| 13 | NKAIN3 | 3.34E-185 | 3.9711072 | Astros_3 |
| 14 | TNIK | 9.71E-184 | 2.8036265 | Astros_3 |
| 15 | PREX2 | 2.53E-183 | 4.2558837 | Astros_3 |
| 16 | FGFR3 | 1.12E-182 | 4.238942 | Astros_3 |
| 17 | PITPNC1 | 4.38E-181 | 3.2206223 | Astros_3 |
| 18 | PTGDS | 4.60E-172 | 3.0966551 | Astros_3 |
| 19 | RORA | 1.31E-171 | 2.2907608 | Astros_3 |
| 20 | MIR99AHG | 5.77E-163 | 2.1363704 | Astros_3 |
| 21 | CADM1 | 4.69E-162 | 2.362016 | Astros_3 |
| 22 | CTNNA2 | 4.88E-162 | 2.118841 | Astros_3 |
| 23 | CDH20 | 1.23E-160 | 3.2537904 | Astros_3 |
| 24 | FBXL7 | 4.30E-159 | 3.5721812 | Astros_3 |
| 25 | NFIA | 1.25E-156 | 2.8595662 | Astros_3 |
| 26 | SLC4A4 | 1.53E-155 | 3.785862 | Astros_3 |
| 27 | RP11-436D23.1 | 1.07E-154 | 3.0093439 | Astros_3 |
| 28 | CST3 | 1.06E-153 | 3.5171263 | Astros_3 |
| 29 | NDRG2 | 1.06E-153 | 3.2851875 | Astros_3 |
| 30 | QKI | 1.39E-151 | 2.3490953 | Astros_3 |
| 31 | MSI2 | 7.09E-144 | 3.0917695 | Astros_3 |
| 32 | APOE | 2.52E-141 | 4.0670166 | Astros_3 |
| 33 | MACF1 | 2.57E-139 | 2.0246303 | Astros_3 |
| 34 | NEAT1 | 4.35E-137 | 2.9694862 | Astros_3 |
| 35 | NRXN1 | 4.62E-137 | 1.572387 | Astros_3 |
| 36 | C1orf61 | 5.97E-137 | 3.0684767 | Astros_3 |
| 37 | ATP1A2 | 1.20E-136 | 4.138092 | Astros_3 |
| 38 | PTPRZ1 | 4.72E-134 | 3.046097 | Astros_3 |
| 39 | TRPS1 | 4.52E-130 | 2.7610312 | Astros_3 |
| 40 | WWOX | 3.27E-126 | 2.121624 | Astros_3 |
| 41 | RP11-384F7.2 | 1.17E-124 | 1.8810855 | Astros_3 |
| 42 | DLC1 | 2.15E-124 | 2.8409688 | Astros_3 |
| 43 | HPSE2 | 1.34E-116 | 4.0126824 | Astros_3 |
| 44 | RYR3 | 8.72E-116 | 2.9838402 | Astros_3 |
| 45 | ACSS1 | 4.77E-111 | 3.9798105 | Astros_3 |
| 46 | SFXN5 | 4.40E-108 | 2.8821237 | Astros_3 |
| 47 | PLPP3 | 1.90E-103 | 4.09786 | Astros_3 |
| 48 | CTNND2 | 2.70E-103 | 1.4619615 | Astros_3 |
| 49 | PARD3 | 1.05E-101 | 3.2123458 | Astros_3 |
| 50 | LRRC16A | 2.34E-100 | 2.504982 | Astros_3 |
| 51 | ZNRF3 | 1.31E-99 | 3.4993844 | Astros_3 |
| 52 | PTN | 2.21E-99 | 3.305112 | Astros_3 |
| 53 | MAML2 | 3.72E-98 | 3.0364199 | Astros_3 |
| 54 | GJA1 | 4.82E-95 | 4.301305 | Astros_3 |

| | | | | |
|---|---|---|---|---|
| 55 | GRAMD3 | 7.15E-95 | 3.8570817 | Astros_3 |
| 56 | ITPR2 | 1.71E-94 | 2.4627445 | Astros_3 |
| 57 | CPE | 3.54E-94 | 1.9703718 | Astros_3 |
| 58 | NHSL1 | 2.32E-93 | 4.4603887 | Astros_3 |
| 59 | PON2 | 9.21E-93 | 4.2151613 | Astros_3 |
| 60 | BMPR1B | 1.46E-92 | 3.9826338 | Astros_3 |
| 61 | PHLPP1 | 3.11E-92 | 2.326286 | Astros_3 |
| 62 | PPP2R2B | 3.17E-92 | 1.4340136 | Astros_3 |
| 63 | SPARCL1 | 1.42E-89 | 1.7548847 | Astros_3 |
| 64 | PARD3B | 5.00E-89 | 3.5796623 | Astros_3 |
| 65 | ATP1B2 | 1.31E-88 | 3.2281017 | Astros_3 |
| 66 | TTYH1 | 1.69E-88 | 2.782027 | Astros_3 |
| 67 | CLU | 4.11E-87 | 1.9729648 | Astros_3 |
| 68 | AQP4 | 1.81E-86 | 4.463992 | Astros_3 |
| 69 | AHCYL2 | 2.14E-86 | 2.6605663 | Astros_3 |
| 70 | PCDH9 | 3.85E-86 | 1.157153 | Astros_3 |
| 71 | MT2A | 7.13E-86 | 3.5948853 | Astros_3 |
| 72 | ZNF98 | 1.74E-85 | 4.4469967 | Astros_3 |
| 73 | ERBB4 | 3.95E-85 | 1.8571738 | Astros_3 |
| 74 | GLIS3 | 7.35E-85 | 3.0931668 | Astros_3 |
| 75 | ADD3 | 4.10E-79 | 2.3183079 | Astros_3 |
| 76 | CACNB2 | 3.72E-78 | 1.6650019 | Astros_3 |
| 77 | HIF3A | 4.48E-78 | 4.0051603 | Astros_3 |
| 78 | PAMR1 | 2.61E-76 | 4.0188546 | Astros_3 |
| 79 | GABRB1 | 8.84E-76 | 1.6392412 | Astros_3 |
| 80 | SLCO1C1 | 3.35E-75 | 4.7645674 | Astros_3 |
| 81 | DOCK4 | 5.10E-75 | 1.5524932 | Astros_3 |
| 82 | NTRK2 | 1.62E-73 | 1.260641 | Astros_3 |
| 83 | SASH1 | 4.02E-73 | 2.7247217 | Astros_3 |
| 84 | SOX5 | 1.64E-72 | 1.8283527 | Astros_3 |
| 85 | PRKG1 | 1.84E-72 | 1.8821231 | Astros_3 |
| 86 | LIFR | 2.15E-72 | 3.066713 | Astros_3 |
| 87 | APC | 6.81E-72 | 2.339522 | Astros_3 |
| 88 | SLC7A11 | 7.75E-72 | 4.38406 | Astros_3 |
| 89 | ABLIM1 | 9.59E-72 | 1.9934746 | Astros_3 |
| 90 | ARHGAP24 | 1.10E-71 | 3.3477778 | Astros_3 |
| 91 | FMN2 | 2.64E-71 | 1.4634223 | Astros_3 |
| 92 | KANK1 | 1.72E-70 | 3.13331 | Astros_3 |
| 93 | F3 | 2.89E-70 | 4.0833097 | Astros_3 |
| 94 | SOX6 | 2.15E-67 | 2.1319315 | Astros_3 |
| 95 | SLC25A18 | 3.59E-67 | 3.6899884 | Astros_3 |
| 96 | FHIT | 9.29E-67 | 1.6372836 | Astros_3 |
| 97 | LRP4 | 1.14E-66 | 1.3169692 | Astros_3 |
| 98 | LRIG1 | 3.05E-66 | 4.057287 | Astros_3 |
| 99 | MGST1 | 3.07E-66 | 2.2208576 | Astros_3 |
| 0 | CLDN5 | 6.98E-47 | 10.189488 | Endo |
| 1 | B2M | 2.90E-43 | 5.515596 | Endo |
| 2 | FLT1 | 9.40E-36 | 7.6711845 | Endo |
| 3 | EPAS1 | 1.58E-35 | 6.972641 | Endo |
| 4 | DLC1 | 8.97E-30 | 3.2220523 | Endo |
| 5 | RGS5 | 4.62E-29 | 5.744089 | Endo |
| 6 | MT2A | 4.24E-28 | 4.503874 | Endo |
| 7 | IGFBP7 | 4.72E-26 | 5.9689026 | Endo |

| | | | | |
|---|---|---|---|---|
| 8 | NEAT1 | 3.83E-25 | 2.883368 | Endo |
| 9 | BSG | 2.64E-24 | 3.1975338 | Endo |
| 10 | EBF1 | 3.78E-24 | 7.6182046 | Endo |
| 11 | RBMS3 | 5.86E-24 | 3.0961459 | Endo |
| 12 | ATP10A | 1.86E-23 | 6.7210417 | Endo |
| 13 | DUSP1 | 3.58E-23 | 4.8650184 | Endo |
| 14 | IFITM3 | 1.05E-22 | 6.3513813 | Endo |
| 15 | IFI27 | 3.00E-22 | 8.113928 | Endo |
| 16 | ATP1A2 | 5.09E-22 | 4.158728 | Endo |
| 17 | PTPRG | 7.93E-22 | 1.6805135 | Endo |
| 18 | A2M | 1.40E-21 | 6.0604434 | Endo |
| 19 | COBLL1 | 1.64E-20 | 4.771038 | Endo |
| 20 | ITM2A | 1.06E-19 | 6.180942 | Endo |
| 21 | ABCB1 | 9.40E-19 | 5.2938213 | Endo |
| 22 | LHFP | 1.78E-18 | 2.9356825 | Endo |
| 23 | ARHGAP29 | 4.40E-18 | 4.3256707 | Endo |
| 24 | HLA-E | 1.54E-17 | 6.682569 | Endo |
| 25 | EPS8 | 4.24E-17 | 4.52857 | Endo |
| 26 | ITIH5 | 1.07E-15 | 7.4440103 | Endo |
| 27 | HLA-B | 1.21E-15 | 5.350135 | Endo |
| 28 | FN1 | 2.30E-15 | 4.5147104 | Endo |
| 29 | TMSB4X | 3.33E-14 | 1.5618732 | Endo |
| 30 | ADIRF | 4.35E-14 | 6.405926 | Endo |
| 31 | NDUFA4L2 | 1.68E-13 | 6.4207473 | Endo |
| 32 | PTMA | 1.68E-13 | 2.0831788 | Endo |
| 33 | MECOM | 2.05E-13 | 4.1770697 | Endo |
| 34 | SPARC | 5.61E-13 | 5.0787168 | Endo |
| 35 | HIGD1B | 6.58E-13 | 7.001621 | Endo |
| 36 | TIMP3 | 1.11E-12 | 3.267348 | Endo |
| 37 | TMSB10 | 3.18E-12 | 1.9397882 | Endo |
| 38 | ADGRF5 | 3.71E-12 | 5.0582542 | Endo |
| 39 | APOLD1 | 8.62E-12 | 5.1611423 | Endo |
| 40 | HSPA1A | 1.39E-11 | 3.775697 | Endo |
| 41 | SLC7A5 | 2.21E-11 | 4.0428963 | Endo |
| 42 | EGFL7 | 5.98E-11 | 3.448205 | Endo |
| 43 | LEF1 | 6.27E-11 | 7.7594137 | Endo |
| 44 | SLC6A12 | 6.46E-11 | 7.6383142 | Endo |
| 45 | MFSD2A | 7.37E-11 | 6.569793 | Endo |
| 46 | H3F3B | 8.16E-11 | 2.6587079 | Endo |
| 47 | THSD4 | 9.19E-11 | 3.820986 | Endo |
| 48 | COLEC12 | 1.22E-10 | 5.520958 | Endo |
| 49 | ABCG2 | 1.31E-10 | 6.8993616 | Endo |
| 50 | CALD1 | 1.75E-10 | 3.0385897 | Endo |
| 51 | SLC7A1 | 2.24E-10 | 3.654594 | Endo |
| 52 | VWF | 2.49E-10 | 5.5174036 | Endo |
| 53 | HES4 | 4.52E-10 | 3.1192994 | Endo |
| 54 | UBC | 5.76E-10 | 2.234159 | Endo |
| 55 | LAMA2 | 6.35E-10 | 2.5861077 | Endo |
| 56 | GPCPD1 | 6.78E-10 | 3.7290957 | Endo |
| 57 | EEF1A1 | 8.66E-10 | 2.5137937 | Endo |
| 58 | ID1 | 1.07E-09 | 6.630014 | Endo |
| 59 | HLA-A | 2.58E-09 | 3.8134837 | Endo |
| 60 | STOM | 2.60E-09 | 5.060805 | Endo |

| | | | | |
|---|---|---|---|---|
| 61 | ITGA6 | 5.89E-09 | 4.772994 | Endo |
| 62 | PARD3 | 7.72E-09 | 2.7049582 | Endo |
| 63 | VIM | 8.29E-09 | 5.6148562 | Endo |
| 64 | SRGN | 1.17E-08 | 6.1284094 | Endo |
| 65 | NOTCH3 | 2.18E-08 | 7.132186 | Endo |
| 66 | ACTB | 2.85E-08 | 1.2528312 | Endo |
| 67 | HLA-C | 3.95E-08 | 4.567409 | Endo |
| 68 | SLC12A7 | 5.09E-08 | 6.646599 | Endo |
| 69 | PDGFRB | 5.31E-08 | 4.2081823 | Endo |
| 70 | ENG | 5.72E-08 | 5.183542 | Endo |
| 71 | PTN | 6.23E-08 | 2.607994 | Endo |
| 72 | LGALS1 | 6.32E-08 | 2.90673 | Endo |
| 73 | UTRN | 6.61E-08 | 2.6361592 | Endo |
| 74 | SLC2A1 | 1.09E-07 | 5.2793336 | Endo |
| 75 | MYL12A | 1.17E-07 | 5.549085 | Endo |
| 76 | KLF2 | 1.24E-07 | 6.319425 | Endo |
| 77 | PODXL | 1.27E-07 | 4.8429875 | Endo |
| 78 | IFI44L | 1.46E-07 | 6.727232 | Endo |
| 79 | SLC38A11 | 1.94E-07 | 3.5354064 | Endo |
| 80 | DCN | 2.07E-07 | 5.7086368 | Endo |
| 81 | ZBTB20 | 3.26E-07 | 1.5894375 | Endo |
| 82 | ID3 | 4.67E-07 | 5.7107086 | Endo |
| 83 | ELOVL7 | 5.02E-07 | 3.6579418 | Endo |
| 84 | WWTR1 | 7.07E-07 | 3.958647 | Endo |
| 85 | TJP1 | 1.09E-06 | 1.5942881 | Endo |
| 86 | CGNL1 | 1.64E-06 | 5.3343716 | Endo |
| 87 | TNS1 | 2.17E-06 | 4.6393647 | Endo |
| 88 | ITGA1 | 2.59E-06 | 4.8027425 | Endo |
| 89 | PTEN | 3.66E-06 | 1.9290915 | Endo |
| 90 | IFI6 | 3.67E-06 | 3.4139597 | Endo |
| 91 | BST2 | 4.68E-06 | 6.427172 | Endo |
| 92 | ST6GALNAC3 | 5.20E-06 | 2.5269487 | Endo |
| 93 | SLC30A10 | 5.66E-06 | 3.769734 | Endo |
| 94 | FLI1 | 6.28E-06 | 5.867047 | Endo |
| 95 | PRKCH | 6.28E-06 | 4.830459 | Endo |
| 96 | SLC9A3R2 | 8.78E-06 | 3.9722323 | Endo |
| 97 | ERG | 9.81E-06 | 4.463079 | Endo |
| 98 | PTRF | 1.36E-05 | 5.795132 | Endo |
| 99 | GGT5 | 1.42E-05 | 7.4036784 | Endo |
| 0 | TSHZ2 | 8.50E-191 | 4.464842 | Ex_1_L5_6 |
| 1 | ASIC2 | 4.69E-184 | 3.2514412 | Ex_1_L5_6 |
| 2 | HTR2C | 2.00E-171 | 6.5019064 | Ex_1_L5_6 |
| 3 | ZNF385D | 3.08E-160 | 3.545742 | Ex_1_L5_6 |
| 4 | HS3ST4 | 1.33E-149 | 3.3542757 | Ex_1_L5_6 |
| 5 | KIAA1456 | 8.96E-137 | 3.2262225 | Ex_1_L5_6 |
| 6 | ITGA8 | 1.15E-136 | 4.8768435 | Ex_1_L5_6 |
| 7 | OLFM3 | 7.70E-123 | 2.4253345 | Ex_1_L5_6 |
| 8 | TLE4 | 1.08E-122 | 3.122198 | Ex_1_L5_6 |
| 9 | IFNG-AS1 | 7.91E-115 | 6.300284 | Ex_1_L5_6 |
| 10 | TRHDE | 1.36E-102 | 2.6133397 | Ex_1_L5_6 |
| 11 | TOX | 3.07E-102 | 2.5120428 | Ex_1_L5_6 |
| 12 | PRR16 | 1.67E-99 | 2.7967596 | Ex_1_L5_6 |
| 13 | LUZP2 | 3.27E-99 | 2.7494884 | Ex_1_L5_6 |

| | | | | |
|---|---|---|---|---|
| 14 | DAB1 | 3.76E-96 | 2.0363708 | Ex_1_L5_6 |
| 15 | KCNT2 | 4.45E-96 | 2.428228 | Ex_1_L5_6 |
| 16 | GRM8 | 5.64E-95 | 2.7289023 | Ex_1_L5_6 |
| 17 | RP4-678D15.1 | 7.93E-95 | 3.9600365 | Ex_1_L5_6 |
| 18 | ALCAM | 1.36E-92 | 2.3699868 | Ex_1_L5_6 |
| 19 | GRIP1 | 1.88E-91 | 2.1767027 | Ex_1_L5_6 |
| 20 | VWC2L | 9.48E-90 | 3.0332842 | Ex_1_L5_6 |
| 21 | CDH6 | 7.12E-89 | 3.8794553 | Ex_1_L5_6 |
| 22 | CHRM2 | 1.02E-88 | 3.6086853 | Ex_1_L5_6 |
| 23 | CRYM | 4.73E-87 | 2.9134367 | Ex_1_L5_6 |
| 24 | LRP1B | 7.68E-87 | 1.6788812 | Ex_1_L5_6 |
| 25 | NLGN1 | 4.20E-86 | 1.6509477 | Ex_1_L5_6 |
| 26 | RP11-420N3.2 | 2.04E-85 | 3.0424356 | Ex_1_L5_6 |
| 27 | TLL1 | 4.80E-85 | 3.7318897 | Ex_1_L5_6 |
| 28 | ADCY2 | 2.10E-83 | 1.7590276 | Ex_1_L5_6 |
| 29 | LHFPL3 | 1.06E-82 | 2.241931 | Ex_1_L5_6 |
| 30 | RYR2 | 2.30E-82 | 1.6812469 | Ex_1_L5_6 |
| 31 | PCP4 | 1.51E-79 | 2.7700002 | Ex_1_L5_6 |
| 32 | DCC | 2.46E-79 | 2.0128522 | Ex_1_L5_6 |
| 33 | ETV1 | 2.66E-79 | 2.8319018 | Ex_1_L5_6 |
| 34 | ZNF385B | 3.71E-79 | 1.7700806 | Ex_1_L5_6 |
| 35 | KCTD8 | 4.88E-79 | 2.3625941 | Ex_1_L5_6 |
| 36 | NXPH2 | 6.27E-79 | 4.336754 | Ex_1_L5_6 |
| 37 | TMEM155 | 3.78E-78 | 3.0516982 | Ex_1_L5_6 |
| 38 | DCLK1 | 4.58E-78 | 1.5146918 | Ex_1_L5_6 |
| 39 | CHN2 | 1.13E-77 | 2.5427818 | Ex_1_L5_6 |
| 40 | MYRIP | 2.82E-76 | 1.9828118 | Ex_1_L5_6 |
| 41 | SLC24A2 | 8.18E-75 | 1.5971729 | Ex_1_L5_6 |
| 42 | PCDH11Y | 8.76E-74 | 2.3449898 | Ex_1_L5_6 |
| 43 | FGFR1 | 2.17E-73 | 3.3276527 | Ex_1_L5_6 |
| 44 | DOCK4 | 2.51E-73 | 1.6615797 | Ex_1_L5_6 |
| 45 | DIRAS2 | 2.20E-72 | 2.361098 | Ex_1_L5_6 |
| 46 | MDGA2 | 2.59E-72 | 1.5389816 | Ex_1_L5_6 |
| 47 | MGAT4C | 3.65E-72 | 1.6566981 | Ex_1_L5_6 |
| 48 | FOXP2 | 1.12E-71 | 2.2489762 | Ex_1_L5_6 |
| 49 | SNTG1 | 8.40E-71 | 1.5801679 | Ex_1_L5_6 |
| 50 | CAMK2D | 1.39E-70 | 1.8074651 | Ex_1_L5_6 |
| 51 | DLC1 | 1.19E-69 | 1.7954518 | Ex_1_L5_6 |
| 52 | LINC00937 | 9.93E-69 | 2.7689295 | Ex_1_L5_6 |
| 53 | COL12A1 | 1.37E-68 | 3.3525903 | Ex_1_L5_6 |
| 54 | RANBP17 | 3.59E-66 | 2.3620417 | Ex_1_L5_6 |
| 55 | BCL11B | 3.72E-65 | 2.606041 | Ex_1_L5_6 |
| 56 | SORCS2 | 4.03E-65 | 2.8369346 | Ex_1_L5_6 |
| 57 | CHSY3 | 5.01E-65 | 1.6087474 | Ex_1_L5_6 |
| 58 | NKAIN2 | 7.04E-65 | 1.531078 | Ex_1_L5_6 |
| 59 | STRBP | 8.84E-65 | 1.5532236 | Ex_1_L5_6 |
| 60 | EML6 | 2.38E-64 | 1.776851 | Ex_1_L5_6 |
| 61 | SDK1 | 4.05E-64 | 1.9609636 | Ex_1_L5_6 |
| 62 | ROBO3 | 9.33E-63 | 4.091567 | Ex_1_L5_6 |
| 63 | KIAA1217 | 5.40E-62 | 1.6930131 | Ex_1_L5_6 |
| 64 | SEMA3E | 5.55E-61 | 3.507978 | Ex_1_L5_6 |
| 65 | MAPK10 | 5.92E-61 | 1.3622181 | Ex_1_L5_6 |
| 66 | MLLT4 | 6.45E-61 | 1.9564475 | Ex_1_L5_6 |

| | | | | |
|---|---|---|---|---|
| 67 | CDH18 | 6.65E-61 | 1.5447074 | Ex_1_L5_6 |
| 68 | DPP10 | 9.98E-61 | 1.6041306 | Ex_1_L5_6 |
| 69 | SPOCK1 | 1.53E-60 | 1.5423001 | Ex_1_L5_6 |
| 70 | CPNE4 | 1.77E-60 | 1.9533318 | Ex_1_L5_6 |
| 71 | RP11-586K2.1 | 3.94E-60 | 2.0035155 | Ex_1_L5_6 |
| 72 | ERC2 | 7.27E-59 | 1.3908815 | Ex_1_L5_6 |
| 73 | NRXN1 | 2.37E-58 | 1.231938 | Ex_1_L5_6 |
| 74 | CD36 | 4.32E-58 | 4.825446 | Ex_1_L5_6 |
| 75 | LINGO2 | 5.01E-58 | 1.5421093 | Ex_1_L5_6 |
| 76 | ZNF385D-AS2 | 7.64E-58 | 4.6465883 | Ex_1_L5_6 |
| 77 | MEG3 | 6.99E-57 | 1.2612255 | Ex_1_L5_6 |
| 78 | GRM3 | 1.84E-56 | 1.6971174 | Ex_1_L5_6 |
| 79 | GHR | 2.11E-56 | 2.7337048 | Ex_1_L5_6 |
| 80 | CDH13 | 3.53E-56 | 1.6691551 | Ex_1_L5_6 |
| 81 | SEMA5B | 2.05E-55 | 3.1901045 | Ex_1_L5_6 |
| 82 | CSMD3 | 4.32E-55 | 1.3408992 | Ex_1_L5_6 |
| 83 | RORA | 9.74E-55 | 1.3940325 | Ex_1_L5_6 |
| 84 | KCNB2 | 1.02E-54 | 1.6872061 | Ex_1_L5_6 |
| 85 | DIP2A | 1.47E-54 | 2.2864168 | Ex_1_L5_6 |
| 86 | DSCAM | 3.47E-54 | 1.354567 | Ex_1_L5_6 |
| 87 | CDH11 | 3.91E-54 | 2.1745963 | Ex_1_L5_6 |
| 88 | XKR6 | 1.52E-53 | 1.5007524 | Ex_1_L5_6 |
| 89 | LRRTM3 | 2.01E-53 | 1.3712993 | Ex_1_L5_6 |
| 90 | CNTN4 | 4.46E-53 | 1.4396725 | Ex_1_L5_6 |
| 91 | GRIN3A | 6.77E-53 | 2.35575 | Ex_1_L5_6 |
| 92 | ERC1 | 2.63E-52 | 1.4864762 | Ex_1_L5_6 |
| 93 | DPY19L1 | 2.91E-52 | 2.292044 | Ex_1_L5_6 |
| 94 | GALNT13 | 1.22E-51 | 1.5533785 | Ex_1_L5_6 |
| 95 | CLSTN2 | 5.78E-51 | 1.5346174 | Ex_1_L5_6 |
| 96 | TSHZ3 | 6.50E-51 | 2.3662202 | Ex_1_L5_6 |
| 97 | RALGPS2 | 5.64E-50 | 2.3406847 | Ex_1_L5_6 |
| 98 | CDH8 | 1.25E-49 | 1.6413107 | Ex_1_L5_6 |
| 99 | CADPS | 1.26E-48 | 1.2957246 | Ex_1_L5_6 |
| 0 | MAP1B | 0 | 3.2726734 | Ex_2_L5 |
| 30 | NAPB | 0 | 2.6916761 | Ex_2_L5 |
| 29 | UCHL1 | 0 | 2.4374688 | Ex_2_L5 |
| 28 | RTN1 | 0 | 2.0389464 | Ex_2_L5 |
| 27 | TMSB10 | 0 | 2.8752701 | Ex_2_L5 |
| 26 | ATP1B1 | 0 | 2.392414 | Ex_2_L5 |
| 25 | PEBP1 | 0 | 2.5813794 | Ex_2_L5 |
| 23 | RTN3 | 0 | 2.1999552 | Ex_2_L5 |
| 22 | DYNLL1 | 0 | 2.694175 | Ex_2_L5 |
| 21 | ACTB | 0 | 2.4644754 | Ex_2_L5 |
| 20 | ARPP19 | 0 | 2.6712794 | Ex_2_L5 |
| 19 | THY1 | 0 | 2.7386167 | Ex_2_L5 |
| 18 | FTH1 | 0 | 2.4911036 | Ex_2_L5 |
| 17 | RGS4 | 0 | 2.9579666 | Ex_2_L5 |
| 16 | TMSB4X | 0 | 2.6377907 | Ex_2_L5 |
| 24 | NGFRAP1 | 0 | 2.5007665 | Ex_2_L5 |
| 14 | ENC1 | 0 | 2.7562284 | Ex_2_L5 |
| 13 | GAPDH | 0 | 2.782088 | Ex_2_L5 |
| 12 | YWHAG | 0 | 2.698037 | Ex_2_L5 |
| 11 | HSP90AA1 | 0 | 2.9401984 | Ex_2_L5 |

| | | | | |
|---|---|---|---|---|
| 10 | NRGN | 0 | 2.8166723 | Ex_2_L5 |
| 9 | STMN2 | 0 | 3.1601486 | Ex_2_L5 |
| 8 | NEFL | 0 | 3.7597826 | Ex_2_L5 |
| 7 | CHN1 | 0 | 2.686525 | Ex_2_L5 |
| 15 | SEPW1 | 0 | 2.7239816 | Ex_2_L5 |
| 6 | TUBA1B | 0 | 3.0786347 | Ex_2_L5 |
| 5 | CALM3 | 0 | 2.920467 | Ex_2_L5 |
| 4 | YWHAH | 0 | 3.0893176 | Ex_2_L5 |
| 3 | CALM1 | 0 | 2.8844593 | Ex_2_L5 |
| 2 | VSNL1 | 0 | 3.125045 | Ex_2_L5 |
| 1 | SNAP25 | 0 | 2.731889 | Ex_2_L5 |
| 31 | STMN1 | 2.02E-306 | 2.4707255 | Ex_2_L5 |
| 32 | BEX1 | 7.55E-304 | 2.7150269 | Ex_2_L5 |
| 33 | PKM | 1.89E-303 | 2.5080504 | Ex_2_L5 |
| 34 | CFL1 | 2.90E-302 | 2.7015154 | Ex_2_L5 |
| 35 | TUBB2A | 4.41E-299 | 3.160548 | Ex_2_L5 |
| 36 | LINC00657 | 4.66E-298 | 2.3865106 | Ex_2_L5 |
| 37 | RAB3A | 7.22E-297 | 2.8867602 | Ex_2_L5 |
| 38 | ALDOA | 2.14E-295 | 2.7162142 | Ex_2_L5 |
| 39 | NEFM | 3.64E-295 | 4.1024284 | Ex_2_L5 |
| 40 | DPYSL2 | 1.33E-293 | 2.171366 | Ex_2_L5 |
| 41 | RTN4 | 2.78E-289 | 2.100786 | Ex_2_L5 |
| 42 | EIF4A2 | 4.20E-289 | 2.1661136 | Ex_2_L5 |
| 43 | C14orf2 | 4.42E-287 | 2.6777353 | Ex_2_L5 |
| 44 | CKB | 1.70E-285 | 2.2299159 | Ex_2_L5 |
| 45 | YWHAB | 7.36E-282 | 2.5741317 | Ex_2_L5 |
| 46 | ATP6V1B2 | 4.20E-279 | 2.518138 | Ex_2_L5 |
| 47 | HSP90AB1 | 4.35E-279 | 2.2872927 | Ex_2_L5 |
| 48 | TPI1 | 5.94E-274 | 2.7667928 | Ex_2_L5 |
| 49 | UQCRH | 2.79E-273 | 2.6432793 | Ex_2_L5 |
| 50 | OLFM1 | 3.59E-273 | 2.3326986 | Ex_2_L5 |
| 51 | ENO2 | 5.88E-270 | 2.2379436 | Ex_2_L5 |
| 52 | SNCG | 1.09E-267 | 3.5468204 | Ex_2_L5 |
| 53 | CLU | 6.02E-267 | 2.4601262 | Ex_2_L5 |
| 54 | SKP1 | 1.24E-266 | 2.4840524 | Ex_2_L5 |
| 55 | LMO4 | 3.04E-265 | 2.3977137 | Ex_2_L5 |
| 56 | GAP43 | 3.58E-265 | 2.436442 | Ex_2_L5 |
| 57 | BASP1 | 8.89E-264 | 1.9970527 | Ex_2_L5 |
| 58 | PPIA | 1.29E-258 | 2.3933294 | Ex_2_L5 |
| 59 | RAB6B | 8.68E-256 | 2.6019938 | Ex_2_L5 |
| 60 | PCSK1N | 6.88E-254 | 2.4226725 | Ex_2_L5 |
| 61 | ACTG1 | 8.24E-254 | 2.2976682 | Ex_2_L5 |
| 62 | SNCB | 6.43E-251 | 2.3870263 | Ex_2_L5 |
| 63 | PCMT1 | 2.67E-248 | 2.7032995 | Ex_2_L5 |
| 64 | HINT1 | 3.05E-247 | 2.7050848 | Ex_2_L5 |
| 65 | SOD1 | 8.70E-247 | 2.840215 | Ex_2_L5 |
| 66 | PRNP | 2.17E-244 | 2.0588284 | Ex_2_L5 |
| 67 | SYP | 4.95E-243 | 2.067795 | Ex_2_L5 |
| 68 | SARAF | 1.77E-242 | 2.2976298 | Ex_2_L5 |
| 69 | YWHAZ | 1.12E-238 | 2.1095498 | Ex_2_L5 |
| 70 | FAIM2 | 1.69E-238 | 2.1563225 | Ex_2_L5 |
| 71 | TAGLN3 | 1.90E-238 | 2.4324932 | Ex_2_L5 |
| 72 | CREG2 | 3.58E-238 | 2.6555867 | Ex_2_L5 |

| | | | | |
|---|---|---|---|---|
| 73 | CLSTN1 | 1.55E-236 | 1.9876236 | Ex_2_L5 |
| 74 | TSPYL1 | 2.83E-235 | 2.2093554 | Ex_2_L5 |
| 75 | GNG3 | 1.01E-230 | 2.7771318 | Ex_2_L5 |
| 76 | SULT4A1 | 8.12E-229 | 2.325044 | Ex_2_L5 |
| 77 | MLLT11 | 2.07E-228 | 2.7170432 | Ex_2_L5 |
| 78 | PGAM1 | 1.99E-227 | 2.5014875 | Ex_2_L5 |
| 79 | ATP1A1 | 6.23E-226 | 2.2373617 | Ex_2_L5 |
| 80 | LYNX1 | 1.29E-224 | 2.6822717 | Ex_2_L5 |
| 81 | NDUFA4 | 1.32E-224 | 2.4301455 | Ex_2_L5 |
| 82 | CCDC85B | 3.94E-220 | 2.3757322 | Ex_2_L5 |
| 83 | SLC17A7 | 5.10E-220 | 2.0752294 | Ex_2_L5 |
| 84 | KLC1 | 1.94E-219 | 2.1645212 | Ex_2_L5 |
| 85 | GHITM | 4.15E-216 | 2.7368152 | Ex_2_L5 |
| 86 | UQCR10 | 7.19E-216 | 2.4760985 | Ex_2_L5 |
| 87 | NDRG4 | 5.55E-215 | 1.9225029 | Ex_2_L5 |
| 88 | PJA2 | 2.72E-214 | 1.9815555 | Ex_2_L5 |
| 89 | SNCA | 5.16E-213 | 2.1645575 | Ex_2_L5 |
| 90 | SCG5 | 1.69E-212 | 2.4785614 | Ex_2_L5 |
| 91 | PREPL | 1.03E-210 | 1.7767693 | Ex_2_L5 |
| 92 | BNIP3 | 3.80E-210 | 2.7380283 | Ex_2_L5 |
| 93 | CEND1 | 2.75E-209 | 2.2815647 | Ex_2_L5 |
| 94 | ATP6V0E2 | 1.81E-208 | 2.3840299 | Ex_2_L5 |
| 95 | NCDN | 3.60E-206 | 2.1986988 | Ex_2_L5 |
| 96 | NDFIP1 | 3.56E-205 | 1.867918 | Ex_2_L5 |
| 97 | SRP14 | 7.51E-205 | 2.430558 | Ex_2_L5 |
| 98 | PSAP | 1.13E-204 | 1.9124153 | Ex_2_L5 |
| 99 | SYT4 | 2.99E-204 | 2.6495357 | Ex_2_L5 |
| 0 | CHN1 | 0 | 1.7578629 | Ex_3_L4_5 |
| 54 | HPCA | 0 | 2.0180476 | Ex_3_L4_5 |
| 53 | PRNP | 0 | 1.1498964 | Ex_3_L4_5 |
| 52 | ENO2 | 0 | 1.1192173 | Ex_3_L4_5 |
| 51 | STXBP1 | 0 | 1.1546209 | Ex_3_L4_5 |
| 50 | PJA2 | 0 | 1.1660157 | Ex_3_L4_5 |
| 48 | PEBP1 | 0 | 1.1835257 | Ex_3_L4_5 |
| 47 | KLC1 | 0 | 1.222929 | Ex_3_L4_5 |
| 46 | CAMK2A | 0 | 1.1859711 | Ex_3_L4_5 |
| 45 | PDP1 | 0 | 1.5092311 | Ex_3_L4_5 |
| 44 | PRKAR1B | 0 | 1.338706 | Ex_3_L4_5 |
| 43 | GAPDH | 0 | 1.176094 | Ex_3_L4_5 |
| 42 | SYN1 | 0 | 1.5766116 | Ex_3_L4_5 |
| 41 | ALDOA | 0 | 1.2701417 | Ex_3_L4_5 |
| 40 | RTN1 | 0 | 1.0772985 | Ex_3_L4_5 |
| 39 | NAPB | 0 | 1.2973037 | Ex_3_L4_5 |
| 55 | SV2B | 0 | 1.2489523 | Ex_3_L4_5 |
| 38 | YWHAB | 0 | 1.3147652 | Ex_3_L4_5 |
| 56 | BEX1 | 0 | 1.274943 | Ex_3_L4_5 |
| 58 | TOLLIP | 0 | 1.3881866 | Ex_3_L4_5 |
| 73 | FXYD7 | 0 | 1.489567 | Ex_3_L4_5 |
| 72 | NPTN | 0 | 1.1053067 | Ex_3_L4_5 |
| 71 | NGFRAP1 | 0 | 1.0970589 | Ex_3_L4_5 |
| 70 | PPP3CA | 0 | 1.0677631 | Ex_3_L4_5 |
| 69 | SYP | 0 | 1.0742569 | Ex_3_L4_5 |
| 68 | NEFL | 0 | 1.2855179 | Ex_3_L4_5 |

| | | | | |
|---|---|---|---|---|
| 67 | EEF1A2 | 0 | 1.4651793 | Ex_3_L4_5 |
| 66 | SULT4A1 | 0 | 1.3082476 | Ex_3_L4_5 |
| 65 | PPP2R1A | 0 | 1.4152169 | Ex_3_L4_5 |
| 64 | RP11-115D19.1 | 0 | 1.7035158 | Ex_3_L4_5 |
| 63 | CLSTN1 | 0 | 1.0992674 | Ex_3_L4_5 |
| 62 | UCHL1 | 0 | 1.0915836 | Ex_3_L4_5 |
| 61 | ATP6V1B2 | 0 | 1.2427075 | Ex_3_L4_5 |
| 60 | RTN3 | 0 | 1.027762 | Ex_3_L4_5 |
| 59 | RORB | 0 | 1.7404575 | Ex_3_L4_5 |
| 57 | PREPL | 0 | 1.0942224 | Ex_3_L4_5 |
| 37 | YWHAZ | 0 | 1.2123188 | Ex_3_L4_5 |
| 49 | EIF4A2 | 0 | 1.1430165 | Ex_3_L4_5 |
| 35 | TUBB4A | 0 | 1.7319396 | Ex_3_L4_5 |
| 15 | VSNL1 | 0 | 1.3550173 | Ex_3_L4_5 |
| 14 | LMO4 | 0 | 1.4693613 | Ex_3_L4_5 |
| 13 | STMN1 | 0 | 1.4554207 | Ex_3_L4_5 |
| 12 | CALM1 | 0 | 1.253782 | Ex_3_L4_5 |
| 11 | YWHAG | 0 | 1.3816471 | Ex_3_L4_5 |
| 10 | PPIA | 0 | 1.5384876 | Ex_3_L4_5 |
| 9 | ARPP19 | 0 | 1.4295185 | Ex_3_L4_5 |
| 8 | NUAK1 | 0 | 1.5846708 | Ex_3_L4_5 |
| 7 | OLFM1 | 0 | 1.5392809 | Ex_3_L4_5 |
| 36 | MAP1B | 0 | 1.1066419 | Ex_3_L4_5 |
| 5 | PHYHIP | 0 | 1.7491605 | Ex_3_L4_5 |
| 4 | SNCB | 0 | 1.7997266 | Ex_3_L4_5 |
| 3 | SNAP25 | 0 | 1.3591661 | Ex_3_L4_5 |
| 2 | SLC17A7 | 0 | 1.9171213 | Ex_3_L4_5 |
| 1 | NRGN | 0 | 1.9496295 | Ex_3_L4_5 |
| 16 | RGS4 | 0 | 1.4360058 | Ex_3_L4_5 |
| 17 | YWHAH | 0 | 1.3439842 | Ex_3_L4_5 |
| 6 | TUBA1B | 0 | 1.5279644 | Ex_3_L4_5 |
| 19 | CALM3 | 0 | 1.2845217 | Ex_3_L4_5 |
| 34 | TAGLN3 | 0 | 1.369549 | Ex_3_L4_5 |
| 33 | GDI1 | 0 | 1.3002949 | Ex_3_L4_5 |
| 32 | NDRG4 | 0 | 1.2484161 | Ex_3_L4_5 |
| 31 | THY1 | 0 | 1.2994692 | Ex_3_L4_5 |
| 30 | FAIM2 | 0 | 1.3124379 | Ex_3_L4_5 |
| 29 | ACTB | 0 | 1.2335223 | Ex_3_L4_5 |
| 27 | SNCA | 0 | 1.3448393 | Ex_3_L4_5 |
| 28 | PIK3R1 | 0 | 1.2759157 | Ex_3_L4_5 |
| 25 | ENC1 | 0 | 1.3992952 | Ex_3_L4_5 |
| 18 | TSPYL1 | 0 | 1.4129723 | Ex_3_L4_5 |
| 24 | HSP90AB1 | 0 | 1.2993941 | Ex_3_L4_5 |
| 23 | NCDN | 0 | 1.423489 | Ex_3_L4_5 |
| 22 | CFL1 | 0 | 1.4166117 | Ex_3_L4_5 |
| 21 | TMSB10 | 0 | 1.4905617 | Ex_3_L4_5 |
| 20 | BASP1 | 0 | 1.242622 | Ex_3_L4_5 |
| 26 | SYT1 | 0 | 1.1006912 | Ex_3_L4_5 |
| 74 | CCDC85B | 4.94E-304 | 1.2790846 | Ex_3_L4_5 |
| 75 | C1orf115 | 1.07E-301 | 1.6455544 | Ex_3_L4_5 |
| 76 | ZBTB18 | 1.37E-300 | 1.3068882 | Ex_3_L4_5 |
| 77 | TSPAN7 | 1.87E-300 | 1.1453687 | Ex_3_L4_5 |
| 78 | CCK | 8.43E-297 | 1.2800704 | Ex_3_L4_5 |

| | | | | |
|---|---|---|---|---|
| 79 | CABP1 | 1.90E-294 | 1.0872735 | Ex_3_L4_5 |
| 80 | SLC39A10 | 4.05E-293 | 1.1545677 | Ex_3_L4_5 |
| 81 | ACTG1 | 4.21E-291 | 1.0523615 | Ex_3_L4_5 |
| 82 | RAB3A | 8.25E-291 | 1.2886034 | Ex_3_L4_5 |
| 83 | SEPW1 | 8.92E-290 | 1.0301714 | Ex_3_L4_5 |
| 84 | LINC00657 | 9.53E-290 | 1.0279533 | Ex_3_L4_5 |
| 85 | SERINC3 | 2.41E-289 | 1.3298206 | Ex_3_L4_5 |
| 86 | DYNLL1 | 1.00E-288 | 1.1111453 | Ex_3_L4_5 |
| 87 | TPI1 | 1.00E-288 | 1.2176472 | Ex_3_L4_5 |
| 88 | ATP1A1 | 9.15E-288 | 1.0885361 | Ex_3_L4_5 |
| 89 | IDS | 5.73E-286 | 0.9464441 | Ex_3_L4_5 |
| 90 | CEND1 | 6.00E-286 | 1.206084 | Ex_3_L4_5 |
| 91 | STMN2 | 2.47E-284 | 1.1525701 | Ex_3_L4_5 |
| 92 | DIRAS2 | 3.62E-283 | 1.416161 | Ex_3_L4_5 |
| 93 | ATP2B2 | 1.30E-282 | 1.04754 | Ex_3_L4_5 |
| 94 | MORF4L1 | 3.61E-282 | 1.0719305 | Ex_3_L4_5 |
| 95 | CAMK2N1 | 5.76E-282 | 1.0104551 | Ex_3_L4_5 |
| 96 | DNM1 | 2.84E-280 | 0.9990594 | Ex_3_L4_5 |
| 97 | JUND | 8.52E-280 | 1.2377062 | Ex_3_L4_5 |
| 98 | DYNLL2 | 4.08E-276 | 1.2310814 | Ex_3_L4_5 |
| 99 | CADPS2 | 1.40E-275 | 1.177863 | Ex_3_L4_5 |
| 0 | KCNIP4 | 9.95E-162 | 2.7668371 | Ex_4_L6 |
| 1 | ZNF804B | 3.34E-149 | 3.5048172 | Ex_4_L6 |
| 2 | RGS12 | 2.09E-141 | 3.8025205 | Ex_4_L6 |
| 3 | SORBS2 | 2.53E-135 | 3.041385 | Ex_4_L6 |
| 4 | ERC2 | 1.10E-130 | 2.3572226 | Ex_4_L6 |
| 5 | LSAMP | 2.80E-126 | 1.7872365 | Ex_4_L6 |
| 6 | KCTD16 | 3.11E-123 | 2.4938886 | Ex_4_L6 |
| 7 | RP11-384F7.2 | 9.68E-118 | 2.3296115 | Ex_4_L6 |
| 8 | SOX5 | 1.11E-117 | 2.668392 | Ex_4_L6 |
| 9 | CADPS | 2.85E-115 | 2.1742754 | Ex_4_L6 |
| 10 | NTNG2 | 9.29E-113 | 4.15079 | Ex_4_L6 |
| 11 | TMEM132D | 1.16E-111 | 2.383982 | Ex_4_L6 |
| 12 | CDH12 | 1.21E-106 | 2.394663 | Ex_4_L6 |
| 13 | FRAS1 | 2.79E-106 | 2.836203 | Ex_4_L6 |
| 14 | PDZRN4 | 3.39E-106 | 2.891687 | Ex_4_L6 |
| 15 | PDE1A | 5.54E-106 | 2.3970644 | Ex_4_L6 |
| 16 | CADPS2 | 1.28E-104 | 2.5057921 | Ex_4_L6 |
| 17 | HS3ST4 | 1.44E-103 | 2.730438 | Ex_4_L6 |
| 18 | CUX1 | 8.94E-103 | 2.817225 | Ex_4_L6 |
| 19 | NRG1 | 1.01E-101 | 2.5325446 | Ex_4_L6 |
| 20 | SYNPR | 1.47E-95 | 2.745354 | Ex_4_L6 |
| 21 | CLSTN2 | 1.53E-92 | 2.2687204 | Ex_4_L6 |
| 22 | SEMA6D | 4.01E-85 | 2.4056463 | Ex_4_L6 |
| 23 | MGAT5 | 4.11E-84 | 2.3681405 | Ex_4_L6 |
| 24 | UNC5C | 5.41E-83 | 2.0979807 | Ex_4_L6 |
| 25 | CPNE4 | 7.22E-82 | 2.394727 | Ex_4_L6 |
| 26 | NWD2 | 1.65E-80 | 2.973046 | Ex_4_L6 |
| 27 | TENM4 | 5.53E-78 | 2.0512738 | Ex_4_L6 |
| 28 | RP11-586K2.1 | 5.53E-78 | 2.5801942 | Ex_4_L6 |
| 29 | SATB2 | 8.76E-78 | 2.0339983 | Ex_4_L6 |
| 30 | RIMS2 | 8.34E-77 | 1.6471553 | Ex_4_L6 |
| 31 | THEMIS | 2.82E-74 | 4.2689238 | Ex_4_L6 |

| | | | | |
|---|---|---|---|---|
| 32 | INPP4B | 5.49E-73 | 1.9829932 | Ex_4_L6 |
| 33 | SPOCK1 | 6.56E-73 | 1.768275 | Ex_4_L6 |
| 34 | CACNA1E | 5.09E-72 | 2.0598931 | Ex_4_L6 |
| 35 | MMP16 | 1.60E-71 | 1.7701368 | Ex_4_L6 |
| 36 | GNG2 | 2.81E-71 | 2.4770367 | Ex_4_L6 |
| 37 | ARAP2 | 7.39E-70 | 1.8707459 | Ex_4_L6 |
| 38 | DLG2 | 2.18E-68 | 1.2479215 | Ex_4_L6 |
| 39 | DGKI | 2.55E-68 | 1.6706971 | Ex_4_L6 |
| 40 | SYT1 | 3.26E-68 | 1.2717685 | Ex_4_L6 |
| 41 | PCDH9 | 6.44E-68 | 1.2426683 | Ex_4_L6 |
| 42 | RBFOX1 | 1.01E-67 | 1.5071638 | Ex_4_L6 |
| 43 | GALNT14 | 3.42E-67 | 3.546716 | Ex_4_L6 |
| 44 | MCTP2 | 5.71E-66 | 5.064094 | Ex_4_L6 |
| 45 | NR4A2 | 8.51E-66 | 4.357556 | Ex_4_L6 |
| 46 | UNC5D | 2.02E-65 | 1.699172 | Ex_4_L6 |
| 47 | MAGI3 | 3.62E-65 | 2.0893247 | Ex_4_L6 |
| 48 | TENM3 | 2.74E-63 | 1.9751076 | Ex_4_L6 |
| 49 | DACH1 | 4.70E-63 | 2.8153422 | Ex_4_L6 |
| 50 | AK5 | 2.07E-61 | 1.5470946 | Ex_4_L6 |
| 51 | ZNF804A | 3.90E-61 | 1.9606524 | Ex_4_L6 |
| 52 | ENOX1 | 1.75E-60 | 1.5930705 | Ex_4_L6 |
| 53 | RP11-739G5.1 | 6.71E-60 | 3.741658 | Ex_4_L6 |
| 54 | PLEKHA5 | 7.60E-60 | 1.5085257 | Ex_4_L6 |
| 55 | TRHDE | 2.71E-59 | 2.0178561 | Ex_4_L6 |
| 56 | CSMD1 | 7.57E-59 | 1.4240885 | Ex_4_L6 |
| 57 | POSTN | 2.58E-58 | 5.303145 | Ex_4_L6 |
| 58 | CA10 | 6.20E-58 | 1.7631689 | Ex_4_L6 |
| 59 | CSGALNACT1 | 1.04E-57 | 1.967668 | Ex_4_L6 |
| 60 | IQCJ-SCHIP1 | 1.33E-57 | 1.6499835 | Ex_4_L6 |
| 61 | PCDH9-AS2 | 1.05E-55 | 1.760732 | Ex_4_L6 |
| 62 | HPSE2 | 1.12E-55 | 2.391218 | Ex_4_L6 |
| 63 | ITGB8 | 4.18E-55 | 2.9218276 | Ex_4_L6 |
| 64 | SORCS3 | 1.74E-53 | 1.7840728 | Ex_4_L6 |
| 65 | RALYL | 9.86E-53 | 1.625358 | Ex_4_L6 |
| 66 | PLD5 | 1.15E-52 | 2.3273203 | Ex_4_L6 |
| 67 | LSAMP-AS1 | 9.07E-52 | 1.9003466 | Ex_4_L6 |
| 68 | PCSK2 | 1.20E-51 | 1.7558124 | Ex_4_L6 |
| 69 | PPFIBP1 | 3.06E-51 | 1.8113502 | Ex_4_L6 |
| 70 | ZNF385B | 4.07E-51 | 1.4805176 | Ex_4_L6 |
| 71 | KCNMB2-AS1 | 1.74E-50 | 1.8348247 | Ex_4_L6 |
| 72 | DSCAML1 | 9.20E-50 | 1.8646282 | Ex_4_L6 |
| 73 | CHSY3 | 1.24E-49 | 1.4487954 | Ex_4_L6 |
| 74 | SEZ6L | 6.46E-49 | 1.6650202 | Ex_4_L6 |
| 75 | KALRN | 1.85E-48 | 1.3324457 | Ex_4_L6 |
| 76 | TRPC5 | 9.96E-48 | 2.684828 | Ex_4_L6 |
| 77 | GNB4 | 1.04E-47 | 2.9750926 | Ex_4_L6 |
| 78 | CTD-2537O9.1 | 1.04E-47 | 1.4704571 | Ex_4_L6 |
| 79 | CNTN4 | 2.06E-47 | 1.403133 | Ex_4_L6 |
| 80 | CDH2 | 2.34E-46 | 1.4333276 | Ex_4_L6 |
| 81 | MAP2K1 | 3.58E-46 | 1.5271987 | Ex_4_L6 |
| 82 | LARGE | 2.78E-44 | 1.3709438 | Ex_4_L6 |
| 83 | KCNQ3 | 1.00E-43 | 1.3135592 | Ex_4_L6 |
| 84 | NBEA | 1.09E-43 | 1.1948719 | Ex_4_L6 |

| | | | | |
|---|---|---|---|---|
| 85 | CTNNA3 | 2.36E-43 | 1.3881072 | Ex_4_L6 |
| 86 | TANC2 | 1.09E-42 | 1.2902777 | Ex_4_L6 |
| 87 | SETBP1 | 2.23E-42 | 1.5155869 | Ex_4_L6 |
| 88 | NPY1R | 3.42E-42 | 2.6584609 | Ex_4_L6 |
| 89 | RXFP1 | 5.26E-42 | 1.8515946 | Ex_4_L6 |
| 90 | PLXDC2 | 1.57E-41 | 1.4349042 | Ex_4_L6 |
| 91 | RIT2 | 2.15E-41 | 1.9387785 | Ex_4_L6 |
| 92 | NELL2 | 4.09E-41 | 1.3133441 | Ex_4_L6 |
| 93 | CPNE5 | 1.12E-40 | 2.2479718 | Ex_4_L6 |
| 94 | SORBS1 | 2.03E-40 | 1.3170905 | Ex_4_L6 |
| 95 | SFMBT2 | 2.51E-40 | 1.918179 | Ex_4_L6 |
| 96 | AFF3 | 2.83E-40 | 1.3025478 | Ex_4_L6 |
| 97 | FRMD4A | 6.38E-40 | 1.1879085 | Ex_4_L6 |
| 98 | CACNA1C | 8.30E-40 | 1.2604674 | Ex_4_L6 |
| 99 | SGCZ | 1.45E-39 | 1.3129605 | Ex_4_L6 |
| 0 | SNAP25 | 0 | 1.936642 | Ex_5_L5 |
| 1 | CHN1 | 0 | 2.1356988 | Ex_5_L5 |
| 2 | CALM1 | 1.55E-305 | 2.0340233 | Ex_5_L5 |
| 3 | MAP1B | 5.97E-305 | 2.2486188 | Ex_5_L5 |
| 4 | NRGN | 5.68E-263 | 2.171745 | Ex_5_L5 |
| 5 | ATP1B1 | 2.47E-154 | 1.6236942 | Ex_5_L5 |
| 6 | VSNL1 | 2.13E-146 | 1.8900363 | Ex_5_L5 |
| 7 | CALM3 | 6.49E-119 | 1.873014 | Ex_5_L5 |
| 8 | ENC1 | 5.18E-109 | 1.8052185 | Ex_5_L5 |
| 9 | YWHAH | 1.15E-107 | 1.8690811 | Ex_5_L5 |
| 10 | TUBA1B | 1.49E-104 | 1.9097413 | Ex_5_L5 |
| 11 | RTN1 | 3.54E-75 | 1.1294371 | Ex_5_L5 |
| 12 | RTN4 | 5.33E-75 | 1.202125 | Ex_5_L5 |
| 13 | ARPP19 | 2.77E-74 | 1.6993245 | Ex_5_L5 |
| 14 | ACTB | 7.19E-73 | 1.3887044 | Ex_5_L5 |
| 15 | RTN3 | 1.26E-72 | 1.2477332 | Ex_5_L5 |
| 16 | YWHAG | 8.38E-67 | 1.7203001 | Ex_5_L5 |
| 17 | TMSB4X | 5.53E-64 | 1.3615943 | Ex_5_L5 |
| 18 | HSP90AA1 | 1.06E-62 | 1.6529423 | Ex_5_L5 |
| 19 | GAPDH | 9.93E-58 | 1.5014162 | Ex_5_L5 |
| 20 | TMSB10 | 4.36E-56 | 1.7622181 | Ex_5_L5 |
| 21 | FTH1 | 4.96E-52 | 1.1438608 | Ex_5_L5 |
| 22 | STMN1 | 1.63E-50 | 1.5728582 | Ex_5_L5 |
| 23 | EIF4A2 | 1.48E-44 | 1.3705353 | Ex_5_L5 |
| 24 | LINC00657 | 4.50E-40 | 1.5208234 | Ex_5_L5 |
| 25 | NEFL | 5.95E-39 | 2.2562778 | Ex_5_L5 |
| 26 | SYT1 | 1.28E-37 | 0.46682283 | Ex_5_L5 |
| 27 | NGFRAP1 | 2.43E-33 | 1.4065096 | Ex_5_L5 |
| 28 | STMN2 | 6.39E-30 | 1.7799641 | Ex_5_L5 |
| 29 | SEPW1 | 1.10E-28 | 1.4870561 | Ex_5_L5 |
| 30 | PRNP | 2.39E-28 | 1.3392085 | Ex_5_L5 |
| 31 | RGS4 | 1.12E-27 | 1.7137369 | Ex_5_L5 |
| 32 | PEBP1 | 7.28E-24 | 1.3516644 | Ex_5_L5 |
| 33 | THY1 | 2.90E-21 | 1.5298822 | Ex_5_L5 |
| 34 | DYNLL1 | 6.00E-21 | 1.3870109 | Ex_5_L5 |
| 35 | SLC17A7 | 7.38E-21 | 1.4205472 | Ex_5_L5 |
| 36 | HSP90AB1 | 3.76E-18 | 1.3155136 | Ex_5_L5 |
| 37 | DPYSL2 | 1.89E-16 | 1.0656787 | Ex_5_L5 |

| | | | | |
|---|---|---|---|---|
| 38 | CAMK2A | 2.35E-16 | 1.03282 | Ex_5_L5 |
| 39 | FAIM2 | 4.81E-16 | 1.3706375 | Ex_5_L5 |
| 40 | PREPL | 1.27E-15 | 1.0891659 | Ex_5_L5 |
| 41 | IDS | 1.91E-14 | 0.97207654 | Ex_5_L5 |
| 42 | ENO2 | 2.40E-14 | 1.1765143 | Ex_5_L5 |
| 43 | CFL1 | 3.13E-13 | 1.4851528 | Ex_5_L5 |
| 44 | PPIA | 8.15E-13 | 1.4716595 | Ex_5_L5 |
| 45 | GPM6A | 8.71E-13 | 0.24681006 | Ex_5_L5 |
| 46 | BASP1 | 1.52E-12 | 0.937095 | Ex_5_L5 |
| 47 | NDRG4 | 3.35E-11 | 1.158793 | Ex_5_L5 |
| 48 | ALDOA | 4.67E-11 | 1.4381675 | Ex_5_L5 |
| 49 | UCHL1 | 8.48E-11 | 1.1855094 | Ex_5_L5 |
| 50 | CLU | 6.22E-10 | 1.192205 | Ex_5_L5 |
| 51 | CKB | 1.24E-09 | 0.8710215 | Ex_5_L5 |
| 52 | TUBB2A | 8.80E-09 | 1.645953 | Ex_5_L5 |
| 53 | BEX1 | 2.20E-08 | 1.4912001 | Ex_5_L5 |
| 54 | SPARCL1 | 5.35E-08 | 0.96576315 | Ex_5_L5 |
| 55 | PKM | 7.26E-08 | 1.2319809 | Ex_5_L5 |
| 56 | OLFM1 | 9.04E-07 | 1.3016015 | Ex_5_L5 |
| 57 | NEFM | 1.41E-06 | 2.2098494 | Ex_5_L5 |
| 58 | CLSTN1 | 1.50E-06 | 1.1319873 | Ex_5_L5 |
| 59 | PSAP | 4.16E-06 | 1.1095271 | Ex_5_L5 |
| 60 | NCDN | 5.94E-06 | 1.4460765 | Ex_5_L5 |
| 61 | CCK | 1.00E-05 | 1.2018671 | Ex_5_L5 |
| 62 | CAMK2N1 | 1.44E-05 | 0.9651696 | Ex_5_L5 |
| 63 | TSPYL1 | 1.56E-05 | 1.342719 | Ex_5_L5 |
| 64 | SYP | 2.90E-05 | 1.0710055 | Ex_5_L5 |
| 65 | LINGO1 | 3.50E-05 | 0.20465544 | Ex_5_L5 |
| 66 | SNCB | 0.00042859 | 1.2984741 | Ex_5_L5 |
| 67 | PPP3CA | 0.0009298 | 0.29456583 | Ex_5_L5 |
| 68 | NAPB | 0.002508049 | 1.3114314 | Ex_5_L5 |
| 69 | PHYHIP | 0.002754221 | 1.2254851 | Ex_5_L5 |
| 70 | TSPAN7 | 0.00564353 | 1.1521529 | Ex_5_L5 |
| 71 | SNCG | 0.009284971 | 1.5541818 | Ex_5_L5 |
| 72 | ATP1A1 | 0.009829427 | 1.2442394 | Ex_5_L5 |
| 73 | PCP4 | 0.022871978 | 1.6522676 | Ex_5_L5 |
| 74 | ACTG1 | 0.043643106 | 1.1169537 | Ex_5_L5 |
| 75 | PRKCB | 0.044946751 | 0.4080532 | Ex_5_L5 |
| 76 | RAB3A | 0.045884086 | 1.4164566 | Ex_5_L5 |
| 77 | ATP6V1B2 | 0.065405152 | 1.262162 | Ex_5_L5 |
| 78 | LMO4 | 0.069166325 | 1.176729 | Ex_5_L5 |
| 79 | MAP1A | 0.086625079 | 1.401635 | Ex_5_L5 |
| 80 | SNCA | 0.111001546 | 1.0993297 | Ex_5_L5 |
| 81 | YWHAB | 0.182743648 | 1.2774615 | Ex_5_L5 |
| 82 | PJA2 | 0.232715415 | 1.0449541 | Ex_5_L5 |
| 83 | HINT1 | 0.23659722 | 1.3797466 | Ex_5_L5 |
| 84 | PCSK1N | 0.245083289 | 1.0851716 | Ex_5_L5 |
| 85 | ATP2B1 | 0.358388733 | 0.3603296 | Ex_5_L5 |
| 86 | ITM2B | 0.414499868 | 0.865919 | Ex_5_L5 |
| 87 | NEFH | 0.418452642 | 1.8685794 | Ex_5_L5 |
| 88 | TOLLIP | 0.497335183 | 1.4065658 | Ex_5_L5 |
| 89 | TPI1 | 0.523124839 | 1.246344 | Ex_5_L5 |
| 90 | KLC1 | 0.642971986 | 1.0998912 | Ex_5_L5 |

| | | | | |
|---|---|---|---|---|
| 91 | C14orf2 | 0.690773132 | 1.1811377 | Ex_5_L5 |
| 92 | DIRAS2 | 0.89819701 | 1.4568571 | Ex_5_L5 |
| 93 | SKP1 | 0.941651146 | 1.147651 | Ex_5_L5 |
| 94 | UQCRH | 1 | 1.2717704 | Ex_5_L5 |
| 95 | TUBB | 1 | 1.4393318 | Ex_5_L5 |
| 96 | RAB6B | 1 | 1.3153934 | Ex_5_L5 |
| 97 | SOD1 | 1 | 1.3374825 | Ex_5_L5 |
| 98 | LYNX1 | 1 | 1.368784 | Ex_5_L5 |
| 99 | CARTPT | 1 | 1.9142498 | Ex_5_L5 |
| 0 | TSHZ2 | 0 | 4.18267 | Ex_6_L4_6 |
| 1 | IL1RAPL2 | 0 | 3.1580646 | Ex_6_L4_6 |
| 2 | DCC | 0 | 2.7263277 | Ex_6_L4_6 |
| 3 | RORB | 1.56E-263 | 2.6041205 | Ex_6_L4_6 |
| 4 | PTPRD | 2.16E-239 | 1.7379482 | Ex_6_L4_6 |
| 5 | RP4-678D15.1 | 1.96E-237 | 4.123525 | Ex_6_L4_6 |
| 6 | POU6F2 | 1.01E-225 | 2.3457656 | Ex_6_L4_6 |
| 7 | KHDRBS2 | 4.20E-214 | 1.7620586 | Ex_6_L4_6 |
| 8 | NKAIN2 | 6.99E-211 | 1.7164627 | Ex_6_L4_6 |
| 9 | FOXP2 | 3.79E-210 | 2.7581596 | Ex_6_L4_6 |
| 10 | RALYL | 1.62E-202 | 1.8567036 | Ex_6_L4_6 |
| 11 | ADGRL2 | 3.16E-192 | 1.7201948 | Ex_6_L4_6 |
| 12 | CTNNA2 | 7.45E-179 | 1.547385 | Ex_6_L4_6 |
| 13 | KHDRBS3 | 5.69E-173 | 1.6329954 | Ex_6_L4_6 |
| 14 | FAM19A2 | 2.84E-172 | 1.727069 | Ex_6_L4_6 |
| 15 | TOX | 3.06E-171 | 2.3176339 | Ex_6_L4_6 |
| 16 | PDZRN4 | 6.96E-168 | 2.4429188 | Ex_6_L4_6 |
| 17 | CHSY3 | 1.93E-166 | 1.6449469 | Ex_6_L4_6 |
| 18 | UNC5D | 3.18E-160 | 1.625171 | Ex_6_L4_6 |
| 19 | NELL2 | 1.23E-159 | 1.587705 | Ex_6_L4_6 |
| 20 | OPCML | 7.74E-146 | 1.3376641 | Ex_6_L4_6 |
| 21 | CNTN5 | 1.46E-141 | 1.5394306 | Ex_6_L4_6 |
| 22 | SNTG1 | 8.36E-139 | 1.4272146 | Ex_6_L4_6 |
| 23 | CPNE4 | 9.37E-138 | 2.0337076 | Ex_6_L4_6 |
| 24 | FOXP1 | 1.62E-137 | 1.4345809 | Ex_6_L4_6 |
| 25 | EFNA5 | 2.96E-137 | 1.509129 | Ex_6_L4_6 |
| 26 | KCNH5 | 3.41E-137 | 2.0080476 | Ex_6_L4_6 |
| 27 | CCSER1 | 8.45E-137 | 1.3382869 | Ex_6_L4_6 |
| 28 | KCNQ5 | 3.47E-136 | 1.4715568 | Ex_6_L4_6 |
| 29 | NEGR1 | 1.99E-135 | 1.3573681 | Ex_6_L4_6 |
| 30 | CTC-340A15.2 | 9.85E-134 | 2.621993 | Ex_6_L4_6 |
| 31 | FSTL4 | 1.30E-130 | 1.8581029 | Ex_6_L4_6 |
| 32 | KCNIP4 | 3.14E-130 | 1.7692211 | Ex_6_L4_6 |
| 33 | THRB | 8.63E-129 | 1.3054262 | Ex_6_L4_6 |
| 34 | PCLO | 1.86E-127 | 1.3473679 | Ex_6_L4_6 |
| 35 | SLC44A5 | 1.51E-126 | 1.6035258 | Ex_6_L4_6 |
| 36 | LRFN5 | 3.22E-118 | 1.3024505 | Ex_6_L4_6 |
| 37 | RYR2 | 6.41E-117 | 1.2908684 | Ex_6_L4_6 |
| 38 | ADGRB3 | 8.13E-114 | 1.086664 | Ex_6_L4_6 |
| 39 | FSTL5 | 1.73E-113 | 1.641824 | Ex_6_L4_6 |
| 40 | IQCJ-SCHIP1 | 1.98E-111 | 1.36457 | Ex_6_L4_6 |
| 41 | PDE4D | 2.17E-110 | 1.2649862 | Ex_6_L4_6 |
| 42 | PDE1A | 1.12E-109 | 1.3569819 | Ex_6_L4_6 |
| 43 | RGS7 | 4.82E-107 | 1.2073588 | Ex_6_L4_6 |

| | | | | |
|---|---|---|---|---|
| 44 | OLFM3 | 1.45E-104 | 1.3500687 | Ex_6_L4_6 |
| 45 | AGBL4 | 6.40E-104 | 1.2145313 | Ex_6_L4_6 |
| 46 | TENM2 | 7.60E-104 | 1.282302 | Ex_6_L4_6 |
| 47 | CAMK1D | 9.78E-103 | 1.2479986 | Ex_6_L4_6 |
| 48 | RP11-475O6.1 | 1.09E-102 | 2.3943403 | Ex_6_L4_6 |
| 49 | GRM7 | 2.24E-101 | 1.2675439 | Ex_6_L4_6 |
| 50 | NRXN3 | 3.58E-101 | 1.1223332 | Ex_6_L4_6 |
| 51 | NRXN1 | 3.64E-98 | 1.0713191 | Ex_6_L4_6 |
| 52 | EPHA4 | 1.60E-97 | 1.3547212 | Ex_6_L4_6 |
| 53 | PDE10A | 2.01E-97 | 1.334863 | Ex_6_L4_6 |
| 54 | RP11-707A18.1 | 1.64E-96 | 2.8494828 | Ex_6_L4_6 |
| 55 | RASAL2 | 2.24E-96 | 1.1446939 | Ex_6_L4_6 |
| 56 | HECW1 | 6.37E-96 | 1.3296431 | Ex_6_L4_6 |
| 57 | SYT1 | 1.01E-94 | 1.0280284 | Ex_6_L4_6 |
| 58 | VWC2L | 7.13E-94 | 2.3705716 | Ex_6_L4_6 |
| 59 | LDB2 | 1.41E-93 | 1.2018244 | Ex_6_L4_6 |
| 60 | FAM155A | 5.81E-93 | 1.0851834 | Ex_6_L4_6 |
| 61 | TRHDE | 2.40E-92 | 1.5957834 | Ex_6_L4_6 |
| 62 | R3HDM1 | 1.04E-89 | 1.1385788 | Ex_6_L4_6 |
| 63 | CTC-535M15.2 | 1.97E-88 | 2.4694648 | Ex_6_L4_6 |
| 64 | RXFP1 | 5.16E-88 | 1.9299952 | Ex_6_L4_6 |
| 65 | NRG3 | 5.27E-88 | 1.0785389 | Ex_6_L4_6 |
| 66 | KCNH1 | 8.75E-86 | 1.2175996 | Ex_6_L4_6 |
| 67 | FLRT2 | 3.48E-85 | 1.1209825 | Ex_6_L4_6 |
| 68 | MSC-AS1 | 8.78E-85 | 2.4594145 | Ex_6_L4_6 |
| 69 | RIMS1 | 1.19E-84 | 1.0934665 | Ex_6_L4_6 |
| 70 | KCNB2 | 5.83E-84 | 1.3788292 | Ex_6_L4_6 |
| 71 | DGKI | 1.18E-83 | 1.1221094 | Ex_6_L4_6 |
| 72 | RARB | 1.62E-83 | 2.1801941 | Ex_6_L4_6 |
| 73 | GUCY1A2 | 1.38E-82 | 1.2900926 | Ex_6_L4_6 |
| 74 | RP11-586K2.1 | 6.02E-82 | 1.7172955 | Ex_6_L4_6 |
| 75 | MEF2C | 1.13E-81 | 1.0152004 | Ex_6_L4_6 |
| 76 | CDH10 | 1.48E-81 | 1.2218962 | Ex_6_L4_6 |
| 77 | MMP16 | 1.11E-80 | 1.1214391 | Ex_6_L4_6 |
| 78 | NOVA1 | 5.06E-80 | 1.0559491 | Ex_6_L4_6 |
| 79 | CELF2 | 1.90E-79 | 1.0441408 | Ex_6_L4_6 |
| 80 | NRG1 | 4.89E-79 | 1.2049023 | Ex_6_L4_6 |
| 81 | PTPRT | 7.76E-79 | 1.289914 | Ex_6_L4_6 |
| 82 | SATB2 | 1.31E-78 | 1.2541667 | Ex_6_L4_6 |
| 83 | RP11-384F7.2 | 1.86E-78 | 1.0400653 | Ex_6_L4_6 |
| 84 | KCNQ3 | 3.21E-78 | 1.0714874 | Ex_6_L4_6 |
| 85 | HERC1 | 2.02E-77 | 1.0434347 | Ex_6_L4_6 |
| 86 | ZNF385B | 1.11E-76 | 1.1485251 | Ex_6_L4_6 |
| 87 | SCN3A | 1.85E-74 | 1.2718174 | Ex_6_L4_6 |
| 88 | CACNB4 | 2.02E-74 | 1.0354147 | Ex_6_L4_6 |
| 89 | MKL2 | 6.88E-74 | 1.0467993 | Ex_6_L4_6 |
| 90 | LINGO2 | 2.47E-71 | 1.1236973 | Ex_6_L4_6 |
| 91 | GABRB3 | 2.67E-71 | 1.0312786 | Ex_6_L4_6 |
| 92 | SLC8A1 | 4.56E-71 | 1.0089451 | Ex_6_L4_6 |
| 93 | PCDH11X | 6.23E-71 | 1.4485282 | Ex_6_L4_6 |
| 94 | CSMD3 | 9.54E-71 | 0.97021896 | Ex_6_L4_6 |
| 95 | RP11-32K4.1 | 1.49E-70 | 1.8254737 | Ex_6_L4_6 |
| 96 | SV2B | 1.43E-68 | 1.0790259 | Ex_6_L4_6 |

| | | | | |
|---|---|---|---|---|
| 97 | UBE2E3 | 1.13E-67 | 1.299073 | Ex_6_L4_6 |
| 98 | GPATCH8 | 3.80E-67 | 1.0728446 | Ex_6_L4_6 |
| 99 | MLIP | 2.99E-66 | 1.4548241 | Ex_6_L4_6 |
| 0 | PHACTR1 | 0 | 2.076664 | Ex_7_L4_6 |
| 15 | AGBL4 | 0 | 1.6293453 | Ex_7_L4_6 |
| 14 | CAMK2D | 0 | 1.9692222 | Ex_7_L4_6 |
| 13 | PTPRD | 0 | 1.6181005 | Ex_7_L4_6 |
| 12 | CLSTN2 | 0 | 1.9933753 | Ex_7_L4_6 |
| 10 | KCNH7 | 0 | 1.9382879 | Ex_7_L4_6 |
| 9 | RYR2 | 0 | 1.728588 | Ex_7_L4_6 |
| 8 | PDE1A | 0 | 1.9571694 | Ex_7_L4_6 |
| 11 | HS3ST4 | 0 | 2.2758284 | Ex_7_L4_6 |
| 6 | RALYL | 0 | 1.9798769 | Ex_7_L4_6 |
| 5 | CTC-340A15.2 | 0 | 3.5567353 | Ex_7_L4_6 |
| 4 | IL1RAPL2 | 0 | 2.8519459 | Ex_7_L4_6 |
| 3 | CADPS2 | 0 | 2.3705087 | Ex_7_L4_6 |
| 2 | NKAIN2 | 0 | 1.9638011 | Ex_7_L4_6 |
| 1 | NRG1 | 0 | 2.4658668 | Ex_7_L4_6 |
| 7 | CTC-535M15.2 | 0 | 3.5294073 | Ex_7_L4_6 |
| 16 | KCNQ5 | 3.60E-304 | 1.7099786 | Ex_7_L4_6 |
| 17 | CPNE4 | 1.34E-301 | 2.1085143 | Ex_7_L4_6 |
| 18 | CADM2 | 2.08E-300 | 1.1539779 | Ex_7_L4_6 |
| 19 | CCDC85A | 2.00E-290 | 2.0859675 | Ex_7_L4_6 |
| 20 | DPP10 | 1.66E-279 | 1.7960923 | Ex_7_L4_6 |
| 21 | CCSER1 | 6.91E-278 | 1.4920017 | Ex_7_L4_6 |
| 22 | PTPRT | 3.39E-274 | 1.7581741 | Ex_7_L4_6 |
| 23 | MDGA2 | 7.44E-272 | 1.4903176 | Ex_7_L4_6 |
| 24 | FSTL4 | 7.20E-268 | 2.02308 | Ex_7_L4_6 |
| 25 | ZNF385B | 2.39E-267 | 1.6389995 | Ex_7_L4_6 |
| 26 | RORB | 2.73E-258 | 1.8963044 | Ex_7_L4_6 |
| 27 | FOXP2 | 5.39E-253 | 2.1871088 | Ex_7_L4_6 |
| 28 | KHDRBS3 | 1.55E-248 | 1.5271016 | Ex_7_L4_6 |
| 29 | KCNIP4 | 9.95E-246 | 1.865431 | Ex_7_L4_6 |
| 30 | ST6GALNAC5 | 3.61E-237 | 1.661436 | Ex_7_L4_6 |
| 31 | CDH10 | 8.43E-236 | 1.614871 | Ex_7_L4_6 |
| 32 | PDE4D | 2.28E-235 | 1.3971505 | Ex_7_L4_6 |
| 33 | ARPP21 | 3.57E-234 | 1.4861157 | Ex_7_L4_6 |
| 34 | SYT1 | 1.83E-233 | 1.1677717 | Ex_7_L4_6 |
| 35 | PCDH7 | 4.68E-232 | 1.5221726 | Ex_7_L4_6 |
| 36 | FRMPD4 | 7.00E-229 | 1.3851069 | Ex_7_L4_6 |
| 37 | FAM19A1 | 6.05E-227 | 1.7690679 | Ex_7_L4_6 |
| 38 | NEGR1 | 1.38E-224 | 1.3896726 | Ex_7_L4_6 |
| 39 | CNTN3 | 2.21E-223 | 1.7180701 | Ex_7_L4_6 |
| 40 | HECW1 | 8.96E-218 | 1.5435438 | Ex_7_L4_6 |
| 41 | LRP1B | 3.25E-215 | 1.3647488 | Ex_7_L4_6 |
| 42 | KCNB2 | 8.64E-215 | 1.6310266 | Ex_7_L4_6 |
| 43 | RXFP1 | 1.06E-214 | 2.138289 | Ex_7_L4_6 |
| 44 | CELF4 | 2.07E-209 | 1.3832562 | Ex_7_L4_6 |
| 45 | FMN1 | 9.27E-209 | 1.49304 | Ex_7_L4_6 |
| 46 | PCLO | 5.57E-205 | 1.310639 | Ex_7_L4_6 |
| 47 | RIMS1 | 4.32E-201 | 1.3055103 | Ex_7_L4_6 |
| 48 | CTNNA2 | 4.64E-200 | 1.3122088 | Ex_7_L4_6 |
| 49 | STXBP5L | 3.02E-197 | 1.338259 | Ex_7_L4_6 |

| | | | | |
|---|---|---|---|---|
| 50 | BCL11A | 9.65E-197 | 1.5000442 | Ex_7_L4_6 |
| 51 | GRM7 | 2.16E-196 | 1.4025309 | Ex_7_L4_6 |
| 52 | NCALD | 6.33E-196 | 1.3441002 | Ex_7_L4_6 |
| 53 | CHSY3 | 7.42E-195 | 1.3877451 | Ex_7_L4_6 |
| 54 | OPCML | 9.26E-194 | 1.2689613 | Ex_7_L4_6 |
| 55 | DMD | 1.21E-191 | 1.2840487 | Ex_7_L4_6 |
| 56 | FAM155A | 4.51E-191 | 1.1767659 | Ex_7_L4_6 |
| 57 | CNTN5 | 1.03E-190 | 1.412759 | Ex_7_L4_6 |
| 58 | CDK14 | 4.73E-188 | 1.2985438 | Ex_7_L4_6 |
| 59 | FAT3 | 9.30E-188 | 1.4105752 | Ex_7_L4_6 |
| 60 | ROBO2 | 3.26E-186 | 1.5312378 | Ex_7_L4_6 |
| 61 | RBFOX1 | 9.73E-186 | 1.3331513 | Ex_7_L4_6 |
| 62 | LRRK1 | 4.12E-185 | 2.3882728 | Ex_7_L4_6 |
| 63 | ARL15 | 3.19E-184 | 1.4666027 | Ex_7_L4_6 |
| 64 | CHN2 | 2.52E-182 | 1.9195453 | Ex_7_L4_6 |
| 65 | POU6F2 | 1.56E-181 | 1.5378172 | Ex_7_L4_6 |
| 66 | BRINP3 | 3.88E-180 | 1.4604164 | Ex_7_L4_6 |
| 67 | WBSCR17 | 1.53E-179 | 1.2842729 | Ex_7_L4_6 |
| 68 | EPHA3 | 3.01E-179 | 2.0253315 | Ex_7_L4_6 |
| 69 | RP1-34H18.1 | 1.15E-177 | 1.4235802 | Ex_7_L4_6 |
| 70 | KCNT2 | 2.41E-176 | 1.5107741 | Ex_7_L4_6 |
| 71 | TENM4 | 1.87E-174 | 1.4733065 | Ex_7_L4_6 |
| 72 | COL11A1 | 8.28E-174 | 1.7896444 | Ex_7_L4_6 |
| 73 | SLC8A1 | 3.51E-173 | 1.2469095 | Ex_7_L4_6 |
| 74 | SORCS1 | 2.46E-171 | 1.4346849 | Ex_7_L4_6 |
| 75 | TLL1 | 2.11E-170 | 3.099081 | Ex_7_L4_6 |
| 76 | ERC2 | 5.87E-170 | 1.1855172 | Ex_7_L4_6 |
| 77 | DSCAM | 2.82E-168 | 1.1791768 | Ex_7_L4_6 |
| 78 | TMEFF2 | 2.29E-166 | 1.254645 | Ex_7_L4_6 |
| 79 | R3HDM1 | 5.01E-165 | 1.2224374 | Ex_7_L4_6 |
| 80 | THRB | 5.92E-165 | 1.1619562 | Ex_7_L4_6 |
| 81 | CDH8 | 2.30E-164 | 1.4311829 | Ex_7_L4_6 |
| 82 | KCNQ3 | 1.52E-163 | 1.1961017 | Ex_7_L4_6 |
| 83 | GRID2 | 2.19E-163 | 1.3144996 | Ex_7_L4_6 |
| 84 | DNAH6 | 2.89E-162 | 2.2033172 | Ex_7_L4_6 |
| 85 | RP11-116O11.1 | 3.07E-162 | 1.7429461 | Ex_7_L4_6 |
| 86 | ADGRL3 | 4.42E-159 | 1.0915931 | Ex_7_L4_6 |
| 87 | KCNC2 | 4.22E-156 | 1.1375967 | Ex_7_L4_6 |
| 88 | KALRN | 1.48E-154 | 1.1633685 | Ex_7_L4_6 |
| 89 | LHFP | 8.28E-154 | 1.5583769 | Ex_7_L4_6 |
| 90 | PID1 | 1.68E-152 | 1.5610415 | Ex_7_L4_6 |
| 91 | RORA | 4.71E-148 | 1.119573 | Ex_7_L4_6 |
| 92 | IQCJ-SCHIP1 | 1.54E-147 | 1.2491536 | Ex_7_L4_6 |
| 93 | XKR6 | 5.25E-147 | 1.2150477 | Ex_7_L4_6 |
| 94 | PRKG1 | 3.35E-146 | 1.2916948 | Ex_7_L4_6 |
| 95 | RAPGEF4 | 6.94E-146 | 1.1259519 | Ex_7_L4_6 |
| 96 | LARGE | 2.35E-145 | 1.1781511 | Ex_7_L4_6 |
| 97 | GABRG3 | 7.30E-145 | 1.171354 | Ex_7_L4_6 |
| 98 | GRIK2 | 4.94E-143 | 1.0965986 | Ex_7_L4_6 |
| 99 | BMPER | 5.73E-143 | 1.3274697 | Ex_7_L4_6 |
| 0 | HS3ST4 | 0 | 4.008965 | Ex_8_L5_6 |
| 1 | KIAA1217 | 0 | 3.1856716 | Ex_8_L5_6 |
| 2 | TLE4 | 0 | 3.4986646 | Ex_8_L5_6 |

| | | | | |
|---|---|---|---|---|
| 3 | LRP1B | 8.83E-306 | 2.096632 | Ex_8_L5_6 |
| 4 | MDGA2 | 7.33E-294 | 2.0870695 | Ex_8_L5_6 |
| 5 | ASIC2 | 6.31E-290 | 2.6052723 | Ex_8_L5_6 |
| 6 | ROBO2 | 1.26E-285 | 2.5658677 | Ex_8_L5_6 |
| 7 | DLC1 | 1.82E-283 | 2.7661502 | Ex_8_L5_6 |
| 8 | PDZRN4 | 2.05E-280 | 3.0176046 | Ex_8_L5_6 |
| 9 | SLC8A1 | 2.93E-266 | 2.071543 | Ex_8_L5_6 |
| 10 | PTPRD | 5.46E-260 | 1.8500485 | Ex_8_L5_6 |
| 11 | RALYL | 4.70E-253 | 2.1324356 | Ex_8_L5_6 |
| 12 | SORBS2 | 5.03E-227 | 2.1389391 | Ex_8_L5_6 |
| 13 | MCTP1 | 2.63E-216 | 2.0379305 | Ex_8_L5_6 |
| 14 | TMEFF2 | 4.82E-216 | 2.077838 | Ex_8_L5_6 |
| 15 | SEMA3E | 6.36E-216 | 4.3703103 | Ex_8_L5_6 |
| 16 | DPP10 | 1.05E-215 | 1.9485018 | Ex_8_L5_6 |
| 17 | PCDH11X | 6.62E-207 | 2.5115662 | Ex_8_L5_6 |
| 18 | TRPM3 | 1.39E-205 | 2.6614797 | Ex_8_L5_6 |
| 19 | FAM155A | 2.94E-201 | 1.4590204 | Ex_8_L5_6 |
| 20 | CADPS | 7.86E-197 | 1.6641346 | Ex_8_L5_6 |
| 21 | RYR3 | 6.47E-192 | 2.3871665 | Ex_8_L5_6 |
| 22 | ZFPM2 | 1.27E-191 | 1.9835901 | Ex_8_L5_6 |
| 23 | CELF2 | 1.05E-187 | 1.5062871 | Ex_8_L5_6 |
| 24 | EPHA5 | 6.91E-186 | 2.1506834 | Ex_8_L5_6 |
| 25 | FOXP2 | 3.29E-183 | 2.619727 | Ex_8_L5_6 |
| 26 | RYR2 | 5.79E-183 | 1.6603315 | Ex_8_L5_6 |
| 27 | ARPP21 | 6.83E-183 | 1.7306994 | Ex_8_L5_6 |
| 28 | FRMPD4 | 6.52E-181 | 1.6025132 | Ex_8_L5_6 |
| 29 | NRXN1 | 7.81E-178 | 1.3887783 | Ex_8_L5_6 |
| 30 | NFIB | 1.37E-176 | 1.8601421 | Ex_8_L5_6 |
| 31 | CSMD1 | 3.62E-174 | 1.5170889 | Ex_8_L5_6 |
| 32 | PCDH11Y | 3.71E-173 | 2.2204084 | Ex_8_L5_6 |
| 33 | PDE1A | 3.18E-170 | 1.7708766 | Ex_8_L5_6 |
| 34 | DGKI | 5.92E-168 | 1.6737592 | Ex_8_L5_6 |
| 35 | OLFM3 | 2.54E-163 | 1.791124 | Ex_8_L5_6 |
| 36 | SLC35F1 | 6.34E-161 | 1.6548536 | Ex_8_L5_6 |
| 37 | LINC01122 | 3.52E-158 | 1.9593192 | Ex_8_L5_6 |
| 38 | MEIS2 | 4.79E-152 | 2.4041748 | Ex_8_L5_6 |
| 39 | PHACTR1 | 5.01E-151 | 1.5181358 | Ex_8_L5_6 |
| 40 | DGKG | 1.39E-149 | 2.2125719 | Ex_8_L5_6 |
| 41 | KHDRBS3 | 4.63E-149 | 1.5519868 | Ex_8_L5_6 |
| 42 | CNTNAP5 | 3.65E-145 | 1.6181166 | Ex_8_L5_6 |
| 43 | ANKS1B | 3.45E-136 | 1.2664338 | Ex_8_L5_6 |
| 44 | MMP16 | 1.20E-132 | 1.4358089 | Ex_8_L5_6 |
| 45 | AFF3 | 2.28E-131 | 1.5008758 | Ex_8_L5_6 |
| 46 | CDH2 | 2.32E-131 | 1.5325963 | Ex_8_L5_6 |
| 47 | DMD | 1.12E-129 | 1.3739537 | Ex_8_L5_6 |
| 48 | ATP2B1 | 4.38E-128 | 1.3228388 | Ex_8_L5_6 |
| 49 | NKAIN2 | 5.02E-128 | 1.4436954 | Ex_8_L5_6 |
| 50 | SYNPO2 | 9.20E-128 | 2.2701871 | Ex_8_L5_6 |
| 51 | PCSK5 | 4.58E-127 | 3.0207412 | Ex_8_L5_6 |
| 52 | GABRG3 | 1.04E-123 | 1.4799365 | Ex_8_L5_6 |
| 53 | DOCK4 | 7.70E-123 | 1.3476839 | Ex_8_L5_6 |
| 54 | CLSTN2 | 1.48E-122 | 1.5643163 | Ex_8_L5_6 |
| 55 | LUZP2 | 1.31E-121 | 1.7794263 | Ex_8_L5_6 |

| | | | | |
|---|---|---|---|---|
| 56 | BCL11A | 1.53E-120 | 1.5262872 | Ex_8_L5_6 |
| 57 | CACNA2D3 | 9.86E-120 | 1.4321129 | Ex_8_L5_6 |
| 58 | RXFP1 | 2.94E-119 | 2.15193 | Ex_8_L5_6 |
| 59 | KALRN | 4.13E-118 | 1.3190763 | Ex_8_L5_6 |
| 60 | MIR137HG | 1.13E-116 | 1.6268713 | Ex_8_L5_6 |
| 61 | DLG2 | 1.15E-116 | 1.0895042 | Ex_8_L5_6 |
| 62 | GARNL3 | 7.68E-116 | 1.7916218 | Ex_8_L5_6 |
| 63 | BMPER | 1.76E-115 | 1.6372174 | Ex_8_L5_6 |
| 64 | NEGR1 | 1.86E-115 | 1.3016789 | Ex_8_L5_6 |
| 65 | PDE4D | 2.43E-112 | 1.300234 | Ex_8_L5_6 |
| 66 | CAMK2D | 6.12E-111 | 1.5275252 | Ex_8_L5_6 |
| 67 | PAK7 | 2.51E-110 | 1.6053438 | Ex_8_L5_6 |
| 68 | RP1-34H18.1 | 3.18E-107 | 1.4695321 | Ex_8_L5_6 |
| 69 | SV2B | 1.22E-105 | 1.334938 | Ex_8_L5_6 |
| 70 | SORCS3 | 2.25E-103 | 1.6650205 | Ex_8_L5_6 |
| 71 | PCLO | 6.17E-103 | 1.2166798 | Ex_8_L5_6 |
| 72 | ERC2 | 3.06E-102 | 1.232479 | Ex_8_L5_6 |
| 73 | SEMA3A | 6.55E-101 | 2.3046181 | Ex_8_L5_6 |
| 74 | DIRAS2 | 9.78E-99 | 1.6736083 | Ex_8_L5_6 |
| 75 | NELL2 | 9.78E-99 | 1.2339407 | Ex_8_L5_6 |
| 76 | FUT9 | 2.23E-98 | 1.3513926 | Ex_8_L5_6 |
| 77 | GRIA3 | 1.28E-96 | 1.2028347 | Ex_8_L5_6 |
| 78 | PRKCB | 2.91E-96 | 1.1838864 | Ex_8_L5_6 |
| 79 | NFIA | 3.86E-96 | 1.458425 | Ex_8_L5_6 |
| 80 | RGS7 | 4.80E-96 | 1.2003914 | Ex_8_L5_6 |
| 81 | PCDH9 | 5.23E-96 | 0.9684421 | Ex_8_L5_6 |
| 82 | DSCAM | 1.16E-95 | 1.1668662 | Ex_8_L5_6 |
| 83 | KCNMB4 | 3.25E-94 | 1.6941742 | Ex_8_L5_6 |
| 84 | SH3GL2 | 1.71E-93 | 1.2373515 | Ex_8_L5_6 |
| 85 | PRKG1 | 2.45E-93 | 1.3530617 | Ex_8_L5_6 |
| 86 | SEZ6L | 3.21E-93 | 1.49968 | Ex_8_L5_6 |
| 87 | PTPRR | 6.79E-93 | 1.6853878 | Ex_8_L5_6 |
| 88 | DCLK1 | 1.24E-92 | 1.1151304 | Ex_8_L5_6 |
| 89 | GPR158 | 2.95E-92 | 1.2827451 | Ex_8_L5_6 |
| 90 | NRG3 | 7.68E-92 | 1.0908377 | Ex_8_L5_6 |
| 91 | LARGE | 5.51E-90 | 1.1639028 | Ex_8_L5_6 |
| 92 | LMO7 | 1.80E-89 | 1.5707525 | Ex_8_L5_6 |
| 93 | CDH11 | 4.76E-89 | 1.9007149 | Ex_8_L5_6 |
| 94 | RIMS2 | 7.87E-89 | 1.1444836 | Ex_8_L5_6 |
| 95 | KIAA1456 | 9.28E-88 | 1.6645217 | Ex_8_L5_6 |
| 96 | NAV3 | 2.43E-87 | 1.1108813 | Ex_8_L5_6 |
| 97 | TANC2 | 2.68E-87 | 1.122114 | Ex_8_L5_6 |
| 98 | STXBP5 | 3.39E-87 | 1.2112995 | Ex_8_L5_6 |
| 99 | AQP4-AS1 | 3.57E-87 | 1.6308495 | Ex_8_L5_6 |
| 0 | ADRA1A | 2.81E-30 | 3.5513434 | Ex_9_L5_6 |
| 1 | BCL11B | 2.35E-28 | 3.0051217 | Ex_9_L5_6 |
| 2 | GRIK2 | 1.83E-27 | 2.2620754 | Ex_9_L5_6 |
| 3 | ROBO2 | 5.30E-27 | 2.3680897 | Ex_9_L5_6 |
| 4 | FAM19A1 | 7.63E-24 | 2.179513 | Ex_9_L5_6 |
| 5 | COL24A1 | 7.63E-24 | 2.7010663 | Ex_9_L5_6 |
| 6 | TOX | 1.27E-22 | 2.1992066 | Ex_9_L5_6 |
| 7 | VAT1L | 1.74E-22 | 3.3013098 | Ex_9_L5_6 |
| 8 | GRM8 | 1.90E-22 | 2.6019073 | Ex_9_L5_6 |

| | | | | |
|---|---|---|---|---|
| 9 | ESRRG | 5.26E-22 | 2.1598234 | Ex_9_L5_6 |
| 10 | NRG1 | 1.41E-21 | 2.1662784 | Ex_9_L5_6 |
| 11 | NRP1 | 1.81E-21 | 2.8731043 | Ex_9_L5_6 |
| 12 | PTCHD1-AS | 7.83E-20 | 2.29803 | Ex_9_L5_6 |
| 13 | LMO7 | 5.97E-19 | 1.9619505 | Ex_9_L5_6 |
| 14 | PCSK6 | 6.55E-19 | 2.6970227 | Ex_9_L5_6 |
| 15 | SLC35F3 | 7.14E-19 | 1.9085513 | Ex_9_L5_6 |
| 16 | HOMER1 | 1.22E-18 | 1.8390096 | Ex_9_L5_6 |
| 17 | ASIC2 | 2.15E-18 | 1.741382 | Ex_9_L5_6 |
| 18 | EPHA6 | 1.20E-17 | 1.7095424 | Ex_9_L5_6 |
| 19 | PDE1C | 1.20E-17 | 1.9368205 | Ex_9_L5_6 |
| 20 | TMTC1 | 1.35E-17 | 1.9459658 | Ex_9_L5_6 |
| 21 | HDAC9 | 2.25E-17 | 1.5719736 | Ex_9_L5_6 |
| 22 | GRIA3 | 3.04E-17 | 1.460654 | Ex_9_L5_6 |
| 23 | AJ006998.2 | 3.41E-17 | 2.5885758 | Ex_9_L5_6 |
| 24 | AGBL4 | 3.73E-17 | 1.4254849 | Ex_9_L5_6 |
| 25 | CLEC2L | 1.74E-16 | 2.3944836 | Ex_9_L5_6 |
| 26 | CBLN2 | 1.88E-16 | 2.0911243 | Ex_9_L5_6 |
| 27 | MEIS2 | 1.99E-16 | 1.9535075 | Ex_9_L5_6 |
| 28 | DCLK1 | 2.78E-16 | 1.3018547 | Ex_9_L5_6 |
| 29 | FAM155A | 5.43E-16 | 1.0750347 | Ex_9_L5_6 |
| 30 | FAM135B | 5.93E-16 | 1.8163537 | Ex_9_L5_6 |
| 31 | SPHKAP | 7.00E-16 | 2.3143036 | Ex_9_L5_6 |
| 32 | NFIB | 7.00E-16 | 1.6538019 | Ex_9_L5_6 |
| 33 | KCNIP4 | 7.17E-16 | 1.5813425 | Ex_9_L5_6 |
| 34 | KIAA1456 | 4.16E-15 | 1.789661 | Ex_9_L5_6 |
| 35 | MYO16 | 4.54E-15 | 1.9874057 | Ex_9_L5_6 |
| 36 | CORO6 | 4.85E-15 | 1.7292197 | Ex_9_L5_6 |
| 37 | COL5A2 | 6.32E-15 | 2.2900693 | Ex_9_L5_6 |
| 38 | ZNF804B | 1.17E-14 | 1.6924323 | Ex_9_L5_6 |
| 39 | CDH20 | 1.58E-14 | 1.5766343 | Ex_9_L5_6 |
| 40 | PEX5L | 1.67E-14 | 1.3957993 | Ex_9_L5_6 |
| 41 | ADAMTSL3 | 2.11E-14 | 2.6723576 | Ex_9_L5_6 |
| 42 | STRBP | 3.11E-14 | 1.318706 | Ex_9_L5_6 |
| 43 | FAT3 | 6.06E-14 | 1.4558064 | Ex_9_L5_6 |
| 44 | KCNN2 | 2.54E-13 | 2.1024456 | Ex_9_L5_6 |
| 45 | FAM189A1 | 2.54E-13 | 1.601151 | Ex_9_L5_6 |
| 46 | STK39 | 2.74E-13 | 1.5342537 | Ex_9_L5_6 |
| 47 | ARL15 | 3.41E-13 | 1.5871495 | Ex_9_L5_6 |
| 48 | CABP1 | 5.15E-13 | 1.3260082 | Ex_9_L5_6 |
| 49 | RYR3 | 5.19E-13 | 1.5891342 | Ex_9_L5_6 |
| 50 | DGKH | 5.49E-13 | 1.9620792 | Ex_9_L5_6 |
| 51 | DSCAML1 | 1.08E-12 | 1.579438 | Ex_9_L5_6 |
| 52 | CRYM | 1.51E-12 | 1.8416697 | Ex_9_L5_6 |
| 53 | PTPRN2 | 1.75E-12 | 1.2826056 | Ex_9_L5_6 |
| 54 | KCNH7 | 1.84E-12 | 1.4152453 | Ex_9_L5_6 |
| 55 | DIAPH2 | 3.56E-12 | 1.4755639 | Ex_9_L5_6 |
| 56 | SLC8A1 | 3.56E-12 | 1.228586 | Ex_9_L5_6 |
| 57 | MAST4 | 4.26E-12 | 1.2516228 | Ex_9_L5_6 |
| 58 | NTNG1 | 4.40E-12 | 1.629103 | Ex_9_L5_6 |
| 59 | GRIN2A | 5.12E-12 | 1.1795955 | Ex_9_L5_6 |
| 60 | MEG3 | 7.76E-12 | 1.097818 | Ex_9_L5_6 |
| 61 | RP11-32K4.1 | 1.00E-11 | 1.7974119 | Ex_9_L5_6 |

| | | | | |
|---|---|---|---|---|
| 62 | HCN1 | 1.17E-11 | 1.2470006 | Ex_9_L5_6 |
| 63 | LRP1B | 1.52E-11 | 1.1152526 | Ex_9_L5_6 |
| 64 | NKAIN2 | 2.21E-11 | 1.2074665 | Ex_9_L5_6 |
| 65 | SLC39A11 | 2.27E-11 | 1.512939 | Ex_9_L5_6 |
| 66 | TMTC2 | 2.35E-11 | 1.4015634 | Ex_9_L5_6 |
| 67 | GPC6 | 4.71E-11 | 1.4415118 | Ex_9_L5_6 |
| 68 | UNC80 | 5.55E-11 | 1.2703013 | Ex_9_L5_6 |
| 69 | FOCAD | 6.88E-11 | 1.226944 | Ex_9_L5_6 |
| 70 | SLC26A4 | 7.70E-11 | 2.3937879 | Ex_9_L5_6 |
| 71 | ANO4 | 9.24E-11 | 1.1711931 | Ex_9_L5_6 |
| 72 | ADAMTS19 | 9.86E-11 | 1.6000684 | Ex_9_L5_6 |
| 73 | RP11-624C23.1 | 1.36E-10 | 1.7006711 | Ex_9_L5_6 |
| 75 | SLIT3 | 1.41E-10 | 1.6242704 | Ex_9_L5_6 |
| 74 | VWA3A | 1.41E-10 | 2.4221902 | Ex_9_L5_6 |
| 76 | ZNF385B | 1.84E-10 | 1.2317797 | Ex_9_L5_6 |
| 77 | FRMD4A | 2.11E-10 | 1.0525993 | Ex_9_L5_6 |
| 78 | NEBL | 3.12E-10 | 1.1586509 | Ex_9_L5_6 |
| 79 | C4orf22 | 3.19E-10 | 2.303907 | Ex_9_L5_6 |
| 80 | RBFOX1 | 3.34E-10 | 1.1200298 | Ex_9_L5_6 |
| 81 | CNTN3 | 3.38E-10 | 1.3072674 | Ex_9_L5_6 |
| 82 | TXK | 3.79E-10 | 3.487941 | Ex_9_L5_6 |
| 83 | STXBP5-AS1 | 4.04E-10 | 1.3218362 | Ex_9_L5_6 |
| 84 | SNX29 | 4.05E-10 | 1.447954 | Ex_9_L5_6 |
| 85 | ARHGEF3 | 4.27E-10 | 1.198468 | Ex_9_L5_6 |
| 86 | NLGN1 | 4.43E-10 | 1.0803542 | Ex_9_L5_6 |
| 87 | ETV1 | 4.60E-10 | 1.7440349 | Ex_9_L5_6 |
| 88 | PDE1A | 5.01E-10 | 1.2252382 | Ex_9_L5_6 |
| 89 | TEAD1 | 5.34E-10 | 1.4570733 | Ex_9_L5_6 |
| 90 | LHFP | 5.50E-10 | 1.2504594 | Ex_9_L5_6 |
| 91 | CCSER1 | 6.51E-10 | 1.0583297 | Ex_9_L5_6 |
| 92 | RAPGEF5 | 6.56E-10 | 1.1128325 | Ex_9_L5_6 |
| 93 | CDHR3 | 7.00E-10 | 1.8825905 | Ex_9_L5_6 |
| 94 | FNBP1L | 7.94E-10 | 1.4736819 | Ex_9_L5_6 |
| 95 | MAGI1 | 8.41E-10 | 1.0737773 | Ex_9_L5_6 |
| 96 | SRPK1 | 1.05E-09 | 1.5757918 | Ex_9_L5_6 |
| 97 | GULP1 | 1.37E-09 | 1.4587003 | Ex_9_L5_6 |
| 98 | SCN4B | 1.37E-09 | 2.345662 | Ex_9_L5_6 |
| 99 | NEGR1 | 1.47E-09 | 1.081104 | Ex_9_L5_6 |
| 0 | KCNIP4 | 0 | 2.4499843 | Ex_10_L2_4 |
| 72 | DLGAP1 | 0 | 0.9907236 | Ex_10_L2_4 |
| 71 | NELL2 | 0 | 1.1507666 | Ex_10_L2_4 |
| 70 | AJ006998.2 | 0 | 2.3006294 | Ex_10_L2_4 |
| 69 | GRM7 | 0 | 1.1435975 | Ex_10_L2_4 |
| 68 | MIR137HG | 0 | 1.5046195 | Ex_10_L2_4 |
| 67 | RFX3 | 0 | 1.3089585 | Ex_10_L2_4 |
| 66 | CCSER1 | 0 | 1.0670229 | Ex_10_L2_4 |
| 65 | SATB2 | 0 | 1.3420539 | Ex_10_L2_4 |
| 64 | RP11-30J20.1 | 0 | 2.3920379 | Ex_10_L2_4 |
| 63 | CACNB2 | 0 | 1.1298302 | Ex_10_L2_4 |
| 62 | ARHGAP26 | 0 | 1.1701986 | Ex_10_L2_4 |
| 61 | GRIA4 | 0 | 1.1598212 | Ex_10_L2_4 |
| 60 | ST6GALNAC5 | 0 | 1.3059257 | Ex_10_L2_4 |
| 59 | ERICH1-AS1 | 0 | 1.5090834 | Ex_10_L2_4 |

| | | | | |
|---|---|---|---|---|
| 58 | SIPA1L1 | 0 | 1.1000134 | Ex_10_L2_4 |
| 57 | TENM2 | 0 | 1.1517566 | Ex_10_L2_4 |
| 56 | LINC01250 | 0 | 1.8824081 | Ex_10_L2_4 |
| 55 | ANKS1B | 0 | 0.98727494 | Ex_10_L2_4 |
| 54 | PDZD2 | 0 | 1.4645382 | Ex_10_L2_4 |
| 53 | MSRA | 0 | 1.2165864 | Ex_10_L2_4 |
| 52 | MLIP | 0 | 1.8522062 | Ex_10_L2_4 |
| 73 | FAM153B | 0 | 1.3777978 | Ex_10_L2_4 |
| 51 | TMEM178B | 0 | 1.2562928 | Ex_10_L2_4 |
| 74 | FAT3 | 0 | 1.2616866 | Ex_10_L2_4 |
| 76 | FRMD4A | 0 | 0.98991865 | Ex_10_L2_4 |
| 97 | AC114765.1 | 0 | 1.867282 | Ex_10_L2_4 |
| 96 | PVRL3 | 0 | 1.5465589 | Ex_10_L2_4 |
| 95 | DPYD | 0 | 1.1208905 | Ex_10_L2_4 |
| 94 | GPC6 | 0 | 1.2268155 | Ex_10_L2_4 |
| 93 | DGKI | 0 | 1.0636735 | Ex_10_L2_4 |
| 92 | CACNA2D1 | 0 | 1.0246284 | Ex_10_L2_4 |
| 91 | ZFPM2 | 0 | 1.21601 | Ex_10_L2_4 |
| 90 | NEGR1 | 0 | 1.0175489 | Ex_10_L2_4 |
| 89 | MED12L | 0 | 1.2700996 | Ex_10_L2_4 |
| 88 | OLFM3 | 0 | 1.2097197 | Ex_10_L2_4 |
| 87 | KCNH7 | 0 | 1.1845217 | Ex_10_L2_4 |
| 86 | EPB41L2 | 0 | 1.1936021 | Ex_10_L2_4 |
| 85 | TANC2 | 0 | 1.080013 | Ex_10_L2_4 |
| 84 | HOMER1 | 0 | 1.2893544 | Ex_10_L2_4 |
| 83 | EPHA6 | 0 | 1.2445621 | Ex_10_L2_4 |
| 82 | CDH9 | 0 | 1.373207 | Ex_10_L2_4 |
| 81 | CNKSR2 | 0 | 1.1435038 | Ex_10_L2_4 |
| 80 | ERC2 | 0 | 1.0406213 | Ex_10_L2_4 |
| 79 | RIMS2 | 0 | 1.0011352 | Ex_10_L2_4 |
| 78 | RYR2 | 0 | 1.0701367 | Ex_10_L2_4 |
| 77 | CUX2 | 0 | 2.2333097 | Ex_10_L2_4 |
| 75 | GRIA2 | 0 | 0.92726463 | Ex_10_L2_4 |
| 50 | SYN3 | 0 | 1.705138 | Ex_10_L2_4 |
| 49 | TMEM132D | 0 | 1.2506307 | Ex_10_L2_4 |
| 48 | FAM155A | 0 | 1.0267868 | Ex_10_L2_4 |
| 21 | CDH18 | 0 | 1.6410792 | Ex_10_L2_4 |
| 20 | CA10 | 0 | 1.891829 | Ex_10_L2_4 |
| 19 | LINGO2 | 0 | 1.6229713 | Ex_10_L2_4 |
| 18 | PTPRD | 0 | 1.394033 | Ex_10_L2_4 |
| 17 | CBLN2 | 0 | 2.4746385 | Ex_10_L2_4 |
| 16 | ATRNL1 | 0 | 1.4378798 | Ex_10_L2_4 |
| 15 | STXBP5L | 0 | 1.5129632 | Ex_10_L2_4 |
| 14 | AC011288.2 | 0 | 2.3682127 | Ex_10_L2_4 |
| 13 | R3HDM1 | 0 | 1.5587804 | Ex_10_L2_4 |
| 12 | RALYL | 0 | 1.7540809 | Ex_10_L2_4 |
| 11 | LDB2 | 0 | 1.6940919 | Ex_10_L2_4 |
| 10 | KCNQ5 | 0 | 1.6505245 | Ex_10_L2_4 |
| 9 | FAM19A1 | 0 | 2.0646517 | Ex_10_L2_4 |
| 8 | PDE4D | 0 | 1.4799819 | Ex_10_L2_4 |
| 7 | HS6ST3 | 0 | 1.8781737 | Ex_10_L2_4 |
| 6 | RBFOX1 | 0 | 1.5600232 | Ex_10_L2_4 |
| 5 | DGKB | 0 | 1.9419206 | Ex_10_L2_4 |

| | | | | |
|---|---|---|---|---|
| 4 | CSMD1 | 0 | 1.5198427 | Ex_10_L2_4 |
| 3 | IQCJ-SCHIP1 | 0 | 1.9446595 | Ex_10_L2_4 |
| 2 | LRRTM4 | 0 | 1.7604989 | Ex_10_L2_4 |
| 1 | MEG3 | 0 | 1.4579382 | Ex_10_L2_4 |
| 22 | CELF2 | 0 | 1.2649049 | Ex_10_L2_4 |
| 23 | RGS6 | 0 | 1.6492687 | Ex_10_L2_4 |
| 24 | ADGRB3 | 0 | 1.0913407 | Ex_10_L2_4 |
| 25 | PTPRK | 0 | 1.6143394 | Ex_10_L2_4 |
| 47 | KCTD16 | 0 | 1.245625 | Ex_10_L2_4 |
| 46 | ARPP21 | 0 | 1.2476074 | Ex_10_L2_4 |
| 45 | ZNF804B | 0 | 1.5476649 | Ex_10_L2_4 |
| 44 | KCNH1 | 0 | 1.4100358 | Ex_10_L2_4 |
| 43 | OPCML | 0 | 1.1447369 | Ex_10_L2_4 |
| 42 | ASIC2 | 0 | 1.2543768 | Ex_10_L2_4 |
| 41 | ADGRL2 | 0 | 1.3115579 | Ex_10_L2_4 |
| 40 | NRG3 | 0 | 1.0692071 | Ex_10_L2_4 |
| 39 | LRFN5 | 0 | 1.2722671 | Ex_10_L2_4 |
| 38 | PLCB1 | 0 | 1.0744331 | Ex_10_L2_4 |
| 98 | EPHA4 | 0 | 1.1340284 | Ex_10_L2_4 |
| 37 | AGBL4 | 0 | 1.2327126 | Ex_10_L2_4 |
| 35 | RGS7 | 0 | 1.186922 | Ex_10_L2_4 |
| 34 | CTD-2537O9.1 | 0 | 1.4361229 | Ex_10_L2_4 |
| 33 | FLRT2 | 0 | 1.3449495 | Ex_10_L2_4 |
| 32 | KHDRBS2 | 0 | 1.3188771 | Ex_10_L2_4 |
| 31 | DLG2 | 0 | 0.9885175 | Ex_10_L2_4 |
| 30 | PHACTR1 | 0 | 1.3243395 | Ex_10_L2_4 |
| 29 | CACNA2D3 | 0 | 1.4623513 | Ex_10_L2_4 |
| 28 | KALRN | 0 | 1.2998387 | Ex_10_L2_4 |
| 27 | CHRM3 | 0 | 1.4130839 | Ex_10_L2_4 |
| 26 | CDH12 | 0 | 1.6145151 | Ex_10_L2_4 |
| 36 | PLEKHA5 | 0 | 1.2855432 | Ex_10_L2_4 |
| 99 | SORCS1 | 0 | 1.1824181 | Ex_10_L2_4 |
| 0 | FGF13 | 0 | 4.539409 | Inhib_1 |
| 16 | PTCHD4 | 0 | 3.7320743 | Inhib_1 |
| 15 | GAD2 | 0 | 4.060084 | Inhib_1 |
| 14 | NXPH1 | 0 | 3.0730307 | Inhib_1 |
| 13 | GRIN2A | 0 | 2.166875 | Inhib_1 |
| 12 | MGAT4C | 0 | 2.5718062 | Inhib_1 |
| 10 | MYO16 | 0 | 3.3911998 | Inhib_1 |
| 9 | SLC6A1 | 0 | 3.7740357 | Inhib_1 |
| 11 | GRIP1 | 0 | 2.8240814 | Inhib_1 |
| 7 | DNER | 0 | 3.3489976 | Inhib_1 |
| 6 | RP11-123O10.4 | 0 | 3.263291 | Inhib_1 |
| 5 | GRIK1 | 0 | 3.6260664 | Inhib_1 |
| 4 | FGF14 | 0 | 2.086485 | Inhib_1 |
| 3 | ADARB2 | 0 | 4.3153563 | Inhib_1 |
| 2 | KIT | 0 | 6.378995 | Inhib_1 |
| 1 | SGCZ | 0 | 3.670339 | Inhib_1 |
| 8 | FBXL7 | 0 | 3.8120785 | Inhib_1 |
| 17 | NFIB | 7.10E-306 | 2.5358198 | Inhib_1 |
| 18 | MACROD2 | 9.23E-306 | 2.129062 | Inhib_1 |
| 19 | PRELID2 | 2.28E-285 | 4.5649614 | Inhib_1 |
| 20 | PTPRT | 4.81E-283 | 2.4124677 | Inhib_1 |

| | | | | |
|---|---|---|---|---|
| 21 | FSTL5 | 1.11E-273 | 2.8207927 | Inhib_1 |
| 22 | ZNF536 | 2.03E-273 | 2.862412 | Inhib_1 |
| 23 | EYA4 | 2.02E-271 | 5.503683 | Inhib_1 |
| 24 | UNC13C | 4.76E-266 | 3.1585922 | Inhib_1 |
| 25 | IL1RAPL1 | 2.14E-264 | 1.786539 | Inhib_1 |
| 26 | KCNC2 | 1.51E-258 | 2.0934064 | Inhib_1 |
| 27 | GRIA4 | 6.40E-256 | 2.0994134 | Inhib_1 |
| 28 | RBMS3 | 4.00E-252 | 2.8003035 | Inhib_1 |
| 29 | RAB3C | 2.16E-244 | 2.312395 | Inhib_1 |
| 30 | ALK | 4.99E-239 | 3.2239554 | Inhib_1 |
| 31 | SLC6A1-AS1 | 2.81E-237 | 3.060737 | Inhib_1 |
| 32 | KCNAB1 | 2.95E-233 | 2.4149392 | Inhib_1 |
| 33 | ATP8A2 | 1.37E-231 | 1.9335515 | Inhib_1 |
| 34 | KAZN | 2.75E-230 | 2.0188904 | Inhib_1 |
| 35 | GRIK2 | 2.11E-229 | 1.8659562 | Inhib_1 |
| 36 | TMEM132D | 1.72E-221 | 1.9430641 | Inhib_1 |
| 37 | PTPRM | 7.22E-218 | 2.3148165 | Inhib_1 |
| 38 | DLX6-AS1 | 1.12E-215 | 3.25557 | Inhib_1 |
| 39 | MTUS2 | 3.55E-215 | 1.9760348 | Inhib_1 |
| 40 | GRIN3A | 7.95E-215 | 3.3032916 | Inhib_1 |
| 41 | NTNG1 | 3.86E-211 | 2.80148 | Inhib_1 |
| 42 | DPP6 | 1.24E-209 | 1.5926937 | Inhib_1 |
| 43 | SV2C | 4.71E-200 | 4.036624 | Inhib_1 |
| 44 | GAD1 | 4.26E-194 | 2.5153883 | Inhib_1 |
| 45 | ERBB4 | 1.66E-193 | 2.156399 | Inhib_1 |
| 46 | LUZP2 | 2.09E-189 | 2.4520097 | Inhib_1 |
| 47 | SPHKAP | 1.19E-188 | 3.2264504 | Inhib_1 |
| 48 | NRIP3 | 6.20E-185 | 3.3447366 | Inhib_1 |
| 49 | GRM5 | 1.87E-183 | 1.4926289 | Inhib_1 |
| 50 | TRPC3 | 8.52E-183 | 4.8709064 | Inhib_1 |
| 51 | LSAMP | 4.12E-179 | 1.2248986 | Inhib_1 |
| 52 | EXT1 | 1.32E-177 | 1.8784517 | Inhib_1 |
| 53 | BCL11B | 4.94E-172 | 2.9551165 | Inhib_1 |
| 54 | SGK1 | 1.95E-171 | 2.152038 | Inhib_1 |
| 55 | LAMP5 | 3.14E-169 | 4.0755625 | Inhib_1 |
| 56 | DAB1 | 4.98E-167 | 1.6106961 | Inhib_1 |
| 57 | SYT1 | 7.40E-165 | 1.1948186 | Inhib_1 |
| 58 | PIP5K1B | 6.66E-163 | 2.5141242 | Inhib_1 |
| 59 | UBASH3B | 5.87E-161 | 3.1421616 | Inhib_1 |
| 60 | UNC5C | 5.27E-156 | 1.6637293 | Inhib_1 |
| 61 | DOCK10 | 3.07E-155 | 2.0631394 | Inhib_1 |
| 62 | ZNF385D | 5.91E-155 | 1.9396276 | Inhib_1 |
| 63 | CACNA2D1 | 1.43E-153 | 1.7420274 | Inhib_1 |
| 64 | ARL4C | 1.14E-152 | 2.5026147 | Inhib_1 |
| 65 | PKP2 | 6.85E-152 | 4.8104286 | Inhib_1 |
| 66 | ZEB2 | 2.48E-150 | 1.5520833 | Inhib_1 |
| 67 | CDH13 | 3.66E-149 | 2.0580945 | Inhib_1 |
| 68 | MTSS1 | 7.40E-149 | 2.4231825 | Inhib_1 |
| 69 | RPH3A | 7.95E-148 | 1.8603195 | Inhib_1 |
| 70 | UNC5D | 8.12E-148 | 1.7249639 | Inhib_1 |
| 71 | AP1S2 | 6.04E-144 | 3.0299215 | Inhib_1 |
| 72 | GABRB2 | 2.59E-143 | 1.3875521 | Inhib_1 |
| 73 | CNTNAP4 | 3.95E-138 | 2.1943984 | Inhib_1 |

| | | | | |
|---|---|---|---|---|
| 74 | NXPH2 | 8.22E-138 | 4.4694853 | Inhib_1 |
| 75 | CNTN5 | 5.95E-137 | 1.5292892 | Inhib_1 |
| 76 | CDK14 | 3.96E-135 | 1.4407352 | Inhib_1 |
| 77 | TIAM1 | 2.00E-134 | 1.8354349 | Inhib_1 |
| 78 | RGS7 | 3.31E-133 | 1.3065279 | Inhib_1 |
| 79 | KCNIP1 | 1.77E-131 | 3.0652847 | Inhib_1 |
| 80 | SCN1A | 8.96E-131 | 1.4113138 | Inhib_1 |
| 81 | LRRC7 | 8.90E-129 | 1.2510954 | Inhib_1 |
| 82 | CCSER1 | 1.63E-126 | 1.2678454 | Inhib_1 |
| 83 | KIRREL3 | 5.96E-125 | 1.6443847 | Inhib_1 |
| 84 | DGKD | 2.72E-124 | 2.8701792 | Inhib_1 |
| 85 | TMTC2 | 3.58E-123 | 1.6754656 | Inhib_1 |
| 86 | SPOCK1 | 2.16E-121 | 1.3581042 | Inhib_1 |
| 87 | NRG1 | 3.18E-120 | 1.6980324 | Inhib_1 |
| 88 | SPOCK2 | 2.68E-118 | 1.7288625 | Inhib_1 |
| 89 | MCTP1 | 1.04E-117 | 1.4594082 | Inhib_1 |
| 90 | KIAA1211 | 7.98E-116 | 2.8530478 | Inhib_1 |
| 91 | WWP1 | 4.61E-115 | 2.264925 | Inhib_1 |
| 92 | NYAP2 | 5.48E-114 | 2.4668312 | Inhib_1 |
| 93 | FAM110B | 1.61E-113 | 2.2157207 | Inhib_1 |
| 94 | PDE8B | 1.69E-111 | 1.705446 | Inhib_1 |
| 95 | ANKRD55 | 9.54E-110 | 3.6362095 | Inhib_1 |
| 96 | IGSF11 | 2.80E-109 | 2.2006118 | Inhib_1 |
| 97 | RORA | 6.49E-109 | 1.2058457 | Inhib_1 |
| 98 | PDGFD | 7.63E-109 | 4.5669746 | Inhib_1 |
| 99 | HECW2 | 1.22E-108 | 1.3004813 | Inhib_1 |
| 0 | ADARB2 | 0 | 4.801573 | Inhib_2_VIP |
| 1 | ERBB4 | 0 | 3.4349444 | Inhib_2_VIP |
| 2 | GALNTL6 | 0 | 3.0544052 | Inhib_2_VIP |
| 3 | SYNPR | 0 | 3.3504002 | Inhib_2_VIP |
| 4 | RGS12 | 0 | 3.5449312 | Inhib_2_VIP |
| 5 | ROBO1 | 0 | 2.431804 | Inhib_2_VIP |
| 6 | ROBO2 | 0 | 2.0817235 | Inhib_2_VIP |
| 7 | THSD7A | 0 | 3.3223324 | Inhib_2_VIP |
| 8 | DLX6-AS1 | 0 | 3.765646 | Inhib_2_VIP |
| 9 | CNTNAP2 | 0 | 1.6120083 | Inhib_2_VIP |
| 10 | NRXN3 | 4.97E-290 | 1.3276458 | Inhib_2_VIP |
| 11 | VIP | 8.52E-276 | 6.613277 | Inhib_2_VIP |
| 12 | PLD5 | 3.34E-273 | 2.9576564 | Inhib_2_VIP |
| 13 | TCF4 | 1.62E-250 | 1.2717156 | Inhib_2_VIP |
| 14 | GRIK2 | 7.85E-245 | 1.6170561 | Inhib_2_VIP |
| 15 | CXCL14 | 3.22E-235 | 4.035236 | Inhib_2_VIP |
| 16 | NPAS3 | 2.74E-216 | 1.7241079 | Inhib_2_VIP |
| 17 | DAB1 | 4.21E-213 | 1.5609576 | Inhib_2_VIP |
| 18 | PROX1 | 8.32E-207 | 4.389212 | Inhib_2_VIP |
| 19 | CNR1 | 2.28E-201 | 3.4909112 | Inhib_2_VIP |
| 20 | SLC24A3 | 1.27E-200 | 2.9546034 | Inhib_2_VIP |
| 21 | KCNT2 | 1.25E-197 | 2.2030551 | Inhib_2_VIP |
| 22 | FGF14 | 3.03E-192 | 1.1634754 | Inhib_2_VIP |
| 23 | NRXN1 | 2.43E-191 | 1.0754167 | Inhib_2_VIP |
| 24 | GRM7 | 3.05E-190 | 1.424357 | Inhib_2_VIP |
| 25 | NLGN1 | 1.36E-187 | 1.1755859 | Inhib_2_VIP |
| 26 | SCG2 | 5.11E-186 | 2.684606 | Inhib_2_VIP |

| | | | | |
|---|---|---|---|---|
| 27 | GAD1 | 2.41E-179 | 2.130581 | Inhib_2_VIP |
| 28 | DSCAM | 5.23E-178 | 1.259693 | Inhib_2_VIP |
| 29 | SNTG1 | 6.81E-170 | 1.2387807 | Inhib_2_VIP |
| 30 | GALNT13 | 3.52E-165 | 1.837948 | Inhib_2_VIP |
| 31 | CNTN4 | 6.71E-165 | 1.3277779 | Inhib_2_VIP |
| 32 | C8orf34 | 1.93E-161 | 2.3972774 | Inhib_2_VIP |
| 33 | ZNF536 | 4.10E-161 | 2.0161583 | Inhib_2_VIP |
| 34 | ARL4C | 1.41E-156 | 2.3288436 | Inhib_2_VIP |
| 35 | INPP4B | 1.46E-150 | 1.8339125 | Inhib_2_VIP |
| 36 | ZNF385D | 1.00E-149 | 1.5987116 | Inhib_2_VIP |
| 37 | FRMD4A | 5.07E-148 | 1.062358 | Inhib_2_VIP |
| 38 | SDK1 | 1.92E-142 | 1.9204403 | Inhib_2_VIP |
| 39 | CHRM3 | 3.89E-141 | 1.2303487 | Inhib_2_VIP |
| 40 | CSMD1 | 1.08E-124 | 1.0120245 | Inhib_2_VIP |
| 41 | CALB2 | 9.64E-123 | 5.64455 | Inhib_2_VIP |
| 42 | DNER | 3.69E-117 | 1.5419661 | Inhib_2_VIP |
| 43 | VWC2L | 1.88E-116 | 2.5194879 | Inhib_2_VIP |
| 44 | AUTS2 | 3.83E-116 | 0.96705765 | Inhib_2_VIP |
| 45 | DOCK10 | 1.30E-113 | 1.5964088 | Inhib_2_VIP |
| 46 | L3MBTL4 | 1.47E-113 | 2.3347268 | Inhib_2_VIP |
| 47 | ANO4 | 6.74E-112 | 1.576628 | Inhib_2_VIP |
| 48 | NR2F2-AS1 | 7.43E-112 | 4.3630624 | Inhib_2_VIP |
| 49 | GRID2 | 1.63E-111 | 1.253811 | Inhib_2_VIP |
| 50 | PDE3A | 2.43E-109 | 3.681671 | Inhib_2_VIP |
| 51 | KAZN | 1.60E-108 | 1.2395179 | Inhib_2_VIP |
| 52 | TAC3 | 1.61E-105 | 5.591078 | Inhib_2_VIP |
| 53 | LRP1B | 2.30E-99 | 0.8951958 | Inhib_2_VIP |
| 54 | CNTN5 | 2.60E-99 | 1.2760249 | Inhib_2_VIP |
| 55 | SOBP | 4.07E-99 | 1.3163468 | Inhib_2_VIP |
| 56 | CNTNAP4 | 4.92E-99 | 1.9363059 | Inhib_2_VIP |
| 57 | NFIB | 5.81E-96 | 1.1856046 | Inhib_2_VIP |
| 58 | GPHN | 8.23E-95 | 0.89965755 | Inhib_2_VIP |
| 59 | MIR325HG | 1.32E-92 | 2.5768294 | Inhib_2_VIP |
| 60 | ZNF804A | 3.95E-89 | 1.5506105 | Inhib_2_VIP |
| 61 | RP11-123O10.4 | 3.50E-88 | 1.4140491 | Inhib_2_VIP |
| 62 | MIR99AHG | 1.14E-85 | 0.87459517 | Inhib_2_VIP |
| 63 | PTPRE | 1.66E-85 | 1.6591724 | Inhib_2_VIP |
| 64 | SPOCK3 | 7.41E-84 | 1.1402929 | Inhib_2_VIP |
| 65 | MYT1L | 2.38E-83 | 0.9416605 | Inhib_2_VIP |
| 66 | TTC28 | 9.60E-83 | 1.3549858 | Inhib_2_VIP |
| 67 | SLC44A5 | 6.11E-81 | 1.2435783 | Inhib_2_VIP |
| 68 | LINC01322 | 1.25E-79 | 1.5959959 | Inhib_2_VIP |
| 69 | CHRNA2 | 2.03E-79 | 6.0066104 | Inhib_2_VIP |
| 70 | SLC6A1 | 1.61E-78 | 1.4932214 | Inhib_2_VIP |
| 71 | IGF1 | 5.17E-78 | 3.0557027 | Inhib_2_VIP |
| 72 | TIMP2 | 2.78E-75 | 1.646076 | Inhib_2_VIP |
| 73 | LSAMP | 6.53E-75 | 0.6844491 | Inhib_2_VIP |
| 74 | REV3L | 1.37E-73 | 1.1525501 | Inhib_2_VIP |
| 75 | CACNA1D | 1.85E-73 | 1.3607594 | Inhib_2_VIP |
| 76 | ASIC2 | 5.44E-73 | 0.914759 | Inhib_2_VIP |
| 77 | AP1S2 | 7.14E-73 | 2.2197201 | Inhib_2_VIP |
| 78 | MACROD2 | 8.69E-73 | 0.81607896 | Inhib_2_VIP |
| 79 | ATP1B1 | 2.10E-68 | 0.7807034 | Inhib_2_VIP |

| | | | | |
|---|---|---|---|---|
| 80 | SEZ6L | 4.93E-68 | 1.2815453 | Inhib_2_VIP |
| 81 | ADAMTS6 | 1.36E-64 | 3.1962628 | Inhib_2_VIP |
| 82 | NHS | 5.62E-64 | 1.9127539 | Inhib_2_VIP |
| 83 | GABRG3 | 5.90E-64 | 1.0451499 | Inhib_2_VIP |
| 84 | GRIP1 | 9.99E-64 | 1.2149981 | Inhib_2_VIP |
| 85 | FSTL5 | 2.33E-63 | 1.5214082 | Inhib_2_VIP |
| 86 | MYRIP | 4.37E-63 | 1.2331116 | Inhib_2_VIP |
| 87 | PARD3 | 1.39E-62 | 1.7264262 | Inhib_2_VIP |
| 88 | NR3C2 | 1.81E-62 | 1.265757 | Inhib_2_VIP |
| 89 | ZEB2 | 4.14E-62 | 0.8441566 | Inhib_2_VIP |
| 90 | RP11-384F7.2 | 5.66E-62 | 0.77661556 | Inhib_2_VIP |
| 91 | GRIK1 | 7.41E-62 | 1.1746678 | Inhib_2_VIP |
| 92 | CRH | 7.82E-62 | 3.2679257 | Inhib_2_VIP |
| 93 | LRRC4C | 2.00E-61 | 0.7342704 | Inhib_2_VIP |
| 94 | HDAC9 | 6.13E-61 | 0.8893904 | Inhib_2_VIP |
| 95 | SHISA8 | 1.56E-59 | 4.871413 | Inhib_2_VIP |
| 96 | DLX1 | 2.25E-59 | 3.2461696 | Inhib_2_VIP |
| 97 | GNAS | 5.61E-59 | 0.88449943 | Inhib_2_VIP |
| 98 | THSD7B | 1.04E-58 | 3.3441303 | Inhib_2_VIP |
| 99 | SLC6A1-AS1 | 1.11E-58 | 1.3452685 | Inhib_2_VIP |
| 0 | RELN | 3.44E-214 | 7.410138 | Inhib_3_SST |
| 1 | ADARB2 | 2.98E-179 | 4.5599365 | Inhib_3_SST |
| 2 | GRIK2 | 7.39E-165 | 2.9150243 | Inhib_3_SST |
| 3 | CXCL14 | 9.59E-161 | 5.955423 | Inhib_3_SST |
| 4 | FGF14 | 5.96E-134 | 1.9973066 | Inhib_3_SST |
| 5 | CNTN5 | 9.89E-120 | 2.585959 | Inhib_3_SST |
| 6 | GALNTL6 | 2.61E-119 | 2.8296216 | Inhib_3_SST |
| 7 | RP11-123O10.4 | 1.01E-118 | 2.9385927 | Inhib_3_SST |
| 8 | ERBB4 | 2.97E-117 | 3.1381464 | Inhib_3_SST |
| 9 | INPP4B | 1.19E-115 | 3.3890376 | Inhib_3_SST |
| 10 | FSTL5 | 8.37E-112 | 3.2597685 | Inhib_3_SST |
| 11 | GRIK1 | 1.87E-106 | 2.9716568 | Inhib_3_SST |
| 12 | DAB1 | 1.98E-105 | 2.2287145 | Inhib_3_SST |
| 13 | ZNF385D | 5.42E-101 | 2.8034666 | Inhib_3_SST |
| 14 | DLX6-AS1 | 3.91E-99 | 3.639159 | Inhib_3_SST |
| 15 | GRIP1 | 4.94E-97 | 2.4598308 | Inhib_3_SST |
| 16 | ROBO2 | 2.53E-89 | 2.2240875 | Inhib_3_SST |
| 17 | NXPH1 | 1.86E-87 | 2.8433328 | Inhib_3_SST |
| 18 | PLD5 | 7.70E-87 | 2.9627588 | Inhib_3_SST |
| 19 | SPOCK3 | 5.88E-86 | 2.393475 | Inhib_3_SST |
| 20 | DOCK10 | 7.38E-86 | 2.69511 | Inhib_3_SST |
| 21 | DNER | 3.48E-85 | 2.5530298 | Inhib_3_SST |
| 22 | PTPRM | 4.87E-85 | 2.433474 | Inhib_3_SST |
| 23 | CNTNAP2 | 8.15E-85 | 1.7213459 | Inhib_3_SST |
| 24 | MGAT4C | 1.10E-83 | 2.0727613 | Inhib_3_SST |
| 25 | HS3ST5 | 2.38E-79 | 3.0966687 | Inhib_3_SST |
| 26 | MYO16 | 3.50E-76 | 2.7068312 | Inhib_3_SST |
| 27 | ROBO1 | 7.94E-74 | 2.0139923 | Inhib_3_SST |
| 28 | CTB-107G13.1 | 3.65E-72 | 6.810575 | Inhib_3_SST |
| 29 | ZNF536 | 5.57E-72 | 2.3198576 | Inhib_3_SST |
| 30 | CNR1 | 8.89E-71 | 3.3123019 | Inhib_3_SST |
| 31 | RGS12 | 4.22E-69 | 2.4657156 | Inhib_3_SST |
| 32 | RBMS3 | 1.34E-66 | 2.5378618 | Inhib_3_SST |

| | | | | |
|---|---|---|---|---|
| 33 | KCNC2 | 1.08E-64 | 1.7014459 | Inhib_3_SST |
| 34 | EPHA6 | 2.93E-63 | 1.8457642 | Inhib_3_SST |
| 35 | SLC6A1 | 4.96E-63 | 2.55532 | Inhib_3_SST |
| 36 | GAD2 | 1.15E-62 | 3.1051953 | Inhib_3_SST |
| 37 | KAZN | 1.23E-62 | 1.740171 | Inhib_3_SST |
| 38 | MACROD2 | 2.89E-60 | 1.5940065 | Inhib_3_SST |
| 39 | PRR16 | 1.37E-59 | 2.5762825 | Inhib_3_SST |
| 40 | CSMD3 | 1.91E-59 | 1.478711 | Inhib_3_SST |
| 41 | GAD1 | 2.97E-58 | 2.2189858 | Inhib_3_SST |
| 42 | CHRM3 | 3.14E-57 | 1.5343024 | Inhib_3_SST |
| 43 | RAB3C | 4.21E-57 | 1.75149 | Inhib_3_SST |
| 44 | LINGO2 | 1.95E-55 | 1.5857489 | Inhib_3_SST |
| 45 | AC074363.1 | 4.52E-55 | 2.0685873 | Inhib_3_SST |
| 46 | NFIB | 8.49E-54 | 1.6318779 | Inhib_3_SST |
| 47 | ALK | 1.56E-53 | 2.814555 | Inhib_3_SST |
| 48 | SGK1 | 2.15E-53 | 2.0466585 | Inhib_3_SST |
| 49 | FRAS1 | 1.89E-51 | 2.2942417 | Inhib_3_SST |
| 50 | AP1S2 | 3.34E-51 | 2.8887784 | Inhib_3_SST |
| 51 | FGF13 | 3.75E-51 | 2.3825555 | Inhib_3_SST |
| 52 | SGCZ | 1.13E-50 | 2.226512 | Inhib_3_SST |
| 53 | UNC5D | 3.98E-49 | 1.638573 | Inhib_3_SST |
| 54 | GRIA2 | 2.85E-47 | 1.1820022 | Inhib_3_SST |
| 55 | MAML3 | 4.73E-47 | 2.7103174 | Inhib_3_SST |
| 56 | CCDC85A | 1.07E-46 | 1.9166602 | Inhib_3_SST |
| 57 | SLC6A1-AS1 | 1.45E-46 | 2.311117 | Inhib_3_SST |
| 58 | PCDH15 | 1.30E-45 | 1.9406419 | Inhib_3_SST |
| 59 | CHSY3 | 1.89E-45 | 1.5619453 | Inhib_3_SST |
| 60 | NCAM2 | 7.35E-45 | 1.4700109 | Inhib_3_SST |
| 61 | NR2F2-AS1 | 7.15E-44 | 3.9028056 | Inhib_3_SST |
| 62 | SCG2 | 2.23E-42 | 2.2495105 | Inhib_3_SST |
| 63 | FREM1 | 9.82E-40 | 3.8546715 | Inhib_3_SST |
| 64 | PIP5K1B | 1.80E-38 | 2.176815 | Inhib_3_SST |
| 65 | CNTNAP4 | 1.83E-38 | 2.176798 | Inhib_3_SST |
| 66 | SHISA9 | 1.72E-37 | 1.4809381 | Inhib_3_SST |
| 67 | EGFR | 2.16E-37 | 3.3188775 | Inhib_3_SST |
| 68 | RP11-223C24.1 | 1.70E-36 | 3.0571592 | Inhib_3_SST |
| 69 | ADRA1A | 3.18E-35 | 3.0668633 | Inhib_3_SST |
| 70 | MCTP1 | 1.45E-34 | 1.4172505 | Inhib_3_SST |
| 71 | SORCS3 | 8.32E-34 | 1.8676987 | Inhib_3_SST |
| 72 | ARL4C | 1.49E-33 | 1.9805151 | Inhib_3_SST |
| 73 | IL1RAPL2 | 5.37E-33 | 1.8532312 | Inhib_3_SST |
| 74 | TOX3 | 1.98E-32 | 3.2890851 | Inhib_3_SST |
| 75 | XKR4 | 3.04E-32 | 1.4030122 | Inhib_3_SST |
| 76 | MTSS1 | 3.96E-32 | 1.9287984 | Inhib_3_SST |
| 77 | KCNAB1 | 5.36E-32 | 1.5255439 | Inhib_3_SST |
| 78 | NYAP2 | 5.54E-32 | 2.373766 | Inhib_3_SST |
| 79 | IL1RAPL1 | 8.54E-32 | 1.0284522 | Inhib_3_SST |
| 80 | GABRB1 | 9.97E-32 | 1.0953276 | Inhib_3_SST |
| 81 | GABRB2 | 1.86E-31 | 1.0638087 | Inhib_3_SST |
| 82 | EXT1 | 2.00E-31 | 1.4643521 | Inhib_3_SST |
| 83 | HDAC9 | 2.22E-31 | 1.2130822 | Inhib_3_SST |
| 84 | C8orf34 | 1.33E-30 | 1.9480021 | Inhib_3_SST |
| 85 | PLPPR5 | 1.49E-30 | 2.0567162 | Inhib_3_SST |

| | | | | |
|---|---|---|---|---|
| 86 | FAM110B | 2.14E-30 | 1.933418 | Inhib_3_SST |
| 87 | DGKB | 6.14E-30 | 1.1429664 | Inhib_3_SST |
| 88 | GRM8 | 6.86E-30 | 2.0339222 | Inhib_3_SST |
| 89 | SST | 1.59E-29 | 3.0307908 | Inhib_3_SST |
| 90 | BCL11B | 1.78E-29 | 2.1997724 | Inhib_3_SST |
| 91 | NPAS3 | 2.06E-29 | 1.1699262 | Inhib_3_SST |
| 92 | QKI | 3.30E-29 | 0.87941563 | Inhib_3_SST |
| 93 | GRIN3A | 1.69E-28 | 2.210673 | Inhib_3_SST |
| 94 | FNBP1L | 2.70E-28 | 1.9937453 | Inhib_3_SST |
| 95 | CACNA1B | 3.06E-28 | 1.1612635 | Inhib_3_SST |
| 96 | SPOCK1 | 6.45E-28 | 1.1061946 | Inhib_3_SST |
| 97 | TCF4 | 9.44E-28 | 0.9201434 | Inhib_3_SST |
| 98 | ABI1 | 9.44E-28 | 1.6331947 | Inhib_3_SST |
| 99 | SV2C | 1.20E-27 | 3.0719647 | Inhib_3_SST |
| 0 | FOS | 0.002349597 | 6.21449 | Inhib_4_SST |
| 1 | GRIK1 | 0.002930997 | 5.2366433 | Inhib_4_SST |
| 2 | SPOCK3 | 0.050188712 | 3.098489 | Inhib_4_SST |
| 3 | ZNF385D | 0.050188712 | 3.2464461 | Inhib_4_SST |
| 4 | SST | 0.122397124 | 6.3037515 | Inhib_4_SST |
| 5 | RBFOX1 | 0.12306227 | 1.8262248 | Inhib_4_SST |
| 6 | JUNB | 0.130379647 | 4.492704 | Inhib_4_SST |
| 7 | EGR3 | 0.130379647 | 4.464183 | Inhib_4_SST |
| 8 | EGR1 | 0.140407085 | 4.1168065 | Inhib_4_SST |
| 9 | DUSP1 | 0.140407085 | 4.1599264 | Inhib_4_SST |
| 10 | ELAVL2 | 0.165698683 | 2.4396377 | Inhib_4_SST |
| 11 | LINC01122 | 0.173387194 | 2.5759234 | Inhib_4_SST |
| 12 | NXPH1 | 0.201289349 | 3.1806455 | Inhib_4_SST |
| 13 | BCL11A | 0.288896796 | 2.2247314 | Inhib_4_SST |
| 14 | NPY | 0.347306022 | 6.6680408 | Inhib_4_SST |
| 15 | NETO2 | 0.381970034 | 3.1117258 | Inhib_4_SST |
| 16 | ROBO2 | 0.427103224 | 2.456508 | Inhib_4_SST |
| 17 | NR4A1 | 0.427103224 | 4.66261 | Inhib_4_SST |
| 18 | CADPS | 0.427103224 | 1.8641225 | Inhib_4_SST |
| 19 | TENM1 | 0.450355752 | 2.8089745 | Inhib_4_SST |
| 20 | ARX | 0.485654279 | 4.0598655 | Inhib_4_SST |
| 21 | RUNX1T1 | 0.578340943 | 1.9563954 | Inhib_4_SST |
| 22 | NRXN3 | 0.594110689 | 1.4481182 | Inhib_4_SST |
| 23 | PDIA2 | 0.594110689 | 2.2962804 | Inhib_4_SST |
| 24 | HSPA1A | 0.633493625 | 3.7133334 | Inhib_4_SST |
| 25 | KIAA1217 | 0.68222634 | 2.2213612 | Inhib_4_SST |
| 26 | SGCZ | 0.68222634 | 3.1580644 | Inhib_4_SST |
| 27 | ELOVL5 | 0.749696587 | 3.5958724 | Inhib_4_SST |
| 28 | VGF | 0.755362006 | 3.5759106 | Inhib_4_SST |
| 29 | WLS | 0.755362006 | 3.4189296 | Inhib_4_SST |
| 30 | SOX6 | 0.774965367 | 2.4641027 | Inhib_4_SST |
| 31 | DPYSL5 | 0.812349574 | 3.3080704 | Inhib_4_SST |
| 32 | TNIK | 0.889911977 | 1.7568779 | Inhib_4_SST |
| 33 | ZNF608 | 0.98616496 | 3.0971904 | Inhib_4_SST |
| 34 | ROBO1 | 0.98616496 | 2.3928525 | Inhib_4_SST |
| 69 | EGR2 | 1 | 5.9902983 | Inhib_4_SST |
| 70 | LRP8 | 1 | 3.0387657 | Inhib_4_SST |
| 71 | PCLO | 1 | 1.2524709 | Inhib_4_SST |
| 72 | GRID2 | 1 | 1.9412632 | Inhib_4_SST |

| | | | | |
|---|---|---|---|---|
| 73 | NTRK2 | 1 | 1.3822819 | Inhib_4_SST |
| 74 | TSPYL2 | 1 | 1.950128 | Inhib_4_SST |
| 75 | PTPN4 | 1 | 1.9319189 | Inhib_4_SST |
| 76 | SHTN1 | 1 | 1.9399729 | Inhib_4_SST |
| 77 | MAP7D1 | 1 | 2.3799732 | Inhib_4_SST |
| 78 | APLP2 | 1 | 1.5540607 | Inhib_4_SST |
| 79 | PRNP | 1 | 1.3498534 | Inhib_4_SST |
| 80 | CTD-2537O9.1 | 1 | 1.4941429 | Inhib_4_SST |
| 81 | ATP6V1A | 1 | 1.5314585 | Inhib_4_SST |
| 82 | CYTH3 | 1 | 2.2043931 | Inhib_4_SST |
| 83 | PWAR6 | 1 | 1.3216169 | Inhib_4_SST |
| 84 | STXBP6 | 1 | 2.6078525 | Inhib_4_SST |
| 85 | CDH9 | 1 | 1.8875915 | Inhib_4_SST |
| 86 | LIMCH1 | 1 | 1.2471821 | Inhib_4_SST |
| 87 | RP11-123O10.4 | 1 | 1.9135909 | Inhib_4_SST |
| 88 | MRPL41 | 1 | 1.2221482 | Inhib_4_SST |
| 89 | SLC24A3 | 1 | 2.1897418 | Inhib_4_SST |
| 90 | FAM135B | 1 | 1.5454088 | Inhib_4_SST |
| 91 | RND3 | 1 | 3.61287 | Inhib_4_SST |
| 92 | IGSF8 | 1 | 1.4894868 | Inhib_4_SST |
| 93 | NOS1 | 1 | 4.0113754 | Inhib_4_SST |
| 94 | ANKRD36C | 1 | 1.3348179 | Inhib_4_SST |
| 95 | PCDH15 | 1 | 1.7682943 | Inhib_4_SST |
| 96 | CPT1C | 1 | 2.0493052 | Inhib_4_SST |
| 97 | KIZ | 1 | 2.128012 | Inhib_4_SST |
| 68 | KIF26B | 1 | 3.1970897 | Inhib_4_SST |
| 67 | SMARCA2 | 1 | 1.5989857 | Inhib_4_SST |
| 49 | THOC2 | 1 | 2.1106832 | Inhib_4_SST |
| 65 | NCOA2 | 1 | 1.5208024 | Inhib_4_SST |
| 35 | RASGRF2 | 1 | 1.7183758 | Inhib_4_SST |
| 36 | LINC00599 | 1 | 1.861266 | Inhib_4_SST |
| 37 | GAD1 | 1 | 2.901789 | Inhib_4_SST |
| 38 | TPD52L2 | 1 | 2.867183 | Inhib_4_SST |
| 39 | RNF157 | 1 | 2.3756979 | Inhib_4_SST |
| 40 | ARC | 1 | 4.5813093 | Inhib_4_SST |
| 41 | PHF20L1 | 1 | 1.8591967 | Inhib_4_SST |
| 42 | TCF4 | 1 | 1.3772413 | Inhib_4_SST |
| 43 | TENM3 | 1 | 2.075845 | Inhib_4_SST |
| 44 | EIF4A2 | 1 | 1.5044861 | Inhib_4_SST |
| 45 | ARHGAP20 | 1 | 2.5018244 | Inhib_4_SST |
| 46 | NAV2 | 1 | 1.7789277 | Inhib_4_SST |
| 47 | RBMS3 | 1 | 1.705714 | Inhib_4_SST |
| 48 | KCTD16 | 1 | 1.4791816 | Inhib_4_SST |
| 66 | GRIK2 | 1 | 1.3567588 | Inhib_4_SST |
| 98 | SYNPR | 1 | 1.5494747 | Inhib_4_SST |
| 51 | COL7A1 | 1 | 3.5364594 | Inhib_4_SST |
| 52 | MIR181A1HG | 1 | 2.1298347 | Inhib_4_SST |
| 53 | PTPRM | 1 | 1.661473 | Inhib_4_SST |
| 54 | CACNG8 | 1 | 1.9348547 | Inhib_4_SST |
| 55 | BACH1 | 1 | 3.3511858 | Inhib_4_SST |
| 56 | MPDZ | 1 | 1.4835486 | Inhib_4_SST |
| 57 | SV2A | 1 | 1.2619113 | Inhib_4_SST |
| 58 | DLGAP1 | 1 | 1.2389535 | Inhib_4_SST |

| | | | | |
|---|---|---|---|---|
| 59 | DNM2 | 1 | 3.187824 | Inhib_4_SST |
| 60 | XKR4 | 1 | 1.6930987 | Inhib_4_SST |
| 61 | DNM3 | 1 | 1.4108377 | Inhib_4_SST |
| 62 | NRIP3 | 1 | 2.8011298 | Inhib_4_SST |
| 63 | NCAM2 | 1 | 1.529204 | Inhib_4_SST |
| 64 | ADCY8 | 1 | 2.7479682 | Inhib_4_SST |
| 50 | MXI1 | 1 | 2.1263304 | Inhib_4_SST |
| 99 | PCBP4 | 1 | 1.9675305 | Inhib_4_SST |
| 0 | ERBB4 | 0 | 4.3909163 | Inhib_5 |
| 28 | MAGI1 | 0 | 1.6323351 | Inhib_5 |
| 27 | CADPS | 0 | 1.6594051 | Inhib_5 |
| 26 | MAGI2 | 0 | 1.4026402 | Inhib_5 |
| 25 | IL1RAPL1 | 0 | 1.6491156 | Inhib_5 |
| 23 | KCNAB1 | 0 | 2.3331504 | Inhib_5 |
| 22 | TENM3 | 0 | 2.3689318 | Inhib_5 |
| 21 | KLF12 | 0 | 2.3290892 | Inhib_5 |
| 20 | PAM | 0 | 2.146853 | Inhib_5 |
| 19 | KCND2 | 0 | 1.9213438 | Inhib_5 |
| 18 | PRKG1 | 0 | 2.2403138 | Inhib_5 |
| 17 | SPOCK3 | 0 | 2.3418508 | Inhib_5 |
| 16 | MYO16 | 0 | 2.9716156 | Inhib_5 |
| 15 | CNTNAP3B | 0 | 3.1953838 | Inhib_5 |
| 24 | ASTN2 | 0 | 2.0366607 | Inhib_5 |
| 13 | PTPRM | 0 | 2.8141947 | Inhib_5 |
| 12 | SOX6 | 0 | 3.170347 | Inhib_5 |
| 11 | SLIT2 | 0 | 2.787581 | Inhib_5 |
| 10 | ZNF804A | 0 | 3.2496212 | Inhib_5 |
| 9 | DLGAP1 | 0 | 1.9118167 | Inhib_5 |
| 8 | DPP10 | 0 | 2.8245125 | Inhib_5 |
| 7 | RP11-123O10.4 | 0 | 3.07077 | Inhib_5 |
| 6 | KIAA1217 | 0 | 3.1514266 | Inhib_5 |
| 5 | FGF12 | 0 | 2.1922512 | Inhib_5 |
| 14 | GRIP1 | 0 | 2.4697852 | Inhib_5 |
| 4 | ZNF385D | 0 | 3.4456642 | Inhib_5 |
| 3 | CNTNAP2 | 0 | 2.2800043 | Inhib_5 |
| 2 | NXPH1 | 0 | 3.8116605 | Inhib_5 |
| 1 | KCNC2 | 0 | 2.898765 | Inhib_5 |
| 29 | EPHA6 | 2.80E-301 | 2.178535 | Inhib_5 |
| 30 | ANO4 | 1.72E-300 | 2.2461805 | Inhib_5 |
| 31 | RP11-444D3.1 | 2.89E-295 | 2.2967992 | Inhib_5 |
| 32 | BTBD11 | 3.67E-294 | 3.578428 | Inhib_5 |
| 33 | VWC2 | 1.41E-291 | 2.7649517 | Inhib_5 |
| 34 | ZNF536 | 1.74E-291 | 2.2956922 | Inhib_5 |
| 35 | GAD1 | 2.17E-288 | 2.4734395 | Inhib_5 |
| 36 | KAZN | 2.47E-286 | 1.7308921 | Inhib_5 |
| 37 | CNTNAP3 | 4.49E-286 | 3.4428155 | Inhib_5 |
| 38 | RBMS3 | 3.62E-276 | 2.3633811 | Inhib_5 |
| 39 | FAM19A2 | 1.16E-273 | 1.760131 | Inhib_5 |
| 40 | XKR4 | 2.33E-264 | 1.8117977 | Inhib_5 |
| 41 | TMEM132C | 3.31E-263 | 2.9289618 | Inhib_5 |
| 42 | TENM1 | 2.18E-259 | 2.5851912 | Inhib_5 |
| 43 | CNTN5 | 1.70E-257 | 1.7817792 | Inhib_5 |
| 44 | ZMAT4 | 1.06E-254 | 2.0704381 | Inhib_5 |

| | | | | |
|---|---|---|---|---|
| 45 | SGCZ | 3.58E-250 | 2.2079997 | Inhib_5 |
| 46 | FRMD5 | 3.75E-249 | 1.5578716 | Inhib_5 |
| 47 | LRRC4C | 3.08E-248 | 1.5543268 | Inhib_5 |
| 48 | TENM2 | 4.56E-248 | 1.5589496 | Inhib_5 |
| 49 | AC010127.3 | 1.87E-243 | 2.1762428 | Inhib_5 |
| 50 | SUPT3H | 7.64E-243 | 1.6832399 | Inhib_5 |
| 51 | SDK1 | 3.96E-240 | 2.1395733 | Inhib_5 |
| 52 | ESRRG | 1.64E-230 | 1.769458 | Inhib_5 |
| 53 | SNRPN | 1.40E-229 | 2.183336 | Inhib_5 |
| 54 | SLC44A5 | 1.46E-226 | 1.7306377 | Inhib_5 |
| 55 | ADAMTS17 | 1.36E-222 | 3.3244576 | Inhib_5 |
| 56 | HCN1 | 5.28E-220 | 1.4713415 | Inhib_5 |
| 57 | KIF26B | 1.09E-218 | 2.590035 | Inhib_5 |
| 58 | RBFOX1 | 2.36E-216 | 1.4186307 | Inhib_5 |
| 59 | ELAVL2 | 1.87E-209 | 1.7075952 | Inhib_5 |
| 60 | ENOX1 | 2.24E-209 | 1.4437504 | Inhib_5 |
| 61 | NEAT1 | 2.35E-207 | 2.327634 | Inhib_5 |
| 62 | RPS6KA2 | 3.23E-207 | 1.8693926 | Inhib_5 |
| 63 | TMEM108 | 3.62E-205 | 1.5174952 | Inhib_5 |
| 64 | GAD2 | 2.15E-202 | 2.545417 | Inhib_5 |
| 65 | TIAM1 | 4.07E-199 | 1.8292805 | Inhib_5 |
| 66 | RASGRF2 | 1.55E-197 | 1.6234939 | Inhib_5 |
| 67 | SLC6A1-AS1 | 1.99E-193 | 2.1480184 | Inhib_5 |
| 68 | KCTD8 | 4.65E-193 | 1.7264174 | Inhib_5 |
| 69 | TAC1 | 6.22E-192 | 3.0953267 | Inhib_5 |
| 70 | GRIA4 | 9.70E-192 | 1.399101 | Inhib_5 |
| 71 | ATRNL1 | 6.09E-190 | 1.3303963 | Inhib_5 |
| 72 | DTNA | 1.05E-189 | 1.2773614 | Inhib_5 |
| 73 | CACNB4 | 5.46E-189 | 1.3219577 | Inhib_5 |
| 74 | KCNMB2-AS1 | 4.69E-188 | 1.835035 | Inhib_5 |
| 75 | NHS | 3.96E-187 | 2.5304208 | Inhib_5 |
| 76 | CNTNAP5 | 8.25E-187 | 1.5085404 | Inhib_5 |
| 77 | SLC4A10 | 2.83E-175 | 1.2404541 | Inhib_5 |
| 78 | INPP4B | 3.52E-171 | 1.5195752 | Inhib_5 |
| 79 | SHISA9 | 4.77E-171 | 1.5357763 | Inhib_5 |
| 80 | FHIT | 2.33E-170 | 1.375722 | Inhib_5 |
| 81 | OSBPL3 | 4.61E-170 | 1.8939914 | Inhib_5 |
| 82 | GABRG3 | 1.14E-168 | 1.4518633 | Inhib_5 |
| 83 | SAMD5 | 1.15E-164 | 2.8644726 | Inhib_5 |
| 84 | LRRTM4 | 4.17E-160 | 1.3586705 | Inhib_5 |
| 85 | PTCHD4 | 4.50E-160 | 2.1479964 | Inhib_5 |
| 86 | PPARGC1A | 6.74E-159 | 2.053142 | Inhib_5 |
| 87 | TRPC4 | 1.69E-157 | 3.17778 | Inhib_5 |
| 88 | SETBP1 | 1.77E-157 | 1.4215698 | Inhib_5 |
| 89 | OXR1 | 8.13E-157 | 1.1814487 | Inhib_5 |
| 90 | GRM5 | 1.98E-156 | 1.1667564 | Inhib_5 |
| 91 | PLCXD3 | 3.99E-155 | 2.5774257 | Inhib_5 |
| 92 | MMP16 | 2.06E-151 | 1.239407 | Inhib_5 |
| 93 | LRRFIP1 | 9.33E-151 | 1.5357162 | Inhib_5 |
| 94 | MEF2C | 9.38E-149 | 1.1076093 | Inhib_5 |
| 95 | SLC9A9 | 1.84E-148 | 2.147879 | Inhib_5 |
| 96 | KLHL5 | 6.27E-147 | 2.2849386 | Inhib_5 |
| 97 | MALAT1 | 4.71E-146 | 1.0674903 | Inhib_5 |

| | | | | |
|---|---|---|---|---|
| 98 | ANK1 | 6.42E-145 | 2.7924843 | Inhib_5 |
| 99 | UNC5D | 3.05E-143 | 1.2525486 | Inhib_5 |
| 0 | GRIK1 | 0 | 5.008804 | Inhib_6_SST |
| 1 | NXPH1 | 0 | 3.5003872 | Inhib_6_SST |
| 2 | GRIK2 | 0 | 2.536988 | Inhib_6_SST |
| 3 | SYNPR | 0 | 3.4584284 | Inhib_6_SST |
| 4 | SST | 0 | 6.1398005 | Inhib_6_SST |
| 5 | KIAA1217 | 0 | 2.9387765 | Inhib_6_SST |
| 6 | ROBO2 | 0 | 2.6937494 | Inhib_6_SST |
| 7 | SPOCK3 | 0 | 3.0619628 | Inhib_6_SST |
| 8 | ROBO1 | 0 | 2.8697674 | Inhib_6_SST |
| 9 | CNTNAP2 | 8.28E-305 | 1.9546367 | Inhib_6_SST |
| 10 | XKR4 | 6.96E-295 | 2.435025 | Inhib_6_SST |
| 11 | NRXN3 | 1.13E-275 | 1.6834093 | Inhib_6_SST |
| 12 | RP11-123O10.4 | 6.82E-244 | 2.539598 | Inhib_6_SST |
| 13 | ASTN2 | 1.23E-233 | 2.1007938 | Inhib_6_SST |
| 14 | PCDH11X | 2.02E-228 | 2.9089708 | Inhib_6_SST |
| 15 | RUNX1T1 | 1.08E-222 | 2.0716352 | Inhib_6_SST |
| 16 | SOX6 | 3.56E-221 | 2.7596521 | Inhib_6_SST |
| 17 | ELAVL2 | 1.55E-218 | 2.1813657 | Inhib_6_SST |
| 18 | CDH9 | 8.08E-213 | 2.5224724 | Inhib_6_SST |
| 19 | RBFOX1 | 1.08E-212 | 1.6043308 | Inhib_6_SST |
| 20 | PCDH15 | 1.19E-205 | 2.839449 | Inhib_6_SST |
| 21 | TENM3 | 1.81E-204 | 2.3405452 | Inhib_6_SST |
| 22 | NPAS3 | 1.21E-195 | 2.109823 | Inhib_6_SST |
| 23 | TRHDE | 1.82E-191 | 2.796332 | Inhib_6_SST |
| 24 | PAM | 1.03E-187 | 1.9727356 | Inhib_6_SST |
| 25 | GAD1 | 3.88E-179 | 2.551346 | Inhib_6_SST |
| 26 | GRID2 | 1.13E-177 | 1.8834931 | Inhib_6_SST |
| 27 | OXR1 | 4.57E-172 | 1.4222293 | Inhib_6_SST |
| 28 | GRIP1 | 1.03E-170 | 2.0533478 | Inhib_6_SST |
| 29 | PTPRM | 1.50E-169 | 2.1607304 | Inhib_6_SST |
| 30 | PLCH1 | 1.50E-169 | 2.682458 | Inhib_6_SST |
| 31 | NETO1 | 5.15E-161 | 2.0416212 | Inhib_6_SST |
| 32 | STXBP6 | 9.27E-160 | 2.695019 | Inhib_6_SST |
| 33 | GRIN3A | 7.79E-159 | 3.2665014 | Inhib_6_SST |
| 34 | RAB3C | 6.00E-153 | 1.7215096 | Inhib_6_SST |
| 35 | CDH13 | 3.59E-150 | 2.0367515 | Inhib_6_SST |
| 36 | TIAM1 | 8.52E-148 | 2.0350595 | Inhib_6_SST |
| 37 | ZNF385D | 9.45E-140 | 1.9963682 | Inhib_6_SST |
| 38 | KCNMB2-AS1 | 1.66E-139 | 2.1302853 | Inhib_6_SST |
| 39 | CADPS | 8.52E-138 | 1.3496783 | Inhib_6_SST |
| 40 | KCNC2 | 3.92E-137 | 1.4849082 | Inhib_6_SST |
| 41 | MGAT4C | 7.70E-137 | 1.6923785 | Inhib_6_SST |
| 42 | GPC6 | 1.78E-133 | 1.7818232 | Inhib_6_SST |
| 43 | PCDH11Y | 2.34E-133 | 2.484908 | Inhib_6_SST |
| 44 | RBMS3 | 2.23E-129 | 2.1151948 | Inhib_6_SST |
| 45 | DLGAP1 | 4.13E-128 | 1.2320694 | Inhib_6_SST |
| 46 | TENM2 | 2.79E-124 | 1.3686045 | Inhib_6_SST |
| 47 | ST6GALNAC5 | 2.18E-121 | 1.5993037 | Inhib_6_SST |
| 48 | SHISA6 | 8.39E-115 | 2.667278 | Inhib_6_SST |
| 49 | KIF26B | 1.51E-112 | 2.558589 | Inhib_6_SST |
| 50 | SLC8A1 | 1.67E-112 | 1.2847271 | Inhib_6_SST |

| | | | | |
|---|---|---|---|---|
| 51 | GRIA3 | 5.45E-111 | 1.2815629 | Inhib_6_SST |
| 52 | AUTS2 | 8.34E-111 | 1.1625812 | Inhib_6_SST |
| 53 | COL25A1 | 7.67E-110 | 3.5498385 | Inhib_6_SST |
| 54 | WLS | 5.04E-109 | 2.813723 | Inhib_6_SST |
| 55 | RASGRF2 | 3.45E-108 | 1.5920931 | Inhib_6_SST |
| 56 | LIMCH1 | 1.34E-107 | 1.196598 | Inhib_6_SST |
| 57 | LHFPL3 | 6.47E-103 | 1.8800339 | Inhib_6_SST |
| 58 | ADGRL2 | 7.24E-102 | 1.3107928 | Inhib_6_SST |
| 59 | GRIA1 | 5.93E-100 | 1.3316355 | Inhib_6_SST |
| 60 | TCF4 | 4.55E-95 | 1.0076889 | Inhib_6_SST |
| 61 | TMTC1 | 4.72E-95 | 1.8736409 | Inhib_6_SST |
| 62 | PCLO | 8.15E-95 | 1.1326504 | Inhib_6_SST |
| 63 | TENM1 | 2.64E-93 | 2.1884754 | Inhib_6_SST |
| 64 | DPP6 | 3.27E-92 | 1.089748 | Inhib_6_SST |
| 65 | SNRPN | 3.18E-88 | 1.7919328 | Inhib_6_SST |
| 66 | KLHL5 | 3.24E-88 | 2.2308352 | Inhib_6_SST |
| 67 | DAB1 | 1.41E-87 | 1.2919837 | Inhib_6_SST |
| 68 | BACH1 | 3.10E-86 | 2.6165528 | Inhib_6_SST |
| 69 | CACNA2D3 | 3.10E-86 | 1.2887454 | Inhib_6_SST |
| 70 | LINC01322 | 1.53E-80 | 1.6600765 | Inhib_6_SST |
| 71 | RAB3B | 6.22E-80 | 3.6088405 | Inhib_6_SST |
| 72 | AAK1 | 2.70E-79 | 1.0973802 | Inhib_6_SST |
| 73 | TIMP2 | 9.03E-76 | 1.9092631 | Inhib_6_SST |
| 74 | GRIN2B | 4.21E-75 | 0.9969018 | Inhib_6_SST |
| 75 | GAD2 | 1.82E-73 | 2.0650291 | Inhib_6_SST |
| 76 | ADCY8 | 5.44E-70 | 2.220764 | Inhib_6_SST |
| 77 | PITPNC1 | 1.86E-66 | 1.2752434 | Inhib_6_SST |
| 78 | DAPK1 | 2.25E-66 | 1.4660113 | Inhib_6_SST |
| 79 | TNIK | 1.34E-65 | 1.0022274 | Inhib_6_SST |
| 80 | FRMD5 | 1.39E-65 | 0.96974945 | Inhib_6_SST |
| 81 | GRIK3 | 2.20E-65 | 2.5155003 | Inhib_6_SST |
| 82 | BCL11A | 7.56E-65 | 1.3560053 | Inhib_6_SST |
| 83 | TMSB4X | 4.57E-64 | 0.9237217 | Inhib_6_SST |
| 84 | NETO2 | 2.11E-62 | 1.8167403 | Inhib_6_SST |
| 85 | PAWR | 8.94E-62 | 3.6257463 | Inhib_6_SST |
| 86 | SLC24A3 | 3.89E-61 | 2.037588 | Inhib_6_SST |
| 87 | SMYD3 | 6.79E-61 | 0.9446469 | Inhib_6_SST |
| 88 | NCOA2 | 1.34E-60 | 1.228088 | Inhib_6_SST |
| 89 | SLC24A2 | 1.85E-60 | 0.86766136 | Inhib_6_SST |
| 90 | RPS6KA2 | 1.57E-59 | 1.3578677 | Inhib_6_SST |
| 91 | ANKRD36 | 5.20E-59 | 1.1151564 | Inhib_6_SST |
| 92 | ANKRD36C | 7.40E-59 | 1.1583863 | Inhib_6_SST |
| 93 | CHRM3 | 1.86E-58 | 0.9644813 | Inhib_6_SST |
| 94 | MAGI1 | 5.19E-58 | 0.851958 | Inhib_6_SST |
| 95 | ADGRL3 | 7.93E-58 | 0.8469482 | Inhib_6_SST |
| 96 | ADCY2 | 1.40E-57 | 1.0929767 | Inhib_6_SST |
| 97 | TMTC2 | 9.76E-57 | 1.2111641 | Inhib_6_SST |
| 98 | LINC00693 | 3.39E-55 | 1.9908838 | Inhib_6_SST |
| 99 | MYT1L | 7.42E-55 | 0.8437947 | Inhib_6_SST |
| 0 | ERBB4 | 0 | 3.8109565 | Inhib_7_PVALB |
| 1 | GAD1 | 9.39E-299 | 3.6304529 | Inhib_7_PVALB |
| 2 | TAC1 | 2.86E-283 | 4.632765 | Inhib_7_PVALB |
| 3 | KCNC2 | 1.37E-281 | 2.617183 | Inhib_7_PVALB |

| | | | | |
|---|---|---|---|---|
| 4 | ZNF385D | 6.70E-280 | 3.1923168 | Inhib_7_PVALB |
| 5 | NXPH1 | 6.15E-253 | 3.1431618 | Inhib_7_PVALB |
| 6 | ZNF804A | 1.40E-234 | 2.936613 | Inhib_7_PVALB |
| 7 | CPLX1 | 1.18E-231 | 2.7292533 | Inhib_7_PVALB |
| 8 | GAD2 | 4.21E-224 | 3.4958518 | Inhib_7_PVALB |
| 9 | SOX6 | 2.15E-205 | 2.61331 | Inhib_7_PVALB |
| 10 | ATP1B1 | 3.01E-203 | 2.1055315 | Inhib_7_PVALB |
| 11 | PTPRM | 1.48E-200 | 2.3503509 | Inhib_7_PVALB |
| 12 | KIAA1217 | 1.24E-199 | 2.3060844 | Inhib_7_PVALB |
| 13 | DPP10 | 2.61E-192 | 2.2820914 | Inhib_7_PVALB |
| 14 | PAM | 2.46E-190 | 2.0563989 | Inhib_7_PVALB |
| 15 | SLIT2 | 1.48E-189 | 2.2610528 | Inhib_7_PVALB |
| 16 | SPARCL1 | 2.00E-186 | 2.1062453 | Inhib_7_PVALB |
| 17 | RP11-123O10.4 | 4.05E-183 | 2.3283155 | Inhib_7_PVALB |
| 18 | SPOCK3 | 4.34E-181 | 2.1316388 | Inhib_7_PVALB |
| 19 | ELAVL2 | 8.72E-177 | 2.0299306 | Inhib_7_PVALB |
| 20 | FGF12 | 5.60E-175 | 1.6185713 | Inhib_7_PVALB |
| 21 | SLC6A1 | 1.60E-162 | 2.377839 | Inhib_7_PVALB |
| 22 | CNTNAP3B | 1.60E-162 | 2.5447536 | Inhib_7_PVALB |
| 23 | BTBD11 | 2.85E-162 | 3.2550166 | Inhib_7_PVALB |
| 24 | CNTNAP2 | 1.75E-160 | 1.6204723 | Inhib_7_PVALB |
| 25 | SV2A | 2.18E-159 | 1.997654 | Inhib_7_PVALB |
| 26 | KCNAB1 | 9.88E-159 | 2.0488887 | Inhib_7_PVALB |
| 27 | GABRB2 | 1.33E-156 | 1.6563268 | Inhib_7_PVALB |
| 28 | TENM1 | 8.63E-152 | 2.457423 | Inhib_7_PVALB |
| 29 | DNER | 3.81E-148 | 2.103577 | Inhib_7_PVALB |
| 30 | MYO16 | 1.39E-145 | 2.3033376 | Inhib_7_PVALB |
| 31 | SNCG | 8.14E-145 | 2.346671 | Inhib_7_PVALB |
| 32 | ATP1A3 | 2.19E-144 | 1.9963828 | Inhib_7_PVALB |
| 33 | PVALB | 8.88E-144 | 4.904617 | Inhib_7_PVALB |
| 34 | TENM3 | 2.62E-142 | 1.9176968 | Inhib_7_PVALB |
| 35 | LANCL1 | 2.40E-139 | 2.0227785 | Inhib_7_PVALB |
| 36 | RBMS3 | 7.96E-138 | 2.1341732 | Inhib_7_PVALB |
| 37 | GRIP1 | 2.00E-132 | 1.770069 | Inhib_7_PVALB |
| 38 | GABRA1 | 4.53E-129 | 1.6514144 | Inhib_7_PVALB |
| 39 | GAPDH | 5.56E-128 | 1.5993768 | Inhib_7_PVALB |
| 40 | SERPINI1 | 2.47E-125 | 1.6352237 | Inhib_7_PVALB |
| 41 | KLF12 | 3.81E-125 | 1.7885991 | Inhib_7_PVALB |
| 42 | VWC2 | 8.07E-125 | 2.2811785 | Inhib_7_PVALB |
| 43 | SLC38A1 | 3.50E-123 | 1.9172189 | Inhib_7_PVALB |
| 44 | NDRG4 | 1.19E-116 | 1.502184 | Inhib_7_PVALB |
| 45 | TIMP2 | 2.52E-116 | 2.1648877 | Inhib_7_PVALB |
| 46 | TMEM132C | 5.50E-116 | 2.3661087 | Inhib_7_PVALB |
| 47 | ZNF536 | 1.62E-115 | 1.7885427 | Inhib_7_PVALB |
| 48 | CNTNAP3 | 8.53E-112 | 2.7566752 | Inhib_7_PVALB |
| 49 | UQCRH | 1.13E-111 | 1.6652464 | Inhib_7_PVALB |
| 50 | DLGAP1 | 1.38E-110 | 1.279312 | Inhib_7_PVALB |
| 51 | CYCS | 6.50E-109 | 1.937311 | Inhib_7_PVALB |
| 52 | PKM | 1.63E-106 | 1.4220197 | Inhib_7_PVALB |
| 53 | RGS5 | 1.86E-106 | 4.024554 | Inhib_7_PVALB |
| 54 | MEF2C | 3.67E-106 | 1.2688504 | Inhib_7_PVALB |
| 55 | DYNLL2 | 8.65E-106 | 1.6041006 | Inhib_7_PVALB |
| 56 | KCNIP2 | 3.24E-105 | 2.3332624 | Inhib_7_PVALB |

| | | | | |
|---|---|---|---|---|
| 57 | GNAS | 5.04E-105 | 1.4983709 | Inhib_7_PVALB |
| 58 | PPARGC1A | 5.78E-105 | 2.095334 | Inhib_7_PVALB |
| 59 | SNCB | 7.90E-105 | 1.4886763 | Inhib_7_PVALB |
| 60 | PRKG1 | 1.46E-104 | 1.5184981 | Inhib_7_PVALB |
| 61 | APLP2 | 3.87E-104 | 1.4933693 | Inhib_7_PVALB |
| 62 | CADPS | 8.46E-104 | 1.307106 | Inhib_7_PVALB |
| 63 | ARL4C | 8.43E-103 | 2.0678868 | Inhib_7_PVALB |
| 64 | CEND1 | 1.45E-102 | 1.5490123 | Inhib_7_PVALB |
| 65 | RP11-444D3.1 | 1.86E-102 | 1.7645907 | Inhib_7_PVALB |
| 66 | SDK1 | 1.64E-101 | 1.6852566 | Inhib_7_PVALB |
| 67 | KCNC1 | 2.53E-100 | 2.3986764 | Inhib_7_PVALB |
| 68 | TIAM1 | 3.14E-100 | 1.6263211 | Inhib_7_PVALB |
| 69 | ANK1 | 2.83E-99 | 2.7727993 | Inhib_7_PVALB |
| 70 | PNMA2 | 2.39E-98 | 1.4300332 | Inhib_7_PVALB |
| 71 | KLHL5 | 1.49E-97 | 2.191208 | Inhib_7_PVALB |
| 72 | PGAM1 | 1.12E-96 | 1.6089427 | Inhib_7_PVALB |
| 73 | VSTM2A | 1.40E-96 | 1.9247863 | Inhib_7_PVALB |
| 74 | PPIP5K2 | 3.30E-96 | 1.9590169 | Inhib_7_PVALB |
| 75 | KCND2 | 1.28E-95 | 1.3053697 | Inhib_7_PVALB |
| 76 | SPOCK2 | 1.15E-94 | 1.6125095 | Inhib_7_PVALB |
| 77 | DIRAS1 | 8.33E-93 | 1.8806024 | Inhib_7_PVALB |
| 78 | MAP1A | 5.03E-92 | 1.4024482 | Inhib_7_PVALB |
| 79 | HSP90AB1 | 6.17E-92 | 1.2654693 | Inhib_7_PVALB |
| 80 | GNAL | 1.08E-91 | 1.9300131 | Inhib_7_PVALB |
| 81 | RAB3A | 1.30E-90 | 1.5494945 | Inhib_7_PVALB |
| 82 | PLCXD3 | 7.92E-90 | 2.426432 | Inhib_7_PVALB |
| 83 | LBH | 1.82E-89 | 3.8617475 | Inhib_7_PVALB |
| 84 | ANO4 | 1.49E-88 | 1.4365655 | Inhib_7_PVALB |
| 85 | HCN1 | 1.53E-88 | 1.2526726 | Inhib_7_PVALB |
| 86 | MAGI1 | 6.87E-88 | 1.1728733 | Inhib_7_PVALB |
| 87 | CHCHD10 | 1.37E-87 | 1.5677965 | Inhib_7_PVALB |
| 88 | PEBP1 | 5.46E-87 | 1.2351463 | Inhib_7_PVALB |
| 89 | SAT1 | 5.48E-87 | 1.7299758 | Inhib_7_PVALB |
| 90 | LHX6 | 1.49E-86 | 3.7105725 | Inhib_7_PVALB |
| 91 | GPX4 | 1.74E-86 | 1.449443 | Inhib_7_PVALB |
| 92 | NPPC | 9.27E-85 | 4.5034533 | Inhib_7_PVALB |
| 93 | NGFRAP1 | 2.18E-84 | 1.1850029 | Inhib_7_PVALB |
| 94 | COX7C | 2.89E-84 | 1.457237 | Inhib_7_PVALB |
| 95 | RAB3C | 3.58E-84 | 1.2759295 | Inhib_7_PVALB |
| 96 | PCSK1N | 5.90E-84 | 1.2598922 | Inhib_7_PVALB |
| 97 | ASTN2 | 1.18E-83 | 1.2270931 | Inhib_7_PVALB |
| 98 | SLC6A17 | 6.15E-83 | 1.5337771 | Inhib_7_PVALB |
| 99 | NDUFA4 | 6.73E-83 | 1.4291042 | Inhib_7_PVALB |
| 0 | CNTN5 | 5.15E-187 | 4.4060535 | Inhib_8_PVALB |
| 1 | ZNF385D | 7.04E-177 | 4.483847 | Inhib_8_PVALB |
| 2 | DPP10 | 2.09E-166 | 3.4105628 | Inhib_8_PVALB |
| 3 | THSD7A | 8.37E-157 | 4.192926 | Inhib_8_PVALB |
| 4 | SDK1 | 2.23E-138 | 3.4289348 | Inhib_8_PVALB |
| 5 | RORA | 7.31E-130 | 2.6097536 | Inhib_8_PVALB |
| 6 | ERBB4 | 8.21E-127 | 3.5254931 | Inhib_8_PVALB |
| 7 | TRPS1 | 2.60E-113 | 2.9653885 | Inhib_8_PVALB |
| 8 | FSTL5 | 2.93E-109 | 2.9340568 | Inhib_8_PVALB |
| 9 | LHFPL3 | 5.42E-107 | 3.1104295 | Inhib_8_PVALB |

| | | | | |
|---|---|---|---|---|
| 10 | TENM1 | 3.02E-101 | 3.1901793 | Inhib_8_PVALB |
| 11 | CNTNAP3B | 5.98E-99 | 3.2400064 | Inhib_8_PVALB |
| 12 | CA8 | 1.28E-98 | 5.2963963 | Inhib_8_PVALB |
| 13 | ALK | 1.97E-95 | 3.534646 | Inhib_8_PVALB |
| 14 | CALN1 | 4.27E-95 | 2.8676574 | Inhib_8_PVALB |
| 15 | GRIA4 | 1.03E-89 | 2.029927 | Inhib_8_PVALB |
| 16 | MDGA2 | 1.24E-89 | 1.8645586 | Inhib_8_PVALB |
| 17 | ZNF804A | 1.27E-89 | 2.7228026 | Inhib_8_PVALB |
| 18 | GABRG3 | 5.55E-89 | 2.2063084 | Inhib_8_PVALB |
| 19 | MYO16 | 1.60E-88 | 2.916227 | Inhib_8_PVALB |
| 20 | TMEM132D | 6.23E-87 | 2.1113923 | Inhib_8_PVALB |
| 21 | PVALB | 9.27E-87 | 4.969492 | Inhib_8_PVALB |
| 22 | FRMPD4 | 9.92E-87 | 1.9263284 | Inhib_8_PVALB |
| 23 | ZNF536 | 4.63E-86 | 2.8300958 | Inhib_8_PVALB |
| 24 | DNER | 1.34E-85 | 2.5334651 | Inhib_8_PVALB |
| 25 | SLC9A9 | 2.37E-84 | 3.245906 | Inhib_8_PVALB |
| 26 | TIAM1 | 4.48E-84 | 2.4596095 | Inhib_8_PVALB |
| 27 | LRRC4C | 1.01E-82 | 1.8918973 | Inhib_8_PVALB |
| 28 | RP11-123O10.4 | 1.07E-80 | 2.455 | Inhib_8_PVALB |
| 29 | GRIA1 | 4.24E-80 | 2.0601575 | Inhib_8_PVALB |
| 30 | GULP1 | 5.79E-80 | 2.868626 | Inhib_8_PVALB |
| 31 | FAM19A2 | 2.52E-77 | 2.0114636 | Inhib_8_PVALB |
| 32 | KCNAB1 | 2.52E-77 | 2.362693 | Inhib_8_PVALB |
| 33 | CRH | 5.82E-77 | 4.3103285 | Inhib_8_PVALB |
| 34 | PLD5 | 1.04E-75 | 2.7269926 | Inhib_8_PVALB |
| 35 | PLCXD3 | 1.88E-75 | 3.3103297 | Inhib_8_PVALB |
| 36 | SLC6A1 | 8.61E-73 | 2.6960344 | Inhib_8_PVALB |
| 37 | EDIL3 | 5.38E-70 | 1.9154423 | Inhib_8_PVALB |
| 38 | SGCD | 6.28E-69 | 2.0241823 | Inhib_8_PVALB |
| 39 | TMTC1 | 8.39E-69 | 2.4460797 | Inhib_8_PVALB |
| 40 | ZNF385D-AS2 | 2.99E-68 | 5.4673505 | Inhib_8_PVALB |
| 41 | NACC2 | 1.18E-65 | 3.6384645 | Inhib_8_PVALB |
| 42 | CNTNAP2 | 2.20E-65 | 1.5650922 | Inhib_8_PVALB |
| 43 | TMEM132C | 2.70E-64 | 2.8111415 | Inhib_8_PVALB |
| 44 | DAB1 | 1.29E-61 | 1.7564132 | Inhib_8_PVALB |
| 45 | RBMS3 | 1.91E-61 | 2.2933793 | Inhib_8_PVALB |
| 46 | KCNC2 | 4.20E-61 | 1.6580474 | Inhib_8_PVALB |
| 47 | DLGAP1 | 2.06E-60 | 1.4057977 | Inhib_8_PVALB |
| 48 | ADGRB3 | 1.04E-58 | 1.2784542 | Inhib_8_PVALB |
| 49 | ASTN2 | 6.15E-58 | 1.6125734 | Inhib_8_PVALB |
| 50 | ADCY8 | 7.65E-58 | 2.9791098 | Inhib_8_PVALB |
| 51 | MEF2C | 2.26E-57 | 1.3713115 | Inhib_8_PVALB |
| 52 | NPNT | 3.17E-57 | 5.812977 | Inhib_8_PVALB |
| 53 | HPSE2 | 4.75E-55 | 3.054742 | Inhib_8_PVALB |
| 54 | TOX | 7.91E-54 | 2.101588 | Inhib_8_PVALB |
| 55 | DLX6-AS1 | 1.43E-52 | 2.5830424 | Inhib_8_PVALB |
| 56 | OSBPL6 | 9.30E-52 | 1.6577486 | Inhib_8_PVALB |
| 57 | CNTNAP3 | 1.32E-51 | 3.089563 | Inhib_8_PVALB |
| 58 | UNC5C | 1.36E-51 | 1.6537807 | Inhib_8_PVALB |
| 59 | ANK1 | 1.45E-51 | 3.2670631 | Inhib_8_PVALB |
| 60 | CNTNAP5 | 4.69E-51 | 1.6060271 | Inhib_8_PVALB |
| 61 | FUT9 | 6.39E-51 | 1.5502093 | Inhib_8_PVALB |
| 62 | VWC2 | 2.09E-50 | 2.4057162 | Inhib_8_PVALB |

| | | | | |
|---|---|---|---|---|
| 63 | MALAT1 | 9.73E-50 | 1.2460681 | Inhib_8_PVALB |
| 64 | AUTS2 | 2.20E-48 | 1.2993836 | Inhib_8_PVALB |
| 65 | SLC6A1-AS1 | 7.04E-48 | 2.3191402 | Inhib_8_PVALB |
| 66 | SLC4A10 | 1.82E-47 | 1.3492833 | Inhib_8_PVALB |
| 67 | COL21A1 | 1.90E-47 | 3.4987185 | Inhib_8_PVALB |
| 68 | CDH12 | 3.29E-47 | 1.5779945 | Inhib_8_PVALB |
| 69 | GABRD | 6.14E-46 | 2.2062042 | Inhib_8_PVALB |
| 70 | RASSF8 | 4.57E-45 | 3.1220655 | Inhib_8_PVALB |
| 71 | SPARCL1 | 4.73E-45 | 1.4555967 | Inhib_8_PVALB |
| 72 | TENM4 | 5.36E-43 | 1.7558355 | Inhib_8_PVALB |
| 73 | SOX6 | 2.95E-42 | 1.8543651 | Inhib_8_PVALB |
| 74 | FAM110B | 2.95E-42 | 2.2118936 | Inhib_8_PVALB |
| 75 | SLITRK5 | 3.65E-42 | 2.0479481 | Inhib_8_PVALB |
| 76 | GRIP1 | 5.32E-42 | 1.8173187 | Inhib_8_PVALB |
| 77 | ATP1B1 | 8.26E-42 | 1.3668827 | Inhib_8_PVALB |
| 78 | HCN1 | 7.60E-41 | 1.3422756 | Inhib_8_PVALB |
| 79 | MIR4500HG | 8.67E-41 | 2.6933136 | Inhib_8_PVALB |
| 80 | VSTM2A | 2.66E-40 | 2.1195118 | Inhib_8_PVALB |
| 81 | KIAA1456 | 2.66E-40 | 2.1576633 | Inhib_8_PVALB |
| 82 | RASGRF2 | 4.27E-40 | 1.5983638 | Inhib_8_PVALB |
| 83 | NFIB | 1.85E-39 | 1.4385189 | Inhib_8_PVALB |
| 84 | AC074363.1 | 1.92E-39 | 1.8136439 | Inhib_8_PVALB |
| 85 | UNC5D | 2.42E-39 | 1.3580625 | Inhib_8_PVALB |
| 86 | GRID2 | 5.76E-39 | 1.3493006 | Inhib_8_PVALB |
| 87 | AC010127.3 | 9.01E-39 | 1.8842868 | Inhib_8_PVALB |
| 88 | OSBPL3 | 1.85E-37 | 1.8221262 | Inhib_8_PVALB |
| 89 | NBEA | 3.44E-37 | 1.1444776 | Inhib_8_PVALB |
| 90 | ZNF804B | 4.25E-37 | 1.5512722 | Inhib_8_PVALB |
| 91 | FAM19A4 | 7.46E-37 | 4.143803 | Inhib_8_PVALB |
| 92 | SLC24A3 | 1.67E-36 | 2.2115314 | Inhib_8_PVALB |
| 93 | CSMD3 | 1.03E-35 | 1.1788093 | Inhib_8_PVALB |
| 94 | SRGAP1 | 1.14E-35 | 2.8741767 | Inhib_8_PVALB |
| 95 | SCG2 | 1.93E-35 | 2.1586568 | Inhib_8_PVALB |
| 96 | FGF13 | 3.09E-35 | 1.9021231 | Inhib_8_PVALB |
| 97 | FGF12 | 4.77E-35 | 1.1493858 | Inhib_8_PVALB |
| 98 | CADPS | 7.19E-35 | 1.1557513 | Inhib_8_PVALB |
| 99 | MAST4 | 9.17E-34 | 1.2386358 | Inhib_8_PVALB |
| 0 | PLXDC2 | 8.23E-211 | 4.7061124 | Micro_Macro |
| 1 | C10orf11 | 1.05E-140 | 5.344235 | Micro_Macro |
| 2 | DOCK4 | 6.81E-123 | 3.047695 | Micro_Macro |
| 3 | ARHGAP24 | 1.82E-118 | 5.358873 | Micro_Macro |
| 4 | FRMD4A | 9.41E-102 | 1.8049898 | Micro_Macro |
| 5 | SFMBT2 | 1.71E-72 | 4.1375313 | Micro_Macro |
| 6 | MEF2C | 6.09E-72 | 1.4151944 | Micro_Macro |
| 7 | ITPR2 | 3.55E-63 | 3.2837956 | Micro_Macro |
| 8 | DOCK8 | 2.58E-61 | 6.3986826 | Micro_Macro |
| 9 | SRGAP2 | 4.48E-59 | 3.6633496 | Micro_Macro |
| 10 | RP11-624C23.1 | 2.07E-56 | 3.745069 | Micro_Macro |
| 11 | MEF2A | 1.03E-55 | 2.7234755 | Micro_Macro |
| 12 | APBB1IP | 1.81E-49 | 6.7682004 | Micro_Macro |
| 13 | ST6GAL1 | 1.33E-48 | 4.2642236 | Micro_Macro |
| 14 | LPAR6 | 6.71E-39 | 5.946847 | Micro_Macro |
| 15 | ANKRD44 | 5.35E-35 | 3.1250565 | Micro_Macro |

| | | | | |
|---|---|---|---|---|
| 16 | CSF1R | 9.07E-34 | 6.730591 | Micro_Macro |
| 17 | ADAM28 | 9.04E-33 | 6.433434 | Micro_Macro |
| 18 | MAML2 | 8.66E-32 | 3.011663 | Micro_Macro |
| 19 | SRGAP2B | 5.73E-31 | 3.3762343 | Micro_Macro |
| 20 | P2RY12 | 1.54E-30 | 5.8425655 | Micro_Macro |
| 21 | ST6GALNAC3 | 7.07E-29 | 3.156423 | Micro_Macro |
| 22 | FYB | 2.47E-27 | 4.8008614 | Micro_Macro |
| 23 | HS3ST4 | 4.09E-24 | 2.3509953 | Micro_Macro |
| 24 | ZFP36L1 | 2.19E-23 | 4.2377834 | Micro_Macro |
| 25 | SLC9A9 | 3.62E-21 | 2.862494 | Micro_Macro |
| 26 | TBXAS1 | 2.36E-19 | 5.917872 | Micro_Macro |
| 27 | PTPRC | 6.72E-19 | 7.200221 | Micro_Macro |
| 28 | CHST11 | 1.33E-18 | 2.8717349 | Micro_Macro |
| 29 | TMSB4X | 3.64E-18 | 1.2232293 | Micro_Macro |
| 30 | CD74 | 5.84E-18 | 6.6556935 | Micro_Macro |
| 31 | C3 | 2.02E-17 | 5.83719 | Micro_Macro |
| 32 | ZFHX3 | 4.86E-17 | 3.3407786 | Micro_Macro |
| 33 | FOXN3 | 5.19E-17 | 2.3894198 | Micro_Macro |
| 34 | SLC1A3 | 1.03E-16 | 2.1457565 | Micro_Macro |
| 35 | AOAH | 1.52E-16 | 4.5276265 | Micro_Macro |
| 36 | ATP8B4 | 1.63E-16 | 4.316464 | Micro_Macro |
| 37 | SYNDIG1 | 1.12E-15 | 2.7400641 | Micro_Macro |
| 38 | RP11-480C22.1 | 6.63E-15 | 6.3711805 | Micro_Macro |
| 39 | CSF2RA | 2.23E-14 | 5.763562 | Micro_Macro |
| 40 | SAMSN1 | 8.62E-14 | 6.027854 | Micro_Macro |
| 41 | INPP5D | 2.66E-13 | 5.015127 | Micro_Macro |
| 42 | A2M | 3.89E-13 | 4.915354 | Micro_Macro |
| 43 | NAV3 | 1.18E-12 | 1.5248195 | Micro_Macro |
| 44 | SLCO2B1 | 2.11E-12 | 5.3187375 | Micro_Macro |
| 45 | QKI | 2.21E-12 | 1.2295066 | Micro_Macro |
| 46 | CX3CR1 | 2.42E-12 | 7.839364 | Micro_Macro |
| 47 | PALD1 | 3.63E-12 | 4.880949 | Micro_Macro |
| 48 | MAML3 | 5.11E-12 | 2.952406 | Micro_Macro |
| 49 | NEAT1 | 7.42E-12 | 1.8370541 | Micro_Macro |
| 50 | SRGAP2C | 1.32E-11 | 2.7571335 | Micro_Macro |
| 51 | ELMO1 | 1.43E-11 | 1.7708604 | Micro_Macro |
| 52 | B3GNT5 | 2.36E-11 | 5.4179044 | Micro_Macro |
| 53 | BNC2 | 3.31E-11 | 4.6294518 | Micro_Macro |
| 54 | DIAPH2 | 4.97E-11 | 2.0340438 | Micro_Macro |
| 55 | EPB41L2 | 9.68E-11 | 1.7033662 | Micro_Macro |
| 56 | SLC8A1 | 1.14E-10 | 0.85476536 | Micro_Macro |
| 57 | RHBDF2 | 1.86E-10 | 5.94474 | Micro_Macro |
| 58 | SPP1 | 2.83E-10 | 3.398083 | Micro_Macro |
| 59 | ITGAX | 4.66E-10 | 6.0844254 | Micro_Macro |
| 60 | RP11-556E13.1 | 4.98E-10 | 5.595988 | Micro_Macro |
| 61 | KCNQ3 | 5.52E-10 | 1.4029982 | Micro_Macro |
| 62 | ABCC4 | 9.30E-10 | 3.8551626 | Micro_Macro |
| 63 | MALAT1 | 1.09E-09 | 0.58243763 | Micro_Macro |
| 64 | BMP2K | 1.20E-09 | 2.782011 | Micro_Macro |
| 65 | SYK | 1.92E-09 | 7.0607014 | Micro_Macro |
| 66 | AKAP13 | 2.08E-09 | 2.7062569 | Micro_Macro |
| 67 | MAF | 2.99E-09 | 2.967782 | Micro_Macro |
| 68 | MSR1 | 3.58E-09 | 4.9704456 | Micro_Macro |

| | | | | |
|---|---|---|---|---|
| 69 | LPCAT2 | 3.59E-09 | 4.8105464 | Micro_Macro |
| 70 | ZNF710 | 7.05E-09 | 3.684892 | Micro_Macro |
| 71 | SORL1 | 8.85E-09 | 1.5998609 | Micro_Macro |
| 72 | SAT1 | 1.66E-08 | 2.2305048 | Micro_Macro |
| 73 | LHFPL2 | 1.70E-08 | 3.4451737 | Micro_Macro |
| 74 | CYFIP1 | 6.17E-08 | 3.1172664 | Micro_Macro |
| 75 | LDLRAD4 | 8.66E-08 | 1.6287266 | Micro_Macro |
| 76 | LAPTM5 | 1.77E-07 | 6.307696 | Micro_Macro |
| 77 | OLR1 | 1.85E-07 | 5.567798 | Micro_Macro |
| 78 | DISC1 | 2.67E-07 | 3.4472277 | Micro_Macro |
| 79 | CPED1 | 3.24E-07 | 4.1037006 | Micro_Macro |
| 80 | ARHGAP22 | 4.42E-07 | 2.779676 | Micro_Macro |
| 81 | TGFBR2 | 1.80E-06 | 5.314548 | Micro_Macro |
| 82 | PICALM | 2.30E-06 | 1.6956265 | Micro_Macro |
| 83 | CSF3R | 2.99E-06 | 6.693391 | Micro_Macro |
| 84 | BLNK | 1.00E-05 | 5.2382426 | Micro_Macro |
| 85 | TGFBR1 | 1.09E-05 | 3.8020904 | Micro_Macro |
| 86 | TYROBP | 1.26E-05 | 7.1175404 | Micro_Macro |
| 87 | LST1 | 1.37E-05 | 7.050578 | Micro_Macro |
| 88 | RUNX1 | 1.58E-05 | 2.5164611 | Micro_Macro |
| 89 | C10orf54 | 1.59E-05 | 4.3345356 | Micro_Macro |
| 90 | FGD4 | 1.91E-05 | 2.5907738 | Micro_Macro |
| 91 | CD86 | 2.03E-05 | 7.003238 | Micro_Macro |
| 92 | IFNGR1 | 3.61E-05 | 3.047708 | Micro_Macro |
| 93 | SLC4A7 | 4.58E-05 | 2.2948072 | Micro_Macro |
| 94 | FLI1 | 5.27E-05 | 5.5884247 | Micro_Macro |
| 95 | FCHSD2 | 5.57E-05 | 1.6851604 | Micro_Macro |
| 96 | C1QB | 5.76E-05 | 6.4191623 | Micro_Macro |
| 97 | FMNL3 | 5.92E-05 | 3.4032848 | Micro_Macro |
| 98 | RP11-13N12.1 | 6.64E-05 | 5.1987476 | Micro_Macro |
| 99 | MYO1F | 7.02E-05 | 4.42274 | Micro_Macro |
| 0 | FTH1 | 8.12E-71 | 3.7958026 | Mix_1 |
| 1 | CALM1 | 8.12E-71 | 3.11475 | Mix_1 |
| 2 | FTL | 6.41E-69 | 4.4130583 | Mix_1 |
| 3 | MAP1A | 4.78E-65 | 3.748428 | Mix_1 |
| 4 | CAMK2N1 | 5.28E-65 | 3.2618036 | Mix_1 |
| 5 | MAP2 | 7.65E-65 | 2.6290805 | Mix_1 |
| 6 | PCSK1N | 4.28E-64 | 3.4849808 | Mix_1 |
| 7 | DHFR | 1.49E-63 | 4.9690022 | Mix_1 |
| 8 | IDS | 1.91E-63 | 2.8963356 | Mix_1 |
| 9 | TUBA1B | 6.56E-63 | 3.148321 | Mix_1 |
| 10 | RPS15 | 7.88E-60 | 3.3216078 | Mix_1 |
| 11 | SEPW1 | 2.25E-59 | 2.8854716 | Mix_1 |
| 12 | MTRNR2L8 | 5.34E-58 | 5.5272946 | Mix_1 |
| 13 | TMSB4X | 5.34E-58 | 2.7861352 | Mix_1 |
| 14 | CST3 | 5.34E-58 | 4.4823666 | Mix_1 |
| 15 | MBP | 1.46E-56 | 3.5654218 | Mix_1 |
| 16 | GAPDH | 1.06E-55 | 2.7373757 | Mix_1 |
| 17 | NDUFA4 | 1.77E-55 | 2.9511182 | Mix_1 |
| 18 | GUK1 | 2.34E-55 | 2.904147 | Mix_1 |
| 19 | NDRG2 | 7.59E-52 | 3.8849912 | Mix_1 |
| 20 | RPL15 | 8.02E-52 | 3.0888548 | Mix_1 |
| 21 | LINC00657 | 1.03E-51 | 2.5234818 | Mix_1 |

| | | | | |
|---|---|---|---|---|
| 22 | CLU | 4.69E-50 | 2.933711 | Mix_1 |
| 23 | GFAP | 5.85E-50 | 4.942676 | Mix_1 |
| 24 | COX7C | 1.89E-49 | 3.0251365 | Mix_1 |
| 25 | TMSB10 | 3.47E-49 | 2.8580747 | Mix_1 |
| 26 | FAM107A | 3.76E-49 | 4.980157 | Mix_1 |
| 27 | EEF1A1 | 2.03E-48 | 3.365714 | Mix_1 |
| 28 | RPL32 | 2.68E-48 | 3.1641755 | Mix_1 |
| 29 | SLC1A2 | 3.11E-48 | 3.3695595 | Mix_1 |
| 30 | CALM3 | 1.07E-47 | 2.4539764 | Mix_1 |
| 31 | CCNI | 4.18E-47 | 2.5773585 | Mix_1 |
| 32 | MT2A | 6.27E-47 | 4.2699456 | Mix_1 |
| 33 | AC105402.4 | 6.30E-47 | 4.7291703 | Mix_1 |
| 34 | EEF2 | 6.40E-47 | 3.2331638 | Mix_1 |
| 35 | RPL41 | 9.04E-47 | 3.176264 | Mix_1 |
| 36 | CHN1 | 9.19E-47 | 2.2222679 | Mix_1 |
| 37 | PPM1G | 1.76E-46 | 3.0489411 | Mix_1 |
| 38 | MT3 | 2.37E-46 | 3.4540122 | Mix_1 |
| 39 | MTURN | 1.73E-45 | 2.9971192 | Mix_1 |
| 40 | APOE | 1.77E-45 | 4.081544 | Mix_1 |
| 41 | RPS14 | 1.80E-45 | 3.0142841 | Mix_1 |
| 42 | NRGN | 2.80E-45 | 2.4231386 | Mix_1 |
| 43 | RPL36 | 2.96E-45 | 3.2246144 | Mix_1 |
| 44 | RPL34 | 1.71E-44 | 3.178083 | Mix_1 |
| 45 | RPLP1 | 1.97E-44 | 3.291884 | Mix_1 |
| 46 | ACTB | 2.26E-44 | 2.2009873 | Mix_1 |
| 47 | PDZD4 | 4.37E-44 | 3.270406 | Mix_1 |
| 48 | APOO | 4.45E-44 | 2.9996848 | Mix_1 |
| 49 | RPL13 | 5.67E-44 | 2.9763134 | Mix_1 |
| 50 | SERF2 | 4.24E-43 | 2.932396 | Mix_1 |
| 51 | CKB | 7.99E-43 | 2.1612 | Mix_1 |
| 52 | ARPP19 | 1.40E-42 | 2.3141675 | Mix_1 |
| 53 | RPL38 | 1.72E-42 | 3.2253137 | Mix_1 |
| 54 | RPS19 | 3.20E-42 | 3.3494706 | Mix_1 |
| 55 | GLUL | 8.09E-42 | 3.506268 | Mix_1 |
| 56 | AES | 1.49E-40 | 2.8620174 | Mix_1 |
| 57 | SMAD2 | 2.13E-40 | 3.2502334 | Mix_1 |
| 58 | COX5B | 2.53E-40 | 3.1215298 | Mix_1 |
| 59 | RPL13A | 2.58E-40 | 3.326487 | Mix_1 |
| 60 | HSP90AA1 | 1.77E-39 | 2.2189052 | Mix_1 |
| 61 | PEBP1 | 2.09E-39 | 2.257921 | Mix_1 |
| 62 | COX4I1 | 2.15E-39 | 2.6202 | Mix_1 |
| 63 | KIF5A | 7.30E-39 | 2.5686605 | Mix_1 |
| 64 | DPYSL2 | 3.21E-38 | 1.9306968 | Mix_1 |
| 65 | RPL28 | 8.43E-38 | 2.8906574 | Mix_1 |
| 66 | GAP43 | 9.60E-38 | 2.4214656 | Mix_1 |
| 67 | PTMA | 4.10E-37 | 2.2159693 | Mix_1 |
| 68 | MAP1B | 5.59E-37 | 1.9112946 | Mix_1 |
| 69 | OOEP | 6.09E-37 | 3.5901845 | Mix_1 |
| 70 | CAMK2A | 6.20E-37 | 1.977763 | Mix_1 |
| 71 | CCDC85B | 8.95E-37 | 2.7046926 | Mix_1 |
| 72 | PMP2 | 9.68E-37 | 4.9146543 | Mix_1 |
| 73 | ALDOA | 1.25E-36 | 2.3262653 | Mix_1 |
| 74 | RPL8 | 3.27E-36 | 3.0618544 | Mix_1 |

| | | | | |
|---|---|---|---|---|
| 75 | GPM6B | 5.79E-36 | 1.9448076 | Mix_1 |
| 76 | C9orf16 | 8.32E-36 | 2.8158076 | Mix_1 |
| 77 | SNAP25 | 8.70E-36 | 1.6999443 | Mix_1 |
| 78 | PEA15 | 1.54E-35 | 3.1699347 | Mix_1 |
| 79 | FBXL16 | 4.84E-35 | 3.0882711 | Mix_1 |
| 80 | RPSA | 1.65E-34 | 3.336155 | Mix_1 |
| 81 | TXNRD1 | 9.18E-34 | 2.7804236 | Mix_1 |
| 82 | C14orf2 | 5.17E-33 | 2.4101186 | Mix_1 |
| 83 | RPL35 | 1.37E-32 | 2.7827423 | Mix_1 |
| 84 | RPS18 | 1.71E-32 | 3.2668052 | Mix_1 |
| 85 | BAIAP2L1 | 2.44E-32 | 3.5585673 | Mix_1 |
| 86 | RPL21 | 2.84E-32 | 3.135488 | Mix_1 |
| 87 | MORF4L1 | 3.09E-32 | 1.9833788 | Mix_1 |
| 88 | RPS27A | 4.48E-32 | 2.8537965 | Mix_1 |
| 89 | UBB | 5.04E-32 | 2.5982952 | Mix_1 |
| 90 | YWHAH | 5.05E-32 | 1.9440842 | Mix_1 |
| 91 | RPS25 | 6.34E-32 | 2.66301 | Mix_1 |
| 92 | CFL1 | 1.82E-31 | 2.0858047 | Mix_1 |
| 93 | SOD1 | 8.95E-31 | 2.537868 | Mix_1 |
| 94 | RPS27 | 9.93E-31 | 2.901303 | Mix_1 |
| 95 | RPL37A | 1.39E-30 | 2.6216242 | Mix_1 |
| 96 | COX7A2 | 1.47E-30 | 2.824431 | Mix_1 |
| 97 | RPS6 | 1.54E-30 | 3.3365128 | Mix_1 |
| 98 | CA11 | 1.54E-30 | 2.4252717 | Mix_1 |
| 99 | DYNLL1 | 2.67E-30 | 1.9855559 | Mix_1 |
| 0 | DHFR | 0 | 6.149673 | Mix_2 |
| 1 | IDS | 8.69E-257 | 2.9569304 | Mix_2 |
| 2 | FTH1 | 7.64E-251 | 2.9109185 | Mix_2 |
| 3 | CALM1 | 5.08E-232 | 2.3702366 | Mix_2 |
| 4 | CAMK2N1 | 7.30E-228 | 3.0438457 | Mix_2 |
| 5 | FTL | 2.22E-192 | 3.3871486 | Mix_2 |
| 6 | CST3 | 3.51E-181 | 4.199651 | Mix_2 |
| 7 | MBP | 6.53E-137 | 3.2216566 | Mix_2 |
| 8 | NDRG2 | 7.98E-130 | 3.6341107 | Mix_2 |
| 9 | NRGN | 3.44E-128 | 1.9591798 | Mix_2 |
| 10 | MAP2 | 1.17E-124 | 1.7222304 | Mix_2 |
| 11 | PCSK1N | 1.18E-120 | 2.8955479 | Mix_2 |
| 12 | CAMK2A | 4.81E-120 | 2.2415981 | Mix_2 |
| 13 | MTRNR2L8 | 1.01E-110 | 5.6892776 | Mix_2 |
| 14 | TMSB4X | 2.34E-106 | 1.930648 | Mix_2 |
| 15 | GLUL | 9.93E-84 | 3.110196 | Mix_2 |
| 16 | TUBA1B | 2.24E-83 | 2.0482626 | Mix_2 |
| 17 | AC105402.4 | 4.27E-78 | 4.271215 | Mix_2 |
| 18 | KIF5A | 6.79E-76 | 2.7602963 | Mix_2 |
| 19 | SLC1A2 | 2.42E-70 | 2.5855434 | Mix_2 |
| 20 | RPS15 | 9.49E-67 | 2.5461636 | Mix_2 |
| 21 | CHN1 | 2.91E-65 | 1.1968147 | Mix_2 |
| 22 | MAP1A | 7.87E-65 | 2.7830806 | Mix_2 |
| 23 | PDZD4 | 6.31E-61 | 3.0624998 | Mix_2 |
| 24 | MTRNR2L12 | 4.52E-55 | 5.499267 | Mix_2 |
| 25 | SNAP25 | 7.98E-53 | 0.7965452 | Mix_2 |
| 26 | APOE | 1.01E-51 | 3.2453341 | Mix_2 |
| 27 | GFAP | 2.93E-51 | 4.0755157 | Mix_2 |

| | | | | |
|---|---|---|---|---|
| 28 | FAM107A | 9.38E-48 | 4.343446 | Mix_2 |
| 29 | GAPDH | 2.01E-44 | 1.5984378 | Mix_2 |
| 30 | RPL34 | 4.62E-43 | 2.5563943 | Mix_2 |
| 31 | ACTB | 3.97E-41 | 1.3143252 | Mix_2 |
| 32 | EEF2 | 5.27E-38 | 2.7590303 | Mix_2 |
| 33 | SMAD2 | 7.27E-38 | 2.9546843 | Mix_2 |
| 34 | CLU | 4.09E-36 | 1.8263643 | Mix_2 |
| 35 | LINC00657 | 5.14E-34 | 1.6503532 | Mix_2 |
| 36 | RPL41 | 6.53E-31 | 2.544192 | Mix_2 |
| 37 | SEPW1 | 4.04E-30 | 1.7114081 | Mix_2 |
| 38 | PMP2 | 4.94E-29 | 4.2767744 | Mix_2 |
| 39 | CKB | 1.12E-27 | 1.2885876 | Mix_2 |
| 40 | CALM3 | 2.38E-26 | 1.4039705 | Mix_2 |
| 41 | GUK1 | 2.09E-24 | 2.103509 | Mix_2 |
| 42 | TMSB10 | 9.66E-24 | 1.5548733 | Mix_2 |
| 43 | MTURN | 8.21E-22 | 2.3119202 | Mix_2 |
| 44 | NSMF | 2.80E-21 | 2.264372 | Mix_2 |
| 45 | MT2A | 2.13E-20 | 2.9122906 | Mix_2 |
| 46 | RPL13 | 2.20E-20 | 2.099433 | Mix_2 |
| 47 | PPM1G | 2.94E-18 | 2.2792077 | Mix_2 |
| 48 | RPLP1 | 3.33E-18 | 2.4805057 | Mix_2 |
| 49 | DDN | 4.45E-17 | 3.4888153 | Mix_2 |
| 50 | MT3 | 4.75E-17 | 2.5688236 | Mix_2 |
| 51 | RPS19 | 2.64E-16 | 2.4059434 | Mix_2 |
| 52 | NDUFA4 | 3.29E-16 | 1.8087772 | Mix_2 |
| 53 | ARPP19 | 5.28E-15 | 1.2233884 | Mix_2 |
| 54 | FBXL16 | 1.47E-14 | 2.6570036 | Mix_2 |
| 55 | CCNI | 4.09E-14 | 1.6762263 | Mix_2 |
| 56 | CBX6 | 4.72E-14 | 2.2520826 | Mix_2 |
| 57 | GPM6B | 7.90E-14 | 1.164922 | Mix_2 |
| 58 | EEF1A1 | 1.28E-13 | 2.2655902 | Mix_2 |
| 59 | NCS1 | 1.71E-13 | 2.4459438 | Mix_2 |
| 60 | ENHO | 1.83E-13 | 3.227062 | Mix_2 |
| 61 | GPM6A | 4.23E-13 | 0.40211144 | Mix_2 |
| 62 | RPL32 | 7.11E-13 | 2.0538368 | Mix_2 |
| 63 | RPL21 | 2.81E-12 | 2.478226 | Mix_2 |
| 64 | CCDC85B | 5.06E-12 | 1.9345894 | Mix_2 |
| 65 | S100A1 | 2.59E-11 | 3.785807 | Mix_2 |
| 66 | OOEP | 4.74E-11 | 2.6237783 | Mix_2 |
| 67 | S100B | 6.06E-11 | 2.8329086 | Mix_2 |
| 68 | PALM | 2.66E-10 | 2.4193022 | Mix_2 |
| 69 | RPL28 | 3.25E-10 | 2.1293182 | Mix_2 |
| 70 | RPL15 | 3.43E-10 | 1.8683764 | Mix_2 |
| 71 | SCD | 5.76E-10 | 2.48006 | Mix_2 |
| 72 | RPS18 | 5.78E-10 | 2.4849787 | Mix_2 |
| 73 | COX7C | 2.14E-09 | 1.8184172 | Mix_2 |
| 74 | PAQR6 | 8.69E-09 | 3.15468 | Mix_2 |
| 75 | HIPK2 | 5.67E-08 | 2.358127 | Mix_2 |
| 76 | RPL36 | 2.56E-07 | 2.003872 | Mix_2 |
| 77 | RPS27A | 9.49E-07 | 2.0294049 | Mix_2 |
| 78 | DLG4 | 1.32E-06 | 1.9672914 | Mix_2 |
| 79 | AES | 2.00E-06 | 1.8180078 | Mix_2 |
| 80 | SLC17A7 | 2.06E-06 | 1.1705465 | Mix_2 |

| | | | | |
|---|---|---|---|---|
| 81 | RPL13A | 2.93E-06 | 2.0195522 | Mix_2 |
| 82 | RPS27 | 4.49E-06 | 2.0627196 | Mix_2 |
| 83 | JUND | 5.98E-06 | 1.6604216 | Mix_2 |
| 84 | RPL35 | 8.04E-06 | 1.9976531 | Mix_2 |
| 85 | MT1E | 1.16E-05 | 3.117548 | Mix_2 |
| 86 | PPP1R9B | 1.27E-05 | 2.5170765 | Mix_2 |
| 87 | LINGO1 | 1.53E-05 | 0.26816663 | Mix_2 |
| 88 | PLEKHB1 | 1.60E-05 | 3.1331477 | Mix_2 |
| 89 | RPS14 | 1.99E-05 | 1.89833 | Mix_2 |
| 90 | PTN | 2.02E-05 | 1.9829084 | Mix_2 |
| 91 | RPL38 | 5.23E-05 | 2.1022851 | Mix_2 |
| 92 | SOWAHA | 8.85E-05 | 3.2539303 | Mix_2 |
| 93 | RPS2 | 9.50E-05 | 2.0397608 | Mix_2 |
| 94 | KIF1A | 0.00010628 | 1.3151592 | Mix_2 |
| 95 | SLC1A3 | 0.000116465 | 1.2753994 | Mix_2 |
| 96 | ZNF358 | 0.000198415 | 2.864887 | Mix_2 |
| 97 | RPL8 | 0.000297246 | 1.9679649 | Mix_2 |
| 98 | TUBB2B | 0.000382429 | 2.392484 | Mix_2 |
| 99 | PSD | 0.00052231 | 1.848529 | Mix_2 |
| 0 | TBC1D3P1-DHX40P1 | 3.40E-44 | 5.022703 | Mix_3 |
| 1 | RPL32 | 1.65E-31 | 3.2549648 | Mix_3 |
| 2 | ATP1B1 | 4.39E-17 | 1.7990551 | Mix_3 |
| 3 | SNAP25 | 6.52E-14 | 1.0270523 | Mix_3 |
| 4 | CALM1 | 5.85E-12 | 1.2379546 | Mix_3 |
| 5 | MAP1B | 1.21E-08 | 1.2429075 | Mix_3 |
| 6 | GAPDH | 5.85E-05 | 1.315735 | Mix_3 |
| 7 | PKM | 0.000880095 | 1.593827 | Mix_3 |
| 8 | TAC1 | 0.00172953 | 3.0748034 | Mix_3 |
| 9 | HSP90AA1 | 0.002461348 | 1.3209444 | Mix_3 |
| 10 | SYT1 | 0.012191218 | 0.3264533 | Mix_3 |
| 11 | PVALB | 0.040641106 | 3.2970479 | Mix_3 |
| 12 | GAD1 | 0.044531052 | 1.9870152 | Mix_3 |
| 13 | NDRG4 | 0.105585549 | 1.1671947 | Mix_3 |
| 14 | SPARCL1 | 0.131927018 | 1.2081629 | Mix_3 |
| 15 | CALM3 | 0.141380787 | 1.0323505 | Mix_3 |
| 16 | LINC00657 | 0.153561601 | 1.0571314 | Mix_3 |
| 17 | CKB | 0.21135417 | 0.78994256 | Mix_3 |
| 18 | NGFRAP1 | 0.264917834 | 1.0345154 | Mix_3 |
| 19 | ENO2 | 0.266310633 | 1.070641 | Mix_3 |
| 20 | TMSB4X | 0.267445733 | 0.7736831 | Mix_3 |
| 21 | FAIM2 | 0.328240374 | 1.1320729 | Mix_3 |
| 22 | RGS5 | 0.434786169 | 2.832009 | Mix_3 |
| 23 | SNCG | 0.529471818 | 1.609539 | Mix_3 |
| 24 | HSP90AB1 | 0.584525149 | 0.9661454 | Mix_3 |
| 25 | RTN4 | 0.6724769 | 0.46723166 | Mix_3 |
| 26 | EIF4A2 | 0.700282842 | 0.79962194 | Mix_3 |
| 27 | PEBP1 | 0.715940086 | 0.9235361 | Mix_3 |
| 28 | VSNL1 | 0.729507955 | 0.7032211 | Mix_3 |
| 29 | CPLX1 | 0.768733164 | 1.5420605 | Mix_3 |
| 30 | ACTB | 0.848512257 | 0.591378 | Mix_3 |
| 31 | IDS | 0.89633119 | 0.6662175 | Mix_3 |
| 79 | AC113189.5 | 1 | 2.6301975 | Mix_3 |
| 78 | RGR | 1 | 2.5427496 | Mix_3 |

| | | | | |
|---|---|---|---|---|
| 77 | RP5-1048B16.1 | 1 | 2.6836975 | Mix_3 |
| 76 | SLC6A17 | 1 | 1.2867875 | Mix_3 |
| 75 | OAZ3 | 1 | 3.3270485 | Mix_3 |
| 74 | FNDC5 | 1 | 2.0039072 | Mix_3 |
| 70 | LGI2 | 1 | 2.2183084 | Mix_3 |
| 72 | RASSF1-AS1 | 1 | 3.8752809 | Mix_3 |
| 71 | ARTN | 1 | 3.5340843 | Mix_3 |
| 80 | PRSS56 | 1 | 3.1630645 | Mix_3 |
| 69 | RAB3B | 1 | 1.7598573 | Mix_3 |
| 68 | RP11-329B9.5 | 1 | 3.6357138 | Mix_3 |
| 67 | NAPB | 1 | 1.0402153 | Mix_3 |
| 73 | RP1-45N11.1 | 1 | 3.6834943 | Mix_3 |
| 81 | LINC01018 | 1 | 2.6303332 | Mix_3 |
| 97 | CTD-2240J17.3 | 1 | 6.4220095 | Mix_3 |
| 83 | RP6-74O6.6 | 1 | 2.680227 | Mix_3 |
| 66 | CRACR2B | 1 | 3.6034558 | Mix_3 |
| 96 | LMNB2 | 1 | 2.0016274 | Mix_3 |
| 95 | GGTLC1 | 1 | 6.6995177 | Mix_3 |
| 94 | CTD-2062F14.3 | 1 | 8.731205 | Mix_3 |
| 93 | RP11-205K6.1 | 1 | 9.925631 | Mix_3 |
| 92 | RP11-77K12.9 | 1 | 2.6660266 | Mix_3 |
| 82 | CDT1 | 1 | 3.1149623 | Mix_3 |
| 91 | DSCR9 | 1 | 2.7699566 | Mix_3 |
| 89 | SPIN4 | 1 | 2.3687985 | Mix_3 |
| 88 | CTB-58E17.5 | 1 | 3.0917554 | Mix_3 |
| 87 | RP11-259K5.1 | 1 | 2.497455 | Mix_3 |
| 86 | RNF122 | 1 | 2.7867541 | Mix_3 |
| 85 | DNM1 | 1 | 0.7007892 | Mix_3 |
| 84 | MAGEL2 | 1 | 2.3222368 | Mix_3 |
| 90 | C1QTNF1 | 1 | 3.0230355 | Mix_3 |
| 65 | STMN1 | 1 | 0.65663856 | Mix_3 |
| 49 | FST | 1 | 4.894886 | Mix_3 |
| 63 | SLC32A1 | 1 | 2.0361364 | Mix_3 |
| 32 | NDUFA4 | 1 | 1.2124515 | Mix_3 |
| 33 | LINGO1 | 1 | 0.19378813 | Mix_3 |
| 34 | CHN1 | 1 | 0.3609068 | Mix_3 |
| 35 | UQCRH | 1 | 1.3144541 | Mix_3 |
| 36 | CAMK2N1 | 1 | 0.74211794 | Mix_3 |
| 37 | TPI1 | 1 | 1.3157271 | Mix_3 |
| 38 | RTN3 | 1 | 0.499233 | Mix_3 |
| 39 | PCSK1N | 1 | 1.1854359 | Mix_3 |
| 40 | GABRA1 | 1 | 1.2126026 | Mix_3 |
| 41 | C1QL1 | 1 | 2.3246233 | Mix_3 |
| 42 | MORF4L1 | 1 | 1.1453974 | Mix_3 |
| 43 | HAPLN4 | 1 | 1.8508955 | Mix_3 |
| 44 | NPPC | 1 | 2.439783 | Mix_3 |
| 45 | SNCB | 1 | 1.0372071 | Mix_3 |
| 46 | ALDOA | 1 | 0.92588305 | Mix_3 |
| 47 | SLC6A1 | 1 | 1.1494154 | Mix_3 |
| 48 | GPR35 | 1 | 3.2389953 | Mix_3 |
| 62 | ZNF629 | 1 | 2.011306 | Mix_3 |
| 61 | SSSCA1 | 1 | 2.2909958 | Mix_3 |
| 60 | COX7C | 1 | 1.1046202 | Mix_3 |

| | | | | |
|---|---|---|---|---|
| 59 | CHURC1 | 1 | 2.8137343 | Mix_3 |
| 58 | NCDN | 1 | 1.0579033 | Mix_3 |
| 57 | QPRT | 1 | 2.243455 | Mix_3 |
| 64 | SCGB3A1 | 1 | 4.3953795 | Mix_3 |
| 56 | CRHBP | 1 | 1.9376211 | Mix_3 |
| 54 | MTA2 | 1 | 1.8292756 | Mix_3 |
| 53 | ARX | 1 | 1.918964 | Mix_3 |
| 52 | CEND1 | 1 | 1.1861974 | Mix_3 |
| 51 | GJD2 | 1 | 2.3914335 | Mix_3 |
| 50 | CAMK2N2 | 1 | 1.7188277 | Mix_3 |
| 98 | RP5-930J4.2 | 1 | 6.60588 | Mix_3 |
| 55 | NRGN | 1 | 0.4551886 | Mix_3 |
| 99 | RP11-73E17.2 | 1 | 6.5938673 | Mix_3 |
| 0 | PTGDS | 3.40E-12 | 1.2712408 | Mix_4 |
| 1 | NRGN | 7.62E-12 | 1.082071 | Mix_4 |
| 2 | GRIN1 | 2.15E-11 | 1.0733602 | Mix_4 |
| 3 | MBP | 2.43E-09 | 1.2262135 | Mix_4 |
| 4 | MEG3 | 7.05E-08 | 0.65906525 | Mix_4 |
| 5 | CERCAM | 1.21E-06 | 1.7715064 | Mix_4 |
| 6 | NDRG2 | 3.80E-06 | 1.3115078 | Mix_4 |
| 7 | KIFC2 | 8.23E-05 | 1.3123256 | Mix_4 |
| 8 | MIR219A2 | 0.000320659 | 1.2842468 | Mix_4 |
| 9 | CAMK2A | 0.000379293 | 0.733111 | Mix_4 |
| 10 | SLC17A7 | 0.000479002 | 0.913466 | Mix_4 |
| 11 | GLUL | 0.000794284 | 1.0708772 | Mix_4 |
| 12 | CARNS1 | 0.001363974 | 1.6333971 | Mix_4 |
| 13 | SLC1A2 | 0.002678201 | 0.73664296 | Mix_4 |
| 14 | CST3 | 0.004194765 | 1.0908095 | Mix_4 |
| 15 | GFAP | 0.005089844 | 1.8801996 | Mix_4 |
| 16 | PLP1 | 0.007297734 | 0.5693867 | Mix_4 |
| 17 | ABCA2 | 0.010575791 | 0.9020244 | Mix_4 |
| 18 | CKB | 0.028826741 | 0.5644266 | Mix_4 |
| 19 | TMEM59L | 0.051734101 | 0.7096395 | Mix_4 |
| 20 | NSMF | 0.053733048 | 1.0506672 | Mix_4 |
| 21 | COL5A3 | 0.054058768 | 1.0996611 | Mix_4 |
| 22 | IDS | 0.056623437 | 0.6156104 | Mix_4 |
| 23 | KIAA0930 | 0.057668929 | 0.7655322 | Mix_4 |
| 24 | PKD1 | 0.063975228 | 0.78314435 | Mix_4 |
| 25 | DNM1 | 0.065721117 | 0.61123085 | Mix_4 |
| 26 | CLU | 0.067570146 | 0.6715028 | Mix_4 |
| 27 | PHLDB1 | 0.080581821 | 1.120695 | Mix_4 |
| 28 | ACAP3 | 0.082519829 | 0.8722175 | Mix_4 |
| 29 | CHN1 | 0.096317817 | 0.3731108 | Mix_4 |
| 30 | SH3GLB2 | 0.11619386 | 1.1009277 | Mix_4 |
| 31 | CAMK2N1 | 0.116675094 | 0.54203486 | Mix_4 |
| 32 | CDK5R1 | 0.129542868 | 0.81444526 | Mix_4 |
| 33 | NOXA1 | 0.136342548 | 1.1918976 | Mix_4 |
| 34 | SLC1A3 | 0.137108596 | 0.5815255 | Mix_4 |
| 35 | CABP1 | 0.153710312 | 0.5702845 | Mix_4 |
| 36 | SNAP25 | 0.156302172 | 0.29348937 | Mix_4 |
| 37 | LINC00599 | 0.17162926 | 0.75465155 | Mix_4 |
| 38 | ADGRB2 | 0.181731608 | 0.806644 | Mix_4 |
| 39 | ARPP19 | 0.186623593 | 0.5211241 | Mix_4 |

| | | | | |
|---|---|---|---|---|
| 40 | PLXNB1 | 0.190146038 | 1.0913649 | Mix_4 |
| 41 | MAPK11 | 0.209724226 | 1.1938976 | Mix_4 |
| 42 | FGFR3 | 0.212515542 | 0.9859723 | Mix_4 |
| 43 | RAPGEF3 | 0.212515542 | 1.4107397 | Mix_4 |
| 44 | PABPN1 | 0.217298835 | 0.8025924 | Mix_4 |
| 45 | ENC1 | 0.242230926 | 0.48421556 | Mix_4 |
| 46 | SRRM2 | 0.270181163 | 0.7604068 | Mix_4 |
| 47 | PHYHIP | 0.270898651 | 0.6581473 | Mix_4 |
| 49 | BASP1 | 0.295997296 | 0.4697567 | Mix_4 |
| 48 | LENG8 | 0.295997296 | 0.83462626 | Mix_4 |
| 50 | PFKL | 0.309866618 | 1.1901952 | Mix_4 |
| 51 | CIRBP | 0.312422853 | 0.7029872 | Mix_4 |
| 52 | MAPK8IP1 | 0.330536178 | 0.9559004 | Mix_4 |
| 53 | MAPK8IP2 | 0.349990856 | 1.1066724 | Mix_4 |
| 54 | GABBR1 | 0.353405908 | 0.92300564 | Mix_4 |
| 55 | CHD5 | 0.386696985 | 0.8259329 | Mix_4 |
| 56 | KIF1A | 0.393984398 | 0.6962977 | Mix_4 |
| 57 | ACADVL | 0.440614431 | 0.8591398 | Mix_4 |
| 58 | CRIP2 | 0.450445623 | 1.0150495 | Mix_4 |
| 59 | SHC2 | 0.46297987 | 1.2034065 | Mix_4 |
| 60 | CALM1 | 0.473271921 | 0.36687076 | Mix_4 |
| 61 | RIMS3 | 0.521557901 | 0.9935703 | Mix_4 |
| 62 | MT2A | 0.549692523 | 1.0225577 | Mix_4 |
| 63 | DMPK | 0.561081838 | 1.4085683 | Mix_4 |
| 64 | ENTPD6 | 0.610492701 | 1.1686388 | Mix_4 |
| 65 | MEIS3 | 0.649182874 | 1.0832101 | Mix_4 |
| 66 | INF2 | 0.656594528 | 1.0247192 | Mix_4 |
| 67 | APOE | 0.664102384 | 0.5373942 | Mix_4 |
| 68 | HAPLN2 | 0.670209359 | 1.1840371 | Mix_4 |
| 69 | MRPL41 | 0.671655799 | 0.77237254 | Mix_4 |
| 70 | NISCH | 0.771900115 | 0.6764712 | Mix_4 |
| 71 | ANKRD24 | 0.779841589 | 1.2466391 | Mix_4 |
| 72 | MT3 | 0.892560417 | 0.96898425 | Mix_4 |
| 73 | PCED1A | 0.894372114 | 1.4281081 | Mix_4 |
| 74 | MVD | 0.932242457 | 1.0048454 | Mix_4 |
| 75 | CORO6 | 0.977474339 | 0.7085243 | Mix_4 |
| 76 | ADAM11 | 0.979496168 | 0.8948901 | Mix_4 |
| 77 | COQ4 | 0.986199825 | 1.1802005 | Mix_4 |
| 97 | GAPDH | 1 | 0.37096423 | Mix_4 |
| 96 | NDUFV1 | 1 | 1.0941454 | Mix_4 |
| 95 | HIP1R | 1 | 0.55241984 | Mix_4 |
| 94 | AKAP8L | 1 | 0.84604865 | Mix_4 |
| 93 | INTS1 | 1 | 0.860856 | Mix_4 |
| 92 | LGI4 | 1 | 0.6883485 | Mix_4 |
| 91 | DPP7 | 1 | 0.9233514 | Mix_4 |
| 90 | PCBP4 | 1 | 0.87139386 | Mix_4 |
| 89 | SREBF1 | 1 | 0.96355414 | Mix_4 |
| 88 | S100B | 1 | 0.82545954 | Mix_4 |
| 98 | MOG | 1 | 1.2050824 | Mix_4 |
| 86 | YWHAH | 1 | 0.4357955 | Mix_4 |
| 85 | GBA | 1 | 1.1568646 | Mix_4 |
| 84 | ZMIZ2 | 1 | 1.1338898 | Mix_4 |
| 83 | STMN1 | 1 | 0.45328128 | Mix_4 |

| | | | | |
|---|---|---|---|---|
| 82 | YJEFN3 | 1 | 1.1152138 | Mix_4 |
| 81 | LINGO1 | 1 | 0.21040979 | Mix_4 |
| 80 | PPFIA4 | 1 | 0.906635 | Mix_4 |
| 79 | FBXW7 | 1 | 0.31714642 | Mix_4 |
| 78 | MICALL2 | 1 | 1.095973 | Mix_4 |
| 87 | RHPN1 | 1 | 0.7863463 | Mix_4 |
| 99 | QDPR | 1 | 0.67658544 | Mix_4 |
| 0 | CNTNAP2 | 2.74E-123 | 1.3063523 | Mix_5 |
| 1 | RBFOX1 | 6.05E-85 | 1.1215373 | Mix_5 |
| 2 | ROBO2 | 8.05E-74 | 1.3818024 | Mix_5 |
| 3 | GRIK1 | 3.82E-69 | 2.2880154 | Mix_5 |
| 4 | DLGAP1 | 3.82E-69 | 0.9723098 | Mix_5 |
| 5 | NRXN3 | 2.02E-67 | 0.90321976 | Mix_5 |
| 6 | ERBB4 | 6.23E-63 | 1.566059 | Mix_5 |
| 7 | NXPH1 | 5.83E-53 | 1.8687403 | Mix_5 |
| 8 | ZNF385D | 1.57E-48 | 1.5781989 | Mix_5 |
| 9 | KIAA1217 | 3.39E-46 | 1.5515431 | Mix_5 |
| 10 | MEG3 | 7.15E-31 | 0.55903673 | Mix_5 |
| 11 | RP11-123O10.4 | 3.28E-20 | 1.2258216 | Mix_5 |
| 12 | LINGO1 | 1.39E-15 | 0.57108784 | Mix_5 |
| 13 | SOX6 | 4.97E-15 | 1.2568437 | Mix_5 |
| 14 | SPOCK3 | 7.36E-15 | 1.0412762 | Mix_5 |
| 15 | DPP10 | 1.09E-14 | 0.6359143 | Mix_5 |
| 16 | TENM2 | 2.09E-13 | 0.5405411 | Mix_5 |
| 17 | ROBO1 | 2.88E-13 | 0.9775348 | Mix_5 |
| 18 | KCNC2 | 4.33E-12 | 0.7468857 | Mix_5 |
| 19 | PTPRM | 6.91E-12 | 1.0772573 | Mix_5 |
| 20 | TCF4 | 9.67E-12 | 0.4786606 | Mix_5 |
| 21 | RASGEF1B | 1.11E-11 | 0.6701458 | Mix_5 |
| 22 | DAB1 | 2.79E-11 | 0.8063957 | Mix_5 |
| 23 | GRIP1 | 4.20E-11 | 1.0938699 | Mix_5 |
| 24 | GRIN1 | 7.40E-11 | 0.6092028 | Mix_5 |
| 25 | ZNF536 | 9.79E-11 | 1.1774199 | Mix_5 |
| 26 | GRIK2 | 1.29E-10 | 0.62267184 | Mix_5 |
| 27 | TIAM1 | 4.39E-10 | 1.3308338 | Mix_5 |
| 28 | TENM3 | 2.11E-09 | 1.0251415 | Mix_5 |
| 29 | NRG3 | 1.29E-08 | 0.30963904 | Mix_5 |
| 30 | KIF26B | 3.51E-08 | 1.6261274 | Mix_5 |
| 31 | RBMS3 | 4.69E-08 | 1.0592128 | Mix_5 |
| 32 | SGCZ | 6.03E-08 | 0.8444071 | Mix_5 |
| 33 | ZNF804A | 1.12E-07 | 0.99111277 | Mix_5 |
| 34 | CNTN5 | 1.30E-07 | 0.60702574 | Mix_5 |
| 35 | FGF14 | 1.50E-07 | 0.30869702 | Mix_5 |
| 36 | FRMD4A | 2.53E-07 | 0.32473952 | Mix_5 |
| 37 | AUTS2 | 6.24E-07 | 0.48532563 | Mix_5 |
| 38 | PCDH15 | 1.54E-06 | 0.7934209 | Mix_5 |
| 39 | FGF12 | 1.62E-05 | 0.27705306 | Mix_5 |
| 40 | OPCML | 2.35E-05 | 0.250586 | Mix_5 |
| 41 | CADPS | 4.00E-05 | 0.46141228 | Mix_5 |
| 42 | RASGRF2 | 4.14E-05 | 0.8380451 | Mix_5 |
| 43 | UPP1 | 4.44E-05 | 1.3674282 | Mix_5 |
| 44 | LINC00599 | 8.23E-05 | 0.7002714 | Mix_5 |
| 45 | ANK1 | 8.59E-05 | 1.6946836 | Mix_5 |

| | | | | |
|---|---|---|---|---|
| 46 | PLCB1 | 0.000171054 | 0.16218884 | Mix_5 |
| 47 | KAZN | 0.00017784 | 0.68276703 | Mix_5 |
| 48 | PCDH11X | 0.000248421 | 0.9498621 | Mix_5 |
| 49 | XKR4 | 0.000255843 | 0.7883904 | Mix_5 |
| 50 | GAD1 | 0.000307158 | 0.57678014 | Mix_5 |
| 51 | ATRNL1 | 0.000477439 | 0.25948727 | Mix_5 |
| 52 | PABPN1 | 0.00048533 | 0.80722374 | Mix_5 |
| 53 | EPHA6 | 0.000590274 | 0.5914398 | Mix_5 |
| 54 | SYNPR | 0.00067892 | 0.75449944 | Mix_5 |
| 55 | FO538757.2 | 0.000841695 | 0.8057233 | Mix_5 |
| 56 | TAC1 | 0.001019271 | 1.5336448 | Mix_5 |
| 57 | BTBD11 | 0.001312269 | 1.338757 | Mix_5 |
| 58 | ASTN2 | 0.001362355 | 0.6012655 | Mix_5 |
| 59 | MAF | 0.001387398 | 1.893686 | Mix_5 |
| 60 | SLC26A3 | 0.001620235 | 0.48545435 | Mix_5 |
| 61 | SHC2 | 0.001730944 | 0.9307578 | Mix_5 |
| 62 | SLC24A3 | 0.001971727 | 1.2106003 | Mix_5 |
| 63 | OXR1 | 0.002539762 | 0.21907134 | Mix_5 |
| 64 | CORO6 | 0.002568703 | 0.8183914 | Mix_5 |
| 65 | CSMD1 | 0.002923165 | 0.25251362 | Mix_5 |
| 66 | THSD7A | 0.003167077 | 0.8485133 | Mix_5 |
| 67 | TENM1 | 0.00347826 | 1.057409 | Mix_5 |
| 68 | PDIA2 | 0.003564909 | 0.9552806 | Mix_5 |
| 69 | DLX6-AS1 | 0.009055765 | 0.633267 | Mix_5 |
| 70 | SDK1 | 0.012476681 | 0.6503032 | Mix_5 |
| 71 | QTRT1 | 0.016070996 | 0.75065 | Mix_5 |
| 72 | GRIP2 | 0.019968218 | 1.4182558 | Mix_5 |
| 73 | VAV2 | 0.023459145 | 1.5408201 | Mix_5 |
| 74 | MRPL41 | 0.025907006 | 0.7587241 | Mix_5 |
| 75 | SNTG1 | 0.026105943 | 0.22273417 | Mix_5 |
| 76 | ADGRB3 | 0.028121649 | 0.11858686 | Mix_5 |
| 77 | PKD1 | 0.029030779 | 0.5073778 | Mix_5 |
| 78 | ALK | 0.030267267 | 1.014765 | Mix_5 |
| 79 | DPP7 | 0.031686923 | 0.78696054 | Mix_5 |
| 80 | KCNT1 | 0.034484202 | 0.78639525 | Mix_5 |
| 81 | RPS6KA2 | 0.036115133 | 0.8384214 | Mix_5 |
| 82 | CCSER1 | 0.03834249 | 0.17137557 | Mix_5 |
| 83 | GRM5 | 0.039744222 | 0.20660733 | Mix_5 |
| 84 | TMEM63B | 0.040801041 | 0.8903964 | Mix_5 |
| 85 | TMEM132C | 0.042533478 | 0.9535083 | Mix_5 |
| 86 | CACNA1A | 0.051062068 | 0.5739654 | Mix_5 |
| 87 | MYO16 | 0.057114381 | 0.59607315 | Mix_5 |
| 88 | ABCA7 | 0.05871519 | 1.000756 | Mix_5 |
| 89 | NHS | 0.060742669 | 0.9990763 | Mix_5 |
| 90 | TLE2 | 0.064150752 | 1.2333777 | Mix_5 |
| 91 | EYS | 0.066724899 | 0.9542935 | Mix_5 |
| 92 | KCNAB1 | 0.071044162 | 0.5875166 | Mix_5 |
| 93 | ADARB2 | 0.086650728 | 0.3781793 | Mix_5 |
| 94 | HNRNPL | 0.091567437 | 1.2178205 | Mix_5 |
| 95 | CKMT1B | 0.092690643 | 0.7308838 | Mix_5 |
| 96 | VARS2 | 0.093058269 | 0.9413767 | Mix_5 |
| 97 | SPTBN4 | 0.114449589 | 0.57699007 | Mix_5 |
| 98 | SRRM3 | 0.11515474 | 0.70312303 | Mix_5 |

| | | | | |
|---|---|---|---|---|
| 99 | HRH3 | 0.131389303 | 1.283216 | Mix_5 |
| 0 | LHFPL3 | 0 | 5.410449 | OPCs_1 |
| 1 | PCDH15 | 0 | 4.4312315 | OPCs_1 |
| 2 | LRRC4C | 2.77E-272 | 2.5962434 | OPCs_1 |
| 3 | PTPRZ1 | 1.40E-257 | 4.5492296 | OPCs_1 |
| 4 | DSCAM | 4.19E-255 | 2.7516625 | OPCs_1 |
| 5 | TNR | 3.54E-143 | 3.2795815 | OPCs_1 |
| 6 | VCAN | 2.87E-138 | 5.0726085 | OPCs_1 |
| 7 | NLGN1 | 2.94E-134 | 1.5675609 | OPCs_1 |
| 8 | LUZP2 | 2.65E-115 | 3.2331429 | OPCs_1 |
| 9 | OPCML | 3.13E-113 | 1.3088955 | OPCs_1 |
| 10 | GRID2 | 3.61E-107 | 1.8758427 | OPCs_1 |
| 11 | LRP1B | 2.18E-105 | 1.2805352 | OPCs_1 |
| 12 | NRXN1 | 7.30E-100 | 1.135571 | OPCs_1 |
| 13 | SOX6 | 9.91E-90 | 2.9221616 | OPCs_1 |
| 14 | MMP16 | 1.85E-86 | 2.1444697 | OPCs_1 |
| 15 | EPN2 | 6.47E-75 | 3.134673 | OPCs_1 |
| 16 | TNK2 | 1.05E-72 | 2.6905313 | OPCs_1 |
| 17 | RP4-668E10.4 | 9.20E-70 | 5.3398843 | OPCs_1 |
| 18 | AGAP1 | 7.88E-67 | 2.133869 | OPCs_1 |
| 19 | MAML2 | 9.67E-67 | 3.0677352 | OPCs_1 |
| 20 | DPP6 | 2.43E-60 | 1.3212583 | OPCs_1 |
| 21 | ADGRL3 | 1.27E-58 | 0.9928698 | OPCs_1 |
| 22 | CHST11 | 1.91E-56 | 3.2492616 | OPCs_1 |
| 23 | LSAMP | 8.65E-56 | 0.76169705 | OPCs_1 |
| 24 | NXPH1 | 2.75E-55 | 2.1829512 | OPCs_1 |
| 25 | LRRTM4 | 1.84E-54 | 1.0289706 | OPCs_1 |
| 26 | XYLT1 | 1.09E-53 | 2.621645 | OPCs_1 |
| 27 | KCND2 | 1.10E-51 | 1.4639189 | OPCs_1 |
| 28 | SEMA5A | 1.94E-51 | 3.4575253 | OPCs_1 |
| 29 | NOVA1 | 9.88E-51 | 1.8034021 | OPCs_1 |
| 30 | ZBTB20 | 1.70E-49 | 1.9524682 | OPCs_1 |
| 31 | NTM | 1.82E-46 | 1.2152386 | OPCs_1 |
| 32 | PCDH9 | 4.44E-44 | 0.7094479 | OPCs_1 |
| 33 | CSMD1 | 2.63E-37 | 0.7846231 | OPCs_1 |
| 34 | QKI | 4.24E-37 | 1.4193712 | OPCs_1 |
| 35 | CA10 | 2.34E-36 | 1.8662804 | OPCs_1 |
| 36 | OLIG1 | 9.94E-35 | 3.6605222 | OPCs_1 |
| 37 | SMOC1 | 9.26E-32 | 4.147999 | OPCs_1 |
| 38 | CSMD3 | 1.91E-30 | 1.0182333 | OPCs_1 |
| 39 | NAV1 | 7.97E-30 | 2.0360198 | OPCs_1 |
| 40 | SCD5 | 3.40E-29 | 1.9073758 | OPCs_1 |
| 41 | ERBB4 | 1.12E-27 | 1.0135311 | OPCs_1 |
| 42 | NPAS3 | 5.89E-27 | 1.1142414 | OPCs_1 |
| 43 | FGF14 | 1.41E-26 | 0.5856649 | OPCs_1 |
| 44 | COL9A1 | 1.90E-26 | 4.6421375 | OPCs_1 |
| 45 | KIF13A | 5.48E-26 | 2.2110412 | OPCs_1 |
| 46 | COL20A1 | 2.08E-24 | 5.3619957 | OPCs_1 |
| 47 | SLC35F1 | 4.58E-24 | 1.5175073 | OPCs_1 |
| 48 | BCAN | 4.79E-24 | 2.7830086 | OPCs_1 |
| 49 | NCKAP5 | 1.40E-22 | 1.9979725 | OPCs_1 |
| 50 | BRINP3 | 1.56E-22 | 1.6731199 | OPCs_1 |
| 51 | DCC | 1.78E-22 | 1.386773 | OPCs_1 |

| | | | | |
|---|---|---|---|---|
| 52 | GRIK1 | 1.93E-22 | 1.4847668 | OPCs_1 |
| 53 | PDGFRA | 8.59E-22 | 4.5391073 | OPCs_1 |
| 54 | HIP1R | 1.62E-20 | 2.5369635 | OPCs_1 |
| 55 | FGF12 | 2.27E-20 | 0.588339 | OPCs_1 |
| 56 | MDGA2 | 2.61E-18 | 0.7192318 | OPCs_1 |
| 57 | KAT2B | 3.97E-18 | 2.4594095 | OPCs_1 |
| 58 | PLPPR1 | 4.49E-18 | 2.6266649 | OPCs_1 |
| 59 | PDE4B | 1.96E-17 | 1.228334 | OPCs_1 |
| 60 | NCAM2 | 2.38E-17 | 0.76837325 | OPCs_1 |
| 61 | NKAIN3 | 1.80E-16 | 1.9368753 | OPCs_1 |
| 62 | C1orf61 | 2.10E-16 | 1.8759209 | OPCs_1 |
| 63 | OPHN1 | 1.58E-15 | 2.0410101 | OPCs_1 |
| 64 | GALNT13 | 2.32E-15 | 1.6001502 | OPCs_1 |
| 65 | RPL13 | 2.39E-15 | 1.6007409 | OPCs_1 |
| 66 | LINC00511 | 7.43E-14 | 2.61296 | OPCs_1 |
| 67 | ZNF462 | 4.68E-13 | 2.3092644 | OPCs_1 |
| 68 | SNTG1 | 1.13E-12 | 0.56606346 | OPCs_1 |
| 69 | FERMT1 | 7.03E-12 | 4.712152 | OPCs_1 |
| 70 | NFIA | 8.11E-12 | 1.3765434 | OPCs_1 |
| 71 | MEGF11 | 2.15E-11 | 3.2306166 | OPCs_1 |
| 72 | SNX22 | 5.63E-11 | 4.2563586 | OPCs_1 |
| 73 | OLIG2 | 2.95E-10 | 3.870467 | OPCs_1 |
| 74 | SGCZ | 4.70E-10 | 0.9775379 | OPCs_1 |
| 75 | RORA | 5.45E-10 | 0.7473117 | OPCs_1 |
| 76 | SDC3 | 3.05E-09 | 1.981412 | OPCs_1 |
| 77 | BCAS1 | 4.54E-09 | 2.2106905 | OPCs_1 |
| 78 | APOD | 4.76E-09 | 2.4981718 | OPCs_1 |
| 79 | ZCCHC24 | 6.09E-09 | 2.9671495 | OPCs_1 |
| 80 | KAZN | 7.39E-09 | 1.0259312 | OPCs_1 |
| 81 | CDH20 | 1.07E-08 | 1.3887727 | OPCs_1 |
| 82 | FHIT | 1.12E-08 | 1.1769118 | OPCs_1 |
| 83 | ALCAM | 1.36E-08 | 1.2846197 | OPCs_1 |
| 84 | CADM2 | 2.44E-08 | 0.22211115 | OPCs_1 |
| 85 | KCNMB2-AS1 | 2.47E-08 | 1.4893551 | OPCs_1 |
| 86 | RP11-436D23.1 | 2.52E-08 | 1.3563803 | OPCs_1 |
| 87 | USP24 | 3.31E-08 | 1.8951839 | OPCs_1 |
| 88 | CSPG4 | 4.97E-08 | 4.978462 | OPCs_1 |
| 89 | LRRN1 | 5.67E-08 | 2.218173 | OPCs_1 |
| 90 | MYT1 | 6.05E-08 | 3.5874515 | OPCs_1 |
| 91 | PTPRG | 1.14E-07 | 0.6430303 | OPCs_1 |
| 92 | FAM155A | 1.31E-07 | 0.27267167 | OPCs_1 |
| 93 | GRM7 | 2.39E-07 | 0.67818636 | OPCs_1 |
| 94 | CCDC50 | 2.45E-07 | 2.5639355 | OPCs_1 |
| 95 | C10orf11 | 3.17E-07 | 1.9040555 | OPCs_1 |
| 96 | DISC1 | 3.37E-07 | 2.9540741 | OPCs_1 |
| 97 | ARHGAP42 | 3.70E-07 | 2.945598 | OPCs_1 |
| 98 | NOVA1-AS1 | 4.60E-07 | 3.526852 | OPCs_1 |
| 99 | WWOX | 4.84E-07 | 0.99699384 | OPCs_1 |
| 0 | LHFPL3 | 5.37E-198 | 5.9833264 | OPCs_2 |
| 1 | PTPRZ1 | 3.32E-195 | 5.4515433 | OPCs_2 |
| 2 | PCDH15 | 1.91E-192 | 5.1017346 | OPCs_2 |
| 3 | LRRC4C | 4.18E-186 | 3.336106 | OPCs_2 |
| 4 | DSCAM | 1.51E-185 | 3.5259595 | OPCs_2 |

| | | | | |
|---|---|---|---|---|
| 5 | TNR | 6.31E-185 | 4.461929 | OPCs_2 |
| 6 | VCAN | 6.18E-170 | 6.091983 | OPCs_2 |
| 7 | LUZP2 | 1.06E-163 | 4.3367004 | OPCs_2 |
| 8 | NLGN1 | 8.08E-155 | 2.4509678 | OPCs_2 |
| 9 | NOVA1 | 1.22E-138 | 2.829038 | OPCs_2 |
| 10 | MMP16 | 1.52E-138 | 2.9566224 | OPCs_2 |
| 11 | SEMA5A | 4.63E-133 | 4.6216245 | OPCs_2 |
| 12 | SOX6 | 1.44E-131 | 3.8777645 | OPCs_2 |
| 13 | SLC35F1 | 1.07E-129 | 2.853516 | OPCs_2 |
| 14 | CHST11 | 3.03E-124 | 4.2238774 | OPCs_2 |
| 15 | GRID2 | 4.48E-123 | 2.61623 | OPCs_2 |
| 16 | EPN2 | 1.02E-122 | 3.8331969 | OPCs_2 |
| 17 | MAML2 | 9.17E-122 | 4.141552 | OPCs_2 |
| 18 | RP4-668E10.4 | 1.78E-121 | 6.396218 | OPCs_2 |
| 19 | AGAP1 | 4.74E-119 | 2.8005066 | OPCs_2 |
| 20 | XYLT1 | 7.66E-117 | 3.488489 | OPCs_2 |
| 21 | LRP1B | 4.63E-116 | 2.0068972 | OPCs_2 |
| 22 | NRXN1 | 2.41E-114 | 1.7391939 | OPCs_2 |
| 23 | SCD5 | 8.31E-114 | 3.0203776 | OPCs_2 |
| 24 | ZBTB20 | 1.61E-111 | 3.0107558 | OPCs_2 |
| 25 | DPP6 | 2.97E-111 | 2.1110811 | OPCs_2 |
| 26 | KCND2 | 7.54E-111 | 2.3147585 | OPCs_2 |
| 27 | OPCML | 3.00E-109 | 1.8610531 | OPCs_2 |
| 28 | KCNMB2-AS1 | 1.62E-107 | 3.101981 | OPCs_2 |
| 29 | GALNT13 | 5.43E-100 | 2.804276 | OPCs_2 |
| 30 | KIF13A | 5.03E-99 | 3.3264594 | OPCs_2 |
| 31 | BRINP3 | 8.37E-99 | 2.736311 | OPCs_2 |
| 32 | OPHN1 | 9.29E-99 | 3.324312 | OPCs_2 |
| 33 | PDE4B | 2.00E-98 | 2.4420116 | OPCs_2 |
| 34 | NTM | 1.24E-96 | 1.9806038 | OPCs_2 |
| 35 | CA10 | 1.62E-94 | 2.7480834 | OPCs_2 |
| 36 | QKI | 2.03E-94 | 2.339685 | OPCs_2 |
| 37 | ADGRL3 | 2.25E-94 | 1.7385811 | OPCs_2 |
| 38 | NXPH1 | 2.84E-93 | 2.9659283 | OPCs_2 |
| 39 | NPAS3 | 2.38E-90 | 2.2074544 | OPCs_2 |
| 40 | DCC | 1.41E-89 | 2.4156296 | OPCs_2 |
| 41 | NRCAM | 8.76E-87 | 1.8335866 | OPCs_2 |
| 42 | LSAMP | 5.25E-85 | 1.4295545 | OPCs_2 |
| 43 | OLIG1 | 1.12E-82 | 4.5945272 | OPCs_2 |
| 44 | PLPPR1 | 7.60E-82 | 3.8654735 | OPCs_2 |
| 45 | LINC00511 | 7.40E-81 | 3.9622884 | OPCs_2 |
| 46 | MDGA2 | 2.02E-80 | 1.7504894 | OPCs_2 |
| 47 | FHIT | 4.31E-77 | 2.3088584 | OPCs_2 |
| 48 | ZSWIM6 | 7.08E-77 | 2.3126884 | OPCs_2 |
| 49 | RORA | 7.49E-76 | 1.7815084 | OPCs_2 |
| 50 | ZFPM2 | 2.06E-75 | 2.2077563 | OPCs_2 |
| 51 | PDZD2 | 3.74E-75 | 2.2936475 | OPCs_2 |
| 52 | KAZN | 1.22E-74 | 1.9834135 | OPCs_2 |
| 53 | SMOC1 | 7.19E-74 | 5.0247173 | OPCs_2 |
| 54 | NFIA | 3.77E-72 | 2.366833 | OPCs_2 |
| 55 | CSMD3 | 1.29E-71 | 1.6987938 | OPCs_2 |
| 56 | ZEB1 | 5.40E-71 | 2.4074259 | OPCs_2 |
| 57 | NCAM2 | 1.43E-70 | 1.699626 | OPCs_2 |

| | | | | |
|---|---|---|---|---|
| 58 | SOX5 | 3.99E-70 | 2.152337 | OPCs_2 |
| 59 | SEZ6L | 1.59E-67 | 2.4743268 | OPCs_2 |
| 60 | PDGFRA | 4.42E-67 | 5.838708 | OPCs_2 |
| 61 | DGKG | 5.34E-67 | 2.8377786 | OPCs_2 |
| 62 | KAT2B | 2.03E-66 | 3.620963 | OPCs_2 |
| 63 | NKAIN3 | 3.34E-66 | 3.1071968 | OPCs_2 |
| 64 | ASTN2 | 1.03E-65 | 1.9011582 | OPCs_2 |
| 65 | GRIK1 | 1.78E-65 | 2.3414755 | OPCs_2 |
| 66 | NAV1 | 6.20E-64 | 2.3518999 | OPCs_2 |
| 67 | ZNF462 | 2.23E-63 | 3.4568598 | OPCs_2 |
| 68 | PCDH9 | 1.36E-61 | 1.2034156 | OPCs_2 |
| 69 | LRRTM4 | 1.80E-61 | 1.6130339 | OPCs_2 |
| 70 | BCAN | 5.50E-60 | 3.3607802 | OPCs_2 |
| 71 | ABHD2 | 1.27E-59 | 2.64356 | OPCs_2 |
| 72 | APOD | 1.56E-59 | 4.303578 | OPCs_2 |
| 73 | GRM7 | 1.89E-59 | 1.6614747 | OPCs_2 |
| 74 | REV3L | 9.59E-59 | 2.256202 | OPCs_2 |
| 75 | SNTG1 | 1.12E-58 | 1.536152 | OPCs_2 |
| 76 | CDH20 | 1.65E-58 | 2.4354882 | OPCs_2 |
| 77 | WWOX | 2.81E-58 | 1.878486 | OPCs_2 |
| 78 | RP11-436D23.1 | 5.27E-58 | 2.1803763 | OPCs_2 |
| 79 | TMEM132C | 8.48E-58 | 3.1173646 | OPCs_2 |
| 80 | BCAS1 | 3.12E-57 | 3.6803255 | OPCs_2 |
| 81 | NLGN4Y | 6.47E-57 | 2.140524 | OPCs_2 |
| 82 | SPATA6 | 8.46E-57 | 2.8868895 | OPCs_2 |
| 83 | NCKAP5 | 2.35E-56 | 2.677607 | OPCs_2 |
| 84 | PID1 | 3.19E-56 | 2.619018 | OPCs_2 |
| 85 | APBB2 | 5.13E-56 | 2.2556047 | OPCs_2 |
| 86 | ALCAM | 9.33E-56 | 2.2539685 | OPCs_2 |
| 87 | NLGN4X | 2.47E-55 | 2.266383 | OPCs_2 |
| 88 | EDIL3 | 3.29E-53 | 1.9436179 | OPCs_2 |
| 89 | ERBB4 | 3.71E-53 | 1.7918408 | OPCs_2 |
| 90 | ADARB2 | 1.04E-51 | 2.208304 | OPCs_2 |
| 91 | PRKCA | 2.36E-50 | 1.7648981 | OPCs_2 |
| 92 | PTN | 2.66E-50 | 3.0188704 | OPCs_2 |
| 93 | RP11-384F7.2 | 3.50E-50 | 1.4885794 | OPCs_2 |
| 94 | SGCZ | 3.73E-50 | 1.8163036 | OPCs_2 |
| 95 | COL9A1 | 3.73E-50 | 5.020197 | OPCs_2 |
| 96 | 01-Mar | 1.28E-49 | 1.6118789 | OPCs_2 |
| 97 | CHL1 | 1.99E-49 | 1.5948275 | OPCs_2 |
| 98 | MEGF11 | 6.87E-49 | 4.3191094 | OPCs_2 |
| 99 | GPC6 | 1.38E-48 | 1.7820427 | OPCs_2 |
| 0 | PLP1 | 1.10E-120 | 5.854277 | Oligos_1 |
| 1 | CTNNA3 | 3.58E-114 | 3.8100977 | Oligos_1 |
| 2 | ST18 | 9.39E-112 | 5.284247 | Oligos_1 |
| 3 | MBP | 9.39E-112 | 4.4354076 | Oligos_1 |
| 4 | QKI | 3.27E-111 | 3.9780228 | Oligos_1 |
| 5 | SLC44A1 | 3.32E-106 | 3.6859384 | Oligos_1 |
| 6 | MOBP | 1.21E-104 | 5.3099666 | Oligos_1 |
| 7 | PIP4K2A | 9.55E-100 | 4.024533 | Oligos_1 |
| 8 | PDE4B | 1.90E-97 | 3.2280796 | Oligos_1 |
| 9 | RNF220 | 2.09E-97 | 4.1660337 | Oligos_1 |
| 10 | TMEM144 | 2.02E-93 | 4.6653533 | Oligos_1 |

| | | | | |
|---|---|---|---|---|
| 11 | TF | 3.73E-93 | 4.6768513 | Oligos_1 |
| 12 | EDIL3 | 1.46E-89 | 3.1888523 | Oligos_1 |
| 13 | MAN2A1 | 2.43E-89 | 3.3746114 | Oligos_1 |
| 14 | TMTC2 | 2.37E-87 | 3.4305763 | Oligos_1 |
| 15 | DOCK10 | 2.34E-83 | 3.822787 | Oligos_1 |
| 16 | IL1RAPL1 | 1.55E-82 | 2.2344797 | Oligos_1 |
| 17 | CLDN11 | 1.64E-81 | 4.201706 | Oligos_1 |
| 18 | PLCL1 | 2.59E-81 | 3.1408222 | Oligos_1 |
| 19 | CRYAB | 5.18E-81 | 4.869841 | Oligos_1 |
| 20 | FMNL2 | 5.53E-81 | 2.4781072 | Oligos_1 |
| 21 | MAP7 | 5.28E-80 | 2.8182034 | Oligos_1 |
| 22 | TMEM165 | 1.73E-79 | 3.6833317 | Oligos_1 |
| 23 | PCDH9 | 2.39E-79 | 1.8053099 | Oligos_1 |
| 24 | MIR219A2 | 1.14E-78 | 4.201359 | Oligos_1 |
| 25 | FRMD4B | 3.42E-77 | 4.156839 | Oligos_1 |
| 26 | DPYD | 2.69E-76 | 3.0986128 | Oligos_1 |
| 27 | CNP | 2.22E-75 | 4.221687 | Oligos_1 |
| 28 | TTLL7 | 4.11E-74 | 2.549769 | Oligos_1 |
| 29 | NCKAP5 | 4.15E-74 | 3.8039365 | Oligos_1 |
| 30 | DNM3 | 3.85E-72 | 2.2192705 | Oligos_1 |
| 31 | ENPP2 | 5.37E-71 | 4.7631583 | Oligos_1 |
| 32 | GPM6B | 1.29E-70 | 2.295639 | Oligos_1 |
| 33 | FRYL | 2.74E-70 | 3.2735987 | Oligos_1 |
| 34 | ZBTB20 | 3.73E-70 | 2.9336023 | Oligos_1 |
| 35 | SLC24A2 | 5.54E-70 | 2.298719 | Oligos_1 |
| 36 | SHTN1 | 1.24E-67 | 2.7955241 | Oligos_1 |
| 37 | ZEB2 | 2.48E-67 | 2.437933 | Oligos_1 |
| 38 | PHLPP1 | 2.50E-66 | 3.0587828 | Oligos_1 |
| 39 | SCD | 5.55E-66 | 4.328766 | Oligos_1 |
| 40 | SLAIN1 | 8.09E-66 | 3.4017038 | Oligos_1 |
| 41 | TCF12 | 1.91E-65 | 2.8323028 | Oligos_1 |
| 42 | FRMD5 | 2.19E-65 | 2.4481564 | Oligos_1 |
| 43 | UNC5C | 4.35E-65 | 2.6018357 | Oligos_1 |
| 44 | PPP1R14A | 7.94E-64 | 5.043246 | Oligos_1 |
| 45 | LINC00609 | 3.15E-63 | 5.342594 | Oligos_1 |
| 46 | PTGDS | 1.03E-62 | 2.8791869 | Oligos_1 |
| 47 | SPOCK3 | 1.25E-62 | 2.688204 | Oligos_1 |
| 48 | UGT8 | 1.40E-62 | 4.8457046 | Oligos_1 |
| 49 | SPP1 | 1.56E-62 | 4.726975 | Oligos_1 |
| 50 | GAB1 | 2.43E-62 | 3.2150254 | Oligos_1 |
| 51 | PPP2R2B | 5.71E-62 | 2.0742445 | Oligos_1 |
| 52 | CARNS1 | 9.55E-62 | 4.1285734 | Oligos_1 |
| 53 | MAP4K4 | 6.37E-61 | 2.5827937 | Oligos_1 |
| 54 | GSN | 1.85E-60 | 4.0064654 | Oligos_1 |
| 55 | DLG1 | 9.19E-60 | 2.3426542 | Oligos_1 |
| 56 | SEPP1 | 9.32E-59 | 5.034703 | Oligos_1 |
| 57 | CDK18 | 2.56E-57 | 3.9426985 | Oligos_1 |
| 58 | ELMO1 | 1.16E-56 | 2.5472217 | Oligos_1 |
| 59 | SYNJ2 | 2.01E-55 | 3.5533946 | Oligos_1 |
| 60 | ABCA2 | 3.29E-53 | 2.6920447 | Oligos_1 |
| 61 | MAGI2 | 1.14E-52 | 1.4906219 | Oligos_1 |
| 62 | FUT8 | 1.45E-52 | 2.4790173 | Oligos_1 |
| 63 | SIK3 | 2.57E-52 | 1.8141505 | Oligos_1 |

| | | | | |
|---|---|---|---|---|
| 64 | PDE8A | 3.68E-52 | 3.2193627 | Oligos_1 |
| 65 | NEAT1 | 6.42E-52 | 2.7633462 | Oligos_1 |
| 66 | C10orf90 | 8.79E-52 | 4.466444 | Oligos_1 |
| 67 | DOCK5 | 1.73E-51 | 4.076381 | Oligos_1 |
| 68 | ARHGAP21 | 5.55E-51 | 2.2037702 | Oligos_1 |
| 69 | 07-Sep | 6.38E-50 | 1.8856308 | Oligos_1 |
| 70 | MARCKSL1 | 1.69E-49 | 3.7403765 | Oligos_1 |
| 71 | ERBB2IP | 1.29E-48 | 2.796536 | Oligos_1 |
| 72 | NKAIN2 | 4.51E-48 | 1.7207893 | Oligos_1 |
| 73 | CREB5 | 5.87E-48 | 3.0741398 | Oligos_1 |
| 74 | DNAJC6 | 5.53E-47 | 2.1130197 | Oligos_1 |
| 75 | CDH20 | 7.59E-47 | 2.766718 | Oligos_1 |
| 76 | PLEKHH1 | 9.51E-47 | 4.4054537 | Oligos_1 |
| 77 | WSB1 | 2.42E-46 | 1.858798 | Oligos_1 |
| 78 | SLC5A11 | 5.20E-46 | 4.8068185 | Oligos_1 |
| 79 | PEX5L | 7.27E-46 | 1.7393124 | Oligos_1 |
| 80 | S100B | 7.72E-46 | 4.27844 | Oligos_1 |
| 81 | ANLN | 1.12E-45 | 4.741355 | Oligos_1 |
| 82 | TUBA1A | 1.20E-45 | 2.4910152 | Oligos_1 |
| 83 | CNTNAP4 | 1.20E-45 | 3.1237965 | Oligos_1 |
| 84 | CERCAM | 1.40E-45 | 3.104958 | Oligos_1 |
| 85 | MTUS1 | 1.65E-45 | 3.5140426 | Oligos_1 |
| 86 | CLDND1 | 3.14E-45 | 3.491484 | Oligos_1 |
| 87 | CLASP2 | 4.67E-45 | 1.535085 | Oligos_1 |
| 88 | FTH1 | 4.64E-44 | 1.7384053 | Oligos_1 |
| 89 | LMCD1-AS1 | 1.66E-43 | 3.540602 | Oligos_1 |
| 90 | PXK | 1.85E-43 | 3.4314268 | Oligos_1 |
| 91 | USP54 | 6.91E-42 | 3.2763941 | Oligos_1 |
| 92 | TJP1 | 1.27E-41 | 1.8297001 | Oligos_1 |
| 93 | ABCA8 | 2.61E-41 | 4.7670393 | Oligos_1 |
| 94 | LANCL1 | 4.60E-41 | 2.6005344 | Oligos_1 |
| 95 | FAM107B | 7.28E-41 | 4.1613274 | Oligos_1 |
| 96 | MVB12B | 1.65E-40 | 3.1553192 | Oligos_1 |
| 97 | DST | 4.10E-40 | 1.4807159 | Oligos_1 |
| 98 | NPAS3 | 7.14E-40 | 1.6731957 | Oligos_1 |
| 99 | BCAS1 | 8.13E-40 | 3.80733 | Oligos_1 |
| 0 | GPR17 | 3.04E-14 | 9.858534 | Oligos_2 |
| 1 | BCAS1 | 3.04E-14 | 7.1389923 | Oligos_2 |
| 2 | TMEM108 | 2.65E-13 | 4.0281863 | Oligos_2 |
| 3 | MIR219A2 | 4.01E-13 | 5.470532 | Oligos_2 |
| 4 | MAML2 | 7.54E-12 | 5.033277 | Oligos_2 |
| 5 | BCAN | 1.69E-11 | 5.394993 | Oligos_2 |
| 6 | PKP4 | 1.69E-11 | 3.731648 | Oligos_2 |
| 7 | FYN | 2.94E-11 | 4.73862 | Oligos_2 |
| 8 | TNS3 | 4.44E-11 | 6.4543653 | Oligos_2 |
| 9 | SIRT2 | 2.24E-10 | 4.9178233 | Oligos_2 |
| 11 | TNR | 2.48E-10 | 3.8547351 | Oligos_2 |
| 10 | PPP2R2B | 2.48E-10 | 3.212308 | Oligos_2 |
| 12 | SEMA5A | 1.79E-09 | 5.299152 | Oligos_2 |
| 13 | LSAMP | 2.04E-09 | 1.9565979 | Oligos_2 |
| 14 | KAZN | 2.66E-09 | 3.910444 | Oligos_2 |
| 15 | SMOC1 | 4.73E-09 | 6.0861764 | Oligos_2 |
| 16 | SOX6 | 8.29E-09 | 4.326186 | Oligos_2 |

| | | | | |
|---|---|---|---|---|
| 17 | DSCAM | 8.30E-09 | 2.8314245 | Oligos_2 |
| 18 | FRMD5 | 1.30E-08 | 2.7979743 | Oligos_2 |
| 19 | ADAM33 | 2.46E-08 | 6.2918277 | Oligos_2 |
| 20 | QKI | 3.32E-08 | 3.2168448 | Oligos_2 |
| 21 | EPHB1 | 3.32E-08 | 4.4462485 | Oligos_2 |
| 22 | EPB41L2 | 5.27E-08 | 3.0909305 | Oligos_2 |
| 23 | VCAN | 1.06E-07 | 5.0029597 | Oligos_2 |
| 24 | SLC44A1 | 1.41E-07 | 3.0929193 | Oligos_2 |
| 25 | MBP | 3.26E-07 | 3.3334243 | Oligos_2 |
| 26 | TRIO | 4.84E-07 | 2.4922583 | Oligos_2 |
| 27 | SOX8 | 5.71E-07 | 4.879811 | Oligos_2 |
| 28 | FRMD4B | 6.12E-07 | 4.009395 | Oligos_2 |
| 29 | DLG1 | 6.50E-07 | 2.9421475 | Oligos_2 |
| 30 | ZBTB20 | 1.24E-06 | 2.898121 | Oligos_2 |
| 31 | TCF7L2 | 1.96E-06 | 4.321659 | Oligos_2 |
| 32 | COL11A1 | 3.40E-06 | 3.2097356 | Oligos_2 |
| 33 | NFASC | 3.50E-06 | 3.20151 | Oligos_2 |
| 34 | PLP1 | 4.22E-06 | 3.8051472 | Oligos_2 |
| 35 | SEMA5B | 4.49E-06 | 4.4609265 | Oligos_2 |
| 36 | SERINC5 | 4.86E-06 | 3.808717 | Oligos_2 |
| 37 | PCDH15 | 5.62E-06 | 3.098815 | Oligos_2 |
| 38 | PRICKLE1 | 8.51E-06 | 2.9061406 | Oligos_2 |
| 39 | CNP | 8.85E-06 | 3.5262735 | Oligos_2 |
| 40 | FRMD4A | 1.17E-05 | 3.0843918 | Oligos_2 |
| 41 | LINC01170 | 1.58E-05 | 4.98651 | Oligos_2 |
| 42 | SEMA4D | 1.64E-05 | 4.1838403 | Oligos_2 |
| 43 | PTGDS | 1.94E-05 | 2.6785426 | Oligos_2 |
| 44 | SGCD | 2.52E-05 | 2.491669 | Oligos_2 |
| 45 | TNK2 | 3.76E-05 | 2.9206417 | Oligos_2 |
| 46 | ZEB2 | 3.93E-05 | 2.2731638 | Oligos_2 |
| 47 | BMPER | 4.83E-05 | 3.1266603 | Oligos_2 |
| 48 | DDX5 | 5.14E-05 | 2.367004 | Oligos_2 |
| 49 | OPCML | 7.18E-05 | 1.4619232 | Oligos_2 |
| 50 | UGT8 | 0.00011843 | 4.2266827 | Oligos_2 |
| 51 | NCKAP5 | 0.00011843 | 3.210884 | Oligos_2 |
| 52 | OPHN1 | 0.000132112 | 3.3378966 | Oligos_2 |
| 53 | SOX10 | 0.000143152 | 5.9760194 | Oligos_2 |
| 54 | POLR2F | 0.000143152 | 3.740635 | Oligos_2 |
| 55 | MARCKSL1 | 0.000144307 | 3.4324064 | Oligos_2 |
| 56 | CADM1 | 0.000175718 | 2.0437036 | Oligos_2 |
| 57 | SOX4 | 0.000208722 | 5.4626217 | Oligos_2 |
| 58 | FIGN | 0.000208722 | 3.3320873 | Oligos_2 |
| 59 | PTPRJ | 0.00024012 | 2.8222928 | Oligos_2 |
| 60 | LUZP2 | 0.000242791 | 3.006092 | Oligos_2 |
| 61 | PLPPR1 | 0.000252892 | 4.076724 | Oligos_2 |
| 62 | CDK18 | 0.000314457 | 3.645989 | Oligos_2 |
| 63 | ARHGEF7 | 0.000337486 | 1.9414669 | Oligos_2 |
| 64 | CRB1 | 0.00035018 | 4.0102386 | Oligos_2 |
| 65 | RNF144A | 0.000391736 | 2.7402966 | Oligos_2 |
| 66 | LHFPL3 | 0.000407363 | 3.3328915 | Oligos_2 |
| 67 | ARHGAP5 | 0.000475163 | 2.6585402 | Oligos_2 |
| 68 | GNB4 | 0.000478706 | 4.2592998 | Oligos_2 |
| 69 | PCDH11X | 0.000478706 | 2.701984 | Oligos_2 |

| | | | | |
|---|---|---|---|---|
| 70 | P2RX7 | 0.000479351 | 3.5736535 | Oligos_2 |
| 71 | IL1RAPL1 | 0.000614068 | 1.3914765 | Oligos_2 |
| 72 | HIPK2 | 0.000690623 | 3.1891036 | Oligos_2 |
| 73 | RP11-89N17.4 | 0.001394921 | 11.475275 | Oligos_2 |
| 74 | GRID1 | 0.001406575 | 2.2471402 | Oligos_2 |
| 75 | NCAM1 | 0.001512147 | 1.6581277 | Oligos_2 |
| 76 | TMEM132B | 0.001621301 | 1.6955763 | Oligos_2 |
| 77 | OLIG1 | 0.00175711 | 4.1179533 | Oligos_2 |
| 78 | PPP1R16B | 0.002050605 | 3.0985904 | Oligos_2 |
| 79 | MDGA2 | 0.002070929 | 1.5256267 | Oligos_2 |
| 80 | PHYHIPL | 0.002683919 | 2.4724426 | Oligos_2 |
| 81 | MMP16 | 0.002693293 | 2.0094137 | Oligos_2 |
| 82 | SCD5 | 0.002746692 | 2.3134978 | Oligos_2 |
| 83 | ABTB2 | 0.002849364 | 5.240189 | Oligos_2 |
| 84 | TCF12 | 0.003078385 | 2.198358 | Oligos_2 |
| 85 | ANKRD44 | 0.003083022 | 2.6498582 | Oligos_2 |
| 86 | DNER | 0.003083022 | 2.1143882 | Oligos_2 |
| 87 | NLGN1 | 0.003430727 | 1.3333929 | Oligos_2 |
| 88 | PDE4B | 0.00370742 | 2.2057793 | Oligos_2 |
| 89 | TMEM163 | 0.003751664 | 3.9372761 | Oligos_2 |
| 90 | TSC22D4 | 0.003941433 | 3.217035 | Oligos_2 |
| 91 | CERCAM | 0.004791069 | 2.7162266 | Oligos_2 |
| 92 | CADM4 | 0.004859238 | 3.4197676 | Oligos_2 |
| 93 | LINC00511 | 0.00486458 | 3.4634159 | Oligos_2 |
| 94 | SCRG1 | 0.004948728 | 4.016106 | Oligos_2 |
| 95 | SGK1 | 0.005164341 | 2.708062 | Oligos_2 |
| 96 | KANK1 | 0.005222939 | 3.299302 | Oligos_2 |
| 97 | ZNF469 | 0.005700359 | 6.600515 | Oligos_2 |
| 98 | MOB3B | 0.005746344 | 2.843453 | Oligos_2 |
| 99 | GALNT13 | 0.007322293 | 2.2314785 | Oligos_2 |
| 0 | MBP | 0 | 4.304231 | Oligos_3 |
| 13 | PPP2R2B | 0 | 1.7635033 | Oligos_3 |
| 12 | PIP4K2A | 0 | 3.8114605 | Oligos_3 |
| 10 | PTGDS | 0 | 2.8818562 | Oligos_3 |
| 9 | MIR219A2 | 0 | 5.1916113 | Oligos_3 |
| 8 | CERCAM | 0 | 5.106231 | Oligos_3 |
| 7 | SLC44A1 | 0 | 3.256061 | Oligos_3 |
| 11 | PDE4B | 0 | 2.636305 | Oligos_3 |
| 5 | CTNNA3 | 0 | 3.1679113 | Oligos_3 |
| 4 | IL1RAPL1 | 0 | 2.1669762 | Oligos_3 |
| 3 | QKI | 0 | 3.57202 | Oligos_3 |
| 2 | PCDH9 | 0 | 1.8521734 | Oligos_3 |
| 1 | PLP1 | 0 | 5.947841 | Oligos_3 |
| 6 | ST18 | 0 | 5.9010925 | Oligos_3 |
| 14 | TMEM144 | 5.12E-300 | 4.902759 | Oligos_3 |
| 15 | RNF220 | 6.42E-298 | 3.58183 | Oligos_3 |
| 16 | CLDN11 | 3.35E-294 | 4.184939 | Oligos_3 |
| 17 | EDIL3 | 2.21E-270 | 2.6473203 | Oligos_3 |
| 18 | SIK3 | 4.52E-270 | 1.8422768 | Oligos_3 |
| 19 | MAN2A1 | 4.29E-264 | 2.6810725 | Oligos_3 |
| 20 | TTLL7 | 7.27E-256 | 2.3052492 | Oligos_3 |
| 21 | PHLPP1 | 3.54E-250 | 3.0646865 | Oligos_3 |
| 22 | TMTC2 | 4.99E-250 | 2.7584622 | Oligos_3 |

| | | | | |
|---|---|---|---|---|
| 23 | DOCK10 | 4.22E-241 | 3.2559965 | Oligos_3 |
| 24 | MOBP | 9.39E-231 | 4.8015614 | Oligos_3 |
| 25 | TF | 4.29E-228 | 4.103391 | Oligos_3 |
| 26 | SLC24A2 | 3.22E-225 | 1.696154 | Oligos_3 |
| 27 | ZBTB20 | 5.36E-216 | 2.384147 | Oligos_3 |
| 28 | CARNS1 | 1.63E-211 | 4.899292 | Oligos_3 |
| 29 | ABCA2 | 6.39E-211 | 2.9432595 | Oligos_3 |
| 30 | MAGI2 | 4.45E-201 | 0.84745526 | Oligos_3 |
| 31 | CNP | 4.69E-200 | 3.9278622 | Oligos_3 |
| 32 | FRMD5 | 8.77E-192 | 1.9101843 | Oligos_3 |
| 33 | TMEM165 | 1.24E-184 | 3.2788591 | Oligos_3 |
| 34 | KCNH8 | 3.59E-169 | 4.470925 | Oligos_3 |
| 35 | FMNL2 | 1.92E-157 | 1.4636873 | Oligos_3 |
| 36 | FRMD4B | 2.98E-151 | 3.7294946 | Oligos_3 |
| 37 | SPOCK3 | 8.69E-150 | 2.0626411 | Oligos_3 |
| 38 | CREB5 | 1.05E-149 | 3.1395295 | Oligos_3 |
| 39 | NCKAP5 | 1.08E-149 | 2.9970818 | Oligos_3 |
| 40 | MYRF | 8.02E-135 | 5.291235 | Oligos_3 |
| 41 | ERBB2IP | 1.87E-126 | 2.6221912 | Oligos_3 |
| 42 | SHTN1 | 3.37E-125 | 2.388589 | Oligos_3 |
| 43 | TCF12 | 8.20E-124 | 2.2295406 | Oligos_3 |
| 44 | ENPP2 | 3.33E-122 | 4.741956 | Oligos_3 |
| 45 | LAMP2 | 2.18E-118 | 3.4910877 | Oligos_3 |
| 46 | DOCK5 | 2.62E-114 | 4.279959 | Oligos_3 |
| 47 | PHLDB1 | 1.38E-109 | 3.4588857 | Oligos_3 |
| 48 | LPAR1 | 2.01E-104 | 4.7616115 | Oligos_3 |
| 49 | DLG2 | 7.77E-103 | 0.51861966 | Oligos_3 |
| 50 | GPM6B | 7.21E-99 | 1.376742 | Oligos_3 |
| 51 | MOG | 1.08E-97 | 5.1307836 | Oligos_3 |
| 52 | PLCL1 | 2.93E-97 | 2.0592158 | Oligos_3 |
| 53 | SLCO1A2 | 5.26E-97 | 5.207808 | Oligos_3 |
| 54 | UGT8 | 1.84E-95 | 4.6060476 | Oligos_3 |
| 55 | LINC01608 | 1.90E-94 | 5.6776004 | Oligos_3 |
| 56 | ZEB2 | 4.71E-90 | 1.5951632 | Oligos_3 |
| 57 | SCD | 1.31E-87 | 3.460065 | Oligos_3 |
| 58 | FRYL | 1.32E-85 | 2.349375 | Oligos_3 |
| 59 | PPP1R14A | 2.15E-83 | 4.6319537 | Oligos_3 |
| 60 | CNDP1 | 1.32E-82 | 5.2595267 | Oligos_3 |
| 61 | C10orf90 | 2.90E-82 | 4.2934804 | Oligos_3 |
| 62 | ELMO1 | 1.47E-80 | 1.9623629 | Oligos_3 |
| 63 | BCAS1 | 1.54E-79 | 3.5537713 | Oligos_3 |
| 64 | DNM3 | 1.62E-79 | 1.2521791 | Oligos_3 |
| 65 | SLAIN1 | 2.31E-79 | 2.5885139 | Oligos_3 |
| 66 | GSN | 3.46E-77 | 3.169191 | Oligos_3 |
| 67 | ANK3 | 4.58E-77 | 0.95637435 | Oligos_3 |
| 68 | NCAM2 | 1.32E-76 | 0.96024346 | Oligos_3 |
| 69 | MALAT1 | 2.49E-74 | 0.66896665 | Oligos_3 |
| 70 | CLDND1 | 5.10E-74 | 3.0049238 | Oligos_3 |
| 71 | COL4A5 | 1.23E-73 | 4.427425 | Oligos_3 |
| 72 | SPP1 | 2.25E-71 | 3.7627468 | Oligos_3 |
| 73 | DLG1 | 1.23E-66 | 1.5076953 | Oligos_3 |
| 74 | MAP7 | 2.34E-64 | 1.7869042 | Oligos_3 |
| 75 | HHIP | 4.62E-64 | 3.3289928 | Oligos_3 |

| | | | | |
|---|---|---|---|---|
| 76 | DAAM2 | 1.76E-63 | 3.5500126 | Oligos_3 |
| 77 | 07-Sep | 1.01E-62 | 1.2515025 | Oligos_3 |
| 78 | CDK18 | 4.52E-62 | 3.2760477 | Oligos_3 |
| 79 | FAM107B | 1.80E-61 | 4.1742344 | Oligos_3 |
| 80 | UNC5C | 5.65E-60 | 1.6859434 | Oligos_3 |
| 81 | CADM2 | 7.98E-60 | 0.32184383 | Oligos_3 |
| 82 | MAG | 9.38E-60 | 5.1077547 | Oligos_3 |
| 83 | QDPR | 4.48E-58 | 2.461047 | Oligos_3 |
| 84 | NKAIN2 | 2.14E-54 | 0.60909176 | Oligos_3 |
| 85 | SEMA3B | 1.83E-52 | 3.6963418 | Oligos_3 |
| 86 | GPR37 | 6.72E-52 | 4.439332 | Oligos_3 |
| 87 | PDE1C | 1.83E-51 | 2.139777 | Oligos_3 |
| 88 | DPYD | 8.08E-51 | 1.7415882 | Oligos_3 |
| 89 | CNTN2 | 3.32E-50 | 2.9726896 | Oligos_3 |
| 90 | PLLP | 8.11E-50 | 3.9763074 | Oligos_3 |
| 91 | NEAT1 | 7.46E-48 | 1.6335096 | Oligos_3 |
| 92 | SGK1 | 1.27E-46 | 2.0608244 | Oligos_3 |
| 93 | 01-Mar | 1.65E-46 | 1.2392763 | Oligos_3 |
| 94 | CRYAB | 2.25E-46 | 2.7892957 | Oligos_3 |
| 95 | SEPP1 | 3.11E-46 | 4.1098638 | Oligos_3 |
| 96 | FBXL7 | 1.89E-45 | 1.9086041 | Oligos_3 |
| 97 | ABCA8 | 1.13E-44 | 4.938488 | Oligos_3 |
| 98 | MAP4K4 | 4.01E-44 | 1.6305306 | Oligos_3 |
| 99 | ZNF536 | 5.90E-44 | 1.9007375 | Oligos_3 |
