## Supplementary Table 2 for "CellDART: Cell type inference by domain adaptation of single-cell and spatial transcriptomic data"

| **Table 2** | The top 100 marker genes for each cell cluster in mouse brain single-cell dataset obtained from primary visual cortex and anterior lateral motor cortex. The genes were ranked based on the Benjamini-Hochberg adjusted p-values. The name of the cells are based on the metadata provided by the paper (Tasic, B. et al. Shared and distinct transcriptomic cell types across neocortical areas. *Nature* **563**, 72-78 (2018)). | | | |
|---|---|---|---|---|
| Number | Genes | Adjusted p-value | Log fold change | Cluster |
| 0 | Slc1a3 | 0 | 14.274208 | Astro |
| 1 | Acsbg1 | 0 | 12.911855 | Astro |
| 2 | Slco1c1 | 0 | 14.211376 | Astro |
| 3 | Ptprz1 | 0 | 9.446946 | Astro |
| 4 | Fam107a | 0 | 11.986991 | Astro |
| 5 | Malat1 | 0 | 2.4290671 | Astro |
| 6 | Dbi | 0 | 4.5168085 | Astro |
| 7 | Gstm2-ps1 | 0 | 7.54976 | Astro |
| 8 | Hepacam | 0 | 10.789352 | Astro |
| 9 | Gm3764 | 0 | 5.596883 | Astro |
| 10 | Sox9 | 0 | 8.725215 | Astro |
| 11 | Cst3 | 0 | 4.613614 | Astro |
| 12 | Qk | 0 | 6.246723 | Astro |
| 13 | Slc27a1 | 0 | 7.091719 | Astro |
| 14 | Cd63 | 0 | 8.064554 | Astro |
| 15 | Fgfr3 | 0 | 11.2232065 | Astro |
| 16 | Il18 | 0 | 8.462346 | Astro |
| 17 | Tsc22d4 | 0 | 9.206058 | Astro |
| 18 | Slc7a10 | 0 | 12.701325 | Astro |
| 19 | Son | 0 | 2.4091733 | Astro |
| 20 | S100a1 | 0 | 9.563618 | Astro |
| 21 | Luzp2 | 0 | 7.788631 | Astro |
| 22 | Mertk | 0 | 11.581311 | Astro |
| 23 | Msmo1 | 0 | 4.8771753 | Astro |
| 24 | Apoe | 0 | 12.930961 | Astro |
| 25 | Sat1 | 0 | 4.673065 | Astro |
| 26 | Gstm1 | 0 | 10.901285 | Astro |
| 27 | Clu | 0 | 4.116528 | Astro |
| 28 | Ntsr2 | 0 | 12.645241 | Astro |
| 29 | Gpr37l1 | 0 | 14.68437 | Astro |
| 30 | Ppap2b | 0 | 12.252937 | Astro |
| 31 | Htra1 | 0 | 9.822572 | Astro |
| 32 | Gja1 | 0 | 14.385135 | Astro |
| 33 | Slc1a2 | 0 | 6.5059643 | Astro |
| 34 | Mt2 | 0 | 11.003161 | Astro |
| 35 | Aldoc | 0 | 10.4162855 | Astro |
| 36 | Mt1 | 0 | 7.753265 | Astro |
| 37 | Mt3 | 0 | 3.3202364 | Astro |

| | | | | |
|---|---|---|---|---|
| 38 | Cldn10 | 0 | 13.440437 | Astro |
| 39 | Pla2g7 | 0 | 11.28443 | Astro |
| 40 | F3 | 0 | 12.612741 | Astro |
| 41 | Prdx6 | 0 | 6.533071 | Astro |
| 42 | Gjb6 | 0 | 14.510004 | Astro |
| 43 | S1pr1 | 0 | 14.1941395 | Astro |
| 44 | Glul | 0 | 6.906565 | Astro |
| 45 | Bcan | 0 | 10.34494 | Astro |
| 46 | Mfge8 | 0 | 13.38221 | Astro |
| 47 | Atp1a2 | 0 | 12.146331 | Astro |
| 48 | Gstm5 | 0 | 3.902836 | Astro |
| 49 | Ndrg2 | 0 | 8.488193 | Astro |
| 50 | Tlcd1 | 6.14E-304 | 12.151042 | Astro |
| 51 | Atp1b2 | 1.45E-303 | 4.3587604 | Astro |
| 52 | Btbd17 | 3.34E-302 | 10.370472 | Astro |
| 53 | Zbtb20 | 6.89E-301 | 8.157679 | Astro |
| 54 | Slc38a3 | 2.52E-299 | 11.954609 | Astro |
| 55 | Acsl3 | 5.98E-298 | 4.277006 | Astro |
| 56 | Mmd2 | 9.31E-297 | 7.0277543 | Astro |
| 57 | Appl2 | 1.59E-294 | 7.910785 | Astro |
| 58 | Prdx1 | 2.19E-294 | 2.6231618 | Astro |
| 59 | Cmtm5 | 1.96E-292 | 11.169573 | Astro |
| 60 | Mlc1 | 3.09E-292 | 12.268967 | Astro |
| 61 | Slc39a12 | 6.99E-291 | 12.822182 | Astro |
| 62 | Vcam1 | 2.51E-290 | 10.720152 | Astro |
| 63 | S100a16 | 5.73E-289 | 10.752025 | Astro |
| 64 | Cd63-ps | 1.08E-287 | 4.7810335 | Astro |
| 65 | Lcat | 5.53E-287 | 12.612626 | Astro |
| 66 | Csrp1 | 7.68E-285 | 8.109918 | Astro |
| 67 | Nat8 | 1.17E-284 | 12.282455 | Astro |
| 68 | Car2 | 1.74E-282 | 8.504835 | Astro |
| 69 | Fxyd1 | 5.62E-280 | 6.56365 | Astro |
| 70 | Dbx2 | 1.45E-277 | 10.510903 | Astro |
| 71 | Hmgn3 | 2.42E-277 | 2.5272176 | Astro |
| 72 | Slc4a4 | 3.88E-277 | 7.671261 | Astro |
| 73 | Fam20a | 4.06E-277 | 8.96129 | Astro |
| 74 | Ttyh1 | 2.00E-275 | 3.6714222 | Astro |
| 75 | Sdc4 | 1.04E-274 | 9.420905 | Astro |
| 76 | Fermt2 | 2.37E-274 | 6.185341 | Astro |
| 77 | Daam2 | 9.86E-274 | 9.520581 | Astro |
| 78 | Aqp4 | 1.91E-273 | 12.957885 | Astro |
| 79 | Cml1 | 9.25E-271 | 6.6435556 | Astro |
| 80 | Rsrp1 | 1.76E-270 | 1.9717891 | Astro |
| 81 | Prex2 | 9.86E-267 | 8.604738 | Astro |
| 82 | Nkain4 | 4.50E-265 | 6.5771074 | Astro |
| 83 | Dclk1 | 6.64E-265 | 2.591302 | Astro |
| 84 | Dio2 | 1.38E-262 | 10.132815 | Astro |
| 85 | Slc25a18 | 2.13E-262 | 10.584597 | Astro |
| 86 | Sfxn5 | 3.07E-261 | 5.402913 | Astro |
| 87 | Ptplb | 1.34E-260 | 5.7348166 | Astro |

| | | | | |
|---|---|---|---|---|
| 88 | Vegfa | 2.22E-260 | 5.0421495 | Astro |
| 89 | Gng5 | 7.45E-260 | 5.89426 | Astro |
| 90 | Slc9a3r1 | 1.41E-259 | 11.0562525 | Astro |
| 91 | Cspg5 | 1.55E-259 | 2.7436461 | Astro |
| 92 | Tril | 8.44E-259 | 10.529548 | Astro |
| 93 | Gm12222 | 5.49E-258 | 4.2850246 | Astro |
| 94 | Id4 | 5.20E-257 | 7.723402 | Astro |
| 95 | Timp4 | 5.59E-256 | 7.8818736 | Astro |
| 96 | Sparcl1 | 9.53E-256 | 3.341848 | Astro |
| 97 | Macf1 | 8.88E-255 | 2.7072437 | Astro |
| 98 | Nfia | 1.25E-254 | 5.7832522 | Astro |
| 99 | Ntrk2 | 7.59E-254 | 2.865673 | Astro |
| 0 | Sdcbp | 8.27E-65 | 1.98193 | Batch Grouping |
| 1 | Pou3f1 | 1.66E-62 | 6.55609 | Batch Grouping |
| 2 | Serpine2 | 6.20E-62 | 6.6125593 | Batch Grouping |
| 3 | LOC102634502 | 1.02E-58 | 5.531138 | Batch Grouping |
| 4 | Gm6789 | 6.87E-58 | 3.1627858 | Batch Grouping |
| 5 | Tmem163 | 1.81E-52 | 6.19174 | Batch Grouping |
| 6 | Chst8 | 2.91E-52 | 5.5039053 | Batch Grouping |
| 7 | Grp | 4.91E-52 | 6.634044 | Batch Grouping |
| 8 | Acer2 | 1.32E-50 | 5.0643177 | Batch Grouping |
| 9 | Mt1 | 8.36E-49 | 3.2623336 | Batch Grouping |
| 10 | Gmpr | 8.36E-49 | 2.682851 | Batch Grouping |
| 11 | Fezf2 | 3.76E-48 | 6.0384283 | Batch Grouping |
| 12 | Gm19410 | 5.15E-47 | 5.2114754 | Batch Grouping |
| 13 | Meis2 | 9.64E-47 | 5.608861 | Batch Grouping |
| 14 | Kcng1 | 2.85E-46 | 5.5372286 | Batch Grouping |
| 15 | Igsf21 | 4.74E-46 | 4.900316 | Batch Grouping |
| 16 | Tcerg1l | 1.28E-44 | 5.205834 | Batch Grouping |
| 17 | Gm2164 | 1.77E-44 | 5.701642 | Batch Grouping |
| 18 | Sh3bgrl3 | 4.47E-44 | 1.571868 | Batch Grouping |
| 19 | 1700037H04Rik | 1.80E-42 | 2.009801 | Batch Grouping |
| 20 | Vsig2 | 2.39E-42 | 3.5305004 | Batch Grouping |
| 21 | Fam19a1 | 3.02E-42 | 5.996327 | Batch Grouping |
| 22 | Mt2 | 3.02E-42 | 4.4874005 | Batch Grouping |
| 23 | Calm2 | 5.85E-42 | 0.98928326 | Batch Grouping |
| 24 | Tekt5 | 1.21E-41 | 5.530959 | Batch Grouping |
| 25 | Dtnbp1 | 1.50E-41 | 2.4470952 | Batch Grouping |
| 26 | Jsrp1 | 1.70E-41 | 4.5096765 | Batch Grouping |
| 27 | Id2 | 2.13E-41 | 4.2278905 | Batch Grouping |
| 28 | Pcp4 | 4.67E-41 | 5.4870825 | Batch Grouping |
| 29 | Qrfpr | 3.07E-40 | 6.2849803 | Batch Grouping |
| 30 | Slc20a1 | 3.37E-40 | 2.8511539 | Batch Grouping |
| 31 | Bhlhe40 | 1.97E-39 | 4.4778523 | Batch Grouping |
| 32 | Pik3r6 | 4.34E-39 | 5.2500215 | Batch Grouping |
| 33 | Gstp1 | 8.22E-39 | 0.9541636 | Batch Grouping |
| 34 | Tmem150c | 2.08E-38 | 3.8609676 | Batch Grouping |
| 35 | Pex5l | 1.68E-37 | 3.3829057 | Batch Grouping |
| 36 | Bend5 | 2.49E-37 | 3.6780548 | Batch Grouping |
| 37 | Slc5a5 | 1.72E-36 | 4.6554174 | Batch Grouping |

| | | | | |
|---|---|---|---|---|
| 38 | Ier5l | 4.14E-36 | 2.8039343 | Batch Grouping |
| 39 | B230206L02Rik | 7.08E-36 | 4.523649 | Batch Grouping |
| 40 | Npr3 | 1.06E-35 | 4.954122 | Batch Grouping |
| 41 | Cyb5r1 | 1.06E-35 | 3.440509 | Batch Grouping |
| 42 | Tiam2 | 1.27E-35 | 3.8514578 | Batch Grouping |
| 43 | Myh7 | 1.32E-35 | 4.926088 | Batch Grouping |
| 44 | Rasgrf1 | 2.72E-35 | 2.2659972 | Batch Grouping |
| 45 | Kif22 | 1.14E-34 | 2.972156 | Batch Grouping |
| 46 | Ckmt1 | 1.56E-34 | 1.464448 | Batch Grouping |
| 47 | Rgma | 1.56E-34 | 3.4129639 | Batch Grouping |
| 48 | Slit3 | 1.60E-34 | 3.2731912 | Batch Grouping |
| 49 | Tenm3 | 2.57E-34 | 3.6015894 | Batch Grouping |
| 50 | Chn1 | 2.63E-34 | 1.7756441 | Batch Grouping |
| 51 | Enc1 | 4.30E-34 | 3.4349446 | Batch Grouping |
| 52 | Ptk2b | 7.67E-34 | 4.448748 | Batch Grouping |
| 53 | Syt17 | 7.76E-34 | 4.4171515 | Batch Grouping |
| 54 | Crym | 8.21E-34 | 6.10533 | Batch Grouping |
| 55 | Ephx1 | 1.19E-33 | 4.703078 | Batch Grouping |
| 56 | Gm36916 | 6.07E-33 | 3.2279513 | Batch Grouping |
| 57 | Mchr1 | 9.65E-33 | 3.8296688 | Batch Grouping |
| 58 | Btg3 | 1.43E-32 | 2.4877667 | Batch Grouping |
| 59 | Gnb5 | 2.27E-32 | 1.4694318 | Batch Grouping |
| 60 | Pak1 | 5.91E-32 | 1.8123316 | Batch Grouping |
| 61 | Mgp | 7.79E-32 | 5.6648946 | Batch Grouping |
| 62 | Lmo1 | 1.03E-31 | 3.9382317 | Batch Grouping |
| 63 | Arpc5 | 1.08E-31 | 1.5573627 | Batch Grouping |
| 64 | Lmo7 | 1.43E-31 | 3.5129762 | Batch Grouping |
| 65 | Dcdc2a | 1.58E-31 | 4.108056 | Batch Grouping |
| 66 | Fgfr1 | 3.76E-31 | 3.6763809 | Batch Grouping |
| 67 | Bhlhe41 | 4.37E-31 | 2.9130774 | Batch Grouping |
| 68 | Tanc1 | 1.07E-30 | 3.968072 | Batch Grouping |
| 69 | Casz1 | 1.41E-30 | 3.3698537 | Batch Grouping |
| 70 | Nrp1 | 1.90E-30 | 4.098777 | Batch Grouping |
| 71 | Myo5b | 2.51E-30 | 4.0609703 | Batch Grouping |
| 72 | Rbm24 | 2.59E-30 | 3.4112258 | Batch Grouping |
| 73 | Mas1 | 3.51E-30 | 4.621208 | Batch Grouping |
| 74 | Galnt9 | 4.30E-30 | 3.4613278 | Batch Grouping |
| 75 | Fam20a | 5.82E-30 | 3.9474947 | Batch Grouping |
| 76 | Nrg1 | 7.23E-30 | 3.3933833 | Batch Grouping |
| 77 | Tspan17 | 1.69E-29 | 3.0050495 | Batch Grouping |
| 78 | Prkcg | 1.88E-29 | 3.351887 | Batch Grouping |
| 79 | Ctsz | 2.07E-29 | 3.7644439 | Batch Grouping |
| 80 | Arhgdib | 2.31E-29 | 4.5783496 | Batch Grouping |
| 81 | Ryr1 | 2.33E-29 | 4.0605454 | Batch Grouping |
| 82 | ND3 | 3.38E-29 | 1.0936569 | Batch Grouping |
| 83 | ND1 | 3.45E-29 | 0.70689625 | Batch Grouping |
| 84 | Prss12 | 3.91E-29 | 4.012462 | Batch Grouping |
| 85 | Igfbp6 | 7.63E-29 | 4.7463956 | Batch Grouping |
| 86 | Nefl | 1.03E-28 | 3.6073232 | Batch Grouping |
| 87 | Gsta4 | 1.20E-28 | 3.6480896 | Batch Grouping |

| | | | | |
|---|---|---|---|---|
| 88 | Fam105a | 1.60E-28 | 3.512525 | Batch Grouping |
| 89 | Gap43 | 1.91E-28 | 2.0953963 | Batch Grouping |
| 90 | Gm34583 | 2.19E-28 | 4.588051 | Batch Grouping |
| 91 | C1ql3 | 3.84E-28 | 3.9356632 | Batch Grouping |
| 92 | Rapgef5 | 4.53E-28 | 3.1226652 | Batch Grouping |
| 93 | A830021M18Rik | 4.53E-28 | 3.6823213 | Batch Grouping |
| 94 | Galnt18 | 4.64E-28 | 3.2706394 | Batch Grouping |
| 95 | Esd | 5.10E-28 | 1.0892463 | Batch Grouping |
| 96 | Fkbp1a | 5.82E-28 | 0.8381173 | Batch Grouping |
| 97 | Tmem91 | 5.92E-28 | 3.7253025 | Batch Grouping |
| 98 | Tmsb4x | 6.47E-28 | 1.0237873 | Batch Grouping |
| 99 | Car12 | 6.56E-28 | 4.574206 | Batch Grouping |
| 0 | Ramp1 | 1.66E-07 | 5.990899 | CR |
| 1 | H3f3b | 1.66E-07 | 1.7762026 | CR |
| 2 | Trp73 | 1.66E-07 | 13.47879 | CR |
| 3 | Reln | 1.66E-07 | 9.33519 | CR |
| 4 | Ndnf | 2.56E-07 | 11.5281 | CR |
| 5 | Rps19 | 4.55E-07 | 1.9873161 | CR |
| 6 | Lhx1os | 5.25E-07 | 15.813924 | CR |
| 7 | Lhx5 | 5.25E-07 | 13.6278925 | CR |
| 8 | Gm27199 | 5.25E-07 | 12.631134 | CR |
| 9 | Gm15427 | 5.48E-07 | 2.0517836 | CR |
| 10 | Cacna2d2 | 6.58E-07 | 6.4861655 | CR |
| 11 | Syndig1l | 8.68E-07 | 8.599677 | CR |
| 12 | Ptma | 8.85E-07 | 1.4656489 | CR |
| 13 | Gm11223 | 1.21E-06 | 2.106205 | CR |
| 14 | Rnpc3 | 1.25E-06 | 2.9997382 | CR |
| 15 | Rps29 | 1.32E-06 | 1.2893865 | CR |
| 16 | Stmn1-rs2 | 1.65E-06 | 2.2488685 | CR |
| 17 | Malat1 | 1.91E-06 | 1.5380827 | CR |
| 18 | Rpl37a | 1.93E-06 | 1.299804 | CR |
| 19 | Gm4149 | 1.97E-06 | 1.7279944 | CR |
| 20 | Stmn1 | 2.08E-06 | 1.8111552 | CR |
| 21 | Rps27 | 2.43E-06 | 1.7510824 | CR |
| 22 | Tmem163 | 2.60E-06 | 7.5504556 | CR |
| 23 | Rpl37 | 2.90E-06 | 1.4813868 | CR |
| 24 | Emx2 | 3.61E-06 | 9.104825 | CR |
| 25 | Rpl7 | 4.18E-06 | 1.3294308 | CR |
| 26 | Cetn3 | 4.48E-06 | 1.566 | CR |
| 27 | Rps23 | 4.55E-06 | 1.2867173 | CR |
| 28 | Gm10076 | 5.06E-06 | 1.5162952 | CR |
| 29 | Twsg1 | 5.07E-06 | 5.5279393 | CR |
| 30 | Gm6485 | 5.07E-06 | 1.8108797 | CR |
| 31 | Rps8 | 5.50E-06 | 1.126416 | CR |
| 32 | Marcks | 5.63E-06 | 2.5160725 | CR |
| 33 | Rpl18a | 6.66E-06 | 1.3788326 | CR |
| 34 | Tbr1 | 7.54E-06 | 4.9479556 | CR |
| 35 | H3f3a | 7.58E-06 | 1.116428 | CR |
| 36 | Msi2 | 8.48E-06 | 3.7908418 | CR |
| 37 | Cxcl12 | 9.54E-06 | 7.920961 | CR |

| | | | | |
|---|---|---|---|---|
| 38 | Uba52 | 1.26E-05 | 1.1067381 | CR |
| 39 | Phpt1 | 1.47E-05 | 1.2958254 | CR |
| 40 | Gm10269 | 1.50E-05 | 2.0422463 | CR |
| 41 | Vbp1 | 2.42E-05 | 1.6254082 | CR |
| 42 | Gm2830 | 2.44E-05 | 2.9628499 | CR |
| 43 | Eef1a1 | 2.58E-05 | 1.2448398 | CR |
| 44 | Gas5 | 2.59E-05 | 1.7527174 | CR |
| 45 | Rpl30 | 2.59E-05 | 1.3202765 | CR |
| 46 | Gm8960 | 3.07E-05 | 2.0065193 | CR |
| 47 | Nr2f2 | 3.48E-05 | 5.4222875 | CR |
| 48 | Rpl37rt | 3.53E-05 | 1.2377051 | CR |
| 49 | Gm5905 | 4.11E-05 | 1.6476927 | CR |
| 50 | Tmem176b | 4.34E-05 | 7.902608 | CR |
| 51 | Clstn2 | 4.57E-05 | 3.8575578 | CR |
| 52 | Rps14 | 5.35E-05 | 1.1817894 | CR |
| 53 | Anapc11 | 5.40E-05 | 1.2635084 | CR |
| 54 | Rps19-ps6 | 6.19E-05 | 2.0082326 | CR |
| 55 | Pfdn5 | 6.80E-05 | 0.86777306 | CR |
| 56 | Snhg11 | 7.78E-05 | 1.652651 | CR |
| 57 | Cd274 | 8.05E-05 | 8.649834 | CR |
| 58 | Rpl36 | 9.19E-05 | 1.2531471 | CR |
| 59 | Ifitm2 | 9.96E-05 | 6.651039 | CR |
| 60 | Ppia | 9.96E-05 | 0.72885585 | CR |
| 61 | Dach1 | 9.99E-05 | 8.141358 | CR |
| 62 | Rpl41 | 0.000119891 | 0.9318233 | CR |
| 63 | Gm6863 | 0.000119961 | 1.1382104 | CR |
| 64 | Diablo | 0.000133419 | 3.3851993 | CR |
| 65 | Rpl13 | 0.000140878 | 0.99466014 | CR |
| 66 | Nfib | 0.000143312 | 3.329948 | CR |
| 67 | Gm7618 | 0.000144143 | 2.2277234 | CR |
| 68 | Gm13456 | 0.00016644 | 1.533905 | CR |
| 69 | Gm7808 | 0.000173948 | 1.5828496 | CR |
| 70 | Ftl1 | 0.000185237 | 1.1708257 | CR |
| 71 | Rpl35 | 0.000200917 | 1.2705239 | CR |
| 72 | Thrsp | 0.000201775 | 5.4091325 | CR |
| 73 | Tubb4b-ps1 | 0.000212752 | 1.7658753 | CR |
| 74 | LOC105244017 | 0.000213758 | 1.8219523 | CR |
| 75 | Rps9 | 0.000221491 | 0.95802337 | CR |
| 76 | Zbtb20 | 0.000239362 | 5.0634007 | CR |
| 77 | Snora30 | 0.000242768 | 3.4087572 | CR |
| 78 | Ebf3 | 0.000249128 | 12.271501 | CR |
| 79 | Rpl32 | 0.000252732 | 1.1798761 | CR |
| 80 | Stmn1-rs1 | 0.000256294 | 2.2499456 | CR |
| 81 | Cacna2d1 | 0.000261423 | 4.146599 | CR |
| 82 | Btg1 | 0.000269114 | 2.591639 | CR |
| 83 | Prdx2 | 0.0002765 | 1.0366321 | CR |
| 84 | Nhlh2 | 0.000290413 | 9.555491 | CR |
| 85 | Thsd7b | 0.000297382 | 6.5703673 | CR |
| 86 | Gm11469 | 0.000298363 | 1.441591 | CR |
| 87 | 2700094K13Rik | 0.00030557 | 1.805964 | CR |

| | | | | |
|---|---|---|---|---|
| 88 | Cxcr4 | 0.00032207 | 7.684543 | CR |
| 89 | Gm10132 | 0.000347155 | 2.0627842 | CR |
| 90 | Gm6063 | 0.000351213 | 0.84298074 | CR |
| 91 | Naca | 0.000354756 | 0.93645185 | CR |
| 92 | Itm2c | 0.000367474 | 1.0879728 | CR |
| 93 | Gm14303 | 0.000374469 | 1.1384394 | CR |
| 94 | Slc17a6 | 0.000385753 | 6.174703 | CR |
| 95 | Gm17241 | 0.000385753 | 2.0580442 | CR |
| 96 | Tubb4b | 0.000400922 | 1.1368171 | CR |
| 97 | Fam159b | 0.000442042 | 8.247244 | CR |
| 98 | Son | 0.000444226 | 1.1661658 | CR |
| 99 | 1500009L16Rik | 0.000454467 | 3.4342039 | CR |
| 0 | Sparcl1 | 3.75E-34 | 3.1224499 | Doublet |
| 1 | Rgs5 | 5.35E-27 | 8.016713 | Doublet |
| 2 | Wls | 8.40E-24 | 4.310743 | Doublet |
| 3 | 04-Sep | 1.33E-21 | 2.7050803 | Doublet |
| 4 | Gng12 | 5.36E-20 | 4.1689277 | Doublet |
| 5 | Prdx1 | 9.07E-20 | 1.5027114 | Doublet |
| 6 | Tshz2 | 1.17E-19 | 5.8702865 | Doublet |
| 7 | Hist1h2bc | 2.86E-19 | 3.5722096 | Doublet |
| 8 | Dmd | 4.94E-19 | 2.498834 | Doublet |
| 9 | Ptma | 1.81E-18 | 0.9548932 | Doublet |
| 10 | Cd81 | 2.82E-18 | 0.9850287 | Doublet |
| 11 | Pitpnc1 | 3.36E-17 | 3.1769173 | Doublet |
| 12 | Mfge8 | 6.00E-17 | 7.3060665 | Doublet |
| 13 | Ftl1 | 1.70E-16 | 1.1450528 | Doublet |
| 14 | Epas1 | 3.34E-16 | 6.9034405 | Doublet |
| 15 | Gnai2 | 2.14E-15 | 2.0185523 | Doublet |
| 16 | Fermt2 | 3.98E-15 | 3.6425025 | Doublet |
| 17 | Ftl2 | 4.00E-15 | 1.5037867 | Doublet |
| 18 | Myl9 | 5.46E-15 | 8.179342 | Doublet |
| 19 | Atp1a2 | 6.05E-15 | 6.137676 | Doublet |
| 20 | Sat1 | 6.93E-15 | 2.1936033 | Doublet |
| 21 | Glul | 1.05E-14 | 2.5830896 | Doublet |
| 22 | Laptm4a | 1.81E-14 | 1.238223 | Doublet |
| 23 | Pdgfrb | 2.42E-14 | 6.4771805 | Doublet |
| 24 | Gng5 | 3.17E-14 | 3.9547133 | Doublet |
| 25 | Gpcpd1 | 4.12E-14 | 1.5592533 | Doublet |
| 26 | H3f3a | 4.30E-14 | 0.8570291 | Doublet |
| 27 | Rarres2 | 6.22E-14 | 7.134447 | Doublet |
| 28 | Dusp1 | 6.73E-14 | 2.7156703 | Doublet |
| 29 | Apoe | 7.60E-14 | 5.057124 | Doublet |
| 30 | Slco1c1 | 9.56E-14 | 7.280057 | Doublet |
| 31 | Cst3 | 1.21E-13 | 1.6570083 | Doublet |
| 32 | Rbpms | 1.33E-13 | 6.518381 | Doublet |
| 33 | Art3 | 2.49E-13 | 7.2125025 | Doublet |
| 34 | Tmco1 | 3.06E-13 | 1.0481037 | Doublet |
| 35 | Pttg1ip | 3.59E-13 | 3.443898 | Doublet |
| 36 | Cyb5a | 4.37E-13 | 1.171893 | Doublet |
| 37 | Serpinh1 | 4.93E-13 | 6.162554 | Doublet |

| | | | | |
|---|---|---|---|---|
| 38 | Fxyd5 | 5.99E-13 | 3.6190684 | Doublet |
| 39 | Filip1l | 7.37E-13 | 5.625272 | Doublet |
| 40 | Ilk | 1.01E-12 | 1.3871714 | Doublet |
| 41 | Sparc | 1.07E-12 | 6.9794135 | Doublet |
| 42 | Axl | 1.22E-12 | 5.8909 | Doublet |
| 43 | Gm21399 | 1.36E-12 | 1.3949479 | Doublet |
| 44 | Sepp1 | 1.71E-12 | 6.609062 | Doublet |
| 45 | Rnr2 | 1.71E-12 | 0.7291789 | Doublet |
| 46 | Aldh2 | 1.97E-12 | 2.2239764 | Doublet |
| 47 | Sash1 | 2.15E-12 | 3.4932399 | Doublet |
| 48 | Ccdc12 | 2.42E-12 | 0.56586856 | Doublet |
| 49 | Id3 | 2.98E-12 | 5.5914674 | Doublet |
| 50 | 1810037I17Rik | 3.06E-12 | 0.6642268 | Doublet |
| 51 | Zfp36l1 | 3.09E-12 | 5.5327764 | Doublet |
| 52 | Mt1 | 3.17E-12 | 2.4448106 | Doublet |
| 53 | Rgs4 | 3.81E-12 | 2.6222928 | Doublet |
| 54 | S1pr1 | 3.91E-12 | 6.6049685 | Doublet |
| 55 | Son | 3.99E-12 | 0.8777424 | Doublet |
| 56 | Gng11 | 3.99E-12 | 6.943824 | Doublet |
| 57 | Ahnak | 4.92E-12 | 5.2820024 | Doublet |
| 58 | Vim | 5.61E-12 | 6.058855 | Doublet |
| 59 | Fry | 5.64E-12 | 1.7469411 | Doublet |
| 60 | Ifitm2 | 6.13E-12 | 5.2165637 | Doublet |
| 61 | Gm7204 | 1.03E-11 | 1.3681622 | Doublet |
| 62 | Tagln2 | 1.15E-11 | 6.603002 | Doublet |
| 63 | Pon2 | 1.24E-11 | 5.8570185 | Doublet |
| 64 | Ndrg2 | 1.31E-11 | 3.2930205 | Doublet |
| 65 | Ifitm3 | 1.79E-11 | 7.0879927 | Doublet |
| 66 | Neat1 | 2.03E-11 | 5.259353 | Doublet |
| 67 | Gm15776 | 2.09E-11 | 3.8310425 | Doublet |
| 68 | Higd1b | 2.49E-11 | 8.390972 | Doublet |
| 69 | Gstm2-ps1 | 2.68E-11 | 3.4043832 | Doublet |
| 70 | Hspb1 | 3.13E-11 | 7.1515636 | Doublet |
| 71 | S100a13 | 3.29E-11 | 5.0625653 | Doublet |
| 72 | Itga1 | 4.57E-11 | 6.976955 | Doublet |
| 73 | Ptk2 | 5.04E-11 | 1.6647227 | Doublet |
| 74 | Sod3 | 5.86E-11 | 6.6955843 | Doublet |
| 75 | Slc38a11 | 6.01E-11 | 7.8478947 | Doublet |
| 76 | Cox4i2 | 6.23E-11 | 7.1500793 | Doublet |
| 77 | Tspo | 6.40E-11 | 6.0600376 | Doublet |
| 78 | Cnn3 | 6.97E-11 | 3.3789227 | Doublet |
| 79 | Rell1 | 8.04E-11 | 3.1737547 | Doublet |
| 80 | Irak2 | 1.13E-10 | 3.5747585 | Doublet |
| 81 | Vtn | 1.24E-10 | 8.411682 | Doublet |
| 82 | Myo10 | 1.39E-10 | 4.0303645 | Doublet |
| 83 | Cyr61 | 1.46E-10 | 4.5217657 | Doublet |
| 84 | Hsp25-ps1 | 1.79E-10 | 5.5027356 | Doublet |
| 85 | Atox1 | 1.89E-10 | 0.6643933 | Doublet |
| 86 | Tbx3 | 2.00E-10 | 6.55056 | Doublet |
| 87 | Eogt | 2.07E-10 | 4.2573094 | Doublet |

| | | | | |
|---|---|---|---|---|
| 88 | Gm15427 | 2.17E-10 | 0.65386945 | Doublet |
| 89 | Gsta4 | 2.44E-10 | 2.5389016 | Doublet |
| 90 | Tm4sf1 | 3.19E-10 | 7.0789886 | Doublet |
| 91 | Atp6v0e | 3.19E-10 | 2.050589 | Doublet |
| 92 | Nfkbia | 3.65E-10 | 2.548496 | Doublet |
| 93 | Pole4 | 3.73E-10 | 0.8860949 | Doublet |
| 94 | P2ry14 | 3.75E-10 | 6.0982943 | Doublet |
| 95 | Bgn | 3.83E-10 | 6.778779 | Doublet |
| 96 | Rhoc | 3.92E-10 | 4.659768 | Doublet |
| 97 | Jun | 4.66E-10 | 3.2756917 | Doublet |
| 98 | Ptrf | 4.74E-10 | 5.8235483 | Doublet |
| 99 | Ndufa4l2 | 5.32E-10 | 7.027952 | Doublet |
| 0 | Cldn5 | 1.67E-105 | 17.579025 | Endo |
| 1 | Bsg | 1.67E-105 | 4.7468324 | Endo |
| 2 | Ly6c1 | 1.67E-105 | 18.11196 | Endo |
| 3 | Flt1 | 1.67E-105 | 16.249687 | Endo |
| 4 | Abcb1a | 1.67E-105 | 13.295869 | Endo |
| 5 | Ly6a | 1.67E-105 | 17.615965 | Endo |
| 6 | Pltp | 1.67E-105 | 14.755315 | Endo |
| 7 | Ly6c2 | 1.67E-105 | 12.317044 | Endo |
| 8 | Ly6e | 7.10E-105 | 4.2584004 | Endo |
| 9 | Epas1 | 1.00E-103 | 12.118248 | Endo |
| 10 | Slco1a4 | 3.48E-103 | 17.156248 | Endo |
| 11 | Eltd1 | 3.48E-103 | 16.839075 | Endo |
| 12 | Sparc | 4.12E-103 | 13.434359 | Endo |
| 13 | Egfl7 | 4.66E-103 | 7.676469 | Endo |
| 14 | Slc2a1 | 1.23E-102 | 9.764671 | Endo |
| 15 | B2m | 3.00E-102 | 5.293576 | Endo |
| 16 | Id3 | 6.79E-102 | 11.730968 | Endo |
| 17 | Id1 | 9.77E-102 | 12.861944 | Endo |
| 18 | Srgn | 9.77E-102 | 14.338893 | Endo |
| 19 | Ifitm3 | 1.60E-101 | 13.547443 | Endo |
| 20 | Itm2a | 3.09E-101 | 12.701847 | Endo |
| 21 | Igfbp7 | 3.25E-101 | 10.961868 | Endo |
| 22 | Hspb1 | 2.39E-100 | 13.187025 | Endo |
| 23 | Hsp25-ps1 | 2.47E-100 | 8.954566 | Endo |
| 24 | Vwa1 | 2.92E-100 | 9.827453 | Endo |
| 25 | Ramp2 | 5.20E-100 | 9.753706 | Endo |
| 26 | Cgnl1 | 9.71E-100 | 11.205723 | Endo |
| 27 | Wfdc1 | 1.31E-99 | 14.203685 | Endo |
| 28 | Esam | 3.37E-99 | 11.475822 | Endo |
| 29 | Ptprb | 3.37E-99 | 12.758001 | Endo |
| 30 | Klf2 | 6.06E-99 | 11.743693 | Endo |
| 31 | Eng | 7.02E-99 | 11.114922 | Endo |
| 32 | Pglyrp1 | 1.14E-98 | 15.409949 | Endo |
| 33 | Myl12a | 1.29E-98 | 7.4534354 | Endo |
| 34 | Gpr116 | 1.35E-98 | 15.001684 | Endo |
| 35 | Gng5 | 3.58E-98 | 7.3288236 | Endo |
| 36 | Slco1c1 | 9.44E-98 | 13.378256 | Endo |
| 37 | Sgms1 | 1.72E-97 | 7.971572 | Endo |

| | | | | |
|---|---|---|---|---|
| 38 | Slc9a3r2 | 2.08E-97 | 7.17054 | Endo |
| 39 | Atox1 | 3.18E-97 | 3.0292778 | Endo |
| 40 | Ahnak | 6.89E-96 | 10.208049 | Endo |
| 41 | Lef1 | 6.93E-96 | 12.051192 | Endo |
| 42 | Pecam1 | 2.60E-95 | 14.342764 | Endo |
| 43 | Fn1 | 7.25E-95 | 13.352939 | Endo |
| 44 | Crip2 | 1.48E-94 | 5.1461554 | Endo |
| 45 | Anxa3 | 2.28E-94 | 13.435392 | Endo |
| 46 | Tek | 3.38E-94 | 11.986804 | Endo |
| 47 | Gng11 | 4.74E-94 | 11.703766 | Endo |
| 48 | Lsr | 6.22E-94 | 11.503678 | Endo |
| 49 | Ly6e-ps1 | 9.83E-94 | 4.06163 | Endo |
| 50 | Sox18 | 5.42E-93 | 11.295472 | Endo |
| 51 | Abcg2 | 5.91E-93 | 9.166358 | Endo |
| 52 | Sepp1 | 1.04E-92 | 12.165012 | Endo |
| 53 | Nostrin | 1.40E-92 | 12.416829 | Endo |
| 54 | S1pr1 | 2.09E-92 | 11.226425 | Endo |
| 55 | Tmem204 | 2.70E-92 | 11.733855 | Endo |
| 56 | Tmsb4x | 4.21E-92 | 1.997625 | Endo |
| 57 | Ifitm2 | 4.42E-92 | 9.255832 | Endo |
| 58 | 9430020K01Rik | 5.66E-92 | 6.973946 | Endo |
| 59 | Myl6 | 1.37E-91 | 1.7224079 | Endo |
| 60 | Zfp36l1 | 1.40E-91 | 9.728485 | Endo |
| 61 | Ptrf | 8.86E-91 | 9.615698 | Endo |
| 62 | Cyyr1 | 1.21E-89 | 12.313558 | Endo |
| 63 | Tsc22d1 | 1.30E-89 | 2.8124962 | Endo |
| 64 | Ftl1 | 2.01E-89 | 2.0801044 | Endo |
| 65 | Paqr5 | 1.35E-88 | 11.820166 | Endo |
| 66 | Podxl | 1.04E-87 | 10.053718 | Endo |
| 67 | Ctsh | 1.05E-87 | 12.590664 | Endo |
| 68 | Clic4 | 1.28E-87 | 9.025038 | Endo |
| 69 | Fcgrt | 1.62E-87 | 11.703184 | Endo |
| 70 | Icam2 | 1.81E-87 | 14.844834 | Endo |
| 71 | Ctla2a | 1.93E-87 | 15.783712 | Endo |
| 72 | Emcn | 4.10E-87 | 13.932962 | Endo |
| 73 | Sptbn1 | 4.79E-87 | 2.7853417 | Endo |
| 74 | Vim | 6.17E-87 | 10.351045 | Endo |
| 75 | Utrn | 2.09E-86 | 6.262456 | Endo |
| 76 | H3f3a | 4.84E-86 | 1.7178088 | Endo |
| 77 | Nfkbia | 8.87E-86 | 6.5617623 | Endo |
| 78 | Kank3 | 2.03E-85 | 8.116089 | Endo |
| 79 | Arhgap31 | 4.64E-85 | 7.3569956 | Endo |
| 80 | S100a13 | 8.27E-84 | 9.267011 | Endo |
| 81 | Cdh5 | 1.11E-83 | 11.984475 | Endo |
| 82 | Tpm4 | 1.31E-83 | 7.5889306 | Endo |
| 83 | Fxyd5 | 1.69E-83 | 7.907572 | Endo |
| 84 | Ltbp4 | 5.93E-83 | 7.453964 | Endo |
| 85 | Ets1 | 7.34E-83 | 8.999996 | Endo |
| 86 | Kdr | 8.18E-83 | 13.504622 | Endo |
| 87 | Eogt | 1.59E-82 | 8.649793 | Endo |

| | | | | |
|---|---|---|---|---|
| 88 | Acvrl1 | 2.31E-82 | 10.339233 | Endo |
| 89 | Serinc3 | 2.63E-82 | 2.3877952 | Endo |
| 90 | Gpcpd1 | 9.02E-82 | 3.3897433 | Endo |
| 91 | Ccdc141 | 1.63E-81 | 10.397397 | Endo |
| 92 | Ly6i | 1.83E-80 | 9.144091 | Endo |
| 93 | Ndrg1 | 4.85E-79 | 9.224159 | Endo |
| 94 | Klf4 | 2.21E-78 | 10.865626 | Endo |
| 95 | Slc22a8 | 2.90E-78 | 12.717227 | Endo |
| 96 | Rasgrp3 | 3.10E-78 | 10.37504 | Endo |
| 97 | Gnai2 | 3.38E-78 | 4.112298 | Endo |
| 98 | St3gal6 | 8.25E-78 | 7.5145187 | Endo |
| 99 | BC028528 | 8.07E-77 | 9.5493355 | Endo |
| 0 | Ywhag | 2.40E-110 | 2.5293107 | High Intronic |
| 1 | Dgkh | 4.44E-110 | 4.287782 | High Intronic |
| 2 | LOC105246694 | 4.44E-110 | 7.771146 | High Intronic |
| 3 | Slc12a5 | 2.07E-109 | 3.017755 | High Intronic |
| 4 | Cdk5r1 | 2.26E-109 | 3.4850137 | High Intronic |
| 5 | LOC105245046 | 2.26E-109 | 3.2517827 | High Intronic |
| 6 | Epha4 | 2.34E-109 | 5.8136153 | High Intronic |
| 7 | Sort1 | 5.10E-109 | 3.82795 | High Intronic |
| 8 | Kif1a | 5.41E-109 | 2.1692767 | High Intronic |
| 9 | Kcnb1 | 9.36E-109 | 3.7889686 | High Intronic |
| 10 | Plcxd2 | 1.53E-108 | 6.4379563 | High Intronic |
| 11 | Srcin1 | 2.07E-108 | 3.1717348 | High Intronic |
| 12 | Thy1 | 6.22E-108 | 2.5206988 | High Intronic |
| 13 | Ptgfrn | 6.96E-108 | 6.8414063 | High Intronic |
| 14 | Miat | 6.97E-108 | 4.8138766 | High Intronic |
| 15 | Dos | 3.48E-107 | 3.190194 | High Intronic |
| 16 | Kcnma1 | 3.48E-107 | 3.2953084 | High Intronic |
| 17 | Scn8a | 3.48E-107 | 2.8003666 | High Intronic |
| 18 | LOC105245638 | 7.04E-107 | 4.0215516 | High Intronic |
| 19 | D430041D05Rik | 7.61E-107 | 4.487109 | High Intronic |
| 20 | Slc6a17 | 2.88E-106 | 2.9380763 | High Intronic |
| 21 | LOC101056001 | 5.29E-106 | 7.022377 | High Intronic |
| 22 | Kalrn | 1.32E-105 | 3.6461735 | High Intronic |
| 23 | Nsg2 | 1.40E-105 | 2.2190866 | High Intronic |
| 24 | Kcnk2 | 1.53E-105 | 5.155339 | High Intronic |
| 25 | Nuak1 | 6.90E-105 | 5.3341293 | High Intronic |
| 26 | Strbp | 7.85E-105 | 2.2547026 | High Intronic |
| 27 | Nlgn2 | 7.85E-105 | 3.7119305 | High Intronic |
| 28 | Celf2 | 7.85E-105 | 2.175451 | High Intronic |
| 29 | Bsn | 2.17E-104 | 3.234397 | High Intronic |
| 30 | Syt7 | 2.17E-104 | 3.7912755 | High Intronic |
| 31 | Ptprd | 3.80E-104 | 3.4592447 | High Intronic |
| 32 | 9530091C08Rik | 7.27E-104 | 7.526781 | High Intronic |
| 33 | Gfra2 | 1.21E-103 | 5.662164 | High Intronic |
| 34 | Whrn | 1.41E-103 | 7.0207405 | High Intronic |
| 35 | Sorl1 | 1.44E-103 | 4.0233145 | High Intronic |
| 36 | Bai1 | 1.56E-103 | 3.2407143 | High Intronic |
| 37 | Rab6b | 1.65E-103 | 2.2693584 | High Intronic |

| | | | | |
|---|---|---|---|---|
| 38 | Mapt | 5.38E-103 | 1.9192002 | High Intronic |
| 39 | Camk2n1 | 6.18E-103 | 3.4299204 | High Intronic |
| 40 | N28178 | 8.81E-103 | 4.313856 | High Intronic |
| 41 | Wbscr17 | 9.18E-103 | 5.7912903 | High Intronic |
| 42 | R3hdm1 | 2.21E-102 | 3.5323052 | High Intronic |
| 43 | Plxnd1 | 9.58E-102 | 6.6084404 | High Intronic |
| 44 | Pde2a | 1.26E-101 | 3.1281853 | High Intronic |
| 45 | Celsr2 | 1.69E-101 | 2.9766588 | High Intronic |
| 46 | Lars2 | 3.66E-101 | 2.123067 | High Intronic |
| 47 | Cacna1a | 4.45E-101 | 3.6051984 | High Intronic |
| 48 | Mir6240 | 4.74E-101 | 4.2356424 | High Intronic |
| 49 | Grin1 | 4.74E-101 | 2.442615 | High Intronic |
| 50 | Rorb | 8.08E-101 | 7.696785 | High Intronic |
| 51 | Spred2 | 1.49E-100 | 4.303033 | High Intronic |
| 52 | Ptms | 3.83E-100 | 1.9256072 | High Intronic |
| 53 | Cadm3 | 4.31E-100 | 3.04586 | High Intronic |
| 54 | Erc2 | 8.77E-100 | 3.5298846 | High Intronic |
| 55 | 5031426D15Rik | 1.33E-99 | 6.2345576 | High Intronic |
| 56 | Pgm2l1 | 1.54E-99 | 2.900239 | High Intronic |
| 57 | Camk2a | 2.94E-99 | 3.4617574 | High Intronic |
| 58 | Lmtk2 | 6.11E-99 | 3.8976061 | High Intronic |
| 59 | Nrxn2 | 9.23E-99 | 3.086417 | High Intronic |
| 60 | Arpp21 | 1.18E-98 | 5.1385803 | High Intronic |
| 61 | Bai2 | 1.49E-98 | 4.18187 | High Intronic |
| 62 | Dpysl2 | 1.90E-98 | 1.9438907 | High Intronic |
| 63 | A930011O12Rik | 2.41E-98 | 2.926634 | High Intronic |
| 64 | Grin2b | 2.84E-98 | 1.9672918 | High Intronic |
| 65 | Ncan | 3.52E-98 | 3.5993598 | High Intronic |
| 66 | Pvrl1 | 4.95E-98 | 4.980825 | High Intronic |
| 67 | Endou | 1.18E-97 | 8.722289 | High Intronic |
| 68 | LOC101056014 | 1.29E-97 | 3.2024875 | High Intronic |
| 69 | Dlgap4 | 1.36E-97 | 2.4650702 | High Intronic |
| 70 | Arhgap26 | 2.21E-97 | 3.0012252 | High Intronic |
| 71 | R3hdm2 | 2.77E-97 | 3.0178683 | High Intronic |
| 72 | Gpr158 | 4.10E-97 | 4.134208 | High Intronic |
| 73 | Spock2 | 4.10E-97 | 2.208752 | High Intronic |
| 74 | Clstn1 | 5.45E-97 | 1.9963261 | High Intronic |
| 75 | Kcnq1ot1 | 1.32E-96 | 3.2424679 | High Intronic |
| 76 | Trim9 | 1.64E-96 | 2.6948912 | High Intronic |
| 77 | Nrcam | 2.00E-96 | 2.518771 | High Intronic |
| 78 | LOC102634502 | 2.21E-96 | 6.3673205 | High Intronic |
| 79 | Gnaq | 2.21E-96 | 2.753477 | High Intronic |
| 80 | Sel1l3 | 4.20E-96 | 4.5822916 | High Intronic |
| 81 | Kif1b | 5.82E-96 | 1.7476544 | High Intronic |
| 82 | Slc1a2 | 8.21E-96 | 4.625451 | High Intronic |
| 83 | Prrt1 | 9.93E-96 | 2.3959503 | High Intronic |
| 84 | Kcna2 | 1.07E-95 | 2.9867616 | High Intronic |
| 85 | Kcnh1 | 1.44E-95 | 4.971434 | High Intronic |
| 86 | Mical2 | 1.67E-95 | 3.1345432 | High Intronic |
| 87 | Mical3 | 5.21E-95 | 4.2814083 | High Intronic |

| | | | | |
|---|---|---|---|---|
| 88 | Lamc2 | 9.16E-95 | 7.0810595 | High Intronic |
| 89 | Coro2b | 1.03E-94 | 3.5048552 | High Intronic |
| 90 | LOC105243541 | 1.54E-94 | 6.8056173 | High Intronic |
| 91 | Atp2b2 | 1.72E-94 | 2.156082 | High Intronic |
| 92 | Pip4k2b | 1.78E-94 | 3.8953524 | High Intronic |
| 93 | Myo5a | 2.62E-94 | 2.165501 | High Intronic |
| 94 | Tenm4 | 2.85E-94 | 4.0600057 | High Intronic |
| 95 | Krt20 | 2.85E-94 | 6.9853454 | High Intronic |
| 96 | Ptprn | 1.67E-93 | 2.4412758 | High Intronic |
| 97 | Meg3 | 1.96E-93 | 2.498304 | High Intronic |
| 98 | Syne1 | 2.48E-93 | 2.3704226 | High Intronic |
| 99 | Pde4d | 2.61E-93 | 3.202786 | High Intronic |
| 0 | Ppp3ca | 0 | 2.2589562 | L2/3 IT |
| 1 | Prex1 | 0 | 4.4366946 | L2/3 IT |
| 2 | Tspan13 | 0 | 1.5459038 | L2/3 IT |
| 3 | Gria3 | 0 | 2.8927615 | L2/3 IT |
| 4 | Cinp | 0 | 2.1820517 | L2/3 IT |
| 5 | Atp2b1 | 0 | 1.8714474 | L2/3 IT |
| 6 | Synpo | 0 | 3.3675563 | L2/3 IT |
| 7 | Chst1 | 0 | 2.891755 | L2/3 IT |
| 8 | Gsta4 | 0 | 4.365176 | L2/3 IT |
| 9 | Rasl10a | 0 | 5.4429107 | L2/3 IT |
| 10 | Dgkb | 0 | 4.289161 | L2/3 IT |
| 11 | Nrgn | 0 | 4.7673078 | L2/3 IT |
| 12 | Pcsk2 | 0 | 2.9031498 | L2/3 IT |
| 13 | Dusp18 | 0 | 4.846562 | L2/3 IT |
| 14 | Ctxn1 | 0 | 3.5520997 | L2/3 IT |
| 15 | Fam131a | 0 | 2.7437356 | L2/3 IT |
| 16 | Gucy1a3 | 0 | 4.4962873 | L2/3 IT |
| 17 | Kcnv1 | 0 | 5.10065 | L2/3 IT |
| 18 | Nell2 | 0 | 3.381532 | L2/3 IT |
| 19 | Rgs7 | 0 | 2.3907373 | L2/3 IT |
| 20 | Palmd | 0 | 5.0757284 | L2/3 IT |
| 21 | Lhx2 | 0 | 4.8170605 | L2/3 IT |
| 22 | 2010300C02Rik | 0 | 3.841654 | L2/3 IT |
| 23 | Sorbs2 | 0 | 2.83865 | L2/3 IT |
| 24 | Pop5 | 0 | 1.6752318 | L2/3 IT |
| 25 | Pdzrn3 | 0 | 4.6389146 | L2/3 IT |
| 26 | Grasp | 0 | 4.0741973 | L2/3 IT |
| 27 | Dact2 | 0 | 3.2908149 | L2/3 IT |
| 28 | Ppp1r1a | 0 | 2.072663 | L2/3 IT |
| 29 | Rps6ka2 | 0 | 4.1079626 | L2/3 IT |
| 30 | Fam84a | 0 | 4.238161 | L2/3 IT |
| 31 | 9130024F11Rik | 0 | 3.5965345 | L2/3 IT |
| 32 | Plk2 | 0 | 2.4385524 | L2/3 IT |
| 33 | Tsc22d1 | 0 | 1.4451748 | L2/3 IT |
| 34 | Gm20063 | 0 | 4.8530664 | L2/3 IT |
| 35 | Evc2 | 0 | 5.749741 | L2/3 IT |
| 36 | Cyp46a1 | 0 | 1.9347434 | L2/3 IT |
| 37 | Cnksr2 | 0 | 3.397045 | L2/3 IT |

| | | | | |
|---|---|---|---|---|
| 38 | A830009L08Rik | 0 | 4.573567 | L2/3 IT |
| 39 | Rasgrf2 | 0 | 4.18347 | L2/3 IT |
| 40 | 11-Sep | 0 | 1.4909586 | L2/3 IT |
| 41 | Nov | 0 | 4.8174143 | L2/3 IT |
| 42 | Gm30687 | 0 | 4.6609735 | L2/3 IT |
| 43 | LOC102634502 | 0 | 4.330797 | L2/3 IT |
| 44 | Gm34996 | 0 | 5.165784 | L2/3 IT |
| 45 | Rtn4rl1 | 0 | 4.0773625 | L2/3 IT |
| 46 | Actr3b | 0 | 2.292116 | L2/3 IT |
| 47 | Lrrtm4 | 0 | 3.5180705 | L2/3 IT |
| 48 | Gm12371 | 0 | 6.239425 | L2/3 IT |
| 49 | Cpne6 | 0 | 3.1018603 | L2/3 IT |
| 50 | Atp2b4 | 0 | 5.009262 | L2/3 IT |
| 51 | Gsg1l | 0 | 5.3824573 | L2/3 IT |
| 52 | Gpr88 | 0 | 6.3320827 | L2/3 IT |
| 53 | Schip1 | 0 | 2.6732142 | L2/3 IT |
| 54 | Ptk2b | 0 | 5.0606627 | L2/3 IT |
| 55 | Calb1 | 0 | 8.019546 | L2/3 IT |
| 56 | Arl15 | 0 | 3.4280024 | L2/3 IT |
| 57 | Chn1 | 0 | 2.0717173 | L2/3 IT |
| 58 | Arpp21 | 0 | 4.832925 | L2/3 IT |
| 59 | Lamp5 | 0 | 7.5711584 | L2/3 IT |
| 60 | Tesc | 0 | 5.3578186 | L2/3 IT |
| 61 | Fam19a1 | 0 | 7.0831356 | L2/3 IT |
| 62 | Pvrl3 | 0 | 5.6101484 | L2/3 IT |
| 63 | Itpka | 0 | 5.3926 | L2/3 IT |
| 64 | Enpp2 | 0 | 5.8777614 | L2/3 IT |
| 65 | Pdp1 | 0 | 3.9894755 | L2/3 IT |
| 66 | Mapk4 | 0 | 4.9259667 | L2/3 IT |
| 67 | Rilpl1 | 0 | 4.089237 | L2/3 IT |
| 68 | B230216N24Rik | 0 | 5.212615 | L2/3 IT |
| 69 | Cacng3 | 0 | 3.3560903 | L2/3 IT |
| 70 | Myh7 | 0 | 7.049229 | L2/3 IT |
| 71 | Fkbp1a | 0 | 1.4511598 | L2/3 IT |
| 72 | Atp1a1 | 0 | 2.2638116 | L2/3 IT |
| 73 | Tsnax | 0 | 2.3116002 | L2/3 IT |
| 74 | Stard8 | 0 | 6.3800235 | L2/3 IT |
| 75 | Igfbp6 | 0 | 5.859942 | L2/3 IT |
| 76 | Cux2 | 0 | 4.2947984 | L2/3 IT |
| 77 | Baiap2 | 0 | 4.10981 | L2/3 IT |
| 78 | C730002L08Rik | 0 | 5.453882 | L2/3 IT |
| 79 | Camk2n1 | 0 | 2.4775963 | L2/3 IT |
| 80 | Meis2 | 0 | 5.6958375 | L2/3 IT |
| 81 | Pkig | 0 | 1.7165887 | L2/3 IT |
| 82 | Btg3 | 0 | 3.2033741 | L2/3 IT |
| 83 | BC030499 | 0 | 5.6233544 | L2/3 IT |
| 84 | Ppp3r1 | 0 | 1.6499763 | L2/3 IT |
| 85 | Arpp19 | 0 | 2.5535393 | L2/3 IT |
| 86 | Actn1 | 0 | 2.750193 | L2/3 IT |
| 87 | Dusp14 | 0 | 3.7885594 | L2/3 IT |

| | | | | |
|---|---|---|---|---|
| 88 | Kctd1 | 0 | 3.9923794 | L2/3 IT |
| 89 | Wfs1 | 0 | 5.5346026 | L2/3 IT |
| 90 | Arpc2 | 0 | 1.366812 | L2/3 IT |
| 91 | Olfm1 | 0 | 2.178884 | L2/3 IT |
| 92 | Mapk1 | 0 | 1.3866067 | L2/3 IT |
| 93 | Tpm1 | 0 | 1.783683 | L2/3 IT |
| 94 | Neurod1 | 0 | 5.396538 | L2/3 IT |
| 95 | Rgl1 | 0 | 3.6049616 | L2/3 IT |
| 96 | Enc1 | 0 | 4.0502257 | L2/3 IT |
| 97 | Rgs14 | 0 | 5.3693137 | L2/3 IT |
| 98 | Kctd4 | 0 | 5.2193666 | L2/3 IT |
| 99 | Map9 | 0 | 1.855811 | L2/3 IT |
| 0 | Nrxn1 | 0 | 2.1956315 | L4 |
| 1 | Extl3 | 0 | 3.3742664 | L4 |
| 2 | Neurod1 | 0 | 5.2101545 | L4 |
| 3 | Rims2 | 0 | 2.067165 | L4 |
| 4 | Hecw1 | 0 | 2.674302 | L4 |
| 5 | Dkkl1 | 0 | 5.9779077 | L4 |
| 6 | Lingo1 | 0 | 4.653838 | L4 |
| 7 | Pamr1 | 0 | 5.8488407 | L4 |
| 8 | Syt13 | 0 | 2.331786 | L4 |
| 9 | Ramp1 | 0 | 4.000455 | L4 |
| 10 | Syne1 | 0 | 1.9683295 | L4 |
| 11 | Gfra2 | 0 | 4.6790285 | L4 |
| 12 | BC030499 | 0 | 5.611172 | L4 |
| 13 | Unc5d | 0 | 4.6295996 | L4 |
| 14 | Grm2 | 0 | 5.054442 | L4 |
| 15 | Osbpl1a | 0 | 2.460989 | L4 |
| 16 | Exph5 | 0 | 5.177297 | L4 |
| 17 | Gpr158 | 0 | 3.5483625 | L4 |
| 18 | N28178 | 0 | 3.6480396 | L4 |
| 19 | Nrn1 | 0 | 6.1040854 | L4 |
| 20 | Kcnb1 | 0 | 2.7392325 | L4 |
| 21 | Homer1 | 0 | 3.13547 | L4 |
| 22 | Nrgn | 0 | 4.7826977 | L4 |
| 23 | Slc1a2 | 0 | 3.995874 | L4 |
| 24 | Nuak1 | 0 | 3.911679 | L4 |
| 25 | B2m | 0 | 2.4980514 | L4 |
| 26 | Kcnh5 | 0 | 4.816619 | L4 |
| 27 | Tpt1-ps3 | 0 | 1.0466344 | L4 |
| 28 | Cd302 | 0 | 4.2980084 | L4 |
| 29 | Lrrk2 | 0 | 3.853592 | L4 |
| 30 | Grm7 | 0 | 3.3926187 | L4 |
| 31 | Mras | 0 | 2.541743 | L4 |
| 32 | Cadps2 | 0 | 4.929852 | L4 |
| 33 | Adora1 | 0 | 4.075166 | L4 |
| 34 | Lzts3 | 0 | 3.3056405 | L4 |
| 35 | Lrrc4c | 0 | 2.4094956 | L4 |
| 36 | Fam212b | 0 | 4.5494986 | L4 |
| 37 | Cux1 | 0 | 3.7718403 | L4 |

| | | | | |
|---|---|---|---|---|
| 38 | Bai1 | 0 | 2.3441331 | L4 |
| 39 | Ier5 | 0 | 4.095599 | L4 |
| 40 | Asap1 | 0 | 2.0059805 | L4 |
| 41 | Satb2 | 0 | 4.560628 | L4 |
| 42 | Cpne8 | 0 | 4.238524 | L4 |
| 43 | Cnksr2 | 0 | 3.528125 | L4 |
| 44 | Kcnma1 | 0 | 2.4972675 | L4 |
| 45 | Glcci1 | 0 | 3.7205787 | L4 |
| 46 | Arc | 0 | 5.450852 | L4 |
| 47 | Rasl11b | 0 | 4.4168158 | L4 |
| 48 | AI593442 | 0 | 2.6380436 | L4 |
| 49 | Ptn | 0 | 5.0647306 | L4 |
| 50 | Egr1 | 0 | 4.623042 | L4 |
| 51 | Egr3 | 0 | 5.4745502 | L4 |
| 52 | Cux2 | 0 | 4.9394937 | L4 |
| 53 | R3hdm2 | 0 | 2.6921525 | L4 |
| 54 | LOC101056001 | 0 | 5.7590213 | L4 |
| 55 | Mical2 | 0 | 2.8863142 | L4 |
| 56 | Atp2b2 | 0 | 2.0552123 | L4 |
| 57 | Tmem145 | 0 | 3.7594233 | L4 |
| 58 | Camk4 | 0 | 4.23498 | L4 |
| 59 | Whrn | 0 | 6.0988703 | L4 |
| 60 | Brinp3 | 0 | 5.6102867 | L4 |
| 61 | Plcxd2 | 0 | 5.406191 | L4 |
| 62 | Cbln4 | 0 | 7.484536 | L4 |
| 63 | Foxp1 | 0 | 3.7569568 | L4 |
| 64 | Atp1a1 | 0 | 2.5461888 | L4 |
| 65 | Rgs4 | 0 | 5.075087 | L4 |
| 66 | Arpp21 | 0 | 5.0060067 | L4 |
| 67 | Camk2n1 | 0 | 3.2389383 | L4 |
| 68 | Rspo1 | 0 | 8.454195 | L4 |
| 69 | Fmnl1 | 0 | 4.506553 | L4 |
| 70 | R3hdm1 | 0 | 3.324493 | L4 |
| 71 | Rorb | 0 | 7.7750397 | L4 |
| 72 | Kalrn | 0 | 3.0100684 | L4 |
| 73 | Mast3 | 0 | 4.3818917 | L4 |
| 74 | Dnajc21 | 0 | 3.6386268 | L4 |
| 75 | S100a10 | 0 | 5.5914707 | L4 |
| 76 | Golga7b | 0 | 2.9620137 | L4 |
| 77 | Camk2a | 0 | 2.9592223 | L4 |
| 78 | Stx1a | 0 | 3.4292345 | L4 |
| 79 | Mef2c | 0 | 1.9808133 | L4 |
| 80 | Nrep | 0 | 3.7079341 | L4 |
| 81 | Pdp1 | 0 | 3.6794658 | L4 |
| 82 | Epha4 | 0 | 4.695211 | L4 |
| 83 | Brinp1 | 0 | 2.9528623 | L4 |
| 84 | Osbp2 | 0 | 3.4083288 | L4 |
| 85 | Slc17a6 | 0 | 6.6784186 | L4 |
| 86 | Tenm4 | 0 | 3.2877388 | L4 |
| 87 | Krt12 | 0 | 5.2697577 | L4 |

| | | | | |
|---|---|---|---|---|
| 88 | Ppp3ca | 0 | 1.9150063 | L4 |
| 89 | Ptprd | 0 | 2.8180037 | L4 |
| 90 | Igfn1 | 0 | 6.5380707 | L4 |
| 91 | Tenm2 | 0 | 3.198212 | L4 |
| 92 | 1110008P14Rik | 0 | 2.7384803 | L4 |
| 93 | Hivep2 | 0 | 2.4533346 | L4 |
| 94 | Cpne9 | 0 | 6.076431 | L4 |
| 95 | Cdh12 | 0 | 4.494685 | L4 |
| 96 | Coch | 0 | 6.1191506 | L4 |
| 97 | Rims3 | 0 | 3.9082186 | L4 |
| 98 | Nudt4 | 0 | 2.846978 | L4 |
| 99 | Crhr1 | 0 | 4.7723737 | L4 |
| 0 | Krt12 | 0 | 5.7243915 | L5 IT |
| 1 | Bok | 0 | 3.9251978 | L5 IT |
| 2 | Lin7a | 0 | 3.3827174 | L5 IT |
| 3 | Ap3m2 | 0 | 2.4024987 | L5 IT |
| 4 | Crym | 0 | 6.276164 | L5 IT |
| 5 | Tagln3 | 0 | 1.3827771 | L5 IT |
| 6 | Mef2c | 0 | 1.856948 | L5 IT |
| 7 | Coch | 0 | 5.0768228 | L5 IT |
| 8 | Hs3st2 | 0 | 5.1099787 | L5 IT |
| 9 | C2cd4b | 0 | 3.952552 | L5 IT |
| 10 | Gfra2 | 0 | 4.3815484 | L5 IT |
| 11 | Basp1 | 0 | 1.4507147 | L5 IT |
| 12 | Tnnc1 | 0 | 6.39366 | L5 IT |
| 13 | Gm20063 | 0 | 4.957448 | L5 IT |
| 14 | Tuba4a | 0 | 1.6647606 | L5 IT |
| 15 | Arpp19 | 0 | 2.4311953 | L5 IT |
| 16 | Cobl | 0 | 4.379489 | L5 IT |
| 17 | Ptn | 0 | 4.9401393 | L5 IT |
| 18 | C730002L08Rik | 0 | 5.036544 | L5 IT |
| 19 | Syn1 | 0 | 1.6258621 | L5 IT |
| 20 | Wnt4 | 0 | 4.650207 | L5 IT |
| 21 | Cd83 | 0 | 5.0434055 | L5 IT |
| 22 | Camkk2 | 0 | 2.6501837 | L5 IT |
| 23 | Slc39a10 | 0 | 2.9200375 | L5 IT |
| 24 | Slc30a3 | 0 | 4.237647 | L5 IT |
| 25 | Tpd52l1 | 0 | 4.0521603 | L5 IT |
| 26 | Agbl4 | 0 | 3.318897 | L5 IT |
| 27 | Ywhah | 0 | 1.2400987 | L5 IT |
| 28 | Pkib | 0 | 4.1466904 | L5 IT |
| 29 | Cck | 0 | 5.3641505 | L5 IT |
| 30 | Adcyap1 | 0 | 5.2326484 | L5 IT |
| 31 | Prdm8 | 0 | 4.305224 | L5 IT |
| 32 | Pip4k2a | 0 | 3.4527526 | L5 IT |
| 33 | Fras1 | 0 | 4.2859206 | L5 IT |
| 34 | Ptgfrn | 0 | 3.9450994 | L5 IT |
| 35 | Boc | 0 | 4.2309275 | L5 IT |
| 36 | Tspan5 | 0 | 1.6272845 | L5 IT |
| 37 | Dusp14 | 0 | 3.4415119 | L5 IT |

| | | | | |
|---|---|---|---|---|
| 38 | Ppp3r1 | 0 | 1.4376225 | L5 IT |
| 39 | Stmn2 | 0 | 1.4744916 | L5 IT |
| 40 | Cdyl2 | 0 | 3.6742482 | L5 IT |
| 41 | Glt8d2 | 0 | 4.519013 | L5 IT |
| 42 | Sv2b | 0 | 5.2111106 | L5 IT |
| 43 | Klf10 | 0 | 4.612071 | L5 IT |
| 44 | Phyhip | 0 | 2.3697724 | L5 IT |
| 45 | Fhod3 | 0 | 3.5231457 | L5 IT |
| 46 | Gm35810 | 0 | 4.5763397 | L5 IT |
| 47 | Lingo1 | 0 | 4.3972692 | L5 IT |
| 48 | Foxp1 | 0 | 3.2743347 | L5 IT |
| 49 | Lhx2 | 0 | 4.5696254 | L5 IT |
| 50 | Shisa4 | 0 | 2.103386 | L5 IT |
| 51 | Adra1d | 0 | 3.6670551 | L5 IT |
| 52 | Cpne9 | 0 | 5.7742205 | L5 IT |
| 53 | Cnih3 | 0 | 4.5507264 | L5 IT |
| 54 | LOC101056001 | 0 | 5.194727 | L5 IT |
| 55 | 6330403A02Rik | 0 | 4.845911 | L5 IT |
| 56 | Rgs4 | 0 | 4.7029104 | L5 IT |
| 57 | Camk4 | 0 | 3.9791508 | L5 IT |
| 58 | Grm2 | 0 | 5.2631125 | L5 IT |
| 59 | Pamr1 | 0 | 6.165286 | L5 IT |
| 60 | Nrn1 | 0 | 6.431351 | L5 IT |
| 61 | Rtn1 | 0 | 1.2429998 | L5 IT |
| 62 | Snca | 0 | 2.7934797 | L5 IT |
| 63 | S100a10 | 0 | 5.689857 | L5 IT |
| 64 | Ly6e | 0 | 2.0487533 | L5 IT |
| 65 | Hpca | 0 | 3.0631597 | L5 IT |
| 66 | Tcrb | 0 | 5.689356 | L5 IT |
| 67 | Scube1 | 0 | 5.435196 | L5 IT |
| 68 | Lmo4 | 0 | 3.7198443 | L5 IT |
| 69 | Neurod6 | 0 | 6.6778097 | L5 IT |
| 70 | Stx1a | 0 | 3.6808667 | L5 IT |
| 71 | Dkkl1 | 0 | 6.7894993 | L5 IT |
| 72 | Itpka | 0 | 5.330865 | L5 IT |
| 73 | 05-Sep | 0 | 1.8769021 | L5 IT |
| 74 | Kcnv1 | 0 | 5.377618 | L5 IT |
| 75 | Sh3gl2 | 0 | 2.319423 | L5 IT |
| 76 | Tusc3 | 0 | 1.523156 | L5 IT |
| 77 | Cdk14 | 0 | 2.9942768 | L5 IT |
| 78 | Ovol2 | 0 | 4.6264405 | L5 IT |
| 79 | Cfl1 | 0 | 0.6263922 | L5 IT |
| 80 | Pgm2l1 | 0 | 2.319 | L5 IT |
| 81 | Stmn1 | 0 | 1.5270116 | L5 IT |
| 82 | Fat3 | 0 | 4.377044 | L5 IT |
| 83 | Osbpl1a | 0 | 2.3789344 | L5 IT |
| 84 | Dkk3 | 0 | 4.723692 | L5 IT |
| 85 | Grm3 | 0 | 4.1872067 | L5 IT |
| 86 | Gm13601 | 0 | 3.9235494 | L5 IT |
| 87 | Lhfp | 0 | 4.6729627 | L5 IT |

| | | | | |
|---|---|---|---|---|
| 88 | Tspan17 | 0 | 3.5716677 | L5 IT |
| 89 | Car10 | 0 | 4.1533856 | L5 IT |
| 90 | Lin7b | 0 | 1.975826 | L5 IT |
| 91 | Rtn4r | 0 | 4.6767817 | L5 IT |
| 92 | Kcnk4 | 0 | 4.08046 | L5 IT |
| 93 | Fam212b | 0 | 4.812341 | L5 IT |
| 94 | Epha4 | 0 | 4.5003014 | L5 IT |
| 95 | Slc17a7 | 0 | 5.9017243 | L5 IT |
| 96 | Myl4 | 0 | 6.048156 | L5 IT |
| 97 | Rorb | 0 | 5.732735 | L5 IT |
| 98 | Dnajc21 | 0 | 3.1799765 | L5 IT |
| 99 | St6galnac5 | 0 | 4.2279396 | L5 IT |
| 0 | Fam19a1 | 0 | 8.179921 | L5 PT |
| 1 | Chn1 | 0 | 1.9461819 | L5 PT |
| 2 | Sdhb | 0 | 0.8832347 | L5 PT |
| 3 | Nkiras1 | 0 | 1.198404 | L5 PT |
| 4 | Atp5g1 | 0 | 1.0873361 | L5 PT |
| 5 | Dtnbp1 | 0 | 2.639751 | L5 PT |
| 6 | Adcyap1 | 0 | 6.045408 | L5 PT |
| 7 | Kcnn2 | 0 | 3.7665036 | L5 PT |
| 8 | Gm34583 | 0 | 6.63009 | L5 PT |
| 9 | Kcng1 | 0 | 5.782039 | L5 PT |
| 10 | Bcl11b | 0 | 4.3903313 | L5 PT |
| 11 | Tox | 0 | 4.7023616 | L5 PT |
| 12 | Gng13 | 0 | 2.0725756 | L5 PT |
| 13 | Mal2 | 0 | 2.2218575 | L5 PT |
| 14 | Bhlhe40 | 0 | 5.1338964 | L5 PT |
| 15 | Serpine2 | 0 | 6.1090016 | L5 PT |
| 16 | Meis2 | 0 | 5.74133 | L5 PT |
| 17 | Mgst3 | 0 | 1.8598229 | L5 PT |
| 18 | Bcl6 | 0 | 5.020265 | L5 PT |
| 19 | Ckmt1 | 0 | 2.1118593 | L5 PT |
| 20 | Gm19410 | 0 | 6.721095 | L5 PT |
| 21 | Fezf2 | 0 | 7.1948767 | L5 PT |
| 22 | Npr3 | 0 | 7.257237 | L5 PT |
| 23 | Chst8 | 0 | 6.4727526 | L5 PT |
| 24 | Fam84b | 0 | 7.228047 | L5 PT |
| 25 | Pou3f1 | 0 | 7.158306 | L5 PT |
| 26 | Tmem163 | 0 | 6.8848367 | L5 PT |
| 27 | Gmpr | 0 | 2.9815776 | L5 PT |
| 28 | Gm2164 | 0 | 7.296124 | L5 PT |
| 29 | Prkcg | 0 | 4.1823215 | L5 PT |
| 30 | Mpc2 | 0 | 1.3371704 | L5 PT |
| 31 | Pex5l | 0 | 3.962546 | L5 PT |
| 32 | Gap43 | 0 | 2.9738731 | L5 PT |
| 33 | Arhgdig | 1.54E-307 | 1.8868484 | L5 PT |
| 34 | Pcsk5 | 3.12E-306 | 4.9617243 | L5 PT |
| 35 | Bend5 | 9.14E-306 | 4.01695 | L5 PT |
| 36 | Lamp5 | 1.21E-304 | 7.066248 | L5 PT |
| 37 | Gnb5 | 2.27E-304 | 1.7226558 | L5 PT |

| | | | | |
|---|---|---|---|---|
| 38 | Polr2h | 5.09E-304 | 2.1912322 | L5 PT |
| 39 | Atp2b1 | 2.59E-301 | 1.935655 | L5 PT |
| 40 | Sdcbp | 2.90E-299 | 1.6050088 | L5 PT |
| 41 | Cacna1h | 2.62E-297 | 4.981756 | L5 PT |
| 42 | Gm36916 | 2.65E-295 | 3.8697371 | L5 PT |
| 43 | Uqcr11 | 2.79E-293 | 0.90922695 | L5 PT |
| 44 | Ldb2 | 1.54E-292 | 4.33865 | L5 PT |
| 45 | Fam131a | 4.54E-290 | 2.691679 | L5 PT |
| 46 | Sec14l1 | 1.61E-289 | 2.5225468 | L5 PT |
| 47 | Slc8a1 | 1.66E-285 | 2.489418 | L5 PT |
| 48 | Enc1 | 2.72E-285 | 3.8083308 | L5 PT |
| 49 | Vat1l | 1.60E-284 | 4.199891 | L5 PT |
| 50 | Sh3bgrl3 | 4.21E-284 | 1.5097852 | L5 PT |
| 51 | Sorl1 | 7.78E-284 | 3.0785272 | L5 PT |
| 52 | Atp5k | 5.04E-280 | 0.9991339 | L5 PT |
| 53 | Rapgef5 | 9.51E-280 | 3.6207097 | L5 PT |
| 54 | Ccl27a | 2.73E-279 | 1.8122642 | L5 PT |
| 55 | Ldhb | 7.05E-279 | 1.0438561 | L5 PT |
| 56 | Cdh22 | 4.48E-278 | 4.367546 | L5 PT |
| 57 | Tspan13 | 1.66E-277 | 1.5423205 | L5 PT |
| 58 | Nos1ap | 1.75E-277 | 2.8060083 | L5 PT |
| 59 | Etv1 | 3.36E-277 | 4.997356 | L5 PT |
| 60 | Etv5 | 3.84E-276 | 2.8745055 | L5 PT |
| 61 | Galnt16 | 8.77E-276 | 3.744744 | L5 PT |
| 62 | Ntng1 | 1.64E-275 | 4.6048975 | L5 PT |
| 63 | Atp5o | 2.44E-274 | 0.9197249 | L5 PT |
| 64 | 1110008P14Rik | 7.80E-272 | 2.453411 | L5 PT |
| 65 | Uqcr10 | 5.86E-271 | 0.932837 | L5 PT |
| 66 | Tubb3 | 1.72E-269 | 1.5901026 | L5 PT |
| 67 | Atp5j2 | 4.72E-269 | 0.95782936 | L5 PT |
| 68 | Nefm | 5.05E-269 | 4.510323 | L5 PT |
| 69 | Rgma | 1.85E-266 | 3.8200192 | L5 PT |
| 70 | Tekt5 | 1.21E-264 | 5.485729 | L5 PT |
| 71 | Slc5a5 | 2.77E-263 | 4.804499 | L5 PT |
| 72 | Atp5f1 | 6.25E-261 | 0.7888438 | L5 PT |
| 73 | Ryr3 | 2.34E-260 | 4.2970862 | L5 PT |
| 74 | Parm1 | 2.98E-260 | 5.0248175 | L5 PT |
| 75 | Mas1 | 7.51E-259 | 5.1940436 | L5 PT |
| 76 | Sgpp2 | 5.54E-257 | 4.6504393 | L5 PT |
| 77 | Esd | 3.76E-255 | 1.22201 | L5 PT |
| 78 | Usmg5 | 2.13E-254 | 0.7756476 | L5 PT |
| 79 | 2410015M20Rik | 1.48E-251 | 1.0286607 | L5 PT |
| 80 | Pigp | 3.00E-251 | 2.0079439 | L5 PT |
| 81 | Efr3a | 3.24E-251 | 1.797701 | L5 PT |
| 82 | Cds1 | 1.16E-250 | 2.5972211 | L5 PT |
| 83 | Tiam2 | 1.89E-250 | 4.0079107 | L5 PT |
| 84 | Calm2 | 3.10E-250 | 0.9372546 | L5 PT |
| 85 | Cox6c | 4.01E-250 | 0.7444574 | L5 PT |
| 86 | Gpr123 | 1.25E-248 | 2.445009 | L5 PT |
| 87 | Nme1 | 1.25E-248 | 0.99794704 | L5 PT |

| | | | | |
|---|---|---|---|---|
| 88 | Tmem150c | 1.28E-248 | 3.8749125 | L5 PT |
| 89 | Hspa12a | 7.51E-248 | 2.9073517 | L5 PT |
| 90 | Ndufa3 | 1.14E-247 | 0.9015308 | L5 PT |
| 91 | Ppm1e | 2.09E-247 | 2.12832 | L5 PT |
| 92 | Ina | 1.90E-246 | 2.286095 | L5 PT |
| 93 | Ndufab1 | 1.48E-245 | 0.8289623 | L5 PT |
| 94 | Timm17a | 3.09E-244 | 0.967851 | L5 PT |
| 95 | Atox1 | 3.24E-243 | 1.1584026 | L5 PT |
| 96 | Scn4b | 2.66E-242 | 4.91411 | L5 PT |
| 97 | Pop5 | 1.94E-241 | 1.5021503 | L5 PT |
| 98 | 2010107E04Rik | 2.98E-241 | 0.9121919 | L5 PT |
| 99 | Gm32960 | 1.62E-239 | 5.821919 | L5 PT |
| 0 | Foxp2 | 0 | 9.379705 | L6 CT |
| 1 | Dynll1 | 0 | 1.3117678 | L6 CT |
| 2 | Emb | 0 | 3.509111 | L6 CT |
| 3 | Adcy1 | 0 | 2.3691905 | L6 CT |
| 4 | Cdh18 | 0 | 4.2715197 | L6 CT |
| 5 | Jup | 0 | 4.1819086 | L6 CT |
| 6 | Ncald | 0 | 2.0134807 | L6 CT |
| 7 | Adora1 | 0 | 4.061856 | L6 CT |
| 8 | Pde1a | 0 | 4.916502 | L6 CT |
| 9 | Ldha | 0 | 1.598549 | L6 CT |
| 10 | Mctp1 | 0 | 3.8973157 | L6 CT |
| 11 | Fxyd7 | 0 | 3.3634088 | L6 CT |
| 12 | Fut9 | 0 | 2.6780188 | L6 CT |
| 13 | Chgb | 0 | 2.0301158 | L6 CT |
| 14 | Sez6 | 0 | 3.7233942 | L6 CT |
| 15 | Baiap2 | 0 | 3.9522777 | L6 CT |
| 16 | Cpe | 0 | 1.0954452 | L6 CT |
| 17 | Ina | 0 | 2.4487517 | L6 CT |
| 18 | Galnt9 | 0 | 4.0155363 | L6 CT |
| 19 | Npdc1 | 0 | 1.1766893 | L6 CT |
| 20 | Rprml | 0 | 4.328545 | L6 CT |
| 21 | Hpcal4 | 0 | 2.7622604 | L6 CT |
| 22 | Diras2 | 0 | 3.584083 | L6 CT |
| 23 | Ptk2 | 0 | 2.8465157 | L6 CT |
| 24 | Htr1f | 0 | 4.2285647 | L6 CT |
| 25 | Gm26974 | 0 | 4.5712614 | L6 CT |
| 26 | Arpc3 | 0 | 1.1529547 | L6 CT |
| 27 | Gm13306 | 0 | 2.3223572 | L6 CT |
| 28 | Rasgrp1 | 0 | 4.2790103 | L6 CT |
| 29 | Itm2c | 0 | 1.2856473 | L6 CT |
| 30 | Kcnk1 | 0 | 1.8757974 | L6 CT |
| 31 | Pdlim1 | 0 | 5.2404976 | L6 CT |
| 32 | Klf9 | 0 | 2.2471018 | L6 CT |
| 33 | Il11ra1 | 0 | 4.1668506 | L6 CT |
| 34 | Igh | 0 | 4.1067305 | L6 CT |
| 35 | Ddah1 | 0 | 3.980261 | L6 CT |
| 36 | Slc1a2 | 0 | 3.5040083 | L6 CT |
| 37 | Ier5 | 0 | 3.8872993 | L6 CT |

| | | | | |
|---|---|---|---|---|
| 38 | Ptk2b | 0 | 4.661718 | L6 CT |
| 39 | Serinc2 | 0 | 4.6719913 | L6 CT |
| 40 | Brk1 | 0 | 1.1142737 | L6 CT |
| 41 | Nfib | 0 | 3.4670577 | L6 CT |
| 42 | Kcnip3 | 0 | 4.161297 | L6 CT |
| 43 | Grik3 | 0 | 4.1864367 | L6 CT |
| 44 | Dkk3 | 0 | 4.4613976 | L6 CT |
| 45 | Necab3 | 0 | 3.4269803 | L6 CT |
| 46 | Spsb1 | 0 | 4.779257 | L6 CT |
| 47 | Ogfrl1 | 0 | 1.7592517 | L6 CT |
| 48 | B3galt2 | 0 | 3.8195722 | L6 CT |
| 49 | Tuba4a | 0 | 1.74275 | L6 CT |
| 50 | Efhd2 | 0 | 2.0660577 | L6 CT |
| 51 | Nrp1 | 0 | 5.6218467 | L6 CT |
| 52 | Nptx1 | 0 | 5.3210316 | L6 CT |
| 53 | Ipcef1 | 0 | 5.14583 | L6 CT |
| 54 | Nos1ap | 0 | 3.3145688 | L6 CT |
| 55 | Col5a1 | 0 | 6.9928083 | L6 CT |
| 56 | Nxph3 | 0 | 5.929796 | L6 CT |
| 57 | Pcp4 | 0 | 6.2209716 | L6 CT |
| 58 | Ramp3 | 0 | 6.000732 | L6 CT |
| 59 | Ccl27a | 0 | 2.2292912 | L6 CT |
| 60 | Crym | 0 | 7.790106 | L6 CT |
| 61 | Rab26 | 0 | 4.6579103 | L6 CT |
| 62 | Syt6 | 0 | 7.6178594 | L6 CT |
| 63 | Garnl3 | 0 | 4.108695 | L6 CT |
| 64 | Tle4 | 0 | 5.488406 | L6 CT |
| 65 | Gadd45a | 0 | 5.9624143 | L6 CT |
| 66 | Prkcb | 0 | 2.4766467 | L6 CT |
| 67 | Ttc9b | 0 | 3.014966 | L6 CT |
| 68 | Hs3st4 | 0 | 6.6250114 | L6 CT |
| 69 | Tcrb | 0 | 6.744749 | L6 CT |
| 70 | 3110035E14Rik | 0 | 6.407568 | L6 CT |
| 71 | Rprm | 0 | 7.976703 | L6 CT |
| 72 | Me3 | 0 | 3.6158822 | L6 CT |
| 73 | Arhgap25 | 0 | 6.6852317 | L6 CT |
| 74 | Tbr1 | 0 | 5.0643225 | L6 CT |
| 75 | Scg3 | 0 | 2.1730123 | L6 CT |
| 76 | Lpgat1 | 0 | 2.1240664 | L6 CT |
| 77 | Rtn4 | 0 | 1.8306766 | L6 CT |
| 78 | Ephb1 | 0 | 4.4187236 | L6 CT |
| 79 | Fezf2 | 0 | 6.039657 | L6 CT |
| 80 | Hs3st2 | 0 | 5.3649516 | L6 CT |
| 81 | Nfia | 0 | 4.7876163 | L6 CT |
| 82 | Igsf21 | 0 | 5.048937 | L6 CT |
| 83 | Prkcg | 0 | 4.045952 | L6 CT |
| 84 | H2-T23 | 0 | 5.456293 | L6 CT |
| 85 | Ano3 | 0 | 4.5945497 | L6 CT |
| 86 | Gm30512 | 0 | 5.080966 | L6 CT |
| 87 | Atp2b1 | 0 | 2.005065 | L6 CT |

| | | | | |
|---|---|---|---|---|
| 88 | Ppp1r1b | 0 | 5.134635 | L6 CT |
| 89 | 07-Sep | 0 | 1.0476378 | L6 CT |
| 90 | Sptbn5 | 0 | 5.6964173 | L6 CT |
| 91 | Igfbp4 | 0 | 5.5549483 | L6 CT |
| 92 | Gm30620 | 0 | 9.283937 | L6 CT |
| 93 | Med10 | 0 | 2.10052 | L6 CT |
| 94 | Zfpm2 | 0 | 6.110513 | L6 CT |
| 95 | Gng12 | 0 | 5.454433 | L6 CT |
| 96 | Rell1 | 0 | 5.3412485 | L6 CT |
| 97 | Ctxn1 | 0 | 3.7097058 | L6 CT |
| 98 | Cyth2 | 0 | 1.9642112 | L6 CT |
| 99 | Mmp17 | 0 | 3.6023197 | L6 CT |
| 0 | Tesc | 0 | 6.1208262 | L6 IT |
| 1 | Adamts3 | 0 | 4.040233 | L6 IT |
| 2 | Mapk11 | 0 | 3.831221 | L6 IT |
| 3 | Satb2 | 0 | 4.322363 | L6 IT |
| 4 | 9130024F11Rik | 0 | 3.4473708 | L6 IT |
| 5 | Sv2b | 0 | 5.078162 | L6 IT |
| 6 | Blnk | 0 | 5.624689 | L6 IT |
| 7 | Slit3 | 0 | 3.3385887 | L6 IT |
| 8 | Crtac1 | 0 | 3.942713 | L6 IT |
| 9 | Mmp17 | 0 | 3.6437721 | L6 IT |
| 10 | Neurod6 | 0 | 5.921069 | L6 IT |
| 11 | Hpca | 0 | 2.6547372 | L6 IT |
| 12 | B3galt2 | 0 | 3.4987726 | L6 IT |
| 13 | Ppp1r1a | 0 | 2.0984201 | L6 IT |
| 14 | Cacna1e | 0 | 3.0580547 | L6 IT |
| 15 | 2010111I01Rik | 0 | 3.1688678 | L6 IT |
| 16 | Gm11549 | 0 | 5.048822 | L6 IT |
| 17 | Cpne5 | 0 | 4.104628 | L6 IT |
| 18 | Rasgrp1 | 0 | 4.427593 | L6 IT |
| 19 | Nptx1 | 0 | 4.5433764 | L6 IT |
| 20 | Ldb2 | 0 | 4.280329 | L6 IT |
| 21 | 3110035E14Rik | 0 | 5.2089276 | L6 IT |
| 22 | H2-T23 | 0 | 4.4569592 | L6 IT |
| 23 | Hs3st2 | 0 | 4.9428334 | L6 IT |
| 24 | 02-Mar | 0 | 2.3273535 | L6 IT |
| 25 | Sema3e | 0 | 4.5151668 | L6 IT |
| 26 | Osr1 | 0 | 7.8588915 | L6 IT |
| 27 | Pfkl | 0 | 2.4774587 | L6 IT |
| 28 | Sstr2 | 0 | 4.3258686 | L6 IT |
| 29 | Sorcs3 | 0 | 4.0051765 | L6 IT |
| 30 | Gnb4 | 0 | 4.0552416 | L6 IT |
| 31 | Sdk2 | 0 | 3.5887873 | L6 IT |
| 32 | Fmnl1 | 0 | 3.358501 | L6 IT |
| 33 | Nnat | 0 | 5.2314672 | L6 IT |
| 34 | Pdia5 | 0 | 4.683288 | L6 IT |
| 35 | Itm2c | 0 | 1.2034081 | L6 IT |
| 36 | Ssbp3 | 0 | 1.8252546 | L6 IT |
| 37 | Nrn1 | 0 | 5.792523 | L6 IT |

| | | | | |
|---|---|---|---|---|
| 38 | Extl1 | 0 | 3.3006723 | L6 IT |
| 39 | Fam105a | 0 | 3.7437384 | L6 IT |
| 40 | Pfkp | 0 | 1.5028409 | L6 IT |
| 41 | Pdrg1 | 0 | 2.0444534 | L6 IT |
| 42 | Baiap2 | 0 | 3.8698044 | L6 IT |
| 43 | Mapk1 | 0 | 1.1102738 | L6 IT |
| 44 | Tenm3 | 0 | 3.5690985 | L6 IT |
| 45 | Gpm6b | 0 | 2.0690908 | L6 IT |
| 46 | Rhou | 0 | 3.5773275 | L6 IT |
| 47 | Olfm1 | 0 | 1.9760149 | L6 IT |
| 48 | Ak4 | 0 | 4.0738316 | L6 IT |
| 49 | Mgat5 | 0 | 3.4472444 | L6 IT |
| 50 | Gpr22 | 0 | 2.612762 | L6 IT |
| 51 | Pcsk2 | 0 | 3.0218647 | L6 IT |
| 52 | Dok5 | 0 | 3.6801758 | L6 IT |
| 53 | Rtn4rl2 | 0 | 5.0672994 | L6 IT |
| 54 | Atp2b4 | 0 | 5.1375203 | L6 IT |
| 55 | Col6a1 | 0 | 6.635112 | L6 IT |
| 56 | Tpm1 | 0 | 1.7576485 | L6 IT |
| 57 | Ociad2 | 0 | 2.9532294 | L6 IT |
| 58 | Igfbp6 | 0 | 6.0626073 | L6 IT |
| 59 | C1ql3 | 0 | 5.4075747 | L6 IT |
| 60 | Hpcal4 | 0 | 2.9830842 | L6 IT |
| 61 | Nell2 | 0 | 3.7136545 | L6 IT |
| 62 | Slc26a4 | 0 | 6.589594 | L6 IT |
| 63 | Eno1 | 0 | 1.2484723 | L6 IT |
| 64 | Pter | 0 | 4.9033604 | L6 IT |
| 65 | Ak5 | 0 | 4.035195 | L6 IT |
| 66 | Bmp3 | 0 | 6.3776517 | L6 IT |
| 67 | Igsf21 | 0 | 5.6590915 | L6 IT |
| 68 | Schip1 | 0 | 2.8428905 | L6 IT |
| 69 | Cck | 0 | 5.9251776 | L6 IT |
| 70 | Gm2694 | 0 | 6.3003044 | L6 IT |
| 71 | Rasl10a | 0 | 6.9578934 | L6 IT |
| 72 | Eno1b | 0 | 1.3412758 | L6 IT |
| 73 | Enc1 | 0 | 3.9357803 | L6 IT |
| 74 | Fhl2 | 0 | 4.1639028 | L6 IT |
| 75 | Ensa | 0 | 1.362688 | L6 IT |
| 76 | Ttc9b | 0 | 2.2398467 | L6 IT |
| 77 | Snca | 0 | 2.4946074 | L6 IT |
| 78 | Fkbp1a | 0 | 0.9672349 | L6 IT |
| 79 | Mas1 | 0 | 5.2893567 | L6 IT |
| 80 | Pdlim1 | 0 | 5.331043 | L6 IT |
| 81 | Rtn4 | 0 | 1.7445936 | L6 IT |
| 82 | Rtn4r | 0 | 4.4079137 | L6 IT |
| 83 | Lmo4 | 0 | 3.2928834 | L6 IT |
| 84 | Slc17a7 | 0 | 5.648625 | L6 IT |
| 85 | Lingo1 | 0 | 4.5350876 | L6 IT |
| 86 | Car12 | 0 | 5.376652 | L6 IT |
| 87 | Brk1 | 0 | 1.1340504 | L6 IT |

| | | | | |
|---|---|---|---|---|
| 88 | Cdh9 | 0 | 5.09092 | L6 IT |
| 89 | Dynll1 | 0 | 1.3295848 | L6 IT |
| 90 | Gm4735 | 0 | 1.1840237 | L6 IT |
| 91 | Aldoa | 0 | 0.7204515 | L6 IT |
| 92 | Fancd2 | 0 | 5.199032 | L6 IT |
| 93 | Atp2b1 | 0 | 1.8819344 | L6 IT |
| 94 | Ldha | 0 | 1.6472403 | L6 IT |
| 95 | Sh3bgrl3 | 0 | 1.5792315 | L6 IT |
| 96 | Slc7a4 | 0 | 3.2984042 | L6 IT |
| 97 | Ctxn1 | 0 | 3.6695821 | L6 IT |
| 98 | Necab3 | 0 | 3.544079 | L6 IT |
| 99 | Otub2 | 0 | 3.406186 | L6 IT |
| 0 | Ctgf | 1.83E-287 | 10.33918 | L6b |
| 1 | Nxph4 | 5.56E-283 | 11.166246 | L6b |
| 2 | Cplx3 | 1.20E-282 | 10.459437 | L6b |
| 3 | Nxph3 | 2.57E-260 | 6.9823146 | L6b |
| 4 | Phyhipl | 1.51E-257 | 2.069336 | L6b |
| 5 | Tmem163 | 1.37E-252 | 8.019105 | L6b |
| 6 | Gng12 | 8.79E-252 | 6.3956776 | L6b |
| 7 | LOC105246064 | 4.55E-245 | 7.1870136 | L6b |
| 8 | Atp6ap2 | 2.06E-243 | 1.4525133 | L6b |
| 9 | Fam163b | 3.73E-242 | 3.9200058 | L6b |
| 10 | Gap43 | 3.53E-241 | 3.0046482 | L6b |
| 11 | Hs3st4 | 4.07E-239 | 6.266187 | L6b |
| 12 | Cnih2 | 1.24E-235 | 2.1360304 | L6b |
| 13 | 3110035E14Rik | 9.19E-234 | 6.0593233 | L6b |
| 14 | Tle4 | 1.53E-233 | 5.41477 | L6b |
| 15 | Rab26 | 4.58E-232 | 4.642221 | L6b |
| 16 | Kcnip3 | 1.35E-228 | 4.97003 | L6b |
| 17 | Ttc9b | 2.31E-223 | 2.847743 | L6b |
| 18 | 2900011O08Rik | 1.35E-222 | 1.8361003 | L6b |
| 19 | Kcnmb4 | 7.25E-222 | 3.4628475 | L6b |
| 20 | Ly6g6e | 2.77E-220 | 8.577792 | L6b |
| 21 | Pcp4 | 1.24E-218 | 6.112787 | L6b |
| 22 | Pcsk5 | 1.68E-217 | 5.997608 | L6b |
| 23 | 07-Sep | 1.37E-216 | 1.1950865 | L6b |
| 24 | Prss12 | 1.51E-216 | 6.0434775 | L6b |
| 25 | Chga | 2.15E-216 | 2.2089782 | L6b |
| 26 | Pak1 | 1.96E-215 | 2.4266403 | L6b |
| 27 | Cd164 | 4.46E-212 | 3.6444852 | L6b |
| 28 | Slc8a1 | 2.58E-210 | 2.8742568 | L6b |
| 29 | Rai14 | 6.10E-210 | 6.0957212 | L6b |
| 30 | Fut9 | 5.63E-206 | 2.969872 | L6b |
| 31 | Slc1a2 | 5.83E-205 | 4.1018724 | L6b |
| 32 | Drd1 | 1.67E-202 | 5.767463 | L6b |
| 33 | Gm32392 | 2.07E-201 | 6.4198976 | L6b |
| 34 | Rogdi | 2.69E-201 | 1.5043888 | L6b |
| 35 | Bag1 | 6.42E-201 | 2.010969 | L6b |
| 36 | Cpe | 2.60E-200 | 1.43913 | L6b |
| 37 | Clic5 | 8.27E-200 | 8.17085 | L6b |

| | | | | |
|---|---|---|---|---|
| 38 | Serpini1 | 3.95E-193 | 3.017613 | L6b |
| 39 | Lhfpl3 | 6.30E-193 | 5.702078 | L6b |
| 40 | Cfl1 | 1.59E-192 | 0.7692349 | L6b |
| 41 | Mpi | 2.98E-192 | 2.9821131 | L6b |
| 42 | Inpp4b | 9.31E-192 | 6.351055 | L6b |
| 43 | Gadd45a | 1.25E-190 | 5.2914186 | L6b |
| 44 | Cdh18 | 2.04E-189 | 4.9803224 | L6b |
| 45 | Srcin1 | 1.14E-188 | 2.0918968 | L6b |
| 46 | D630023F18Rik | 1.87E-187 | 5.1020885 | L6b |
| 47 | Atp6ap1l | 7.56E-187 | 5.6131215 | L6b |
| 48 | Ndrg3 | 1.23E-186 | 1.4755342 | L6b |
| 49 | St3gal5 | 2.90E-186 | 2.8061576 | L6b |
| 50 | Tubb2a | 1.54E-185 | 1.1961024 | L6b |
| 51 | Gfra4 | 1.85E-184 | 2.0443437 | L6b |
| 52 | Usp46 | 2.03E-184 | 3.319652 | L6b |
| 53 | Fxyd7 | 5.68E-183 | 3.4182987 | L6b |
| 54 | Lman1l | 2.75E-182 | 6.473178 | L6b |
| 55 | Atp2b4 | 1.40E-180 | 5.066171 | L6b |
| 56 | Sdk2 | 1.56E-180 | 4.572278 | L6b |
| 57 | Tuba1a | 2.15E-179 | 0.9896737 | L6b |
| 58 | Pde1a | 5.25E-179 | 5.0890102 | L6b |
| 59 | Cyth2 | 2.44E-178 | 1.9724209 | L6b |
| 60 | Pls3 | 2.73E-178 | 2.6084235 | L6b |
| 61 | Hpcal4 | 5.34E-177 | 2.8664184 | L6b |
| 62 | Gabra5 | 9.32E-177 | 4.7720757 | L6b |
| 63 | 1700001L19Rik | 5.00E-175 | 4.126069 | L6b |
| 64 | Hpcal1 | 1.72E-173 | 5.0516133 | L6b |
| 65 | Aldh1b1 | 6.50E-173 | 5.0308127 | L6b |
| 66 | Dtnb | 4.73E-172 | 2.9839525 | L6b |
| 67 | Caln1 | 7.10E-170 | 3.8744864 | L6b |
| 68 | Jup | 9.15E-170 | 4.480842 | L6b |
| 69 | Tmsb10 | 5.70E-169 | 2.2987645 | L6b |
| 70 | Pgrmc1 | 1.79E-168 | 1.2796758 | L6b |
| 71 | Mctp1 | 9.88E-167 | 4.047938 | L6b |
| 72 | Tmem158 | 5.69E-166 | 3.1896253 | L6b |
| 73 | Oprm1 | 6.00E-165 | 4.856179 | L6b |
| 74 | Nptx1 | 4.95E-164 | 4.8227406 | L6b |
| 75 | Brk1 | 5.60E-164 | 1.2833524 | L6b |
| 76 | Pbx1 | 9.06E-164 | 2.710188 | L6b |
| 77 | Ssbp3 | 1.69E-162 | 2.0758405 | L6b |
| 78 | Tmem40 | 2.96E-162 | 6.642143 | L6b |
| 79 | Tusc3 | 1.88E-161 | 1.5488231 | L6b |
| 80 | Lmo3 | 7.45E-161 | 3.835145 | L6b |
| 81 | Trp53i11 | 1.17E-160 | 5.1038785 | L6b |
| 82 | Celf4 | 2.16E-160 | 1.4070563 | L6b |
| 83 | Ipcef1 | 2.32E-160 | 4.5874104 | L6b |
| 84 | Itm2c | 5.97E-160 | 1.3733029 | L6b |
| 85 | Rgs20 | 1.27E-159 | 3.8017626 | L6b |
| 86 | Svil | 1.89E-159 | 5.3252506 | L6b |
| 87 | Sulf1 | 2.93E-159 | 5.681577 | L6b |

| | | | | |
|---|---|---|---|---|
| 88 | Igsf21 | 4.71E-159 | 4.8164916 | L6b |
| 89 | Zfpm2 | 1.13E-158 | 5.16159 | L6b |
| 90 | Hsbp1 | 8.72E-158 | 0.777333 | L6b |
| 91 | Sez6 | 2.58E-157 | 3.67526 | L6b |
| 92 | Cyb5a | 1.55E-156 | 1.6946323 | L6b |
| 93 | Mmp16 | 4.73E-156 | 3.5931072 | L6b |
| 94 | Bcl11a | 6.29E-156 | 2.5773888 | L6b |
| 95 | Gm9844 | 1.19E-155 | 2.1978152 | L6b |
| 96 | Actg1 | 3.63E-155 | 0.8750731 | L6b |
| 97 | Tmod1 | 3.66E-155 | 3.8351865 | L6b |
| 98 | Mllt11 | 1.24E-153 | 1.2962507 | L6b |
| 99 | Sh3gl3 | 9.99E-153 | 3.3969994 | L6b |
| 0 | Dner | 0 | 5.272085 | Lamp5 |
| 1 | Eepd1 | 0 | 4.1090403 | Lamp5 |
| 2 | Cntnap2 | 0 | 3.0648522 | Lamp5 |
| 3 | Tsc22d1 | 0 | 1.7556598 | Lamp5 |
| 4 | Mapk3 | 0 | 3.2432492 | Lamp5 |
| 5 | Nr2f2 | 0 | 5.2680745 | Lamp5 |
| 6 | Rgs10 | 0 | 4.886741 | Lamp5 |
| 7 | Rora | 0 | 3.1849182 | Lamp5 |
| 8 | Ogfrl1 | 0 | 2.22817 | Lamp5 |
| 9 | Frmpd4 | 0 | 2.7433474 | Lamp5 |
| 10 | Mgll | 0 | 2.3202186 | Lamp5 |
| 11 | Spock1 | 0 | 2.475845 | Lamp5 |
| 12 | Rapgef4 | 0 | 1.9703357 | Lamp5 |
| 13 | Sv2c | 0 | 5.7651377 | Lamp5 |
| 14 | Nfix | 0 | 2.9296556 | Lamp5 |
| 15 | Afap1 | 0 | 4.705707 | Lamp5 |
| 16 | Ptchd2 | 0 | 4.894961 | Lamp5 |
| 17 | Tmeff1 | 0 | 2.9068108 | Lamp5 |
| 18 | Erbb4 | 0 | 6.342992 | Lamp5 |
| 19 | Sema5a | 0 | 5.687699 | Lamp5 |
| 20 | Kcnn2 | 0 | 3.63033 | Lamp5 |
| 21 | Thrsp | 0 | 5.371114 | Lamp5 |
| 22 | Sh3bgrl | 0 | 2.995951 | Lamp5 |
| 23 | Crtac1 | 0 | 4.2547126 | Lamp5 |
| 24 | Homer2 | 0 | 3.8209548 | Lamp5 |
| 25 | Pcbp3 | 0 | 2.2451437 | Lamp5 |
| 26 | Zcchc18 | 0 | 1.2529839 | Lamp5 |
| 27 | Rimbp2 | 0 | 2.9786365 | Lamp5 |
| 28 | Tox3 | 0 | 4.317045 | Lamp5 |
| 29 | Chst7 | 0 | 6.933304 | Lamp5 |
| 30 | Igf1 | 0 | 5.4196444 | Lamp5 |
| 31 | Serpini1 | 0 | 2.53051 | Lamp5 |
| 32 | Plekho1 | 0 | 2.5839374 | Lamp5 |
| 33 | Slc32a1 | 0 | 4.398383 | Lamp5 |
| 34 | Ppp1r2 | 0 | 1.5437949 | Lamp5 |
| 35 | Necab2 | 0 | 4.657628 | Lamp5 |
| 36 | Lancl1 | 0 | 1.253069 | Lamp5 |
| 37 | Ptpro | 0 | 4.377507 | Lamp5 |

| | | | | |
|---|---|---|---|---|
| 38 | Fmo1 | 0 | 7.507155 | Lamp5 |
| 39 | Maf | 0 | 4.352202 | Lamp5 |
| 40 | Slc25a17 | 0 | 2.1052465 | Lamp5 |
| 41 | Maged1 | 0 | 1.3603541 | Lamp5 |
| 42 | Slc39a6 | 0 | 3.461035 | Lamp5 |
| 43 | Reln | 0 | 6.2250977 | Lamp5 |
| 44 | Nmnat2 | 0 | 1.7272093 | Lamp5 |
| 45 | Prox1 | 0 | 5.1011515 | Lamp5 |
| 46 | Slc2a13 | 0 | 2.8048635 | Lamp5 |
| 47 | Sox2ot | 0 | 4.9074597 | Lamp5 |
| 48 | Pld5 | 0 | 5.717196 | Lamp5 |
| 49 | Pde11a | 0 | 7.7512107 | Lamp5 |
| 50 | Hdac9 | 0 | 4.041462 | Lamp5 |
| 51 | Zfp536 | 0 | 5.7475715 | Lamp5 |
| 52 | Dlx6os1 | 0 | 6.903718 | Lamp5 |
| 53 | Pip5k1b | 0 | 5.415047 | Lamp5 |
| 54 | Id2 | 0 | 4.78302 | Lamp5 |
| 55 | Cryab | 0 | 6.1502457 | Lamp5 |
| 56 | Igsf11 | 0 | 5.352938 | Lamp5 |
| 57 | Ptprm | 0 | 5.773989 | Lamp5 |
| 58 | Lmo2 | 0 | 6.188536 | Lamp5 |
| 59 | Tnfaip8l3 | 0 | 7.708152 | Lamp5 |
| 60 | Npy | 0 | 10.088898 | Lamp5 |
| 61 | Ngf | 0 | 6.980342 | Lamp5 |
| 62 | Kctd6 | 0 | 3.612049 | Lamp5 |
| 63 | Lamp5 | 0 | 7.751709 | Lamp5 |
| 64 | Kit | 0 | 8.292356 | Lamp5 |
| 65 | Slc6a1 | 0 | 7.1077857 | Lamp5 |
| 66 | Hapln1 | 0 | 9.035785 | Lamp5 |
| 67 | Fgf13 | 0 | 4.6347237 | Lamp5 |
| 68 | Cplx3 | 0 | 9.721323 | Lamp5 |
| 69 | Gad1 | 0 | 8.523781 | Lamp5 |
| 70 | Rab3c | 0 | 3.3054745 | Lamp5 |
| 71 | Gad2 | 0 | 7.9433885 | Lamp5 |
| 72 | Parm1 | 0 | 5.9310455 | Lamp5 |
| 73 | Sema3c | 0 | 6.8381424 | Lamp5 |
| 74 | Resp18 | 0 | 3.2248623 | Lamp5 |
| 75 | Nfib | 0 | 4.0363154 | Lamp5 |
| 76 | Nyap2 | 0 | 4.0839043 | Lamp5 |
| 77 | Bcl11b | 0 | 4.2242107 | Lamp5 |
| 78 | Fgf9 | 0 | 3.48928 | Lamp5 |
| 79 | Spock3 | 0 | 3.8198736 | Lamp5 |
| 80 | Trpc5 | 0 | 4.6576743 | Lamp5 |
| 81 | Arl4c | 0 | 4.4898586 | Lamp5 |
| 82 | Tnnt1 | 0 | 7.982083 | Lamp5 |
| 83 | Gabrb2 | 0 | 2.340536 | Lamp5 |
| 84 | Krt1 | 0 | 4.544824 | Lamp5 |
| 85 | Rgs7bp | 0 | 1.7382944 | Lamp5 |
| 86 | Grik1 | 0 | 4.506535 | Lamp5 |
| 87 | Adra1a | 0 | 5.1145673 | Lamp5 |

| | | | | |
|---|---|---|---|---|
| 88 | Gad1-ps | 0 | 3.1262913 | Lamp5 |
| 89 | Cpne7 | 0 | 5.696211 | Lamp5 |
| 90 | 2900055J20Rik | 0 | 2.8745627 | Lamp5 |
| 91 | Klhl13 | 0 | 6.1920476 | Lamp5 |
| 92 | A830018L16Rik | 0 | 3.0490918 | Lamp5 |
| 93 | Adarb2 | 0 | 7.3662896 | Lamp5 |
| 94 | Dlx1as | 0 | 5.563658 | Lamp5 |
| 95 | Tspan7 | 0 | 1.1656075 | Lamp5 |
| 96 | Pnoc | 0 | 7.497781 | Lamp5 |
| 97 | Zmat4 | 0 | 4.3789887 | Lamp5 |
| 98 | Crispld1 | 0 | 6.820142 | Lamp5 |
| 99 | Osbpl8 | 0 | 2.1401262 | Lamp5 |
| 0 | Rnr2 | 4.67E-53 | 1.7962486 | Low Quality |
| 1 | COX1 | 3.14E-50 | 1.266103 | Low Quality |
| 2 | CYTB | 9.50E-47 | 1.3303863 | Low Quality |
| 3 | ND4 | 2.11E-46 | 1.4288906 | Low Quality |
| 4 | Rnr1 | 1.30E-44 | 1.859949 | Low Quality |
| 5 | Gm6581 | 4.21E-43 | 1.4230634 | Low Quality |
| 6 | TrnQ | 6.39E-43 | 1.9890516 | Low Quality |
| 7 | ND5 | 3.19E-41 | 1.0623606 | Low Quality |
| 8 | ND6 | 6.54E-40 | 1.2212945 | Low Quality |
| 9 | ND2 | 9.67E-38 | 1.1611369 | Low Quality |
| 10 | Gm15266 | 3.19E-34 | 1.3662248 | Low Quality |
| 11 | ND3 | 3.83E-30 | 1.2779305 | Low Quality |
| 12 | Apoe | 3.18E-27 | 5.3236856 | Low Quality |
| 13 | ND1 | 8.52E-26 | 0.7306352 | Low Quality |
| 14 | LOC100503946 | 8.52E-23 | 1.3143748 | Low Quality |
| 15 | Mt2 | 4.64E-22 | 4.856639 | Low Quality |
| 16 | COX2 | 2.16E-21 | 1.3872689 | Low Quality |
| 17 | COX3 | 2.75E-21 | 1.3239795 | Low Quality |
| 18 | ATP6 | 4.30E-21 | 1.2815057 | Low Quality |
| 19 | Rn18s-rs5 | 2.09E-20 | 1.3474948 | Low Quality |
| 20 | LOC105246169 | 9.72E-18 | 1.7335308 | Low Quality |
| 21 | TrnV | 5.85E-16 | 3.7003345 | Low Quality |
| 22 | LOC105245046 | 8.20E-16 | 1.179258 | Low Quality |
| 23 | TrnA | 1.56E-15 | 2.5596771 | Low Quality |
| 24 | Ftl1 | 2.10E-15 | -0.16934772 | Low Quality |
| 25 | Gm8129 | 2.56E-15 | 0.6118086 | Low Quality |
| 26 | TrnP | 1.47E-14 | 1.2761883 | Low Quality |
| 27 | Gm20594 | 2.72E-14 | 2.5025458 | Low Quality |
| 28 | Gm33085 | 5.41E-14 | 1.8194934 | Low Quality |
| 29 | TrnM | 7.58E-14 | 2.5731244 | Low Quality |
| 30 | Gm9683 | 1.63E-13 | 3.3726506 | Low Quality |
| 31 | LOC105245638 | 2.74E-13 | 1.8813407 | Low Quality |
| 32 | Gm10116 | 3.66E-13 | 1.4210795 | Low Quality |
| 33 | TrnC | 2.69E-12 | 2.3727546 | Low Quality |
| 34 | Gm12164 | 5.02E-12 | 1.747178 | Low Quality |
| 35 | ATP8 | 1.87E-11 | 1.3185102 | Low Quality |
| 36 | Fth1 | 3.10E-11 | -0.31705603 | Low Quality |
| 37 | TrnL1 | 3.92E-11 | 2.9157445 | Low Quality |

| | | | | |
|---|---|---|---|---|
| 38 | Gm11478 | 1.11E-10 | 1.167199 | Low Quality |
| 39 | Col7a1 | 2.27E-10 | 4.081364 | Low Quality |
| 40 | Gm6789 | 4.99E-10 | 2.6374927 | Low Quality |
| 41 | Igk | 7.35E-10 | 1.5269464 | Low Quality |
| 42 | Gm4617 | 1.43E-09 | 0.85900855 | Low Quality |
| 43 | Ptma | 1.75E-09 | 0.079633705 | Low Quality |
| 44 | Dap | 2.95E-09 | 3.0540173 | Low Quality |
| 45 | Lars2 | 8.03E-09 | 0.7180746 | Low Quality |
| 46 | Gm4735 | 1.74E-08 | 0.46479908 | Low Quality |
| 47 | Mt1 | 1.92E-08 | 1.6200355 | Low Quality |
| 48 | Mir6240 | 1.94E-08 | 2.2529664 | Low Quality |
| 49 | TrnS2 | 2.52E-08 | 3.8540573 | Low Quality |
| 50 | TrnI | 2.97E-08 | 2.5665681 | Low Quality |
| 51 | Rps27rt | 3.25E-08 | 0.81048375 | Low Quality |
| 52 | Ezr | 3.69E-08 | 3.1620843 | Low Quality |
| 53 | Cnn3 | 4.83E-08 | 2.0795937 | Low Quality |
| 54 | Il11ra1 | 5.18E-08 | 2.017662 | Low Quality |
| 55 | Gm11361 | 5.31E-08 | 1.2687792 | Low Quality |
| 56 | Gm14303 | 6.32E-08 | 0.32215774 | Low Quality |
| 57 | Vsig2 | 7.72E-08 | 1.4390268 | Low Quality |
| 58 | Rpl13a | 8.40E-08 | -0.41024056 | Low Quality |
| 59 | Gm7204 | 9.30E-08 | 1.140215 | Low Quality |
| 60 | Prdx1 | 1.11E-07 | -0.5156132 | Low Quality |
| 61 | TrnE | 1.29E-07 | 1.4648442 | Low Quality |
| 62 | Gm5879 | 1.69E-07 | 1.3181486 | Low Quality |
| 63 | TrnF | 1.94E-07 | 2.8564198 | Low Quality |
| 64 | Ifi30 | 1.94E-07 | 3.421905 | Low Quality |
| 65 | S100b | 3.06E-07 | 2.6815398 | Low Quality |
| 66 | Gm26756 | 3.61E-07 | 3.350898 | Low Quality |
| 67 | Lat | 5.22E-07 | 4.274949 | Low Quality |
| 68 | Baiap2 | 5.38E-07 | 0.865952 | Low Quality |
| 69 | Gm11189 | 5.63E-07 | 1.8949986 | Low Quality |
| 70 | Gm20746 | 8.05E-07 | 1.3709948 | Low Quality |
| 71 | Gm3953 | 9.14E-07 | 2.0176573 | Low Quality |
| 72 | Camk2a | 1.24E-06 | 0.5635996 | Low Quality |
| 73 | Gm21399 | 1.43E-06 | 1.089541 | Low Quality |
| 74 | Gstm2-ps1 | 1.48E-06 | 3.1820695 | Low Quality |
| 75 | Id2 | 2.14E-06 | 0.91155446 | Low Quality |
| 76 | Bai1 | 2.94E-06 | 0.4740622 | Low Quality |
| 77 | Mir6236 | 3.37E-06 | 0.7851628 | Low Quality |
| 78 | Gm8814 | 3.65E-06 | 1.1558611 | Low Quality |
| 79 | Rps23 | 4.25E-06 | -0.60018164 | Low Quality |
| 80 | LOC105245621 | 4.61E-06 | 1.2716047 | Low Quality |
| 81 | Gm18173 | 4.78E-06 | 1.713263 | Low Quality |
| 82 | Rps14 | 5.61E-06 | -0.62136734 | Low Quality |
| 83 | Slc1a2 | 6.44E-06 | 1.1500709 | Low Quality |
| 84 | TrnD | 7.62E-06 | 4.589399 | Low Quality |
| 85 | Nmb | 8.50E-06 | 2.8441558 | Low Quality |
| 86 | Gstm1 | 9.82E-06 | 2.823284 | Low Quality |
| 87 | Rpl35 | 1.18E-05 | -0.3344731 | Low Quality |

| | | | | |
|---|---|---|---|---|
| 88 | Cdt1 | 1.24E-05 | 3.1553495 | Low Quality |
| 89 | Ssbp4 | 1.36E-05 | -0.26783958 | Low Quality |
| 90 | Tlcd2 | 1.37E-05 | 4.554798 | Low Quality |
| 91 | Csrp1 | 1.78E-05 | 2.195735 | Low Quality |
| 92 | Gm17936 | 2.11E-05 | 2.2849545 | Low Quality |
| 93 | TrnL2 | 2.15E-05 | 3.5433602 | Low Quality |
| 94 | Gm12892 | 2.40E-05 | 0.38460264 | Low Quality |
| 95 | Gm18728 | 2.64E-05 | 1.7354684 | Low Quality |
| 96 | Oxt | 2.83E-05 | 5.100859 | Low Quality |
| 97 | Cox6a1 | 2.85E-05 | -0.7285907 | Low Quality |
| 98 | Ppp1r1b | 3.39E-05 | 1.5796982 | Low Quality |
| 99 | Mdk | 3.48E-05 | 2.977959 | Low Quality |
| 0 | Fcer1g | 3.74E-83 | 17.27102 | Macrophage |
| 1 | C1qa | 3.74E-83 | 18.16257 | Macrophage |
| 2 | C1qc | 3.74E-83 | 18.65388 | Macrophage |
| 3 | C1qb | 3.74E-83 | 18.779028 | Macrophage |
| 4 | Fcgr3 | 3.74E-83 | 16.011482 | Macrophage |
| 5 | Ctss | 3.74E-83 | 18.427763 | Macrophage |
| 6 | Tyrobp | 3.74E-83 | 17.346004 | Macrophage |
| 7 | Tmsb4x | 1.07E-82 | 3.0944514 | Macrophage |
| 8 | B2m | 2.02E-82 | 6.179138 | Macrophage |
| 9 | Fcrls | 4.26E-82 | 16.957088 | Macrophage |
| 10 | Csf1r | 4.26E-82 | 17.173004 | Macrophage |
| 11 | Lgmn | 4.26E-82 | 5.11481 | Macrophage |
| 12 | Laptm5 | 4.26E-82 | 16.811726 | Macrophage |
| 13 | Cst3 | 5.06E-82 | 6.3549447 | Macrophage |
| 14 | Unc93b1 | 9.91E-82 | 13.775535 | Macrophage |
| 15 | Ly86 | 6.50E-81 | 16.713326 | Macrophage |
| 16 | Aif1 | 8.23E-81 | 12.641487 | Macrophage |
| 17 | Cyba | 1.22E-80 | 13.3355 | Macrophage |
| 18 | Ftl1 | 2.22E-80 | 3.3210316 | Macrophage |
| 19 | Ctsd | 2.23E-80 | 5.0583463 | Macrophage |
| 20 | Cd53 | 1.17E-79 | 15.092299 | Macrophage |
| 21 | 4632428N05Rik | 1.77E-79 | 14.222475 | Macrophage |
| 22 | Pld4 | 1.03E-78 | 15.932829 | Macrophage |
| 23 | P2ry12 | 1.80E-78 | 15.621141 | Macrophage |
| 24 | Zfp36 | 5.07E-78 | 13.54961 | Macrophage |
| 25 | Serinc3 | 7.28E-78 | 3.3707266 | Macrophage |
| 26 | Txnip | 1.57E-77 | 13.241795 | Macrophage |
| 27 | Sepp1 | 1.74E-77 | 12.804172 | Macrophage |
| 28 | Itm2b | 2.34E-77 | 1.9441341 | Macrophage |
| 29 | Rnase4 | 2.91E-77 | 13.492336 | Macrophage |
| 30 | Rnaset2a | 3.33E-76 | 6.350267 | Macrophage |
| 31 | Zfhx3 | 3.98E-76 | 9.664404 | Macrophage |
| 32 | Gpr34 | 4.39E-76 | 13.567857 | Macrophage |
| 33 | Rnaset2b | 8.69E-76 | 5.9547644 | Macrophage |
| 34 | Hexb | 5.40E-75 | 9.540816 | Macrophage |
| 35 | Trem2 | 7.60E-75 | 12.324406 | Macrophage |
| 36 | Grn | 1.93E-74 | 5.8474607 | Macrophage |
| 37 | Cx3cr1 | 2.15E-74 | 16.707466 | Macrophage |

| | | | | |
|---|---|---|---|---|
| 38 | Ltc4s | 3.19E-74 | 13.262836 | Macrophage |
| 39 | Zfp36l1 | 4.37E-74 | 10.234196 | Macrophage |
| 40 | Trf | 4.98E-74 | 11.939297 | Macrophage |
| 41 | AF251705 | 3.77E-73 | 14.152801 | Macrophage |
| 42 | Ctsz | 7.68E-73 | 6.994981 | Macrophage |
| 43 | Cd68 | 2.17E-72 | 10.589472 | Macrophage |
| 44 | Gm14303 | 5.47E-71 | 2.4066782 | Macrophage |
| 45 | Sat1 | 1.00E-70 | 4.646158 | Macrophage |
| 46 | Ftl2 | 1.21E-70 | 3.1621096 | Macrophage |
| 47 | Rgs10 | 3.95E-70 | 6.7954826 | Macrophage |
| 48 | Cyth4 | 1.13E-69 | 13.286673 | Macrophage |
| 49 | Fyb | 9.11E-69 | 14.772811 | Macrophage |
| 50 | Hmha1 | 9.86E-69 | 14.271204 | Macrophage |
| 51 | Hpgds | 3.11E-68 | 10.875006 | Macrophage |
| 52 | Lyn | 6.09E-68 | 11.779037 | Macrophage |
| 53 | Rps29 | 6.26E-68 | 2.0630043 | Macrophage |
| 54 | Spi1 | 1.08E-67 | 14.965279 | Macrophage |
| 55 | Mertk | 2.67E-67 | 10.286898 | Macrophage |
| 56 | Gng5 | 3.06E-67 | 6.3804154 | Macrophage |
| 57 | Slco2b1 | 6.35E-67 | 12.282058 | Macrophage |
| 58 | Marcks | 1.35E-66 | 3.4488637 | Macrophage |
| 59 | Gm10116 | 1.81E-66 | 3.4151769 | Macrophage |
| 60 | Pfn1 | 2.06E-66 | 1.9067116 | Macrophage |
| 61 | Entpd1 | 2.79E-66 | 10.086884 | Macrophage |
| 62 | Kctd12 | 8.53E-66 | 7.8266654 | Macrophage |
| 63 | Emr1 | 1.84E-65 | 15.498029 | Macrophage |
| 64 | Jun | 3.54E-65 | 7.4843726 | Macrophage |
| 65 | Junb | 5.02E-65 | 6.3234143 | Macrophage |
| 66 | Clic1 | 7.70E-65 | 10.253839 | Macrophage |
| 67 | Selplg | 1.12E-64 | 15.091084 | Macrophage |
| 68 | P2ry13 | 3.27E-64 | 14.400961 | Macrophage |
| 69 | Ptpn18 | 3.63E-64 | 12.943416 | Macrophage |
| 70 | Ivns1abp | 5.82E-64 | 4.211828 | Macrophage |
| 71 | Ctsh | 1.61E-63 | 12.049917 | Macrophage |
| 72 | Gm12164 | 2.32E-63 | 3.673639 | Macrophage |
| 73 | Ly6e | 4.07E-63 | 2.8544755 | Macrophage |
| 74 | Itgb5 | 7.98E-63 | 10.355321 | Macrophage |
| 75 | Rhob | 9.46E-63 | 4.147983 | Macrophage |
| 76 | Maf | 1.10E-61 | 6.1465554 | Macrophage |
| 77 | Lpcat2 | 1.38E-61 | 10.155346 | Macrophage |
| 78 | Gpx1 | 1.50E-61 | 2.922501 | Macrophage |
| 79 | Rab3il1 | 1.72E-61 | 9.17294 | Macrophage |
| 80 | H3f3b | 2.33E-61 | 2.3422046 | Macrophage |
| 81 | Ccdc152 | 3.30E-61 | 6.8280516 | Macrophage |
| 82 | Gpr183 | 4.60E-61 | 10.57026 | Macrophage |
| 83 | Gm8814 | 1.04E-60 | 3.4429326 | Macrophage |
| 84 | Cfh | 2.01E-60 | 11.778444 | Macrophage |
| 85 | Ubc | 5.47E-60 | 1.8166832 | Macrophage |
| 86 | Siglech | 5.77E-60 | 16.195185 | Macrophage |
| 87 | Hk2 | 9.23E-60 | 9.973263 | Macrophage |

| | | | | |
|---|---|---|---|---|
| 88 | Ifngr1 | 1.33E-59 | 8.12127 | Macrophage |
| 89 | Inpp5d | 1.83E-59 | 9.177308 | Macrophage |
| 90 | Hpgd | 1.95E-59 | 10.83566 | Macrophage |
| 91 | Gm7618 | 1.08E-58 | 3.3651981 | Macrophage |
| 92 | Arpc1b | 1.35E-58 | 8.390062 | Macrophage |
| 93 | Tgfbr1 | 2.35E-58 | 8.743285 | Macrophage |
| 94 | Rgs2 | 3.63E-58 | 4.433374 | Macrophage |
| 95 | Arhgdib | 6.07E-58 | 9.190225 | Macrophage |
| 96 | Crybb1 | 8.30E-58 | 13.482924 | Macrophage |
| 97 | Btg2 | 1.28E-57 | 7.804659 | Macrophage |
| 98 | Gm20746 | 1.50E-57 | 3.516986 | Macrophage |
| 99 | Glul | 2.01E-57 | 4.228671 | Macrophage |
| 0 | H3f3b | 7.45E-26 | 1.7268059 | Meis2 |
| 1 | Pcbp3 | 1.11E-25 | 3.8557677 | Meis2 |
| 2 | Gng4 | 1.83E-23 | 5.532091 | Meis2 |
| 3 | Meis2 | 4.04E-22 | 6.6736445 | Meis2 |
| 4 | Sp8 | 8.88E-22 | 6.2615376 | Meis2 |
| 5 | Btg1 | 1.51E-21 | 3.808544 | Meis2 |
| 6 | H3f3a | 1.79E-21 | 1.4079708 | Meis2 |
| 7 | Penk | 3.46E-21 | 8.560697 | Meis2 |
| 8 | Pcp4l1 | 4.31E-21 | 6.0289054 | Meis2 |
| 9 | Frmd7 | 2.56E-20 | 10.248264 | Meis2 |
| 10 | Ube2e3 | 3.78E-20 | 2.1871717 | Meis2 |
| 11 | Gm6485 | 4.15E-20 | 1.7406992 | Meis2 |
| 12 | Cpne6 | 7.05E-20 | 5.5639796 | Meis2 |
| 13 | Scgn | 1.71E-19 | 11.207891 | Meis2 |
| 14 | Ckb | 2.94E-19 | 1.4502919 | Meis2 |
| 15 | Ck-ps3 | 3.83E-19 | 1.8654296 | Meis2 |
| 16 | Map3k1 | 6.78E-19 | 4.6105733 | Meis2 |
| 17 | G630016G05Rik | 9.46E-19 | 5.693485 | Meis2 |
| 18 | Shisa8 | 1.45E-18 | 7.596503 | Meis2 |
| 19 | Hist3h2a | 2.69E-18 | 2.2887542 | Meis2 |
| 20 | Grb2 | 5.14E-18 | 1.9929765 | Meis2 |
| 21 | Pbx3 | 5.97E-18 | 7.2813573 | Meis2 |
| 22 | Ptpro | 1.53E-17 | 5.9387283 | Meis2 |
| 23 | Trim17 | 1.95E-17 | 2.2803745 | Meis2 |
| 24 | Rsrp1 | 4.10E-17 | 1.0031189 | Meis2 |
| 25 | Gm12892 | 4.40E-17 | 1.518523 | Meis2 |
| 26 | Ybx1 | 1.17E-16 | 1.336521 | Meis2 |
| 27 | Clk1 | 1.34E-16 | 1.6871529 | Meis2 |
| 28 | Synpr | 1.38E-16 | 6.4494376 | Meis2 |
| 29 | Tuba1a | 3.77E-16 | 0.97151375 | Meis2 |
| 30 | Apc | 6.47E-16 | 2.174954 | Meis2 |
| 31 | Pcp4 | 9.10E-16 | 4.945935 | Meis2 |
| 32 | Ptma | 2.22E-15 | 1.0416259 | Meis2 |
| 33 | Myo16 | 2.23E-15 | 5.526175 | Meis2 |
| 34 | Hmgn2 | 3.10E-15 | 1.552637 | Meis2 |
| 35 | Amigo2 | 3.17E-15 | 7.4049463 | Meis2 |
| 36 | Cited2 | 3.94E-15 | 2.5998044 | Meis2 |
| 37 | Nrip3 | 4.04E-15 | 3.7253098 | Meis2 |

| | | | | |
|---|---|---|---|---|
| 38 | Malat1 | 5.33E-15 | 1.0499008 | Meis2 |
| 39 | Mt1 | 1.81E-14 | 2.959922 | Meis2 |
| 40 | 1700019L22Rik | 2.43E-14 | 6.225634 | Meis2 |
| 41 | Dlx5 | 8.54E-14 | 4.371073 | Meis2 |
| 42 | Itm2c | 1.26E-13 | 1.2024941 | Meis2 |
| 43 | Sez6 | 1.35E-13 | 3.1475074 | Meis2 |
| 44 | Mtfp1 | 2.16E-13 | 1.679034 | Meis2 |
| 45 | Gpsm1 | 3.00E-13 | 5.604398 | Meis2 |
| 46 | Selm | 3.22E-13 | 1.1968935 | Meis2 |
| 47 | Podxl2 | 3.32E-13 | 1.7240285 | Meis2 |
| 48 | Dgkb | 3.87E-13 | 3.968748 | Meis2 |
| 49 | Fam184a | 5.00E-13 | 3.949181 | Meis2 |
| 50 | Pbx1 | 5.36E-13 | 2.558641 | Meis2 |
| 51 | Marcks | 8.08E-13 | 2.0878878 | Meis2 |
| 52 | Th | 1.55E-12 | 9.210231 | Meis2 |
| 53 | Epha5 | 1.69E-12 | 2.42152 | Meis2 |
| 54 | Hmgn1 | 2.05E-12 | 1.0869638 | Meis2 |
| 55 | Lgr5 | 3.59E-12 | 7.188122 | Meis2 |
| 56 | Ddx5 | 4.14E-12 | 0.93818843 | Meis2 |
| 57 | Hnrnpc | 1.97E-11 | 0.4474904 | Meis2 |
| 58 | Zfp385b | 3.20E-11 | 2.3041854 | Meis2 |
| 59 | Dcx | 5.06E-11 | 3.5930152 | Meis2 |
| 60 | Syt6 | 5.20E-11 | 6.2136436 | Meis2 |
| 61 | Hist1h2bc | 6.01E-11 | 3.9538462 | Meis2 |
| 62 | Isoc1 | 7.77E-11 | 5.1484327 | Meis2 |
| 63 | Rbm39 | 8.91E-11 | 0.86907184 | Meis2 |
| 64 | Hnrnpa2b1 | 1.04E-10 | 0.6980943 | Meis2 |
| 65 | Pkig | 1.07E-10 | 1.1316416 | Meis2 |
| 66 | Meis1 | 1.09E-10 | 6.0944476 | Meis2 |
| 67 | Sp9 | 1.45E-10 | 3.9632764 | Meis2 |
| 68 | Ahi1 | 1.77E-10 | 1.4771533 | Meis2 |
| 69 | Rpl13 | 1.77E-10 | 0.7902563 | Meis2 |
| 70 | Dclk2 | 1.80E-10 | 3.47985 | Meis2 |
| 71 | Hap1 | 1.90E-10 | 3.7989566 | Meis2 |
| 72 | Pnmal2 | 1.98E-10 | 1.498291 | Meis2 |
| 73 | Gm13889 | 2.03E-10 | 2.9044247 | Meis2 |
| 74 | Adamts19 | 2.16E-10 | 9.203993 | Meis2 |
| 75 | Ck-ps1 | 2.30E-10 | 2.0487473 | Meis2 |
| 76 | Celf4 | 2.41E-10 | 1.2769296 | Meis2 |
| 77 | Dync1i1 | 2.50E-10 | 1.6838905 | Meis2 |
| 78 | Celf2 | 2.60E-10 | 1.06834 | Meis2 |
| 79 | Cacna1c | 3.77E-10 | 2.7224298 | Meis2 |
| 80 | Ftl1 | 4.96E-10 | 0.8123904 | Meis2 |
| 81 | Gad1 | 7.22E-10 | 6.157554 | Meis2 |
| 82 | Ppp1cc | 8.01E-10 | 0.9790801 | Meis2 |
| 83 | Dlx1 | 1.34E-09 | 3.6843483 | Meis2 |
| 84 | Rpp25 | 1.44E-09 | 3.503388 | Meis2 |
| 85 | Srsf7 | 1.92E-09 | 0.6543161 | Meis2 |
| 86 | Gm7554 | 2.03E-09 | 2.772372 | Meis2 |
| 87 | H3f3a-ps2 | 2.28E-09 | 1.5703899 | Meis2 |

| | | | | |
|---|---|---|---|---|
| 88 | Emp3 | 3.36E-09 | 6.6503386 | Meis2 |
| 89 | Eno2 | 4.59E-09 | 0.6778277 | Meis2 |
| 90 | Psip1 | 4.62E-09 | 0.8032024 | Meis2 |
| 91 | Ubb | 6.82E-09 | 0.39226124 | Meis2 |
| 92 | Ccdc109b | 8.09E-09 | 5.3714566 | Meis2 |
| 93 | Atp2b4 | 8.11E-09 | 3.442773 | Meis2 |
| 94 | Sall3 | 8.28E-09 | 7.0751915 | Meis2 |
| 95 | Son | 1.32E-08 | 0.803734 | Meis2 |
| 96 | Grm2 | 1.33E-08 | 3.824761 | Meis2 |
| 97 | Atp5g2 | 1.58E-08 | 0.51888895 | Meis2 |
| 98 | COX2 | 1.61E-08 | 1.0833082 | Meis2 |
| 99 | Kcnh6 | 1.80E-08 | 2.2127194 | Meis2 |
| 0 | Tshz2 | 0 | 10.878432 | NP |
| 1 | Vwc2l | 0 | 6.905071 | NP |
| 2 | Gsta4 | 0 | 4.8928714 | NP |
| 3 | Sla2 | 0 | 7.979017 | NP |
| 4 | Lmo3 | 0 | 4.3099456 | NP |
| 5 | Neto2 | 0 | 4.594931 | NP |
| 6 | Meis2 | 0 | 6.2814093 | NP |
| 7 | Tle4 | 0 | 5.335202 | NP |
| 8 | Gm11223 | 0 | 1.9107449 | NP |
| 9 | Slit1 | 0 | 5.4143353 | NP |
| 10 | Stard5 | 0 | 6.65663 | NP |
| 11 | Adcy2 | 0 | 4.3483963 | NP |
| 12 | Rprm | 0 | 6.457656 | NP |
| 13 | Stmn1 | 0 | 1.9689063 | NP |
| 14 | Il11ra1 | 0 | 6.7955194 | NP |
| 15 | Etv1 | 0 | 7.396979 | NP |
| 16 | Nxph3 | 0 | 7.3734584 | NP |
| 17 | Gm5454 | 0 | 5.4183106 | NP |
| 18 | Tmem159 | 0 | 6.35411 | NP |
| 19 | Fezf2 | 0 | 7.6127944 | NP |
| 20 | Myl4 | 0 | 8.537749 | NP |
| 21 | Efr3a | 0 | 2.791498 | NP |
| 22 | Lypd1 | 0 | 8.219433 | NP |
| 23 | Olfm3 | 0 | 6.4783845 | NP |
| 24 | Sec62 | 0 | 1.4240088 | NP |
| 25 | Uap1 | 1.07E-303 | 2.9753692 | NP |
| 26 | Trp53i11 | 2.10E-300 | 5.7933784 | NP |
| 27 | Fdps | 7.32E-298 | 2.2900825 | NP |
| 28 | Dync1i1 | 1.36E-297 | 2.1742272 | NP |
| 29 | Rnf152 | 3.55E-297 | 5.8460617 | NP |
| 30 | Ift57 | 2.49E-295 | 2.5444903 | NP |
| 31 | Plcxd2 | 1.38E-294 | 4.987351 | NP |
| 32 | Spon1 | 3.75E-292 | 5.919289 | NP |
| 33 | Amigo2 | 1.04E-290 | 6.921702 | NP |
| 34 | Pcp4 | 9.13E-281 | 5.897216 | NP |
| 35 | LOC105244376 | 7.24E-279 | 8.78527 | NP |
| 36 | Prkca | 8.82E-279 | 3.211461 | NP |
| 37 | Rap1gds1 | 1.10E-278 | 2.097057 | NP |

| | | | | |
|---|---|---|---|---|
| 38 | Crym | 1.76E-278 | 7.188455 | NP |
| 39 | Kcnmb4 | 1.79E-276 | 3.187967 | NP |
| 40 | Camk2d | 1.30E-272 | 3.4317937 | NP |
| 41 | Stxbp2 | 2.32E-272 | 4.4442315 | NP |
| 42 | Trpc3 | 2.63E-272 | 5.244662 | NP |
| 43 | Nrsn2 | 2.21E-271 | 4.1353283 | NP |
| 44 | Pamr1 | 1.62E-270 | 5.892152 | NP |
| 45 | Hlf | 1.28E-269 | 3.385624 | NP |
| 46 | Ctgf | 6.94E-267 | 6.4417644 | NP |
| 47 | Marcks | 5.05E-266 | 2.3484662 | NP |
| 48 | Mdh1 | 4.22E-259 | 1.0498351 | NP |
| 49 | Fut9 | 3.43E-257 | 2.776549 | NP |
| 50 | Kcna6 | 1.20E-256 | 2.8034232 | NP |
| 51 | Diras2 | 1.78E-253 | 3.8584826 | NP |
| 52 | Myzap | 1.40E-252 | 9.15083 | NP |
| 53 | P4ha1 | 2.39E-252 | 3.1684196 | NP |
| 54 | Trhr | 2.03E-249 | 10.188203 | NP |
| 55 | Ccdc12 | 6.07E-249 | 1.5698781 | NP |
| 56 | Kcnk2 | 6.44E-242 | 3.9886131 | NP |
| 57 | Shisa4 | 5.15E-240 | 2.138871 | NP |
| 58 | Msmo1 | 6.62E-238 | 2.4592957 | NP |
| 59 | Cxx1c | 7.55E-233 | 1.2581403 | NP |
| 60 | Kremen1 | 1.65E-232 | 5.5348096 | NP |
| 61 | Tmsb10 | 4.79E-227 | 1.9904286 | NP |
| 62 | Gm2163 | 5.59E-227 | 2.6365252 | NP |
| 63 | Chgb | 2.31E-225 | 2.0529997 | NP |
| 64 | Col12a1 | 1.63E-223 | 6.012377 | NP |
| 65 | Ssbp2 | 1.02E-222 | 1.6600144 | NP |
| 66 | Unc13b | 4.71E-221 | 3.917624 | NP |
| 67 | Mgat4c | 4.05E-220 | 4.631801 | NP |
| 68 | Tecr | 1.42E-219 | 0.7527594 | NP |
| 69 | Wbp5 | 1.63E-219 | 1.2682129 | NP |
| 70 | B3glct | 1.67E-217 | 4.041258 | NP |
| 71 | Cd47 | 1.49E-216 | 1.1530744 | NP |
| 72 | Scai | 2.27E-216 | 2.7724276 | NP |
| 73 | Pcdh17 | 2.37E-215 | 3.4107628 | NP |
| 74 | Bcl11b | 2.00E-214 | 3.8797677 | NP |
| 75 | Pid1 | 1.69E-213 | 3.6050806 | NP |
| 76 | 5830416P10Rik | 1.98E-213 | 6.3935804 | NP |
| 77 | Lor | 1.87E-211 | 3.0802202 | NP |
| 78 | Neurod6 | 1.09E-210 | 5.7584896 | NP |
| 79 | Scg3 | 2.35E-208 | 1.7589178 | NP |
| 80 | Camk1 | 2.94E-208 | 1.6675187 | NP |
| 81 | Rit2 | 6.02E-206 | 1.7606431 | NP |
| 82 | Cib2 | 9.99E-203 | 2.5778782 | NP |
| 83 | Foxp1 | 6.93E-202 | 3.042659 | NP |
| 84 | Bcat1 | 1.41E-201 | 3.03046 | NP |
| 85 | Cdc5l | 2.22E-200 | 1.7530499 | NP |
| 86 | Adora1 | 4.67E-200 | 3.7939034 | NP |
| 87 | Oaz1 | 6.89E-200 | 0.6404475 | NP |

| | | | | |
|---|---|---|---|---|
| 88 | G630016G05Rik | 5.86E-199 | 3.7854083 | NP |
| 89 | Cpne4 | 1.94E-198 | 4.2857018 | NP |
| 90 | Fdft1 | 2.68E-197 | 2.048223 | NP |
| 91 | Hs3st4 | 5.23E-197 | 4.3738213 | NP |
| 92 | Mgst3 | 1.67E-196 | 1.5308026 | NP |
| 93 | Serpini1 | 1.38E-192 | 2.4209428 | NP |
| 94 | Irak2 | 1.96E-192 | 4.5664115 | NP |
| 95 | Gmpr | 2.15E-192 | 2.3082175 | NP |
| 96 | Dusp5 | 5.43E-192 | 3.7221537 | NP |
| 97 | Aes | 1.86E-184 | 0.78428245 | NP |
| 98 | LOC102638890 | 1.88E-184 | 4.5414224 | NP |
| 99 | Gm32412 | 1.95E-184 | 5.5463266 | NP |
| 0 | Tsc22d4 | 1.06E-121 | 4.918315 | No Class |
| 1 | S100a16 | 9.83E-111 | 6.05481 | No Class |
| 2 | Apoe | 2.22E-110 | 5.0325685 | No Class |
| 3 | Cd9 | 8.55E-104 | 7.084992 | No Class |
| 4 | Dbi | 2.75E-98 | 1.657472 | No Class |
| 5 | Plekhb1 | 2.64E-95 | 4.8543153 | No Class |
| 6 | Csrp1 | 3.48E-94 | 4.468008 | No Class |
| 7 | Cmtm5 | 4.48E-93 | 6.3244724 | No Class |
| 8 | Olig1 | 1.91E-91 | 6.831245 | No Class |
| 9 | Gng5 | 1.87E-88 | 2.977068 | No Class |
| 10 | S100a1 | 2.04E-88 | 4.2887306 | No Class |
| 11 | Car2 | 1.49E-82 | 4.4790525 | No Class |
| 12 | Cd63 | 4.62E-82 | 4.027551 | No Class |
| 13 | S100a13 | 1.35E-81 | 4.2951283 | No Class |
| 14 | Qk | 2.53E-79 | 2.9163287 | No Class |
| 15 | Phgdh | 6.44E-77 | 3.9257512 | No Class |
| 16 | Mbp | 1.24E-76 | 3.9088914 | No Class |
| 17 | Bcas1 | 1.99E-76 | 6.8262935 | No Class |
| 18 | Mt1 | 4.04E-76 | 2.3943026 | No Class |
| 19 | Cnp | 1.11E-72 | 4.749875 | No Class |
| 20 | Plp1 | 6.82E-72 | 6.6900153 | No Class |
| 21 | Hist1h2bc | 7.49E-70 | 2.8466089 | No Class |
| 22 | Gltp | 7.49E-70 | 4.0856133 | No Class |
| 23 | Ptgds | 1.38E-69 | 5.186974 | No Class |
| 24 | Rhog | 2.69E-69 | 4.740856 | No Class |
| 25 | Ppap2b | 7.74E-68 | 3.8797538 | No Class |
| 26 | Ftl1 | 2.19E-65 | 0.7213474 | No Class |
| 27 | Kcnj10 | 1.23E-64 | 5.269391 | No Class |
| 28 | Gstm1 | 4.81E-62 | 3.3037167 | No Class |
| 29 | Pllp | 4.47E-59 | 4.3787966 | No Class |
| 30 | Gatm | 9.05E-58 | 3.8128078 | No Class |
| 31 | Dock10 | 4.29E-57 | 2.9315917 | No Class |
| 32 | Mobp | 1.12E-56 | 4.679862 | No Class |
| 33 | Tspan15 | 2.25E-56 | 4.5743704 | No Class |
| 34 | Wscd1 | 4.64E-56 | 3.00474 | No Class |
| 35 | Sirt2 | 8.91E-55 | 1.6155542 | No Class |
| 36 | Trf | 1.28E-54 | 5.9962926 | No Class |
| 37 | Sox10 | 1.38E-54 | 6.800449 | No Class |

| | | | | |
|---|---|---|---|---|
| 38 | Cd81 | 3.99E-54 | 0.7822305 | No Class |
| 39 | Gjc3 | 2.84E-53 | 7.182736 | No Class |
| 40 | Litaf | 3.20E-53 | 4.781818 | No Class |
| 41 | Tspan2 | 4.11E-53 | 3.5195684 | No Class |
| 42 | Metrn | 1.11E-52 | 3.7288535 | No Class |
| 43 | St18 | 3.51E-52 | 4.7871494 | No Class |
| 44 | Clic4 | 3.85E-52 | 3.4990711 | No Class |
| 45 | Psat1 | 8.12E-52 | 2.5427506 | No Class |
| 46 | Abhd4 | 8.12E-51 | 3.2430012 | No Class |
| 47 | Cers2 | 1.38E-50 | 2.8642843 | No Class |
| 48 | Gpr37l1 | 6.91E-50 | 4.753418 | No Class |
| 49 | Cryab | 2.99E-49 | 3.0265646 | No Class |
| 50 | Hadh | 4.46E-49 | 3.6084151 | No Class |
| 51 | Epas1 | 4.51E-49 | 3.8628216 | No Class |
| 52 | Lamp1 | 4.66E-49 | 0.83912945 | No Class |
| 53 | Elovl1 | 1.71E-48 | 3.7204072 | No Class |
| 54 | Pla2g16 | 7.67E-48 | 4.7457767 | No Class |
| 55 | Phldb1 | 1.54E-47 | 3.1517508 | No Class |
| 56 | Gsn | 1.54E-47 | 3.7060432 | No Class |
| 57 | Cnn3 | 2.99E-47 | 2.7339473 | No Class |
| 58 | Lap3 | 7.20E-47 | 2.3903713 | No Class |
| 59 | Cercam | 7.20E-47 | 3.8611472 | No Class |
| 60 | Npc2 | 1.17E-46 | 1.8182894 | No Class |
| 61 | Cldn11 | 6.94E-46 | 6.9569516 | No Class |
| 62 | Carhsp1 | 9.55E-46 | 4.2038083 | No Class |
| 63 | Lims2 | 9.74E-46 | 4.514063 | No Class |
| 64 | Stxbp3a | 1.08E-45 | 2.9606218 | No Class |
| 65 | Ugt8a | 1.19E-45 | 5.73312 | No Class |
| 66 | Mag | 1.34E-45 | 6.5592937 | No Class |
| 67 | S100b | 3.09E-45 | 3.050094 | No Class |
| 68 | 04-Sep | 4.45E-45 | 1.7738031 | No Class |
| 69 | Chd7 | 5.06E-45 | 3.014047 | No Class |
| 70 | Mt2 | 5.59E-45 | 2.6693206 | No Class |
| 71 | Gstm7 | 8.91E-45 | 2.895145 | No Class |
| 72 | Fa2h | 9.34E-45 | 6.75762 | No Class |
| 73 | Ndrg2 | 9.47E-45 | 2.7757657 | No Class |
| 74 | Slc1a3 | 9.47E-45 | 3.598716 | No Class |
| 75 | Ppp1r14a | 2.64E-44 | 5.6222105 | No Class |
| 76 | Hepacam | 3.12E-44 | 3.9526339 | No Class |
| 77 | Scrg1 | 1.37E-43 | 4.24226 | No Class |
| 78 | Cd82 | 1.81E-43 | 5.923028 | No Class |
| 79 | Olig2 | 3.59E-43 | 5.8939676 | No Class |
| 80 | Zbtb20 | 7.32E-43 | 2.499719 | No Class |
| 81 | Fermt2 | 9.82E-43 | 2.1113813 | No Class |
| 82 | Arpc1b | 1.04E-42 | 3.51331 | No Class |
| 83 | Grb14 | 5.05E-42 | 2.6312218 | No Class |
| 84 | CYTB | 6.65E-42 | 0.5450004 | No Class |
| 85 | Mfge8 | 7.51E-42 | 3.8258767 | No Class |
| 86 | Gal3st1 | 7.51E-42 | 3.336787 | No Class |
| 87 | Rab31 | 9.48E-42 | 2.3847911 | No Class |

| | | | | |
|---|---|---|---|---|
| 88 | Gpr37 | 1.10E-40 | 4.2420635 | No Class |
| 89 | Atp1a2 | 2.13E-40 | 3.021 | No Class |
| 90 | Cpm | 2.13E-40 | 3.6361134 | No Class |
| 91 | Tmbim1 | 2.57E-40 | 3.3748136 | No Class |
| 92 | Mcam | 1.24E-39 | 5.439919 | No Class |
| 93 | Il18 | 5.19E-39 | 2.3858418 | No Class |
| 94 | Arhgef10 | 9.53E-39 | 3.4153802 | No Class |
| 95 | Pdlim2 | 1.54E-38 | 4.7176495 | No Class |
| 96 | 2810468N07Rik | 3.72E-38 | 2.394447 | No Class |
| 97 | Plekha2 | 5.07E-38 | 2.5527477 | No Class |
| 98 | Erbb2ip | 7.34E-38 | 2.2955418 | No Class |
| 99 | Myo6 | 9.00E-38 | 2.4140155 | No Class |
| 0 | Olig1 | 3.89E-116 | 12.954681 | Oligo |
| 1 | Pllp | 4.24E-114 | 10.95058 | Oligo |
| 2 | Sox10 | 1.45E-113 | 13.246759 | Oligo |
| 3 | Dbi | 2.78E-113 | 5.1781898 | Oligo |
| 4 | Cd9 | 1.83E-111 | 12.938216 | Oligo |
| 5 | Qk | 1.17E-110 | 6.991934 | Oligo |
| 6 | S100a16 | 2.77E-110 | 11.449196 | Oligo |
| 7 | Cd81 | 1.46E-109 | 2.60794 | Oligo |
| 8 | Gatm | 8.76E-105 | 9.696034 | Oligo |
| 9 | Gjc3 | 2.39E-103 | 13.204174 | Oligo |
| 10 | Phldb1 | 2.96E-101 | 8.107514 | Oligo |
| 11 | Scd2 | 3.46E-100 | 4.405299 | Oligo |
| 12 | Gm12222 | 1.25E-99 | 4.994621 | Oligo |
| 13 | Tsc22d4 | 1.53E-97 | 8.991424 | Oligo |
| 14 | Enpp2 | 3.66E-96 | 7.2939653 | Oligo |
| 15 | Plekhb1 | 1.50E-94 | 9.145475 | Oligo |
| 16 | Npc1 | 4.15E-93 | 4.3045716 | Oligo |
| 17 | Cmtm5 | 7.47E-93 | 11.462966 | Oligo |
| 18 | Bcas1 | 2.01E-91 | 11.266663 | Oligo |
| 19 | Cercam | 5.77E-90 | 8.945714 | Oligo |
| 20 | Ugt8a | 7.24E-90 | 11.133194 | Oligo |
| 21 | S100a13 | 1.88E-88 | 8.703247 | Oligo |
| 22 | Kcnj10 | 5.43E-86 | 9.545408 | Oligo |
| 23 | S100a1 | 9.65E-84 | 8.314813 | Oligo |
| 24 | Cd9-ps | 6.19E-83 | 6.9943824 | Oligo |
| 25 | Ftl1 | 7.80E-83 | 1.6400133 | Oligo |
| 26 | 2810468N07Rik | 7.63E-81 | 6.3633137 | Oligo |
| 27 | Cnp | 1.05E-80 | 9.562953 | Oligo |
| 28 | Cd63 | 1.04E-79 | 6.5605097 | Oligo |
| 29 | Olig2 | 1.93E-79 | 10.629103 | Oligo |
| 30 | Gltp | 3.88E-79 | 8.061337 | Oligo |
| 31 | Sirt2 | 5.96E-79 | 3.964173 | Oligo |
| 32 | Hepacam | 1.16E-77 | 7.890831 | Oligo |
| 33 | Grb14 | 2.44E-77 | 7.326058 | Oligo |
| 34 | Malat1 | 4.42E-77 | 1.3510035 | Oligo |
| 35 | Plp1 | 8.32E-77 | 12.602878 | Oligo |
| 36 | Jam3 | 1.40E-75 | 6.0072217 | Oligo |
| 37 | Fa2h | 4.43E-75 | 11.574962 | Oligo |

| | | | | |
|---|---|---|---|---|
| 38 | Cers2 | 7.48E-75 | 6.655675 | Oligo |
| 39 | Ptma | 2.32E-73 | 1.752074 | Oligo |
| 40 | Rhog | 6.29E-71 | 9.048545 | Oligo |
| 41 | Wscd1 | 7.35E-71 | 5.9515133 | Oligo |
| 42 | Gng5 | 1.03E-70 | 5.0739045 | Oligo |
| 43 | H3f3a | 1.12E-70 | 1.2606068 | Oligo |
| 44 | Ptgds | 3.97E-69 | 10.641807 | Oligo |
| 45 | Cd63-ps | 1.19E-68 | 3.6305478 | Oligo |
| 46 | Apoe | 1.72E-67 | 8.001794 | Oligo |
| 47 | Lamp1 | 6.78E-67 | 1.9941521 | Oligo |
| 48 | Gal3st1 | 1.23E-66 | 7.2739325 | Oligo |
| 49 | Tmem258 | 1.80E-66 | 1.4544561 | Oligo |
| 50 | Cdc42ep1 | 4.18E-66 | 8.465 | Oligo |
| 51 | Litaf | 1.38E-65 | 8.792407 | Oligo |
| 52 | Arhgef10 | 3.13E-65 | 7.071656 | Oligo |
| 53 | Rnf13 | 7.21E-65 | 2.7044854 | Oligo |
| 54 | Myrf | 7.81E-65 | 9.786589 | Oligo |
| 55 | Cryab | 1.31E-64 | 6.8142548 | Oligo |
| 56 | Prdx1 | 5.30E-64 | 1.6615056 | Oligo |
| 57 | Slc44a1 | 6.03E-63 | 5.5968328 | Oligo |
| 58 | Dock10 | 1.89E-62 | 6.2039742 | Oligo |
| 59 | Serinc5 | 5.35E-62 | 5.624491 | Oligo |
| 60 | Trf | 2.48E-61 | 11.204493 | Oligo |
| 61 | Ddr1 | 5.39E-61 | 5.165238 | Oligo |
| 62 | Ttyh2 | 2.02E-60 | 8.088001 | Oligo |
| 63 | 1700047M11Rik | 5.36E-60 | 10.364959 | Oligo |
| 64 | Mag | 1.61E-59 | 11.21688 | Oligo |
| 65 | Gm4617 | 2.62E-59 | 2.223879 | Oligo |
| 66 | Gpm6b | 2.78E-59 | 2.391333 | Oligo |
| 67 | Tmbim6 | 3.65E-59 | 1.7270118 | Oligo |
| 68 | Gsn | 4.26E-59 | 7.239747 | Oligo |
| 69 | Cldn11 | 5.28E-59 | 11.61154 | Oligo |
| 70 | Tmem88b | 1.28E-57 | 8.393133 | Oligo |
| 71 | Csrp1 | 1.61E-57 | 7.2529263 | Oligo |
| 72 | Hist1h2bc | 2.31E-57 | 4.8006525 | Oligo |
| 73 | Ftl2 | 3.25E-57 | 1.7199504 | Oligo |
| 74 | Rab31 | 5.26E-57 | 5.145797 | Oligo |
| 75 | Tnfaip6 | 7.59E-57 | 8.086938 | Oligo |
| 76 | Ermn | 7.90E-57 | 11.286227 | Oligo |
| 77 | Gm7204 | 8.13E-57 | 2.1797054 | Oligo |
| 78 | Mog | 1.15E-56 | 11.216276 | Oligo |
| 79 | Gab1 | 3.75E-56 | 7.8716874 | Oligo |
| 80 | Nkx6-2 | 4.32E-56 | 10.256294 | Oligo |
| 81 | Taldo1 | 8.57E-56 | 2.4229255 | Oligo |
| 82 | Pla2g16 | 1.39E-55 | 8.733156 | Oligo |
| 83 | Gm21399 | 2.00E-55 | 2.2185593 | Oligo |
| 84 | Sec11c | 3.18E-55 | 1.907734 | Oligo |
| 85 | Apod | 3.18E-55 | 9.685853 | Oligo |
| 86 | Plxnb3 | 1.49E-54 | 9.785857 | Oligo |
| 87 | Degs1 | 1.99E-54 | 1.8492707 | Oligo |

| | | | | |
|---|---|---|---|---|
| 88 | CYTB | 1.03E-53 | 0.90048695 | Oligo |
| 89 | Carhsp1 | 1.05E-53 | 7.391201 | Oligo |
| 90 | Josd2 | 1.36E-53 | 3.2506607 | Oligo |
| 91 | Opalin | 1.08E-52 | 11.56505 | Oligo |
| 92 | Gng11 | 1.23E-51 | 8.534472 | Oligo |
| 93 | Mt1 | 1.49E-51 | 3.3144164 | Oligo |
| 94 | Tspan2 | 1.53E-51 | 6.6108418 | Oligo |
| 95 | Mal | 1.61E-51 | 11.118335 | Oligo |
| 96 | Tspan15 | 3.01E-51 | 7.5392327 | Oligo |
| 97 | Gpr37 | 4.16E-51 | 8.278544 | Oligo |
| 98 | Ptn | 5.84E-51 | 4.426413 | Oligo |
| 99 | Phgdh | 9.25E-51 | 6.136504 | Oligo |
| 0 | Atp13a5 | 1.34E-18 | 15.646597 | Peri |
| 1 | Higd1b | 1.34E-18 | 15.815601 | Peri |
| 2 | Vtn | 1.34E-18 | 17.827215 | Peri |
| 3 | Ndufa4l2 | 1.34E-18 | 13.687341 | Peri |
| 4 | Ifitm1 | 1.34E-18 | 14.504967 | Peri |
| 5 | Rgs5 | 1.34E-18 | 13.001669 | Peri |
| 6 | Igfbp7 | 1.50E-18 | 12.261171 | Peri |
| 7 | Cald1 | 1.50E-18 | 11.956551 | Peri |
| 8 | Ifitm2 | 2.05E-18 | 10.805882 | Peri |
| 9 | Myl9 | 2.05E-18 | 14.9197855 | Peri |
| 10 | Laptm4a | 2.35E-18 | 3.5728335 | Peri |
| 11 | Itm2b | 2.73E-18 | 1.8922306 | Peri |
| 12 | Ifitm3 | 3.70E-18 | 12.60828 | Peri |
| 13 | Epas1 | 6.49E-18 | 10.6080675 | Peri |
| 14 | Cox4i2 | 6.62E-18 | 12.84759 | Peri |
| 15 | Sod3 | 6.66E-18 | 14.135922 | Peri |
| 16 | Atp1a2 | 6.83E-18 | 11.60475 | Peri |
| 17 | Art3 | 6.83E-18 | 13.207199 | Peri |
| 18 | Sparc | 6.83E-18 | 11.926032 | Peri |
| 19 | Pdgfrb | 7.08E-18 | 11.941524 | Peri |
| 20 | Hspb1 | 9.22E-18 | 12.734727 | Peri |
| 21 | Hsp25-ps1 | 9.34E-18 | 8.551531 | Peri |
| 22 | Mfge8 | 1.03E-17 | 11.652906 | Peri |
| 23 | Mgp | 1.06E-17 | 11.577969 | Peri |
| 24 | Nbl1 | 1.29E-17 | 8.108578 | Peri |
| 25 | Ptn | 1.58E-17 | 6.98202 | Peri |
| 26 | Kcnj8 | 2.08E-17 | 16.273169 | Peri |
| 27 | Rarres2 | 2.54E-17 | 12.127863 | Peri |
| 28 | Gatm | 2.60E-17 | 9.1583805 | Peri |
| 29 | Myl12a | 2.67E-17 | 6.9707894 | Peri |
| 30 | Gng11 | 3.05E-17 | 12.397197 | Peri |
| 31 | 04-Sep | 3.73E-17 | 5.4964533 | Peri |
| 32 | Cst3 | 4.46E-17 | 2.9067008 | Peri |
| 33 | Ppp1r14a | 5.17E-17 | 11.002545 | Peri |
| 34 | Rbpms | 5.53E-17 | 10.621697 | Peri |
| 35 | H3f3b | 6.45E-17 | 1.8897086 | Peri |
| 36 | Pla1a | 1.29E-16 | 13.931993 | Peri |
| 37 | Ackr3 | 1.57E-16 | 9.587221 | Peri |

| | | | | |
|---|---|---|---|---|
| 38 | Tm4sf1 | 2.20E-16 | 12.998775 | Peri |
| 39 | Plat | 2.23E-16 | 8.203257 | Peri |
| 40 | P2ry14 | 2.62E-16 | 12.97576 | Peri |
| 41 | Tagln2 | 2.98E-16 | 11.736747 | Peri |
| 42 | Coro1b | 4.81E-16 | 4.6657624 | Peri |
| 43 | Pitpnc1 | 9.96E-16 | 5.807191 | Peri |
| 44 | Abcc9 | 1.22E-15 | 14.091102 | Peri |
| 45 | Tmsb4x | 1.28E-15 | 1.8740828 | Peri |
| 46 | Slc6a20a | 1.79E-15 | 13.568853 | Peri |
| 47 | Slc38a11 | 1.81E-15 | 13.103518 | Peri |
| 48 | Cp | 1.91E-15 | 9.744163 | Peri |
| 49 | Adap2 | 2.06E-15 | 10.63662 | Peri |
| 50 | Lamc1 | 2.85E-15 | 7.1344004 | Peri |
| 51 | Colec12 | 2.88E-15 | 11.22468 | Peri |
| 52 | Gjc1 | 3.26E-15 | 10.571155 | Peri |
| 53 | Cd63 | 3.47E-15 | 7.1154685 | Peri |
| 54 | Axl | 4.79E-15 | 9.871382 | Peri |
| 55 | Serinc3 | 6.79E-15 | 2.3972316 | Peri |
| 56 | Cgnl1 | 7.70E-15 | 8.778164 | Peri |
| 57 | Bloc1s1 | 8.44E-15 | 2.4254847 | Peri |
| 58 | S100a11 | 1.16E-14 | 8.966038 | Peri |
| 59 | Tmem204 | 1.18E-14 | 9.657501 | Peri |
| 60 | Atp2a3 | 1.51E-14 | 11.989381 | Peri |
| 61 | Itga1 | 2.40E-14 | 11.015883 | Peri |
| 62 | Ftl1 | 2.54E-14 | 1.6315701 | Peri |
| 63 | Zak | 2.72E-14 | 7.210902 | Peri |
| 64 | Gng5 | 3.35E-14 | 6.164835 | Peri |
| 65 | Serpinh1 | 3.42E-14 | 10.329387 | Peri |
| 66 | Cd81 | 3.58E-14 | 1.44096 | Peri |
| 67 | Zfp36l1 | 4.19E-14 | 8.426231 | Peri |
| 68 | Gm7676 | 4.24E-14 | 6.852838 | Peri |
| 69 | Malat1 | 4.36E-14 | 1.7364156 | Peri |
| 70 | Apold1 | 5.88E-14 | 8.311449 | Peri |
| 71 | Dmd | 9.33E-14 | 3.9166973 | Peri |
| 72 | S1pr3 | 9.88E-14 | 8.085825 | Peri |
| 73 | 11-Sep | 1.11E-13 | 2.4151535 | Peri |
| 74 | Filip1l | 1.65E-13 | 9.085829 | Peri |
| 75 | Gm13861 | 2.12E-13 | 10.858261 | Peri |
| 76 | Phlda1 | 2.12E-13 | 4.4551716 | Peri |
| 77 | Tbx3 | 2.45E-13 | 8.919156 | Peri |
| 78 | Cd63-ps | 2.64E-13 | 4.003284 | Peri |
| 79 | Jund | 2.83E-13 | 2.8120682 | Peri |
| 80 | Car4 | 4.40E-13 | 7.0784082 | Peri |
| 81 | Myo1b | 6.86E-13 | 6.901028 | Peri |
| 82 | B2m | 7.58E-13 | 2.8889487 | Peri |
| 83 | Anxa5 | 8.31E-13 | 5.6812253 | Peri |
| 84 | Hes1 | 9.58E-13 | 5.6485085 | Peri |
| 85 | Ace2 | 9.68E-13 | 13.857953 | Peri |
| 86 | Tnfaip1 | 9.90E-13 | 4.3695345 | Peri |
| 87 | Enpep | 1.12E-12 | 12.4357 | Peri |

| | | | | |
|---|---|---|---|---|
| 88 | Ggt5 | 1.17E-12 | 11.723578 | Peri |
| 89 | Mylk | 1.31E-12 | 7.4377866 | Peri |
| 90 | Aspn | 1.54E-12 | 12.36647 | Peri |
| 91 | Prdx1 | 1.61E-12 | 1.9011979 | Peri |
| 92 | Sdc2 | 1.83E-12 | 5.044868 | Peri |
| 93 | Rgs4 | 2.33E-12 | 4.5940127 | Peri |
| 94 | Dock6 | 2.91E-12 | 7.8282566 | Peri |
| 95 | Gucy1b3 | 3.09E-12 | 3.3128016 | Peri |
| 96 | Zbtb20 | 3.13E-12 | 6.9049034 | Peri |
| 97 | Ece1 | 3.81E-12 | 4.553116 | Peri |
| 98 | Arhgdib | 4.67E-12 | 8.300804 | Peri |
| 99 | Actb | 4.91E-12 | 1.0784159 | Peri |
| 0 | Pvalb | 0 | 11.412964 | Pvalb |
| 1 | Cntnap4 | 0 | 5.4028163 | Pvalb |
| 2 | Cacna2d2 | 0 | 4.8413944 | Pvalb |
| 3 | Man1c1 | 0 | 4.8558846 | Pvalb |
| 4 | Paip2 | 0 | 1.3168322 | Pvalb |
| 5 | Ndufv2 | 0 | 0.9770267 | Pvalb |
| 6 | Tmx2 | 0 | 1.2503897 | Pvalb |
| 7 | Atp5o | 0 | 1.010557 | Pvalb |
| 8 | Sncb | 0 | 1.425069 | Pvalb |
| 9 | Asns | 0 | 2.168752 | Pvalb |
| 10 | Cox7b | 0 | 1.1023136 | Pvalb |
| 11 | Ndufb5 | 0 | 1.182743 | Pvalb |
| 12 | Pacsin2 | 0 | 4.2558975 | Pvalb |
| 13 | Uqcrh | 0 | 1.0650527 | Pvalb |
| 14 | Ndufs2 | 0 | 1.3021389 | Pvalb |
| 15 | Atp5f1 | 0 | 0.9383399 | Pvalb |
| 16 | Esrrg | 0 | 4.5512934 | Pvalb |
| 17 | Laptm4b | 0 | 2.912713 | Pvalb |
| 18 | Btbd11 | 0 | 5.8749857 | Pvalb |
| 19 | Cemip | 0 | 5.799569 | Pvalb |
| 20 | Uqcrc1 | 0 | 1.1448073 | Pvalb |
| 21 | Ankrd29 | 0 | 4.4939084 | Pvalb |
| 22 | Kcnk3 | 0 | 4.145286 | Pvalb |
| 23 | Timm8a1 | 0 | 3.02652 | Pvalb |
| 24 | Slc6a1 | 0 | 6.5355844 | Pvalb |
| 25 | Nars | 0 | 1.4610684 | Pvalb |
| 26 | Rcan2 | 0 | 3.3255317 | Pvalb |
| 27 | Cox4i1 | 0 | 0.7580069 | Pvalb |
| 28 | Glrx5 | 0 | 1.5897782 | Pvalb |
| 29 | Vdac2 | 0 | 0.9563972 | Pvalb |
| 30 | Nxph1 | 0 | 7.631834 | Pvalb |
| 31 | Prss23 | 0 | 6.5979805 | Pvalb |
| 32 | Cox6b1 | 0 | 0.8800954 | Pvalb |
| 33 | Idh3b | 0 | 1.0618836 | Pvalb |
| 34 | Sparcl1 | 0 | 2.966286 | Pvalb |
| 35 | Fam210b | 0 | 4.8281426 | Pvalb |
| 36 | Zfp385a | 0 | 3.3059947 | Pvalb |
| 37 | Ghitm | 0 | 1.3449569 | Pvalb |

| | | | | |
|---|---|---|---|---|
| 38 | Flt3 | 0 | 5.043082 | Pvalb |
| 39 | Ank1 | 0 | 4.387715 | Pvalb |
| 40 | Kcns3 | 0 | 6.1372137 | Pvalb |
| 41 | Mdh2 | 0 | 0.9051448 | Pvalb |
| 42 | Idh3a | 0 | 1.3312167 | Pvalb |
| 43 | Ppp1cc | 0 | 1.9394597 | Pvalb |
| 44 | Ndufa8 | 0 | 0.9831678 | Pvalb |
| 45 | Ndufb8 | 0 | 0.96302265 | Pvalb |
| 46 | Lpl | 0 | 7.1511154 | Pvalb |
| 47 | Gapdh | 0 | 0.6057199 | Pvalb |
| 48 | Gpr176 | 0 | 5.898357 | Pvalb |
| 49 | Cartpt | 0 | 7.582122 | Pvalb |
| 50 | Scn1a | 0 | 2.8121161 | Pvalb |
| 51 | Atp5g3 | 0 | 1.001828 | Pvalb |
| 52 | Atp5b | 0 | 0.9572388 | Pvalb |
| 53 | Slc25a3 | 0 | 0.91930234 | Pvalb |
| 54 | Slc25a5 | 0 | 1.3076682 | Pvalb |
| 55 | Atp5c1 | 0 | 1.0632843 | Pvalb |
| 56 | Cyc1 | 0 | 1.2636374 | Pvalb |
| 57 | Slc25a4 | 0 | 0.8674748 | Pvalb |
| 58 | Fndc5 | 0 | 3.6832056 | Pvalb |
| 59 | Cend1 | 0 | 1.9103914 | Pvalb |
| 60 | Atp1b1 | 0 | 1.6954659 | Pvalb |
| 61 | Nek7 | 0 | 8.044111 | Pvalb |
| 62 | Rnd2 | 0 | 5.9987664 | Pvalb |
| 63 | Kcnc1 | 0 | 4.4095645 | Pvalb |
| 64 | Gm13629 | 0 | 6.6600013 | Pvalb |
| 65 | Kcnc2 | 0 | 5.4821777 | Pvalb |
| 66 | Ldhb | 0 | 1.6759908 | Pvalb |
| 67 | Mdh1 | 0 | 1.3657168 | Pvalb |
| 68 | Phlda1 | 0 | 4.730376 | Pvalb |
| 69 | Got1 | 0 | 2.009418 | Pvalb |
| 70 | Cox6a2 | 0 | 8.997437 | Pvalb |
| 71 | Cplx1 | 0 | 3.4788973 | Pvalb |
| 72 | Tac1 | 0 | 11.64089 | Pvalb |
| 73 | Kcnab3 | 0 | 5.30424 | Pvalb |
| 74 | Ifitm10 | 0 | 4.6526465 | Pvalb |
| 75 | Gm5514 | 0 | 1.5910362 | Pvalb |
| 76 | Cox5a | 0 | 1.1373028 | Pvalb |
| 77 | Cycs | 0 | 1.02148 | Pvalb |
| 78 | Sox6 | 0 | 6.643503 | Pvalb |
| 79 | Tmem132c | 0 | 6.715785 | Pvalb |
| 80 | Uqcrfs1 | 0 | 1.1597111 | Pvalb |
| 81 | Hcn1 | 0 | 3.605839 | Pvalb |
| 82 | Lhx6 | 0 | 6.8387113 | Pvalb |
| 83 | Ndufa6 | 0 | 1.1254321 | Pvalb |
| 84 | Atp5a1 | 0 | 0.9876362 | Pvalb |
| 85 | Inpp5j | 0 | 4.71558 | Pvalb |
| 86 | Ndrg4 | 0 | 1.6049248 | Pvalb |
| 87 | Sars | 0 | 1.6059798 | Pvalb |

| | | | | |
|---|---|---|---|---|
| 88 | Pcp4l1 | 0 | 6.8147345 | Pvalb |
| 89 | Nog | 0 | 5.8517566 | Pvalb |
| 90 | St3gal6 | 0 | 6.4534926 | Pvalb |
| 91 | Selm | 0 | 1.7158242 | Pvalb |
| 92 | Ndufa12 | 0 | 1.2476192 | Pvalb |
| 93 | Atp5j | 0 | 1.0228034 | Pvalb |
| 94 | Fgf12 | 0 | 2.259502 | Pvalb |
| 95 | Tm6sf1 | 0 | 4.7600203 | Pvalb |
| 96 | Erbb4 | 0 | 7.1341734 | Pvalb |
| 97 | Gabrd | 0 | 4.464662 | Pvalb |
| 98 | Chchd10 | 0 | 1.4043326 | Pvalb |
| 99 | Snhg4 | 0 | 3.922948 | Pvalb |
| 0 | Tagln | 7.35E-69 | 16.673727 | SMC |
| 1 | Tpm2 | 7.35E-69 | 13.4696045 | SMC |
| 2 | Myl6 | 7.35E-69 | 3.7909694 | SMC |
| 3 | Myl9 | 7.35E-69 | 17.443481 | SMC |
| 4 | Acta2 | 7.35E-69 | 17.129553 | SMC |
| 5 | Gm5526 | 7.35E-69 | 4.1908774 | SMC |
| 6 | Myh11 | 7.35E-69 | 15.961194 | SMC |
| 7 | Tpm1 | 7.35E-69 | 5.7830014 | SMC |
| 8 | Crip1 | 7.35E-69 | 16.591537 | SMC |
| 9 | Mustn1 | 7.35E-69 | 12.075568 | SMC |
| 10 | Wtip | 7.35E-69 | 10.84344 | SMC |
| 11 | Vim | 8.17E-69 | 13.402409 | SMC |
| 12 | Filip1l | 8.17E-69 | 13.215634 | SMC |
| 13 | Cald1 | 8.17E-69 | 12.172344 | SMC |
| 14 | Ptrf | 9.10E-69 | 12.694455 | SMC |
| 15 | Gm10080 | 9.10E-69 | 3.857819 | SMC |
| 16 | Zak | 9.93E-69 | 10.436342 | SMC |
| 17 | Igfbp7 | 1.10E-68 | 12.283396 | SMC |
| 18 | Fxyd1 | 1.10E-68 | 8.875441 | SMC |
| 19 | S100a11 | 1.14E-68 | 12.036873 | SMC |
| 20 | Prkcdbp | 1.72E-68 | 7.1906853 | SMC |
| 21 | Dstn | 2.18E-68 | 4.021543 | SMC |
| 22 | Csrp1 | 2.24E-68 | 10.833484 | SMC |
| 23 | Gm13889 | 3.32E-68 | 6.841754 | SMC |
| 24 | Zfhx3 | 6.58E-68 | 10.235973 | SMC |
| 25 | Hspb1 | 7.72E-68 | 13.2400055 | SMC |
| 26 | Mylk | 8.54E-68 | 12.097872 | SMC |
| 27 | Sncg | 8.54E-68 | 14.105173 | SMC |
| 28 | Hsp25-ps1 | 9.25E-68 | 8.872146 | SMC |
| 29 | Gm7665 | 1.87E-67 | 8.732703 | SMC |
| 30 | Rbpms | 1.87E-67 | 12.544336 | SMC |
| 31 | Sparcl1 | 2.34E-67 | 4.79152 | SMC |
| 32 | Actb | 2.55E-67 | 2.2950726 | SMC |
| 33 | Gng11 | 3.85E-67 | 12.304293 | SMC |
| 34 | Rras | 7.13E-67 | 9.642382 | SMC |
| 35 | Epas1 | 7.13E-67 | 11.451254 | SMC |
| 36 | Mgp | 9.21E-67 | 11.448852 | SMC |
| 37 | Ppp1r14a | 1.63E-66 | 11.9873 | SMC |

| | | | | |
|---|---|---|---|---|
| 38 | Flna | 1.80E-66 | 9.804313 | SMC |
| 39 | Hspb2 | 2.09E-66 | 11.426714 | SMC |
| 40 | CYTB | 2.95E-66 | 2.0737095 | SMC |
| 41 | Ifitm2 | 3.64E-66 | 9.941287 | SMC |
| 42 | Pdlim3 | 4.14E-66 | 12.044357 | SMC |
| 43 | LOC102635661 | 7.78E-66 | 4.0319343 | SMC |
| 44 | Fth1 | 7.38E-65 | 1.7874249 | SMC |
| 45 | Sparc | 7.38E-65 | 12.134938 | SMC |
| 46 | Mfge8 | 9.82E-65 | 12.4172 | SMC |
| 47 | Gm8894 | 1.11E-64 | 3.9604683 | SMC |
| 48 | Pfn1 | 1.19E-64 | 2.2093554 | SMC |
| 49 | Map3k7cl | 1.85E-64 | 12.80868 | SMC |
| 50 | LOC102637129 | 2.79E-64 | 6.817258 | SMC |
| 51 | Tagln2 | 7.21E-64 | 11.782239 | SMC |
| 52 | Mcam | 8.51E-64 | 12.741971 | SMC |
| 53 | Pln | 8.88E-64 | 12.368128 | SMC |
| 54 | Crip2 | 1.58E-63 | 4.8298798 | SMC |
| 55 | Ifitm3 | 1.97E-63 | 11.857956 | SMC |
| 56 | Ppp1r12a | 2.10E-63 | 3.5896413 | SMC |
| 57 | Phldb2 | 3.15E-63 | 11.392613 | SMC |
| 58 | Aspn | 3.43E-63 | 15.086963 | SMC |
| 59 | Bcam | 3.43E-63 | 11.017034 | SMC |
| 60 | Slc38a11 | 3.68E-63 | 14.616942 | SMC |
| 61 | Lmod1 | 4.78E-63 | 11.876171 | SMC |
| 62 | Tinagl1 | 2.22E-62 | 12.328762 | SMC |
| 63 | Timp3 | 2.96E-62 | 9.416485 | SMC |
| 64 | Des | 4.19E-62 | 14.058633 | SMC |
| 65 | H3f3a | 4.22E-62 | 2.0667083 | SMC |
| 66 | Ahnak | 4.39E-62 | 9.2602825 | SMC |
| 67 | Myl12a | 5.09E-62 | 6.1725845 | SMC |
| 68 | Gm13861 | 5.72E-62 | 12.681709 | SMC |
| 69 | Gng5 | 9.98E-62 | 6.6077538 | SMC |
| 70 | Rgs5 | 1.07E-61 | 10.656623 | SMC |
| 71 | Cnn2 | 1.82E-61 | 11.067461 | SMC |
| 72 | Pde3a | 4.08E-61 | 11.014592 | SMC |
| 73 | Tns1 | 4.49E-61 | 8.682143 | SMC |
| 74 | Serping1 | 4.55E-61 | 12.708363 | SMC |
| 75 | Cd9 | 9.51E-61 | 12.022409 | SMC |
| 76 | Tgfb1i1 | 1.45E-60 | 7.0902166 | SMC |
| 77 | ND4 | 1.49E-60 | 1.9375067 | SMC |
| 78 | Gja4 | 1.94E-60 | 10.849408 | SMC |
| 79 | Lgals1 | 2.49E-60 | 9.353406 | SMC |
| 80 | Utrn | 4.12E-60 | 6.2891097 | SMC |
| 81 | COX1 | 4.72E-60 | 1.6304629 | SMC |
| 82 | Rsu1 | 6.40E-60 | 5.998413 | SMC |
| 83 | Mir143hg | 6.53E-60 | 13.046372 | SMC |
| 84 | Itgb1 | 9.24E-60 | 7.0914645 | SMC |
| 85 | Crim1 | 1.14E-59 | 7.188181 | SMC |
| 86 | Cryab | 1.14E-59 | 6.7462883 | SMC |
| 87 | Rbpms2 | 1.25E-59 | 12.096452 | SMC |

| | | | | |
|---|---|---|---|---|
| 88 | Notch3 | 1.78E-59 | 8.690692 | SMC |
| 89 | Cst3 | 9.65E-59 | 2.5842342 | SMC |
| 90 | Palld | 1.12E-58 | 9.372497 | SMC |
| 91 | Lamb2 | 1.25E-58 | 10.751229 | SMC |
| 92 | Rap1a | 1.62E-58 | 3.5244954 | SMC |
| 93 | Tm4sf1 | 1.97E-58 | 11.977773 | SMC |
| 94 | Ybx1 | 2.06E-58 | 2.5022845 | SMC |
| 95 | Snhg18 | 2.45E-58 | 10.978345 | SMC |
| 96 | Rhoc | 5.54E-58 | 8.646476 | SMC |
| 97 | Tuba1c | 8.29E-58 | 8.815059 | SMC |
| 98 | Prss23 | 1.12E-57 | 8.296174 | SMC |
| 99 | Mob2 | 1.32E-57 | 4.1920724 | SMC |
| 0 | Nov | 1.05E-51 | 8.310271 | Serpinf1 |
| 1 | Gsto1 | 3.77E-51 | 3.7747304 | Serpinf1 |
| 2 | Kcnip1 | 4.65E-48 | 7.0731363 | Serpinf1 |
| 3 | Enpp2 | 6.43E-48 | 7.747691 | Serpinf1 |
| 4 | Cnr1 | 5.62E-46 | 5.707516 | Serpinf1 |
| 5 | Ndufa13 | 1.26E-43 | 1.5207275 | Serpinf1 |
| 6 | Necab2 | 1.39E-42 | 6.6296377 | Serpinf1 |
| 7 | Elmo1 | 3.10E-42 | 4.182321 | Serpinf1 |
| 8 | Ap1s2 | 5.01E-42 | 5.4657774 | Serpinf1 |
| 9 | E530001K10Rik | 5.11E-41 | 6.1579785 | Serpinf1 |
| 10 | Sp8 | 6.43E-41 | 7.44547 | Serpinf1 |
| 11 | Nt5dc2 | 1.13E-39 | 7.5590425 | Serpinf1 |
| 12 | Adarb2 | 1.47E-39 | 8.049888 | Serpinf1 |
| 13 | Dner | 1.72E-39 | 4.786816 | Serpinf1 |
| 14 | Mt1 | 2.89E-39 | 4.4182124 | Serpinf1 |
| 15 | Tcerg1l | 1.52E-38 | 6.239918 | Serpinf1 |
| 16 | Cntn4 | 3.27E-38 | 4.4049807 | Serpinf1 |
| 17 | Camk2d | 5.23E-38 | 3.7750745 | Serpinf1 |
| 18 | Rgs8 | 5.23E-38 | 5.596217 | Serpinf1 |
| 19 | Luzp2 | 5.10E-37 | 6.592258 | Serpinf1 |
| 20 | Lppr1 | 7.26E-37 | 5.674209 | Serpinf1 |
| 21 | 2900055J20Rik | 1.19E-36 | 3.0469012 | Serpinf1 |
| 22 | Slc44a5 | 2.19E-36 | 6.016938 | Serpinf1 |
| 23 | Ogfrl1 | 3.85E-36 | 2.6980255 | Serpinf1 |
| 24 | Pcdh20 | 6.27E-36 | 6.7901483 | Serpinf1 |
| 25 | Nr3c2 | 1.03E-35 | 4.351824 | Serpinf1 |
| 26 | Tnfaip8l3 | 1.51E-35 | 7.5138364 | Serpinf1 |
| 27 | Cacng5 | 1.93E-35 | 6.7567105 | Serpinf1 |
| 28 | Cck | 2.66E-35 | 5.588516 | Serpinf1 |
| 29 | Pnoc | 9.94E-35 | 7.5362105 | Serpinf1 |
| 30 | Tcf4 | 3.03E-34 | 1.7789638 | Serpinf1 |
| 31 | Zbtb20 | 3.75E-34 | 6.2328196 | Serpinf1 |
| 32 | Shisa9 | 7.95E-34 | 4.3788056 | Serpinf1 |
| 33 | Dlx1as | 1.01E-33 | 5.662262 | Serpinf1 |
| 34 | Mt2 | 1.12E-33 | 6.2514496 | Serpinf1 |
| 35 | Rgs7bp | 2.12E-33 | 1.9503092 | Serpinf1 |
| 36 | Alcam | 2.69E-33 | 4.086992 | Serpinf1 |
| 37 | Sema3c | 2.01E-32 | 6.5970445 | Serpinf1 |

| | | | | |
|---|---|---|---|---|
| 38 | Igf1 | 2.10E-32 | 6.70748 | Serpinf1 |
| 39 | Rgs16 | 3.56E-32 | 5.499942 | Serpinf1 |
| 40 | Magi1 | 3.66E-32 | 2.6930854 | Serpinf1 |
| 41 | Fhl1 | 3.66E-32 | 3.615399 | Serpinf1 |
| 42 | Gnao1 | 4.74E-32 | 1.4761177 | Serpinf1 |
| 43 | Rgs10 | 4.95E-32 | 5.523548 | Serpinf1 |
| 44 | Dlx1 | 5.51E-32 | 5.384212 | Serpinf1 |
| 45 | Gpx4 | 1.07E-31 | 0.8673704 | Serpinf1 |
| 46 | Serpinf1 | 2.22E-31 | 8.837185 | Serpinf1 |
| 47 | Tuba8 | 5.85E-31 | 5.078621 | Serpinf1 |
| 48 | Cpne7 | 5.86E-31 | 5.5529075 | Serpinf1 |
| 49 | Qpct | 1.53E-30 | 5.0907016 | Serpinf1 |
| 50 | Kcnip4 | 3.76E-30 | 3.3963292 | Serpinf1 |
| 51 | Cd24a | 4.57E-30 | 7.3113456 | Serpinf1 |
| 52 | Plekhg1 | 4.78E-30 | 6.465808 | Serpinf1 |
| 53 | Ppp1r2 | 5.32E-30 | 1.9742956 | Serpinf1 |
| 54 | Dlx6os1 | 6.09E-30 | 6.3231993 | Serpinf1 |
| 55 | Gpr12 | 6.48E-30 | 3.8948982 | Serpinf1 |
| 56 | Gng4 | 7.96E-30 | 3.9904504 | Serpinf1 |
| 57 | Prox1 | 1.05E-29 | 5.8140807 | Serpinf1 |
| 58 | Enho | 1.11E-29 | 2.8154373 | Serpinf1 |
| 59 | Maf | 1.29E-29 | 4.8208103 | Serpinf1 |
| 60 | Kctd16 | 1.39E-29 | 3.850575 | Serpinf1 |
| 61 | Cplx2 | 1.49E-29 | 3.38434 | Serpinf1 |
| 62 | Eps8 | 1.49E-29 | 5.592486 | Serpinf1 |
| 63 | Sox1 | 1.64E-29 | 4.9838405 | Serpinf1 |
| 64 | Zbtb16 | 1.67E-29 | 2.8861082 | Serpinf1 |
| 65 | Scg3 | 2.10E-29 | 1.9189631 | Serpinf1 |
| 66 | Gad1 | 2.19E-29 | 7.371929 | Serpinf1 |
| 67 | Npy | 3.36E-29 | 8.191606 | Serpinf1 |
| 68 | Gm17750 | 4.37E-29 | 5.3965034 | Serpinf1 |
| 69 | Ccnh | 5.81E-29 | 1.4838929 | Serpinf1 |
| 70 | Htra1 | 1.01E-28 | 5.366123 | Serpinf1 |
| 71 | Nrxn3 | 1.09E-28 | 2.2349453 | Serpinf1 |
| 72 | Dhx32 | 2.55E-28 | 3.738796 | Serpinf1 |
| 73 | Bcat1 | 3.94E-28 | 3.273018 | Serpinf1 |
| 74 | Nrip3 | 5.58E-28 | 4.0924745 | Serpinf1 |
| 75 | A530058N18Rik | 7.03E-28 | 4.8862677 | Serpinf1 |
| 76 | Larp1b | 8.05E-28 | 4.0258965 | Serpinf1 |
| 77 | Mgat5b | 8.30E-28 | 4.224172 | Serpinf1 |
| 78 | Sntg1 | 1.31E-27 | 3.2298872 | Serpinf1 |
| 79 | Slc24a3 | 1.44E-27 | 3.7716458 | Serpinf1 |
| 80 | Sez6 | 1.85E-27 | 3.6705787 | Serpinf1 |
| 81 | Resp18 | 3.33E-27 | 2.7421052 | Serpinf1 |
| 82 | Aplp2 | 3.61E-27 | 1.2117838 | Serpinf1 |
| 83 | Scg2 | 1.14E-26 | 2.8879945 | Serpinf1 |
| 84 | Rab3c | 1.28E-26 | 2.4706454 | Serpinf1 |
| 85 | Rasgrf2 | 1.61E-26 | 4.193126 | Serpinf1 |
| 86 | G0s2 | 1.96E-26 | 4.56149 | Serpinf1 |
| 87 | Sh3bgrl | 2.29E-26 | 3.1158211 | Serpinf1 |

| | | | | |
|---|---|---|---|---|
| 88 | Cnn3 | 2.50E-26 | 4.563603 | Serpinf1 |
| 89 | Vstm2a | 3.11E-26 | 2.7984548 | Serpinf1 |
| 90 | Tmem55a | 3.11E-26 | 2.178145 | Serpinf1 |
| 91 | Kcnn2 | 3.17E-26 | 3.5403988 | Serpinf1 |
| 92 | Shisa8 | 4.12E-26 | 5.9787793 | Serpinf1 |
| 93 | Zfp536 | 6.39E-26 | 4.851638 | Serpinf1 |
| 94 | Gng2 | 8.80E-26 | 2.0522594 | Serpinf1 |
| 95 | Car2 | 9.52E-26 | 5.5890226 | Serpinf1 |
| 96 | Fgf1 | 2.43E-25 | 5.328331 | Serpinf1 |
| 97 | Fam163a | 3.98E-25 | 5.4399223 | Serpinf1 |
| 98 | Frmd7 | 4.82E-25 | 7.58713 | Serpinf1 |
| 99 | Fam174a | 1.12E-24 | 1.5829947 | Serpinf1 |
| 0 | Cnr1 | 2.14E-156 | 6.956153 | Sncg |
| 1 | Lppr1 | 1.68E-148 | 6.8850727 | Sncg |
| 2 | Npas1 | 7.37E-146 | 8.542158 | Sncg |
| 3 | Gng2 | 2.94E-145 | 3.0717432 | Sncg |
| 4 | Kcnip1 | 2.66E-135 | 6.783791 | Sncg |
| 5 | Nrxn3 | 1.62E-133 | 2.8337567 | Sncg |
| 6 | Sncg | 6.10E-131 | 11.537137 | Sncg |
| 7 | Adarb2 | 2.45E-130 | 8.458506 | Sncg |
| 8 | Necab1 | 5.84E-130 | 4.719888 | Sncg |
| 9 | Kctd12 | 6.63E-128 | 7.062844 | Sncg |
| 10 | Cxcl14 | 1.36E-125 | 9.970744 | Sncg |
| 11 | Sema3c | 1.33E-124 | 7.674816 | Sncg |
| 12 | Dlx1as | 1.70E-124 | 6.6092815 | Sncg |
| 13 | Enho | 2.05E-121 | 3.4557428 | Sncg |
| 14 | Egln3 | 6.56E-121 | 7.4374213 | Sncg |
| 15 | Tnfaip8l3 | 1.58E-120 | 8.504573 | Sncg |
| 16 | Gpx4 | 2.67E-120 | 1.1183695 | Sncg |
| 17 | Cplx2 | 2.40E-119 | 3.826661 | Sncg |
| 18 | Qpct | 4.38E-119 | 5.733985 | Sncg |
| 19 | Htr3a | 5.58E-118 | 9.635294 | Sncg |
| 20 | Npas3 | 1.55E-116 | 6.145454 | Sncg |
| 21 | Dlx6os1 | 6.96E-116 | 7.1889434 | Sncg |
| 22 | Nrip3 | 2.54E-114 | 4.624327 | Sncg |
| 23 | Sp8 | 2.87E-114 | 5.9849467 | Sncg |
| 24 | Elmod1 | 1.15E-113 | 2.325995 | Sncg |
| 25 | Celf6 | 1.15E-113 | 4.325639 | Sncg |
| 26 | Shisa9 | 1.05E-112 | 4.5850983 | Sncg |
| 27 | Jam2 | 2.73E-112 | 7.1553693 | Sncg |
| 28 | Rwdd3 | 2.73E-112 | 5.5594616 | Sncg |
| 29 | Rgs12 | 8.43E-112 | 7.6562557 | Sncg |
| 30 | 2610001J05Rik | 9.20E-112 | 2.2048774 | Sncg |
| 31 | Fxyd6 | 3.66E-111 | 4.926269 | Sncg |
| 32 | Gng4 | 6.38E-109 | 4.585572 | Sncg |
| 33 | Nap1l5 | 1.02E-108 | 2.2106652 | Sncg |
| 34 | Fgf1 | 2.70E-108 | 6.9031844 | Sncg |
| 35 | Cadps2 | 6.12E-108 | 6.4848847 | Sncg |
| 36 | Pcdh20 | 1.57E-107 | 6.5777965 | Sncg |
| 37 | Megf10 | 3.60E-105 | 7.7789683 | Sncg |

| | | | | |
|---|---|---|---|---|
| 38 | Eps8 | 6.67E-105 | 6.710718 | Sncg |
| 39 | Rnd2 | 7.90E-105 | 5.1477375 | Sncg |
| 40 | Morf4l2 | 2.52E-104 | 1.1712004 | Sncg |
| 41 | Slc44a5 | 2.39E-101 | 5.6308904 | Sncg |
| 42 | 03-Mar | 4.82E-101 | 5.8849025 | Sncg |
| 43 | Col25a1 | 4.48E-100 | 5.812977 | Sncg |
| 44 | Cck | 8.90E-100 | 5.7616544 | Sncg |
| 45 | Dner | 1.72E-99 | 4.341646 | Sncg |
| 46 | Qdpr | 3.34E-99 | 1.9456762 | Sncg |
| 47 | Atp6v1g1 | 8.22E-99 | 1.1798211 | Sncg |
| 48 | Cntnap2 | 2.76E-98 | 3.413074 | Sncg |
| 49 | Nr2f2 | 7.85E-98 | 6.1970544 | Sncg |
| 50 | Pnoc | 2.66E-96 | 6.99543 | Sncg |
| 51 | Pcp4l1 | 8.06E-96 | 6.4054837 | Sncg |
| 52 | Arhgap24 | 8.19E-96 | 5.926583 | Sncg |
| 53 | Cygb | 1.57E-95 | 3.600312 | Sncg |
| 54 | Frem1 | 5.37E-95 | 7.6396637 | Sncg |
| 55 | Gm31271 | 8.02E-94 | 6.3558598 | Sncg |
| 56 | Tsn | 1.77E-93 | 1.3406947 | Sncg |
| 57 | Ciapin1 | 2.16E-92 | 2.630428 | Sncg |
| 58 | 1700019L22Rik | 7.78E-92 | 5.345288 | Sncg |
| 59 | Sh3bgrl | 8.61E-92 | 3.3340325 | Sncg |
| 60 | Cpne2 | 9.58E-91 | 6.3861017 | Sncg |
| 61 | Ache | 1.01E-90 | 3.5789974 | Sncg |
| 62 | Acot7 | 1.28E-90 | 1.3466457 | Sncg |
| 63 | Atp1b3 | 1.33E-90 | 1.8788494 | Sncg |
| 64 | Unc5b | 3.50E-90 | 5.975153 | Sncg |
| 65 | Igf1 | 7.56E-90 | 6.4394464 | Sncg |
| 66 | Abat | 1.69E-89 | 4.031489 | Sncg |
| 67 | P2ry1 | 2.01E-89 | 8.286612 | Sncg |
| 68 | Serpina12 | 2.62E-89 | 5.847917 | Sncg |
| 69 | G0s2 | 2.24E-88 | 4.8504033 | Sncg |
| 70 | A530058N18Rik | 2.40E-88 | 4.979109 | Sncg |
| 71 | Vstm2a | 1.47E-87 | 2.9083803 | Sncg |
| 72 | Nr3c2 | 3.70E-87 | 3.7434406 | Sncg |
| 73 | Gnao1 | 4.57E-87 | 1.3520536 | Sncg |
| 74 | Necab2 | 3.84E-86 | 5.5422053 | Sncg |
| 75 | Ndufa13 | 4.45E-86 | 1.6234015 | Sncg |
| 76 | Grip1 | 8.86E-85 | 3.6964731 | Sncg |
| 77 | Bend4 | 1.31E-84 | 3.7979805 | Sncg |
| 78 | Clic4 | 3.05E-84 | 5.6063385 | Sncg |
| 79 | Rgmb | 3.18E-84 | 3.0580847 | Sncg |
| 80 | Krt1 | 1.10E-83 | 4.2679377 | Sncg |
| 81 | Gm7338 | 3.41E-83 | 1.1728442 | Sncg |
| 82 | Atf4 | 7.13E-83 | 1.683476 | Sncg |
| 83 | Cd81 | 1.38E-82 | 1.0661433 | Sncg |
| 84 | Stk32c | 2.96E-82 | 3.633105 | Sncg |
| 85 | Slc25a1 | 5.10E-82 | 3.4943225 | Sncg |
| 86 | Cited2 | 8.09E-82 | 2.7336574 | Sncg |
| 87 | Elovl5 | 1.89E-81 | 4.464898 | Sncg |

| | | | | |
|---|---|---|---|---|
| 88 | Klhl13 | 7.78E-81 | 5.5522685 | Sncg |
| 89 | Galnt14 | 9.24E-81 | 4.5969954 | Sncg |
| 90 | Wnt5a | 9.43E-81 | 5.944559 | Sncg |
| 91 | E530001K10Rik | 1.52E-80 | 4.7342505 | Sncg |
| 92 | Serpina3n | 1.83E-80 | 5.3320394 | Sncg |
| 93 | Dlx2 | 8.73E-80 | 4.5782123 | Sncg |
| 94 | Vat1 | 2.57E-79 | 3.4997973 | Sncg |
| 95 | Cacng4 | 4.61E-79 | 4.4827137 | Sncg |
| 96 | App | 6.51E-79 | 1.5170028 | Sncg |
| 97 | Fstl5 | 3.19E-78 | 4.862361 | Sncg |
| 98 | Sqstm1 | 5.99E-78 | 1.0042226 | Sncg |
| 99 | Rragb | 2.23E-77 | 2.0658386 | Sncg |
| 0 | Sst | 0 | 14.250828 | Sst |
| 1 | Fhl1 | 0 | 3.2164388 | Sst |
| 2 | Kcnip1 | 0 | 5.7521744 | Sst |
| 3 | Ugdh | 0 | 3.315862 | Sst |
| 4 | Utrn | 0 | 3.567668 | Sst |
| 5 | Atp1b1 | 0 | 1.3080834 | Sst |
| 6 | Slc32a1 | 0 | 4.5184345 | Sst |
| 7 | Gm14164 | 0 | 2.5170038 | Sst |
| 8 | Dleu7 | 0 | 3.7910464 | Sst |
| 9 | Vstm2a | 0 | 2.6718974 | Sst |
| 10 | Neto2 | 0 | 3.7000148 | Sst |
| 11 | Chd3os | 0 | 1.4645066 | Sst |
| 12 | Dlx5 | 0 | 4.63754 | Sst |
| 13 | Cdh13 | 0 | 4.7231765 | Sst |
| 14 | Gm13889 | 0 | 3.7075474 | Sst |
| 15 | Cacna2d2 | 0 | 4.0376506 | Sst |
| 16 | Cgref1 | 0 | 3.6071985 | Sst |
| 17 | Dnm3 | 0 | 1.9026699 | Sst |
| 18 | Slc35g2 | 0 | 3.102764 | Sst |
| 19 | Npy | 0 | 7.029762 | Sst |
| 20 | Scoc | 0 | 1.2590607 | Sst |
| 21 | Rcan2 | 0 | 3.118028 | Sst |
| 22 | Cntnap3 | 0 | 5.0034432 | Sst |
| 23 | Ifit2 | 0 | 5.38508 | Sst |
| 24 | Inafm1 | 0 | 2.485458 | Sst |
| 25 | Reln | 0 | 5.6468716 | Sst |
| 26 | Samd5 | 0 | 3.4441109 | Sst |
| 27 | Unc13c | 0 | 6.290667 | Sst |
| 28 | Rnh1 | 0 | 2.5926402 | Sst |
| 29 | Npas1 | 0 | 5.486045 | Sst |
| 30 | Snx8 | 0 | 3.806254 | Sst |
| 31 | Gprasp2 | 0 | 2.1631145 | Sst |
| 32 | Gad2 | 0 | 6.54067 | Sst |
| 33 | Ptprm | 0 | 4.24593 | Sst |
| 34 | Psme1 | 0 | 1.87025 | Sst |
| 35 | Dlx6 | 0 | 3.3230157 | Sst |
| 36 | Vstm2b | 0 | 3.4162378 | Sst |
| 37 | Smarcd3 | 0 | 2.247684 | Sst |

| | | | | |
|---|---|---|---|---|
| 38 | Col19a1 | 0 | 4.206503 | Sst |
| 39 | Peg3 | 0 | 2.245882 | Sst |
| 40 | Frmd4b | 0 | 4.3620353 | Sst |
| 41 | Zfp385a | 0 | 2.7408721 | Sst |
| 42 | Ly6h | 0 | 1.4772491 | Sst |
| 43 | Nsg1 | 0 | 1.2631536 | Sst |
| 44 | Lgals1 | 0 | 6.0264635 | Sst |
| 45 | Tmeff2 | 0 | 2.7187304 | Sst |
| 46 | Ngfrap1 | 0 | 1.1113415 | Sst |
| 47 | Gng3 | 0 | 1.2468474 | Sst |
| 48 | Syngr3 | 0 | 1.6533616 | Sst |
| 49 | Crhbp | 0 | 7.403076 | Sst |
| 50 | Pfn2 | 0 | 1.0873228 | Sst |
| 51 | Lypd6 | 0 | 7.4605646 | Sst |
| 52 | Tmem130 | 0 | 2.6163964 | Sst |
| 53 | Arx | 0 | 5.5878816 | Sst |
| 54 | Rasgrp2 | 0 | 4.897266 | Sst |
| 55 | Ache | 0 | 3.831716 | Sst |
| 56 | Grm1 | 0 | 5.334351 | Sst |
| 57 | Nxph1 | 0 | 7.926072 | Sst |
| 58 | Satb1 | 0 | 4.44931 | Sst |
| 59 | Tmem91 | 0 | 4.7755103 | Sst |
| 60 | Elfn1 | 0 | 6.332425 | Sst |
| 61 | Mafb | 0 | 6.1151505 | Sst |
| 62 | Rpp25 | 0 | 5.5323358 | Sst |
| 63 | Klf5 | 0 | 4.967047 | Sst |
| 64 | Lypd6b | 0 | 6.9948754 | Sst |
| 65 | Sox6 | 0 | 6.9587116 | Sst |
| 66 | Nipsnap3b | 0 | 4.416046 | Sst |
| 67 | Nrsn2 | 0 | 4.6119637 | Sst |
| 68 | Rbp4 | 0 | 6.688812 | Sst |
| 69 | Grin3a | 0 | 7.5746274 | Sst |
| 70 | Lhx6 | 0 | 7.9957476 | Sst |
| 71 | Rab3b | 0 | 6.7796674 | Sst |
| 72 | Synpr | 0 | 7.6902676 | Sst |
| 73 | Masp1 | 0 | 4.875034 | Sst |
| 74 | Zcchc12 | 0 | 6.2291427 | Sst |
| 75 | Calb1 | 0 | 7.3517423 | Sst |
| 76 | Stxbp6 | 0 | 4.8040257 | Sst |
| 77 | Lurap1l | 0 | 2.982557 | Sst |
| 78 | Nrip2 | 0 | 4.357664 | Sst |
| 79 | Camk2n2 | 0 | 2.653392 | Sst |
| 80 | Trhde | 0 | 4.6745806 | Sst |
| 81 | Mgat4c | 0 | 4.760606 | Sst |
| 82 | Grik3 | 0 | 4.355991 | Sst |
| 83 | Eef1e1 | 0 | 1.917947 | Sst |
| 84 | Kcnc2 | 0 | 4.225035 | Sst |
| 85 | Nap1l5 | 0 | 1.9374659 | Sst |
| 86 | Glrx | 0 | 2.163194 | Sst |
| 87 | Rab27b | 0 | 3.2232907 | Sst |

| | | | | |
|---|---|---|---|---|
| 88 | Rgs17 | 0 | 2.3616288 | Sst |
| 89 | Laptm4b | 0 | 2.6622877 | Sst |
| 90 | Dgkg | 0 | 3.4008822 | Sst |
| 91 | Ifitm10 | 0 | 4.021205 | Sst |
| 92 | Pitpnc1 | 0 | 4.004429 | Sst |
| 93 | Lgmn | 0 | 1.810892 | Sst |
| 94 | Neto1 | 0 | 3.307765 | Sst |
| 95 | Elavl2 | 0 | 2.8257873 | Sst |
| 96 | Kctd8 | 0 | 6.052156 | Sst |
| 97 | Zwint | 0 | 1.3248196 | Sst |
| 98 | Slc5a5 | 0 | 4.958974 | Sst |
| 99 | Grik1 | 0 | 4.244884 | Sst |
| 0 | Apod | 1.90E-83 | 16.407696 | VLMC |
| 1 | Laptm4a | 1.90E-83 | 4.3699417 | VLMC |
| 2 | Ifitm2 | 1.45E-82 | 10.876615 | VLMC |
| 3 | Ptn | 3.53E-81 | 7.829719 | VLMC |
| 4 | Nupr1 | 3.68E-81 | 9.897647 | VLMC |
| 5 | Serping1 | 3.68E-81 | 14.384064 | VLMC |
| 6 | Apoe | 1.93E-80 | 12.912892 | VLMC |
| 7 | Col1a2 | 2.12E-80 | 14.673577 | VLMC |
| 8 | S100a11 | 2.13E-80 | 10.762166 | VLMC |
| 9 | Cd63 | 2.89E-80 | 8.59538 | VLMC |
| 10 | Serpinf1 | 3.80E-80 | 13.286699 | VLMC |
| 11 | Igfbp2 | 5.33E-80 | 13.068801 | VLMC |
| 12 | Cped1 | 6.19E-80 | 11.979099 | VLMC |
| 13 | Cxcl12 | 8.15E-80 | 10.21835 | VLMC |
| 14 | Dcn | 1.26E-79 | 16.568377 | VLMC |
| 15 | Ifitm3 | 1.47E-79 | 13.023707 | VLMC |
| 16 | Sparc | 3.96E-79 | 12.500664 | VLMC |
| 17 | Ptgds | 5.16E-79 | 14.949978 | VLMC |
| 18 | Ftl1 | 8.01E-78 | 2.7077372 | VLMC |
| 19 | Slc6a13 | 1.65E-77 | 16.91033 | VLMC |
| 20 | Igf2 | 1.65E-77 | 16.272503 | VLMC |
| 21 | Emp3 | 1.85E-77 | 11.097799 | VLMC |
| 22 | Pcolce | 2.23E-77 | 14.254008 | VLMC |
| 23 | Timp3 | 2.92E-77 | 10.127554 | VLMC |
| 24 | Cd63-ps | 4.40E-77 | 5.2180586 | VLMC |
| 25 | Zic1 | 8.38E-77 | 11.817486 | VLMC |
| 26 | Nbl1 | 1.47E-76 | 7.5900345 | VLMC |
| 27 | Zfp36l1 | 1.51E-76 | 9.716042 | VLMC |
| 28 | Vtn | 1.55E-76 | 15.428864 | VLMC |
| 29 | Atp1a2 | 5.06E-76 | 11.619926 | VLMC |
| 30 | Cfh | 1.08E-75 | 13.76197 | VLMC |
| 31 | Foxc1 | 2.15E-74 | 11.805659 | VLMC |
| 32 | Itm2b | 2.48E-74 | 1.5334007 | VLMC |
| 33 | Myl12a | 7.08E-74 | 6.4307013 | VLMC |
| 34 | Colec12 | 9.83E-74 | 12.979215 | VLMC |
| 35 | Slc6a20a | 5.53E-73 | 15.160074 | VLMC |
| 36 | Cp | 6.73E-73 | 10.920576 | VLMC |
| 37 | Arhgap29 | 6.90E-73 | 9.02366 | VLMC |

| | | | | |
|---|---|---|---|---|
| 38 | Itm2a | 1.86E-72 | 9.189937 | VLMC |
| 39 | Bgn | 1.93E-72 | 12.941518 | VLMC |
| 40 | Atp1b3 | 2.17E-72 | 3.394633 | VLMC |
| 41 | Cald1 | 3.08E-72 | 8.6596775 | VLMC |
| 42 | Gm7665 | 4.15E-72 | 7.2847147 | VLMC |
| 43 | Prelp | 5.65E-72 | 12.150881 | VLMC |
| 44 | Ftl2 | 6.31E-72 | 2.7665043 | VLMC |
| 45 | Pltp | 1.06E-71 | 11.245265 | VLMC |
| 46 | Slc1a3 | 1.45E-71 | 11.113808 | VLMC |
| 47 | Anxa2 | 4.15E-71 | 10.614903 | VLMC |
| 48 | Col1a1 | 8.49E-71 | 13.035321 | VLMC |
| 49 | Slc7a11 | 1.33E-70 | 11.297819 | VLMC |
| 50 | Fcgrt | 4.16E-70 | 10.942843 | VLMC |
| 51 | Id3 | 6.15E-70 | 10.112305 | VLMC |
| 52 | Aldh1a2 | 6.83E-70 | 16.419067 | VLMC |
| 53 | Cst3 | 2.10E-68 | 2.404504 | VLMC |
| 54 | Serpinh1 | 2.45E-68 | 10.403067 | VLMC |
| 55 | Gja1 | 3.23E-68 | 10.049342 | VLMC |
| 56 | Gng5 | 4.66E-68 | 6.2380877 | VLMC |
| 57 | Vamp5 | 1.05E-67 | 11.171202 | VLMC |
| 58 | Glul | 2.72E-66 | 4.684374 | VLMC |
| 59 | Ctsl | 9.94E-66 | 2.610737 | VLMC |
| 60 | Slc22a6 | 1.79E-65 | 16.85827 | VLMC |
| 61 | Ctsh | 3.04E-65 | 11.774093 | VLMC |
| 62 | B2m | 3.35E-65 | 3.9338865 | VLMC |
| 63 | Gpx8 | 9.42E-65 | 9.548645 | VLMC |
| 64 | LOC102637129 | 3.17E-64 | 5.3717117 | VLMC |
| 65 | Olfml3 | 3.78E-64 | 12.203359 | VLMC |
| 66 | Isyna1 | 4.15E-64 | 6.309657 | VLMC |
| 67 | Pdgfra | 1.41E-63 | 10.806719 | VLMC |
| 68 | Vcam1 | 1.67E-63 | 10.49941 | VLMC |
| 69 | Cgnl1 | 2.24E-63 | 8.218824 | VLMC |
| 70 | Pcca | 2.35E-63 | 5.9205546 | VLMC |
| 71 | Lum | 2.76E-63 | 16.332699 | VLMC |
| 72 | Efemp1 | 3.02E-63 | 10.652866 | VLMC |
| 73 | Gjb2 | 5.13E-63 | 13.711938 | VLMC |
| 74 | LOC100861618 | 8.06E-63 | 10.605427 | VLMC |
| 75 | Gstm1 | 1.75E-62 | 7.919214 | VLMC |
| 76 | Anxa5 | 4.77E-62 | 5.462823 | VLMC |
| 77 | Ogn | 7.01E-62 | 13.773615 | VLMC |
| 78 | Lpar1 | 1.83E-61 | 8.204672 | VLMC |
| 79 | Gm10116 | 2.66E-61 | 2.6894069 | VLMC |
| 80 | Slc22a8 | 6.02E-61 | 12.372645 | VLMC |
| 81 | Lgals1 | 8.06E-61 | 8.105876 | VLMC |
| 82 | Itih5 | 8.81E-61 | 10.6276655 | VLMC |
| 83 | Cd81 | 8.99E-61 | 1.7900066 | VLMC |
| 84 | Cnn2 | 4.18E-60 | 10.035216 | VLMC |
| 85 | Slc13a4 | 6.61E-60 | 13.885454 | VLMC |
| 86 | Islr | 2.16E-59 | 9.58293 | VLMC |
| 87 | Sepp1 | 2.57E-59 | 9.574301 | VLMC |

| | | | | |
|---|---|---|---|---|
| 88 | Slc3a2 | 3.00E-59 | 2.9684784 | VLMC |
| 89 | Phlda1 | 4.28E-59 | 4.7625527 | VLMC |
| 90 | Trf | 6.67E-59 | 10.735312 | VLMC |
| 91 | Cpq | 4.46E-58 | 9.959476 | VLMC |
| 92 | Cmbl | 4.68E-58 | 8.566605 | VLMC |
| 93 | Rcn3 | 8.34E-58 | 10.951226 | VLMC |
| 94 | Tbx18 | 2.17E-57 | 11.602339 | VLMC |
| 95 | Tnfsf12 | 3.31E-57 | 6.8148956 | VLMC |
| 96 | Fxyd1 | 3.32E-57 | 5.860918 | VLMC |
| 97 | Mdk | 3.71E-57 | 8.683446 | VLMC |
| 98 | Rbp1 | 4.03E-57 | 10.334151 | VLMC |
| 99 | Gjb6 | 1.15E-56 | 11.636535 | VLMC |
| 0 | Ap1s2 | 0 | 5.9219623 | Vip |
| 1 | Hnrnpa2b1 | 0 | 0.64935833 | Vip |
| 2 | Scg2 | 0 | 2.7722223 | Vip |
| 3 | Resp18 | 0 | 2.5767412 | Vip |
| 4 | Sez6 | 0 | 3.2442732 | Vip |
| 5 | Tmem130 | 0 | 2.2071984 | Vip |
| 6 | Cxcl14 | 0 | 7.222315 | Vip |
| 7 | A530058N18Rik | 0 | 4.4219065 | Vip |
| 8 | Enho | 0 | 2.4005828 | Vip |
| 9 | Rgs16 | 0 | 5.0632353 | Vip |
| 10 | Kcnip1 | 0 | 5.3643055 | Vip |
| 11 | Dennd2a | 0 | 3.744151 | Vip |
| 12 | Slc2a13 | 0 | 2.7696955 | Vip |
| 13 | Magi3 | 0 | 3.3126683 | Vip |
| 14 | Srrm4 | 0 | 2.6422944 | Vip |
| 15 | Dlx2 | 0 | 4.4678464 | Vip |
| 16 | Npas3 | 0 | 4.624935 | Vip |
| 17 | Rpp25 | 0 | 4.762993 | Vip |
| 18 | Rgs8 | 0 | 4.446156 | Vip |
| 19 | Bex4 | 0 | 1.9743524 | Vip |
| 20 | Dlx1as | 0 | 4.660394 | Vip |
| 21 | Gad1 | 0 | 7.676919 | Vip |
| 22 | Sema3c | 0 | 5.232242 | Vip |
| 23 | Slc24a3 | 0 | 3.6860914 | Vip |
| 24 | Lurap1l | 0 | 2.6510158 | Vip |
| 25 | Sox2ot | 0 | 4.595591 | Vip |
| 26 | Wsb1 | 0 | 1.577946 | Vip |
| 27 | Dner | 0 | 3.4869342 | Vip |
| 28 | Pcbd1 | 0 | 4.0197387 | Vip |
| 29 | Nrip3 | 0 | 3.47047 | Vip |
| 30 | Galnt14 | 0 | 3.8441133 | Vip |
| 31 | Elmod1 | 0 | 1.6483037 | Vip |
| 32 | Zfp804a | 0 | 3.7894642 | Vip |
| 33 | Gad2 | 0 | 6.3129516 | Vip |
| 34 | Arl4c | 0 | 3.7088943 | Vip |
| 35 | Slc32a1 | 0 | 4.1862206 | Vip |
| 36 | Npas1 | 0 | 5.525597 | Vip |
| 37 | Tac2 | 0 | 8.194211 | Vip |

| | | | | |
|---|---|---|---|---|
| 38 | Nr2e1 | 0 | 5.8505225 | Vip |
| 39 | Gm5087 | 0 | 4.9743114 | Vip |
| 40 | E530001K10Rik | 0 | 4.382616 | Vip |
| 41 | Wbp5 | 0 | 1.1773887 | Vip |
| 42 | Elfn1 | 0 | 4.552415 | Vip |
| 43 | Glra2 | 0 | 4.6666718 | Vip |
| 44 | Ddah1 | 0 | 3.6782677 | Vip |
| 45 | Gm3134 | 0 | 5.742471 | Vip |
| 46 | Cbln2 | 0 | 4.6092877 | Vip |
| 47 | Kit | 0 | 5.605914 | Vip |
| 48 | Htr3a | 0 | 7.903124 | Vip |
| 49 | Cpne2 | 0 | 5.817361 | Vip |
| 50 | Asic1 | 0 | 3.6009681 | Vip |
| 51 | Bex1 | 0 | 2.0896068 | Vip |
| 52 | LOC102632463 | 0 | 7.82361 | Vip |
| 53 | Nap1l5 | 0 | 2.0596142 | Vip |
| 54 | Gpd1 | 0 | 5.14123 | Vip |
| 55 | Vstm2l | 0 | 3.3770244 | Vip |
| 56 | Igsf3 | 0 | 4.6169853 | Vip |
| 57 | Limch1 | 0 | 3.9729204 | Vip |
| 58 | Cnr1 | 0 | 4.664827 | Vip |
| 59 | Crh | 0 | 10.527908 | Vip |
| 60 | Sh3bgrl | 0 | 3.712147 | Vip |
| 61 | Fxyd6 | 0 | 4.782559 | Vip |
| 62 | Dlx6os1 | 0 | 7.3176737 | Vip |
| 63 | Asic4 | 0 | 7.57405 | Vip |
| 64 | Prox1 | 0 | 8.037738 | Vip |
| 65 | Rgs10 | 0 | 6.420411 | Vip |
| 66 | Tcf4 | 0 | 2.03345 | Vip |
| 67 | Dlx1 | 0 | 6.348443 | Vip |
| 68 | Synpr | 0 | 8.526439 | Vip |
| 69 | Adarb2 | 0 | 9.018723 | Vip |
| 70 | Igf1 | 0 | 8.344139 | Vip |
| 71 | Vip | 0 | 14.640301 | Vip |
| 72 | Pthlh | 0 | 8.211665 | Vip |
| 73 | Sall1 | 0 | 7.2498093 | Vip |
| 74 | Cd63 | 0 | 6.065173 | Vip |
| 75 | Sorcs3 | 0 | 4.796174 | Vip |
| 76 | Gm17750 | 0 | 5.5505342 | Vip |
| 77 | Htra1 | 0 | 5.023504 | Vip |
| 78 | Uchl3 | 0 | 1.7209444 | Vip |
| 79 | Fstl5 | 0 | 4.808689 | Vip |
| 80 | Ptprz1 | 0 | 5.6810946 | Vip |
| 81 | Nr2f2 | 0 | 5.3361435 | Vip |
| 82 | Qpct | 0 | 4.679887 | Vip |
| 83 | Adra1b | 0 | 4.4790144 | Vip |
| 84 | Tiam1 | 0 | 3.140927 | Vip |
| 85 | Calb2 | 0 | 9.074859 | Vip |
| 86 | H3f3b | 0 | 0.7900018 | Vip |
| 87 | Erbb4 | 0 | 6.201828 | Vip |

| | | | | |
|---|---|---|---|---|
| 88 | Inpp5f | 0 | 2.5255957 | Vip |
| 89 | Nrxn3 | 0 | 2.197458 | Vip |
| 90 | Zcchc12 | 0 | 5.8905764 | Vip |
| 91 | Grpr | 0 | 9.114876 | Vip |
| 92 | Vstm2a | 0 | 2.9590929 | Vip |
| 93 | Cpne7 | 0 | 5.517805 | Vip |
| 94 | Csmd3 | 0 | 3.6827667 | Vip |
| 95 | Penk | 0 | 8.26366 | Vip |
| 96 | Zfp536 | 0 | 5.3707767 | Vip |
| 97 | Cit | 0 | 2.3687732 | Vip |
| 98 | Npy1r | 0 | 4.688037 | Vip |
| 99 | Rit2 | 0 | 1.5518634 | Vip |
| 0 | Cx3cl1 | 9.98E-70 | 1.7244904 | nan |
| 1 | Mical2 | 5.36E-67 | 2.3479955 | nan |
| 2 | Fscn1 | 4.02E-63 | 1.8516325 | nan |
| 3 | Dmtn | 1.64E-60 | 1.4975702 | nan |
| 4 | Fam131a | 7.36E-60 | 2.5003111 | nan |
| 5 | Eno2 | 2.17E-59 | 1.4364173 | nan |
| 6 | Nefl | 9.91E-58 | 3.5445259 | nan |
| 7 | Pak1 | 1.60E-55 | 1.8749458 | nan |
| 8 | R3hdm4 | 2.09E-55 | 1.7369885 | nan |
| 9 | Brinp1 | 6.23E-54 | 2.262036 | nan |
| 10 | Dnajb5 | 1.34E-53 | 2.885125 | nan |
| 11 | Tmem151a | 1.34E-53 | 1.9173046 | nan |
| 12 | Syp | 1.56E-53 | 1.2629701 | nan |
| 13 | Psap | 4.90E-53 | 0.9681648 | nan |
| 14 | Galnt9 | 1.41E-52 | 3.2257512 | nan |
| 15 | Ncdn | 2.03E-52 | 1.5760326 | nan |
| 16 | Ndrg4 | 6.27E-52 | 1.2486968 | nan |
| 17 | Rtn3 | 1.19E-50 | 0.7963266 | nan |
| 18 | Arf3 | 5.13E-50 | 1.4983784 | nan |
| 19 | Arc | 1.82E-49 | 4.281644 | nan |
| 20 | Nefm | 3.12E-49 | 3.7433958 | nan |
| 21 | Cyfip2 | 4.96E-49 | 1.8873422 | nan |
| 22 | App | 5.99E-49 | 1.0243944 | nan |
| 23 | Extl1 | 8.07E-49 | 2.8823135 | nan |
| 24 | Atp1a3 | 8.63E-49 | 1.6980028 | nan |
| 25 | Gm5559 | 9.83E-49 | 1.0298798 | nan |
| 26 | Rangap1 | 1.29E-48 | 1.7757196 | nan |
| 27 | Stxbp1 | 1.31E-47 | 1.1792629 | nan |
| 28 | Clstn3 | 2.20E-47 | 1.8008147 | nan |
| 29 | Slc4a10 | 3.13E-47 | 1.5662639 | nan |
| 30 | Kifc2 | 4.18E-47 | 1.8205758 | nan |
| 31 | Gpr162 | 6.80E-47 | 1.4340239 | nan |
| 32 | Trim9 | 2.31E-46 | 1.6669666 | nan |
| 33 | Dbn1 | 2.87E-46 | 1.7849208 | nan |
| 34 | Tubb4a | 2.21E-45 | 1.1576864 | nan |
| 35 | Chst8 | 2.95E-45 | 4.3862286 | nan |
| 36 | Ap2a1 | 2.62E-44 | 1.6660663 | nan |
| 37 | Lasp1 | 2.62E-44 | 2.3234148 | nan |

| | | | | |
|---|---|---|---|---|
| 38 | Sult4a1 | 3.21E-44 | 1.2982637 | nan |
| 39 | Apbb1 | 4.82E-44 | 1.3375719 | nan |
| 40 | Cyb561 | 6.51E-44 | 2.1192603 | nan |
| 41 | Chst1 | 2.50E-43 | 2.2745287 | nan |
| 42 | Adora1 | 3.95E-43 | 3.2209737 | nan |
| 43 | Stx1b | 4.13E-43 | 1.5007472 | nan |
| 44 | Chrm1 | 5.02E-43 | 2.4131906 | nan |
| 45 | Hk1 | 5.11E-43 | 1.6076959 | nan |
| 46 | Ptk2b | 6.86E-43 | 3.5696664 | nan |
| 47 | Scamp5 | 9.19E-43 | 1.3632394 | nan |
| 48 | Tmem198 | 1.36E-42 | 2.3641 | nan |
| 49 | Icam5 | 2.87E-42 | 2.7088306 | nan |
| 50 | Tenm2 | 5.49E-42 | 2.198339 | nan |
| 51 | D030054H15Rik | 5.90E-42 | 2.22034 | nan |
| 52 | Synpo | 8.08E-42 | 2.4947007 | nan |
| 53 | Nrcam | 8.56E-42 | 1.5632336 | nan |
| 54 | Dctn1 | 9.52E-42 | 1.4943483 | nan |
| 55 | Etv5 | 2.96E-41 | 2.2939599 | nan |
| 56 | Sirpa | 6.71E-41 | 1.7581016 | nan |
| 57 | Camk2b | 7.15E-41 | 1.2396796 | nan |
| 58 | Tox | 9.16E-41 | 3.145685 | nan |
| 59 | Bhlhe40 | 1.30E-40 | 3.5403628 | nan |
| 60 | Slc17a7 | 2.31E-40 | 4.2087955 | nan |
| 61 | Kcnip3 | 2.38E-40 | 3.2154796 | nan |
| 62 | Cap2 | 4.37E-40 | 2.0880663 | nan |
| 63 | Ptpn5 | 4.72E-40 | 1.8039178 | nan |
| 64 | Dgat2 | 4.76E-40 | 2.2928777 | nan |
| 65 | Zfp365 | 7.37E-40 | 1.9078658 | nan |
| 66 | Prkcg | 1.07E-39 | 2.889859 | nan |
| 67 | Pik3r2 | 1.15E-39 | 2.2158267 | nan |
| 68 | Lppr2 | 1.32E-39 | 1.5844127 | nan |
| 69 | Vamp2 | 2.28E-39 | 0.76175857 | nan |
| 70 | Camk2a | 2.48E-39 | 1.9198983 | nan |
| 71 | 8430419L09Rik | 3.10E-39 | 1.3994089 | nan |
| 72 | Got2 | 4.65E-39 | 1.2262803 | nan |
| 73 | Dlg1 | 4.75E-39 | 2.1093593 | nan |
| 74 | Ndrg3 | 5.00E-39 | 1.0566027 | nan |
| 75 | Magee1 | 8.12E-39 | 1.4181045 | nan |
| 76 | Tiam2 | 1.03E-38 | 3.0189593 | nan |
| 77 | Stx6 | 2.03E-38 | 1.7091976 | nan |
| 78 | Slc25a22 | 6.46E-38 | 1.5094992 | nan |
| 79 | Rnf112 | 7.14E-38 | 2.1183417 | nan |
| 80 | Apba2 | 1.54E-37 | 1.6717159 | nan |
| 81 | Rasgrf1 | 1.66E-37 | 1.8775653 | nan |
| 82 | Nr4a1 | 2.00E-37 | 3.099112 | nan |
| 83 | 05-Sep | 2.21E-37 | 1.2895435 | nan |
| 84 | Slc20a1 | 2.35E-37 | 2.1209822 | nan |
| 85 | Mapk8ip2 | 2.46E-37 | 1.7014598 | nan |
| 86 | Rap1gap | 2.46E-37 | 1.6070963 | nan |
| 87 | Ldb2 | 3.05E-37 | 3.119417 | nan |

| | | | | |
|---|---|---|---|---|
| 88 | 9430020K01Rik | 3.77E-37 | 2.9861085 | nan |
| 89 | Tenm3 | 4.12E-37 | 2.7830837 | nan |
| 90 | Dusp6 | 4.14E-37 | 2.659461 | nan |
| 91 | Tspyl1 | 5.35E-37 | 1.1942362 | nan |
| 92 | Dclk1 | 6.76E-37 | 1.1665971 | nan |
| 93 | Sorl1 | 9.90E-37 | 2.1925685 | nan |
| 94 | Slc6a7 | 2.28E-36 | 2.6481004 | nan |
| 95 | Tmem132a | 3.42E-36 | 2.09609 | nan |
| 96 | Sprn | 3.63E-36 | 2.0678031 | nan |
| 97 | Elmo2 | 4.02E-36 | 2.0613146 | nan |
| 98 | Camkv | 5.05E-36 | 1.8131216 | nan |
| 99 | Tcf25 | 5.07E-36 | 0.8071935 | nan |
