## Supplementary Table 3 for "CellDART: Cell type inference by domain adaptation of single-cell and spatial transcriptomic data"

| **Table 3** | Baseline and clinical characteristic of the patient whom two lung tissues were obtained. *Clinical and pathologic stage were determined by the 8th edition of American Joint Committee on Cancer |
|---|---|
| **Age** | 76 |
| **Sex** | Female |
| **Smoking history** | Never smoker |
| **Cell type** | adenocarcinoma |
| **Tumor location** | Left lower lobe |
| **Surgery** | Lobectomy |
| **Clinical stage** | cT2aN0M0 |
| **Pathologic stage** | pT1cN0M0 |
