## Supplementary Table 4 for "CellDART: Cell type inference by domain adaptation of single-cell and spatial transcriptomic data"

| **Table 4** | The top 100 marker genes for each cell clusters in human normal lung single-cell dataset. The genes were ranked based on the Benjamini-Hochberg adjusted p-values. The name of the cells are based on the metadata provided by the paper (Travaglini, K.J. et al. A molecular cell atlas of the human lung from single-cell RNA sequencing. *Nature* **587**, 619-625 (2020)). | | | |
|---|---|---|---|---|
| Number | Genes | Adjusted p-value | Log fold change | Cluster |
| 0 | DCN | 0 | 169.00246 | Adventitial Fibroblast |
| 1 | FBLN1 | 0 | 68.617966 | Adventitial Fibroblast |
| 2 | LUM | 0 | 43.32164 | Adventitial Fibroblast |
| 3 | C1S | 0 | 17.63311 | Adventitial Fibroblast |
| 4 | COL6A2 | 0 | 15.723071 | Adventitial Fibroblast |
| 5 | C1R | 0 | 20.35082 | Adventitial Fibroblast |
| 6 | COL1A2 | 3.15E-304 | 18.76894 | Adventitial Fibroblast |
| 7 | MGP | 1.22E-297 | 53.51414 | Adventitial Fibroblast |
| 8 | MFAP4 | 1.76E-292 | 18.127424 | Adventitial Fibroblast |
| 9 | RARRES2 | 4.24E-287 | 11.159872 | Adventitial Fibroblast |
| 10 | ADH1B | 1.76E-277 | 14.682549 | Adventitial Fibroblast |
| 11 | LTBP4 | 5.40E-261 | 8.056789 | Adventitial Fibroblast |
| 12 | SERPINF1 | 1.08E-259 | 17.358837 | Adventitial Fibroblast |
| 13 | MMP2 | 6.51E-256 | 9.219447 | Adventitial Fibroblast |
| 14 | PLAC9 | 8.20E-253 | 11.1654215 | Adventitial Fibroblast |
| 15 | SEPP1 | 2.69E-251 | 13.445597 | Adventitial Fibroblast |
| 16 | COL6A1 | 1.15E-249 | 7.5892277 | Adventitial Fibroblast |
| 17 | PCOLCE | 9.31E-247 | 9.87148 | Adventitial Fibroblast |
| 18 | CYR61 | 3.09E-237 | 35.419773 | Adventitial Fibroblast |
| 19 | SOD3 | 9.37E-232 | 7.8457317 | Adventitial Fibroblast |
| 20 | SPARCL1 | 1.96E-231 | 13.201212 | Adventitial Fibroblast |
| 21 | CCDC80 | 1.35E-226 | 22.808495 | Adventitial Fibroblast |
| 22 | CFD | 3.49E-219 | 105.99709 | Adventitial Fibroblast |
| 23 | IGFBP6 | 1.58E-211 | 43.709667 | Adventitial Fibroblast |
| 24 | NNMT | 7.87E-211 | 13.648226 | Adventitial Fibroblast |
| 25 | PMP22 | 1.59E-204 | 7.1837497 | Adventitial Fibroblast |
| 26 | PLTP | 5.72E-198 | 6.907031 | Adventitial Fibroblast |
| 27 | MYC | 1.02E-196 | 9.191546 | Adventitial Fibroblast |
| 28 | AEBP1 | 3.01E-196 | 6.7943025 | Adventitial Fibroblast |
| 29 | C3 | 4.66E-192 | 10.441346 | Adventitial Fibroblast |
| 30 | COL1A1 | 2.05E-188 | 11.002228 | Adventitial Fibroblast |
| 31 | PRELP | 4.37E-180 | 6.7185097 | Adventitial Fibroblast |
| 32 | C7 | 1.05E-179 | 11.877691 | Adventitial Fibroblast |
| 33 | JUNB | 5.04E-179 | 37.590252 | Adventitial Fibroblast |
| 34 | EGR1 | 1.93E-177 | 7.8634505 | Adventitial Fibroblast |
| 35 | CFH | 3.81E-177 | 6.185496 | Adventitial Fibroblast |
| 36 | FSTL1 | 5.17E-176 | 7.8820252 | Adventitial Fibroblast |
| 37 | FXYD1 | 1.15E-175 | 5.5792575 | Adventitial Fibroblast |
| 38 | GSN | 3.01E-173 | 43.786953 | Adventitial Fibroblast |
| 39 | EFEMP1 | 5.41E-171 | 10.676024 | Adventitial Fibroblast |
| 40 | OLFML3 | 3.02E-167 | 6.7719955 | Adventitial Fibroblast |
| 41 | FBLN5 | 5.65E-160 | 5.483748 | Adventitial Fibroblast |
| 42 | ZFP36 | 1.53E-159 | 35.27056 | Adventitial Fibroblast |
| 43 | CLU | 1.35E-158 | 15.347804 | Adventitial Fibroblast |
| 44 | BTG2 | 1.54E-157 | 9.790561 | Adventitial Fibroblast |
| 45 | CALD1 | 4.92E-157 | 3.7203622 | Adventitial Fibroblast |
| 46 | NR4A1 | 3.38E-156 | 8.896727 | Adventitial Fibroblast |
| 47 | SERPING1 | 5.09E-156 | 7.739921 | Adventitial Fibroblast |
| 48 | PPAP2B | 3.67E-155 | 5.326537 | Adventitial Fibroblast |
| 49 | COL6A3 | 4.99E-154 | 6.2845984 | Adventitial Fibroblast |
| 50 | FBLN2 | 2.33E-151 | 8.587626 | Adventitial Fibroblast |
| 51 | FHL1 | 6.24E-148 | 5.826919 | Adventitial Fibroblast |
| 52 | IGFBP7 | 4.80E-145 | 18.014246 | Adventitial Fibroblast |
| 53 | TIMP1 | 3.00E-144 | 22.55048 | Adventitial Fibroblast |
| 54 | TIMP3 | 1.26E-140 | 7.1778846 | Adventitial Fibroblast |

| | | | | |
|---|---|---|---|---|
| 55 | COL3A1 | 2.12E-139 | 9.760446 | Adventitial Fibroblast |
| 56 | ARID5B | 1.67E-137 | 5.2078815 | Adventitial Fibroblast |
| 57 | SERPINA3 | 9.13E-134 | 6.604821 | Adventitial Fibroblast |
| 58 | CPE | 4.03E-131 | 8.127172 | Adventitial Fibroblast |
| 59 | EMP1 | 1.24E-130 | 6.2616267 | Adventitial Fibroblast |
| 60 | CEBPD | 5.04E-128 | 13.223508 | Adventitial Fibroblast |
| 61 | FGF7 | 5.03E-127 | 7.1889157 | Adventitial Fibroblast |
| 62 | ELN | 8.11E-126 | 4.9416666 | Adventitial Fibroblast |
| 63 | PTGDS | 2.18E-125 | 14.372524 | Adventitial Fibroblast |
| 64 | VASN | 3.70E-125 | 5.916349 | Adventitial Fibroblast |
| 65 | FIBIN | 1.10E-124 | 6.503283 | Adventitial Fibroblast |
| 66 | MXRA8 | 2.45E-124 | 5.5992017 | Adventitial Fibroblast |
| 67 | DUSP1 | 4.59E-123 | 22.197784 | Adventitial Fibroblast |
| 68 | PODN | 4.96E-122 | 7.645545 | Adventitial Fibroblast |
| 69 | SOCS3 | 5.72E-122 | 11.225279 | Adventitial Fibroblast |
| 70 | CRISPLD2 | 1.10E-121 | 6.6930084 | Adventitial Fibroblast |
| 71 | CTGF | 1.33E-121 | 18.110153 | Adventitial Fibroblast |
| 72 | SFRP2 | 1.83E-118 | 39.337616 | Adventitial Fibroblast |
| 73 | MEG3 | 1.92E-118 | 8.262818 | Adventitial Fibroblast |
| 74 | CHRDL1 | 1.50E-116 | 8.086965 | Adventitial Fibroblast |
| 75 | A2M | 1.59E-114 | 13.169773 | Adventitial Fibroblast |
| 76 | SPARC | 1.20E-113 | 5.1715918 | Adventitial Fibroblast |
| 77 | RNASE4 | 1.45E-111 | 3.7165766 | Adventitial Fibroblast |
| 78 | ZFP36L1 | 4.43E-110 | 5.500745 | Adventitial Fibroblast |
| 79 | CCL2 | 3.86E-109 | 53.83411 | Adventitial Fibroblast |
| 80 | DPT | 7.89E-108 | 7.08795 | Adventitial Fibroblast |
| 81 | ABI3BP | 5.47E-107 | 4.7392607 | Adventitial Fibroblast |
| 82 | FGFR1 | 4.28E-106 | 4.3777747 | Adventitial Fibroblast |
| 83 | ADAMTS1 | 1.59E-105 | 5.2922125 | Adventitial Fibroblast |
| 84 | CYBRD1 | 1.61E-105 | 4.3331923 | Adventitial Fibroblast |
| 85 | AKAP12 | 1.09E-104 | 4.619515 | Adventitial Fibroblast |
| 86 | RARRES1 | 4.47E-104 | 7.8359103 | Adventitial Fibroblast |
| 87 | WISP2 | 2.15E-103 | 9.562735 | Adventitial Fibroblast |
| 88 | EFEMP2 | 3.15E-100 | 4.125719 | Adventitial Fibroblast |
| 89 | BGN | 4.03E-100 | 3.5227985 | Adventitial Fibroblast |
| 90 | LAPTM4A | 5.52E-100 | 6.202166 | Adventitial Fibroblast |
| 91 | SFRP4 | 9.55E-100 | 35.10838 | Adventitial Fibroblast |
| 92 | SELM | 3.33E-99 | 3.1077304 | Adventitial Fibroblast |
| 93 | HTRA3 | 2.31E-98 | 8.774634 | Adventitial Fibroblast |
| 94 | LMNA | 2.85E-97 | 10.939475 | Adventitial Fibroblast |
| 95 | TCF21 | 3.74E-97 | 5.198577 | Adventitial Fibroblast |
| 96 | TMEM176B | 4.53E-96 | 3.6075652 | Adventitial Fibroblast |
| 97 | FOS | 1.58E-94 | 20.089106 | Adventitial Fibroblast |
| 98 | CTSF | 4.99E-94 | 4.1487994 | Adventitial Fibroblast |
| 99 | CTSK | 1.67E-93 | 6.537733 | Adventitial Fibroblast |
| 0 | TAGLN | 0 | 110.27098 | Airway Smooth Muscle |
| 17 | SPARCL1 | 0 | 18.219484 | Airway Smooth Muscle |
| 16 | BGN | 0 | 9.134907 | Airway Smooth Muscle |
| 15 | MGP | 0 | 16.586779 | Airway Smooth Muscle |
| 14 | PLAC9 | 0 | 8.708839 | Airway Smooth Muscle |
| 13 | SELM | 0 | 11.576464 | Airway Smooth Muscle |
| 12 | ACTG2 | 0 | 12.531518 | Airway Smooth Muscle |
| 10 | MFAP4 | 0 | 13.355396 | Airway Smooth Muscle |
| 9 | MYH11 | 0 | 14.298017 | Airway Smooth Muscle |
| 11 | DSTN | 0 | 25.141073 | Airway Smooth Muscle |
| 7 | CALD1 | 0 | 15.906803 | Airway Smooth Muscle |
| 6 | IGFBP7 | 0 | 53.844624 | Airway Smooth Muscle |
| 5 | SOD3 | 0 | 18.00666 | Airway Smooth Muscle |
| 4 | PPP1R14A | 0 | 14.5787525 | Airway Smooth Muscle |
| 3 | MYL9 | 0 | 54.571735 | Airway Smooth Muscle |
| 2 | TPM2 | 0 | 44.20291 | Airway Smooth Muscle |
| 1 | ACTA2 | 0 | 80.60473 | Airway Smooth Muscle |
| 8 | TPM1 | 0 | 13.501352 | Airway Smooth Muscle |
| 18 | ADIRF | 8.75E-305 | 33.342247 | Airway Smooth Muscle |

| | | | | |
|---|---|---|---|---|
| 19 | CRYAB | 4.31E-303 | 9.1758 | Airway Smooth Muscle |
| 20 | PRKCDBP | 1.16E-302 | 7.6623983 | Airway Smooth Muscle |
| 21 | MFGE8 | 1.52E-292 | 7.926437 | Airway Smooth Muscle |
| 22 | CSRP1 | 7.87E-291 | 9.071665 | Airway Smooth Muscle |
| 23 | CNN1 | 9.68E-280 | 10.359457 | Airway Smooth Muscle |
| 24 | PDLIM3 | 5.49E-267 | 7.1842604 | Airway Smooth Muscle |
| 25 | DES | 4.64E-257 | 13.605358 | Airway Smooth Muscle |
| 26 | PLN | 1.32E-253 | 12.170655 | Airway Smooth Muscle |
| 27 | IGFBP5 | 3.41E-250 | 8.987259 | Airway Smooth Muscle |
| 28 | LMOD1 | 9.16E-249 | 7.7848916 | Airway Smooth Muscle |
| 29 | COL6A2 | 3.25E-247 | 5.239654 | Airway Smooth Muscle |
| 30 | TGFB1I1 | 1.44E-245 | 5.5952554 | Airway Smooth Muscle |
| 31 | FXYD1 | 6.29E-240 | 6.371144 | Airway Smooth Muscle |
| 32 | TINAGL1 | 1.73E-239 | 5.4822364 | Airway Smooth Muscle |
| 33 | CSRP2 | 9.21E-239 | 7.1035924 | Airway Smooth Muscle |
| 34 | AEBP1 | 2.47E-237 | 5.909943 | Airway Smooth Muscle |
| 35 | WFDC1 | 1.15E-232 | 5.96292 | Airway Smooth Muscle |
| 36 | MYLK | 1.77E-232 | 6.5865073 | Airway Smooth Muscle |
| 37 | SDC2 | 9.07E-227 | 5.061749 | Airway Smooth Muscle |
| 38 | ID4 | 6.26E-225 | 6.806478 | Airway Smooth Muscle |
| 39 | C2orf40 | 1.38E-224 | 5.835368 | Airway Smooth Muscle |
| 40 | MCAM | 1.34E-218 | 5.855905 | Airway Smooth Muscle |
| 41 | AOC3 | 1.98E-216 | 5.094225 | Airway Smooth Muscle |
| 42 | MAP1B | 2.56E-214 | 6.537548 | Airway Smooth Muscle |
| 43 | EGFL6 | 1.64E-209 | 6.4025154 | Airway Smooth Muscle |
| 44 | RARRES2 | 4.48E-204 | 4.1223755 | Airway Smooth Muscle |
| 45 | PDLIM7 | 7.10E-204 | 4.7381425 | Airway Smooth Muscle |
| 46 | ANXA6 | 1.84E-202 | 4.2915406 | Airway Smooth Muscle |
| 47 | NEXN | 8.28E-200 | 5.801833 | Airway Smooth Muscle |
| 48 | CRIP2 | 6.71E-198 | 5.842468 | Airway Smooth Muscle |
| 49 | CAV1 | 2.36E-190 | 7.5290203 | Airway Smooth Muscle |
| 50 | PTRF | 3.95E-189 | 4.6984496 | Airway Smooth Muscle |
| 51 | HES4 | 1.11E-183 | 4.330022 | Airway Smooth Muscle |
| 52 | FRZB | 2.57E-183 | 7.769269 | Airway Smooth Muscle |
| 53 | COX7A1 | 1.28E-182 | 4.549683 | Airway Smooth Muscle |
| 54 | FLNA | 6.69E-174 | 5.796167 | Airway Smooth Muscle |
| 55 | C11orf96 | 4.15E-172 | 7.6330943 | Airway Smooth Muscle |
| 56 | 07-Sep | 5.71E-167 | 4.742722 | Airway Smooth Muscle |
| 57 | PDGFRB | 3.38E-166 | 4.5384264 | Airway Smooth Muscle |
| 58 | HSPB1 | 4.92E-162 | 10.427133 | Airway Smooth Muscle |
| 59 | ACTN1 | 3.31E-160 | 3.902293 | Airway Smooth Muscle |
| 60 | DCN | 7.26E-160 | 0.9263545 | Airway Smooth Muscle |
| 61 | TPPP3 | 2.87E-159 | 3.646296 | Airway Smooth Muscle |
| 62 | COX4I2 | 1.45E-158 | 4.8170614 | Airway Smooth Muscle |
| 63 | ISYNA1 | 2.41E-152 | 4.943556 | Airway Smooth Muscle |
| 64 | PRELP | 6.48E-149 | 4.177516 | Airway Smooth Muscle |
| 65 | MYH10 | 2.64E-148 | 5.3978505 | Airway Smooth Muscle |
| 66 | PPP1R12A | 4.67E-148 | 3.7089984 | Airway Smooth Muscle |
| 67 | TNNT2 | 1.58E-145 | 9.989634 | Airway Smooth Muscle |
| 68 | WISP2 | 4.70E-145 | 7.2985277 | Airway Smooth Muscle |
| 69 | ELN | 2.06E-138 | 5.0114236 | Airway Smooth Muscle |
| 70 | LPP | 2.46E-138 | 3.8854244 | Airway Smooth Muscle |
| 71 | SNCG | 1.93E-135 | 6.764453 | Airway Smooth Muscle |
| 72 | ITM2C | 2.22E-135 | 3.157348 | Airway Smooth Muscle |
| 73 | NTN4 | 6.56E-134 | 4.1441684 | Airway Smooth Muscle |
| 74 | RASL12 | 1.94E-132 | 6.2026114 | Airway Smooth Muscle |
| 75 | SGCA | 8.93E-131 | 5.89903 | Airway Smooth Muscle |
| 76 | PLS3 | 1.45E-129 | 3.7163198 | Airway Smooth Muscle |
| 77 | TIMP3 | 9.14E-127 | 3.2198849 | Airway Smooth Muscle |
| 78 | NOTCH3 | 1.70E-126 | 4.998761 | Airway Smooth Muscle |
| 79 | PALLD | 1.46E-124 | 4.5238605 | Airway Smooth Muscle |
| 80 | COL18A1 | 6.40E-123 | 4.556103 | Airway Smooth Muscle |
| 81 | FILIP1L | 6.85E-123 | 3.373886 | Airway Smooth Muscle |
| 82 | SMTN | 4.69E-122 | 4.994295 | Airway Smooth Muscle |

| | | | | |
|---|---|---|---|---|
| 83 | PDK4 | 5.11E-122 | 5.0992565 | Airway Smooth Muscle |
| 84 | CAV2 | 8.27E-120 | 2.6753592 | Airway Smooth Muscle |
| 85 | RCAN2 | 3.72E-119 | 5.9102206 | Airway Smooth Muscle |
| 86 | COL6A1 | 4.47E-119 | 3.6624568 | Airway Smooth Muscle |
| 87 | SBDS | 1.19E-118 | 2.8742237 | Airway Smooth Muscle |
| 88 | GPM6B | 3.56E-115 | 4.3879414 | Airway Smooth Muscle |
| 89 | CPE | 4.61E-115 | 4.2257733 | Airway Smooth Muscle |
| 90 | ADAMTS1 | 1.60E-113 | 4.3578615 | Airway Smooth Muscle |
| 91 | HSPA2 | 1.32E-112 | 4.881104 | Airway Smooth Muscle |
| 92 | TSC22D1 | 1.07E-111 | 4.877411 | Airway Smooth Muscle |
| 93 | C9orf3 | 3.57E-111 | 3.723304 | Airway Smooth Muscle |
| 94 | MRGPRF | 1.79E-110 | 5.6677876 | Airway Smooth Muscle |
| 95 | PPP1R12B | 2.65E-109 | 4.8665066 | Airway Smooth Muscle |
| 96 | ZAK | 5.30E-108 | 3.8398407 | Airway Smooth Muscle |
| 97 | GPX3 | 3.37E-107 | 4.317288 | Airway Smooth Muscle |
| 98 | ITGB1 | 8.69E-105 | 3.2067811 | Airway Smooth Muscle |
| 99 | FHL1 | 2.29E-104 | 4.1714497 | Airway Smooth Muscle |
| 0 | AGER | 0 | 67.772896 | Alveolar Epithelial Type 1 |
| 20 | GPRC5A | 0 | 7.4912076 | Alveolar Epithelial Type 1 |
| 21 | CLIC3 | 0 | 7.7435164 | Alveolar Epithelial Type 1 |
| 22 | TNNC1 | 0 | 8.460309 | Alveolar Epithelial Type 1 |
| 23 | CEACAM6 | 0 | 7.9843545 | Alveolar Epithelial Type 1 |
| 25 | CAV2 | 0 | 6.623309 | Alveolar Epithelial Type 1 |
| 26 | EPCAM | 0 | 5.6774735 | Alveolar Epithelial Type 1 |
| 27 | UPK3B | 0 | 8.630625 | Alveolar Epithelial Type 1 |
| 28 | SPOCK2 | 0 | 6.250268 | Alveolar Epithelial Type 1 |
| 29 | RNASE1 | 0 | 13.671409 | Alveolar Epithelial Type 1 |
| 30 | RAB11FIP1 | 0 | 5.9199643 | Alveolar Epithelial Type 1 |
| 31 | CD55 | 0 | 10.153559 | Alveolar Epithelial Type 1 |
| 32 | SFTA2 | 0 | 3.6679513 | Alveolar Epithelial Type 1 |
| 33 | CLIC5 | 0 | 6.2107 | Alveolar Epithelial Type 1 |
| 34 | C19orf33 | 0 | 4.878337 | Alveolar Epithelial Type 1 |
| 35 | TNFRSF12A | 0 | 6.3024073 | Alveolar Epithelial Type 1 |
| 19 | RTKN2 | 0 | 11.545992 | Alveolar Epithelial Type 1 |
| 18 | KRT8 | 0 | 11.5246935 | Alveolar Epithelial Type 1 |
| 24 | SCEL | 0 | 9.1905365 | Alveolar Epithelial Type 1 |
| 16 | SLC39A8 | 0 | 6.761143 | Alveolar Epithelial Type 1 |
| 1 | EMP2 | 0 | 68.85682 | Alveolar Epithelial Type 1 |
| 2 | KRT7 | 0 | 20.47263 | Alveolar Epithelial Type 1 |
| 17 | FOLR1 | 0 | 9.144173 | Alveolar Epithelial Type 1 |
| 4 | CYP4B1 | 0 | 14.906934 | Alveolar Epithelial Type 1 |
| 5 | TSPAN13 | 0 | 10.43805 | Alveolar Epithelial Type 1 |
| 6 | CAV1 | 0 | 32.119835 | Alveolar Epithelial Type 1 |
| 7 | KRT19 | 0 | 13.679434 | Alveolar Epithelial Type 1 |
| 8 | CLDN18 | 0 | 9.721343 | Alveolar Epithelial Type 1 |
| 3 | LMO7 | 0 | 10.441554 | Alveolar Epithelial Type 1 |
| 10 | TACSTD2 | 0 | 11.16712 | Alveolar Epithelial Type 1 |
| 11 | FXYD3 | 0 | 8.086205 | Alveolar Epithelial Type 1 |
| 9 | KRT18 | 0 | 19.811005 | Alveolar Epithelial Type 1 |
| 12 | HOPX | 0 | 12.979723 | Alveolar Epithelial Type 1 |
| 13 | MYL9 | 0 | 8.512915 | Alveolar Epithelial Type 1 |
| 14 | AQP4 | 0 | 7.6832604 | Alveolar Epithelial Type 1 |
| 15 | ANXA3 | 0 | 8.554716 | Alveolar Epithelial Type 1 |
| 36 | CLDN4 | 1.33E-307 | 5.877732 | Alveolar Epithelial Type 1 |
| 37 | CST6 | 6.91E-305 | 6.3769884 | Alveolar Epithelial Type 1 |
| 38 | ADIRF | 1.13E-304 | 10.848875 | Alveolar Epithelial Type 1 |
| 39 | SUSD2 | 6.09E-304 | 5.566758 | Alveolar Epithelial Type 1 |
| 40 | SEMA3B | 9.16E-288 | 6.267104 | Alveolar Epithelial Type 1 |
| 41 | NKX2-1 | 1.65E-284 | 5.513018 | Alveolar Epithelial Type 1 |
| 42 | CADM1 | 5.06E-270 | 4.535479 | Alveolar Epithelial Type 1 |
| 43 | VEGFA | 1.55E-261 | 4.961303 | Alveolar Epithelial Type 1 |
| 44 | LIMCH1 | 1.48E-260 | 4.2587385 | Alveolar Epithelial Type 1 |
| 45 | GAS6 | 7.32E-258 | 4.1087317 | Alveolar Epithelial Type 1 |
| 46 | ICAM1 | 1.73E-257 | 7.4000793 | Alveolar Epithelial Type 1 |

| | | | | |
|---|---|---|---|---|
| 47 | PEBP4 | 1.29E-254 | 3.4650965 | Alveolar Epithelial Type 1 |
| 48 | PCYOX1 | 1.21E-251 | 4.936222 | Alveolar Epithelial Type 1 |
| 49 | CYR61 | 1.28E-249 | 6.46074 | Alveolar Epithelial Type 1 |
| 50 | SFTA1P | 8.73E-249 | 5.673573 | Alveolar Epithelial Type 1 |
| 51 | CLDN7 | 1.82E-240 | 3.589246 | Alveolar Epithelial Type 1 |
| 52 | PRSS8 | 1.40E-236 | 4.881097 | Alveolar Epithelial Type 1 |
| 53 | EFEMP1 | 1.31E-228 | 3.4010503 | Alveolar Epithelial Type 1 |
| 54 | BCAM | 1.19E-223 | 3.4679902 | Alveolar Epithelial Type 1 |
| 55 | EFNA1 | 2.46E-223 | 4.450241 | Alveolar Epithelial Type 1 |
| 56 | SDC1 | 1.29E-221 | 4.5373716 | Alveolar Epithelial Type 1 |
| 57 | SPINT2 | 1.82E-220 | 4.0224795 | Alveolar Epithelial Type 1 |
| 58 | VSIG2 | 3.76E-219 | 4.3382874 | Alveolar Epithelial Type 1 |
| 59 | AGR3 | 1.44E-217 | 2.7502987 | Alveolar Epithelial Type 1 |
| 60 | CKB | 7.87E-215 | 3.2555206 | Alveolar Epithelial Type 1 |
| 61 | HSD17B6 | 1.36E-212 | 5.0904636 | Alveolar Epithelial Type 1 |
| 62 | SELENBP1 | 6.39E-208 | 2.7867942 | Alveolar Epithelial Type 1 |
| 63 | CD151 | 1.58E-205 | 3.8632143 | Alveolar Epithelial Type 1 |
| 64 | EZR | 2.28E-203 | 4.0120683 | Alveolar Epithelial Type 1 |
| 65 | ANOS1 | 1.11E-202 | 4.9497094 | Alveolar Epithelial Type 1 |
| 66 | CD9 | 2.05E-197 | 7.4666615 | Alveolar Epithelial Type 1 |
| 67 | SEPP1 | 6.88E-197 | 2.615742 | Alveolar Epithelial Type 1 |
| 68 | GKN2 | 2.98E-192 | 4.517986 | Alveolar Epithelial Type 1 |
| 69 | PLLP | 3.05E-192 | 3.9124482 | Alveolar Epithelial Type 1 |
| 70 | KLK11 | 2.82E-190 | 3.5091002 | Alveolar Epithelial Type 1 |
| 71 | LAMB3 | 3.39E-190 | 4.3757243 | Alveolar Epithelial Type 1 |
| 72 | MGLL | 1.40E-188 | 3.6169255 | Alveolar Epithelial Type 1 |
| 73 | PERP | 8.01E-187 | 2.0492592 | Alveolar Epithelial Type 1 |
| 74 | NCKAP5 | 1.09E-183 | 6.146842 | Alveolar Epithelial Type 1 |
| 75 | ALCAM | 3.93E-178 | 2.7794569 | Alveolar Epithelial Type 1 |
| 76 | MSLN | 5.72E-178 | 4.690785 | Alveolar Epithelial Type 1 |
| 77 | SFTA3 | 5.93E-175 | 2.4172134 | Alveolar Epithelial Type 1 |
| 78 | KLF6 | 1.74E-174 | 7.7686615 | Alveolar Epithelial Type 1 |
| 79 | NEDD4L | 1.76E-170 | 3.7502549 | Alveolar Epithelial Type 1 |
| 80 | MS4A15 | 1.08E-169 | 7.1474824 | Alveolar Epithelial Type 1 |
| 81 | CXADR | 6.80E-167 | 3.3688207 | Alveolar Epithelial Type 1 |
| 82 | CYSTM1 | 5.83E-164 | 3.1111443 | Alveolar Epithelial Type 1 |
| 83 | ARHGEF26 | 3.50E-163 | 5.681864 | Alveolar Epithelial Type 1 |
| 84 | ID1 | 7.41E-162 | 3.219477 | Alveolar Epithelial Type 1 |
| 85 | WFS1 | 1.57E-161 | 4.432401 | Alveolar Epithelial Type 1 |
| 86 | APLP2 | 4.31E-158 | 3.3395534 | Alveolar Epithelial Type 1 |
| 87 | C4BPA | 1.68E-157 | 4.3980517 | Alveolar Epithelial Type 1 |
| 88 | DSTN | 2.61E-156 | 4.2200885 | Alveolar Epithelial Type 1 |
| 89 | TMEM265 | 1.42E-154 | 3.5082679 | Alveolar Epithelial Type 1 |
| 90 | S100A10 | 2.06E-154 | 10.015872 | Alveolar Epithelial Type 1 |
| 91 | GGTLC1 | 3.42E-153 | 7.508692 | Alveolar Epithelial Type 1 |
| 92 | NDNF | 4.52E-153 | 4.2247853 | Alveolar Epithelial Type 1 |
| 93 | ELF3 | 2.90E-150 | 2.7021592 | Alveolar Epithelial Type 1 |
| 94 | GJA1 | 3.28E-147 | 3.8569696 | Alveolar Epithelial Type 1 |
| 95 | PDPN | 6.25E-146 | 4.778554 | Alveolar Epithelial Type 1 |
| 96 | FMO2 | 5.40E-145 | 3.4504552 | Alveolar Epithelial Type 1 |
| 97 | PDZK1IP1 | 1.92E-144 | 3.2792776 | Alveolar Epithelial Type 1 |
| 98 | NEDD9 | 2.34E-144 | 2.8154182 | Alveolar Epithelial Type 1 |
| 99 | EPB41L5 | 1.11E-143 | 5.006677 | Alveolar Epithelial Type 1 |
| 0 | NAPSA | 0 | 79.829475 | Alveolar Epithelial Type 2 |
| 72 | ADIRF | 0 | 8.637775 | Alveolar Epithelial Type 2 |
| 71 | CHI3L2 | 0 | 8.860499 | Alveolar Epithelial Type 2 |
| 70 | SPTSSA | 0 | 4.9943185 | Alveolar Epithelial Type 2 |
| 69 | HSD17B6 | 0 | 6.222491 | Alveolar Epithelial Type 2 |
| 68 | F3 | 0 | 5.9882402 | Alveolar Epithelial Type 2 |
| 67 | NKX2-1 | 0 | 5.826446 | Alveolar Epithelial Type 2 |
| 66 | TPPP3 | 0 | 6.937574 | Alveolar Epithelial Type 2 |
| 65 | PLA2G1B | 0 | 8.615446 | Alveolar Epithelial Type 2 |
| 64 | C8orf4 | 0 | 16.00474 | Alveolar Epithelial Type 2 |
| 63 | MFSD2A | 0 | 7.2062516 | Alveolar Epithelial Type 2 |

| | | | | |
|---|---|---|---|---|
| 62 | MID1IP1 | 0 | 7.689152 | Alveolar Epithelial Type 2 |
| 61 | DBI | 0 | 15.011909 | Alveolar Epithelial Type 2 |
| 60 | CXCL2 | 0 | 35.106636 | Alveolar Epithelial Type 2 |
| 59 | CRTAC1 | 0 | 7.4594846 | Alveolar Epithelial Type 2 |
| 58 | SCGB3A2 | 0 | 19.13874 | Alveolar Epithelial Type 2 |
| 57 | EPHX1 | 0 | 6.414254 | Alveolar Epithelial Type 2 |
| 56 | CLDN7 | 0 | 5.531591 | Alveolar Epithelial Type 2 |
| 55 | MRPL14 | 0 | 6.0789266 | Alveolar Epithelial Type 2 |
| 54 | DHCR24 | 0 | 5.5132637 | Alveolar Epithelial Type 2 |
| 53 | FGGY | 0 | 6.609035 | Alveolar Epithelial Type 2 |
| 52 | C11orf96 | 0 | 9.499074 | Alveolar Epithelial Type 2 |
| 73 | TSPAN13 | 0 | 4.4142137 | Alveolar Epithelial Type 2 |
| 51 | FXYD3 | 0 | 4.050796 | Alveolar Epithelial Type 2 |
| 74 | MAL2 | 0 | 4.8889656 | Alveolar Epithelial Type 2 |
| 76 | SLC22A31 | 0 | 7.245191 | Alveolar Epithelial Type 2 |
| 97 | AREG | 0 | 13.014033 | Alveolar Epithelial Type 2 |
| 96 | LRRC75A-AS1 | 0 | 7.792867 | Alveolar Epithelial Type 2 |
| 95 | FASN | 0 | 5.9372425 | Alveolar Epithelial Type 2 |
| 94 | NNMT | 0 | 14.77177 | Alveolar Epithelial Type 2 |
| 93 | XBP1 | 0 | 7.561223 | Alveolar Epithelial Type 2 |
| 92 | MBIP | 0 | 5.1519094 | Alveolar Epithelial Type 2 |
| 91 | CRNDE | 0 | 4.1258554 | Alveolar Epithelial Type 2 |
| 90 | ATP1A1 | 0 | 4.795853 | Alveolar Epithelial Type 2 |
| 89 | SPINT2 | 0 | 4.5833964 | Alveolar Epithelial Type 2 |
| 88 | GSTP1 | 0 | 13.728187 | Alveolar Epithelial Type 2 |
| 87 | PTP4A1 | 0 | 4.541898 | Alveolar Epithelial Type 2 |
| 86 | NRGN | 0 | 5.791453 | Alveolar Epithelial Type 2 |
| 85 | TMEM125 | 0 | 5.457989 | Alveolar Epithelial Type 2 |
| 84 | KRT19 | 0 | 1.9204227 | Alveolar Epithelial Type 2 |
| 83 | C14orf1 | 0 | 4.8119144 | Alveolar Epithelial Type 2 |
| 82 | STEAP4 | 0 | 7.742331 | Alveolar Epithelial Type 2 |
| 81 | RNF145 | 0 | 5.0045414 | Alveolar Epithelial Type 2 |
| 80 | CA2 | 0 | 6.153521 | Alveolar Epithelial Type 2 |
| 79 | RGS16 | 0 | 6.1994643 | Alveolar Epithelial Type 2 |
| 78 | TMEM243 | 0 | 4.6601295 | Alveolar Epithelial Type 2 |
| 77 | CLDN4 | 0 | 4.9240932 | Alveolar Epithelial Type 2 |
| 75 | DCXR | 0 | 4.7517514 | Alveolar Epithelial Type 2 |
| 50 | TSC22D1 | 0 | 12.370776 | Alveolar Epithelial Type 2 |
| 49 | AGR3 | 0 | 8.07288 | Alveolar Epithelial Type 2 |
| 48 | TSTD1 | 0 | 5.161595 | Alveolar Epithelial Type 2 |
| 21 | CYB5A | 0 | 27.706024 | Alveolar Epithelial Type 2 |
| 20 | LAMP3 | 0 | 10.823161 | Alveolar Epithelial Type 2 |
| 19 | MALL | 0 | 11.313709 | Alveolar Epithelial Type 2 |
| 18 | C16orf89 | 0 | 10.919962 | Alveolar Epithelial Type 2 |
| 17 | NPC2 | 0 | 70.0726 | Alveolar Epithelial Type 2 |
| 16 | SELENBP1 | 0 | 11.930705 | Alveolar Epithelial Type 2 |
| 15 | CXCL17 | 0 | 18.430193 | Alveolar Epithelial Type 2 |
| 14 | SFTA3 | 0 | 11.588479 | Alveolar Epithelial Type 2 |
| 13 | AK1 | 0 | 13.65999 | Alveolar Epithelial Type 2 |
| 12 | SLPI | 0 | 85.89338 | Alveolar Epithelial Type 2 |
| 11 | PEBP4 | 0 | 12.466363 | Alveolar Epithelial Type 2 |
| 10 | HOPX | 0 | 17.258358 | Alveolar Epithelial Type 2 |
| 9 | MUC1 | 0 | 16.196512 | Alveolar Epithelial Type 2 |
| 8 | PGC | 0 | 44.917095 | Alveolar Epithelial Type 2 |
| 7 | SLC34A2 | 0 | 16.775818 | Alveolar Epithelial Type 2 |
| 6 | SFTPA1 | 0 | 284.2662 | Alveolar Epithelial Type 2 |
| 5 | SFTPA2 | 0 | 242.87665 | Alveolar Epithelial Type 2 |
| 4 | SFTA2 | 0 | 26.512468 | Alveolar Epithelial Type 2 |
| 3 | SFTPD | 0 | 57.89353 | Alveolar Epithelial Type 2 |
| 2 | SFTPB | 0 | 182.94696 | Alveolar Epithelial Type 2 |
| 1 | SFTPC | 0 | inf | Alveolar Epithelial Type 2 |
| 22 | S100A14 | 0 | 8.581541 | Alveolar Epithelial Type 2 |
| 23 | C4BPA | 0 | 10.393109 | Alveolar Epithelial Type 2 |
| 24 | ABCA3 | 0 | 9.331014 | Alveolar Epithelial Type 2 |

| | | | | |
|---|---|---|---|---|
| 25 | RNASE1 | 0 | 21.73822 | Alveolar Epithelial Type 2 |
| 47 | SLC39A8 | 0 | 6.038752 | Alveolar Epithelial Type 2 |
| 46 | AGR2 | 0 | 6.4814186 | Alveolar Epithelial Type 2 |
| 45 | EPCAM | 0 | 5.495193 | Alveolar Epithelial Type 2 |
| 44 | ELF3 | 0 | 7.939418 | Alveolar Epithelial Type 2 |
| 43 | SDC4 | 0 | 7.8030252 | Alveolar Epithelial Type 2 |
| 42 | WFDC2 | 0 | 0.96067876 | Alveolar Epithelial Type 2 |
| 41 | LPCAT1 | 0 | 6.540475 | Alveolar Epithelial Type 2 |
| 40 | CEBPD | 0 | 19.98786 | Alveolar Epithelial Type 2 |
| 39 | NGFRAP1 | 0 | 5.8498588 | Alveolar Epithelial Type 2 |
| 38 | SDR16C5 | 0 | 7.9660306 | Alveolar Epithelial Type 2 |
| 98 | RPL24 | 0 | 12.452266 | Alveolar Epithelial Type 2 |
| 37 | CTSH | 0 | 19.030994 | Alveolar Epithelial Type 2 |
| 35 | KRT8 | 0 | 8.074457 | Alveolar Epithelial Type 2 |
| 34 | MGST1 | 0 | 10.08782 | Alveolar Epithelial Type 2 |
| 33 | PRDX5 | 0 | 12.201089 | Alveolar Epithelial Type 2 |
| 32 | SEPP1 | 0 | 7.5143704 | Alveolar Epithelial Type 2 |
| 31 | C3 | 0 | 5.953322 | Alveolar Epithelial Type 2 |
| 30 | FOLR1 | 0 | 7.276298 | Alveolar Epithelial Type 2 |
| 29 | WIF1 | 0 | 10.780049 | Alveolar Epithelial Type 2 |
| 28 | LRRK2 | 0 | 8.465556 | Alveolar Epithelial Type 2 |
| 27 | CLDN18 | 0 | 7.473548 | Alveolar Epithelial Type 2 |
| 26 | PIGR | 0 | 8.965513 | Alveolar Epithelial Type 2 |
| 36 | KRT18 | 0 | 8.344837 | Alveolar Epithelial Type 2 |
| 99 | RPS4X | 0 | 26.19505 | Alveolar Epithelial Type 2 |
| 0 | LUM | 0 | 36.642036 | Alveolar Fibroblast |
| 26 | PLAC9 | 0 | 5.147389 | Alveolar Fibroblast |
| 27 | FBLN5 | 0 | 6.189703 | Alveolar Fibroblast |
| 28 | DKK3 | 0 | 5.8842864 | Alveolar Fibroblast |
| 29 | SPARCL1 | 0 | 6.4964085 | Alveolar Fibroblast |
| 30 | TIMP3 | 0 | 10.089243 | Alveolar Fibroblast |
| 31 | LIMCH1 | 0 | 5.2469544 | Alveolar Fibroblast |
| 32 | SCN7A | 0 | 7.419075 | Alveolar Fibroblast |
| 33 | MACF1 | 0 | 5.476509 | Alveolar Fibroblast |
| 34 | PPP1R14A | 0 | 4.7889786 | Alveolar Fibroblast |
| 35 | LMCD1 | 0 | 5.615206 | Alveolar Fibroblast |
| 36 | CFD | 0 | 16.69229 | Alveolar Fibroblast |
| 37 | MYH10 | 0 | 6.5503564 | Alveolar Fibroblast |
| 38 | FIBIN | 0 | 7.8481483 | Alveolar Fibroblast |
| 39 | COL6A1 | 0 | 5.099807 | Alveolar Fibroblast |
| 40 | OLFML3 | 0 | 6.2446995 | Alveolar Fibroblast |
| 41 | CALD1 | 0 | 3.373605 | Alveolar Fibroblast |
| 42 | PPAP2B | 0 | 4.537241 | Alveolar Fibroblast |
| 43 | COL6A3 | 0 | 5.9740214 | Alveolar Fibroblast |
| 44 | ELN | 0 | 6.297402 | Alveolar Fibroblast |
| 25 | C1R | 0 | 6.0527287 | Alveolar Fibroblast |
| 23 | MMP2 | 0 | 6.7392235 | Alveolar Fibroblast |
| 24 | FHL1 | 0 | 6.4664073 | Alveolar Fibroblast |
| 21 | FMO2 | 0 | 6.589103 | Alveolar Fibroblast |
| 1 | DCN | 0 | 33.629066 | Alveolar Fibroblast |
| 2 | RARRES2 | 0 | 16.715433 | Alveolar Fibroblast |
| 3 | MFAP4 | 0 | 21.526491 | Alveolar Fibroblast |
| 4 | A2M | 0 | 21.775763 | Alveolar Fibroblast |
| 5 | SEPP1 | 0 | 17.158995 | Alveolar Fibroblast |
| 6 | CYR61 | 0 | 36.171627 | Alveolar Fibroblast |
| 7 | FBLN1 | 0 | 10.882108 | Alveolar Fibroblast |
| 8 | GPC3 | 0 | 10.7234745 | Alveolar Fibroblast |
| 9 | CTGF | 0 | 31.351013 | Alveolar Fibroblast |
| 22 | LTBP4 | 0 | 5.467371 | Alveolar Fibroblast |
| 10 | PTGDS | 0 | 16.48068 | Alveolar Fibroblast |
| 12 | ADH1B | 0 | 10.207877 | Alveolar Fibroblast |
| 13 | COL6A2 | 0 | 7.8828793 | Alveolar Fibroblast |
| 14 | MGP | 0 | 18.531702 | Alveolar Fibroblast |
| 15 | C1S | 0 | 7.6566405 | Alveolar Fibroblast |

| | | | | |
|---|---|---|---|---|
| 16 | COL1A2 | 0 | 7.268934 | Alveolar Fibroblast |
| 17 | TCF21 | 0 | 8.028131 | Alveolar Fibroblast |
| 18 | PRELP | 0 | 7.3133073 | Alveolar Fibroblast |
| 19 | INMT | 0 | 7.7752414 | Alveolar Fibroblast |
| 20 | SOD3 | 0 | 5.47524 | Alveolar Fibroblast |
| 11 | PMP22 | 0 | 9.02511 | Alveolar Fibroblast |
| 45 | LBH | 1.82E-301 | 4.9376917 | Alveolar Fibroblast |
| 46 | FN1 | 1.46E-299 | 3.017175 | Alveolar Fibroblast |
| 47 | FGFR4 | 1.53E-297 | 7.837258 | Alveolar Fibroblast |
| 48 | C7 | 1.30E-296 | 4.6153493 | Alveolar Fibroblast |
| 49 | AOC3 | 2.03E-283 | 3.8399963 | Alveolar Fibroblast |
| 50 | EMILIN1 | 1.08E-280 | 5.8122516 | Alveolar Fibroblast |
| 51 | MAMDC2 | 1.50E-270 | 7.1570964 | Alveolar Fibroblast |
| 52 | PCOLCE | 2.35E-267 | 4.4232383 | Alveolar Fibroblast |
| 53 | DST | 4.31E-259 | 3.323519 | Alveolar Fibroblast |
| 54 | SLIT2 | 5.63E-258 | 6.35598 | Alveolar Fibroblast |
| 55 | CDH11 | 2.41E-250 | 6.703971 | Alveolar Fibroblast |
| 56 | MXRA8 | 3.53E-244 | 4.895027 | Alveolar Fibroblast |
| 57 | COL1A1 | 8.50E-244 | 4.27101 | Alveolar Fibroblast |
| 58 | FXYD1 | 1.40E-236 | 3.9204586 | Alveolar Fibroblast |
| 59 | NKD2 | 3.41E-234 | 6.822007 | Alveolar Fibroblast |
| 60 | RGCC | 2.64E-231 | 10.488411 | Alveolar Fibroblast |
| 61 | ANGPT1 | 9.39E-229 | 6.0818615 | Alveolar Fibroblast |
| 62 | G0S2 | 5.16E-226 | 5.1586037 | Alveolar Fibroblast |
| 63 | AEBP1 | 8.17E-215 | 4.3391814 | Alveolar Fibroblast |
| 64 | GDF10 | 4.40E-207 | 7.4981923 | Alveolar Fibroblast |
| 65 | ADAMTS8 | 1.37E-206 | 6.1786604 | Alveolar Fibroblast |
| 66 | GPM6B | 2.81E-206 | 4.418919 | Alveolar Fibroblast |
| 67 | COL3A1 | 1.84E-202 | 3.9917827 | Alveolar Fibroblast |
| 68 | PALLD | 4.43E-196 | 3.612608 | Alveolar Fibroblast |
| 69 | SERPING1 | 6.91E-196 | 0.90664315 | Alveolar Fibroblast |
| 70 | SPARC | 1.33E-195 | 2.6397128 | Alveolar Fibroblast |
| 71 | ITGA8 | 2.07E-195 | 6.2658677 | Alveolar Fibroblast |
| 72 | CES1 | 3.28E-192 | -0.49991706 | Alveolar Fibroblast |
| 73 | NPNT | 7.85E-191 | 4.377128 | Alveolar Fibroblast |
| 74 | GPX3 | 1.69E-190 | 4.292093 | Alveolar Fibroblast |
| 75 | CSRP1 | 4.54E-189 | 2.780686 | Alveolar Fibroblast |
| 76 | PLTP | 2.60E-181 | 3.3190515 | Alveolar Fibroblast |
| 77 | BGN | 1.33E-179 | 2.474068 | Alveolar Fibroblast |
| 78 | ARID5B | 5.00E-179 | 3.0702195 | Alveolar Fibroblast |
| 79 | IGFBP7 | 2.48E-177 | 6.318987 | Alveolar Fibroblast |
| 80 | SLC40A1 | 2.57E-173 | 3.7127457 | Alveolar Fibroblast |
| 81 | LAMA4 | 4.51E-170 | 4.3592896 | Alveolar Fibroblast |
| 82 | CCBE1 | 2.30E-166 | 6.371837 | Alveolar Fibroblast |
| 83 | TMEM176B | 1.85E-165 | 2.7953506 | Alveolar Fibroblast |
| 84 | PDLIM3 | 2.41E-165 | 3.1990283 | Alveolar Fibroblast |
| 85 | RGS3 | 1.28E-158 | 3.8327389 | Alveolar Fibroblast |
| 86 | DPT | 9.46E-157 | 4.927806 | Alveolar Fibroblast |
| 87 | MFGE8 | 1.46E-152 | 2.6990707 | Alveolar Fibroblast |
| 88 | ABI3BP | 9.36E-152 | 3.7849448 | Alveolar Fibroblast |
| 89 | HSD11B1 | 7.91E-148 | 5.0937276 | Alveolar Fibroblast |
| 90 | CCDC102B | 1.32E-147 | 4.4649487 | Alveolar Fibroblast |
| 91 | TSPAN8 | 9.54E-146 | 2.402031 | Alveolar Fibroblast |
| 92 | EMP2 | 4.90E-140 | 0.86161697 | Alveolar Fibroblast |
| 93 | CFH | 2.60E-137 | 3.1505556 | Alveolar Fibroblast |
| 94 | CRYAB | 1.39E-136 | 3.0430806 | Alveolar Fibroblast |
| 95 | LTBP2 | 1.47E-136 | 5.001603 | Alveolar Fibroblast |
| 96 | PLEKHH2 | 3.49E-135 | 5.7606335 | Alveolar Fibroblast |
| 97 | MALAT1 | 6.99E-134 | 139.298 | Alveolar Fibroblast |
| 98 | SERPINA3 | 1.67E-133 | 4.16497 | Alveolar Fibroblast |
| 99 | MYL9 | 9.97E-131 | 0.31402287 | Alveolar Fibroblast |
| 0 | MGP | 0 | 31.41044 | Artery |
| 31 | CRIP2 | 0 | 6.561514 | Artery |
| 32 | GPX3 | 0 | 10.580815 | Artery |

| | | | | |
|---|---|---|---|---|
| 33 | CAV1 | 0 | 9.947057 | Artery |
| 34 | RNASE1 | 0 | 8.471519 | Artery |
| 35 | HYAL2 | 0 | 7.5323887 | Artery |
| 36 | HEY1 | 0 | 7.048649 | Artery |
| 37 | JAM2 | 0 | 4.970908 | Artery |
| 38 | TIMP3 | 0 | 11.18199 | Artery |
| 39 | A2M | 0 | 4.5260634 | Artery |
| 40 | SPARCL1 | 0 | 7.7574396 | Artery |
| 41 | GIMAP7 | 0 | 5.733545 | Artery |
| 42 | MMRN2 | 0 | 5.3853693 | Artery |
| 43 | MT1E | 0 | 25.020067 | Artery |
| 44 | CD93 | 0 | 5.1236324 | Artery |
| 45 | ENG | 0 | 4.864122 | Artery |
| 46 | ID3 | 0 | 6.7092423 | Artery |
| 47 | DKK2 | 0 | 10.130396 | Artery |
| 48 | IFITM2 | 0 | 7.5356445 | Artery |
| 50 | EMP1 | 0 | 12.292829 | Artery |
| 51 | LIFR | 0 | 4.6291456 | Artery |
| 52 | AQP1 | 0 | 5.5760374 | Artery |
| 53 | PCAT19 | 0 | 4.398335 | Artery |
| 54 | LRRC32 | 0 | 4.8895044 | Artery |
| 55 | S1PR1 | 0 | 4.8931284 | Artery |
| 56 | EDN1 | 0 | 9.235442 | Artery |
| 30 | IFI27 | 0 | 22.346008 | Artery |
| 29 | NPDC1 | 0 | 5.0436916 | Artery |
| 49 | PTRF | 0 | 4.5987797 | Artery |
| 27 | SRPX | 0 | 6.3396974 | Artery |
| 1 | CLEC3B | 0 | 14.502098 | Artery |
| 2 | C10orf10 | 0 | 28.413794 | Artery |
| 3 | EPAS1 | 0 | 16.876795 | Artery |
| 4 | FAM107A | 0 | 11.273986 | Artery |
| 5 | CLDN5 | 0 | 28.748621 | Artery |
| 6 | CLEC14A | 0 | 9.890047 | Artery |
| 7 | CTNNAL1 | 0 | 8.061562 | Artery |
| 28 | SLC9A3R2 | 0 | 8.843653 | Artery |
| 8 | RAMP2 | 0 | 11.604131 | Artery |
| 9 | IFITM3 | 0 | 23.50929 | Artery |
| 11 | PTPRB | 0 | 6.6997423 | Artery |
| 12 | IGFBP4 | 0 | 7.665243 | Artery |
| 13 | IFITM1 | 0 | 13.617252 | Artery |
| 10 | TFPI | 0 | 7.300688 | Artery |
| 15 | LDB2 | 0 | 5.544684 | Artery |
| 26 | TMEM100 | 0 | 11.546019 | Artery |
| 25 | MT1M | 0 | 23.117567 | Artery |
| 24 | IL33 | 0 | 6.1803193 | Artery |
| 14 | GJA5 | 0 | 8.425084 | Artery |
| 22 | SOX17 | 0 | 8.767679 | Artery |
| 21 | CALCRL | 0 | 6.004264 | Artery |
| 23 | ID1 | 0 | 12.893043 | Artery |
| 19 | TSPAN7 | 0 | 6.1658463 | Artery |
| 18 | ARGLU1 | 0 | 7.0617347 | Artery |
| 17 | CXCL12 | 0 | 8.578133 | Artery |
| 16 | TM4SF1 | 0 | 19.175838 | Artery |
| 20 | VWF | 0 | 5.8208976 | Artery |
| 57 | F8 | 8.36E-308 | 5.118868 | Artery |
| 58 | EFNB2 | 1.03E-305 | 5.396312 | Artery |
| 59 | IGFBP7 | 5.57E-302 | 5.225314 | Artery |
| 60 | BST2 | 1.56E-301 | 6.549067 | Artery |
| 61 | PECAM1 | 3.19E-295 | 5.1193657 | Artery |
| 62 | PRSS23 | 1.08E-294 | 3.9566152 | Artery |
| 63 | MT1A | 1.41E-294 | 21.538351 | Artery |
| 64 | GNG11 | 3.38E-293 | 3.4347408 | Artery |
| 65 | TNFSF10 | 1.62E-292 | 7.6644692 | Artery |
| 66 | MT1X | 3.01E-291 | 33.934517 | Artery |

| | | | | |
|---|---|---|---|---|
| 67 | CDH5 | 3.50E-288 | 3.542284 | Artery |
| 68 | NRN1 | 8.99E-287 | 4.96179 | Artery |
| 69 | PIK3R3 | 1.36E-285 | 5.0478616 | Artery |
| 70 | PALMD | 1.62E-285 | 4.8611326 | Artery |
| 71 | MT2A | 3.18E-284 | 94.44392 | Artery |
| 72 | ENPP2 | 5.88E-269 | 7.130604 | Artery |
| 73 | EGFL7 | 3.71E-267 | 4.3845706 | Artery |
| 74 | ARHGAP29 | 1.70E-262 | 3.554143 | Artery |
| 75 | ITM2A | 2.28E-261 | 3.699993 | Artery |
| 76 | ESAM | 2.59E-260 | 3.121281 | Artery |
| 77 | PLLP | 1.38E-256 | 4.0746236 | Artery |
| 78 | SDPR | 3.06E-256 | 5.0234838 | Artery |
| 79 | IDO1 | 3.50E-246 | 5.8753223 | Artery |
| 80 | NFIB | 1.85E-245 | 3.5438612 | Artery |
| 81 | ASRGL1 | 2.04E-243 | 3.9817648 | Artery |
| 82 | ICAM2 | 1.21E-239 | 4.043078 | Artery |
| 83 | ECSCR | 1.76E-239 | 3.213231 | Artery |
| 84 | STOM | 5.82E-233 | 4.208291 | Artery |
| 85 | IGFBP3 | 4.35E-226 | 6.802594 | Artery |
| 86 | CSRNP1 | 9.15E-225 | 3.4485903 | Artery |
| 87 | GIMAP4 | 6.37E-221 | 3.4806092 | Artery |
| 88 | GPIHBP1 | 1.07E-218 | 3.7238247 | Artery |
| 89 | GIMAP8 | 3.55E-217 | 3.9654627 | Artery |
| 90 | CAV2 | 1.12E-216 | 2.7458513 | Artery |
| 91 | RPGR | 1.38E-213 | 3.9075353 | Artery |
| 92 | TGM2 | 2.38E-211 | 3.3037584 | Artery |
| 93 | BCAM | 4.87E-211 | 2.4673386 | Artery |
| 94 | LTC4S | 9.13E-211 | 3.125249 | Artery |
| 95 | KLF4 | 1.58E-208 | 4.955874 | Artery |
| 96 | SERPINE2 | 4.67E-206 | 7.252672 | Artery |
| 97 | CXCL2 | 6.12E-203 | 20.379177 | Artery |
| 98 | TCF4 | 8.17E-202 | 3.1492383 | Artery |
| 99 | FKBP1A | 1.17E-199 | 6.4111643 | Artery |
| 0 | CD79A | 0 | 13.534601 | B |
| 1 | CD79B | 0 | 9.372321 | B |
| 2 | MS4A1 | 0 | 11.888463 | B |
| 3 | LTB | 0 | 10.247002 | B |
| 4 | PTPRCAP | 4.36E-280 | 5.3454423 | B |
| 5 | LINC00926 | 1.65E-263 | 10.60311 | B |
| 6 | CD37 | 5.29E-256 | 7.809749 | B |
| 7 | CXCR4 | 2.57E-240 | 7.260862 | B |
| 8 | RCSD1 | 5.80E-239 | 5.2226524 | B |
| 9 | RPS27 | 3.94E-233 | 57.562492 | B |
| 10 | BANK1 | 4.57E-224 | 7.648177 | B |
| 11 | LIMD2 | 5.01E-223 | 4.3081923 | B |
| 12 | RPL13AP5 | 1.50E-206 | 5.0155373 | B |
| 13 | CORO1A | 4.84E-197 | 3.6294472 | B |
| 14 | FCMR | 6.51E-188 | 4.9578757 | B |
| 15 | RPS29 | 5.98E-176 | 28.811335 | B |
| 16 | BTG1 | 3.95E-170 | 7.0994377 | B |
| 17 | RPL13A | 7.05E-165 | 30.099138 | B |
| 18 | MEF2C | 1.42E-156 | 3.7344368 | B |
| 19 | SELL | 2.40E-146 | 4.359569 | B |
| 20 | RPL18A | 5.38E-141 | 19.770096 | B |
| 21 | RPSAP58 | 3.16E-140 | 3.2522352 | B |
| 22 | RPL39 | 5.48E-140 | 21.130396 | B |
| 23 | RPL23A | 6.69E-139 | 17.874073 | B |
| 24 | RPLP2 | 9.83E-139 | 23.452675 | B |
| 25 | RPL17 | 6.65E-135 | 6.8799257 | B |
| 26 | RPS8 | 7.49E-133 | 14.96179 | B |
| 27 | RPS23 | 3.72E-129 | 15.895949 | B |
| 28 | EEF2 | 8.66E-129 | 5.263129 | B |
| 29 | TCL1A | 2.09E-128 | 10.250676 | B |
| 30 | RPL34 | 3.69E-128 | 21.723478 | B |

| | | | | |
|---|---|---|---|---|
| 31 | RPL13 | 2.95E-126 | 23.066357 | B |
| 32 | CD48 | 7.77E-126 | 2.7782059 | B |
| 33 | RPS10 | 3.82E-124 | 4.5248923 | B |
| 34 | IGLL5 | 1.35E-122 | 14.203314 | B |
| 35 | VPREB3 | 3.72E-122 | 10.004687 | B |
| 36 | RPS15A | 4.28E-122 | 17.444223 | B |
| 37 | CD52 | 2.64E-121 | -3.7121484 | B |
| 38 | RPS21 | 1.69E-118 | 8.499492 | B |
| 39 | RPL37 | 1.22E-117 | 12.370952 | B |
| 40 | RPSA | 9.64E-115 | 6.978536 | B |
| 41 | GLTSCR2 | 1.02E-112 | 3.093588 | B |
| 42 | RPS18 | 9.30E-112 | 23.614197 | B |
| 43 | RPL30 | 1.03E-108 | 11.915379 | B |
| 44 | EEF1G | 1.57E-108 | 6.7391253 | B |
| 45 | RPL21 | 3.53E-105 | 21.475924 | B |
| 46 | RPS6 | 2.05E-104 | 14.667734 | B |
| 47 | BLK | 1.79E-103 | 7.820028 | B |
| 48 | FAM65B | 8.50E-101 | 2.493165 | B |
| 49 | RPL32 | 4.84E-99 | 15.883908 | B |
| 50 | CLEC2D | 9.83E-98 | 3.3609812 | B |
| 51 | RPS3A | 1.74E-96 | 12.757253 | B |
| 52 | HLA-DOB | 7.98E-96 | 5.5294094 | B |
| 53 | GAS5 | 1.51E-95 | 2.4183807 | B |
| 54 | RPL36A | 5.45E-95 | 2.9961045 | B |
| 55 | RPL18 | 7.56E-95 | 7.2056904 | B |
| 56 | HVCN1 | 5.65E-94 | 3.1260464 | B |
| 57 | RPL3 | 7.75E-93 | 11.298485 | B |
| 58 | RPS17 | 8.58E-91 | 8.4209795 | B |
| 59 | RPS5 | 9.36E-91 | 6.519855 | B |
| 60 | CD74 | 7.92E-90 | -42.494427 | B |
| 61 | ISG20 | 1.10E-89 | 2.13143 | B |
| 62 | ADAM28 | 7.31E-89 | 4.5163546 | B |
| 63 | RPL36 | 1.38E-87 | 8.255377 | B |
| 64 | RPS11 | 1.99E-87 | 6.424669 | B |
| 65 | IRF8 | 2.01E-87 | 2.0220473 | B |
| 66 | RPL35A | 1.90E-85 | 8.508159 | B |
| 67 | GNG7 | 3.72E-85 | 3.9979885 | B |
| 68 | EEF1B2 | 2.12E-82 | 3.6106193 | B |
| 69 | RPL10A | 3.08E-82 | 6.6416397 | B |
| 70 | FCER2 | 2.07E-80 | 8.906951 | B |
| 71 | RPS27A | 3.71E-80 | 12.217805 | B |
| 72 | FCRLA | 5.18E-79 | 9.048195 | B |
| 73 | RPS25 | 1.25E-78 | 8.9518 | B |
| 74 | FAM26F | 1.71E-78 | 2.3325865 | B |
| 75 | LAPTM5 | 2.62E-78 | -0.1067444 | B |
| 76 | RPS2 | 3.02E-78 | 12.402815 | B |
| 77 | POU2F2 | 7.59E-78 | 3.4695604 | B |
| 78 | PRKCB | 9.33E-78 | 3.3368642 | B |
| 79 | RPL11 | 9.83E-77 | 11.582618 | B |
| 80 | RPS7 | 3.42E-76 | 6.2206306 | B |
| 81 | RPS28 | 1.86E-75 | 10.962184 | B |
| 82 | RPS12 | 3.40E-74 | 13.707031 | B |
| 83 | HLA-DQA1 | 1.49E-72 | -5.6104565 | B |
| 84 | RPS4X | 1.63E-72 | 8.493865 | B |
| 85 | RPL4 | 2.13E-72 | 4.2101464 | B |
| 86 | CD19 | 7.15E-72 | 7.4111476 | B |
| 87 | RPL19 | 1.62E-71 | 9.346989 | B |
| 88 | JCHAIN | 2.01E-71 | 10.052399 | B |
| 89 | RPL38 | 3.13E-71 | 4.9551077 | B |
| 90 | RPL41 | 4.19E-71 | 21.009495 | B |
| 91 | HLA-DRA | 1.13E-69 | -70.458145 | B |
| 92 | HLA-DPB1 | 2.81E-69 | -11.205093 | B |
| 93 | RPL5 | 7.80E-69 | 4.4485035 | B |
| 94 | BIRC3 | 3.31E-68 | 1.5182571 | B |

| | | | | |
|---|---|---|---|---|
| 95 | RPS19 | 4.53E-67 | 8.946502 | B |
| 96 | RPL26 | 3.95E-66 | 8.475488 | B |
| 97 | RPL27A | 6.49E-65 | 9.679267 | B |
| 98 | P2RX5 | 7.68E-65 | 6.2028065 | B |
| 99 | EVI2B | 1.63E-64 | 1.0442598 | B |
| 0 | S100A2 | 8.79E-246 | 43.455116 | Basal |
| 1 | KRT17 | 1.08E-236 | 79.29607 | Basal |
| 2 | TACSTD2 | 1.64E-232 | 20.212557 | Basal |
| 3 | MIR205HG | 3.10E-230 | 13.762443 | Basal |
| 4 | PERP | 1.46E-227 | 12.442894 | Basal |
| 5 | KRT19 | 5.85E-223 | 21.81519 | Basal |
| 6 | SERPINF1 | 2.91E-187 | 10.768625 | Basal |
| 7 | IGFBP2 | 6.84E-187 | 11.680516 | Basal |
| 8 | LAMB3 | 1.62E-184 | 7.696941 | Basal |
| 9 | F3 | 1.03E-180 | 11.2111635 | Basal |
| 10 | SFN | 2.58E-179 | 10.677329 | Basal |
| 11 | BCAM | 7.69E-176 | 10.599791 | Basal |
| 12 | FXYD3 | 3.67E-173 | 7.75931 | Basal |
| 13 | KRT18 | 3.60E-169 | 12.689582 | Basal |
| 14 | MPZL2 | 7.10E-168 | 7.2957134 | Basal |
| 15 | KRT15 | 5.54E-167 | 14.269513 | Basal |
| 16 | RPL3 | 9.48E-167 | 59.799744 | Basal |
| 17 | RPL10A | 6.62E-160 | 37.214756 | Basal |
| 18 | KRT8 | 3.81E-157 | 9.103807 | Basal |
| 19 | KLF5 | 3.19E-156 | 6.4494643 | Basal |
| 20 | TNFRSF12A | 2.65E-154 | 8.777553 | Basal |
| 21 | RPS4X | 3.42E-150 | 49.305862 | Basal |
| 22 | EEF1G | 4.73E-149 | 25.929857 | Basal |
| 23 | RPLP1 | 1.24E-148 | 98.39641 | Basal |
| 24 | SPINT2 | 1.49E-147 | 6.8502994 | Basal |
| 25 | LRRC75A-AS1 | 3.55E-146 | 13.817847 | Basal |
| 26 | HMGB3 | 9.48E-146 | 7.8608255 | Basal |
| 27 | TSC22D1 | 2.48E-143 | 13.394496 | Basal |
| 28 | RPS6 | 1.16E-142 | 56.40619 | Basal |
| 29 | CLU | 1.40E-142 | 11.457367 | Basal |
| 30 | ERRFI1 | 2.88E-142 | 7.659099 | Basal |
| 31 | KRT7 | 3.32E-140 | 10.127526 | Basal |
| 32 | CYP4B1 | 9.58E-139 | 6.0683966 | Basal |
| 33 | FHL2 | 1.52E-138 | 7.1610446 | Basal |
| 34 | RPS18 | 2.39E-136 | 69.456276 | Basal |
| 35 | GAS5 | 6.36E-136 | 7.707518 | Basal |
| 36 | RPL7 | 7.50E-136 | 41.715183 | Basal |
| 37 | CYB5A | 1.01E-135 | 17.879559 | Basal |
| 38 | RPL5 | 1.78E-133 | 22.829107 | Basal |
| 39 | RPLP0 | 6.06E-133 | 23.677717 | Basal |
| 40 | AQP3 | 1.78E-129 | 12.180488 | Basal |
| 41 | RPL13A | 5.03E-129 | 59.105915 | Basal |
| 42 | RPS3 | 1.88E-128 | 40.428802 | Basal |
| 43 | RPS5 | 8.46E-128 | 24.941492 | Basal |
| 44 | HNRNPA1 | 4.42E-127 | 12.783893 | Basal |
| 45 | RPL13 | 1.57E-124 | 57.77933 | Basal |
| 46 | WFDC2 | 2.18E-124 | 11.135712 | Basal |
| 47 | YBX3 | 4.97E-123 | 7.128174 | Basal |
| 48 | RPL8 | 2.15E-122 | 33.696815 | Basal |
| 49 | ID1 | 1.71E-119 | 13.00407 | Basal |
| 50 | MGST1 | 1.95E-119 | 6.2241654 | Basal |
| 51 | KRT5 | 2.97E-119 | 10.444992 | Basal |
| 52 | RPL18 | 3.24E-119 | 22.33046 | Basal |
| 53 | GNB2L1 | 5.87E-118 | 20.854683 | Basal |
| 54 | NPM1 | 2.56E-117 | 11.080929 | Basal |
| 55 | RPS8 | 5.60E-117 | 33.684273 | Basal |
| 56 | NGFRAP1 | 2.48E-116 | 4.215194 | Basal |
| 57 | CXCL17 | 4.04E-116 | 5.3635297 | Basal |
| 58 | RPL4 | 4.98E-116 | 15.550509 | Basal |

| | | | | |
|---|---|---|---|---|
| 59 | C19orf33 | 8.02E-114 | 5.507001 | Basal |
| 60 | SOD3 | 9.77E-114 | 4.3770814 | Basal |
| 61 | RPL7A | 8.87E-113 | 22.895424 | Basal |
| 62 | RPS7 | 1.38E-112 | 22.451649 | Basal |
| 63 | CD9 | 4.23E-112 | 18.700699 | Basal |
| 64 | FOLR1 | 8.65E-112 | 4.0788875 | Basal |
| 65 | IER3 | 4.20E-111 | 8.8421955 | Basal |
| 66 | MYC | 4.51E-110 | 6.9534383 | Basal |
| 67 | ALDH3A1 | 9.72E-110 | 7.5392118 | Basal |
| 68 | S100A14 | 1.77E-108 | 5.4631457 | Basal |
| 69 | RPS9 | 9.25E-108 | 29.674046 | Basal |
| 70 | RPS17 | 2.71E-106 | 31.15056 | Basal |
| 71 | RPS3A | 3.08E-106 | 33.337376 | Basal |
| 72 | CLDN1 | 3.56E-106 | 6.3888516 | Basal |
| 73 | IGFBP7 | 4.33E-106 | 7.842317 | Basal |
| 74 | CD24 | 9.65E-105 | 4.862 | Basal |
| 75 | APOD | 9.78E-105 | 8.905338 | Basal |
| 76 | RPL29 | 1.58E-104 | 22.874916 | Basal |
| 77 | EEF2 | 5.41E-104 | 9.937076 | Basal |
| 78 | RPL15 | 8.11E-104 | 31.059776 | Basal |
| 79 | CEBPD | 1.42E-103 | 14.067302 | Basal |
| 80 | RPS2 | 1.54E-101 | 44.151363 | Basal |
| 81 | ZFP36L1 | 4.61E-101 | 7.041439 | Basal |
| 82 | EPCAM | 5.73E-101 | 3.994414 | Basal |
| 83 | SYT8 | 2.32E-100 | 6.88723 | Basal |
| 84 | GLTSCR2 | 2.89E-99 | 5.515154 | Basal |
| 85 | RPL6 | 3.40E-99 | 21.149128 | Basal |
| 86 | RPL14 | 3.68E-99 | 20.294125 | Basal |
| 87 | SDC1 | 1.38E-98 | 4.6113033 | Basal |
| 88 | IFITM1 | 1.67E-98 | 9.827981 | Basal |
| 89 | OAT | 9.75E-98 | 4.1506658 | Basal |
| 90 | RPL24 | 1.20E-97 | 15.250858 | Basal |
| 91 | RPSA | 4.71E-97 | 14.435332 | Basal |
| 92 | BTF3 | 9.19E-97 | 11.459677 | Basal |
| 93 | HSP90AB1 | 9.98E-96 | 7.979882 | Basal |
| 94 | CFH | 2.97E-95 | 3.9339547 | Basal |
| 95 | RPL10 | 2.02E-94 | 57.137173 | Basal |
| 96 | SFTPB | 2.89E-94 | 21.402617 | Basal |
| 97 | RPL35A | 1.04E-93 | 23.58258 | Basal |
| 98 | CP | 7.68E-93 | 5.5829062 | Basal |
| 99 | IMPDH2 | 7.82E-92 | 3.663555 | Basal |
| 0 | TPSB2 | 0 | 259.41547 | Basophil/Mast 1 |
| 19 | HDC | 0 | 6.7398095 | Basophil/Mast 1 |
| 18 | FCER1A | 0 | 5.1943893 | Basophil/Mast 1 |
| 17 | C1orf186 | 0 | 7.950345 | Basophil/Mast 1 |
| 16 | MAOB | 0 | 6.6890516 | Basophil/Mast 1 |
| 15 | CLU | 0 | 5.164462 | Basophil/Mast 1 |
| 14 | CD69 | 0 | 12.364193 | Basophil/Mast 1 |
| 13 | SAMSN1 | 0 | 6.530434 | Basophil/Mast 1 |
| 12 | GATA2 | 0 | 6.985809 | Basophil/Mast 1 |
| 10 | VWA5A | 0 | 7.640216 | Basophil/Mast 1 |
| 11 | KIT | 0 | 8.484905 | Basophil/Mast 1 |
| 8 | RGS2 | 0 | 18.467585 | Basophil/Mast 1 |
| 7 | LTC4S | 0 | 7.7233615 | Basophil/Mast 1 |
| 6 | RGS13 | 0 | 9.043746 | Basophil/Mast 1 |
| 5 | HPGDS | 0 | 9.611063 | Basophil/Mast 1 |
| 4 | RGS1 | 0 | 17.827732 | Basophil/Mast 1 |
| 3 | MS4A2 | 0 | 12.692408 | Basophil/Mast 1 |
| 2 | CPA3 | 0 | 17.824558 | Basophil/Mast 1 |
| 1 | TPSAB1 | 0 | 174.7702 | Basophil/Mast 1 |
| 9 | SLC18A2 | 0 | 9.099983 | Basophil/Mast 1 |
| 20 | SIGLEC17P | 9.94E-306 | 6.040942 | Basophil/Mast 1 |
| 21 | ITM2C | 2.64E-301 | 3.878491 | Basophil/Mast 1 |
| 22 | CALB2 | 1.76E-283 | 6.577281 | Basophil/Mast 1 |

| | | | | |
|---|---|---|---|---|
| 23 | TSC22D3 | 8.59E-283 | 8.08195 | Basophil/Mast 1 |
| 24 | PTGS1 | 5.77E-274 | 5.884309 | Basophil/Mast 1 |
| 25 | IL1RL1 | 3.46E-267 | 2.1989884 | Basophil/Mast 1 |
| 26 | RAB27B | 1.28E-206 | 5.2486367 | Basophil/Mast 1 |
| 27 | TDRD3 | 2.13E-197 | 3.8219461 | Basophil/Mast 1 |
| 28 | RHOH | 1.27E-193 | 3.4066164 | Basophil/Mast 1 |
| 29 | CAPG | 9.48E-189 | -1.9117155 | Basophil/Mast 1 |
| 30 | RAC2 | 7.47E-187 | 2.4679394 | Basophil/Mast 1 |
| 31 | SRGN | 2.39E-185 | 6.8543673 | Basophil/Mast 1 |
| 32 | CKLF | 9.45E-178 | 2.1250498 | Basophil/Mast 1 |
| 33 | LAPTM4A | 1.41E-176 | 4.002324 | Basophil/Mast 1 |
| 34 | HPGD | 2.98E-175 | 2.5104134 | Basophil/Mast 1 |
| 35 | STMN1 | 1.15E-174 | 1.4230665 | Basophil/Mast 1 |
| 36 | FOS | 3.99E-174 | 14.45543 | Basophil/Mast 1 |
| 37 | BATF | 1.88E-173 | 4.659305 | Basophil/Mast 1 |
| 38 | ALOX5AP | 4.70E-173 | -4.297357 | Basophil/Mast 1 |
| 39 | SOCS1 | 2.83E-166 | 2.954154 | Basophil/Mast 1 |
| 40 | CTNNBL1 | 1.06E-163 | 2.621963 | Basophil/Mast 1 |
| 41 | LOC284454 | 1.62E-159 | 3.10673 | Basophil/Mast 1 |
| 42 | CPM | 1.40E-155 | 2.689236 | Basophil/Mast 1 |
| 43 | ANXA1 | 4.42E-146 | 4.674618 | Basophil/Mast 1 |
| 44 | LAT | 1.17E-138 | 3.0285506 | Basophil/Mast 1 |
| 45 | RGS10 | 2.70E-136 | 1.7509279 | Basophil/Mast 1 |
| 46 | PTMA | 2.60E-134 | 8.709771 | Basophil/Mast 1 |
| 47 | CD44 | 7.28E-134 | 1.7971956 | Basophil/Mast 1 |
| 48 | CNRIP1 | 1.81E-130 | 3.698083 | Basophil/Mast 1 |
| 49 | SMYD3 | 4.64E-129 | 3.4250011 | Basophil/Mast 1 |
| 50 | GRAP2 | 3.27E-123 | 4.0318127 | Basophil/Mast 1 |
| 51 | BACE2 | 4.59E-123 | 2.6727295 | Basophil/Mast 1 |
| 52 | VIM | 7.05E-123 | -3.1112196 | Basophil/Mast 1 |
| 53 | LMO4 | 4.30E-122 | 1.9432669 | Basophil/Mast 1 |
| 54 | SLC45A3 | 4.65E-119 | 5.474376 | Basophil/Mast 1 |
| 55 | NTM | 1.25E-115 | 4.4961467 | Basophil/Mast 1 |
| 56 | RPL34 | 1.34E-107 | 9.55265 | Basophil/Mast 1 |
| 57 | SYTL3 | 4.78E-107 | 2.3132896 | Basophil/Mast 1 |
| 58 | SLC26A2 | 7.49E-107 | 2.654026 | Basophil/Mast 1 |
| 59 | H3F3B | 1.36E-105 | 6.1217318 | Basophil/Mast 1 |
| 60 | ARHGDIB | 2.45E-103 | 1.6543792 | Basophil/Mast 1 |
| 61 | C5orf30 | 3.73E-103 | 4.982727 | Basophil/Mast 1 |
| 62 | CTTNBP2 | 1.17E-100 | 6.428373 | Basophil/Mast 1 |
| 63 | SNHG8 | 2.76E-100 | 1.7290264 | Basophil/Mast 1 |
| 64 | FXYD5 | 6.91E-98 | 1.1908565 | Basophil/Mast 1 |
| 65 | FCER1G | 2.34E-97 | -14.242843 | Basophil/Mast 1 |
| 66 | FTH1 | 6.91E-97 | -206.99352 | Basophil/Mast 1 |
| 67 | SLC44A1 | 1.12E-96 | 2.285379 | Basophil/Mast 1 |
| 68 | AREG | 4.87E-94 | 9.1949005 | Basophil/Mast 1 |
| 69 | RENBP | 2.25E-89 | 1.3282561 | Basophil/Mast 1 |
| 70 | LAPTM5 | 6.88E-87 | -1.5331007 | Basophil/Mast 1 |
| 71 | PLIN2 | 5.96E-83 | -0.09537428 | Basophil/Mast 1 |
| 72 | TESPA1 | 2.90E-82 | 4.495335 | Basophil/Mast 1 |
| 73 | GLUL | 5.24E-82 | -1.6576588 | Basophil/Mast 1 |
| 74 | GPR65 | 1.02E-81 | 1.7980227 | Basophil/Mast 1 |
| 75 | LAT2 | 4.99E-79 | 1.5978258 | Basophil/Mast 1 |
| 76 | CD37 | 5.40E-79 | 0.09662132 | Basophil/Mast 1 |
| 77 | SMIM3 | 4.65E-76 | 2.7848442 | Basophil/Mast 1 |
| 78 | CRBN | 1.01E-75 | 1.6410342 | Basophil/Mast 1 |
| 79 | ACSL4 | 1.50E-75 | 1.9137655 | Basophil/Mast 1 |
| 80 | LXN | 1.54E-75 | 2.799129 | Basophil/Mast 1 |
| 81 | RPL36AL | 2.26E-74 | 1.6737154 | Basophil/Mast 1 |
| 82 | FOSB | 4.80E-74 | 1.2572027 | Basophil/Mast 1 |
| 83 | NDST2 | 8.77E-74 | 2.8079448 | Basophil/Mast 1 |
| 84 | EIF3E | 5.68E-73 | 1.2421503 | Basophil/Mast 1 |
| 85 | RPL14 | 9.50E-72 | 3.4439614 | Basophil/Mast 1 |
| 86 | BTK | 2.10E-71 | 1.6483326 | Basophil/Mast 1 |

| | | | | |
|---|---|---|---|---|
| 87 | TMEM176B | 1.10E-69 | 1.8412441 | Basophil/Mast 1 |
| 88 | PLGRKT | 2.28E-69 | 1.4984 | Basophil/Mast 1 |
| 89 | ADRB2 | 8.29E-67 | 1.7567353 | Basophil/Mast 1 |
| 90 | FOXP1 | 3.03E-66 | 1.4562823 | Basophil/Mast 1 |
| 91 | FAM212A | 4.48E-65 | 3.716917 | Basophil/Mast 1 |
| 92 | ALOX5 | 1.06E-61 | -1.0157325 | Basophil/Mast 1 |
| 93 | PHF20 | 2.00E-61 | 1.7889395 | Basophil/Mast 1 |
| 94 | RPL7 | 4.20E-60 | 2.4243157 | Basophil/Mast 1 |
| 95 | LYL1 | 4.98E-60 | 1.7448276 | Basophil/Mast 1 |
| 96 | ARHGEF6 | 1.30E-59 | 2.4219468 | Basophil/Mast 1 |
| 97 | KLRG1 | 4.84E-59 | 2.4311922 | Basophil/Mast 1 |
| 98 | CHN2 | 2.54E-58 | 4.0305204 | Basophil/Mast 1 |
| 99 | RPS4X | 9.32E-58 | 0.66348344 | Basophil/Mast 1 |
| 0 | TPSAB1 | 0 | 169.30643 | Basophil/Mast 2 |
| 1 | TPSB2 | 0 | 228.39511 | Basophil/Mast 2 |
| 2 | CPA3 | 0 | 21.140018 | Basophil/Mast 2 |
| 3 | MS4A2 | 0 | 13.946862 | Basophil/Mast 2 |
| 4 | HPGDS | 0 | 11.232315 | Basophil/Mast 2 |
| 5 | RGS2 | 1.19E-302 | 27.257483 | Basophil/Mast 2 |
| 6 | RGS1 | 1.23E-302 | 23.385254 | Basophil/Mast 2 |
| 7 | CD69 | 1.27E-292 | 21.592945 | Basophil/Mast 2 |
| 8 | RGS13 | 5.32E-292 | 10.594086 | Basophil/Mast 2 |
| 9 | VWA5A | 2.35E-286 | 8.356973 | Basophil/Mast 2 |
| 10 | NFKBIA | 9.32E-260 | 32.720486 | Basophil/Mast 2 |
| 11 | C1orf186 | 1.34E-251 | 7.9593544 | Basophil/Mast 2 |
| 12 | SLC18A2 | 2.38E-250 | 7.9500365 | Basophil/Mast 2 |
| 13 | LMNA | 6.04E-248 | 38.570774 | Basophil/Mast 2 |
| 14 | BIRC3 | 1.91E-247 | 13.740721 | Basophil/Mast 2 |
| 15 | ID2 | 2.49E-243 | 22.53932 | Basophil/Mast 2 |
| 16 | HDC | 2.79E-238 | 8.649108 | Basophil/Mast 2 |
| 17 | IL1RL1 | 4.96E-237 | 5.007257 | Basophil/Mast 2 |
| 18 | SAMSN1 | 9.65E-233 | 8.160961 | Basophil/Mast 2 |
| 19 | GPR65 | 1.44E-230 | 8.286903 | Basophil/Mast 2 |
| 20 | LTC4S | 3.99E-226 | 6.031026 | Basophil/Mast 2 |
| 21 | FCER1A | 2.23E-222 | 5.983845 | Basophil/Mast 2 |
| 22 | KIT | 2.32E-220 | 7.1412373 | Basophil/Mast 2 |
| 23 | CREM | 2.01E-217 | 8.928278 | Basophil/Mast 2 |
| 24 | SRGN | 1.23E-207 | 53.131897 | Basophil/Mast 2 |
| 25 | TNFAIP3 | 1.46E-203 | 6.0768886 | Basophil/Mast 2 |
| 26 | SELK | 1.07E-197 | 9.463466 | Basophil/Mast 2 |
| 27 | CLU | 7.61E-197 | 4.8048906 | Basophil/Mast 2 |
| 28 | CPM | 1.60E-195 | 6.2678275 | Basophil/Mast 2 |
| 29 | LOC101927482 | 9.77E-190 | 11.248655 | Basophil/Mast 2 |
| 30 | TNFRSF9 | 4.11E-186 | 9.001315 | Basophil/Mast 2 |
| 31 | MAOB | 1.81E-183 | 5.789486 | Basophil/Mast 2 |
| 32 | SIGLEC17P | 1.28E-180 | 6.6951447 | Basophil/Mast 2 |
| 33 | RHOH | 3.58E-174 | 5.3902297 | Basophil/Mast 2 |
| 34 | CSF1 | 2.74E-172 | 5.865577 | Basophil/Mast 2 |
| 35 | FOSB | 8.60E-168 | 5.8971033 | Basophil/Mast 2 |
| 36 | DDIT4 | 4.98E-167 | 6.6115766 | Basophil/Mast 2 |
| 37 | CKLF | 4.10E-166 | 4.7377543 | Basophil/Mast 2 |
| 38 | GADD45B | 6.15E-160 | 9.040466 | Basophil/Mast 2 |
| 39 | ANXA1 | 1.39E-157 | 29.261211 | Basophil/Mast 2 |
| 40 | ITM2C | 6.52E-156 | 3.6521556 | Basophil/Mast 2 |
| 41 | AREG | 4.57E-154 | 17.947039 | Basophil/Mast 2 |
| 42 | REL | 5.76E-148 | 4.1637764 | Basophil/Mast 2 |
| 43 | BATF | 1.36E-147 | 4.7026625 | Basophil/Mast 2 |
| 44 | SOCS1 | 2.10E-140 | 4.488644 | Basophil/Mast 2 |
| 45 | ELF1 | 2.28E-138 | 4.206376 | Basophil/Mast 2 |
| 46 | NR4A1 | 9.92E-138 | 4.0906496 | Basophil/Mast 2 |
| 47 | GATA2 | 1.16E-137 | 6.0269804 | Basophil/Mast 2 |
| 48 | LAPTM4A | 4.98E-134 | 6.393433 | Basophil/Mast 2 |
| 49 | SLC2A3 | 2.53E-133 | 4.1603823 | Basophil/Mast 2 |
| 50 | LMO4 | 3.18E-130 | 3.9962556 | Basophil/Mast 2 |

| | | | | |
|---|---|---|---|---|
| 51 | LEO1 | 4.55E-129 | 5.0340223 | Basophil/Mast 2 |
| 52 | CALB2 | 7.78E-128 | 6.710841 | Basophil/Mast 2 |
| 53 | KDM6B | 1.15E-124 | 3.912089 | Basophil/Mast 2 |
| 54 | BHLHE40 | 1.35E-123 | 3.5516207 | Basophil/Mast 2 |
| 55 | BRE-AS1 | 1.78E-122 | 4.9766645 | Basophil/Mast 2 |
| 56 | STK17B | 3.98E-120 | 3.9277084 | Basophil/Mast 2 |
| 57 | H3F3B | 2.49E-117 | 18.1631 | Basophil/Mast 2 |
| 58 | NR4A3 | 2.29E-115 | 5.575275 | Basophil/Mast 2 |
| 59 | SLC45A3 | 4.42E-114 | 6.5382977 | Basophil/Mast 2 |
| 60 | PTGS1 | 5.60E-114 | 5.0077295 | Basophil/Mast 2 |
| 61 | RPL34 | 5.83E-114 | 26.420498 | Basophil/Mast 2 |
| 62 | NFKBIZ | 1.12E-113 | 3.5265055 | Basophil/Mast 2 |
| 63 | HPGD | 1.98E-111 | 3.765852 | Basophil/Mast 2 |
| 64 | PTGS2 | 2.36E-109 | 5.830273 | Basophil/Mast 2 |
| 65 | GLUL | 5.55E-107 | 9.209553 | Basophil/Mast 2 |
| 66 | CCL4L1 | 4.84E-105 | 7.441105 | Basophil/Mast 2 |
| 67 | LOC284454 | 3.94E-104 | 3.7182465 | Basophil/Mast 2 |
| 68 | SDCBP | 3.18E-103 | 4.3950562 | Basophil/Mast 2 |
| 69 | CD83 | 1.24E-102 | 4.101395 | Basophil/Mast 2 |
| 70 | SYAP1 | 1.34E-100 | 4.6369033 | Basophil/Mast 2 |
| 71 | JUNB | 3.92E-100 | 9.054476 | Basophil/Mast 2 |
| 72 | RAB27B | 1.03E-97 | 5.230643 | Basophil/Mast 2 |
| 73 | PLIN2 | 2.54E-93 | 4.2773013 | Basophil/Mast 2 |
| 74 | ELL2 | 1.98E-91 | 3.2885573 | Basophil/Mast 2 |
| 75 | FTH1 | 5.43E-88 | -67.06834 | Basophil/Mast 2 |
| 76 | LAT | 4.39E-86 | 3.3872335 | Basophil/Mast 2 |
| 77 | CAPG | 4.77E-85 | -1.6114855 | Basophil/Mast 2 |
| 78 | DUSP10 | 6.16E-84 | 3.8576508 | Basophil/Mast 2 |
| 79 | RPS4X | 7.66E-84 | 12.98653 | Basophil/Mast 2 |
| 80 | CST7 | 1.58E-79 | 1.923647 | Basophil/Mast 2 |
| 81 | ACSL4 | 1.42E-76 | 2.910253 | Basophil/Mast 2 |
| 82 | SRSF5 | 6.10E-76 | 3.5169873 | Basophil/Mast 2 |
| 83 | PTMA | 5.60E-74 | 12.980499 | Basophil/Mast 2 |
| 84 | ARHGEF6 | 1.07E-73 | 3.573693 | Basophil/Mast 2 |
| 85 | STMN1 | 8.66E-73 | 1.2820988 | Basophil/Mast 2 |
| 86 | HIF1A | 3.33E-72 | 2.5418758 | Basophil/Mast 2 |
| 87 | BACE2 | 3.14E-70 | 2.887806 | Basophil/Mast 2 |
| 88 | MYADM | 4.79E-70 | 2.9391172 | Basophil/Mast 2 |
| 89 | VIM | 1.55E-69 | -0.43676656 | Basophil/Mast 2 |
| 90 | ARHGDIB | 8.03E-69 | 3.7592943 | Basophil/Mast 2 |
| 91 | NR4A2 | 9.55E-68 | 3.2434828 | Basophil/Mast 2 |
| 92 | GRAP2 | 5.07E-67 | 4.2281623 | Basophil/Mast 2 |
| 93 | MCL1 | 2.33E-65 | 3.4318223 | Basophil/Mast 2 |
| 94 | DDX3X | 2.62E-65 | 2.4417748 | Basophil/Mast 2 |
| 95 | SKIL | 2.04E-64 | 2.3251276 | Basophil/Mast 2 |
| 96 | FXYD5 | 2.41E-64 | 2.2401805 | Basophil/Mast 2 |
| 97 | MALAT1 | 3.29E-64 | 181.36827 | Basophil/Mast 2 |
| 98 | PRNP | 5.37E-64 | 2.3504133 | Basophil/Mast 2 |
| 99 | RPS27 | 9.35E-64 | 18.555067 | Basophil/Mast 2 |
| 0 | SPARCL1 | 6.07E-240 | 28.198532 | Bronchial Vessel 1 |
| 1 | GNG11 | 1.09E-203 | 13.692903 | Bronchial Vessel 1 |
| 2 | IGFBP7 | 6.92E-199 | 41.85479 | Bronchial Vessel 1 |
| 3 | SPRY1 | 4.70E-171 | 20.735798 | Bronchial Vessel 1 |
| 4 | TM4SF1 | 6.63E-171 | 26.402304 | Bronchial Vessel 1 |
| 5 | MGP | 1.93E-165 | 16.731281 | Bronchial Vessel 1 |
| 6 | CLEC14A | 1.47E-164 | 6.753722 | Bronchial Vessel 1 |
| 7 | A2M | 1.73E-164 | 8.307788 | Bronchial Vessel 1 |
| 8 | EMCN | 8.03E-164 | 5.7164607 | Bronchial Vessel 1 |
| 9 | VWF | 3.76E-161 | 7.6288047 | Bronchial Vessel 1 |
| 11 | IGFBP4 | 2.57E-159 | 8.652573 | Bronchial Vessel 1 |
| 10 | PLVAP | 2.57E-159 | 11.1895485 | Bronchial Vessel 1 |
| 12 | TCF4 | 2.31E-155 | 6.0188127 | Bronchial Vessel 1 |
| 13 | NPDC1 | 1.77E-153 | 6.493121 | Bronchial Vessel 1 |
| 14 | BCAM | 2.34E-141 | 5.7835712 | Bronchial Vessel 1 |

| | | | | |
|---|---|---|---|---|
| 15 | ADGRL4 | 2.82E-135 | 6.4441366 | Bronchial Vessel 1 |
| 16 | IFITM1 | 6.47E-134 | 12.997908 | Bronchial Vessel 1 |
| 17 | TSPAN7 | 1.15E-133 | 5.879541 | Bronchial Vessel 1 |
| 18 | LDB2 | 9.31E-133 | 4.933194 | Bronchial Vessel 1 |
| 19 | MARCKSL1 | 1.00E-132 | 5.4381065 | Bronchial Vessel 1 |
| 20 | SOCS3 | 7.34E-130 | 18.105167 | Bronchial Vessel 1 |
| 21 | ECSCR | 2.73E-127 | 5.205023 | Bronchial Vessel 1 |
| 22 | IFITM3 | 8.08E-126 | 16.389467 | Bronchial Vessel 1 |
| 23 | RAMP2 | 1.57E-123 | 6.542682 | Bronchial Vessel 1 |
| 24 | ITM2A | 2.72E-123 | 5.281264 | Bronchial Vessel 1 |
| 25 | PALMD | 1.62E-118 | 5.207905 | Bronchial Vessel 1 |
| 26 | PRSS23 | 1.97E-118 | 6.169856 | Bronchial Vessel 1 |
| 27 | CAV1 | 1.20E-117 | 9.692065 | Bronchial Vessel 1 |
| 28 | ACKR1 | 2.52E-116 | 39.713356 | Bronchial Vessel 1 |
| 29 | PTRF | 7.81E-112 | 4.513588 | Bronchial Vessel 1 |
| 30 | MYC | 1.07E-110 | 7.9409995 | Bronchial Vessel 1 |
| 31 | EMP1 | 1.69E-110 | 10.388541 | Bronchial Vessel 1 |
| 32 | CYYR1 | 4.94E-107 | 5.7593975 | Bronchial Vessel 1 |
| 33 | CRIP2 | 1.10E-106 | 5.459977 | Bronchial Vessel 1 |
| 34 | NNMT | 1.03E-103 | 7.773777 | Bronchial Vessel 1 |
| 35 | IFITM2 | 9.43E-103 | 7.5768714 | Bronchial Vessel 1 |
| 36 | S100A16 | 5.47E-102 | 4.0369062 | Bronchial Vessel 1 |
| 37 | JAM2 | 8.88E-101 | 4.6375456 | Bronchial Vessel 1 |
| 38 | ENG | 4.01E-100 | 5.1401415 | Bronchial Vessel 1 |
| 39 | CD93 | 2.27E-98 | 4.0251017 | Bronchial Vessel 1 |
| 40 | NOSTRIN | 1.22E-97 | 3.962833 | Bronchial Vessel 1 |
| 41 | HSPG2 | 6.03E-92 | 5.1390576 | Bronchial Vessel 1 |
| 42 | AQP1 | 1.69E-90 | 6.418563 | Bronchial Vessel 1 |
| 43 | LMCD1 | 2.20E-90 | 4.637232 | Bronchial Vessel 1 |
| 44 | NRN1 | 6.68E-90 | 5.2961054 | Bronchial Vessel 1 |
| 45 | ESAM | 1.84E-88 | 3.269075 | Bronchial Vessel 1 |
| 46 | RAMP3 | 2.70E-88 | 4.435531 | Bronchial Vessel 1 |
| 47 | IFI27 | 3.76E-88 | 8.233011 | Bronchial Vessel 1 |
| 48 | TM4SF18 | 4.96E-86 | 6.5903654 | Bronchial Vessel 1 |
| 49 | CNN3 | 4.86E-85 | 3.7529633 | Bronchial Vessel 1 |
| 50 | CTNNAL1 | 6.19E-83 | 4.297127 | Bronchial Vessel 1 |
| 51 | GIMAP7 | 4.30E-82 | 4.2782 | Bronchial Vessel 1 |
| 52 | CLDN5 | 5.95E-82 | 6.8484926 | Bronchial Vessel 1 |
| 53 | CCDC85B | 1.17E-81 | 3.7981746 | Bronchial Vessel 1 |
| 54 | PCAT19 | 1.49E-79 | 3.333025 | Bronchial Vessel 1 |
| 55 | ETS2 | 1.63E-79 | 4.407595 | Bronchial Vessel 1 |
| 56 | CXorf36 | 1.11E-78 | 5.551012 | Bronchial Vessel 1 |
| 57 | APOLD1 | 1.79E-77 | 6.9424324 | Bronchial Vessel 1 |
| 58 | SPARC | 1.92E-76 | 4.1478577 | Bronchial Vessel 1 |
| 59 | GSN | 3.88E-76 | 6.637815 | Bronchial Vessel 1 |
| 60 | CTGF | 1.91E-75 | 14.771257 | Bronchial Vessel 1 |
| 61 | NFIB | 6.16E-74 | 3.6711562 | Bronchial Vessel 1 |
| 62 | SNCG | 4.92E-73 | 5.3142943 | Bronchial Vessel 1 |
| 63 | VWA1 | 6.63E-73 | 4.5224137 | Bronchial Vessel 1 |
| 64 | VAMP5 | 4.63E-72 | 3.5063336 | Bronchial Vessel 1 |
| 65 | CD59 | 1.06E-71 | 7.5840926 | Bronchial Vessel 1 |
| 66 | APP | 1.42E-71 | 2.6298 | Bronchial Vessel 1 |
| 67 | CLEC3B | 1.47E-71 | 3.4290235 | Bronchial Vessel 1 |
| 68 | CD34 | 9.87E-71 | 4.8560066 | Bronchial Vessel 1 |
| 69 | IL33 | 3.45E-70 | 4.139695 | Bronchial Vessel 1 |
| 70 | ADIRF | 3.45E-70 | 11.645515 | Bronchial Vessel 1 |
| 71 | FKBP1A | 3.19E-68 | 8.443765 | Bronchial Vessel 1 |
| 72 | SDPR | 8.57E-68 | 3.8161838 | Bronchial Vessel 1 |
| 73 | MYCT1 | 2.64E-67 | 4.165432 | Bronchial Vessel 1 |
| 74 | PECAM1 | 1.25E-65 | 3.3853612 | Bronchial Vessel 1 |
| 75 | RDX | 7.35E-65 | 2.932778 | Bronchial Vessel 1 |
| 76 | LPAR6 | 8.86E-65 | 3.4628344 | Bronchial Vessel 1 |
| 77 | CAV2 | 1.68E-64 | 2.6047206 | Bronchial Vessel 1 |
| 78 | GJA1 | 5.78E-64 | 4.3222375 | Bronchial Vessel 1 |

| | | | | |
|---|---|---|---|---|
| 79 | PRCP | 5.97E-64 | 4.2001877 | Bronchial Vessel 1 |
| 80 | RNASE1 | 1.01E-63 | 2.3445687 | Bronchial Vessel 1 |
| 81 | CDKN1A | 5.77E-62 | 5.5577965 | Bronchial Vessel 1 |
| 82 | MEF2C | 9.46E-62 | 3.3252952 | Bronchial Vessel 1 |
| 83 | SOCS2 | 2.43E-61 | 3.4129405 | Bronchial Vessel 1 |
| 84 | CLU | 1.42E-60 | 17.347782 | Bronchial Vessel 1 |
| 85 | ZFP36 | 1.97E-60 | 22.88732 | Bronchial Vessel 1 |
| 86 | TSC22D1 | 2.00E-60 | 7.793817 | Bronchial Vessel 1 |
| 87 | CYR61 | 8.60E-60 | 7.755695 | Bronchial Vessel 1 |
| 88 | MMRN2 | 4.93E-59 | 3.6610918 | Bronchial Vessel 1 |
| 89 | ADAMTS1 | 6.07E-58 | 9.715512 | Bronchial Vessel 1 |
| 90 | EGR1 | 8.59E-58 | 4.5694084 | Bronchial Vessel 1 |
| 91 | TINAGL1 | 2.30E-57 | 2.3669913 | Bronchial Vessel 1 |
| 92 | PVRL2 | 2.43E-57 | 2.9848175 | Bronchial Vessel 1 |
| 93 | EGFL7 | 4.28E-57 | 2.7512355 | Bronchial Vessel 1 |
| 94 | LIFR | 6.16E-57 | 3.456022 | Bronchial Vessel 1 |
| 95 | POSTN | 6.70E-57 | 7.4382687 | Bronchial Vessel 1 |
| 96 | ERG | 1.03E-56 | 3.8097901 | Bronchial Vessel 1 |
| 97 | TMEM88 | 1.63E-56 | 4.211395 | Bronchial Vessel 1 |
| 98 | ADAMTS9 | 7.82E-56 | 4.8478894 | Bronchial Vessel 1 |
| 99 | IFI16 | 1.09E-55 | 3.12389 | Bronchial Vessel 1 |
| 0 | TM4SF1 | 1.05E-113 | 39.475132 | Bronchial Vessel 2 |
| 1 | EMP1 | 3.23E-92 | 12.888003 | Bronchial Vessel 2 |
| 2 | SPARCL1 | 1.11E-83 | 10.288969 | Bronchial Vessel 2 |
| 3 | SOCS3 | 8.52E-82 | 13.55356 | Bronchial Vessel 2 |
| 4 | CLEC14A | 6.54E-71 | 5.344246 | Bronchial Vessel 2 |
| 5 | IFITM3 | 3.34E-69 | 16.077744 | Bronchial Vessel 2 |
| 6 | GNG11 | 2.61E-63 | 5.353558 | Bronchial Vessel 2 |
| 7 | RAMP2 | 6.05E-62 | 5.295913 | Bronchial Vessel 2 |
| 8 | IFITM1 | 2.74E-58 | 10.744243 | Bronchial Vessel 2 |
| 9 | ESAM | 3.21E-54 | 3.5088544 | Bronchial Vessel 2 |
| 10 | CAV1 | 3.86E-49 | 7.0584154 | Bronchial Vessel 2 |
| 11 | PALMD | 2.18E-47 | 4.505625 | Bronchial Vessel 2 |
| 12 | ITM2A | 4.12E-46 | 3.5913079 | Bronchial Vessel 2 |
| 13 | A2M | 9.74E-46 | 3.7940836 | Bronchial Vessel 2 |
| 14 | LDB2 | 3.55E-44 | 3.3189735 | Bronchial Vessel 2 |
| 15 | CD36 | 8.04E-44 | 4.278908 | Bronchial Vessel 2 |
| 16 | MT2A | 4.48E-43 | 82.74148 | Bronchial Vessel 2 |
| 17 | IFI27 | 1.14E-42 | 6.9874444 | Bronchial Vessel 2 |
| 18 | IGFBP4 | 6.41E-42 | 2.8557746 | Bronchial Vessel 2 |
| 19 | SLC2A3 | 1.06E-41 | 4.183803 | Bronchial Vessel 2 |
| 20 | SERTAD1 | 1.75E-41 | 3.2118804 | Bronchial Vessel 2 |
| 21 | ARID5A | 6.74E-41 | 3.727473 | Bronchial Vessel 2 |
| 22 | GJA1 | 9.59E-39 | 4.284362 | Bronchial Vessel 2 |
| 23 | CDKN1A | 7.84E-38 | 3.6826978 | Bronchial Vessel 2 |
| 24 | VWF | 1.14E-37 | 2.71188 | Bronchial Vessel 2 |
| 25 | MT1M | 1.18E-37 | 13.085074 | Bronchial Vessel 2 |
| 26 | EMCN | 1.41E-37 | 3.0374143 | Bronchial Vessel 2 |
| 27 | EPAS1 | 2.80E-37 | 3.5348108 | Bronchial Vessel 2 |
| 28 | PRSS23 | 4.50E-37 | 2.8902514 | Bronchial Vessel 2 |
| 29 | CLEC3B | 5.31E-37 | 3.1391454 | Bronchial Vessel 2 |
| 30 | CRIP2 | 4.02E-36 | 2.7810564 | Bronchial Vessel 2 |
| 31 | IFITM2 | 9.72E-36 | 4.457902 | Bronchial Vessel 2 |
| 32 | TCF4 | 1.99E-35 | 3.305076 | Bronchial Vessel 2 |
| 33 | CD93 | 2.65E-35 | 2.8376467 | Bronchial Vessel 2 |
| 34 | ZFP36 | 5.07E-35 | 16.37718 | Bronchial Vessel 2 |
| 35 | S100A16 | 6.63E-35 | 2.2797918 | Bronchial Vessel 2 |
| 36 | SOX17 | 1.98E-34 | 5.39276 | Bronchial Vessel 2 |
| 37 | TSC22D1 | 2.14E-33 | 6.856778 | Bronchial Vessel 2 |
| 38 | CSRNP1 | 8.24E-33 | 2.8250494 | Bronchial Vessel 2 |
| 39 | THBD | 1.43E-31 | 2.8508642 | Bronchial Vessel 2 |
| 40 | STOM | 1.92E-31 | 3.3161964 | Bronchial Vessel 2 |
| 41 | MGP | 2.95E-31 | 10.12468 | Bronchial Vessel 2 |
| 42 | CLDN5 | 7.31E-31 | 5.072129 | Bronchial Vessel 2 |

| | | | | |
|---|---|---|---|---|
| 43 | BCAM | 2.58E-30 | 2.355664 | Bronchial Vessel 2 |
| 44 | FAM107A | 2.86E-30 | 3.285604 | Bronchial Vessel 2 |
| 45 | NEDD9 | 4.15E-30 | 3.1224144 | Bronchial Vessel 2 |
| 46 | PTRF | 2.28E-29 | 2.1267288 | Bronchial Vessel 2 |
| 47 | CAV2 | 3.00E-29 | 2.691436 | Bronchial Vessel 2 |
| 48 | LRRC32 | 4.71E-29 | 3.4973235 | Bronchial Vessel 2 |
| 49 | RNASE1 | 6.30E-29 | 1.3645585 | Bronchial Vessel 2 |
| 50 | PCDH17 | 1.16E-28 | 3.6014712 | Bronchial Vessel 2 |
| 51 | SLC25A25 | 1.84E-28 | 3.2327008 | Bronchial Vessel 2 |
| 52 | ENG | 2.52E-28 | 2.6417365 | Bronchial Vessel 2 |
| 53 | SDPR | 2.73E-28 | 4.2222424 | Bronchial Vessel 2 |
| 54 | SLC9A3R2 | 2.90E-28 | 2.8989437 | Bronchial Vessel 2 |
| 55 | APOLD1 | 6.57E-28 | 5.3277125 | Bronchial Vessel 2 |
| 56 | CTNNAL1 | 9.82E-28 | 3.106859 | Bronchial Vessel 2 |
| 57 | RASIP1 | 1.77E-27 | 2.492575 | Bronchial Vessel 2 |
| 58 | S1PR1 | 3.44E-27 | 3.0402982 | Bronchial Vessel 2 |
| 59 | ETS2 | 4.53E-27 | 2.5551558 | Bronchial Vessel 2 |
| 60 | C10orf10 | 2.03E-26 | 18.042025 | Bronchial Vessel 2 |
| 61 | NNMT | 2.46E-26 | 5.64081 | Bronchial Vessel 2 |
| 62 | ADGRL4 | 3.84E-26 | 3.077454 | Bronchial Vessel 2 |
| 63 | HYAL2 | 2.06E-25 | 2.7989144 | Bronchial Vessel 2 |
| 64 | ECSCR | 1.20E-24 | 2.177285 | Bronchial Vessel 2 |
| 65 | TSPAN7 | 2.74E-24 | 2.740148 | Bronchial Vessel 2 |
| 66 | JAM2 | 4.66E-24 | 3.003807 | Bronchial Vessel 2 |
| 67 | MT1X | 5.14E-24 | 18.37574 | Bronchial Vessel 2 |
| 68 | CYR61 | 9.58E-24 | 5.3286405 | Bronchial Vessel 2 |
| 69 | NPDC1 | 1.14E-23 | 2.1154923 | Bronchial Vessel 2 |
| 70 | F8 | 1.21E-23 | 3.5221531 | Bronchial Vessel 2 |
| 71 | SOCS2 | 1.53E-23 | 5.0982833 | Bronchial Vessel 2 |
| 72 | LPAR6 | 2.76E-23 | 2.9573 | Bronchial Vessel 2 |
| 73 | CYYR1 | 2.81E-23 | 3.7372859 | Bronchial Vessel 2 |
| 74 | MYCT1 | 3.15E-23 | 3.3559403 | Bronchial Vessel 2 |
| 75 | CDH5 | 4.14E-23 | 2.1585088 | Bronchial Vessel 2 |
| 76 | CD59 | 7.56E-23 | 4.0253778 | Bronchial Vessel 2 |
| 77 | FLT1 | 7.59E-23 | 2.6260571 | Bronchial Vessel 2 |
| 78 | PCAT19 | 9.62E-23 | 2.0646524 | Bronchial Vessel 2 |
| 79 | NFIB | 1.06E-22 | 2.4607043 | Bronchial Vessel 2 |
| 80 | APP | 2.45E-22 | 1.6840414 | Bronchial Vessel 2 |
| 81 | ABCB1 | 9.25E-22 | 5.817893 | Bronchial Vessel 2 |
| 82 | ADM | 2.01E-21 | 4.338523 | Bronchial Vessel 2 |
| 83 | LMCD1 | 3.63E-21 | 2.469683 | Bronchial Vessel 2 |
| 84 | JUNB | 4.39E-21 | 6.5884275 | Bronchial Vessel 2 |
| 85 | DLC1 | 6.72E-21 | 2.5747168 | Bronchial Vessel 2 |
| 86 | ARHGAP29 | 3.35E-20 | 1.7359804 | Bronchial Vessel 2 |
| 87 | CALD1 | 1.27E-19 | 1.0354357 | Bronchial Vessel 2 |
| 88 | CEBPD | 2.57E-19 | 5.463367 | Bronchial Vessel 2 |
| 89 | MARCKSL1 | 3.87E-19 | 2.2090476 | Bronchial Vessel 2 |
| 90 | CALCRL | 6.39E-19 | 1.8733262 | Bronchial Vessel 2 |
| 91 | MMRN2 | 6.81E-19 | 2.853912 | Bronchial Vessel 2 |
| 92 | EGFL7 | 7.50E-19 | 2.0506325 | Bronchial Vessel 2 |
| 93 | SGK1 | 1.42E-18 | 2.2622483 | Bronchial Vessel 2 |
| 94 | TM4SF18 | 2.08E-18 | 5.051832 | Bronchial Vessel 2 |
| 95 | CNN3 | 4.42E-18 | 1.297831 | Bronchial Vessel 2 |
| 96 | RAMP3 | 4.78E-18 | 1.4591203 | Bronchial Vessel 2 |
| 97 | SEC14L1 | 4.93E-18 | 2.3605804 | Bronchial Vessel 2 |
| 98 | TIE1 | 6.38E-18 | 2.3038888 | Bronchial Vessel 2 |
| 99 | IGFBP7 | 1.09E-17 | 3.2885704 | Bronchial Vessel 2 |
| 0 | CD2 | 0 | 6.535717 | CD4+ Memory/Effector T |
| 20 | CD52 | 0 | -5.754465 | CD4+ Memory/Effector T |
| 21 | PRDM1 | 0 | 3.472686 | CD4+ Memory/Effector T |
| 22 | TUBA4A | 0 | 2.9988234 | CD4+ Memory/Effector T |
| 23 | GZMA | 0 | 3.860815 | CD4+ Memory/Effector T |
| 25 | CD40LG | 0 | 6.625562 | CD4+ Memory/Effector T |
| 26 | IL2RG | 0 | 2.0778182 | CD4+ Memory/Effector T |

| | | | | |
|---|---|---|---|---|
| 19 | FYB | 0 | 2.6781945 | CD4+ Memory/Effector T |
| 27 | LAT | 0 | 3.3240755 | CD4+ Memory/Effector T |
| 29 | GPR171 | 0 | 4.730974 | CD4+ Memory/Effector T |
| 30 | TRAT1 | 0 | 5.011085 | CD4+ Memory/Effector T |
| 31 | IFITM1 | 0 | 2.543683 | CD4+ Memory/Effector T |
| 32 | RHOH | 0 | 3.0643065 | CD4+ Memory/Effector T |
| 33 | SPOCK2 | 0 | 2.8055663 | CD4+ Memory/Effector T |
| 34 | CD3G | 0 | 3.8255591 | CD4+ Memory/Effector T |
| 28 | CD6 | 0 | 4.568279 | CD4+ Memory/Effector T |
| 18 | CLEC2D | 0 | 3.6564734 | CD4+ Memory/Effector T |
| 24 | PTPRC | 0 | 2.1077073 | CD4+ Memory/Effector T |
| 16 | TSC22D3 | 0 | 6.6439986 | CD4+ Memory/Effector T |
| 1 | CD3D | 0 | 6.3748546 | CD4+ Memory/Effector T |
| 17 | CD48 | 0 | 2.7784526 | CD4+ Memory/Effector T |
| 3 | CXCR4 | 0 | 11.018285 | CD4+ Memory/Effector T |
| 4 | IL32 | 0 | 11.058491 | CD4+ Memory/Effector T |
| 5 | PTPRCAP | 0 | 4.8638687 | CD4+ Memory/Effector T |
| 6 | ZFP36L2 | 0 | 9.236929 | CD4+ Memory/Effector T |
| 7 | CORO1A | 0 | 4.8060083 | CD4+ Memory/Effector T |
| 8 | CD69 | 0 | 4.615263 | CD4+ Memory/Effector T |
| 2 | CD3E | 0 | 6.020976 | CD4+ Memory/Effector T |
| 10 | LCK | 0 | 4.2184296 | CD4+ Memory/Effector T |
| 9 | BTG1 | 0 | 8.866684 | CD4+ Memory/Effector T |
| 11 | RGS1 | 0 | 7.223717 | CD4+ Memory/Effector T |
| 12 | CCL5 | 0 | 6.4756737 | CD4+ Memory/Effector T |
| 13 | LTB | 0 | 6.163694 | CD4+ Memory/Effector T |
| 14 | ACAP1 | 0 | 3.7287238 | CD4+ Memory/Effector T |
| 15 | LIME1 | 0 | 3.8240492 | CD4+ Memory/Effector T |
| 35 | ICAM3 | 6.13E-298 | 2.248965 | CD4+ Memory/Effector T |
| 36 | PBXIP1 | 1.23E-289 | 2.506536 | CD4+ Memory/Effector T |
| 37 | PIK3IP1 | 8.94E-278 | 2.3234127 | CD4+ Memory/Effector T |
| 38 | KLRB1 | 1.56E-277 | 3.9252205 | CD4+ Memory/Effector T |
| 39 | 01-Sep | 2.79E-274 | 3.0488474 | CD4+ Memory/Effector T |
| 40 | IL7R | 1.54E-262 | 2.0418434 | CD4+ Memory/Effector T |
| 41 | BATF | 1.75E-260 | 2.7348347 | CD4+ Memory/Effector T |
| 42 | PLP2 | 6.58E-258 | 1.6510532 | CD4+ Memory/Effector T |
| 43 | CXCR3 | 2.47E-242 | 4.590926 | CD4+ Memory/Effector T |
| 44 | TRAF3IP3 | 5.30E-241 | 2.9117846 | CD4+ Memory/Effector T |
| 45 | ANXA6 | 1.32E-239 | 2.1085217 | CD4+ Memory/Effector T |
| 46 | GPR183 | 1.42E-237 | 2.6610918 | CD4+ Memory/Effector T |
| 47 | SIT1 | 2.53E-229 | 4.4621716 | CD4+ Memory/Effector T |
| 48 | CXCR6 | 1.01E-224 | 5.536156 | CD4+ Memory/Effector T |
| 49 | EMB | 2.50E-219 | 2.9556017 | CD4+ Memory/Effector T |
| 50 | BCL11B | 5.44E-203 | 3.8759515 | CD4+ Memory/Effector T |
| 51 | LIMD2 | 8.95E-196 | 1.9765868 | CD4+ Memory/Effector T |
| 52 | RPS29 | 3.78E-195 | 12.343711 | CD4+ Memory/Effector T |
| 53 | CYTIP | 3.64E-190 | 1.2664005 | CD4+ Memory/Effector T |
| 54 | ARHGAP15 | 6.96E-189 | 2.4532142 | CD4+ Memory/Effector T |
| 55 | LINC00892 | 1.52E-188 | 6.389757 | CD4+ Memory/Effector T |
| 56 | RPS27 | 3.94E-187 | 19.12656 | CD4+ Memory/Effector T |
| 57 | HCST | 4.19E-186 | 0.6130339 | CD4+ Memory/Effector T |
| 58 | CST7 | 2.17E-178 | 0.9319292 | CD4+ Memory/Effector T |
| 59 | EVL | 5.70E-178 | 0.59403986 | CD4+ Memory/Effector T |
| 60 | LSP1 | 2.59E-174 | 0.0493312 | CD4+ Memory/Effector T |
| 61 | APOBEC3G | 4.52E-173 | 2.3246865 | CD4+ Memory/Effector T |
| 62 | STK17A | 2.19E-172 | 1.7359058 | CD4+ Memory/Effector T |
| 63 | HMHA1 | 2.56E-163 | 2.2021148 | CD4+ Memory/Effector T |
| 64 | RASAL3 | 3.61E-160 | 2.6161067 | CD4+ Memory/Effector T |
| 65 | ARHGDIB | 1.02E-154 | 0.3530986 | CD4+ Memory/Effector T |
| 66 | SAMSN1 | 1.03E-154 | 0.850518 | CD4+ Memory/Effector T |
| 67 | STK17B | 7.00E-150 | 1.4526103 | CD4+ Memory/Effector T |
| 68 | S100A4 | 8.16E-147 | -19.786652 | CD4+ Memory/Effector T |
| 69 | CD96 | 2.04E-146 | 3.5894895 | CD4+ Memory/Effector T |
| 70 | LEPROTL1 | 5.91E-146 | 1.1486183 | CD4+ Memory/Effector T |

| | | | | |
|---|---|---|---|---|
| 71 | RAC2 | 9.59E-146 | 0.8566407 | CD4+ Memory/Effector T |
| 72 | RARRES3 | 1.62E-142 | 1.0833422 | CD4+ Memory/Effector T |
| 73 | SKAP1 | 9.67E-142 | 2.6177232 | CD4+ Memory/Effector T |
| 74 | AIM1 | 2.86E-141 | 2.0724463 | CD4+ Memory/Effector T |
| 75 | ID2 | 2.28E-135 | 2.3672976 | CD4+ Memory/Effector T |
| 76 | FKBP11 | 4.44E-135 | 1.7893687 | CD4+ Memory/Effector T |
| 77 | GZMM | 4.65E-134 | 1.9158791 | CD4+ Memory/Effector T |
| 78 | TAGAP | 1.21E-132 | 2.2555685 | CD4+ Memory/Effector T |
| 79 | XCL1 | 1.98E-132 | 4.449111 | CD4+ Memory/Effector T |
| 80 | PTGER4 | 6.32E-132 | 1.4211591 | CD4+ Memory/Effector T |
| 81 | ETS1 | 1.07E-131 | 1.5612148 | CD4+ Memory/Effector T |
| 82 | DDIT4 | 1.11E-129 | 1.6027342 | CD4+ Memory/Effector T |
| 83 | CD247 | 9.91E-129 | 1.2277846 | CD4+ Memory/Effector T |
| 84 | CD5 | 1.91E-128 | 4.518422 | CD4+ Memory/Effector T |
| 85 | RPL23A | 9.78E-127 | 5.92834 | CD4+ Memory/Effector T |
| 86 | CCDC167 | 4.39E-124 | 1.9387577 | CD4+ Memory/Effector T |
| 87 | FXYD5 | 3.29E-114 | 0.5838029 | CD4+ Memory/Effector T |
| 88 | TBC1D10C | 5.19E-114 | 1.1440196 | CD4+ Memory/Effector T |
| 89 | PCED1B-AS1 | 1.32E-111 | 1.6965654 | CD4+ Memory/Effector T |
| 90 | PRKCQ-AS1 | 2.42E-111 | 2.7074666 | CD4+ Memory/Effector T |
| 91 | EVI2A | 1.06E-109 | 1.157756 | CD4+ Memory/Effector T |
| 92 | ITK | 2.61E-109 | 3.6421227 | CD4+ Memory/Effector T |
| 93 | ALOX5AP | 1.84E-108 | -9.443063 | CD4+ Memory/Effector T |
| 94 | FLT3LG | 1.85E-106 | 2.9149654 | CD4+ Memory/Effector T |
| 95 | GIMAP7 | 3.41E-106 | 1.2037234 | CD4+ Memory/Effector T |
| 96 | CD7 | 4.13E-106 | 1.2025596 | CD4+ Memory/Effector T |
| 97 | SH3BGRL3 | 5.09E-106 | -6.8139033 | CD4+ Memory/Effector T |
| 98 | TNFSF8 | 5.77E-106 | 3.1443646 | CD4+ Memory/Effector T |
| 99 | CDC42SE2 | 1.55E-105 | 1.368882 | CD4+ Memory/Effector T |
| 0 | LTB | 0 | 10.192268 | CD4+ Naive T |
| 1 | RPS27 | 0 | 83.60393 | CD4+ Naive T |
| 2 | CD3E | 0 | 5.427428 | CD4+ Naive T |
| 3 | RPS29 | 0 | 51.21565 | CD4+ Naive T |
| 4 | CD3D | 0 | 4.644739 | CD4+ Naive T |
| 5 | RPS15A | 0 | 34.956593 | CD4+ Naive T |
| 6 | RPL34 | 0 | 44.4274 | CD4+ Naive T |
| 7 | PRKCQ-AS1 | 1.60E-305 | 6.0130844 | CD4+ Naive T |
| 8 | RPS6 | 5.26E-304 | 36.323658 | CD4+ Naive T |
| 9 | RPL13 | 8.96E-302 | 46.950962 | CD4+ Naive T |
| 10 | RPL17 | 9.48E-299 | 12.137712 | CD4+ Naive T |
| 11 | GAS5 | 9.06E-289 | 5.7112017 | CD4+ Naive T |
| 12 | RPL13AP5 | 7.96E-285 | 5.691679 | CD4+ Naive T |
| 13 | RPS12 | 3.25E-283 | 42.29301 | CD4+ Naive T |
| 14 | CCR7 | 1.32E-281 | 6.886894 | CD4+ Naive T |
| 15 | RPL3 | 9.38E-281 | 26.872187 | CD4+ Naive T |
| 16 | RPL32 | 1.43E-279 | 35.94462 | CD4+ Naive T |
| 17 | RPL13A | 4.88E-276 | 38.837696 | CD4+ Naive T |
| 18 | RPLP2 | 6.51E-275 | 35.498627 | CD4+ Naive T |
| 19 | CD48 | 1.81E-273 | 3.7981117 | CD4+ Naive T |
| 20 | RPL30 | 1.68E-272 | 23.882944 | CD4+ Naive T |
| 21 | RPS3 | 3.83E-271 | 25.272106 | CD4+ Naive T |
| 22 | RPS3A | 4.26E-269 | 30.569344 | CD4+ Naive T |
| 23 | RPS25 | 8.89E-267 | 23.045156 | CD4+ Naive T |
| 24 | RPL23A | 1.04E-265 | 24.575815 | CD4+ Naive T |
| 25 | RPL21 | 2.87E-265 | 43.02999 | CD4+ Naive T |
| 26 | RPL39 | 3.64E-265 | 31.48698 | CD4+ Naive T |
| 27 | PTPRCAP | 2.94E-260 | 3.4249713 | CD4+ Naive T |
| 28 | RPL35A | 3.80E-260 | 20.013731 | CD4+ Naive T |
| 29 | RPS18 | 8.35E-259 | 38.81281 | CD4+ Naive T |
| 30 | SELL | 4.65E-257 | 5.110444 | CD4+ Naive T |
| 31 | RPS27A | 4.32E-255 | 30.415924 | CD4+ Naive T |
| 32 | RPL31 | 4.61E-255 | 19.87144 | CD4+ Naive T |
| 33 | RPL36 | 7.61E-255 | 18.506546 | CD4+ Naive T |
| 34 | RPL36A | 2.31E-250 | 5.9146614 | CD4+ Naive T |

| | | | | |
|---|---|---|---|---|
| 35 | RPS10 | 6.94E-249 | 7.1241193 | CD4+ Naive T |
| 36 | RPL4 | 1.02E-243 | 10.589146 | CD4+ Naive T |
| 37 | PIK3IP1 | 6.74E-240 | 3.7872407 | CD4+ Naive T |
| 38 | RPL37 | 1.91E-234 | 18.914602 | CD4+ Naive T |
| 39 | RPS21 | 3.95E-233 | 12.7595005 | CD4+ Naive T |
| 40 | CORO1A | 1.96E-231 | 3.3645513 | CD4+ Naive T |
| 41 | RPSAP58 | 7.96E-231 | 3.8749774 | CD4+ Naive T |
| 42 | GLTSCR2 | 3.05E-230 | 4.756167 | CD4+ Naive T |
| 43 | EEF1G | 1.16E-226 | 12.342824 | CD4+ Naive T |
| 44 | RPSA | 1.27E-226 | 10.45718 | CD4+ Naive T |
| 45 | RPS28 | 1.48E-221 | 23.885336 | CD4+ Naive T |
| 46 | LIMD2 | 9.57E-216 | 3.427089 | CD4+ Naive T |
| 47 | RPL5 | 8.05E-213 | 10.850057 | CD4+ Naive T |
| 48 | BTG1 | 6.37E-211 | 5.725617 | CD4+ Naive T |
| 49 | RPS8 | 1.10E-208 | 18.587484 | CD4+ Naive T |
| 50 | RPL10A | 5.19E-207 | 12.852774 | CD4+ Naive T |
| 51 | RPS14 | 3.49E-205 | 26.475836 | CD4+ Naive T |
| 52 | RPL18A | 8.98E-203 | 20.838198 | CD4+ Naive T |
| 53 | RPS4X | 2.65E-200 | 19.802494 | CD4+ Naive T |
| 54 | RPL38 | 4.40E-200 | 11.862044 | CD4+ Naive T |
| 55 | IFITM1 | 8.54E-194 | 3.9204779 | CD4+ Naive T |
| 56 | RPS14P3 | 2.98E-191 | 3.462542 | CD4+ Naive T |
| 57 | RPL11 | 7.95E-191 | 22.19613 | CD4+ Naive T |
| 58 | LEF1 | 3.07E-190 | 6.3151703 | CD4+ Naive T |
| 59 | CD27 | 5.32E-189 | 4.3861666 | CD4+ Naive T |
| 60 | RPL10 | 1.09E-187 | 37.06955 | CD4+ Naive T |
| 61 | RPS17 | 3.16E-187 | 13.474543 | CD4+ Naive T |
| 62 | RPL7 | 4.26E-186 | 17.09459 | CD4+ Naive T |
| 63 | RPS23 | 1.33E-185 | 17.602905 | CD4+ Naive T |
| 64 | SNHG5 | 2.95E-185 | 4.258543 | CD4+ Naive T |
| 65 | RPL18 | 1.10E-182 | 10.662117 | CD4+ Naive T |
| 66 | IL32 | 4.96E-182 | 3.8599808 | CD4+ Naive T |
| 67 | GIMAP7 | 2.83E-181 | 2.8192117 | CD4+ Naive T |
| 68 | LCK | 1.73E-179 | 3.408683 | CD4+ Naive T |
| 69 | RPL27A | 3.02E-179 | 18.070837 | CD4+ Naive T |
| 70 | CXCR4 | 1.52E-177 | 3.361333 | CD4+ Naive T |
| 71 | PCED1B-AS1 | 3.56E-177 | 3.6624787 | CD4+ Naive T |
| 72 | RPL19 | 8.24E-176 | 17.94233 | CD4+ Naive T |
| 73 | CLEC2D | 1.32E-175 | 3.3998582 | CD4+ Naive T |
| 74 | RPL9 | 8.12E-175 | 16.405539 | CD4+ Naive T |
| 75 | EEF1B2 | 2.15E-173 | 5.6800632 | CD4+ Naive T |
| 76 | EEF2 | 6.55E-172 | 4.99379 | CD4+ Naive T |
| 77 | FYB | 4.48E-170 | 2.8421082 | CD4+ Naive T |
| 78 | RPS2 | 3.22E-169 | 21.343027 | CD4+ Naive T |
| 79 | RPL14 | 8.08E-167 | 11.006241 | CD4+ Naive T |
| 80 | RPL41 | 3.35E-159 | 33.85491 | CD4+ Naive T |
| 81 | RPS13 | 4.36E-159 | 14.580346 | CD4+ Naive T |
| 82 | CD2 | 3.39E-155 | 2.5931587 | CD4+ Naive T |
| 83 | RPS7 | 6.62E-155 | 8.961151 | CD4+ Naive T |
| 84 | TRAF3IP3 | 1.21E-154 | 3.4333067 | CD4+ Naive T |
| 85 | ACAP1 | 4.69E-154 | 3.2644334 | CD4+ Naive T |
| 86 | RPS5 | 1.34E-152 | 8.879581 | CD4+ Naive T |
| 87 | LDHB | 1.00E-151 | 2.4294624 | CD4+ Naive T |
| 88 | LIME1 | 3.04E-151 | 3.0719175 | CD4+ Naive T |
| 89 | EEF1A1 | 2.05E-149 | 39.635994 | CD4+ Naive T |
| 90 | RPL22 | 1.25E-148 | 7.1415224 | CD4+ Naive T |
| 91 | TCF7 | 1.25E-145 | 4.061445 | CD4+ Naive T |
| 92 | EIF3E | 1.96E-145 | 3.1070464 | CD4+ Naive T |
| 93 | CD7 | 1.43E-144 | 2.393543 | CD4+ Naive T |
| 94 | RPL6 | 2.29E-144 | 9.939549 | CD4+ Naive T |
| 95 | RPS15 | 1.41E-143 | 13.06942 | CD4+ Naive T |
| 96 | RPL35 | 5.62E-141 | 9.134025 | CD4+ Naive T |
| 97 | NPM1 | 2.19E-140 | 3.6697128 | CD4+ Naive T |
| 98 | RPL26 | 2.96E-140 | 14.185569 | CD4+ Naive T |

| | | | | |
|---|---|---|---|---|
| 99 | RPS16 | 2.95E-134 | 10.108024 | CD4+ Naive T |
| 0 | CCL5 | 0 | 14.242965 | CD8+ Memory/Effector T |
| 1 | CD3D | 0 | 4.821581 | CD8+ Memory/Effector T |
| 2 | GZMK | 0 | 8.984069 | CD8+ Memory/Effector T |
| 3 | NKG7 | 0 | 7.9363256 | CD8+ Memory/Effector T |
| 4 | CD3E | 0 | 4.7184076 | CD8+ Memory/Effector T |
| 5 | PTPRCAP | 0 | 4.6822853 | CD8+ Memory/Effector T |
| 6 | CXCR4 | 0 | 8.10624 | CD8+ Memory/Effector T |
| 7 | CST7 | 0 | 4.6371074 | CD8+ Memory/Effector T |
| 8 | GZMA | 0 | 5.008802 | CD8+ Memory/Effector T |
| 9 | IL32 | 0 | 6.725637 | CD8+ Memory/Effector T |
| 10 | DUSP2 | 0 | 5.05165 | CD8+ Memory/Effector T |
| 11 | ZFP36L2 | 2.46E-301 | 6.321497 | CD8+ Memory/Effector T |
| 12 | CD2 | 3.34E-273 | 3.5015275 | CD8+ Memory/Effector T |
| 13 | CTSW | 2.89E-270 | 3.057408 | CD8+ Memory/Effector T |
| 14 | PRF1 | 1.32E-239 | 3.0714571 | CD8+ Memory/Effector T |
| 15 | CORO1A | 3.55E-233 | 2.7381663 | CD8+ Memory/Effector T |
| 16 | GZMM | 1.05E-229 | 3.4190917 | CD8+ Memory/Effector T |
| 17 | CD7 | 6.19E-218 | 3.19855 | CD8+ Memory/Effector T |
| 18 | RPS27 | 2.42E-216 | 29.28967 | CD8+ Memory/Effector T |
| 19 | CD8A | 1.12E-213 | 5.2861047 | CD8+ Memory/Effector T |
| 20 | CD48 | 2.97E-205 | 2.6420987 | CD8+ Memory/Effector T |
| 21 | BTG1 | 1.02E-202 | 4.897006 | CD8+ Memory/Effector T |
| 22 | FYN | 1.01E-200 | 3.0844727 | CD8+ Memory/Effector T |
| 23 | LCK | 5.91E-197 | 3.3077385 | CD8+ Memory/Effector T |
| 24 | CLEC2D | 2.80E-185 | 3.2848032 | CD8+ Memory/Effector T |
| 25 | RPS29 | 3.84E-175 | 15.915943 | CD8+ Memory/Effector T |
| 26 | KLRG1 | 4.23E-159 | 4.0432086 | CD8+ Memory/Effector T |
| 27 | CD69 | 4.06E-156 | 2.3797362 | CD8+ Memory/Effector T |
| 28 | LIME1 | 3.47E-153 | 2.8187542 | CD8+ Memory/Effector T |
| 29 | PTPRC | 2.30E-141 | 1.9704188 | CD8+ Memory/Effector T |
| 30 | HCST | 1.91E-136 | 1.4546393 | CD8+ Memory/Effector T |
| 31 | IFITM1 | 1.59E-128 | 1.7751117 | CD8+ Memory/Effector T |
| 32 | CD8B | 5.18E-123 | 4.5904694 | CD8+ Memory/Effector T |
| 33 | GZMH | 8.36E-123 | 3.223404 | CD8+ Memory/Effector T |
| 34 | RPL17 | 1.08E-122 | 4.4101505 | CD8+ Memory/Effector T |
| 35 | RPL23A | 3.22E-122 | 8.779228 | CD8+ Memory/Effector T |
| 36 | ACAP1 | 1.25E-121 | 2.5308805 | CD8+ Memory/Effector T |
| 37 | KLRB1 | 1.02E-119 | 6.374667 | CD8+ Memory/Effector T |
| 38 | LIMD2 | 4.51E-112 | 2.0453198 | CD8+ Memory/Effector T |
| 39 | PIK3R1 | 9.75E-112 | 2.6589 | CD8+ Memory/Effector T |
| 40 | SYTL3 | 5.01E-108 | 2.6011117 | CD8+ Memory/Effector T |
| 41 | ICAM3 | 1.92E-107 | 1.9302182 | CD8+ Memory/Effector T |
| 42 | SPOCK2 | 2.07E-105 | 2.5080829 | CD8+ Memory/Effector T |
| 43 | RPS15A | 6.12E-105 | 8.386616 | CD8+ Memory/Effector T |
| 44 | TARP | 9.33E-105 | 2.603628 | CD8+ Memory/Effector T |
| 45 | CD27 | 2.94E-103 | 3.3308003 | CD8+ Memory/Effector T |
| 46 | SH2D1A | 9.53E-103 | 3.6805384 | CD8+ Memory/Effector T |
| 47 | RPL13AP5 | 1.07E-101 | 1.9361608 | CD8+ Memory/Effector T |
| 48 | RPS3 | 3.60E-96 | 6.5806956 | CD8+ Memory/Effector T |
| 49 | PVRIG | 4.11E-93 | 3.0939438 | CD8+ Memory/Effector T |
| 50 | LTB | 7.60E-93 | 2.6872547 | CD8+ Memory/Effector T |
| 51 | CD247 | 7.00E-91 | 1.6066415 | CD8+ Memory/Effector T |
| 52 | CD3G | 7.30E-89 | 3.0405834 | CD8+ Memory/Effector T |
| 53 | ISG20 | 1.70E-87 | 1.3354572 | CD8+ Memory/Effector T |
| 54 | RPL36A | 3.33E-84 | 1.9100622 | CD8+ Memory/Effector T |
| 55 | PRDM1 | 5.07E-84 | 2.3917773 | CD8+ Memory/Effector T |
| 56 | RUNX3 | 1.15E-81 | 2.605899 | CD8+ Memory/Effector T |
| 57 | RPL13A | 8.54E-80 | 7.121246 | CD8+ Memory/Effector T |
| 58 | GPR171 | 1.42E-78 | 3.223306 | CD8+ Memory/Effector T |
| 59 | ETS1 | 1.67E-74 | 1.7561073 | CD8+ Memory/Effector T |
| 60 | GZMB | 4.30E-74 | 2.1680048 | CD8+ Memory/Effector T |
| 61 | IL7R | 6.55E-74 | 2.7163358 | CD8+ Memory/Effector T |
| 62 | ARL4C | 5.53E-72 | 2.1936922 | CD8+ Memory/Effector T |

| | | | | |
|---|---|---|---|---|
| 63 | SH2D2A | 5.07E-71 | 2.5776846 | CD8+ Memory/Effector T |
| 64 | LYAR | 2.70E-66 | 0.6890147 | CD8+ Memory/Effector T |
| 65 | RPL31 | 5.97E-66 | 3.7370806 | CD8+ Memory/Effector T |
| 66 | IL2RG | 7.61E-65 | 0.9264815 | CD8+ Memory/Effector T |
| 67 | GIMAP7 | 2.40E-64 | 1.1988122 | CD8+ Memory/Effector T |
| 68 | SAMD3 | 1.28E-63 | 2.944682 | CD8+ Memory/Effector T |
| 69 | RPL3 | 2.15E-63 | 4.2747827 | CD8+ Memory/Effector T |
| 70 | RPLP2 | 2.45E-61 | 3.8528607 | CD8+ Memory/Effector T |
| 71 | RPL13 | 5.95E-60 | 6.2199106 | CD8+ Memory/Effector T |
| 72 | LAT | 2.63E-59 | 2.041167 | CD8+ Memory/Effector T |
| 73 | RPS21 | 3.05E-59 | 1.9413283 | CD8+ Memory/Effector T |
| 74 | AKNA | 5.55E-59 | 1.8320106 | CD8+ Memory/Effector T |
| 75 | EMB | 7.18E-59 | 2.171134 | CD8+ Memory/Effector T |
| 76 | KLRD1 | 1.93E-58 | 2.033178 | CD8+ Memory/Effector T |
| 77 | HMHA1 | 8.15E-58 | 1.8627523 | CD8+ Memory/Effector T |
| 78 | BCL11B | 1.91E-57 | 2.9750571 | CD8+ Memory/Effector T |
| 79 | SLC38A1 | 2.57E-57 | 2.168113 | CD8+ Memory/Effector T |
| 80 | RPSA | 2.57E-56 | 1.533208 | CD8+ Memory/Effector T |
| 81 | RPS12 | 3.05E-56 | 4.645649 | CD8+ Memory/Effector T |
| 82 | RARRES3 | 2.43E-55 | 0.7448398 | CD8+ Memory/Effector T |
| 83 | TUBA4A | 5.10E-55 | 1.69921 | CD8+ Memory/Effector T |
| 84 | BIN2 | 7.32E-55 | 1.3776643 | CD8+ Memory/Effector T |
| 85 | TRAT1 | 1.75E-54 | 3.1223297 | CD8+ Memory/Effector T |
| 86 | CDC42SE2 | 3.35E-54 | 1.4062614 | CD8+ Memory/Effector T |
| 87 | LAG3 | 1.01E-53 | 3.8732593 | CD8+ Memory/Effector T |
| 88 | PCED1B-AS1 | 1.62E-53 | 1.7078996 | CD8+ Memory/Effector T |
| 89 | TRAF3IP3 | 2.09E-53 | 2.0293405 | CD8+ Memory/Effector T |
| 90 | MATK | 5.95E-53 | 2.1407802 | CD8+ Memory/Effector T |
| 91 | CMC1 | 1.05E-52 | 2.7791195 | CD8+ Memory/Effector T |
| 92 | 01-Sep | 2.39E-52 | 1.974978 | CD8+ Memory/Effector T |
| 93 | RPL27A | 2.99E-52 | 2.813069 | CD8+ Memory/Effector T |
| 94 | CD6 | 8.95E-48 | 2.5965223 | CD8+ Memory/Effector T |
| 95 | IKZF3 | 2.50E-46 | 2.783726 | CD8+ Memory/Effector T |
| 96 | PBXIP1 | 2.97E-46 | 1.6237631 | CD8+ Memory/Effector T |
| 97 | CD52 | 4.07E-46 | -13.513614 | CD8+ Memory/Effector T |
| 98 | PARP8 | 1.78E-45 | 1.9996618 | CD8+ Memory/Effector T |
| 99 | RPSAP58 | 2.00E-45 | 1.2782978 | CD8+ Memory/Effector T |
| 0 | CCL5 | 0 | 19.295109 | CD8+ Naive T |
| 21 | CD2 | 0 | 3.641736 | CD8+ Naive T |
| 20 | LCK | 0 | 3.445488 | CD8+ Naive T |
| 19 | KLRD1 | 0 | 3.6103325 | CD8+ Naive T |
| 18 | CD7 | 0 | 3.3202252 | CD8+ Naive T |
| 17 | IFITM1 | 0 | 3.6606154 | CD8+ Naive T |
| 16 | GZMM | 0 | 3.8768585 | CD8+ Naive T |
| 15 | GNLY | 0 | 14.33411 | CD8+ Naive T |
| 14 | CD8A | 0 | 5.661076 | CD8+ Naive T |
| 13 | CORO1A | 0 | 3.2957766 | CD8+ Naive T |
| 12 | CXCR4 | 0 | 4.910725 | CD8+ Naive T |
| 11 | PRF1 | 0 | 3.6952026 | CD8+ Naive T |
| 10 | GZMA | 0 | 5.0439453 | CD8+ Naive T |
| 9 | CTSW | 0 | 4.4295883 | CD8+ Naive T |
| 8 | GZMB | 0 | 7.0459833 | CD8+ Naive T |
| 7 | IL32 | 0 | 7.26838 | CD8+ Naive T |
| 6 | CD3E | 0 | 4.7313037 | CD8+ Naive T |
| 5 | CST7 | 0 | 5.939869 | CD8+ Naive T |
| 4 | GZMH | 0 | 7.899955 | CD8+ Naive T |
| 3 | PTPRCAP | 0 | 5.007629 | CD8+ Naive T |
| 2 | NKG7 | 0 | 16.3351 | CD8+ Naive T |
| 1 | CD3D | 0 | 6.270774 | CD8+ Naive T |
| 22 | CD8B | 0 | 5.43251 | CD8+ Naive T |
| 23 | ZFP36L2 | 0 | 4.4282126 | CD8+ Naive T |
| 24 | HCST | 1.65E-302 | 2.4535224 | CD8+ Naive T |
| 25 | LIME1 | 1.09E-262 | 2.9368284 | CD8+ Naive T |
| 26 | KLRG1 | 1.68E-261 | 4.330099 | CD8+ Naive T |

| | | | | |
|---|---|---|---|---|
| 27 | CD3G | 1.73E-261 | 4.055619 | CD8+ Naive T |
| 28 | CD48 | 5.32E-245 | 2.077028 | CD8+ Naive T |
| 29 | CLEC2D | 6.13E-241 | 3.0541265 | CD8+ Naive T |
| 30 | DUSP2 | 1.33E-222 | 2.9825103 | CD8+ Naive T |
| 31 | PTPRC | 5.86E-210 | 1.7390293 | CD8+ Naive T |
| 32 | ACAP1 | 2.22E-203 | 2.6606808 | CD8+ Naive T |
| 33 | CD247 | 1.30E-199 | 1.8813869 | CD8+ Naive T |
| 34 | RPS27 | 3.06E-198 | 15.968521 | CD8+ Naive T |
| 35 | C12orf75 | 8.93E-189 | 1.5200821 | CD8+ Naive T |
| 36 | CMC1 | 4.30E-185 | 3.6515772 | CD8+ Naive T |
| 37 | TARP | 6.53E-184 | 2.2666116 | CD8+ Naive T |
| 38 | KLRB1 | 3.92E-183 | 3.2271564 | CD8+ Naive T |
| 39 | LIMD2 | 2.08E-167 | 1.9430308 | CD8+ Naive T |
| 40 | FGFBP2 | 1.64E-165 | 2.7548807 | CD8+ Naive T |
| 41 | BTG1 | 1.57E-164 | 2.8561428 | CD8+ Naive T |
| 42 | ICAM3 | 9.15E-161 | 1.8922801 | CD8+ Naive T |
| 43 | FYN | 4.11E-160 | 2.1155803 | CD8+ Naive T |
| 44 | APOBEC3G | 1.34E-159 | 2.2975812 | CD8+ Naive T |
| 45 | PRDM1 | 3.13E-157 | 2.5636775 | CD8+ Naive T |
| 46 | RPS29 | 7.27E-149 | 8.007672 | CD8+ Naive T |
| 47 | KLRC2 | 5.50E-145 | 4.9574914 | CD8+ Naive T |
| 48 | RUNX3 | 1.26E-138 | 2.8787928 | CD8+ Naive T |
| 49 | KLRF1 | 2.08E-138 | 2.6535635 | CD8+ Naive T |
| 50 | IL2RG | 2.83E-135 | 1.2314312 | CD8+ Naive T |
| 51 | ISG20 | 7.84E-135 | 1.2291675 | CD8+ Naive T |
| 52 | CD69 | 8.73E-124 | 1.6157451 | CD8+ Naive T |
| 53 | ANXA6 | 3.33E-123 | 1.6677773 | CD8+ Naive T |
| 54 | MATK | 2.48E-120 | 2.534104 | CD8+ Naive T |
| 55 | BIN2 | 1.12E-118 | 1.5875916 | CD8+ Naive T |
| 56 | SYTL3 | 7.99E-118 | 2.2060838 | CD8+ Naive T |
| 57 | PVRIG | 3.44E-117 | 2.961557 | CD8+ Naive T |
| 58 | ETS1 | 2.02E-116 | 1.6855588 | CD8+ Naive T |
| 59 | 01-Sep | 1.83E-115 | 2.2681596 | CD8+ Naive T |
| 60 | SAMD3 | 6.24E-115 | 3.1583455 | CD8+ Naive T |
| 61 | RPS3 | 4.84E-113 | 3.82489 | CD8+ Naive T |
| 62 | CD52 | 4.26E-112 | -12.2014 | CD8+ Naive T |
| 63 | RARRES3 | 7.79E-112 | 0.9149694 | CD8+ Naive T |
| 64 | LAT | 4.76E-111 | 2.22556 | CD8+ Naive T |
| 65 | AKNA | 6.28E-111 | 1.9410874 | CD8+ Naive T |
| 66 | ZAP70 | 5.47E-108 | 2.7966673 | CD8+ Naive T |
| 67 | ITGB2 | 3.60E-106 | 0.52356756 | CD8+ Naive T |
| 68 | SH2D2A | 6.29E-105 | 2.6206372 | CD8+ Naive T |
| 69 | LSP1 | 3.65E-104 | -0.4024021 | CD8+ Naive T |
| 70 | TIGIT | 1.08E-103 | 3.935634 | CD8+ Naive T |
| 71 | RAC2 | 1.90E-102 | 0.7642864 | CD8+ Naive T |
| 72 | RPL23A | 2.56E-102 | 4.0130005 | CD8+ Naive T |
| 73 | SH2D1A | 5.68E-98 | 3.1128883 | CD8+ Naive T |
| 74 | TUBA4A | 1.43E-96 | 1.9566809 | CD8+ Naive T |
| 75 | GNG2 | 9.00E-96 | 2.3786721 | CD8+ Naive T |
| 76 | ARL4C | 1.68E-90 | 2.0006714 | CD8+ Naive T |
| 77 | SPOCK2 | 4.02E-90 | 1.852299 | CD8+ Naive T |
| 78 | RPL17 | 3.43E-87 | 1.5535132 | CD8+ Naive T |
| 79 | ID2 | 1.10E-86 | 1.1015865 | CD8+ Naive T |
| 80 | TRAF3IP3 | 2.96E-85 | 1.9752017 | CD8+ Naive T |
| 81 | S1PR5 | 5.50E-82 | 2.691515 | CD8+ Naive T |
| 82 | PCED1B-AS1 | 1.88E-80 | 1.5228583 | CD8+ Naive T |
| 83 | EMB | 2.12E-80 | 2.2056873 | CD8+ Naive T |
| 84 | RASAL3 | 6.72E-79 | 2.1782093 | CD8+ Naive T |
| 85 | TBC1D10C | 1.93E-76 | 1.0155315 | CD8+ Naive T |
| 86 | FCRL6 | 8.87E-76 | 3.0081053 | CD8+ Naive T |
| 87 | GIMAP7 | 1.03E-74 | 0.87073237 | CD8+ Naive T |
| 88 | STK17A | 3.48E-73 | 1.2171408 | CD8+ Naive T |
| 89 | RPL13AP5 | 2.08E-70 | 0.8787968 | CD8+ Naive T |
| 90 | RPS15A | 1.06E-69 | 2.6179285 | CD8+ Naive T |

| | | | | |
|---|---|---|---|---|
| 91 | IKZF3 | 2.06E-68 | 2.8162766 | CD8+ Naive T |
| 92 | TBX21 | 8.89E-68 | 2.9262755 | CD8+ Naive T |
| 93 | GZMK | 1.90E-66 | 3.4758644 | CD8+ Naive T |
| 94 | CYTIP | 3.48E-66 | 0.65808356 | CD8+ Naive T |
| 95 | SKAP1 | 3.88E-66 | 2.0835238 | CD8+ Naive T |
| 96 | LAG3 | 1.10E-65 | 3.7261026 | CD8+ Naive T |
| 97 | CDC42SE2 | 2.97E-65 | 1.2055596 | CD8+ Naive T |
| 98 | PYHIN1 | 3.95E-65 | 2.8106046 | CD8+ Naive T |
| 99 | DDIT4 | 7.47E-65 | 1.0121351 | CD8+ Naive T |
| 0 | FCN3 | 0 | 21.319874 | Capillary |
| 72 | DUSP6 | 0 | 2.0696316 | Capillary |
| 71 | FENDRR | 0 | 2.8301492 | Capillary |
| 70 | ID3 | 0 | 2.2953079 | Capillary |
| 69 | CX3CL1 | 0 | 3.2336662 | Capillary |
| 68 | ADRB1 | 0 | 3.363555 | Capillary |
| 67 | MT2A | 0 | 15.656328 | Capillary |
| 66 | MT1X | 0 | 9.759165 | Capillary |
| 65 | STXBP6 | 0 | 2.3674512 | Capillary |
| 64 | RASIP1 | 0 | 2.6783874 | Capillary |
| 63 | TSPAN7 | 0 | 2.613809 | Capillary |
| 62 | PTRF | 0 | 1.7988076 | Capillary |
| 61 | LIFR | 0 | 2.83031 | Capillary |
| 60 | IL7R | 0 | 1.5363816 | Capillary |
| 59 | VIPR1 | 0 | 2.9964821 | Capillary |
| 58 | TPM1 | 0 | 1.9976989 | Capillary |
| 57 | ADGRF5 | 0 | 2.4830112 | Capillary |
| 56 | HLA-E | 0 | 5.8635135 | Capillary |
| 55 | NPDC1 | 0 | 1.9609088 | Capillary |
| 54 | PECAM1 | 0 | 2.2141569 | Capillary |
| 53 | CNN3 | 0 | 2.307853 | Capillary |
| 52 | CAV2 | 0 | 1.9873048 | Capillary |
| 73 | ZBTB16 | 0 | 2.255977 | Capillary |
| 51 | ACVRL1 | 0 | 2.287657 | Capillary |
| 74 | A2M | 0 | 0.8261212 | Capillary |
| 76 | CRIP2 | 0 | 1.3821247 | Capillary |
| 97 | THBD | 0 | 1.2004983 | Capillary |
| 96 | NFIB | 0 | 1.8709095 | Capillary |
| 95 | BMPR2 | 0 | 2.031493 | Capillary |
| 94 | GPR146 | 0 | 3.092741 | Capillary |
| 93 | FOXF1 | 0 | 2.4210083 | Capillary |
| 92 | ROBO4 | 0 | 2.3643441 | Capillary |
| 91 | IFITM2 | 0 | 2.2391264 | Capillary |
| 90 | BST2 | 0 | 1.6636509 | Capillary |
| 89 | SLCO4A1 | 0 | 2.8166003 | Capillary |
| 88 | F2RL3 | 0 | 2.8808594 | Capillary |
| 87 | S1PR1 | 0 | 2.3452163 | Capillary |
| 86 | GIMAP7 | 0 | 1.8935357 | Capillary |
| 85 | JAM2 | 0 | 2.5392644 | Capillary |
| 84 | ECSCR | 0 | 1.5698841 | Capillary |
| 83 | TFPI | 0 | 1.3316061 | Capillary |
| 82 | ITM2A | 0 | 1.6014864 | Capillary |
| 81 | SEC14L1 | 0 | 2.0769174 | Capillary |
| 80 | VAMP5 | 0 | 1.7025039 | Capillary |
| 79 | GPX3 | 0 | 1.9389281 | Capillary |
| 78 | TMEM2 | 0 | 2.4047556 | Capillary |
| 77 | SOX7 | 0 | 3.1428096 | Capillary |
| 75 | TCF4 | 0 | 1.9858581 | Capillary |
| 50 | AKAP12 | 0 | 5.2800293 | Capillary |
| 49 | ID1 | 0 | 3.1095803 | Capillary |
| 48 | MT1E | 0 | 5.2808595 | Capillary |
| 21 | IFITM3 | 0 | 9.063852 | Capillary |
| 20 | IFI27 | 0 | 6.305844 | Capillary |
| 19 | NOSTRIN | 0 | 4.590256 | Capillary |
| 18 | FAM107A | 0 | 4.000287 | Capillary |

| | | | | |
|---|---|---|---|---|
| 17 | CALCRL | 0 | 4.122563 | Capillary |
| 16 | CAV1 | 0 | 4.890641 | Capillary |
| 15 | GNG11 | 0 | 3.4761078 | Capillary |
| 14 | C10orf10 | 0 | 6.541989 | Capillary |
| 13 | CLEC14A | 0 | 4.5373044 | Capillary |
| 12 | TIMP3 | 0 | 8.219875 | Capillary |
| 11 | SDPR | 0 | 6.773136 | Capillary |
| 10 | CA4 | 0 | 7.2074666 | Capillary |
| 9 | HYAL2 | 0 | 6.167332 | Capillary |
| 8 | RAMP3 | 0 | 5.159644 | Capillary |
| 7 | EGFL7 | 0 | 5.597739 | Capillary |
| 6 | RAMP2 | 0 | 7.115616 | Capillary |
| 5 | CLEC3B | 0 | 6.69537 | Capillary |
| 4 | EDN1 | 0 | 10.805174 | Capillary |
| 3 | CLDN5 | 0 | 14.297946 | Capillary |
| 2 | EPAS1 | 0 | 9.010767 | Capillary |
| 1 | TMEM100 | 0 | 12.608525 | Capillary |
| 22 | VWF | 0 | 3.8297875 | Capillary |
| 23 | MT1M | 0 | 9.386011 | Capillary |
| 24 | ESAM | 0 | 3.2599366 | Capillary |
| 25 | RNASE1 | 0 | 3.0495477 | Capillary |
| 47 | PTPRB | 0 | 3.1512601 | Capillary |
| 46 | TMEM204 | 0 | 2.5954068 | Capillary |
| 45 | EMP2 | 0 | 1.1767288 | Capillary |
| 44 | SLCO2A1 | 0 | 3.1529164 | Capillary |
| 43 | CD36 | 0 | 3.0757592 | Capillary |
| 42 | TM4SF1 | 0 | 3.4714444 | Capillary |
| 41 | ENG | 0 | 3.1503906 | Capillary |
| 40 | ARHGAP29 | 0 | 2.9536653 | Capillary |
| 39 | SPARC | 0 | 2.87669 | Capillary |
| 38 | LDB2 | 0 | 3.2829232 | Capillary |
| 98 | ADCY4 | 0 | 3.3865492 | Capillary |
| 37 | IGFBP4 | 0 | 2.676496 | Capillary |
| 35 | PCAT19 | 0 | 3.377676 | Capillary |
| 34 | LRRC32 | 0 | 4.146231 | Capillary |
| 33 | CD93 | 0 | 3.497653 | Capillary |
| 32 | SLC9A3R2 | 0 | 3.3299277 | Capillary |
| 31 | CDH5 | 0 | 3.1646733 | Capillary |
| 30 | C8orf4 | 0 | 4.556903 | Capillary |
| 29 | BTNL9 | 0 | 4.360277 | Capillary |
| 28 | SLC6A4 | 0 | 5.0533667 | Capillary |
| 27 | GPIHBP1 | 0 | 4.912988 | Capillary |
| 26 | TNFSF10 | 0 | 4.799554 | Capillary |
| 36 | AQP1 | 0 | 3.378832 | Capillary |
| 99 | PLLP | 0 | 2.4741836 | Capillary |
| 0 | EDNRB | 0 | 12.847466 | Capillary Aerocyte |
| 72 | VIPR1 | 0 | 2.9047153 | Capillary Aerocyte |
| 71 | RHOB | 0 | 5.0224123 | Capillary Aerocyte |
| 70 | ID3 | 0 | 3.1375399 | Capillary Aerocyte |
| 69 | SPARCL1 | 0 | 6.301146 | Capillary Aerocyte |
| 68 | TNFSF10 | 0 | 3.5206318 | Capillary Aerocyte |
| 67 | GALNT18 | 0 | 3.3734539 | Capillary Aerocyte |
| 66 | TUBA1A | 0 | 3.8236332 | Capillary Aerocyte |
| 65 | IFITM3 | 0 | 5.347675 | Capillary Aerocyte |
| 64 | SERPINE1 | 0 | 5.942721 | Capillary Aerocyte |
| 63 | B3GALNT1 | 0 | 4.461809 | Capillary Aerocyte |
| 62 | ECSCR | 0 | 2.514611 | Capillary Aerocyte |
| 61 | FOXF1 | 0 | 3.6004448 | Capillary Aerocyte |
| 60 | ACE | 0 | 3.6338224 | Capillary Aerocyte |
| 59 | EFNB1 | 0 | 3.8378937 | Capillary Aerocyte |
| 58 | IFNGR1 | 0 | 3.826809 | Capillary Aerocyte |
| 57 | PRX | 0 | 3.877421 | Capillary Aerocyte |
| 56 | FENDRR | 0 | 3.7491636 | Capillary Aerocyte |
| 55 | CLEC14A | 0 | 3.0539126 | Capillary Aerocyte |

| | | | | |
|---|---|---|---|---|
| 54 | TBX3 | 0 | 4.1313066 | Capillary Aerocyte |
| 53 | BCAM | 0 | 2.5881832 | Capillary Aerocyte |
| 52 | NPDC1 | 0 | 2.8402507 | Capillary Aerocyte |
| 73 | F2RL3 | 0 | 4.0689373 | Capillary Aerocyte |
| 51 | SLCO2A1 | 0 | 2.9368806 | Capillary Aerocyte |
| 74 | ADGRL4 | 0 | 3.5354054 | Capillary Aerocyte |
| 76 | HLA-C | 0 | 7.06729 | Capillary Aerocyte |
| 97 | MYLK | 0 | 2.7437391 | Capillary Aerocyte |
| 96 | BST2 | 0 | 2.4134398 | Capillary Aerocyte |
| 95 | NES | 0 | 3.5347378 | Capillary Aerocyte |
| 94 | CRIP2 | 0 | 1.6985866 | Capillary Aerocyte |
| 93 | PPFIBP1 | 0 | 2.5606847 | Capillary Aerocyte |
| 92 | FAXDC2 | 0 | 3.8395708 | Capillary Aerocyte |
| 91 | SPTBN1 | 0 | 1.9815797 | Capillary Aerocyte |
| 90 | KANK3 | 0 | 2.8362474 | Capillary Aerocyte |
| 89 | CLEC3B | 0 | 1.1490275 | Capillary Aerocyte |
| 88 | HLA-A | 0 | 5.210342 | Capillary Aerocyte |
| 87 | HLA-B | 0 | 8.740597 | Capillary Aerocyte |
| 86 | INPP1 | 0 | 2.7261102 | Capillary Aerocyte |
| 85 | ADGRL2 | 0 | 3.350045 | Capillary Aerocyte |
| 84 | PDLIM1 | 0 | 4.418858 | Capillary Aerocyte |
| 83 | TSPAN15 | 0 | 2.434001 | Capillary Aerocyte |
| 82 | RASIP1 | 0 | 2.80009 | Capillary Aerocyte |
| 81 | CALD1 | 0 | 1.0980256 | Capillary Aerocyte |
| 80 | ZBTB16 | 0 | 2.622846 | Capillary Aerocyte |
| 79 | CYP3A5 | 0 | 5.174524 | Capillary Aerocyte |
| 78 | SH3BP5 | 0 | 2.2006793 | Capillary Aerocyte |
| 77 | JUN | 0 | 4.5476522 | Capillary Aerocyte |
| 75 | CX3CL1 | 0 | 3.7104208 | Capillary Aerocyte |
| 50 | ANXA3 | 0 | 3.3664343 | Capillary Aerocyte |
| 49 | STXBP6 | 0 | 3.441324 | Capillary Aerocyte |
| 48 | LHFP | 0 | 3.4313521 | Capillary Aerocyte |
| 21 | TIMP3 | 0 | 8.271956 | Capillary Aerocyte |
| 20 | SPARC | 0 | 5.9893656 | Capillary Aerocyte |
| 19 | CNN3 | 0 | 7.10659 | Capillary Aerocyte |
| 18 | S100A3 | 0 | 8.655014 | Capillary Aerocyte |
| 17 | C10orf10 | 0 | 11.257608 | Capillary Aerocyte |
| 16 | TMEM204 | 0 | 6.043276 | Capillary Aerocyte |
| 15 | SOSTDC1 | 0 | 11.383046 | Capillary Aerocyte |
| 14 | ICAM2 | 0 | 6.3936353 | Capillary Aerocyte |
| 13 | APP | 0 | 5.8146505 | Capillary Aerocyte |
| 12 | ADIRF | 0 | 10.986112 | Capillary Aerocyte |
| 11 | ESAM | 0 | 5.443962 | Capillary Aerocyte |
| 10 | CAV1 | 0 | 10.793283 | Capillary Aerocyte |
| 9 | SDPR | 0 | 10.972606 | Capillary Aerocyte |
| 8 | CA4 | 0 | 9.241708 | Capillary Aerocyte |
| 7 | HLA-E | 0 | 29.856754 | Capillary Aerocyte |
| 6 | AQP1 | 0 | 9.914691 | Capillary Aerocyte |
| 5 | CLDN5 | 0 | 25.494051 | Capillary Aerocyte |
| 4 | RAMP2 | 0 | 13.642973 | Capillary Aerocyte |
| 3 | GNG11 | 0 | 10.630638 | Capillary Aerocyte |
| 2 | EMCN | 0 | 9.755757 | Capillary Aerocyte |
| 1 | HPGD | 0 | 19.469482 | Capillary Aerocyte |
| 22 | EMP2 | 0 | 4.3920465 | Capillary Aerocyte |
| 23 | CAV2 | 0 | 4.678018 | Capillary Aerocyte |
| 24 | ITM2A | 0 | 4.6972322 | Capillary Aerocyte |
| 25 | SLC9A3R2 | 0 | 4.9204116 | Capillary Aerocyte |
| 47 | ROBO4 | 0 | 3.977233 | Capillary Aerocyte |
| 46 | ACVRL1 | 0 | 3.2358313 | Capillary Aerocyte |
| 45 | PECAM1 | 0 | 3.9027567 | Capillary Aerocyte |
| 44 | RAMP3 | 0 | 2.9516053 | Capillary Aerocyte |
| 43 | IL32 | 0 | 5.138506 | Capillary Aerocyte |
| 42 | ARHGAP29 | 0 | 3.4590073 | Capillary Aerocyte |
| 41 | TINAGL1 | 0 | 4.525635 | Capillary Aerocyte |

| | | | | |
|---|---|---|---|---|
| 40 | ITM2B | 0 | 16.940983 | Capillary Aerocyte |
| 39 | IGFBP4 | 0 | 3.5128927 | Capillary Aerocyte |
| 38 | GPX3 | 0 | 6.5310254 | Capillary Aerocyte |
| 98 | PCDH12 | 0 | 4.134439 | Capillary Aerocyte |
| 37 | TMEM100 | 0 | 5.52185 | Capillary Aerocyte |
| 35 | TSPAN12 | 0 | 4.5671597 | Capillary Aerocyte |
| 34 | IL1RL1 | 0 | 11.425825 | Capillary Aerocyte |
| 33 | PTRF | 0 | 3.621543 | Capillary Aerocyte |
| 32 | IFI27 | 0 | 9.767226 | Capillary Aerocyte |
| 31 | CDH5 | 0 | 3.9514344 | Capillary Aerocyte |
| 30 | EPAS1 | 0 | 4.827614 | Capillary Aerocyte |
| 29 | FRY | 0 | 5.0842404 | Capillary Aerocyte |
| 28 | HYAL2 | 0 | 4.746441 | Capillary Aerocyte |
| 27 | SGK1 | 0 | 5.8157325 | Capillary Aerocyte |
| 26 | EGFL7 | 0 | 4.505956 | Capillary Aerocyte |
| 36 | TM4SF1 | 0 | 5.8784943 | Capillary Aerocyte |
| 99 | TGFBR3 | 0 | 3.0094125 | Capillary Aerocyte |
| 0 | AQP1 | 0 | 23.172909 | Capillary Intermediate 1 |
| 21 | GPX3 | 0 | 18.454659 | Capillary Intermediate 1 |
| 20 | CDH5 | 0 | 7.6905413 | Capillary Intermediate 1 |
| 19 | SLC9A3R2 | 0 | 12.850289 | Capillary Intermediate 1 |
| 18 | IFITM3 | 0 | 31.046396 | Capillary Intermediate 1 |
| 17 | HPGD | 0 | 20.985365 | Capillary Intermediate 1 |
| 16 | TMEM100 | 0 | 24.748144 | Capillary Intermediate 1 |
| 15 | SPARC | 0 | 13.398896 | Capillary Intermediate 1 |
| 14 | HYAL2 | 0 | 15.308523 | Capillary Intermediate 1 |
| 13 | EDNRB | 0 | 14.776031 | Capillary Intermediate 1 |
| 12 | ESAM | 0 | 10.186729 | Capillary Intermediate 1 |
| 11 | GNG11 | 0 | 17.107952 | Capillary Intermediate 1 |
| 10 | EGFL7 | 0 | 14.840845 | Capillary Intermediate 1 |
| 9 | RAMP2 | 0 | 24.470633 | Capillary Intermediate 1 |
| 8 | CA4 | 0 | 20.848022 | Capillary Intermediate 1 |
| 7 | TMEM204 | 0 | 11.819617 | Capillary Intermediate 1 |
| 6 | TIMP3 | 0 | 30.00583 | Capillary Intermediate 1 |
| 5 | CNN3 | 0 | 15.009841 | Capillary Intermediate 1 |
| 4 | SDPR | 0 | 27.136747 | Capillary Intermediate 1 |
| 3 | EPAS1 | 0 | 27.620035 | Capillary Intermediate 1 |
| 2 | HLA-E | 0 | 61.88857 | Capillary Intermediate 1 |
| 1 | CLDN5 | 0 | 62.3699 | Capillary Intermediate 1 |
| 22 | C10orf10 | 0 | 25.913757 | Capillary Intermediate 1 |
| 23 | EMCN | 0 | 10.029988 | Capillary Intermediate 1 |
| 24 | EMP2 | 1.03E-306 | 11.558775 | Capillary Intermediate 1 |
| 25 | ICAM2 | 1.65E-306 | 10.011324 | Capillary Intermediate 1 |
| 26 | RAMP3 | 3.61E-305 | 8.986082 | Capillary Intermediate 1 |
| 27 | PECAM1 | 6.03E-305 | 11.187641 | Capillary Intermediate 1 |
| 28 | CAV1 | 2.66E-304 | 19.26263 | Capillary Intermediate 1 |
| 29 | FCN3 | 1.29E-300 | 28.862911 | Capillary Intermediate 1 |
| 30 | SLCO2A1 | 1.95E-300 | 7.1612926 | Capillary Intermediate 1 |
| 31 | FAM107A | 8.23E-299 | 8.113661 | Capillary Intermediate 1 |
| 32 | APP | 8.34E-297 | 7.9943447 | Capillary Intermediate 1 |
| 33 | CAV2 | 1.26E-293 | 7.6891007 | Capillary Intermediate 1 |
| 34 | CLEC3B | 4.88E-293 | 14.156995 | Capillary Intermediate 1 |
| 35 | ZBTB16 | 1.57E-292 | 6.372155 | Capillary Intermediate 1 |
| 36 | PTRF | 1.66E-292 | 7.59672 | Capillary Intermediate 1 |
| 37 | ACVRL1 | 1.09E-291 | 7.3663077 | Capillary Intermediate 1 |
| 38 | SGK1 | 2.00E-290 | 9.942455 | Capillary Intermediate 1 |
| 39 | ARHGAP29 | 8.65E-290 | 6.993612 | Capillary Intermediate 1 |
| 40 | MT1M | 7.31E-284 | 22.549526 | Capillary Intermediate 1 |
| 41 | PCAT19 | 5.35E-281 | 6.4563427 | Capillary Intermediate 1 |
| 42 | IGFBP4 | 7.52E-279 | 7.8276668 | Capillary Intermediate 1 |
| 43 | VIPR1 | 1.70E-277 | 6.4932046 | Capillary Intermediate 1 |
| 44 | NPDC1 | 5.67E-276 | 5.831395 | Capillary Intermediate 1 |
| 45 | IFI27 | 2.99E-273 | 39.34881 | Capillary Intermediate 1 |
| 46 | ROBO4 | 2.00E-271 | 6.245639 | Capillary Intermediate 1 |

| | | | | |
|---|---|---|---|---|
| 47 | ADIRF | 2.06E-271 | 17.381233 | Capillary Intermediate 1 |
| 48 | FKBP5 | 1.04E-270 | 7.098932 | Capillary Intermediate 1 |
| 49 | MALAT1 | 1.08E-269 | 516.559 | Capillary Intermediate 1 |
| 50 | PRX | 2.83E-262 | 5.7099543 | Capillary Intermediate 1 |
| 51 | ID3 | 7.65E-262 | 9.956681 | Capillary Intermediate 1 |
| 52 | CX3CL1 | 3.01E-261 | 9.622738 | Capillary Intermediate 1 |
| 53 | CLEC14A | 1.03E-260 | 7.8567777 | Capillary Intermediate 1 |
| 54 | SEC14L1 | 1.47E-260 | 5.9011436 | Capillary Intermediate 1 |
| 55 | SPTBN1 | 1.51E-259 | 6.1387076 | Capillary Intermediate 1 |
| 56 | IFNGR1 | 1.53E-258 | 8.872174 | Capillary Intermediate 1 |
| 57 | IL1RL1 | 1.65E-258 | 15.388116 | Capillary Intermediate 1 |
| 58 | ADGRF5 | 4.78E-258 | 5.892532 | Capillary Intermediate 1 |
| 59 | BTNL9 | 6.12E-257 | 5.882188 | Capillary Intermediate 1 |
| 60 | TINAGL1 | 4.68E-256 | 6.927991 | Capillary Intermediate 1 |
| 61 | TM4SF1 | 1.92E-254 | 16.556408 | Capillary Intermediate 1 |
| 62 | ITM2A | 2.60E-254 | 5.970309 | Capillary Intermediate 1 |
| 63 | LHFP | 6.41E-248 | 5.537314 | Capillary Intermediate 1 |
| 64 | F2RL3 | 1.12E-246 | 6.6897917 | Capillary Intermediate 1 |
| 65 | RHOB | 2.26E-246 | 14.176587 | Capillary Intermediate 1 |
| 66 | MT1E | 1.03E-245 | 18.745989 | Capillary Intermediate 1 |
| 67 | STXBP6 | 3.98E-243 | 5.4536023 | Capillary Intermediate 1 |
| 68 | VAMP5 | 1.01E-240 | 6.4797583 | Capillary Intermediate 1 |
| 69 | PPFIBP1 | 4.02E-240 | 5.380092 | Capillary Intermediate 1 |
| 70 | RASIP1 | 2.01E-238 | 5.4037757 | Capillary Intermediate 1 |
| 71 | FENDRR | 2.27E-238 | 5.4935613 | Capillary Intermediate 1 |
| 72 | SLCO4A1 | 2.27E-238 | 6.622822 | Capillary Intermediate 1 |
| 73 | S100A3 | 4.36E-238 | 8.9415 | Capillary Intermediate 1 |
| 74 | SH3BP5 | 6.63E-238 | 5.4936886 | Capillary Intermediate 1 |
| 75 | CRIP2 | 1.44E-236 | 6.0729237 | Capillary Intermediate 1 |
| 76 | S1PR1 | 2.27E-236 | 5.6205664 | Capillary Intermediate 1 |
| 77 | IL32 | 2.76E-236 | 11.284895 | Capillary Intermediate 1 |
| 78 | PDLIM1 | 5.35E-236 | 17.06371 | Capillary Intermediate 1 |
| 79 | CD99 | 1.07E-233 | 8.13465 | Capillary Intermediate 1 |
| 80 | JUN | 2.13E-232 | 18.016376 | Capillary Intermediate 1 |
| 81 | EFNB1 | 9.90E-231 | 5.382318 | Capillary Intermediate 1 |
| 82 | CSRNP1 | 2.10E-230 | 5.2681875 | Capillary Intermediate 1 |
| 83 | NOSTRIN | 2.59E-230 | 5.0373178 | Capillary Intermediate 1 |
| 84 | ECSCR | 3.58E-229 | 4.7654862 | Capillary Intermediate 1 |
| 85 | TCF4 | 5.22E-229 | 5.510098 | Capillary Intermediate 1 |
| 86 | ID1 | 9.01E-228 | 9.057794 | Capillary Intermediate 1 |
| 87 | IFITM2 | 3.53E-224 | 8.296993 | Capillary Intermediate 1 |
| 88 | EDN1 | 6.32E-224 | 12.687734 | Capillary Intermediate 1 |
| 89 | FLT1 | 7.79E-224 | 5.350481 | Capillary Intermediate 1 |
| 90 | HLA-B | 5.14E-222 | 87.043465 | Capillary Intermediate 1 |
| 91 | HLA-A | 1.16E-221 | 45.532085 | Capillary Intermediate 1 |
| 92 | ENG | 2.05E-221 | 5.9373975 | Capillary Intermediate 1 |
| 93 | ADGRL2 | 3.04E-221 | 5.5357404 | Capillary Intermediate 1 |
| 94 | TNFSF10 | 1.64E-220 | 6.8051505 | Capillary Intermediate 1 |
| 95 | HLA-C | 2.43E-220 | 63.264553 | Capillary Intermediate 1 |
| 96 | ITM2B | 4.37E-220 | 32.67099 | Capillary Intermediate 1 |
| 97 | AKAP12 | 3.20E-217 | 9.90782 | Capillary Intermediate 1 |
| 98 | VWF | 6.21E-216 | 5.4782166 | Capillary Intermediate 1 |
| 99 | LRRC32 | 6.70E-216 | 5.867958 | Capillary Intermediate 1 |
| 0 | SLC9A3R2 | 5.06E-165 | 6.259953 | Capillary Intermediate 2 |
| 1 | FCN3 | 4.38E-123 | 5.3385167 | Capillary Intermediate 2 |
| 2 | PCAT19 | 6.14E-116 | 3.3045712 | Capillary Intermediate 2 |
| 3 | EPAS1 | 1.04E-113 | 4.6141376 | Capillary Intermediate 2 |
| 4 | TMEM100 | 1.07E-113 | 3.2105095 | Capillary Intermediate 2 |
| 5 | SDPR | 3.35E-106 | 2.9279056 | Capillary Intermediate 2 |
| 6 | GNG11 | 6.71E-100 | 2.407761 | Capillary Intermediate 2 |
| 7 | MT1M | 4.36E-99 | 3.5378745 | Capillary Intermediate 2 |
| 8 | PTRF | 6.20E-89 | 2.4089367 | Capillary Intermediate 2 |
| 9 | IFI27 | 9.93E-88 | 3.3065817 | Capillary Intermediate 2 |
| 10 | RAMP2 | 3.06E-85 | 1.6727853 | Capillary Intermediate 2 |

| | | | | |
|---|---|---|---|---|
| 11 | CLDN5 | 9.04E-85 | 0.40765193 | Capillary Intermediate 2 |
| 12 | EGFL7 | 2.54E-79 | 2.1587868 | Capillary Intermediate 2 |
| 13 | SPARC | 9.07E-67 | 1.9747869 | Capillary Intermediate 2 |
| 14 | FAM107A | 1.61E-66 | 1.8005638 | Capillary Intermediate 2 |
| 15 | MT1E | 1.06E-64 | 1.5212328 | Capillary Intermediate 2 |
| 16 | SPTBN1 | 2.57E-53 | 1.9063832 | Capillary Intermediate 2 |
| 17 | IFITM3 | 9.06E-50 | 2.6551082 | Capillary Intermediate 2 |
| 18 | TMEM204 | 9.94E-49 | 1.5626799 | Capillary Intermediate 2 |
| 19 | MT1X | 2.27E-46 | -1.815455 | Capillary Intermediate 2 |
| 20 | CAV1 | 1.42E-45 | -0.19400851 | Capillary Intermediate 2 |
| 21 | ADIRF | 4.54E-45 | 1.1249955 | Capillary Intermediate 2 |
| 22 | CA4 | 9.43E-44 | 0.58217746 | Capillary Intermediate 2 |
| 23 | RAMP3 | 8.19E-41 | 1.1546533 | Capillary Intermediate 2 |
| 24 | C10orf10 | 7.16E-39 | -0.67161804 | Capillary Intermediate 2 |
| 25 | MT2A | 5.04E-36 | -12.238876 | Capillary Intermediate 2 |
| 26 | EDN1 | 7.34E-35 | 0.13800518 | Capillary Intermediate 2 |
| 27 | CNN3 | 3.55E-34 | 0.7161534 | Capillary Intermediate 2 |
| 28 | MYZAP | 1.08E-31 | 2.7870662 | Capillary Intermediate 2 |
| 29 | CCND1 | 5.33E-31 | 1.476309 | Capillary Intermediate 2 |
| 30 | ID1 | 7.88E-28 | 0.10788582 | Capillary Intermediate 2 |
| 31 | SPARCL1 | 3.80E-27 | 0.4199004 | Capillary Intermediate 2 |
| 32 | GPX3 | 9.43E-27 | 0.270956 | Capillary Intermediate 2 |
| 33 | VAMP5 | 1.43E-26 | 1.1005294 | Capillary Intermediate 2 |
| 34 | NOSTRIN | 1.75E-25 | 1.4859481 | Capillary Intermediate 2 |
| 35 | TIMP3 | 5.12E-23 | -0.74386626 | Capillary Intermediate 2 |
| 36 | COX7A1 | 6.19E-22 | 1.5692269 | Capillary Intermediate 2 |
| 37 | ID3 | 1.16E-20 | 0.3315707 | Capillary Intermediate 2 |
| 38 | HPGD | 1.65E-20 | 0.24409045 | Capillary Intermediate 2 |
| 39 | LDB2 | 2.36E-20 | 1.1786484 | Capillary Intermediate 2 |
| 40 | PRX | 3.63E-20 | 1.764921 | Capillary Intermediate 2 |
| 41 | STXBP6 | 6.46E-20 | 1.1335129 | Capillary Intermediate 2 |
| 42 | C8orf4 | 2.44E-19 | -0.6726439 | Capillary Intermediate 2 |
| 43 | IGFBP4 | 2.45E-19 | 0.26583987 | Capillary Intermediate 2 |
| 44 | ARHGAP29 | 7.96E-18 | 0.77293533 | Capillary Intermediate 2 |
| 45 | APP | 1.84E-17 | 0.5999422 | Capillary Intermediate 2 |
| 46 | EMCN | 2.14E-17 | 0.5002195 | Capillary Intermediate 2 |
| 47 | GPIHBP1 | 5.14E-17 | 1.1866455 | Capillary Intermediate 2 |
| 48 | KIAA1462 | 5.82E-16 | 2.4867365 | Capillary Intermediate 2 |
| 49 | CAV2 | 8.32E-16 | 0.15997203 | Capillary Intermediate 2 |
| 50 | CLEC3B | 7.39E-15 | -0.9202804 | Capillary Intermediate 2 |
| 51 | IL32 | 3.31E-14 | -0.64139175 | Capillary Intermediate 2 |
| 52 | HYAL2 | 7.11E-14 | -0.19005513 | Capillary Intermediate 2 |
| 53 | ZEB1 | 1.21E-13 | 2.397142 | Capillary Intermediate 2 |
| 54 | MT1L | 1.22E-13 | 0.38677838 | Capillary Intermediate 2 |
| 55 | EDNRB | 3.27E-13 | -0.27319276 | Capillary Intermediate 2 |
| 56 | S100A3 | 1.07E-12 | 0.7794398 | Capillary Intermediate 2 |
| 57 | CALD1 | 4.26E-12 | -0.5318649 | Capillary Intermediate 2 |
| 58 | MT1A | 7.20E-12 | -0.24128221 | Capillary Intermediate 2 |
| 59 | TPM1 | 1.53E-11 | 0.2072823 | Capillary Intermediate 2 |
| 60 | NPDC1 | 1.65E-11 | 0.3238499 | Capillary Intermediate 2 |
| 61 | AQP1 | 2.09E-11 | -0.6846758 | Capillary Intermediate 2 |
| 62 | RNASE1 | 2.39E-11 | -2.5703294 | Capillary Intermediate 2 |
| 63 | MYLK | 9.16E-11 | 1.2420976 | Capillary Intermediate 2 |
| 64 | AKAP12 | 1.61E-10 | -0.40539473 | Capillary Intermediate 2 |
| 65 | BTNL9 | 2.24E-10 | 0.9007462 | Capillary Intermediate 2 |
| 66 | HSPA12B | 3.17E-10 | 1.9147767 | Capillary Intermediate 2 |
| 67 | NDRG2 | 2.48E-09 | 0.90077 | Capillary Intermediate 2 |
| 68 | CD93 | 4.55E-09 | 0.4724385 | Capillary Intermediate 2 |
| 69 | CRIP2 | 1.23E-08 | -0.04374853 | Capillary Intermediate 2 |
| 70 | SH3BP5 | 1.84E-08 | 0.33249062 | Capillary Intermediate 2 |
| 71 | LHFP | 1.25E-07 | 0.3295803 | Capillary Intermediate 2 |
| 72 | CALCRL | 1.50E-07 | 0.35838386 | Capillary Intermediate 2 |
| 73 | ARRDC2 | 2.25E-07 | 0.9476785 | Capillary Intermediate 2 |
| 74 | C1orf115 | 2.48E-07 | 2.181747 | Capillary Intermediate 2 |

| | | | | |
|---|---|---|---|---|
| 75 | ESAM | 3.97E-07 | -0.2697759 | Capillary Intermediate 2 |
| 76 | RASIP1 | 2.03E-06 | 0.66229856 | Capillary Intermediate 2 |
| 77 | S100A16 | 2.15E-06 | -0.006232567 | Capillary Intermediate 2 |
| 78 | ERG | 2.54E-06 | 1.3197109 | Capillary Intermediate 2 |
| 79 | TM4SF1 | 3.49E-06 | -2.6718826 | Capillary Intermediate 2 |
| 80 | FXYD6 | 4.13E-06 | 0.7522445 | Capillary Intermediate 2 |
| 81 | APOL3 | 4.67E-06 | 0.7772491 | Capillary Intermediate 2 |
| 82 | ECSCR | 6.34E-06 | 0.20223461 | Capillary Intermediate 2 |
| 83 | GIMAP6 | 6.40E-06 | 0.9199885 | Capillary Intermediate 2 |
| 84 | VIPR1 | 7.56E-06 | 0.49639192 | Capillary Intermediate 2 |
| 85 | PPFIBP1 | 1.05E-05 | 0.54258853 | Capillary Intermediate 2 |
| 86 | GIMAP4 | 1.19E-05 | 0.40798956 | Capillary Intermediate 2 |
| 87 | SOSTDC1 | 1.36E-05 | -0.41784644 | Capillary Intermediate 2 |
| 88 | FENDRR | 1.96E-05 | 0.61136407 | Capillary Intermediate 2 |
| 89 | VWF | 2.09E-05 | -0.2518857 | Capillary Intermediate 2 |
| 90 | UACA | 2.29E-05 | 0.82382697 | Capillary Intermediate 2 |
| 91 | MTRNR2L2 | 2.50E-05 | 0.5174399 | Capillary Intermediate 2 |
| 92 | FRY | 0.000149439 | 0.4563339 | Capillary Intermediate 2 |
| 93 | BCL6B | 0.000174823 | 1.7273802 | Capillary Intermediate 2 |
| 94 | PKIG | 0.000189433 | 0.3884664 | Capillary Intermediate 2 |
| 95 | ZBTB16 | 0.000192375 | 0.13158998 | Capillary Intermediate 2 |
| 96 | RGS5 | 0.00023743 | 0.4536336 | Capillary Intermediate 2 |
| 97 | GATA2-AS1 | 0.000248484 | 1.7087948 | Capillary Intermediate 2 |
| 98 | TSPAN12 | 0.000297461 | 0.5930058 | Capillary Intermediate 2 |
| 99 | CLEC14A | 0.000320338 | -0.47224343 | Capillary Intermediate 2 |
| 0 | CAPS | 0 | 71.57877 | Ciliated |
| 72 | TMEM190 | 0 | 48.358105 | Ciliated |
| 71 | FAM81B | 0 | 11.403193 | Ciliated |
| 70 | DYNLL1 | 0 | 41.400288 | Ciliated |
| 69 | CCDC17 | 0 | 11.316905 | Ciliated |
| 68 | DNAH12 | 0 | 11.346198 | Ciliated |
| 67 | CD24 | 0 | 15.299616 | Ciliated |
| 66 | CCDC42B | 0 | 11.249831 | Ciliated |
| 65 | KIF9 | 0 | 9.537592 | Ciliated |
| 64 | DPCD | 0 | 9.085671 | Ciliated |
| 63 | ELF3 | 0 | 13.908641 | Ciliated |
| 62 | PERP | 0 | 9.108116 | Ciliated |
| 61 | CFAP43 | 0 | 11.248068 | Ciliated |
| 60 | CFAP53 | 0 | 11.547516 | Ciliated |
| 59 | SPAG6 | 0 | 11.541566 | Ciliated |
| 58 | IK | 0 | 10.358597 | Ciliated |
| 57 | WDR38 | 0 | 12.080961 | Ciliated |
| 56 | TEKT1 | 0 | 12.233368 | Ciliated |
| 55 | MRPS31 | 0 | 10.952851 | Ciliated |
| 54 | ERICH3 | 0 | 12.789406 | Ciliated |
| 53 | FOXJ1 | 0 | 13.001862 | Ciliated |
| 52 | RSPH9 | 0 | 12.16873 | Ciliated |
| 73 | TSTD1 | 0 | 11.915367 | Ciliated |
| 51 | CLDN3 | 0 | 12.295783 | Ciliated |
| 74 | C2orf40 | 0 | 14.6558895 | Ciliated |
| 76 | CKB | 0 | 11.155525 | Ciliated |
| 97 | SPA17 | 0 | 9.2145605 | Ciliated |
| 96 | CCDC153 | 0 | 9.342178 | Ciliated |
| 95 | TSPAN19 | 0 | 13.599787 | Ciliated |
| 94 | TUBA4B | 0 | 11.110656 | Ciliated |
| 93 | DNPH1 | 0 | 8.741447 | Ciliated |
| 92 | FAM174A | 0 | 7.6560597 | Ciliated |
| 91 | PPAP2C | 0 | 8.200809 | Ciliated |
| 90 | WFDC2 | 0 | 12.109947 | Ciliated |
| 89 | CYSTM1 | 0 | 13.752025 | Ciliated |
| 88 | CD164L2 | 0 | 10.716867 | Ciliated |
| 87 | C11orf97 | 0 | 11.916346 | Ciliated |
| 86 | SPAG17 | 0 | 11.109568 | Ciliated |
| 85 | C21orf59 | 0 | 8.647581 | Ciliated |

| | | | | |
|---|---|---|---|---|
| 84 | CRNDE | 0 | 8.77005 | Ciliated |
| 83 | DMKN | 0 | 8.545271 | Ciliated |
| 82 | HMGN3 | 0 | 17.848595 | Ciliated |
| 81 | LDLRAD1 | 0 | 11.417034 | Ciliated |
| 80 | CDS1 | 0 | 8.959678 | Ciliated |
| 79 | DYNLT1 | 0 | 15.744836 | Ciliated |
| 78 | ATPIF1 | 0 | 17.9802 | Ciliated |
| 77 | AGR2 | 0 | 14.1984215 | Ciliated |
| 75 | NME5 | 0 | 10.739291 | Ciliated |
| 50 | TUBB4B | 0 | 35.7399 | Ciliated |
| 49 | FAM216B | 0 | 12.613646 | Ciliated |
| 48 | LRRC46 | 0 | 11.583592 | Ciliated |
| 21 | CAPSL | 0 | 17.05107 | Ciliated |
| 20 | MORN2 | 0 | 18.24113 | Ciliated |
| 19 | ODF3B | 0 | 20.522942 | Ciliated |
| 18 | DYNLRB2 | 0 | 17.014698 | Ciliated |
| 17 | MS4A8 | 0 | 15.520447 | Ciliated |
| 16 | CCDC78 | 0 | 15.141618 | Ciliated |
| 15 | C5orf49 | 0 | 18.482994 | Ciliated |
| 14 | C11orf88 | 0 | 19.792065 | Ciliated |
| 13 | CETN2 | 0 | 25.395468 | Ciliated |
| 12 | C9orf116 | 0 | 19.455755 | Ciliated |
| 11 | ZMYND10 | 0 | 16.62089 | Ciliated |
| 10 | C1orf194 | 0 | 25.40738 | Ciliated |
| 9 | SNTN | 0 | 22.702768 | Ciliated |
| 8 | LRRIQ1 | 0 | 16.755533 | Ciliated |
| 7 | FAM183A | 0 | 26.657724 | Ciliated |
| 6 | TSPAN1 | 0 | 32.23895 | Ciliated |
| 5 | PIFO | 0 | 20.714834 | Ciliated |
| 4 | C9orf24 | 0 | 43.208294 | Ciliated |
| 3 | RSPH1 | 0 | 25.948633 | Ciliated |
| 2 | TPPP3 | 0 | 81.82945 | Ciliated |
| 1 | C20orf85 | 0 | 39.16726 | Ciliated |
| 22 | FAM92B | 0 | 14.653244 | Ciliated |
| 23 | CCDC146 | 0 | 14.051713 | Ciliated |
| 24 | AGR3 | 0 | 31.173746 | Ciliated |
| 25 | EFHC1 | 0 | 11.3108015 | Ciliated |
| 47 | DNALI1 | 0 | 10.921116 | Ciliated |
| 46 | C9orf135 | 0 | 13.029941 | Ciliated |
| 45 | EZR | 0 | 19.165031 | Ciliated |
| 44 | IFT57 | 0 | 12.475347 | Ciliated |
| 43 | EFCAB1 | 0 | 11.972482 | Ciliated |
| 42 | TCTEX1D2 | 0 | 11.827199 | Ciliated |
| 41 | MORN5 | 0 | 14.314496 | Ciliated |
| 40 | LRRC23 | 0 | 11.34273 | Ciliated |
| 39 | C12orf75 | 0 | 14.022468 | Ciliated |
| 38 | CDHR3 | 0 | 13.232251 | Ciliated |
| 98 | TEKT2 | 0 | 10.735368 | Ciliated |
| 37 | TMC5 | 0 | 16.337057 | Ciliated |
| 35 | FAM229B | 0 | 12.356842 | Ciliated |
| 34 | PSENEN | 0 | 15.5090475 | Ciliated |
| 33 | ROPN1L | 0 | 13.748159 | Ciliated |
| 32 | WDR54 | 0 | 11.434403 | Ciliated |
| 31 | SLC44A4 | 0 | 12.690786 | Ciliated |
| 30 | FXYD3 | 0 | 20.427399 | Ciliated |
| 29 | SMIM22 | 0 | 14.570044 | Ciliated |
| 28 | CCDC170 | 0 | 13.373036 | Ciliated |
| 27 | PRDX5 | 0 | 58.915283 | Ciliated |
| 26 | CFAP126 | 0 | 15.40586 | Ciliated |
| 36 | DNAH5 | 0 | 12.483754 | Ciliated |
| 99 | C11orf70 | 0 | 10.221648 | Ciliated |
| 0 | VCAN | 0 | 11.471735 | Classical Monocyte |
| 21 | TNFSF13B | 0 | 2.7086315 | Classical Monocyte |
| 20 | CORO1A | 0 | 3.038834 | Classical Monocyte |

| | | | | |
|---|---|---|---|---|
| 19 | SELL | 0 | 4.3332753 | Classical Monocyte |
| 18 | AIF1 | 0 | 2.24407 | Classical Monocyte |
| 17 | LST1 | 0 | 2.8081772 | Classical Monocyte |
| 16 | COTL1 | 0 | 3.9816477 | Classical Monocyte |
| 15 | CD48 | 0 | 3.3583255 | Classical Monocyte |
| 14 | CTSS | 0 | 4.3919673 | Classical Monocyte |
| 13 | CSTA | 0 | 3.6089208 | Classical Monocyte |
| 12 | MPEG1 | 0 | 3.9335718 | Classical Monocyte |
| 11 | MS4A6A | 0 | 3.3380566 | Classical Monocyte |
| 10 | MNDA | 0 | 4.2189054 | Classical Monocyte |
| 9 | AP1S2 | 0 | 4.3769794 | Classical Monocyte |
| 8 | CFP | 0 | 4.555487 | Classical Monocyte |
| 7 | LYZ | 0 | 40.2436 | Classical Monocyte |
| 6 | RGS2 | 0 | 5.5549607 | Classical Monocyte |
| 5 | FGL2 | 0 | 4.9251366 | Classical Monocyte |
| 4 | S100A9 | 0 | 88.83918 | Classical Monocyte |
| 3 | S100A8 | 0 | 68.549446 | Classical Monocyte |
| 2 | S100A12 | 0 | 14.797408 | Classical Monocyte |
| 1 | FCN1 | 0 | 10.061 | Classical Monocyte |
| 22 | TYROBP | 0 | -5.327552 | Classical Monocyte |
| 23 | C10orf54 | 0 | 2.6664457 | Classical Monocyte |
| 24 | FYB | 1.50E-297 | 2.6656628 | Classical Monocyte |
| 25 | NUP214 | 2.44E-294 | 2.576534 | Classical Monocyte |
| 26 | CD300E | 4.03E-294 | 4.5797114 | Classical Monocyte |
| 27 | SNHG5 | 2.15E-285 | 3.6190953 | Classical Monocyte |
| 28 | CLEC7A | 1.17E-275 | 2.1769474 | Classical Monocyte |
| 29 | RNASE6 | 2.77E-268 | 2.2770603 | Classical Monocyte |
| 30 | RPL39 | 1.79E-265 | 12.312816 | Classical Monocyte |
| 31 | TYMP | 7.58E-264 | 1.3298008 | Classical Monocyte |
| 32 | CD14 | 1.38E-262 | 3.00163 | Classical Monocyte |
| 33 | CDA | 5.78E-249 | 4.923173 | Classical Monocyte |
| 34 | LILRA5 | 1.55E-242 | 3.2933934 | Classical Monocyte |
| 35 | CLEC4E | 1.63E-226 | 2.359682 | Classical Monocyte |
| 36 | IFI30 | 1.66E-225 | -13.52338 | Classical Monocyte |
| 37 | EVI2B | 2.95E-225 | 1.4306632 | Classical Monocyte |
| 38 | FCER1G | 3.97E-219 | -10.330399 | Classical Monocyte |
| 39 | CPVL | 6.98E-219 | 1.3044552 | Classical Monocyte |
| 40 | TKT | 7.05E-219 | 1.4515995 | Classical Monocyte |
| 41 | LGALS2 | 8.20E-218 | 5.36935 | Classical Monocyte |
| 42 | S100A4 | 1.17E-209 | -13.49177 | Classical Monocyte |
| 43 | FAM65B | 2.98E-208 | 2.5402 | Classical Monocyte |
| 44 | CSF3R | 1.95E-202 | 3.6048837 | Classical Monocyte |
| 45 | FTL | 1.48E-197 | -281.47656 | Classical Monocyte |
| 46 | THBS1 | 5.55E-197 | 4.2872543 | Classical Monocyte |
| 47 | STK17B | 3.15E-195 | 2.0102305 | Classical Monocyte |
| 48 | RPS24 | 1.79E-192 | 8.348516 | Classical Monocyte |
| 49 | FGR | 3.55E-189 | 1.5212475 | Classical Monocyte |
| 50 | EVI2A | 7.95E-185 | 2.0928037 | Classical Monocyte |
| 51 | C14orf2 | 4.27E-184 | 1.666717 | Classical Monocyte |
| 52 | PRKCB | 1.04E-181 | 2.9677742 | Classical Monocyte |
| 53 | METTL9 | 8.11E-179 | 1.9953905 | Classical Monocyte |
| 54 | CYBA | 6.28E-178 | -9.956611 | Classical Monocyte |
| 55 | CYBB | 1.06E-177 | 0.4963489 | Classical Monocyte |
| 56 | FPR1 | 1.27E-177 | 0.94468737 | Classical Monocyte |
| 57 | RBP7 | 2.08E-175 | 3.1794546 | Classical Monocyte |
| 58 | FAM26F | 1.26E-170 | 2.2494667 | Classical Monocyte |
| 59 | TNFRSF1B | 1.72E-170 | 2.1883197 | Classical Monocyte |
| 60 | H3F3AP4 | 3.61E-168 | 1.968871 | Classical Monocyte |
| 61 | NEAT1 | 9.97E-167 | 3.467992 | Classical Monocyte |
| 62 | GABARAP | 1.68E-165 | 1.0263538 | Classical Monocyte |
| 63 | HMGB2 | 2.99E-160 | 1.9154811 | Classical Monocyte |
| 64 | ATP5E | 4.43E-154 | 1.2367216 | Classical Monocyte |
| 65 | LIMD2 | 1.38E-151 | 1.7753129 | Classical Monocyte |
| 66 | SMAP2 | 2.85E-147 | 1.8479562 | Classical Monocyte |

| | | | | |
|---|---|---|---|---|
| 67 | CST3 | 3.87E-146 | -9.888462 | Classical Monocyte |
| 68 | MEGF9 | 6.32E-146 | 2.8558352 | Classical Monocyte |
| 69 | LILRB2 | 1.68E-144 | 2.2357006 | Classical Monocyte |
| 70 | C1orf162 | 2.23E-144 | -1.2129632 | Classical Monocyte |
| 71 | GPX1 | 7.46E-144 | -1.5140547 | Classical Monocyte |
| 72 | GCA | 1.81E-142 | 1.0721549 | Classical Monocyte |
| 73 | SH3BGRL3 | 1.06E-138 | -3.6711771 | Classical Monocyte |
| 74 | STXBP2 | 5.00E-137 | -2.8198962 | Classical Monocyte |
| 75 | GLIPR1 | 8.20E-137 | 1.2630745 | Classical Monocyte |
| 76 | MEF2C | 1.00E-136 | 2.120926 | Classical Monocyte |
| 77 | CYP1B1 | 5.20E-136 | 2.3266976 | Classical Monocyte |
| 78 | CD36 | 1.15E-132 | 1.3289589 | Classical Monocyte |
| 79 | PSAP | 1.60E-132 | -8.302756 | Classical Monocyte |
| 80 | RGS18 | 2.48E-132 | 3.7291284 | Classical Monocyte |
| 81 | CLEC12A | 1.94E-130 | 1.5432479 | Classical Monocyte |
| 82 | SRGN | 6.15E-129 | -0.84900206 | Classical Monocyte |
| 83 | CFD | 1.14E-127 | -4.737105 | Classical Monocyte |
| 84 | IRS2 | 3.75E-127 | 1.8515929 | Classical Monocyte |
| 85 | CD93 | 6.55E-127 | 1.3700999 | Classical Monocyte |
| 86 | ICAM3 | 1.84E-126 | 1.8276416 | Classical Monocyte |
| 87 | NCF2 | 2.75E-125 | 0.27648342 | Classical Monocyte |
| 88 | RPS13 | 4.55E-124 | 4.5062604 | Classical Monocyte |
| 89 | LGALS1 | 5.35E-122 | -3.974688 | Classical Monocyte |
| 90 | RASSF2 | 2.58E-120 | 3.158982 | Classical Monocyte |
| 91 | RNF130 | 1.31E-118 | 0.5719442 | Classical Monocyte |
| 92 | CD1D | 2.60E-118 | 4.1686935 | Classical Monocyte |
| 93 | RPL34 | 1.47E-116 | 6.557925 | Classical Monocyte |
| 94 | VNN2 | 3.34E-116 | 3.4756322 | Classical Monocyte |
| 95 | 01-Mar | 3.40E-115 | 1.9133645 | Classical Monocyte |
| 96 | LCP1 | 2.00E-114 | 0.17148186 | Classical Monocyte |
| 97 | SPI1 | 7.22E-114 | -3.0712957 | Classical Monocyte |
| 98 | APOBEC3A | 9.53E-114 | 4.3773155 | Classical Monocyte |
| 99 | CLEC4A | 6.25E-113 | 1.8996742 | Classical Monocyte |
| 0 | SCGB3A2 | 0 | 726.88293 | Club |
| 20 | KRT18 | 0 | 6.9364424 | Club |
| 19 | TACSTD2 | 0 | 7.279893 | Club |
| 18 | ELF3 | 0 | 7.143965 | Club |
| 17 | C19orf33 | 0 | 6.6032195 | Club |
| 16 | SFTA2 | 0 | 7.4376564 | Club |
| 15 | C16orf89 | 0 | 8.0966425 | Club |
| 14 | FXYD3 | 0 | 6.0947666 | Club |
| 13 | KRT7 | 0 | 8.247163 | Club |
| 12 | MUC1 | 0 | 7.572184 | Club |
| 11 | HOPX | 0 | 12.457659 | Club |
| 9 | PIGR | 0 | 12.40018 | Club |
| 8 | CYB5A | 0 | 29.34571 | Club |
| 7 | RNASE1 | 0 | 28.93415 | Club |
| 6 | FOLR1 | 0 | 11.694155 | Club |
| 5 | CYP2B7P | 0 | 9.195024 | Club |
| 4 | CXCL17 | 0 | 14.077798 | Club |
| 3 | AGR3 | 0 | 18.312517 | Club |
| 2 | SFTPB | 0 | 95.658554 | Club |
| 1 | SLPI | 0 | 186.38095 | Club |
| 10 | WFDC2 | 0 | 15.567117 | Club |
| 21 | CLDN4 | 9.05E-307 | 6.55401 | Club |
| 22 | GPRC5A | 4.48E-301 | 6.485247 | Club |
| 23 | KRT8 | 2.67E-297 | 5.7276855 | Club |
| 24 | SFTA1P | 3.76E-291 | 7.2945147 | Club |
| 25 | CYP4B1 | 1.43E-290 | 5.1328244 | Club |
| 26 | SLC34A2 | 2.24E-282 | 6.8247395 | Club |
| 27 | SCGB3A1 | 4.73E-279 | 328.94614 | Club |
| 28 | MGST1 | 5.22E-273 | 7.38768 | Club |
| 29 | S100A14 | 3.99E-271 | 5.447529 | Club |
| 30 | CEACAM6 | 2.57E-270 | 6.97789 | Club |

| | | | | |
|---|---|---|---|---|
| 31 | SFTA3 | 9.44E-270 | 4.748504 | Club |
| 32 | BCAM | 6.42E-258 | 5.6006527 | Club |
| 33 | SDC4 | 2.23E-245 | 5.892073 | Club |
| 34 | ADIRF | 1.64E-244 | 11.142021 | Club |
| 35 | EPCAM | 2.95E-238 | 4.410237 | Club |
| 36 | KRT19 | 7.50E-236 | 2.6519794 | Club |
| 37 | MALL | 1.25E-234 | 4.236093 | Club |
| 38 | SFTPC | 1.20E-231 | -99.52008 | Club |
| 39 | CLU | 2.50E-228 | 7.238997 | Club |
| 40 | STEAP4 | 3.77E-228 | 6.0724387 | Club |
| 41 | ATP1B1 | 1.01E-225 | 6.7557287 | Club |
| 42 | LMO7 | 6.55E-222 | 4.280436 | Club |
| 43 | SELENBP1 | 2.72E-218 | 4.2523336 | Club |
| 44 | NGFRAP1 | 6.14E-217 | 4.3916073 | Club |
| 45 | SFTPA2 | 5.89E-213 | 8.863946 | Club |
| 46 | RNF145 | 2.20E-212 | 4.513121 | Club |
| 47 | AQP4 | 7.81E-211 | 4.9419074 | Club |
| 48 | PTP4A1 | 2.26E-208 | 5.03529 | Club |
| 49 | EZR | 9.28E-208 | 5.121775 | Club |
| 50 | NKX2-1 | 2.10E-201 | 4.4188066 | Club |
| 51 | LRRC75A-AS1 | 6.59E-201 | 9.353457 | Club |
| 52 | TSTD1 | 1.37E-198 | 3.7888446 | Club |
| 53 | AQP5 | 1.81E-197 | 4.0641847 | Club |
| 54 | IFT57 | 6.85E-197 | 4.0973 | Club |
| 55 | ZFP36L1 | 4.13E-193 | 6.4104643 | Club |
| 56 | MAL2 | 2.75E-190 | 4.028738 | Club |
| 57 | GAS5 | 3.98E-184 | 5.1299906 | Club |
| 58 | SOX4 | 1.16E-182 | 6.830368 | Club |
| 59 | CLDN3 | 1.90E-182 | 3.3989022 | Club |
| 60 | RPS6 | 3.61E-181 | 35.2538 | Club |
| 61 | SFTPA1 | 5.47E-178 | -0.99808997 | Club |
| 62 | SPINT2 | 2.96E-175 | 3.9118645 | Club |
| 63 | SFTPD | 5.73E-175 | 6.701788 | Club |
| 64 | MET | 3.84E-174 | 4.8437233 | Club |
| 65 | KIAA1324 | 9.83E-169 | 5.1831164 | Club |
| 66 | CP | 5.53E-168 | 4.1681137 | Club |
| 67 | AGR2 | 1.70E-167 | 3.0292993 | Club |
| 68 | RPS3 | 5.36E-165 | 30.56096 | Club |
| 69 | RPS18 | 4.62E-163 | 43.73492 | Club |
| 70 | KLK11 | 5.79E-161 | 5.45966 | Club |
| 71 | GSTP1 | 7.85E-160 | 9.428602 | Club |
| 72 | NEDD4L | 3.82E-159 | 4.3016334 | Club |
| 73 | RPL24 | 2.66E-157 | 13.760063 | Club |
| 74 | RPL5 | 2.51E-154 | 14.6163845 | Club |
| 75 | SUSD2 | 1.14E-152 | 4.111599 | Club |
| 76 | CLDN7 | 9.12E-152 | 2.9394953 | Club |
| 77 | RPS14 | 1.01E-151 | 34.321102 | Club |
| 78 | EMP2 | 1.06E-150 | 3.906853 | Club |
| 79 | RPL10A | 1.24E-150 | 17.780788 | Club |
| 80 | PRDX5 | 1.56E-150 | 4.024484 | Club |
| 81 | TSC22D1 | 1.38E-149 | 5.152873 | Club |
| 82 | TMC5 | 1.23E-148 | 3.2074115 | Club |
| 83 | SMIM22 | 6.24E-147 | 2.5757458 | Club |
| 84 | HSD17B6 | 1.33E-145 | 4.7740917 | Club |
| 85 | CEBPD | 1.57E-145 | 10.764683 | Club |
| 86 | CTSE | 4.44E-144 | 6.508109 | Club |
| 87 | RPL18 | 1.66E-142 | 17.05494 | Club |
| 88 | RPS5 | 1.30E-139 | 15.363657 | Club |
| 89 | FOSB | 2.13E-137 | 4.858611 | Club |
| 90 | TMEM265 | 7.54E-135 | 3.6897435 | Club |
| 91 | RPL7A | 4.23E-134 | 15.362003 | Club |
| 92 | RPL13 | 1.06E-133 | 38.181774 | Club |
| 93 | DHCR24 | 3.66E-132 | 3.1848843 | Club |
| 94 | GDF15 | 1.71E-131 | 6.9374566 | Club |

| | | | | |
|---|---|---|---|---|
| 95 | RPS4X | 7.58E-130 | 24.691486 | Club |
| 96 | NCOA7 | 1.27E-129 | 4.203212 | Club |
| 97 | RPS8 | 1.63E-129 | 20.480011 | Club |
| 98 | RPL29 | 1.01E-128 | 15.869829 | Club |
| 99 | DSTN | 6.99E-127 | 4.074949 | Club |
| 0 | TACSTD2 | 1.35E-151 | 34.527184 | Differentiating Basal |
| 1 | KRT19 | 2.87E-149 | 123.30984 | Differentiating Basal |
| 2 | PERP | 5.02E-133 | 17.432234 | Differentiating Basal |
| 3 | CLDN4 | 7.66E-131 | 14.245637 | Differentiating Basal |
| 4 | FXYD3 | 2.27E-130 | 19.965837 | Differentiating Basal |
| 5 | AGR2 | 5.26E-123 | 33.718605 | Differentiating Basal |
| 6 | KRT8 | 1.05E-121 | 30.245813 | Differentiating Basal |
| 7 | S100A2 | 1.17E-120 | 21.754997 | Differentiating Basal |
| 8 | C19orf33 | 1.48E-120 | 10.638224 | Differentiating Basal |
| 9 | KRT7 | 2.51E-120 | 15.094686 | Differentiating Basal |
| 10 | KLF5 | 3.67E-111 | 8.185883 | Differentiating Basal |
| 11 | ELF3 | 4.28E-108 | 12.83616 | Differentiating Basal |
| 12 | WFDC2 | 3.02E-107 | 72.36702 | Differentiating Basal |
| 13 | KRT18 | 1.25E-101 | 21.849562 | Differentiating Basal |
| 14 | RPLP0 | 4.15E-101 | 45.21167 | Differentiating Basal |
| 15 | MIR205HG | 8.76E-101 | 10.279327 | Differentiating Basal |
| 16 | SLPI | 6.09E-100 | 178.34073 | Differentiating Basal |
| 17 | SPINT2 | 2.34E-99 | 11.394262 | Differentiating Basal |
| 18 | AQP3 | 4.26E-97 | 27.570822 | Differentiating Basal |
| 19 | GSTP1 | 2.38E-94 | 76.08856 | Differentiating Basal |
| 20 | MDK | 5.89E-92 | 10.782419 | Differentiating Basal |
| 21 | S100A14 | 6.20E-92 | 8.21013 | Differentiating Basal |
| 22 | HSPB1 | 5.16E-90 | 21.317379 | Differentiating Basal |
| 23 | CXCL17 | 7.35E-89 | 22.12609 | Differentiating Basal |
| 24 | EEF1G | 1.25E-87 | 36.682613 | Differentiating Basal |
| 25 | RPL18 | 1.26E-86 | 45.61605 | Differentiating Basal |
| 26 | RPS5 | 2.89E-86 | 38.35292 | Differentiating Basal |
| 27 | RPL7 | 6.03E-86 | 64.03939 | Differentiating Basal |
| 28 | RPL10A | 2.25E-85 | 56.81171 | Differentiating Basal |
| 29 | CAPS | 3.84E-85 | 5.5020685 | Differentiating Basal |
| 30 | RPS18 | 4.54E-85 | 109.725876 | Differentiating Basal |
| 31 | SERPINB3 | 5.10E-85 | 44.941067 | Differentiating Basal |
| 32 | PPAP2C | 1.29E-83 | 7.108892 | Differentiating Basal |
| 33 | RPL13A | 3.69E-83 | 101.83168 | Differentiating Basal |
| 34 | KRT17 | 1.25E-82 | 18.867937 | Differentiating Basal |
| 35 | SDC1 | 1.27E-82 | 6.0962834 | Differentiating Basal |
| 36 | RPSA | 4.22E-82 | 34.41138 | Differentiating Basal |
| 37 | RAB25 | 5.17E-82 | 6.4771094 | Differentiating Basal |
| 38 | AQP5 | 1.36E-80 | 20.914516 | Differentiating Basal |
| 39 | SFN | 1.36E-80 | 6.1240134 | Differentiating Basal |
| 40 | F3 | 1.69E-80 | 8.898661 | Differentiating Basal |
| 41 | MGST1 | 2.76E-80 | 13.315478 | Differentiating Basal |
| 42 | RPS2 | 2.04E-79 | 88.3338 | Differentiating Basal |
| 43 | RPS4X | 2.28E-79 | 85.76397 | Differentiating Basal |
| 44 | RPL8 | 2.32E-79 | 58.466854 | Differentiating Basal |
| 45 | EHF | 2.86E-79 | 5.995921 | Differentiating Basal |
| 46 | RPL3 | 5.69E-77 | 75.80385 | Differentiating Basal |
| 47 | S100A16 | 1.75E-76 | 6.2773657 | Differentiating Basal |
| 48 | CYP4B1 | 1.18E-75 | 9.155685 | Differentiating Basal |
| 49 | SLC25A6 | 2.34E-74 | 16.376293 | Differentiating Basal |
| 50 | RPL4 | 2.35E-74 | 22.044352 | Differentiating Basal |
| 51 | RPS3 | 1.11E-73 | 64.34341 | Differentiating Basal |
| 52 | LRRC75A-AS1 | 3.68E-73 | 18.982664 | Differentiating Basal |
| 53 | DHRS3 | 5.01E-71 | 5.2347207 | Differentiating Basal |
| 54 | RPL13 | 1.28E-70 | 89.4066 | Differentiating Basal |
| 55 | RPS3A | 1.44E-70 | 54.763424 | Differentiating Basal |
| 56 | OCIAD2 | 1.50E-70 | 4.5781317 | Differentiating Basal |
| 57 | ADH1C | 7.50E-70 | 10.49565 | Differentiating Basal |
| 58 | KLK11 | 1.28E-69 | 9.319327 | Differentiating Basal |

| | | | | |
|---|---|---|---|---|
| 59 | TSPAN1 | 7.31E-69 | 14.358159 | Differentiating Basal |
| 60 | RPL29 | 2.21E-68 | 42.272118 | Differentiating Basal |
| 61 | RPLP1 | 3.18E-68 | 127.67138 | Differentiating Basal |
| 62 | RPL18A | 8.43E-68 | 57.20078 | Differentiating Basal |
| 63 | RPL15 | 1.96E-67 | 60.333282 | Differentiating Basal |
| 64 | S100P | 2.57E-67 | 6.330374 | Differentiating Basal |
| 65 | RPS8 | 4.43E-67 | 43.660595 | Differentiating Basal |
| 66 | ZFP36L1 | 1.42E-66 | 10.26028 | Differentiating Basal |
| 67 | KRT10 | 2.99E-66 | 4.972793 | Differentiating Basal |
| 68 | HMGB3 | 8.00E-66 | 7.382571 | Differentiating Basal |
| 69 | MAL2 | 1.46E-65 | 4.4587865 | Differentiating Basal |
| 70 | EPCAM | 2.26E-65 | 6.33701 | Differentiating Basal |
| 71 | RPL5 | 6.85E-65 | 27.004452 | Differentiating Basal |
| 72 | NGFRAP1 | 2.09E-64 | 5.6073847 | Differentiating Basal |
| 73 | RPS19 | 7.96E-64 | 55.33049 | Differentiating Basal |
| 74 | CD9 | 2.12E-63 | 24.113857 | Differentiating Basal |
| 75 | SYTL1 | 2.65E-63 | 5.0931244 | Differentiating Basal |
| 76 | RPL19 | 2.69E-63 | 45.70124 | Differentiating Basal |
| 77 | HMGA1 | 3.47E-63 | 4.9829826 | Differentiating Basal |
| 78 | HES1 | 5.15E-63 | 7.340392 | Differentiating Basal |
| 79 | PABPC1 | 5.30E-63 | 12.551725 | Differentiating Basal |
| 80 | CLDN7 | 9.21E-63 | 7.3666344 | Differentiating Basal |
| 81 | EEF2 | 9.46E-63 | 15.6545725 | Differentiating Basal |
| 82 | RPS14 | 2.14E-62 | 68.36551 | Differentiating Basal |
| 83 | RPL6 | 2.77E-62 | 32.86046 | Differentiating Basal |
| 84 | RPS9 | 2.95E-62 | 43.673153 | Differentiating Basal |
| 85 | CP | 3.75E-62 | 4.855745 | Differentiating Basal |
| 86 | PRDX2 | 5.19E-62 | 5.750939 | Differentiating Basal |
| 87 | GAS5 | 6.20E-62 | 11.569383 | Differentiating Basal |
| 88 | GNB2L1 | 9.74E-62 | 32.293697 | Differentiating Basal |
| 89 | KRT15 | 1.16E-61 | 8.593507 | Differentiating Basal |
| 90 | PRDX5 | 2.92E-60 | 9.804935 | Differentiating Basal |
| 91 | SFTPC | 6.16E-60 | -188.10146 | Differentiating Basal |
| 92 | SPINT1 | 6.60E-60 | 4.853346 | Differentiating Basal |
| 93 | RPS7 | 2.27E-59 | 34.07071 | Differentiating Basal |
| 94 | RPS6 | 2.43E-59 | 69.66296 | Differentiating Basal |
| 95 | OAT | 2.69E-59 | 5.660084 | Differentiating Basal |
| 96 | BTF3 | 2.64E-58 | 18.17624 | Differentiating Basal |
| 97 | SERPINF1 | 5.48E-58 | 6.013933 | Differentiating Basal |
| 98 | CXCL1 | 8.05E-58 | 9.102583 | Differentiating Basal |
| 99 | SEPW1 | 1.07E-57 | 6.787411 | Differentiating Basal |
| 0 | RGS2 | 1.20E-62 | 13.932151 | EREG+ Dendritic |
| 1 | GPR183 | 1.19E-60 | 9.215867 | EREG+ Dendritic |
| 2 | MS4A6A | 4.36E-43 | 6.475378 | EREG+ Dendritic |
| 3 | NAMPT | 1.03E-42 | 8.299991 | EREG+ Dendritic |
| 4 | SAMSN1 | 2.54E-40 | 5.7016115 | EREG+ Dendritic |
| 5 | SRGN | 2.88E-40 | 40.95188 | EREG+ Dendritic |
| 6 | RAB31 | 1.26E-37 | 4.6034594 | EREG+ Dendritic |
| 7 | AREG | 1.72E-37 | 20.488316 | EREG+ Dendritic |
| 8 | MXD1 | 3.03E-34 | 5.3210626 | EREG+ Dendritic |
| 9 | FGL2 | 2.45E-33 | 4.025353 | EREG+ Dendritic |
| 10 | THBS1 | 5.56E-33 | 8.20844 | EREG+ Dendritic |
| 12 | HLA-DPB1 | 5.98E-33 | 34.31246 | EREG+ Dendritic |
| 11 | FCGR2A | 5.98E-33 | 4.4923615 | EREG+ Dendritic |
| 13 | CD93 | 1.21E-32 | 4.4242463 | EREG+ Dendritic |
| 14 | HIF1A | 1.48E-32 | 4.005087 | EREG+ Dendritic |
| 15 | SERPINB9 | 2.85E-32 | 5.127556 | EREG+ Dendritic |
| 16 | TIMP1 | 3.81E-32 | 27.124968 | EREG+ Dendritic |
| 17 | FCN1 | 5.30E-32 | 6.8314433 | EREG+ Dendritic |
| 18 | CD163 | 2.03E-31 | 4.2328978 | EREG+ Dendritic |
| 19 | RNASE6 | 2.03E-31 | 5.0739017 | EREG+ Dendritic |
| 20 | IER3 | 2.37E-31 | 7.051011 | EREG+ Dendritic |
| 21 | HLA-DQB1 | 5.02E-31 | 13.397564 | EREG+ Dendritic |
| 22 | CD300E | 5.17E-31 | 6.3182244 | EREG+ Dendritic |

| | | | | |
|---|---|---|---|---|
| 23 | VEGFA | 7.49E-31 | 4.5439076 | EREG+ Dendritic |
| 24 | COTL1 | 1.39E-30 | 7.109233 | EREG+ Dendritic |
| 25 | CREM | 1.80E-30 | 4.3974366 | EREG+ Dendritic |
| 26 | PLAUR | 1.80E-30 | 7.064623 | EREG+ Dendritic |
| 27 | CLEC7A | 2.11E-30 | 3.403241 | EREG+ Dendritic |
| 28 | LILRB2 | 2.14E-30 | 4.9269433 | EREG+ Dendritic |
| 29 | VCAN | 3.25E-30 | 6.638826 | EREG+ Dendritic |
| 30 | LYZ | 3.29E-30 | 24.696848 | EREG+ Dendritic |
| 31 | MCL1 | 3.61E-29 | 5.758597 | EREG+ Dendritic |
| 32 | HLA-DRA | 8.78E-29 | 42.691284 | EREG+ Dendritic |
| 33 | FPR3 | 1.50E-28 | 4.713164 | EREG+ Dendritic |
| 34 | CTSB | 1.72E-28 | 5.3756957 | EREG+ Dendritic |
| 35 | RGS1 | 1.72E-28 | 8.232449 | EREG+ Dendritic |
| 36 | ATP5E | 1.75E-28 | 8.43194 | EREG+ Dendritic |
| 37 | ACSL1 | 1.95E-28 | 3.3582542 | EREG+ Dendritic |
| 38 | GPX1 | 1.95E-28 | 10.551048 | EREG+ Dendritic |
| 39 | CLEC10A | 2.70E-28 | 7.6457744 | EREG+ Dendritic |
| 40 | TNFAIP3 | 4.31E-28 | 4.2739286 | EREG+ Dendritic |
| 41 | C5AR1 | 4.91E-28 | 3.5645375 | EREG+ Dendritic |
| 42 | CD14 | 1.32E-27 | 7.2275863 | EREG+ Dendritic |
| 43 | EREG | 1.52E-27 | 6.551226 | EREG+ Dendritic |
| 44 | HLA-DQA1 | 1.60E-27 | 10.42472 | EREG+ Dendritic |
| 45 | LST1 | 2.93E-27 | 4.798851 | EREG+ Dendritic |
| 46 | LITAF | 2.94E-27 | 5.7511296 | EREG+ Dendritic |
| 47 | SAT1 | 3.73E-27 | 21.299654 | EREG+ Dendritic |
| 48 | RGS10 | 5.47E-27 | 3.6051724 | EREG+ Dendritic |
| 49 | MAFB | 6.44E-27 | 4.296158 | EREG+ Dendritic |
| 50 | CSF1R | 2.41E-26 | 3.3387444 | EREG+ Dendritic |
| 51 | CHMP1B | 4.50E-26 | 4.230191 | EREG+ Dendritic |
| 52 | FCER1G | 4.95E-26 | 4.642803 | EREG+ Dendritic |
| 53 | TGFBI | 4.95E-26 | 3.706948 | EREG+ Dendritic |
| 54 | TLR2 | 5.32E-26 | 4.2366667 | EREG+ Dendritic |
| 55 | AIF1 | 5.55E-26 | 4.105819 | EREG+ Dendritic |
| 56 | C1orf162 | 6.12E-26 | 5.2367196 | EREG+ Dendritic |
| 57 | HCLS1 | 6.73E-26 | 3.2601528 | EREG+ Dendritic |
| 58 | CPVL | 1.66E-25 | 3.2558267 | EREG+ Dendritic |
| 59 | TYMP | 1.74E-25 | 4.5129113 | EREG+ Dendritic |
| 60 | CST3 | 4.15E-25 | 18.580261 | EREG+ Dendritic |
| 61 | HLA-DQA2 | 4.74E-25 | 7.595125 | EREG+ Dendritic |
| 62 | ATP13A3 | 5.48E-25 | 3.990131 | EREG+ Dendritic |
| 63 | METRNL | 6.70E-25 | 3.4275503 | EREG+ Dendritic |
| 64 | LCP2 | 3.65E-24 | 3.684598 | EREG+ Dendritic |
| 65 | HLA-DPA1 | 5.65E-24 | 28.186058 | EREG+ Dendritic |
| 66 | FPR1 | 6.59E-24 | 4.340547 | EREG+ Dendritic |
| 67 | PPIF | 8.45E-24 | 5.313919 | EREG+ Dendritic |
| 68 | CEBPB | 8.89E-24 | 7.721114 | EREG+ Dendritic |
| 69 | TYROBP | 1.08E-23 | 1.9440669 | EREG+ Dendritic |
| 70 | CLEC5A | 1.41E-23 | 6.098543 | EREG+ Dendritic |
| 71 | DSE | 2.17E-23 | 3.495739 | EREG+ Dendritic |
| 72 | JARID2 | 2.28E-23 | 3.9386034 | EREG+ Dendritic |
| 73 | RPS26 | 2.28E-23 | 11.855841 | EREG+ Dendritic |
| 75 | IFNGR2 | 3.89E-23 | 3.1933124 | EREG+ Dendritic |
| 74 | UPP1 | 3.89E-23 | 3.334965 | EREG+ Dendritic |
| 76 | MAP2K1 | 8.11E-23 | 2.8339753 | EREG+ Dendritic |
| 77 | IL1B | 1.05E-22 | 11.337382 | EREG+ Dendritic |
| 78 | HCST | 2.54E-22 | 3.9008489 | EREG+ Dendritic |
| 79 | CSF2RA | 3.24E-22 | 3.5365214 | EREG+ Dendritic |
| 80 | LAT2 | 3.67E-22 | 2.953263 | EREG+ Dendritic |
| 81 | HLA-DMB | 4.35E-22 | 2.483482 | EREG+ Dendritic |
| 82 | FTH1 | 5.60E-22 | -30.61042 | EREG+ Dendritic |
| 83 | IFI30 | 6.31E-22 | -4.128716 | EREG+ Dendritic |
| 84 | CDKN1A | 1.12E-21 | 2.9726088 | EREG+ Dendritic |
| 85 | RPS24 | 2.25E-21 | 19.963337 | EREG+ Dendritic |
| 86 | PTPRE | 3.89E-21 | 3.6505396 | EREG+ Dendritic |

| | | | | |
|---|---|---|---|---|
| 87 | LIMS1 | 7.84E-21 | 2.9333997 | EREG+ Dendritic |
| 88 | RPL39 | 1.23E-20 | 17.312963 | EREG+ Dendritic |
| 89 | TNFRSF1B | 1.50E-20 | 3.1812963 | EREG+ Dendritic |
| 90 | LINC00152 | 1.95E-20 | 4.1773763 | EREG+ Dendritic |
| 91 | RILPL2 | 2.24E-20 | 2.6278617 | EREG+ Dendritic |
| 92 | SLC2A3 | 2.48E-20 | 3.0598638 | EREG+ Dendritic |
| 93 | HLA-DRB1 | 2.48E-20 | 24.505714 | EREG+ Dendritic |
| 94 | PFKFB3 | 2.68E-20 | 3.885412 | EREG+ Dendritic |
| 95 | EMP3 | 3.71E-20 | 6.4147487 | EREG+ Dendritic |
| 96 | SERP1 | 4.77E-20 | 2.8876638 | EREG+ Dendritic |
| 97 | LAPTM5 | 5.42E-20 | 3.4367254 | EREG+ Dendritic |
| 98 | ADGRE2 | 5.92E-20 | 4.239473 | EREG+ Dendritic |
| 99 | CMTM6 | 6.81E-20 | 2.658278 | EREG+ Dendritic |
| 0 | CLU | 7.47E-58 | 84.65867 | Fibromyocyte |
| 1 | ACTA2 | 8.34E-58 | 67.160324 | Fibromyocyte |
| 2 | TAGLN | 5.63E-57 | 64.30975 | Fibromyocyte |
| 3 | TPM2 | 1.13E-55 | 18.883337 | Fibromyocyte |
| 4 | MYL9 | 8.15E-55 | 22.612368 | Fibromyocyte |
| 5 | CSRP1 | 8.92E-53 | 10.370475 | Fibromyocyte |
| 6 | MYLK | 2.51E-52 | 12.573694 | Fibromyocyte |
| 7 | SPARCL1 | 8.96E-52 | 21.884438 | Fibromyocyte |
| 8 | BCHE | 8.96E-52 | 13.309469 | Fibromyocyte |
| 9 | CRYAB | 5.11E-51 | 12.578212 | Fibromyocyte |
| 10 | DCN | 1.52E-50 | 17.427645 | Fibromyocyte |
| 11 | CTGF | 9.04E-49 | 56.135204 | Fibromyocyte |
| 12 | CALD1 | 8.15E-48 | 10.396056 | Fibromyocyte |
| 13 | ACTG2 | 1.08E-47 | 18.864937 | Fibromyocyte |
| 14 | ASPN | 1.10E-47 | 11.86421 | Fibromyocyte |
| 15 | CNN1 | 1.15E-47 | 12.958455 | Fibromyocyte |
| 16 | MYH11 | 1.15E-47 | 9.637651 | Fibromyocyte |
| 17 | RAMP1 | 1.06E-46 | 8.745997 | Fibromyocyte |
| 18 | PRELP | 5.88E-46 | 9.547936 | Fibromyocyte |
| 19 | IGFBP7 | 6.05E-45 | 26.893894 | Fibromyocyte |
| 20 | FHL1 | 6.15E-45 | 10.630317 | Fibromyocyte |
| 21 | CKB | 1.41E-44 | 9.414175 | Fibromyocyte |
| 22 | DKK3 | 1.46E-44 | 9.368924 | Fibromyocyte |
| 23 | BGN | 6.52E-44 | 10.57294 | Fibromyocyte |
| 24 | MGP | 3.46E-43 | 14.520166 | Fibromyocyte |
| 25 | COL6A2 | 4.06E-43 | 6.718232 | Fibromyocyte |
| 26 | MFAP4 | 6.72E-43 | 8.518693 | Fibromyocyte |
| 27 | SCARA3 | 2.30E-42 | 7.809654 | Fibromyocyte |
| 28 | LTBP1 | 2.03E-40 | 7.8315816 | Fibromyocyte |
| 29 | AEBP1 | 2.57E-40 | 7.0785084 | Fibromyocyte |
| 30 | COL1A2 | 4.34E-40 | 9.623089 | Fibromyocyte |
| 31 | C9orf3 | 1.63E-38 | 5.486064 | Fibromyocyte |
| 32 | SELM | 2.17E-38 | 6.945098 | Fibromyocyte |
| 33 | CYR61 | 2.43E-38 | 22.118475 | Fibromyocyte |
| 34 | DES | 2.76E-38 | 12.649516 | Fibromyocyte |
| 35 | GEM | 2.82E-38 | 8.687058 | Fibromyocyte |
| 36 | PDLIM3 | 3.34E-38 | 6.846368 | Fibromyocyte |
| 37 | RARRES2 | 2.24E-36 | 6.5119815 | Fibromyocyte |
| 38 | DSTN | 8.10E-36 | 9.726666 | Fibromyocyte |
| 39 | PDLIM7 | 2.15E-35 | 5.859714 | Fibromyocyte |
| 40 | C1S | 3.79E-35 | 5.036002 | Fibromyocyte |
| 41 | COL6A1 | 4.32E-35 | 4.965599 | Fibromyocyte |
| 42 | FILIP1L | 1.44E-34 | 7.556698 | Fibromyocyte |
| 43 | SYNPO2 | 1.69E-34 | 7.98471 | Fibromyocyte |
| 44 | TPM1 | 1.96E-33 | 6.798713 | Fibromyocyte |
| 45 | SDC2 | 2.96E-33 | 4.5878353 | Fibromyocyte |
| 46 | ADAMTS1 | 3.71E-32 | 7.7839627 | Fibromyocyte |
| 47 | CNN3 | 3.41E-31 | 5.1573453 | Fibromyocyte |
| 48 | PPP1R14A | 4.05E-31 | 5.0649686 | Fibromyocyte |
| 49 | WIF1 | 8.26E-30 | 7.2794204 | Fibromyocyte |
| 50 | TGFB1I1 | 1.18E-29 | 5.1953692 | Fibromyocyte |

| | | | | |
|---|---|---|---|---|
| 51 | LMOD1 | 1.24E-29 | 6.2416387 | Fibromyocyte |
| 52 | BTG2 | 2.76E-29 | 10.493586 | Fibromyocyte |
| 53 | PLS3 | 8.23E-29 | 4.4526916 | Fibromyocyte |
| 54 | TSC22D1 | 1.32E-28 | 9.990872 | Fibromyocyte |
| 55 | SBSPON | 2.25E-28 | 7.599267 | Fibromyocyte |
| 56 | ANGPTL1 | 2.81E-27 | 6.248492 | Fibromyocyte |
| 57 | CRIP2 | 1.64E-26 | 3.708398 | Fibromyocyte |
| 58 | GAS6 | 5.38E-26 | 4.02332 | Fibromyocyte |
| 59 | COL3A1 | 2.02E-25 | 5.0160203 | Fibromyocyte |
| 60 | PTN | 2.23E-25 | 5.549183 | Fibromyocyte |
| 61 | FXYD1 | 1.02E-24 | 5.0410748 | Fibromyocyte |
| 62 | C1R | 5.00E-24 | 4.277868 | Fibromyocyte |
| 63 | HSPB1 | 5.55E-24 | 10.607452 | Fibromyocyte |
| 64 | SOD3 | 6.76E-24 | 4.0989833 | Fibromyocyte |
| 65 | GREM2 | 7.50E-24 | 9.666832 | Fibromyocyte |
| 66 | KCNMB1 | 8.02E-24 | 6.082677 | Fibromyocyte |
| 67 | SCX | 8.94E-24 | 10.306992 | Fibromyocyte |
| 68 | SMTN | 1.35E-23 | 5.1495914 | Fibromyocyte |
| 69 | CFH | 1.35E-23 | 4.4671135 | Fibromyocyte |
| 70 | CAV1 | 6.53E-23 | 5.0987554 | Fibromyocyte |
| 71 | COL1A1 | 1.38E-22 | 5.1443987 | Fibromyocyte |
| 72 | LMCD1 | 2.30E-22 | 4.73896 | Fibromyocyte |
| 73 | NDN | 6.32E-22 | 4.116295 | Fibromyocyte |
| 74 | FGF18 | 8.63E-22 | 7.9489465 | Fibromyocyte |
| 75 | RASL12 | 1.59E-21 | 5.60354 | Fibromyocyte |
| 76 | SGCA | 1.59E-21 | 5.7812066 | Fibromyocyte |
| 77 | FBLN1 | 1.89E-21 | 2.4139607 | Fibromyocyte |
| 78 | A2M | 9.52E-21 | 3.1476738 | Fibromyocyte |
| 79 | NEXN | 2.51E-20 | 4.5532017 | Fibromyocyte |
| 80 | MT1X | 4.35E-20 | 11.201084 | Fibromyocyte |
| 81 | TNC | 4.42E-20 | 6.0052056 | Fibromyocyte |
| 82 | ID4 | 2.99E-19 | 4.393339 | Fibromyocyte |
| 83 | EMILIN1 | 5.03E-19 | 4.5846467 | Fibromyocyte |
| 84 | TUBA1A | 9.69E-19 | 5.347815 | Fibromyocyte |
| 85 | FSTL3 | 1.05E-18 | 2.2909784 | Fibromyocyte |
| 86 | NR4A1 | 1.05E-18 | 6.797406 | Fibromyocyte |
| 87 | LUM | 4.76E-18 | 2.4025564 | Fibromyocyte |
| 88 | PRKCDBP | 1.92E-17 | 3.1917217 | Fibromyocyte |
| 89 | LPP | 2.07E-17 | 3.2899387 | Fibromyocyte |
| 90 | NGFRAP1 | 2.62E-17 | 2.4087467 | Fibromyocyte |
| 91 | MXRA8 | 2.72E-17 | 4.3882427 | Fibromyocyte |
| 92 | IGFBP5 | 5.42E-17 | 5.533668 | Fibromyocyte |
| 93 | FSTL1 | 6.09E-17 | 3.8957746 | Fibromyocyte |
| 94 | PKIG | 6.16E-17 | 3.3555043 | Fibromyocyte |
| 95 | GSN | 6.77E-17 | 3.5086129 | Fibromyocyte |
| 96 | COL16A1 | 8.82E-17 | 5.708973 | Fibromyocyte |
| 97 | ANXA6 | 1.70E-16 | 2.9060423 | Fibromyocyte |
| 98 | ACTN1 | 1.76E-16 | 3.3204513 | Fibromyocyte |
| 99 | PALLD | 1.91E-16 | 4.206486 | Fibromyocyte |
| 0 | WFDC2 | 4.73E-99 | 441.7562 | Goblet |
| 1 | KRT19 | 4.83E-99 | 296.06516 | Goblet |
| 2 | AQP5 | 7.52E-99 | 62.83476 | Goblet |
| 3 | S100P | 8.66E-99 | 52.75709 | Goblet |
| 4 | TSPAN1 | 2.32E-98 | 65.71772 | Goblet |
| 5 | AGR2 | 3.20E-98 | 199.65083 | Goblet |
| 6 | LCN2 | 1.77E-97 | 167.2807 | Goblet |
| 7 | CEACAM6 | 1.58E-96 | 27.397522 | Goblet |
| 8 | SLPI | 1.87E-96 | 881.7225 | Goblet |
| 9 | KRT7 | 6.17E-96 | 59.498943 | Goblet |
| 10 | FAM3D | 6.17E-96 | 27.425976 | Goblet |
| 11 | MSMB | 1.30E-94 | 114.3096 | Goblet |
| 12 | CXCL17 | 1.39E-94 | 92.01488 | Goblet |
| 13 | FXYD3 | 1.39E-94 | 63.824123 | Goblet |
| 14 | TACSTD2 | 3.24E-94 | 59.29257 | Goblet |

| | | | | |
|---|---|---|---|---|
| 15 | KRT18 | 3.67E-94 | 57.63084 | Goblet |
| 16 | MDK | 4.18E-94 | 33.983013 | Goblet |
| 17 | SLC44A4 | 8.36E-93 | 12.576586 | Goblet |
| 18 | BPIFB1 | 2.29E-92 | 356.63416 | Goblet |
| 19 | SCGB1A1 | 2.44E-92 | 758.29956 | Goblet |
| 20 | PRSS23 | 8.47E-92 | 44.21116 | Goblet |
| 21 | CD24 | 9.06E-92 | 26.241535 | Goblet |
| 22 | KRT8 | 9.06E-92 | 75.54417 | Goblet |
| 23 | C19orf33 | 4.03E-91 | 24.402605 | Goblet |
| 24 | XBP1 | 2.79E-90 | 46.688854 | Goblet |
| 25 | GSTP1 | 4.37E-90 | 223.46852 | Goblet |
| 26 | CP | 9.17E-90 | 16.895739 | Goblet |
| 27 | CLDN7 | 9.17E-90 | 28.404367 | Goblet |
| 28 | ELF3 | 1.44E-89 | 25.787216 | Goblet |
| 29 | STARD10 | 3.69E-88 | 11.673493 | Goblet |
| 30 | TSPAN8 | 7.05E-88 | 36.264523 | Goblet |
| 31 | CYP2F1 | 2.25E-85 | 20.387888 | Goblet |
| 32 | MUC20 | 2.63E-85 | 16.629559 | Goblet |
| 33 | MUC4 | 3.27E-85 | 13.684989 | Goblet |
| 34 | PERP | 3.33E-85 | 22.815332 | Goblet |
| 35 | CYP4B1 | 5.80E-84 | 17.032492 | Goblet |
| 36 | TFF3 | 6.19E-84 | 81.65992 | Goblet |
| 37 | SERPINB3 | 6.63E-84 | 73.83361 | Goblet |
| 38 | CLDN4 | 1.19E-83 | 20.77421 | Goblet |
| 39 | ALDH1A1 | 1.83E-83 | 29.03654 | Goblet |
| 40 | CAPS | 2.89E-83 | 8.946852 | Goblet |
| 41 | PIGR | 5.19E-83 | 42.00774 | Goblet |
| 42 | S100A16 | 5.83E-83 | 21.400293 | Goblet |
| 43 | ZG16B | 1.21E-82 | 11.005572 | Goblet |
| 44 | MUC1 | 1.83E-82 | 20.43177 | Goblet |
| 45 | ASS1 | 2.71E-82 | 13.299385 | Goblet |
| 46 | SPINT1 | 2.78E-82 | 9.38383 | Goblet |
| 47 | PPAP2C | 2.92E-82 | 13.980297 | Goblet |
| 48 | EPCAM | 3.09E-82 | 16.52675 | Goblet |
| 49 | SPINT2 | 4.07E-82 | 25.820238 | Goblet |
| 50 | SCGB3A1 | 6.09E-82 | 120.02263 | Goblet |
| 51 | ST6GALNAC1 | 1.73E-81 | 12.642986 | Goblet |
| 52 | HMGB3 | 9.99E-81 | 13.67037 | Goblet |
| 53 | MAL2 | 4.42E-80 | 11.7673025 | Goblet |
| 54 | CLDN3 | 6.38E-80 | 18.183977 | Goblet |
| 55 | EHF | 7.19E-80 | 9.513115 | Goblet |
| 56 | KLF5 | 1.20E-79 | 14.362909 | Goblet |
| 57 | AGR3 | 5.00E-79 | 24.991096 | Goblet |
| 58 | TMEM205 | 5.32E-79 | 10.523678 | Goblet |
| 59 | CYP2B7P | 6.03E-79 | 9.947274 | Goblet |
| 60 | F3 | 6.03E-79 | 14.719105 | Goblet |
| 61 | AQP3 | 1.15E-78 | 36.343483 | Goblet |
| 62 | S100A14 | 1.30E-78 | 21.596561 | Goblet |
| 63 | ATP1B1 | 1.80E-78 | 19.652279 | Goblet |
| 64 | MGST1 | 2.06E-78 | 32.87747 | Goblet |
| 65 | HS3ST1 | 3.65E-78 | 10.091167 | Goblet |
| 66 | SLC9A3R1 | 3.99E-78 | 11.952948 | Goblet |
| 67 | CHST9 | 4.74E-78 | 10.446347 | Goblet |
| 68 | ALCAM | 9.85E-78 | 9.799997 | Goblet |
| 69 | VMO1 | 1.68E-77 | 29.797783 | Goblet |
| 70 | SSR4 | 1.91E-77 | 27.68549 | Goblet |
| 71 | EZR | 1.97E-77 | 22.091253 | Goblet |
| 72 | HES4 | 2.67E-77 | 12.961867 | Goblet |
| 73 | CFB | 5.46E-77 | 13.291334 | Goblet |
| 74 | TSPAN13 | 1.85E-76 | 10.430193 | Goblet |
| 75 | CDC42EP5 | 3.13E-76 | 10.529597 | Goblet |
| 76 | CCNO | 5.05E-76 | 18.513771 | Goblet |
| 77 | RPLP0 | 5.71E-76 | 73.88963 | Goblet |
| 78 | HEBP2 | 9.00E-76 | 13.317752 | Goblet |

| | | | | |
|---|---|---|---|---|
| 79 | OAT | 1.89E-75 | 10.483229 | Goblet |
| 80 | PRDX2 | 3.02E-75 | 13.22269 | Goblet |
| 81 | RAB25 | 6.48E-75 | 13.5986395 | Goblet |
| 82 | SMIM22 | 7.96E-75 | 8.282875 | Goblet |
| 83 | SELENBP1 | 1.01E-74 | 11.933135 | Goblet |
| 84 | BIK | 1.45E-74 | 9.631343 | Goblet |
| 85 | NDRG2 | 1.92E-74 | 9.140347 | Goblet |
| 86 | BAG1 | 2.87E-74 | 12.502329 | Goblet |
| 87 | RARRES3 | 3.28E-74 | 19.194157 | Goblet |
| 88 | CLDN10 | 6.44E-74 | 13.055656 | Goblet |
| 89 | TSTA3 | 1.63E-73 | 9.199533 | Goblet |
| 90 | VSIG2 | 2.57E-73 | 9.407529 | Goblet |
| 91 | MISP | 5.37E-73 | 9.361742 | Goblet |
| 92 | NUCB2 | 6.00E-73 | 21.091862 | Goblet |
| 93 | ST14 | 7.14E-73 | 7.9701557 | Goblet |
| 94 | PLAC8 | 7.88E-73 | 20.03866 | Goblet |
| 95 | SOX2 | 1.02E-72 | 11.965024 | Goblet |
| 96 | WFDC21P | 1.56E-72 | 17.254389 | Goblet |
| 97 | EPS8L1 | 1.74E-72 | 8.963161 | Goblet |
| 98 | PPDPF | 2.58E-72 | 58.156643 | Goblet |
| 99 | ST6GAL1 | 4.26E-72 | 8.82173 | Goblet |
| 0 | SEPP1 | 9.88E-144 | 20.753431 | IGSF21+ Dendritic |
| 1 | FOLR2 | 4.99E-139 | 12.835161 | IGSF21+ Dendritic |
| 2 | F13A1 | 3.88E-120 | 9.834774 | IGSF21+ Dendritic |
| 3 | MS4A6A | 4.65E-110 | 8.347639 | IGSF21+ Dendritic |
| 4 | STAB1 | 4.30E-105 | 7.9068737 | IGSF21+ Dendritic |
| 5 | MAFB | 8.30E-95 | 5.7314253 | IGSF21+ Dendritic |
| 6 | TMEM176B | 1.62E-91 | 4.9203486 | IGSF21+ Dendritic |
| 7 | SLC40A1 | 1.58E-89 | 7.2569184 | IGSF21+ Dendritic |
| 8 | RGS1 | 1.57E-87 | 9.546888 | IGSF21+ Dendritic |
| 9 | CSF1R | 1.67E-75 | 4.142092 | IGSF21+ Dendritic |
| 10 | GGTA1P | 4.29E-74 | 3.95262 | IGSF21+ Dendritic |
| 11 | HLA-DPB1 | 1.01E-72 | 22.913496 | IGSF21+ Dendritic |
| 12 | FGL2 | 1.39E-72 | 4.4961925 | IGSF21+ Dendritic |
| 13 | CD14 | 2.89E-72 | 7.0994534 | IGSF21+ Dendritic |
| 14 | CD163 | 5.22E-71 | 3.9889429 | IGSF21+ Dendritic |
| 15 | HLA-DQA1 | 1.14E-69 | 12.13087 | IGSF21+ Dendritic |
| 16 | GPR183 | 5.99E-69 | 4.86795 | IGSF21+ Dendritic |
| 17 | SLCO2B1 | 3.88E-68 | 3.6108785 | IGSF21+ Dendritic |
| 18 | CCL13 | 6.04E-67 | 9.255622 | IGSF21+ Dendritic |
| 19 | C1QC | 1.64E-65 | 1.6913311 | IGSF21+ Dendritic |
| 20 | RNASE1 | 3.53E-65 | 13.111758 | IGSF21+ Dendritic |
| 21 | HLA-DPA1 | 3.55E-64 | 23.47192 | IGSF21+ Dendritic |
| 22 | HLA-DRA | 5.07E-64 | 30.233913 | IGSF21+ Dendritic |
| 24 | FCGR2A | 1.25E-60 | 4.0709023 | IGSF21+ Dendritic |
| 23 | HLA-DQB1 | 1.25E-60 | 7.2888975 | IGSF21+ Dendritic |
| 25 | C1QA | 4.30E-58 | -12.913478 | IGSF21+ Dendritic |
| 26 | C1QB | 8.59E-56 | -22.727955 | IGSF21+ Dendritic |
| 27 | RNASE6 | 9.92E-56 | 3.46851 | IGSF21+ Dendritic |
| 28 | AIF1 | 1.19E-55 | 3.9124591 | IGSF21+ Dendritic |
| 29 | GPR34 | 1.43E-55 | 5.223758 | IGSF21+ Dendritic |
| 30 | HLA-DRB1 | 1.68E-55 | 15.189757 | IGSF21+ Dendritic |
| 31 | LTC4S | 4.52E-54 | 3.7878673 | IGSF21+ Dendritic |
| 32 | CD74 | 1.66E-52 | 29.807623 | IGSF21+ Dendritic |
| 33 | MRC1 | 1.76E-52 | 2.5890913 | IGSF21+ Dendritic |
| 34 | RGS10 | 1.78E-52 | 3.1515195 | IGSF21+ Dendritic |
| 35 | MS4A7 | 2.06E-52 | 1.7515347 | IGSF21+ Dendritic |
| 36 | DAB2 | 2.62E-50 | 2.8268762 | IGSF21+ Dendritic |
| 37 | HLA-DRB5 | 2.28E-49 | 4.5648007 | IGSF21+ Dendritic |
| 38 | TMEM176A | 3.70E-49 | 3.8373752 | IGSF21+ Dendritic |
| 39 | HLA-DRB6 | 1.34E-47 | 5.1213923 | IGSF21+ Dendritic |
| 40 | HLA-DQA2 | 2.71E-47 | 3.556332 | IGSF21+ Dendritic |
| 41 | CST3 | 9.75E-45 | 7.358277 | IGSF21+ Dendritic |
| 42 | HLA-DMA | 2.28E-43 | 2.8470597 | IGSF21+ Dendritic |

| | | | | |
|---|---|---|---|---|
| 43 | FCGR2B | 9.36E-43 | 5.503927 | IGSF21+ Dendritic |
| 44 | MS4A4A | 2.70E-41 | 1.3745202 | IGSF21+ Dendritic |
| 45 | TYROBP | 2.18E-40 | -6.834715 | IGSF21+ Dendritic |
| 46 | KCTD12 | 7.75E-40 | 2.4655755 | IGSF21+ Dendritic |
| 47 | HLA-DMB | 4.62E-39 | 2.4061627 | IGSF21+ Dendritic |
| 48 | FPR3 | 6.79E-39 | 3.4397788 | IGSF21+ Dendritic |
| 49 | LGMN | 1.35E-37 | 4.452985 | IGSF21+ Dendritic |
| 50 | FOS | 4.86E-37 | 9.622794 | IGSF21+ Dendritic |
| 51 | CTSC | 9.81E-37 | -1.9613022 | IGSF21+ Dendritic |
| 52 | GPX1 | 1.04E-36 | 3.9367824 | IGSF21+ Dendritic |
| 53 | PDK4 | 3.73E-36 | 3.2902067 | IGSF21+ Dendritic |
| 54 | CFD | 1.26E-35 | -0.69062364 | IGSF21+ Dendritic |
| 55 | FTL | 2.00E-34 | -223.6983 | IGSF21+ Dendritic |
| 56 | CTSB | 3.15E-32 | -0.5781467 | IGSF21+ Dendritic |
| 57 | LILRB5 | 6.14E-32 | 6.5687666 | IGSF21+ Dendritic |
| 58 | MAF | 1.02E-31 | 3.5191174 | IGSF21+ Dendritic |
| 59 | CCL2 | 4.10E-31 | 6.4298496 | IGSF21+ Dendritic |
| 60 | CTSZ | 1.79E-30 | 0.54907835 | IGSF21+ Dendritic |
| 61 | CLEC10A | 2.46E-30 | 4.312351 | IGSF21+ Dendritic |
| 62 | MARCKS | 6.35E-30 | 1.5636038 | IGSF21+ Dendritic |
| 63 | IGSF21 | 8.28E-30 | 9.316065 | IGSF21+ Dendritic |
| 64 | RBPJ | 9.11E-30 | 2.2363992 | IGSF21+ Dendritic |
| 65 | NPC2 | 1.10E-29 | -0.21536605 | IGSF21+ Dendritic |
| 66 | TGFBI | 1.10E-29 | 1.8107266 | IGSF21+ Dendritic |
| 67 | FCER1G | 1.94E-29 | -9.875451 | IGSF21+ Dendritic |
| 68 | KLF6 | 2.02E-29 | 4.443109 | IGSF21+ Dendritic |
| 69 | C3AR1 | 1.97E-28 | 1.6402669 | IGSF21+ Dendritic |
| 70 | IER3 | 2.76E-27 | 4.2148113 | IGSF21+ Dendritic |
| 71 | MFSD1 | 1.05E-26 | 1.726525 | IGSF21+ Dendritic |
| 72 | HLA-DQB2 | 3.07E-26 | 1.7809063 | IGSF21+ Dendritic |
| 73 | PLTP | 2.21E-25 | 3.0074093 | IGSF21+ Dendritic |
| 74 | FAM26F | 8.02E-25 | 2.4878392 | IGSF21+ Dendritic |
| 75 | CD93 | 4.37E-24 | 2.0931697 | IGSF21+ Dendritic |
| 76 | CD4 | 2.62E-23 | 1.3823892 | IGSF21+ Dendritic |
| 77 | HCLS1 | 1.34E-21 | 1.380108 | IGSF21+ Dendritic |
| 78 | HLA-DOA | 2.57E-21 | 2.3654346 | IGSF21+ Dendritic |
| 79 | MEF2C | 3.52E-20 | 2.236097 | IGSF21+ Dendritic |
| 80 | CLEC7A | 2.01E-19 | 1.8315498 | IGSF21+ Dendritic |
| 81 | CYBA | 2.14E-19 | -9.681186 | IGSF21+ Dendritic |
| 82 | RAB31 | 4.44E-19 | 1.5607382 | IGSF21+ Dendritic |
| 83 | RGS2 | 1.38E-18 | 2.593914 | IGSF21+ Dendritic |
| 84 | MPEG1 | 2.06E-18 | 2.1835213 | IGSF21+ Dendritic |
| 85 | C1orf162 | 2.31E-18 | -0.99794275 | IGSF21+ Dendritic |
| 86 | CPM | 3.91E-18 | 1.8805258 | IGSF21+ Dendritic |
| 87 | HMOX1 | 4.66E-18 | 5.0446835 | IGSF21+ Dendritic |
| 88 | PSAP | 7.36E-18 | -7.201225 | IGSF21+ Dendritic |
| 89 | FCGRT | 9.55E-18 | -1.0244532 | IGSF21+ Dendritic |
| 90 | CD68 | 1.52E-17 | -6.531065 | IGSF21+ Dendritic |
| 91 | SLC1A3 | 1.57E-17 | 2.5122771 | IGSF21+ Dendritic |
| 92 | ADAP2 | 1.99E-17 | 2.56532 | IGSF21+ Dendritic |
| 93 | ZFP36L1 | 8.27E-17 | 1.6217711 | IGSF21+ Dendritic |
| 94 | GAS6 | 1.38E-16 | 1.2689266 | IGSF21+ Dendritic |
| 95 | SPI1 | 2.44E-16 | -2.7415187 | IGSF21+ Dendritic |
| 96 | A2M | 1.40E-15 | 0.54437476 | IGSF21+ Dendritic |
| 97 | EMB | 3.27E-15 | 2.3822348 | IGSF21+ Dendritic |
| 98 | TPT1 | 4.57E-15 | 1.290736 | IGSF21+ Dendritic |
| 99 | VSIG4 | 6.89E-15 | -5.110598 | IGSF21+ Dendritic |
| 0 | LILRB2 | 7.45E-98 | 8.514771 | Intermediate Monocyte |
| 1 | FCN1 | 3.48E-92 | 14.220522 | Intermediate Monocyte |
| 2 | COTL1 | 1.84E-89 | 21.447792 | Intermediate Monocyte |
| 3 | FGL2 | 2.64E-88 | 6.528241 | Intermediate Monocyte |
| 4 | CD300E | 2.36E-84 | 7.6250257 | Intermediate Monocyte |
| 5 | LST1 | 1.29E-82 | 21.773472 | Intermediate Monocyte |
| 6 | LILRA5 | 3.19E-79 | 6.3899765 | Intermediate Monocyte |

| | | | | |
|---|---|---|---|---|
| 7 | VCAN | 4.38E-79 | 8.206407 | Intermediate Monocyte |
| 8 | RGS2 | 3.68E-77 | 11.120435 | Intermediate Monocyte |
| 9 | CD48 | 2.89E-75 | 6.1590943 | Intermediate Monocyte |
| 10 | CFP | 1.59E-74 | 6.7563004 | Intermediate Monocyte |
| 11 | FAM26F | 3.44E-73 | 6.6500587 | Intermediate Monocyte |
| 12 | APOBEC3A | 3.17E-72 | 9.645125 | Intermediate Monocyte |
| 13 | TNFRSF1B | 6.18E-69 | 5.195843 | Intermediate Monocyte |
| 14 | RPS26 | 2.33E-68 | 16.762215 | Intermediate Monocyte |
| 15 | MPEG1 | 4.69E-68 | 5.333975 | Intermediate Monocyte |
| 16 | ADGRE2 | 8.62E-66 | 5.9930873 | Intermediate Monocyte |
| 17 | FGR | 8.62E-66 | 4.98779 | Intermediate Monocyte |
| 18 | C10orf54 | 4.30E-64 | 4.7758026 | Intermediate Monocyte |
| 19 | FAM65B | 6.68E-63 | 5.3368053 | Intermediate Monocyte |
| 20 | TIMP1 | 3.32E-62 | 15.725371 | Intermediate Monocyte |
| 21 | CORO1A | 1.17E-61 | 5.138474 | Intermediate Monocyte |
| 22 | IL1B | 2.97E-61 | 5.763205 | Intermediate Monocyte |
| 23 | PRELID1 | 1.99E-60 | 6.5824733 | Intermediate Monocyte |
| 24 | FYB | 5.46E-58 | 4.3053975 | Intermediate Monocyte |
| 25 | RPS24 | 1.09E-57 | 28.866179 | Intermediate Monocyte |
| 26 | NAMPT | 2.12E-57 | 8.106516 | Intermediate Monocyte |
| 27 | SAT1 | 6.01E-57 | 33.66065 | Intermediate Monocyte |
| 28 | LINC01272 | 5.82E-56 | 6.2831445 | Intermediate Monocyte |
| 29 | PLAC8 | 1.44E-55 | 7.168753 | Intermediate Monocyte |
| 30 | MAFB | 8.87E-55 | 4.5380874 | Intermediate Monocyte |
| 31 | SMAP2 | 1.04E-54 | 4.2962465 | Intermediate Monocyte |
| 32 | AIF1 | 2.42E-54 | 11.215567 | Intermediate Monocyte |
| 33 | IFITM2 | 9.53E-54 | 9.704574 | Intermediate Monocyte |
| 34 | PTPRC | 1.30E-53 | 5.377626 | Intermediate Monocyte |
| 35 | RPL39 | 6.75E-51 | 26.967495 | Intermediate Monocyte |
| 36 | CD300A | 1.25E-50 | 4.308177 | Intermediate Monocyte |
| 37 | ATP5E | 1.79E-50 | 10.445787 | Intermediate Monocyte |
| 38 | RPL37 | 1.90E-50 | 19.43937 | Intermediate Monocyte |
| 39 | STK17B | 7.03E-49 | 3.730703 | Intermediate Monocyte |
| 40 | CTSS | 1.72E-48 | 10.357594 | Intermediate Monocyte |
| 41 | THBS1 | 5.04E-48 | 6.694799 | Intermediate Monocyte |
| 42 | RNASET2 | 5.82E-48 | 4.44863 | Intermediate Monocyte |
| 43 | RPS9 | 2.34E-47 | 22.58079 | Intermediate Monocyte |
| 44 | C1orf162 | 6.51E-47 | 5.748711 | Intermediate Monocyte |
| 45 | SOCS3 | 1.31E-46 | 5.5095077 | Intermediate Monocyte |
| 46 | RPL36A | 5.01E-46 | 4.29409 | Intermediate Monocyte |
| 47 | PSAP | 4.05E-45 | 11.219344 | Intermediate Monocyte |
| 48 | SLC25A6 | 8.01E-45 | 7.1669984 | Intermediate Monocyte |
| 49 | PTPRE | 1.18E-44 | 3.6572986 | Intermediate Monocyte |
| 50 | RPL23 | 1.56E-44 | 9.750491 | Intermediate Monocyte |
| 51 | CLEC7A | 5.35E-44 | 3.4584072 | Intermediate Monocyte |
| 52 | FOS | 5.65E-44 | 13.864803 | Intermediate Monocyte |
| 53 | TYMP | 5.65E-44 | 5.235693 | Intermediate Monocyte |
| 54 | RPL34 | 6.83E-44 | 26.534775 | Intermediate Monocyte |
| 55 | AP1S2 | 9.26E-44 | 3.6399012 | Intermediate Monocyte |
| 56 | PABPC1 | 1.08E-43 | 7.606124 | Intermediate Monocyte |
| 57 | RPL26 | 2.82E-43 | 25.056526 | Intermediate Monocyte |
| 58 | RPL38 | 4.25E-43 | 12.230858 | Intermediate Monocyte |
| 59 | CSTA | 7.07E-43 | 3.519522 | Intermediate Monocyte |
| 60 | LILRA1 | 1.93E-42 | 6.2822886 | Intermediate Monocyte |
| 61 | MS4A6A | 3.05E-42 | 4.3621683 | Intermediate Monocyte |
| 62 | TBXAS1 | 3.92E-42 | 3.3689306 | Intermediate Monocyte |
| 63 | TNFSF13B | 5.48E-42 | 3.9186618 | Intermediate Monocyte |
| 64 | NEAT1 | 1.05E-41 | 12.366865 | Intermediate Monocyte |
| 65 | IFITM3 | 2.74E-41 | 10.763588 | Intermediate Monocyte |
| 66 | CPVL | 2.76E-41 | 3.2082286 | Intermediate Monocyte |
| 67 | ABI3 | 3.33E-41 | 3.6377854 | Intermediate Monocyte |
| 68 | CST3 | 4.42E-41 | 16.335152 | Intermediate Monocyte |
| 69 | CD52 | 4.58E-41 | 6.560236 | Intermediate Monocyte |
| 70 | RPS28 | 4.99E-41 | 20.294128 | Intermediate Monocyte |

| | | | | |
|---|---|---|---|---|
| 71 | RPS27 | 1.30E-40 | 30.158949 | Intermediate Monocyte |
| 72 | RPS8 | 2.74E-40 | 17.216923 | Intermediate Monocyte |
| 73 | SLA | 3.04E-40 | 2.8541784 | Intermediate Monocyte |
| 74 | RPS13 | 3.21E-40 | 18.387985 | Intermediate Monocyte |
| 75 | RGS18 | 4.39E-40 | 4.7869754 | Intermediate Monocyte |
| 76 | S100A8 | 5.72E-40 | 12.207426 | Intermediate Monocyte |
| 77 | IFI30 | 6.49E-40 | 1.8072717 | Intermediate Monocyte |
| 78 | LRRC25 | 1.75E-39 | 3.9633274 | Intermediate Monocyte |
| 79 | ITGA4 | 6.57E-39 | 3.717548 | Intermediate Monocyte |
| 80 | RPL28 | 7.07E-39 | 25.780563 | Intermediate Monocyte |
| 81 | FCER1G | 1.51E-38 | 6.338089 | Intermediate Monocyte |
| 82 | RPL18A | 1.59E-38 | 20.287855 | Intermediate Monocyte |
| 83 | CEBPB | 1.61E-38 | 7.106305 | Intermediate Monocyte |
| 84 | RNASE6 | 1.93E-38 | 3.3405743 | Intermediate Monocyte |
| 85 | S100A9 | 2.39E-38 | 11.686819 | Intermediate Monocyte |
| 86 | RPL27 | 2.60E-38 | 11.170972 | Intermediate Monocyte |
| 87 | SRGN | 2.60E-38 | 14.565139 | Intermediate Monocyte |
| 88 | TYROBP | 6.95E-38 | 6.3184633 | Intermediate Monocyte |
| 89 | RPL41 | 5.56E-37 | 38.281563 | Intermediate Monocyte |
| 90 | RPS29 | 6.44E-37 | 19.267048 | Intermediate Monocyte |
| 91 | RAB24 | 6.52E-37 | 3.052843 | Intermediate Monocyte |
| 92 | RPS17 | 7.58E-37 | 14.870589 | Intermediate Monocyte |
| 93 | LIMD2 | 1.04E-36 | 2.973636 | Intermediate Monocyte |
| 94 | RPL37A | 1.15E-36 | 13.794749 | Intermediate Monocyte |
| 95 | LYZ | 1.19E-36 | 1.1467239 | Intermediate Monocyte |
| 96 | RPS12 | 1.51E-36 | 23.97524 | Intermediate Monocyte |
| 97 | RPS19 | 2.31E-36 | 27.279627 | Intermediate Monocyte |
| 98 | CSF1R | 2.47E-36 | 3.5057962 | Intermediate Monocyte |
| 99 | SERPINB9 | 3.06E-36 | 4.233878 | Intermediate Monocyte |
| 0 | ASCL3 | 1.15E-11 | 42.439842 | Ionocyte |
| 1 | MUC20 | 1.15E-11 | 11.804043 | Ionocyte |
| 2 | HMGB3 | 1.15E-11 | 13.853696 | Ionocyte |
| 3 | CD24 | 1.15E-11 | 24.51554 | Ionocyte |
| 4 | EPCAM | 1.15E-11 | 18.19775 | Ionocyte |
| 5 | SEC11C | 2.80E-11 | 23.37156 | Ionocyte |
| 6 | HEPACAM2 | 2.80E-11 | 14.332232 | Ionocyte |
| 7 | CLCNKB | 2.80E-11 | 16.004414 | Ionocyte |
| 8 | TMEM61 | 2.80E-11 | 16.798496 | Ionocyte |
| 9 | TPD52 | 8.71E-11 | 10.4428625 | Ionocyte |
| 10 | MGST1 | 8.71E-11 | 34.29129 | Ionocyte |
| 12 | PERP | 1.06E-10 | 10.22311 | Ionocyte |
| 11 | KRT18 | 1.06E-10 | 20.199097 | Ionocyte |
| 13 | TFF3 | 1.08E-10 | 15.083667 | Ionocyte |
| 14 | GADD45G | 1.44E-10 | 11.991887 | Ionocyte |
| 15 | IDH2 | 2.01E-10 | 6.190326 | Ionocyte |
| 16 | FOXI1 | 2.31E-10 | 18.872124 | Ionocyte |
| 17 | KRT8 | 2.31E-10 | 23.125362 | Ionocyte |
| 18 | ATP6V1B1 | 2.61E-10 | 11.149368 | Ionocyte |
| 19 | TACSTD2 | 2.61E-10 | 9.183426 | Ionocyte |
| 20 | KRT7 | 2.61E-10 | 17.64517 | Ionocyte |
| 21 | PRDX2 | 2.67E-10 | 9.494939 | Ionocyte |
| 22 | IGF1 | 4.73E-10 | 9.602631 | Ionocyte |
| 23 | CD9 | 1.10E-09 | 53.261417 | Ionocyte |
| 25 | CLDN4 | 1.11E-09 | 8.946066 | Ionocyte |
| 24 | MIF | 1.11E-09 | 18.508938 | Ionocyte |
| 26 | ATP1A1 | 1.67E-09 | 13.564177 | Ionocyte |
| 27 | ATP6V0B | 2.11E-09 | 24.913172 | Ionocyte |
| 28 | CLDN25 | 2.11E-09 | 16.280537 | Ionocyte |
| 29 | MAGED2 | 2.75E-09 | 4.9449954 | Ionocyte |
| 30 | H1F0 | 4.16E-09 | 7.828041 | Ionocyte |
| 31 | PPAP2C | 4.16E-09 | 6.034867 | Ionocyte |
| 32 | SCNN1B | 5.61E-09 | 9.846722 | Ionocyte |
| 33 | AKR1B1 | 7.37E-09 | 9.38266 | Ionocyte |
| 34 | CLDN7 | 8.13E-09 | 5.755345 | Ionocyte |

| | | | | |
|---|---|---|---|---|
| 35 | GOLM1 | 8.92E-09 | 9.732624 | Ionocyte |
| 36 | TUSC3 | 9.25E-09 | 5.6288466 | Ionocyte |
| 37 | MARCKSL1 | 1.10E-08 | 8.694752 | Ionocyte |
| 38 | SPINT1 | 1.22E-08 | 4.8257565 | Ionocyte |
| 39 | SPINT2 | 1.44E-08 | 9.455061 | Ionocyte |
| 40 | PFN2 | 1.49E-08 | 7.0127788 | Ionocyte |
| 41 | GNAS | 1.61E-08 | 8.92394 | Ionocyte |
| 42 | SOX4 | 1.61E-08 | 7.472312 | Ionocyte |
| 43 | H3F3B | 1.61E-08 | 33.40797 | Ionocyte |
| 44 | ATP6V1G3 | 1.79E-08 | 16.820654 | Ionocyte |
| 45 | ST14 | 2.36E-08 | 6.524843 | Ionocyte |
| 46 | HEBP2 | 2.36E-08 | 4.6902995 | Ionocyte |
| 47 | KRT19 | 2.45E-08 | 17.654985 | Ionocyte |
| 48 | CIRBP | 2.52E-08 | 8.882271 | Ionocyte |
| 49 | NGRN | 2.59E-08 | 5.809719 | Ionocyte |
| 50 | HSPB1 | 3.02E-08 | 21.03605 | Ionocyte |
| 51 | RASSF6 | 3.08E-08 | 7.920132 | Ionocyte |
| 52 | C9orf152 | 3.49E-08 | 6.809138 | Ionocyte |
| 53 | HES6 | 3.49E-08 | 12.95744 | Ionocyte |
| 54 | RARRES2 | 3.56E-08 | 17.2328 | Ionocyte |
| 55 | ATP5B | 3.88E-08 | 10.058178 | Ionocyte |
| 56 | FAM3B | 3.88E-08 | 5.4538045 | Ionocyte |
| 57 | ATP1B1 | 4.21E-08 | 9.314454 | Ionocyte |
| 58 | PHB | 4.27E-08 | 5.3233657 | Ionocyte |
| 59 | ATP6V1A | 4.53E-08 | 5.2889795 | Ionocyte |
| 60 | HIGD1A | 4.69E-08 | 4.8587465 | Ionocyte |
| 61 | PHLDA1 | 5.20E-08 | 5.2842736 | Ionocyte |
| 62 | SLC25A6 | 6.54E-08 | 11.028197 | Ionocyte |
| 63 | DST | 7.02E-08 | 4.8887997 | Ionocyte |
| 64 | NCALD | 1.17E-07 | 5.9745274 | Ionocyte |
| 65 | CLCNKA | 1.40E-07 | 11.705185 | Ionocyte |
| 66 | TMPRSS11E | 1.40E-07 | 12.720165 | Ionocyte |
| 67 | CEL | 1.40E-07 | 18.514551 | Ionocyte |
| 68 | DMRT2 | 1.52E-07 | 10.328604 | Ionocyte |
| 69 | PPDPF | 1.67E-07 | 18.590153 | Ionocyte |
| 70 | LDHB | 1.67E-07 | 8.910357 | Ionocyte |
| 71 | FOXP1 | 1.67E-07 | 4.7741947 | Ionocyte |
| 72 | CLDN3 | 1.73E-07 | 7.4892488 | Ionocyte |
| 76 | ATP6V1G1 | 1.91E-07 | 7.853778 | Ionocyte |
| 75 | CNN3 | 1.91E-07 | 5.8397255 | Ionocyte |
| 74 | AP1M2 | 1.91E-07 | 4.7967634 | Ionocyte |
| 73 | CFTR | 1.91E-07 | 8.630673 | Ionocyte |
| 77 | NEURL1 | 1.94E-07 | 7.8013954 | Ionocyte |
| 78 | ATP6V0C | 1.96E-07 | 14.481431 | Ionocyte |
| 79 | AMN | 2.07E-07 | 7.8377967 | Ionocyte |
| 80 | SLC25A11 | 2.39E-07 | 3.9709966 | Ionocyte |
| 81 | SRM | 2.43E-07 | 6.047997 | Ionocyte |
| 82 | ITPR2 | 2.75E-07 | 5.76369 | Ionocyte |
| 83 | APLP2 | 2.86E-07 | 9.26509 | Ionocyte |
| 84 | COX5B | 2.86E-07 | 9.995678 | Ionocyte |
| 85 | LAPTM4B | 3.23E-07 | 4.629091 | Ionocyte |
| 87 | POSTN | 3.25E-07 | 9.31627 | Ionocyte |
| 86 | CENPW | 3.25E-07 | 5.447955 | Ionocyte |
| 88 | SFTPC | 3.33E-07 | 91.639046 | Ionocyte |
| 89 | BIK | 3.48E-07 | 7.800384 | Ionocyte |
| 90 | SYNE2 | 3.48E-07 | 4.0007677 | Ionocyte |
| 91 | SEMA3C | 3.48E-07 | 4.882515 | Ionocyte |
| 92 | CUTA | 3.48E-07 | 5.8389006 | Ionocyte |
| 93 | GSN | 3.57E-07 | 12.479088 | Ionocyte |
| 94 | PDHB | 3.75E-07 | 4.527246 | Ionocyte |
| 95 | ABHD11 | 3.81E-07 | 4.680389 | Ionocyte |
| 96 | GAPDH | 4.22E-07 | 25.876106 | Ionocyte |
| 97 | HMGN1 | 4.24E-07 | 6.186044 | Ionocyte |
| 98 | UPF3A | 4.81E-07 | 4.7046437 | Ionocyte |

| | | | | |
|---|---|---|---|---|
| 99 | SRP9 | 4.81E-07 | 4.516163 | Ionocyte |
| 0 | COL1A1 | 2.58E-20 | 54.40987 | Lipofibroblast |
| 1 | COL3A1 | 2.58E-20 | 26.083845 | Lipofibroblast |
| 2 | COL1A2 | 2.58E-20 | 38.64952 | Lipofibroblast |
| 3 | CRISPLD2 | 3.33E-20 | 10.796258 | Lipofibroblast |
| 4 | C3 | 3.33E-20 | 27.328465 | Lipofibroblast |
| 5 | SERPINF1 | 1.21E-19 | 21.72535 | Lipofibroblast |
| 6 | TIMP1 | 1.49E-19 | 92.19098 | Lipofibroblast |
| 7 | MT2A | 1.65E-19 | 458.85797 | Lipofibroblast |
| 8 | CCDC80 | 3.26E-19 | 12.94809 | Lipofibroblast |
| 9 | C1S | 3.46E-19 | 22.607355 | Lipofibroblast |
| 10 | COL6A2 | 4.63E-19 | 20.929302 | Lipofibroblast |
| 11 | PTGDS | 5.24E-19 | 24.603731 | Lipofibroblast |
| 12 | FBLN1 | 5.58E-19 | 20.909431 | Lipofibroblast |
| 13 | C1R | 7.88E-19 | 19.247923 | Lipofibroblast |
| 14 | NNMT | 7.88E-19 | 23.9872 | Lipofibroblast |
| 15 | C7 | 1.83E-18 | 24.35853 | Lipofibroblast |
| 16 | DCN | 1.92E-18 | 42.909847 | Lipofibroblast |
| 17 | ALDH1A3 | 1.92E-18 | 10.079624 | Lipofibroblast |
| 18 | COL6A3 | 2.07E-18 | 12.732666 | Lipofibroblast |
| 19 | RARRES2 | 2.07E-18 | 8.79485 | Lipofibroblast |
| 20 | PCOLCE | 2.14E-18 | 14.846378 | Lipofibroblast |
| 21 | ADAMTS4 | 2.43E-18 | 10.45014 | Lipofibroblast |
| 22 | ARID5B | 3.29E-18 | 11.756275 | Lipofibroblast |
| 23 | EFEMP1 | 3.29E-18 | 8.69355 | Lipofibroblast |
| 25 | CREM | 4.03E-18 | 16.174973 | Lipofibroblast |
| 24 | COL6A1 | 4.03E-18 | 14.520061 | Lipofibroblast |
| 26 | TFPI2 | 4.59E-18 | 12.775619 | Lipofibroblast |
| 27 | JUNB | 5.54E-18 | 69.941315 | Lipofibroblast |
| 28 | PHLDA1 | 6.95E-18 | 21.244799 | Lipofibroblast |
| 29 | MGP | 7.34E-18 | 31.444557 | Lipofibroblast |
| 30 | MYC | 1.13E-17 | 21.320086 | Lipofibroblast |
| 31 | MAT2A | 1.23E-17 | 13.42485 | Lipofibroblast |
| 32 | IGFBP4 | 1.46E-17 | 11.431482 | Lipofibroblast |
| 33 | RARRES1 | 4.39E-17 | 12.208514 | Lipofibroblast |
| 34 | MT1X | 4.39E-17 | 100.534904 | Lipofibroblast |
| 35 | SRPX | 1.46E-16 | 11.002286 | Lipofibroblast |
| 36 | DPT | 1.50E-16 | 15.018821 | Lipofibroblast |
| 37 | EGR1 | 1.54E-16 | 11.217477 | Lipofibroblast |
| 38 | RGS2 | 1.74E-16 | 20.553705 | Lipofibroblast |
| 39 | SPARCL1 | 2.19E-16 | 17.05723 | Lipofibroblast |
| 40 | NR4A2 | 2.26E-16 | 8.566078 | Lipofibroblast |
| 41 | MMP2 | 7.11E-16 | 10.215284 | Lipofibroblast |
| 42 | OLFML3 | 7.11E-16 | 7.8998213 | Lipofibroblast |
| 43 | PNRC1 | 1.13E-15 | 19.992235 | Lipofibroblast |
| 44 | FST | 1.21E-15 | 16.51819 | Lipofibroblast |
| 45 | HAS1 | 1.44E-15 | 16.388346 | Lipofibroblast |
| 46 | GFPT2 | 1.74E-15 | 8.943106 | Lipofibroblast |
| 47 | MT1M | 1.74E-15 | 30.914862 | Lipofibroblast |
| 48 | IFITM1 | 2.10E-15 | 17.681446 | Lipofibroblast |
| 49 | NAMPT | 4.42E-15 | 12.725204 | Lipofibroblast |
| 50 | EMILIN1 | 6.71E-15 | 7.814979 | Lipofibroblast |
| 51 | SOCS3 | 1.27E-14 | 15.308657 | Lipofibroblast |
| 52 | MCL1 | 1.27E-14 | 21.389803 | Lipofibroblast |
| 53 | SEPP1 | 1.46E-14 | 9.190139 | Lipofibroblast |
| 54 | MEDAG | 1.69E-14 | 10.72882 | Lipofibroblast |
| 55 | PTN | 2.12E-14 | 8.876713 | Lipofibroblast |
| 56 | ZFP36L1 | 4.17E-14 | 14.115853 | Lipofibroblast |
| 57 | APOE | 6.50E-14 | 39.850834 | Lipofibroblast |
| 58 | THBS1 | 7.02E-14 | 9.642715 | Lipofibroblast |
| 59 | CALD1 | 1.24E-13 | 7.416907 | Lipofibroblast |
| 60 | HAS2 | 1.89E-13 | 9.472889 | Lipofibroblast |
| 61 | MT1A | 2.24E-13 | 18.030552 | Lipofibroblast |
| 62 | BGN | 3.47E-13 | 4.934236 | Lipofibroblast |

| | | | | |
|---|---|---|---|---|
| 63 | IFITM3 | 3.47E-13 | 21.004799 | Lipofibroblast |
| 64 | TMEM176B | 3.70E-13 | 5.7258773 | Lipofibroblast |
| 65 | MARCKSL1 | 3.88E-13 | 5.6052504 | Lipofibroblast |
| 66 | APOD | 9.18E-13 | 6.504963 | Lipofibroblast |
| 67 | IGFBP7 | 9.45E-13 | 17.8115 | Lipofibroblast |
| 68 | SPARC | 1.44E-12 | 10.533995 | Lipofibroblast |
| 69 | VEGFA | 1.46E-12 | 5.3632994 | Lipofibroblast |
| 70 | SPTSSA | 1.55E-12 | 5.0968943 | Lipofibroblast |
| 71 | MLLT11 | 1.57E-12 | 10.874473 | Lipofibroblast |
| 72 | ELL2 | 2.82E-12 | 5.913014 | Lipofibroblast |
| 73 | DUSP1 | 3.32E-12 | 44.43988 | Lipofibroblast |
| 74 | NFIL3 | 5.76E-12 | 5.8077807 | Lipofibroblast |
| 75 | CYP1B1 | 1.33E-11 | 11.4823675 | Lipofibroblast |
| 76 | FSTL1 | 1.69E-11 | 4.9945803 | Lipofibroblast |
| 77 | GEM | 1.71E-11 | 7.8319106 | Lipofibroblast |
| 78 | CD248 | 1.95E-11 | 8.484292 | Lipofibroblast |
| 79 | ADAMTS1 | 2.10E-11 | 9.712323 | Lipofibroblast |
| 80 | FGF7 | 2.32E-11 | 16.7573 | Lipofibroblast |
| 81 | GAS1 | 2.42E-11 | 9.0405855 | Lipofibroblast |
| 82 | B4GALT1 | 4.34E-11 | 6.964384 | Lipofibroblast |
| 83 | CEBPB | 4.70E-11 | 19.649216 | Lipofibroblast |
| 84 | CEBPD | 5.51E-11 | 15.231867 | Lipofibroblast |
| 85 | PDLIM4 | 5.65E-11 | 6.3319926 | Lipofibroblast |
| 86 | CXCL12 | 5.87E-11 | 9.2696 | Lipofibroblast |
| 87 | TIPARP | 6.61E-11 | 4.9346585 | Lipofibroblast |
| 88 | WT1 | 8.08E-11 | 10.836349 | Lipofibroblast |
| 89 | C11orf96 | 1.16E-10 | 17.692547 | Lipofibroblast |
| 90 | PLA2G2A | 1.20E-10 | 28.115595 | Lipofibroblast |
| 91 | DDX21 | 1.37E-10 | 4.8976283 | Lipofibroblast |
| 92 | CDKN1A | 1.94E-10 | 6.3717074 | Lipofibroblast |
| 93 | STEAP1 | 2.00E-10 | 6.666526 | Lipofibroblast |
| 94 | CCL2 | 2.02E-10 | 14.891474 | Lipofibroblast |
| 95 | NR4A1 | 2.02E-10 | 5.911422 | Lipofibroblast |
| 96 | PLTP | 2.46E-10 | 5.3124332 | Lipofibroblast |
| 97 | CFI | 2.90E-10 | 6.4580164 | Lipofibroblast |
| 98 | UGCG | 5.48E-10 | 4.4316955 | Lipofibroblast |
| 99 | FGFR1 | 7.06E-10 | 5.4534693 | Lipofibroblast |
| 0 | CCL21 | 1.61E-294 | 220.70868 | Lymphatic |
| 1 | TFF3 | 1.02E-283 | 23.987839 | Lymphatic |
| 2 | IGFBP7 | 3.36E-270 | 62.841244 | Lymphatic |
| 3 | MMRN1 | 5.36E-251 | 11.811371 | Lymphatic |
| 4 | TFPI | 4.71E-224 | 10.194353 | Lymphatic |
| 5 | GNG11 | 1.37E-221 | 16.592602 | Lymphatic |
| 6 | ECSCR | 3.66E-208 | 9.306371 | Lymphatic |
| 7 | PPFIBP1 | 6.47E-187 | 6.74984 | Lymphatic |
| 8 | SDPR | 1.19E-182 | 14.199187 | Lymphatic |
| 9 | NNMT | 3.46E-176 | 10.03296 | Lymphatic |
| 10 | SNCG | 2.45E-166 | 8.143575 | Lymphatic |
| 11 | AKAP12 | 7.69E-162 | 11.044145 | Lymphatic |
| 12 | APP | 8.12E-161 | 5.652345 | Lymphatic |
| 13 | MARCKSL1 | 1.24E-150 | 5.400104 | Lymphatic |
| 14 | CLDN5 | 1.19E-149 | 15.230594 | Lymphatic |
| 15 | ADIRF | 1.21E-147 | 14.304466 | Lymphatic |
| 16 | PRSS23 | 8.23E-145 | 5.5715055 | Lymphatic |
| 17 | PDPN | 1.78E-143 | 8.48373 | Lymphatic |
| 18 | CRIP2 | 1.61E-141 | 5.3769083 | Lymphatic |
| 19 | CLU | 1.27E-135 | 12.041114 | Lymphatic |
| 20 | FSCN1 | 2.91E-134 | 6.385429 | Lymphatic |
| 21 | FXYD6 | 1.08E-131 | 4.937822 | Lymphatic |
| 22 | RAMP2 | 1.08E-130 | 7.2170014 | Lymphatic |
| 23 | IGFBP4 | 1.50E-126 | 5.3304014 | Lymphatic |
| 24 | HYAL2 | 1.18E-124 | 5.76495 | Lymphatic |
| 25 | TM4SF1 | 3.35E-122 | 10.171267 | Lymphatic |
| 26 | PDLIM4 | 1.66E-119 | 5.3802447 | Lymphatic |

| | | | | |
|---|---|---|---|---|
| 27 | CNN3 | 6.29E-119 | 4.8252416 | Lymphatic |
| 28 | KANK3 | 1.22E-117 | 4.3875055 | Lymphatic |
| 29 | THBD | 5.74E-117 | 5.7595177 | Lymphatic |
| 30 | RGS16 | 1.30E-114 | 6.035067 | Lymphatic |
| 31 | EFEMP1 | 4.70E-111 | 7.2029667 | Lymphatic |
| 32 | ELK3 | 1.64E-109 | 4.2625737 | Lymphatic |
| 33 | CALD1 | 7.77E-108 | 2.850444 | Lymphatic |
| 34 | CD59 | 1.17E-107 | 9.714766 | Lymphatic |
| 35 | CAV1 | 3.72E-106 | 8.591575 | Lymphatic |
| 36 | TIMP3 | 2.62E-96 | 3.9749122 | Lymphatic |
| 37 | PROX1 | 6.64E-95 | 8.890895 | Lymphatic |
| 38 | PCAT19 | 3.33E-94 | 4.4460654 | Lymphatic |
| 39 | EMP1 | 2.71E-93 | 5.216862 | Lymphatic |
| 40 | IGF1 | 4.81E-93 | 5.554784 | Lymphatic |
| 41 | RAB11A | 2.77E-91 | 4.138971 | Lymphatic |
| 42 | LAYN | 8.80E-91 | 5.2100124 | Lymphatic |
| 43 | APOLD1 | 2.39E-90 | 5.261721 | Lymphatic |
| 44 | TUBB6 | 8.42E-90 | 3.7600281 | Lymphatic |
| 45 | CD9 | 1.10E-87 | 9.1388645 | Lymphatic |
| 46 | SPHK1 | 2.75E-87 | 4.7407255 | Lymphatic |
| 47 | LYVE1 | 2.82E-85 | 9.290654 | Lymphatic |
| 48 | TM4SF18 | 6.75E-85 | 6.2990594 | Lymphatic |
| 49 | PPP1R2 | 1.01E-84 | 4.5082498 | Lymphatic |
| 50 | ARL4A | 6.25E-84 | 3.3599434 | Lymphatic |
| 51 | GYPC | 2.96E-83 | 4.6600547 | Lymphatic |
| 52 | STMN1 | 4.00E-83 | 2.9142487 | Lymphatic |
| 53 | TIE1 | 8.90E-83 | 3.6531339 | Lymphatic |
| 54 | PMP22 | 6.10E-82 | 4.2018266 | Lymphatic |
| 55 | RHOC | 2.99E-81 | 4.1127295 | Lymphatic |
| 56 | LRRC70 | 6.68E-81 | 6.6748033 | Lymphatic |
| 57 | TSHZ2 | 1.06E-80 | 4.891535 | Lymphatic |
| 58 | HSPB1 | 2.52E-79 | 5.983534 | Lymphatic |
| 59 | OAF | 7.53E-79 | 5.882395 | Lymphatic |
| 60 | CAV2 | 2.20E-75 | 3.009869 | Lymphatic |
| 61 | SEPP1 | 3.65E-75 | 2.06886 | Lymphatic |
| 62 | LAMA4 | 2.10E-74 | 4.5861206 | Lymphatic |
| 63 | SHC1 | 2.07E-72 | 3.4880717 | Lymphatic |
| 64 | MTUS1 | 7.44E-71 | 3.7917578 | Lymphatic |
| 65 | HTRA1 | 2.66E-70 | 4.3360405 | Lymphatic |
| 66 | RASIP1 | 2.00E-69 | 3.1367416 | Lymphatic |
| 67 | CXorf36 | 7.18E-69 | 4.926986 | Lymphatic |
| 68 | TIMP1 | 9.55E-69 | 13.374035 | Lymphatic |
| 69 | IFITM3 | 3.31E-68 | 7.8582835 | Lymphatic |
| 70 | MGST2 | 6.10E-68 | 2.653874 | Lymphatic |
| 71 | PTRF | 2.94E-66 | 2.538085 | Lymphatic |
| 72 | SMAD1 | 4.73E-66 | 4.293593 | Lymphatic |
| 73 | NRP2 | 5.12E-66 | 3.5902753 | Lymphatic |
| 74 | NDRG1 | 5.81E-66 | 3.0464377 | Lymphatic |
| 75 | SYPL1 | 8.37E-66 | 2.5629895 | Lymphatic |
| 76 | DBN1 | 6.92E-65 | 4.5619755 | Lymphatic |
| 77 | TCF4 | 1.39E-64 | 2.9678113 | Lymphatic |
| 78 | EGFL7 | 1.48E-64 | 2.5494182 | Lymphatic |
| 79 | FAM213A | 2.86E-64 | 3.1606677 | Lymphatic |
| 80 | MYC | 5.43E-64 | 3.2680779 | Lymphatic |
| 81 | TSTA3 | 8.18E-64 | 3.0599556 | Lymphatic |
| 82 | NRN1 | 2.22E-63 | 3.6214721 | Lymphatic |
| 83 | TSPAN4 | 6.82E-63 | 2.4389596 | Lymphatic |
| 84 | LIMS1 | 6.84E-63 | 2.8906634 | Lymphatic |
| 85 | ANXA2 | 1.60E-62 | 0.9318018 | Lymphatic |
| 86 | ARID5B | 5.76E-62 | 3.1141996 | Lymphatic |
| 87 | LOC643733 | 1.39E-61 | 5.2867794 | Lymphatic |
| 88 | GPM6A | 3.96E-61 | 5.7088995 | Lymphatic |
| 89 | HOXD8 | 2.08E-60 | 9.288928 | Lymphatic |
| 90 | C16orf62 | 3.06E-60 | 3.9192255 | Lymphatic |

| | | | | |
|---|---|---|---|---|
| 91 | PPAP2A | 3.16E-60 | 3.853738 | Lymphatic |
| 92 | FLT4 | 1.07E-58 | 5.1138115 | Lymphatic |
| 93 | S100A16 | 6.93E-57 | 2.1035516 | Lymphatic |
| 94 | CCDC80 | 7.83E-57 | 2.533768 | Lymphatic |
| 95 | HEBP1 | 1.18E-55 | 2.4502478 | Lymphatic |
| 96 | WFS1 | 1.87E-55 | 3.8752422 | Lymphatic |
| 97 | ADCY4 | 1.95E-55 | 3.3132923 | Lymphatic |
| 98 | EPHX1 | 2.89E-54 | 2.2947977 | Lymphatic |
| 99 | ACTN1 | 3.56E-54 | 1.9345981 | Lymphatic |
| 0 | APOC1 | 0 | 181.43411 | Macrophage |
| 72 | GLRX | 0 | 16.478369 | Macrophage |
| 71 | ALOX5 | 0 | 12.747852 | Macrophage |
| 70 | GLUL | 0 | 31.159958 | Macrophage |
| 69 | LGALS1 | 0 | 46.34331 | Macrophage |
| 68 | LST1 | 0 | 17.59231 | Macrophage |
| 67 | HLA-DPA1 | 0 | 95.89362 | Macrophage |
| 66 | S100A10 | 0 | 71.147644 | Macrophage |
| 65 | HLA-DRB5 | 0 | 66.62494 | Macrophage |
| 64 | TXN | 0 | 34.98885 | Macrophage |
| 63 | CST3 | 0 | 83.0506 | Macrophage |
| 62 | HLA-DRB1 | 0 | 184.71588 | Macrophage |
| 61 | APOE | 0 | 52.225338 | Macrophage |
| 60 | ANXA5 | 0 | 30.863787 | Macrophage |
| 59 | VIM | 0 | 131.81345 | Macrophage |
| 58 | CSTB | 0 | 41.08133 | Macrophage |
| 57 | CD52 | 0 | 72.838806 | Macrophage |
| 56 | GCHFR | 0 | 23.68359 | Macrophage |
| 55 | GSTO1 | 0 | 26.243944 | Macrophage |
| 54 | STXBP2 | 0 | 24.692036 | Macrophage |
| 53 | HLA-DPB1 | 0 | 77.80055 | Macrophage |
| 52 | HLA-DQA1 | 0 | 39.754677 | Macrophage |
| 73 | GPNMB | 0 | 13.61617 | Macrophage |
| 51 | AIF1 | 0 | 24.518702 | Macrophage |
| 74 | NOP10 | 0 | 16.638098 | Macrophage |
| 76 | MS4A4A | 0 | 14.393413 | Macrophage |
| 97 | SLC11A1 | 0 | 9.309889 | Macrophage |
| 96 | FCGR3A | 0 | 17.785234 | Macrophage |
| 95 | ATP6V0C | 0 | 20.702892 | Macrophage |
| 94 | CFD | 0 | 25.704237 | Macrophage |
| 93 | MYL6 | 0 | 85.026505 | Macrophage |
| 92 | CYP27A1 | 0 | 14.767652 | Macrophage |
| 91 | ATP6V0B | 0 | 18.282824 | Macrophage |
| 90 | TYMP | 0 | 16.425726 | Macrophage |
| 89 | VAMP8 | 0 | 21.49398 | Macrophage |
| 88 | CTSL | 0 | 19.12374 | Macrophage |
| 87 | PYCARD | 0 | 13.441596 | Macrophage |
| 86 | YBX1 | 0 | 39.33592 | Macrophage |
| 85 | CYBB | 0 | 9.763806 | Macrophage |
| 84 | HLA-DMA | 0 | 21.60556 | Macrophage |
| 83 | MGST3 | 0 | 27.383127 | Macrophage |
| 82 | HLA-DRB6 | 0 | 57.74248 | Macrophage |
| 81 | FABP5 | 0 | 30.445318 | Macrophage |
| 80 | CSTA | 0 | 14.876277 | Macrophage |
| 79 | LTA4H | 0 | 17.582754 | Macrophage |
| 78 | ATP6V1F | 0 | 17.967485 | Macrophage |
| 77 | GPX1 | 0 | 30.917238 | Macrophage |
| 75 | PFN1 | 0 | 68.59967 | Macrophage |
| 50 | CXCL16 | 0 | 14.81682 | Macrophage |
| 49 | CD74 | 0 | 380.97607 | Macrophage |
| 48 | OAZ1 | 0 | 72.11069 | Macrophage |
| 21 | MRC1 | 0 | 16.494158 | Macrophage |
| 20 | OLR1 | 0 | 17.258327 | Macrophage |
| 19 | CRIP1 | 0 | 166.62836 | Macrophage |
| 18 | MS4A7 | 0 | 22.429052 | Macrophage |

| | | | | |
|---|---|---|---|---|
| 17 | MSR1 | 0 | 17.856321 | Macrophage |
| 16 | HLA-DRA | 0 | 376.54358 | Macrophage |
| 15 | FABP4 | 0 | 162.16832 | Macrophage |
| 14 | GRN | 0 | 75.8091 | Macrophage |
| 13 | MCEMP1 | 0 | 35.238884 | Macrophage |
| 12 | CD68 | 0 | 42.144363 | Macrophage |
| 11 | FCER1G | 0 | 78.404816 | Macrophage |
| 10 | ACP5 | 0 | 33.67315 | Macrophage |
| 9 | IFI30 | 0 | 90.07895 | Macrophage |
| 8 | TYROBP | 0 | 103.15122 | Macrophage |
| 7 | MARCO | 0 | 57.141792 | Macrophage |
| 6 | C1QC | 0 | 87.41689 | Macrophage |
| 5 | C1QB | 0 | 196.10213 | Macrophage |
| 4 | FTH1 | 0 | inf | Macrophage |
| 3 | CTSD | 0 | 83.16369 | Macrophage |
| 2 | C1QA | 0 | 183.6842 | Macrophage |
| 1 | FTL | 0 | inf | Macrophage |
| 22 | S100A11 | 0 | 142.55672 | Macrophage |
| 23 | ALOX5AP | 0 | 51.841934 | Macrophage |
| 24 | LYZ | 0 | 169.34032 | Macrophage |
| 25 | GLIPR2 | 0 | 18.232174 | Macrophage |
| 47 | LAPTM5 | 0 | 23.340717 | Macrophage |
| 46 | SH3BGRL3 | 0 | 53.91654 | Macrophage |
| 45 | TREM1 | 0 | 19.26951 | Macrophage |
| 44 | HLA-DQB1 | 0 | 42.45608 | Macrophage |
| 43 | CTSC | 0 | 34.00143 | Macrophage |
| 42 | CTSZ | 0 | 16.959024 | Macrophage |
| 41 | C1orf162 | 0 | 17.708559 | Macrophage |
| 40 | SERPINA1 | 0 | 58.805984 | Macrophage |
| 39 | CTSS | 0 | 40.853725 | Macrophage |
| 38 | CTSB | 0 | 25.808493 | Macrophage |
| 98 | ARPC1B | 0 | 22.733177 | Macrophage |
| 37 | SNX10 | 0 | 16.43214 | Macrophage |
| 35 | CYBA | 0 | 80.18846 | Macrophage |
| 34 | TSPO | 0 | 50.11005 | Macrophage |
| 33 | LGALS3 | 0 | 90.15527 | Macrophage |
| 32 | S100A4 | 0 | 140.63733 | Macrophage |
| 31 | CAPG | 0 | 31.460281 | Macrophage |
| 30 | PSAP | 0 | 59.329266 | Macrophage |
| 29 | ALDH2 | 0 | 45.85776 | Macrophage |
| 28 | SPI1 | 0 | 22.266829 | Macrophage |
| 27 | VSIG4 | 0 | 31.8928 | Macrophage |
| 26 | FBP1 | 0 | 35.615395 | Macrophage |
| 36 | TMSB4X | 0 | 822.6482 | Macrophage |
| 99 | ARPC3 | 0 | 29.488897 | Macrophage |
| 0 | TIMP1 | 4.47E-16 | 271.68515 | Mesothelial |
| 1 | PDPN | 4.47E-16 | 9.615606 | Mesothelial |
| 2 | CFB | 5.04E-16 | 42.945312 | Mesothelial |
| 3 | NNMT | 1.66E-15 | 49.277077 | Mesothelial |
| 4 | C1S | 4.06E-15 | 27.147736 | Mesothelial |
| 5 | TFPI2 | 5.50E-15 | 18.793156 | Mesothelial |
| 6 | C1R | 5.50E-15 | 22.327246 | Mesothelial |
| 7 | C3 | 5.50E-15 | 63.0861 | Mesothelial |
| 8 | CFI | 5.50E-15 | 11.492811 | Mesothelial |
| 9 | CCL2 | 5.77E-15 | 82.076675 | Mesothelial |
| 10 | MT2A | 5.77E-15 | 462.64264 | Mesothelial |
| 12 | MT1E | 6.63E-15 | 93.16288 | Mesothelial |
| 11 | RARRES1 | 6.63E-15 | 23.894346 | Mesothelial |
| 13 | KRT18 | 6.88E-15 | 87.58626 | Mesothelial |
| 14 | ARID5B | 6.88E-15 | 20.36807 | Mesothelial |
| 15 | PLA2G2A | 7.57E-15 | 101.03218 | Mesothelial |
| 16 | MIF | 7.77E-15 | 54.225815 | Mesothelial |
| 17 | ALDH1A3 | 8.88E-15 | 21.628828 | Mesothelial |
| 18 | CALB2 | 8.88E-15 | 18.49638 | Mesothelial |

| | | | | |
|---|---|---|---|---|
| 19 | COL1A1 | 1.26E-14 | 52.01549 | Mesothelial |
| 20 | COL1A2 | 2.92E-14 | 35.43816 | Mesothelial |
| 21 | COL6A1 | 3.30E-14 | 13.200472 | Mesothelial |
| 22 | COL6A2 | 4.18E-14 | 20.697453 | Mesothelial |
| 23 | EFEMP1 | 5.68E-14 | 12.852804 | Mesothelial |
| 24 | MT1X | 6.36E-14 | 113.16163 | Mesothelial |
| 25 | RARRES2 | 6.78E-14 | 22.391926 | Mesothelial |
| 26 | RDH10 | 6.88E-14 | 9.029666 | Mesothelial |
| 27 | SULF1 | 9.36E-14 | 9.783816 | Mesothelial |
| 28 | COL3A1 | 9.38E-14 | 39.15887 | Mesothelial |
| 29 | CCDC80 | 1.37E-13 | 31.203674 | Mesothelial |
| 30 | IFITM3 | 1.52E-13 | 42.529182 | Mesothelial |
| 31 | RPS4Y1 | 1.80E-13 | 22.368803 | Mesothelial |
| 32 | RPL39 | 2.00E-13 | 126.61997 | Mesothelial |
| 33 | IL6ST | 2.09E-13 | 8.343727 | Mesothelial |
| 34 | ZFAS1 | 2.21E-13 | 18.903479 | Mesothelial |
| 35 | RPL36 | 2.21E-13 | 91.87913 | Mesothelial |
| 36 | RPL17 | 2.89E-13 | 45.111443 | Mesothelial |
| 37 | RPS12 | 2.93E-13 | 167.79718 | Mesothelial |
| 38 | TMEM98 | 3.51E-13 | 10.252428 | Mesothelial |
| 39 | IGFBP6 | 3.51E-13 | 20.479156 | Mesothelial |
| 40 | TNFRSF12A | 4.18E-13 | 14.552417 | Mesothelial |
| 41 | GAS5 | 4.45E-13 | 22.913412 | Mesothelial |
| 42 | DCBLD2 | 4.78E-13 | 6.798636 | Mesothelial |
| 43 | RPL34 | 4.92E-13 | 152.5278 | Mesothelial |
| 44 | RPS19 | 4.92E-13 | 141.41127 | Mesothelial |
| 45 | KRT8 | 5.14E-13 | 39.207138 | Mesothelial |
| 46 | LRRC75A-AS1 | 5.14E-13 | 44.158424 | Mesothelial |
| 47 | PRDX6 | 5.94E-13 | 12.326421 | Mesothelial |
| 48 | PRG4 | 5.96E-13 | 22.093252 | Mesothelial |
| 50 | RPS8 | 6.14E-13 | 95.404785 | Mesothelial |
| 49 | HAS1 | 6.14E-13 | 23.421541 | Mesothelial |
| 51 | GFPT2 | 6.99E-13 | 10.088809 | Mesothelial |
| 52 | LXN | 9.19E-13 | 10.52714 | Mesothelial |
| 53 | RPS18 | 1.00E-12 | 191.98085 | Mesothelial |
| 54 | RPL10A | 1.17E-12 | 70.247055 | Mesothelial |
| 55 | WBP5 | 1.30E-12 | 9.096134 | Mesothelial |
| 56 | ERRFI1 | 1.32E-12 | 10.8800125 | Mesothelial |
| 57 | MGP | 1.52E-12 | 23.115328 | Mesothelial |
| 58 | RPL35 | 2.03E-12 | 71.34773 | Mesothelial |
| 59 | SPARC | 2.03E-12 | 42.04597 | Mesothelial |
| 60 | RPS17 | 2.03E-12 | 77.44242 | Mesothelial |
| 61 | RPL41 | 2.03E-12 | 250.20091 | Mesothelial |
| 63 | RPLP0 | 2.28E-12 | 63.781357 | Mesothelial |
| 62 | RPL36A | 2.28E-12 | 14.181487 | Mesothelial |
| 64 | TPM2 | 2.30E-12 | 11.227269 | Mesothelial |
| 65 | KRT19 | 2.32E-12 | 23.803938 | Mesothelial |
| 66 | C12orf57 | 2.74E-12 | 8.988134 | Mesothelial |
| 67 | PCOLCE | 3.13E-12 | 13.736133 | Mesothelial |
| 68 | RPL32 | 3.13E-12 | 122.36878 | Mesothelial |
| 69 | RPL13A | 3.19E-12 | 175.13051 | Mesothelial |
| 70 | RPL37 | 3.23E-12 | 76.95879 | Mesothelial |
| 71 | RPL7 | 3.46E-12 | 118.461555 | Mesothelial |
| 72 | TM4SF1 | 3.61E-12 | 33.9076 | Mesothelial |
| 73 | SLC25A6 | 3.61E-12 | 23.494694 | Mesothelial |
| 74 | RPL12 | 3.73E-12 | 122.61962 | Mesothelial |
| 75 | HIF1A | 3.89E-12 | 9.307419 | Mesothelial |
| 76 | PTMA | 3.96E-12 | 102.78456 | Mesothelial |
| 77 | CFH | 3.96E-12 | 10.298989 | Mesothelial |
| 78 | LGALS1 | 4.98E-12 | 134.11708 | Mesothelial |
| 79 | RPL5 | 4.98E-12 | 50.5004 | Mesothelial |
| 80 | RPL13 | 5.05E-12 | 169.52805 | Mesothelial |
| 81 | POLR2L | 5.17E-12 | 17.490963 | Mesothelial |
| 82 | GGCT | 5.32E-12 | 6.9924216 | Mesothelial |

| | | | | |
|---|---|---|---|---|
| 83 | GAS1 | 5.71E-12 | 15.687601 | Mesothelial |
| 84 | RPL10 | 5.86E-12 | 208.6985 | Mesothelial |
| 85 | LINC01279 | 6.07E-12 | 9.083219 | Mesothelial |
| 86 | STEAP1 | 6.51E-12 | 10.131902 | Mesothelial |
| 87 | EEF2 | 6.70E-12 | 21.800798 | Mesothelial |
| 88 | IFITM1 | 6.81E-12 | 18.57131 | Mesothelial |
| 89 | RPS15A | 7.02E-12 | 112.72822 | Mesothelial |
| 90 | CALD1 | 7.35E-12 | 12.113905 | Mesothelial |
| 91 | RPS15 | 7.40E-12 | 81.453186 | Mesothelial |
| 92 | RPS6 | 8.30E-12 | 100.944916 | Mesothelial |
| 93 | SNHG5 | 8.40E-12 | 19.751951 | Mesothelial |
| 94 | RPL7A | 8.40E-12 | 58.392357 | Mesothelial |
| 95 | RPL26 | 8.89E-12 | 98.80989 | Mesothelial |
| 96 | UAP1 | 9.34E-12 | 7.0076833 | Mesothelial |
| 97 | RPL35A | 9.71E-12 | 67.77772 | Mesothelial |
| 98 | CPE | 1.11E-11 | 9.79123 | Mesothelial |
| 99 | CLDN1 | 1.12E-11 | 11.583027 | Mesothelial |
| 0 | SCGB3A1 | 0 | inf | Mucous |
| 1 | WFDC2 | 0 | 318.49905 | Mucous |
| 2 | BPIFB1 | 5.87E-305 | 585.5846 | Mucous |
| 3 | LCN2 | 2.70E-303 | 235.87361 | Mucous |
| 4 | CP | 2.42E-302 | 37.31986 | Mucous |
| 5 | PIGR | 4.57E-298 | 67.71106 | Mucous |
| 6 | SLPI | 4.66E-298 | 829.086 | Mucous |
| 7 | SCGB1A1 | 1.12E-295 | inf | Mucous |
| 8 | KLK11 | 1.23E-286 | 18.868025 | Mucous |
| 9 | CXCL17 | 1.05E-284 | 41.992203 | Mucous |
| 10 | KRT7 | 1.45E-284 | 30.851065 | Mucous |
| 11 | TMC5 | 5.31E-281 | 16.53842 | Mucous |
| 12 | C3 | 1.25E-278 | 26.232342 | Mucous |
| 13 | MDK | 1.26E-278 | 18.864548 | Mucous |
| 14 | TMEM45A | 2.07E-278 | 22.206161 | Mucous |
| 15 | FOLR1 | 1.21E-265 | 22.096842 | Mucous |
| 16 | CLU | 5.58E-264 | 49.3905 | Mucous |
| 17 | CYB5A | 4.91E-260 | 58.296906 | Mucous |
| 18 | SAA1 | 5.04E-260 | 223.30696 | Mucous |
| 19 | FXYD3 | 6.47E-259 | 17.60703 | Mucous |
| 20 | TSPAN8 | 7.79E-257 | 13.068964 | Mucous |
| 21 | TACSTD2 | 4.47E-256 | 17.875925 | Mucous |
| 22 | RARRES1 | 1.98E-254 | 28.970127 | Mucous |
| 23 | AGR2 | 8.94E-252 | 29.501617 | Mucous |
| 24 | MUC1 | 3.90E-251 | 16.204979 | Mucous |
| 25 | ATP1B1 | 6.51E-244 | 15.12159 | Mucous |
| 26 | AKR1C1 | 6.91E-244 | 20.067022 | Mucous |
| 27 | MGST1 | 4.32E-243 | 22.509857 | Mucous |
| 28 | SAA2 | 1.06E-242 | 115.57284 | Mucous |
| 29 | SAT1 | 4.14E-242 | 128.38692 | Mucous |
| 30 | CFB | 1.14E-241 | 15.559507 | Mucous |
| 31 | KRT18 | 2.16E-241 | 17.179132 | Mucous |
| 32 | ELF3 | 2.21E-241 | 14.452143 | Mucous |
| 33 | TSPAN1 | 2.23E-238 | 10.736113 | Mucous |
| 34 | MMP7 | 3.90E-238 | 27.536036 | Mucous |
| 35 | PPAP2C | 9.86E-238 | 7.264186 | Mucous |
| 36 | XBP1 | 7.75E-237 | 28.252985 | Mucous |
| 37 | KRT19 | 2.05E-235 | 30.8337 | Mucous |
| 38 | SERPINF1 | 1.04E-234 | 11.215859 | Mucous |
| 39 | HS3ST1 | 1.45E-233 | 9.615686 | Mucous |
| 40 | DUSP23 | 2.05E-231 | 18.593744 | Mucous |
| 41 | C19orf33 | 3.05E-230 | 9.413929 | Mucous |
| 42 | CD55 | 4.99E-230 | 22.641973 | Mucous |
| 43 | AGR3 | 1.90E-228 | 19.165491 | Mucous |
| 44 | STEAP4 | 2.78E-227 | 8.345433 | Mucous |
| 45 | PDZK1IP1 | 7.21E-227 | 14.902624 | Mucous |
| 46 | RNF145 | 2.13E-224 | 10.443712 | Mucous |

| | | | | |
|---|---|---|---|---|
| 47 | ANKRD36C | 2.57E-224 | 16.106628 | Mucous |
| 48 | KRT8 | 3.66E-220 | 13.05278 | Mucous |
| 49 | SSR4 | 5.08E-220 | 20.377335 | Mucous |
| 50 | RPL18A | 6.91E-219 | 84.13158 | Mucous |
| 51 | CEACAM6 | 7.51E-218 | 10.239408 | Mucous |
| 52 | EZR | 1.43E-217 | 13.63212 | Mucous |
| 53 | C16orf89 | 5.17E-215 | 9.133353 | Mucous |
| 54 | SPINT2 | 9.84E-215 | 10.756083 | Mucous |
| 55 | INSR | 1.29E-214 | 8.059031 | Mucous |
| 56 | MUC5B | 1.54E-213 | 22.247877 | Mucous |
| 57 | ALCAM | 5.82E-211 | 7.1629457 | Mucous |
| 58 | PERP | 8.51E-211 | 7.1780252 | Mucous |
| 59 | MUC4 | 1.28E-209 | 8.550914 | Mucous |
| 60 | GSTA1 | 6.15E-209 | 10.591477 | Mucous |
| 61 | EPCAM | 2.45E-208 | 7.5762386 | Mucous |
| 62 | ASS1 | 3.55E-208 | 7.3206716 | Mucous |
| 63 | SLC44A4 | 1.52E-207 | 6.5071373 | Mucous |
| 64 | RPS6 | 4.98E-206 | 75.676125 | Mucous |
| 65 | NCOA7 | 2.23E-205 | 14.612577 | Mucous |
| 66 | LINC00342 | 1.02E-204 | 12.970521 | Mucous |
| 67 | RPL13 | 5.55E-204 | 95.48575 | Mucous |
| 68 | UBD | 1.60E-203 | 23.200409 | Mucous |
| 69 | BACE2 | 3.68E-203 | 8.0822 | Mucous |
| 70 | RPL13A | 2.17E-201 | 97.34534 | Mucous |
| 71 | RPLP0 | 8.36E-201 | 34.735016 | Mucous |
| 72 | SDC4 | 4.36E-200 | 15.59028 | Mucous |
| 73 | S100P | 1.16E-199 | 19.997639 | Mucous |
| 74 | LINC01207 | 2.42E-199 | 8.801607 | Mucous |
| 75 | RPS18 | 2.90E-199 | 100.27623 | Mucous |
| 76 | CYP2B7P | 3.85E-198 | 8.420186 | Mucous |
| 77 | RPS12 | 9.03E-198 | 90.3851 | Mucous |
| 78 | CXCL1 | 2.23E-197 | 62.35493 | Mucous |
| 79 | TMEM205 | 1.10E-196 | 7.217656 | Mucous |
| 80 | RPL18 | 3.62E-196 | 36.66794 | Mucous |
| 81 | CLDN7 | 1.20E-195 | 7.4529634 | Mucous |
| 82 | HES4 | 5.74E-195 | 6.8915095 | Mucous |
| 83 | RHOV | 1.93E-194 | 9.324352 | Mucous |
| 84 | RPL7 | 9.04E-194 | 60.419167 | Mucous |
| 85 | RPLP1 | 3.54E-193 | 116.39772 | Mucous |
| 86 | RPL8 | 6.98E-193 | 53.41318 | Mucous |
| 87 | MIF | 4.63E-192 | 21.408901 | Mucous |
| 88 | SMIM22 | 1.37E-191 | 6.2985682 | Mucous |
| 89 | RPL29 | 7.52E-191 | 40.10914 | Mucous |
| 90 | GSTP1 | 2.60E-190 | 33.91987 | Mucous |
| 91 | SFTPB | 8.02E-190 | 104.80058 | Mucous |
| 92 | RPS8 | 1.09E-189 | 51.435387 | Mucous |
| 93 | RPL10A | 1.43E-189 | 38.503567 | Mucous |
| 94 | RNASE1 | 3.22E-189 | 24.178007 | Mucous |
| 95 | CLDN4 | 1.16E-188 | 10.479023 | Mucous |
| 96 | RPL3 | 1.08E-187 | 60.175175 | Mucous |
| 97 | ARFGEF3 | 1.18E-186 | 6.791667 | Mucous |
| 98 | RPS16 | 7.69E-186 | 48.299442 | Mucous |
| 99 | RPL5 | 1.08E-185 | 30.72287 | Mucous |
| 0 | HLA-DPB1 | 1.26E-64 | 154.20491 | Myeloid Dendritic Type 1 |
| 1 | CSF2RA | 3.99E-62 | 10.636388 | Myeloid Dendritic Type 1 |
| 2 | RGS1 | 3.27E-59 | 21.609125 | Myeloid Dendritic Type 1 |
| 3 | LSP1 | 1.45E-58 | 15.158431 | Myeloid Dendritic Type 1 |
| 4 | HLA-DPA1 | 3.78E-58 | 145.75674 | Myeloid Dendritic Type 1 |
| 5 | HLA-DQA1 | 3.78E-58 | 61.708282 | Myeloid Dendritic Type 1 |
| 6 | HLA-DQB1 | 9.37E-52 | 60.58952 | Myeloid Dendritic Type 1 |
| 7 | GPR183 | 1.18E-51 | 16.135721 | Myeloid Dendritic Type 1 |
| 8 | RGS10 | 5.48E-51 | 8.15137 | Myeloid Dendritic Type 1 |
| 9 | HLA-DQA2 | 7.47E-51 | 22.33638 | Myeloid Dendritic Type 1 |
| 10 | WFDC21P | 1.41E-49 | 26.164694 | Myeloid Dendritic Type 1 |

| | | | | |
|---|---|---|---|---|
| 12 | BASP1 | 1.88E-49 | 8.748354 | Myeloid Dendritic Type 1 |
| 11 | CD74 | 1.88E-49 | 321.77625 | Myeloid Dendritic Type 1 |
| 13 | NAPSB | 2.56E-47 | 12.094909 | Myeloid Dendritic Type 1 |
| 14 | LGALS2 | 3.70E-47 | 11.880948 | Myeloid Dendritic Type 1 |
| 15 | NAP1L1 | 1.85E-46 | 9.515919 | Myeloid Dendritic Type 1 |
| 16 | CXCR4 | 2.45E-45 | 12.761378 | Myeloid Dendritic Type 1 |
| 17 | SERPINB9 | 1.82E-44 | 9.647783 | Myeloid Dendritic Type 1 |
| 18 | HLA-DRA | 1.82E-44 | 218.31288 | Myeloid Dendritic Type 1 |
| 19 | DUSP4 | 3.64E-43 | 8.523402 | Myeloid Dendritic Type 1 |
| 20 | PABPC1 | 4.28E-43 | 15.178643 | Myeloid Dendritic Type 1 |
| 21 | FNBP1 | 2.85E-42 | 4.876284 | Myeloid Dendritic Type 1 |
| 22 | RPS2 | 5.19E-42 | 51.25716 | Myeloid Dendritic Type 1 |
| 23 | EEF1G | 2.17E-41 | 21.101118 | Myeloid Dendritic Type 1 |
| 24 | RPS23 | 2.23E-41 | 37.041836 | Myeloid Dendritic Type 1 |
| 25 | CD83 | 6.79E-41 | 12.207467 | Myeloid Dendritic Type 1 |
| 26 | RPL18A | 1.11E-40 | 36.932697 | Myeloid Dendritic Type 1 |
| 27 | HLA-DRB5 | 1.66E-40 | 37.3412 | Myeloid Dendritic Type 1 |
| 28 | RPS16 | 1.07E-39 | 30.012003 | Myeloid Dendritic Type 1 |
| 29 | HLA-DRB1 | 1.07E-39 | 107.50179 | Myeloid Dendritic Type 1 |
| 30 | CST7 | 4.08E-39 | 13.12533 | Myeloid Dendritic Type 1 |
| 31 | RPL10 | 5.00E-39 | 69.51679 | Myeloid Dendritic Type 1 |
| 32 | RPL4 | 1.92E-38 | 16.032597 | Myeloid Dendritic Type 1 |
| 33 | EEF1B2 | 2.50E-38 | 11.698454 | Myeloid Dendritic Type 1 |
| 34 | HLA-DRB6 | 2.50E-38 | 31.23465 | Myeloid Dendritic Type 1 |
| 35 | SUB1 | 4.27E-38 | 8.020786 | Myeloid Dendritic Type 1 |
| 36 | ADAM8 | 5.56E-38 | 6.2863283 | Myeloid Dendritic Type 1 |
| 37 | RPS24 | 1.12E-37 | 33.7923 | Myeloid Dendritic Type 1 |
| 38 | RPS3A | 2.43E-37 | 35.933285 | Myeloid Dendritic Type 1 |
| 39 | COTL1 | 3.68E-37 | 13.849825 | Myeloid Dendritic Type 1 |
| 40 | EIF1 | 5.97E-37 | 29.278687 | Myeloid Dendritic Type 1 |
| 41 | TMSB10 | 6.54E-37 | 95.20898 | Myeloid Dendritic Type 1 |
| 42 | RPL8 | 8.39E-37 | 26.75356 | Myeloid Dendritic Type 1 |
| 43 | ACTG1 | 4.79E-36 | 30.340582 | Myeloid Dendritic Type 1 |
| 44 | KLF6 | 4.79E-36 | 10.38243 | Myeloid Dendritic Type 1 |
| 45 | SYNGR2 | 8.08E-36 | 6.0087776 | Myeloid Dendritic Type 1 |
| 46 | RPL26 | 3.55E-35 | 33.190796 | Myeloid Dendritic Type 1 |
| 47 | TYMP | 7.63E-35 | 9.5451975 | Myeloid Dendritic Type 1 |
| 48 | RPS9 | 1.31E-34 | 28.528008 | Myeloid Dendritic Type 1 |
| 49 | RPS6 | 1.83E-34 | 32.741253 | Myeloid Dendritic Type 1 |
| 50 | SLC25A6 | 1.90E-34 | 9.837016 | Myeloid Dendritic Type 1 |
| 51 | CMTM6 | 2.62E-34 | 5.3195014 | Myeloid Dendritic Type 1 |
| 52 | UBA52 | 4.23E-34 | 17.959583 | Myeloid Dendritic Type 1 |
| 53 | RPS18 | 5.26E-34 | 45.905563 | Myeloid Dendritic Type 1 |
| 54 | RPS14 | 8.11E-34 | 36.135357 | Myeloid Dendritic Type 1 |
| 55 | RPS7 | 8.20E-34 | 20.1722 | Myeloid Dendritic Type 1 |
| 56 | RPS21 | 9.50E-34 | 17.665546 | Myeloid Dendritic Type 1 |
| 57 | CST3 | 9.99E-34 | 122.01897 | Myeloid Dendritic Type 1 |
| 58 | RPS11 | 2.45E-33 | 18.179647 | Myeloid Dendritic Type 1 |
| 59 | ACTB | 2.82E-33 | 128.40352 | Myeloid Dendritic Type 1 |
| 60 | RPL13 | 3.20E-33 | 39.02694 | Myeloid Dendritic Type 1 |
| 61 | REL | 3.34E-33 | 8.508416 | Myeloid Dendritic Type 1 |
| 62 | SERPINF1 | 4.58E-33 | 4.3048077 | Myeloid Dendritic Type 1 |
| 63 | RPS19 | 9.05E-33 | 39.588108 | Myeloid Dendritic Type 1 |
| 64 | VIM | 9.42E-33 | 57.21876 | Myeloid Dendritic Type 1 |
| 65 | PLEK | 9.43E-33 | 5.190507 | Myeloid Dendritic Type 1 |
| 66 | IGFLR1 | 2.80E-32 | 3.7850666 | Myeloid Dendritic Type 1 |
| 67 | RPS15 | 4.50E-32 | 24.9756 | Myeloid Dendritic Type 1 |
| 68 | RPL28 | 4.68E-32 | 35.909416 | Myeloid Dendritic Type 1 |
| 69 | RPL35 | 7.14E-32 | 18.369242 | Myeloid Dendritic Type 1 |
| 70 | MARCKSL1 | 1.13E-31 | 9.690175 | Myeloid Dendritic Type 1 |
| 71 | RPL36 | 1.70E-31 | 16.989773 | Myeloid Dendritic Type 1 |
| 72 | HLA-DMA | 3.35E-31 | 9.604845 | Myeloid Dendritic Type 1 |
| 74 | RPL9 | 3.52E-31 | 24.84069 | Myeloid Dendritic Type 1 |
| 73 | RPS17 | 3.52E-31 | 21.140476 | Myeloid Dendritic Type 1 |

| | | | | |
|---|---|---|---|---|
| 75 | FAU | 9.14E-31 | 17.48323 | Myeloid Dendritic Type 1 |
| 76 | HLA-DQB2 | 1.10E-30 | 7.993377 | Myeloid Dendritic Type 1 |
| 77 | BTG1 | 1.16E-30 | 21.925415 | Myeloid Dendritic Type 1 |
| 78 | RPL18 | 1.22E-30 | 15.627365 | Myeloid Dendritic Type 1 |
| 79 | ZFAS1 | 1.34E-30 | 6.8767548 | Myeloid Dendritic Type 1 |
| 80 | RPL35A | 1.59E-30 | 22.388382 | Myeloid Dendritic Type 1 |
| 81 | RPS13 | 1.85E-30 | 23.90784 | Myeloid Dendritic Type 1 |
| 82 | SNHG5 | 2.06E-30 | 7.2719607 | Myeloid Dendritic Type 1 |
| 83 | ADAM19 | 2.17E-30 | 7.135202 | Myeloid Dendritic Type 1 |
| 84 | RPL21 | 2.28E-30 | 38.945786 | Myeloid Dendritic Type 1 |
| 85 | RPS29 | 2.63E-30 | 32.026478 | Myeloid Dendritic Type 1 |
| 86 | RPS15A | 3.20E-30 | 26.70432 | Myeloid Dendritic Type 1 |
| 87 | RPS8 | 3.78E-30 | 23.955458 | Myeloid Dendritic Type 1 |
| 88 | RPL15 | 4.87E-30 | 24.772467 | Myeloid Dendritic Type 1 |
| 89 | RPS5 | 5.32E-30 | 14.683911 | Myeloid Dendritic Type 1 |
| 90 | NEAT1 | 7.43E-30 | 17.261732 | Myeloid Dendritic Type 1 |
| 91 | PFN1 | 7.97E-30 | 26.303978 | Myeloid Dendritic Type 1 |
| 92 | RPL13A | 9.53E-30 | 37.630165 | Myeloid Dendritic Type 1 |
| 93 | RPLP1 | 1.21E-29 | 50.054993 | Myeloid Dendritic Type 1 |
| 94 | RPLP0 | 1.25E-29 | 12.938714 | Myeloid Dendritic Type 1 |
| 95 | PTMA | 1.27E-29 | 25.83799 | Myeloid Dendritic Type 1 |
| 96 | GNB2L1 | 1.27E-29 | 13.230234 | Myeloid Dendritic Type 1 |
| 97 | RPL6 | 4.87E-29 | 18.17811 | Myeloid Dendritic Type 1 |
| 98 | RPL37 | 4.94E-29 | 21.476488 | Myeloid Dendritic Type 1 |
| 99 | RPL23A | 5.16E-29 | 25.615456 | Myeloid Dendritic Type 1 |
| 0 | HLA-DPB1 | 2.42E-126 | 124.980835 | Myeloid Dendritic Type 2 |
| 1 | CLEC10A | 3.19E-125 | 14.726264 | Myeloid Dendritic Type 2 |
| 2 | FGL2 | 3.33E-123 | 7.9662 | Myeloid Dendritic Type 2 |
| 3 | FCER1A | 1.83E-117 | 24.722076 | Myeloid Dendritic Type 2 |
| 4 | HLA-DPA1 | 3.91E-111 | 108.063805 | Myeloid Dendritic Type 2 |
| 5 | HLA-DQA1 | 1.07E-109 | 48.46986 | Myeloid Dendritic Type 2 |
| 6 | GPR183 | 4.63E-107 | 13.207848 | Myeloid Dendritic Type 2 |
| 7 | RNASE6 | 1.20E-106 | 8.086956 | Myeloid Dendritic Type 2 |
| 8 | LSP1 | 2.99E-105 | 11.40307 | Myeloid Dendritic Type 2 |
| 9 | CD1C | 1.68E-104 | 16.000498 | Myeloid Dendritic Type 2 |
| 10 | RGS10 | 1.68E-104 | 6.232216 | Myeloid Dendritic Type 2 |
| 11 | HLA-DQB1 | 1.87E-104 | 41.259193 | Myeloid Dendritic Type 2 |
| 12 | CPVL | 2.89E-103 | 10.944588 | Myeloid Dendritic Type 2 |
| 13 | CST3 | 3.26E-103 | 91.099724 | Myeloid Dendritic Type 2 |
| 14 | HLA-DMB | 7.56E-102 | 10.494372 | Myeloid Dendritic Type 2 |
| 15 | RPS24 | 1.14E-100 | 43.483162 | Myeloid Dendritic Type 2 |
| 16 | COTL1 | 1.53E-100 | 15.988633 | Myeloid Dendritic Type 2 |
| 17 | CORO1A | 6.90E-100 | 9.582358 | Myeloid Dendritic Type 2 |
| 18 | RPS23 | 1.10E-99 | 44.50674 | Myeloid Dendritic Type 2 |
| 19 | CXCR4 | 4.81E-99 | 13.371266 | Myeloid Dendritic Type 2 |
| 20 | SNHG5 | 8.14E-98 | 11.055731 | Myeloid Dendritic Type 2 |
| 21 | HLA-DQA2 | 3.34E-97 | 13.121294 | Myeloid Dendritic Type 2 |
| 22 | RPL26 | 1.20E-96 | 46.517952 | Myeloid Dendritic Type 2 |
| 23 | FAM26F | 2.90E-96 | 6.501681 | Myeloid Dendritic Type 2 |
| 24 | NAPSB | 8.60E-96 | 8.587744 | Myeloid Dendritic Type 2 |
| 25 | HLA-DRB6 | 2.57E-95 | 49.14328 | Myeloid Dendritic Type 2 |
| 26 | RPS2 | 8.73E-95 | 59.667248 | Myeloid Dendritic Type 2 |
| 27 | PABPC1 | 1.06E-94 | 15.366042 | Myeloid Dendritic Type 2 |
| 28 | MS4A6A | 2.23E-94 | 14.402457 | Myeloid Dendritic Type 2 |
| 29 | RPS3A | 5.76E-94 | 47.80012 | Myeloid Dendritic Type 2 |
| 30 | EEF1G | 1.12E-92 | 22.7955 | Myeloid Dendritic Type 2 |
| 31 | CD74 | 1.37E-91 | 235.47626 | Myeloid Dendritic Type 2 |
| 32 | RPL4 | 1.50E-91 | 18.36223 | Myeloid Dendritic Type 2 |
| 33 | HLA-DRA | 1.57E-90 | 192.65604 | Myeloid Dendritic Type 2 |
| 34 | AP1S2 | 5.65E-89 | 5.908807 | Myeloid Dendritic Type 2 |
| 35 | RPL18A | 1.17E-88 | 40.986523 | Myeloid Dendritic Type 2 |
| 36 | RPS18 | 1.34E-88 | 59.747173 | Myeloid Dendritic Type 2 |
| 37 | HLA-DRB5 | 3.72E-88 | 36.14931 | Myeloid Dendritic Type 2 |
| 38 | RPL10 | 8.31E-87 | 74.29511 | Myeloid Dendritic Type 2 |

| | | | | |
|---|---|---|---|---|
| 39 | SLC25A6 | 8.27E-86 | 11.578219 | Myeloid Dendritic Type 2 |
| 40 | HLA-DMA | 8.27E-86 | 12.633181 | Myeloid Dendritic Type 2 |
| 41 | HLA-DRB1 | 2.44E-85 | 92.011 | Myeloid Dendritic Type 2 |
| 42 | HLA-DQB2 | 4.31E-85 | 8.241801 | Myeloid Dendritic Type 2 |
| 43 | RPL13 | 6.75E-85 | 51.160435 | Myeloid Dendritic Type 2 |
| 44 | RPS7 | 1.38E-84 | 23.726835 | Myeloid Dendritic Type 2 |
| 45 | STK17B | 1.89E-84 | 5.189092 | Myeloid Dendritic Type 2 |
| 46 | LGALS2 | 8.68E-84 | 9.559346 | Myeloid Dendritic Type 2 |
| 47 | RPL13A | 2.80E-83 | 52.30306 | Myeloid Dendritic Type 2 |
| 48 | RPL18 | 6.06E-83 | 20.58693 | Myeloid Dendritic Type 2 |
| 49 | RPL32 | 2.50E-82 | 40.058807 | Myeloid Dendritic Type 2 |
| 50 | ITGB2 | 3.74E-82 | 7.0345364 | Myeloid Dendritic Type 2 |
| 51 | RPS6 | 3.98E-82 | 39.886707 | Myeloid Dendritic Type 2 |
| 52 | RPS15A | 8.48E-82 | 36.216805 | Myeloid Dendritic Type 2 |
| 53 | RPS4X | 2.84E-81 | 37.18885 | Myeloid Dendritic Type 2 |
| 54 | RPS14 | 5.32E-81 | 43.00967 | Myeloid Dendritic Type 2 |
| 55 | RPS8 | 9.84E-81 | 29.673412 | Myeloid Dendritic Type 2 |
| 56 | RPS13 | 1.06E-80 | 32.539204 | Myeloid Dendritic Type 2 |
| 57 | EEF1B2 | 1.42E-80 | 10.633624 | Myeloid Dendritic Type 2 |
| 58 | RPS17 | 2.74E-80 | 26.464146 | Myeloid Dendritic Type 2 |
| 59 | RPS9 | 1.17E-79 | 29.790403 | Myeloid Dendritic Type 2 |
| 60 | ZFP36L2 | 1.28E-79 | 10.767775 | Myeloid Dendritic Type 2 |
| 61 | RPS11 | 1.53E-79 | 20.601583 | Myeloid Dendritic Type 2 |
| 62 | RPL34 | 1.77E-78 | 43.533398 | Myeloid Dendritic Type 2 |
| 63 | RPL15 | 4.52E-78 | 32.909096 | Myeloid Dendritic Type 2 |
| 64 | RPL39 | 6.57E-78 | 37.673923 | Myeloid Dendritic Type 2 |
| 65 | CSF2RA | 7.67E-78 | 5.513429 | Myeloid Dendritic Type 2 |
| 66 | RPS16 | 1.46E-77 | 26.767694 | Myeloid Dendritic Type 2 |
| 67 | RPL7 | 2.54E-77 | 31.530901 | Myeloid Dendritic Type 2 |
| 68 | RPL10A | 3.34E-77 | 21.597137 | Myeloid Dendritic Type 2 |
| 69 | LIMD2 | 3.61E-77 | 5.11186 | Myeloid Dendritic Type 2 |
| 70 | RPL35A | 6.25E-77 | 23.695515 | Myeloid Dendritic Type 2 |
| 71 | RPS15 | 6.57E-77 | 31.661217 | Myeloid Dendritic Type 2 |
| 72 | EIF3L | 2.30E-76 | 6.0110226 | Myeloid Dendritic Type 2 |
| 73 | RPL23A | 2.93E-76 | 29.613287 | Myeloid Dendritic Type 2 |
| 74 | EEF2 | 4.41E-76 | 9.214699 | Myeloid Dendritic Type 2 |
| 75 | RPL11 | 6.14E-76 | 37.64778 | Myeloid Dendritic Type 2 |
| 76 | GNB2L1 | 8.45E-76 | 17.619999 | Myeloid Dendritic Type 2 |
| 77 | RPL27A | 1.35E-75 | 34.62006 | Myeloid Dendritic Type 2 |
| 78 | RPL19 | 6.12E-75 | 33.55193 | Myeloid Dendritic Type 2 |
| 79 | RPL29 | 9.41E-75 | 20.125711 | Myeloid Dendritic Type 2 |
| 80 | RPL5 | 1.17E-74 | 17.794254 | Myeloid Dendritic Type 2 |
| 81 | GDI2 | 3.12E-74 | 4.5941916 | Myeloid Dendritic Type 2 |
| 82 | RPL7A | 3.62E-74 | 18.883976 | Myeloid Dendritic Type 2 |
| 83 | RPSA | 5.13E-74 | 15.234513 | Myeloid Dendritic Type 2 |
| 84 | RGS2 | 8.70E-74 | 10.11456 | Myeloid Dendritic Type 2 |
| 85 | RPL37 | 2.41E-73 | 25.485645 | Myeloid Dendritic Type 2 |
| 86 | RPS3 | 3.63E-73 | 29.67139 | Myeloid Dendritic Type 2 |
| 87 | RPL35 | 4.37E-73 | 21.019075 | Myeloid Dendritic Type 2 |
| 88 | GLIPR1 | 1.47E-72 | 4.177023 | Myeloid Dendritic Type 2 |
| 89 | IGFLR1 | 1.99E-72 | 4.3742423 | Myeloid Dendritic Type 2 |
| 90 | RPL28 | 1.99E-72 | 38.26227 | Myeloid Dendritic Type 2 |
| 91 | AMICA1 | 5.04E-72 | 4.739721 | Myeloid Dendritic Type 2 |
| 92 | UBA52 | 5.04E-72 | 18.733248 | Myeloid Dendritic Type 2 |
| 93 | RPS5 | 6.39E-72 | 16.486376 | Myeloid Dendritic Type 2 |
| 94 | RPL6 | 7.39E-72 | 23.414228 | Myeloid Dendritic Type 2 |
| 95 | RPLP0 | 1.36E-71 | 14.798737 | Myeloid Dendritic Type 2 |
| 96 | ACTB | 5.75E-71 | 107.93953 | Myeloid Dendritic Type 2 |
| 97 | RPL22 | 1.02E-70 | 14.445199 | Myeloid Dendritic Type 2 |
| 98 | HNRNPA1 | 1.35E-70 | 11.125085 | Myeloid Dendritic Type 2 |
| 99 | RPL17 | 2.44E-70 | 9.633645 | Myeloid Dendritic Type 2 |
| 0 | DCN | 7.94E-144 | 31.709053 | Myofibroblast |
| 1 | ASPN | 2.60E-131 | 17.30726 | Myofibroblast |
| 2 | LUM | 1.36E-126 | 18.442972 | Myofibroblast |

| | | | | |
|---|---|---|---|---|
| 3 | CFH | 6.53E-126 | 10.489023 | Myofibroblast |
| 4 | BGN | 1.74E-121 | 12.806381 | Myofibroblast |
| 5 | CLU | 6.25E-119 | 65.08445 | Myofibroblast |
| 6 | MGP | 3.38E-118 | 39.973614 | Myofibroblast |
| 7 | MFAP4 | 5.37E-115 | 7.8798366 | Myofibroblast |
| 8 | COL1A2 | 2.54E-114 | 14.818647 | Myofibroblast |
| 9 | PRELP | 6.81E-114 | 9.289991 | Myofibroblast |
| 10 | DKK3 | 2.92E-110 | 7.2398305 | Myofibroblast |
| 11 | CTGF | 4.43E-109 | 25.94315 | Myofibroblast |
| 12 | CALD1 | 2.43E-107 | 7.261638 | Myofibroblast |
| 13 | RARRES2 | 8.41E-105 | 7.214081 | Myofibroblast |
| 14 | TAGLN | 1.00E-101 | 12.500251 | Myofibroblast |
| 15 | C1S | 3.15E-98 | 6.9927077 | Myofibroblast |
| 16 | COL6A2 | 3.39E-98 | 8.241182 | Myofibroblast |
| 17 | LTBP1 | 2.05E-95 | 6.567403 | Myofibroblast |
| 18 | TSPAN8 | 3.57E-93 | 6.5464854 | Myofibroblast |
| 19 | FHL1 | 4.65E-88 | 7.253892 | Myofibroblast |
| 20 | SPARCL1 | 2.71E-86 | 7.5065813 | Myofibroblast |
| 21 | ACTA2 | 1.57E-84 | 8.9150095 | Myofibroblast |
| 22 | LTBP2 | 1.96E-84 | 7.1486106 | Myofibroblast |
| 23 | COL3A1 | 2.72E-84 | 13.066629 | Myofibroblast |
| 25 | DPT | 3.00E-82 | 9.377009 | Myofibroblast |
| 24 | COL1A1 | 3.00E-82 | 13.247935 | Myofibroblast |
| 26 | TPM2 | 2.04E-75 | 4.0899525 | Myofibroblast |
| 27 | FILIP1L | 6.22E-73 | 5.1426916 | Myofibroblast |
| 28 | BCHE | 2.09E-71 | 7.857901 | Myofibroblast |
| 29 | CYR61 | 6.12E-71 | 11.257012 | Myofibroblast |
| 30 | MMP2 | 3.61E-70 | 6.8120427 | Myofibroblast |
| 31 | PDLIM3 | 5.47E-69 | 5.2313094 | Myofibroblast |
| 32 | AEBP1 | 1.08E-68 | 5.333468 | Myofibroblast |
| 33 | LMCD1 | 1.15E-68 | 4.597461 | Myofibroblast |
| 34 | C1R | 1.46E-68 | 4.898314 | Myofibroblast |
| 35 | SDC2 | 5.77E-67 | 4.155126 | Myofibroblast |
| 36 | CNN3 | 1.15E-64 | 3.9137025 | Myofibroblast |
| 37 | RARRES1 | 4.40E-63 | 5.8990574 | Myofibroblast |
| 38 | MYL9 | 1.17E-62 | 2.8314948 | Myofibroblast |
| 39 | GEM | 1.66E-62 | 6.799424 | Myofibroblast |
| 40 | SCARA3 | 5.55E-62 | 6.9756417 | Myofibroblast |
| 41 | MXRA8 | 3.40E-59 | 5.519993 | Myofibroblast |
| 42 | GPX3 | 4.71E-56 | 6.0767756 | Myofibroblast |
| 43 | SELM | 1.31E-53 | 3.2763345 | Myofibroblast |
| 44 | FXYD1 | 1.75E-53 | 4.2694597 | Myofibroblast |
| 45 | PCOLCE | 4.89E-53 | 5.2578893 | Myofibroblast |
| 46 | CRYAB | 1.07E-52 | 5.6241055 | Myofibroblast |
| 47 | TSC22D1 | 7.51E-49 | 5.176346 | Myofibroblast |
| 48 | COL6A1 | 2.90E-48 | 5.330641 | Myofibroblast |
| 49 | PLAC9 | 3.27E-48 | 3.5749853 | Myofibroblast |
| 50 | ADH1B | 6.28E-47 | 4.690405 | Myofibroblast |
| 51 | F2R | 7.57E-47 | 4.870433 | Myofibroblast |
| 52 | A2M | 7.91E-47 | 4.7087626 | Myofibroblast |
| 53 | EFEMP2 | 3.98E-46 | 3.9595356 | Myofibroblast |
| 54 | PMP22 | 5.61E-45 | 2.9022217 | Myofibroblast |
| 55 | TNC | 5.65E-45 | 5.995815 | Myofibroblast |
| 56 | TIMP1 | 7.35E-45 | 13.532254 | Myofibroblast |
| 57 | ANGPTL2 | 2.69E-44 | 6.2147264 | Myofibroblast |
| 58 | WIF1 | 1.24E-42 | 5.3228774 | Myofibroblast |
| 59 | TPPP3 | 1.48E-42 | 1.1250427 | Myofibroblast |
| 60 | SPARC | 1.87E-42 | 5.784213 | Myofibroblast |
| 61 | PLS3 | 3.80E-42 | 3.20258 | Myofibroblast |
| 62 | IGFBP6 | 7.69E-42 | 4.1869116 | Myofibroblast |
| 63 | TMEM98 | 8.49E-41 | 3.5865831 | Myofibroblast |
| 64 | ITGBL1 | 1.31E-40 | 6.766789 | Myofibroblast |
| 65 | FN1 | 6.13E-39 | 1.3204472 | Myofibroblast |
| 66 | PALLD | 2.28E-38 | 3.6956754 | Myofibroblast |

| | | | | |
|---|---|---|---|---|
| 67 | MT1X | 5.00E-38 | 26.1128 | Myofibroblast |
| 68 | RERG | 9.49E-38 | 5.2286644 | Myofibroblast |
| 69 | THY1 | 1.67E-37 | 6.034731 | Myofibroblast |
| 70 | PKIG | 1.87E-36 | 2.8502162 | Myofibroblast |
| 71 | NNMT | 1.53E-35 | 4.906992 | Myofibroblast |
| 72 | NGFRAP1 | 3.21E-35 | 2.381246 | Myofibroblast |
| 73 | TGFB1I1 | 3.74E-34 | 2.9213166 | Myofibroblast |
| 74 | NFIB | 3.98E-34 | 2.4869425 | Myofibroblast |
| 75 | PAMR1 | 6.38E-34 | 7.1286864 | Myofibroblast |
| 76 | FBLN1 | 2.79E-33 | 3.0025172 | Myofibroblast |
| 77 | GAS6 | 2.92E-33 | 2.2033648 | Myofibroblast |
| 78 | ELN | 7.32E-33 | 4.467133 | Myofibroblast |
| 79 | MAMDC2 | 8.40E-33 | 5.2441306 | Myofibroblast |
| 80 | RAB34 | 1.25E-32 | 2.241339 | Myofibroblast |
| 81 | FKBP10 | 1.56E-32 | 4.748112 | Myofibroblast |
| 82 | FSTL1 | 1.16E-31 | 3.3912303 | Myofibroblast |
| 83 | CAV1 | 1.95E-31 | 1.6510775 | Myofibroblast |
| 84 | LTBP4 | 4.75E-31 | 2.8884144 | Myofibroblast |
| 85 | SSPN | 5.55E-31 | 3.927918 | Myofibroblast |
| 87 | CLEC11A | 6.45E-31 | 4.0364113 | Myofibroblast |
| 86 | SFRP4 | 6.45E-31 | 3.345292 | Myofibroblast |
| 88 | SPRY1 | 8.66E-31 | 2.843787 | Myofibroblast |
| 89 | COL6A3 | 3.56E-30 | 6.3556547 | Myofibroblast |
| 90 | SEPP1 | 6.05E-30 | 1.8253462 | Myofibroblast |
| 91 | CSRP1 | 8.64E-29 | 2.424511 | Myofibroblast |
| 92 | VCAN | 8.90E-29 | 2.5804038 | Myofibroblast |
| 93 | EMILIN1 | 1.37E-28 | 4.5853066 | Myofibroblast |
| 94 | RCN3 | 2.17E-28 | 4.7745833 | Myofibroblast |
| 95 | VIM | 3.96E-28 | -2.3838181 | Myofibroblast |
| 96 | NDN | 2.35E-27 | 3.0884178 | Myofibroblast |
| 97 | C12orf57 | 4.04E-27 | 1.9100093 | Myofibroblast |
| 98 | PRKCDBP | 9.38E-27 | 2.318175 | Myofibroblast |
| 99 | LGALS1 | 1.24E-26 | 1.2535775 | Myofibroblast |
| 0 | NKG7 | 0 | 26.929928 | Natural Killer |
| 37 | SH2D2A | 0 | 4.0490823 | Natural Killer |
| 38 | RARRES3 | 0 | 1.4857051 | Natural Killer |
| 39 | AREG | 0 | 2.3618653 | Natural Killer |
| 40 | MYOM2 | 0 | 6.0758443 | Natural Killer |
| 41 | LOC374443 | 0 | 3.9033842 | Natural Killer |
| 42 | CHST2 | 0 | 3.080726 | Natural Killer |
| 43 | NCR3 | 0 | 4.5953712 | Natural Killer |
| 44 | RAC2 | 0 | 1.4539435 | Natural Killer |
| 45 | TTC38 | 0 | 3.6315887 | Natural Killer |
| 46 | ZAP70 | 0 | 3.8581583 | Natural Killer |
| 47 | ZFP36L2 | 0 | 1.7839278 | Natural Killer |
| 48 | ISG20 | 0 | 1.5803571 | Natural Killer |
| 50 | HCST | 0 | 0.9570482 | Natural Killer |
| 51 | TXK | 0 | 4.7502027 | Natural Killer |
| 52 | GPR65 | 0 | 1.7811127 | Natural Killer |
| 53 | ADGRG1 | 0 | 3.025222 | Natural Killer |
| 54 | AKNA | 0 | 2.5741026 | Natural Killer |
| 68 | SH2D1B | 0 | 6.135507 | Natural Killer |
| 67 | SAMD3 | 0 | 3.8395648 | Natural Killer |
| 66 | HSH2D | 0 | 3.5774186 | Natural Killer |
| 65 | KLRC3 | 0 | 4.714794 | Natural Killer |
| 64 | PVRIG | 0 | 3.4931087 | Natural Killer |
| 63 | GNG2 | 0 | 3.0026493 | Natural Killer |
| 36 | BTG1 | 0 | 3.2280965 | Natural Killer |
| 62 | TBX21 | 0 | 4.5015845 | Natural Killer |
| 60 | IL2RG | 0 | 1.3185351 | Natural Killer |
| 59 | CD48 | 0 | 1.398744 | Natural Killer |
| 58 | FCRL6 | 0 | 4.425101 | Natural Killer |
| 57 | ARL4C | 0 | 2.716732 | Natural Killer |
| 56 | LOC100130872 | 0 | 3.9161508 | Natural Killer |

| | | | | |
|---|---|---|---|---|
| 55 | FYN | 0 | 2.2128253 | Natural Killer |
| 61 | PYHIN1 | 0 | 4.267638 | Natural Killer |
| 35 | IL2RB | 0 | 4.6479864 | Natural Killer |
| 49 | RUNX3 | 0 | 3.1583388 | Natural Killer |
| 33 | MATK | 0 | 3.856786 | Natural Killer |
| 14 | IFITM1 | 0 | 6.066145 | Natural Killer |
| 13 | PTPRCAP | 0 | 4.4496417 | Natural Killer |
| 12 | CTSW | 0 | 5.9425554 | Natural Killer |
| 11 | GZMA | 0 | 6.3389015 | Natural Killer |
| 10 | KLRF1 | 0 | 7.2456326 | Natural Killer |
| 9 | KLRD1 | 0 | 7.0200286 | Natural Killer |
| 15 | SPON2 | 0 | 6.279954 | Natural Killer |
| 8 | FGFBP2 | 0 | 9.780865 | Natural Killer |
| 6 | KLRB1 | 0 | 8.588076 | Natural Killer |
| 5 | CD7 | 0 | 7.9960504 | Natural Killer |
| 4 | CST7 | 0 | 9.008729 | Natural Killer |
| 3 | GNLY | 0 | 26.895988 | Natural Killer |
| 34 | HOPX | 0 | 0.7152916 | Natural Killer |
| 1 | GZMB | 0 | 15.287183 | Natural Killer |
| 7 | CD247 | 0 | 7.3345656 | Natural Killer |
| 16 | CLIC3 | 0 | 4.743005 | Natural Killer |
| 2 | PRF1 | 0 | 11.473014 | Natural Killer |
| 18 | CCL5 | 0 | 8.447749 | Natural Killer |
| 32 | CMC1 | 0 | 3.5078435 | Natural Killer |
| 31 | FCGR3A | 0 | 0.16037199 | Natural Killer |
| 30 | CXCR4 | 0 | 2.871777 | Natural Killer |
| 29 | ITGB2 | 0 | 2.098442 | Natural Killer |
| 27 | BIN2 | 0 | 2.9200952 | Natural Killer |
| 26 | SYTL3 | 0 | 3.651183 | Natural Killer |
| 25 | LIMD2 | 0 | 2.7341764 | Natural Killer |
| 28 | CORO1A | 0 | 2.1299396 | Natural Killer |
| 23 | S1PR5 | 0 | 5.617368 | Natural Killer |
| 22 | DDIT4 | 0 | 3.097143 | Natural Killer |
| 21 | PLAC8 | 0 | 2.1785967 | Natural Killer |
| 20 | ID2 | 0 | 4.4696107 | Natural Killer |
| 17 | GZMH | 0 | 5.029034 | Natural Killer |
| 19 | GZMM | 0 | 4.4480906 | Natural Killer |
| 24 | DUSP2 | 0 | 4.1000037 | Natural Killer |
| 69 | APMAP | 3.00E-299 | 1.4738934 | Natural Killer |
| 70 | TMIGD2 | 3.74E-296 | 5.5041766 | Natural Killer |
| 71 | CHST12 | 3.01E-289 | 1.6912606 | Natural Killer |
| 72 | LCK | 1.95E-284 | 2.1649363 | Natural Killer |
| 73 | GIMAP7 | 3.95E-281 | 1.1684937 | Natural Killer |
| 74 | PTPN4 | 4.90E-278 | 3.0700939 | Natural Killer |
| 75 | KLRC1 | 2.34E-272 | 4.663581 | Natural Killer |
| 76 | CX3CR1 | 3.63E-268 | 4.4395084 | Natural Killer |
| 77 | C12orf75 | 4.48E-263 | 0.98758334 | Natural Killer |
| 78 | ETS1 | 2.90E-262 | 1.7440062 | Natural Killer |
| 79 | ABHD17A | 4.78E-261 | 1.8091217 | Natural Killer |
| 80 | PTGDR | 4.23E-260 | 4.615263 | Natural Killer |
| 81 | CYTIP | 6.54E-255 | 0.92616767 | Natural Killer |
| 82 | YPEL1 | 1.20E-251 | 3.6568177 | Natural Killer |
| 83 | TRAF3IP3 | 3.92E-245 | 2.2828784 | Natural Killer |
| 84 | PTGER2 | 1.79E-244 | 2.4924703 | Natural Killer |
| 85 | CD69 | 4.30E-235 | 0.5754741 | Natural Killer |
| 86 | TYROBP | 1.37E-233 | -23.006157 | Natural Killer |
| 87 | ICAM3 | 8.41E-231 | 1.4212728 | Natural Killer |
| 88 | SLA2 | 3.14E-227 | 4.2064743 | Natural Killer |
| 89 | ABI3 | 1.50E-222 | 1.5147119 | Natural Killer |
| 90 | C1orf21 | 3.94E-210 | 1.8866298 | Natural Killer |
| 91 | FCMR | 1.49E-204 | 2.6295373 | Natural Killer |
| 92 | ANKRD20A11P | 1.66E-204 | 5.648094 | Natural Killer |
| 93 | IFITM2 | 2.24E-202 | 1.1693217 | Natural Killer |
| 94 | FKBP11 | 9.61E-201 | 1.4852477 | Natural Killer |

| | | | | |
|---|---|---|---|---|
| 95 | SYTL1 | 3.51E-199 | 1.5262645 | Natural Killer |
| 96 | TBC1D10C | 6.07E-198 | 1.0631616 | Natural Killer |
| 97 | PTPRC | 5.83E-195 | 0.26107764 | Natural Killer |
| 98 | PLEKHF1 | 1.16E-189 | 3.2750738 | Natural Killer |
| 99 | CD2 | 2.80E-187 | 1.223343 | Natural Killer |
| 0 | CCL5 | 6.02E-174 | 18.231905 | Natural Killer T |
| 1 | CXCR4 | 5.07E-141 | 9.583726 | Natural Killer T |
| 2 | CTSW | 1.76E-136 | 6.3226748 | Natural Killer T |
| 3 | GNLY | 2.16E-123 | 14.329076 | Natural Killer T |
| 4 | NKG7 | 9.70E-116 | 7.6357265 | Natural Killer T |
| 5 | GZMB | 6.98E-111 | 5.686702 | Natural Killer T |
| 6 | CD7 | 5.17E-103 | 4.8380857 | Natural Killer T |
| 7 | PTPRCAP | 4.46E-100 | 4.3205276 | Natural Killer T |
| 8 | KLRC2 | 2.71E-95 | 6.385977 | Natural Killer T |
| 9 | ZFP36L2 | 8.71E-92 | 6.712756 | Natural Killer T |
| 10 | CST7 | 3.29E-91 | 4.0231123 | Natural Killer T |
| 11 | CD2 | 1.02E-88 | 3.5549443 | Natural Killer T |
| 12 | KLRD1 | 2.25E-86 | 3.8931673 | Natural Killer T |
| 13 | GZMH | 2.28E-85 | 4.8087406 | Natural Killer T |
| 14 | ZNF683 | 7.38E-83 | 8.757736 | Natural Killer T |
| 15 | CORO1A | 2.00E-79 | 3.1603158 | Natural Killer T |
| 16 | BTG1 | 1.09E-78 | 6.415769 | Natural Killer T |
| 17 | PRF1 | 4.25E-72 | 3.2751362 | Natural Killer T |
| 18 | CD69 | 1.14E-68 | 2.4822862 | Natural Killer T |
| 19 | XCL1 | 1.06E-67 | 6.411995 | Natural Killer T |
| 20 | KLRC1 | 1.99E-67 | 5.557342 | Natural Killer T |
| 21 | KLRC3 | 1.16E-61 | 4.9790244 | Natural Killer T |
| 22 | TSC22D3 | 1.53E-60 | 5.118737 | Natural Killer T |
| 23 | RGS1 | 3.53E-59 | 3.7345815 | Natural Killer T |
| 24 | CD8A | 7.91E-59 | 3.970561 | Natural Killer T |
| 25 | ID2 | 1.08E-58 | 3.3254583 | Natural Killer T |
| 26 | XCL2 | 6.28E-55 | 5.428009 | Natural Killer T |
| 27 | LCK | 1.93E-54 | 3.1732671 | Natural Killer T |
| 28 | CD3E | 2.14E-52 | 2.9017184 | Natural Killer T |
| 29 | APOBEC3G | 1.56E-50 | 3.0940745 | Natural Killer T |
| 30 | C12orf75 | 2.75E-50 | 2.1736448 | Natural Killer T |
| 31 | HCST | 5.39E-47 | 2.0703735 | Natural Killer T |
| 32 | HOPX | 8.59E-45 | 1.4842005 | Natural Killer T |
| 33 | TUBA4A | 1.03E-42 | 2.5619423 | Natural Killer T |
| 34 | DUSP2 | 1.87E-39 | 2.9408674 | Natural Killer T |
| 35 | IFITM1 | 8.71E-39 | 2.2873368 | Natural Killer T |
| 36 | ACAP1 | 5.17E-34 | 2.5463622 | Natural Killer T |
| 37 | KRT86 | 7.08E-34 | 8.280742 | Natural Killer T |
| 38 | EMB | 9.80E-34 | 2.87086 | Natural Killer T |
| 39 | LIME1 | 1.08E-33 | 2.4720805 | Natural Killer T |
| 40 | GPR171 | 5.77E-33 | 3.478641 | Natural Killer T |
| 41 | KRT81 | 1.16E-32 | 9.179183 | Natural Killer T |
| 42 | PTPRC | 5.14E-32 | 1.4950739 | Natural Killer T |
| 43 | FCRL6 | 2.54E-31 | 3.7656906 | Natural Killer T |
| 44 | CD247 | 3.71E-31 | 1.9562509 | Natural Killer T |
| 45 | LSP1 | 4.74E-31 | 0.39106393 | Natural Killer T |
| 46 | CD48 | 1.88E-29 | 1.5973582 | Natural Killer T |
| 47 | IL2RG | 2.54E-27 | 1.2691541 | Natural Killer T |
| 48 | CLEC2D | 3.63E-27 | 2.3963969 | Natural Killer T |
| 49 | CD52 | 9.01E-27 | -11.163021 | Natural Killer T |
| 50 | CYTIP | 1.47E-24 | 1.2304543 | Natural Killer T |
| 51 | MATK | 1.71E-24 | 2.5739455 | Natural Killer T |
| 52 | IL2RB | 2.71E-24 | 3.1126807 | Natural Killer T |
| 53 | STK17A | 6.05E-22 | 1.6357352 | Natural Killer T |
| 54 | SAMSN1 | 1.50E-21 | 0.7085468 | Natural Killer T |
| 55 | BATF | 8.87E-20 | 1.9269923 | Natural Killer T |
| 56 | PRDM1 | 1.72E-19 | 1.908975 | Natural Killer T |
| 57 | IL32 | 1.17E-18 | 2.2595336 | Natural Killer T |
| 58 | ITGB2 | 2.19E-18 | 0.28606996 | Natural Killer T |

| | | | | |
|---|---|---|---|---|
| 59 | GZMM | 2.69E-18 | 1.8693279 | Natural Killer T |
| 60 | ANXA6 | 2.81E-18 | 1.5839664 | Natural Killer T |
| 61 | RAC2 | 9.35E-17 | 0.7703094 | Natural Killer T |
| 62 | KLRB1 | 1.16E-16 | 1.5596204 | Natural Killer T |
| 63 | GZMA | 1.36E-16 | 1.760219 | Natural Killer T |
| 64 | ICAM3 | 2.20E-16 | 1.3333458 | Natural Killer T |
| 65 | 01-Sep | 7.41E-16 | 1.9675606 | Natural Killer T |
| 66 | RASAL3 | 1.32E-15 | 2.0851178 | Natural Killer T |
| 67 | SOCS1 | 1.35E-15 | 1.4657812 | Natural Killer T |
| 68 | PARP8 | 4.82E-15 | 2.1246665 | Natural Killer T |
| 69 | IL10RA | 5.11E-15 | 1.3924222 | Natural Killer T |
| 70 | PIK3R1 | 1.34E-13 | 1.7819098 | Natural Killer T |
| 71 | RARRES3 | 1.62E-13 | 0.7202848 | Natural Killer T |
| 72 | ITGB7 | 3.87E-13 | 1.9942684 | Natural Killer T |
| 73 | SH2D2A | 1.38E-12 | 2.2954562 | Natural Killer T |
| 74 | LINC00152 | 1.47E-12 | 1.2096764 | Natural Killer T |
| 75 | PVRIG | 2.70E-12 | 2.2693176 | Natural Killer T |
| 76 | CD3D | 3.60E-11 | 2.7528603 | Natural Killer T |
| 77 | CD96 | 4.40E-11 | 2.744853 | Natural Killer T |
| 78 | CHST12 | 4.40E-11 | 1.2219677 | Natural Killer T |
| 79 | ZAP70 | 7.64E-11 | 2.1445913 | Natural Killer T |
| 80 | CXCR3 | 1.05E-10 | 2.6891544 | Natural Killer T |
| 81 | SKAP1 | 1.91E-10 | 1.9636146 | Natural Killer T |
| 82 | SYTL3 | 2.47E-10 | 1.7430327 | Natural Killer T |
| 83 | ITGA4 | 3.08E-10 | 1.7947041 | Natural Killer T |
| 84 | EVL | 4.55E-10 | -0.09595588 | Natural Killer T |
| 85 | CD53 | 5.22E-10 | -0.6145156 | Natural Killer T |
| 86 | SAMD3 | 5.95E-10 | 2.2843034 | Natural Killer T |
| 87 | PSTPIP1 | 6.31E-10 | 2.2191849 | Natural Killer T |
| 88 | PIK3IP1 | 7.46E-10 | 1.2270616 | Natural Killer T |
| 89 | KLRG1 | 1.05E-09 | 2.2938912 | Natural Killer T |
| 90 | TRAF3IP3 | 1.68E-09 | 1.6043581 | Natural Killer T |
| 91 | CD8B | 2.50E-09 | 2.8182979 | Natural Killer T |
| 92 | NCR3 | 2.78E-09 | 2.5566719 | Natural Killer T |
| 93 | SCML4 | 3.30E-09 | 3.5557036 | Natural Killer T |
| 94 | BIN2 | 4.69E-09 | 1.1529318 | Natural Killer T |
| 95 | HMGB2 | 4.83E-09 | 0.4672709 | Natural Killer T |
| 96 | SH2D1A | 5.82E-09 | 2.3185003 | Natural Killer T |
| 97 | FYB | 1.03E-08 | 0.6443375 | Natural Killer T |
| 98 | TNFRSF18 | 1.53E-08 | 3.2559552 | Natural Killer T |
| 99 | TXNIP | 1.88E-08 | 0.14388123 | Natural Killer T |
| 0 | PCSK1N | 0.000111343 | 92.59768 | Neuroendocrine |
| 1 | BEX1 | 0.000111343 | 18.258684 | Neuroendocrine |
| 2 | SEC11C | 0.000111343 | 124.51391 | Neuroendocrine |
| 3 | CPE | 0.000111343 | 40.830677 | Neuroendocrine |
| 4 | NGFRAP1 | 0.000470187 | 14.943043 | Neuroendocrine |
| 5 | PRDX2 | 0.000470187 | 9.295571 | Neuroendocrine |
| 6 | CHGA | 0.000470187 | 25.792416 | Neuroendocrine |
| 7 | SCGN | 0.000470187 | 23.510572 | Neuroendocrine |
| 8 | SCG2 | 0.000470187 | 40.251698 | Neuroendocrine |
| 9 | SCG5 | 0.000470187 | 26.300467 | Neuroendocrine |
| 10 | UCHL1 | 0.000470187 | 14.046902 | Neuroendocrine |
| 14 | NENF | 0.000588063 | 15.336387 | Neuroendocrine |
| 13 | SLC22A17 | 0.000588063 | 7.3625636 | Neuroendocrine |
| 11 | MEG3 | 0.000588063 | 19.914785 | Neuroendocrine |
| 12 | CIRBP | 0.000588063 | 25.278013 | Neuroendocrine |
| 15 | EIF4A2 | 0.00060334 | 14.873679 | Neuroendocrine |
| 16 | VAMP2 | 0.000933093 | 12.480721 | Neuroendocrine |
| 17 | MS4A8 | 0.000933093 | 6.0044646 | Neuroendocrine |
| 18 | EPCAM | 0.001300547 | 11.234935 | Neuroendocrine |
| 19 | CD24 | 0.001300547 | 10.068142 | Neuroendocrine |
| 20 | BCAM | 0.00156186 | 19.482826 | Neuroendocrine |
| 21 | EGR1 | 0.002045317 | 10.878331 | Neuroendocrine |
| 22 | ERI3 | 0.002192409 | 3.471735 | Neuroendocrine |

| | | | | |
|---|---|---|---|---|
| 23 | FAM46A | 0.002192409 | 5.4579 | Neuroendocrine |
| 24 | NPDC1 | 0.002266868 | 6.970928 | Neuroendocrine |
| 25 | NGRN | 0.002440807 | 5.268857 | Neuroendocrine |
| 26 | CHGB | 0.002462401 | 20.887959 | Neuroendocrine |
| 27 | GNG4 | 0.002462401 | 11.42048 | Neuroendocrine |
| 28 | APLP1 | 0.002462401 | 11.74736 | Neuroendocrine |
| 29 | DUSP26 | 0.002462401 | 9.764194 | Neuroendocrine |
| 30 | APP | 0.002462401 | 6.2157626 | Neuroendocrine |
| 31 | SPINT2 | 0.002617337 | 16.865509 | Neuroendocrine |
| 32 | ITM2C | 0.002617337 | 8.025199 | Neuroendocrine |
| 33 | PKIB | 0.002730899 | 9.342464 | Neuroendocrine |
| 34 | TCEAL2 | 0.002730899 | 8.478562 | Neuroendocrine |
| 35 | TAF7 | 0.003557604 | 3.7502606 | Neuroendocrine |
| 36 | BEX2 | 0.003605397 | 13.038429 | Neuroendocrine |
| 37 | MALAT1 | 0.003938853 | 582.77264 | Neuroendocrine |
| 38 | GABARAPL2 | 0.004140296 | 7.8953896 | Neuroendocrine |
| 39 | C6orf48 | 0.004140296 | 5.3003273 | Neuroendocrine |
| 40 | RASD1 | 0.004140296 | 9.28039 | Neuroendocrine |
| 41 | TMEM176B | 0.004140296 | 15.024147 | Neuroendocrine |
| 42 | TFF3 | 0.00418615 | 24.393894 | Neuroendocrine |
| 43 | TMEM176A | 0.004221073 | 12.247509 | Neuroendocrine |
| 44 | MAP1B | 0.004879107 | 6.3616385 | Neuroendocrine |
| 45 | DPP7 | 0.006043341 | 6.8013325 | Neuroendocrine |
| 46 | ATP6V0E2 | 0.006043341 | 6.2250233 | Neuroendocrine |
| 47 | EID1 | 0.006043341 | 7.415905 | Neuroendocrine |
| 48 | PFN2 | 0.006043341 | 5.4811254 | Neuroendocrine |
| 51 | PLEKHB1 | 0.006064833 | 5.3213415 | Neuroendocrine |
| 49 | CLU | 0.006064833 | 44.90582 | Neuroendocrine |
| 50 | FKBP2 | 0.006064833 | 8.718698 | Neuroendocrine |
| 52 | PTMS | 0.006359371 | 10.700574 | Neuroendocrine |
| 53 | ARL3 | 0.006438012 | 4.5228996 | Neuroendocrine |
| 54 | FXYD6 | 0.006713892 | 11.344936 | Neuroendocrine |
| 55 | C12orf57 | 0.00749397 | 5.7269845 | Neuroendocrine |
| 56 | KRT18 | 0.007509443 | 5.492534 | Neuroendocrine |
| 57 | RPL3 | 0.007509443 | 79.34537 | Neuroendocrine |
| 58 | HSPB1 | 0.008017635 | 31.26424 | Neuroendocrine |
| 59 | STARD10 | 0.008331928 | 4.751526 | Neuroendocrine |
| 60 | P4HTM | 0.010264784 | 5.284157 | Neuroendocrine |
| 61 | CAMLG | 0.010388273 | 4.1453285 | Neuroendocrine |
| 62 | STMN1 | 0.010388273 | 5.8296285 | Neuroendocrine |
| 73 | DIRAS3 | 0.010867875 | 40.084263 | Neuroendocrine |
| 72 | SCG3 | 0.010867875 | 12.444246 | Neuroendocrine |
| 71 | TAGLN3 | 0.010867875 | 13.044522 | Neuroendocrine |
| 70 | SYT4 | 0.010867875 | 10.352311 | Neuroendocrine |
| 69 | DDC | 0.010867875 | 14.164325 | Neuroendocrine |
| 74 | TUBB2B | 0.010867875 | 11.213155 | Neuroendocrine |
| 67 | SLC35D3 | 0.010867875 | 13.350196 | Neuroendocrine |
| 66 | CPLX2 | 0.010867875 | 15.519799 | Neuroendocrine |
| 65 | INA | 0.010867875 | 14.557177 | Neuroendocrine |
| 64 | TUSC3 | 0.010867875 | 5.4099555 | Neuroendocrine |
| 68 | PTPRN | 0.010867875 | 15.85744 | Neuroendocrine |
| 63 | ZBTB20 | 0.010867875 | 4.89378 | Neuroendocrine |
| 77 | MAGEF1 | 0.011480716 | 4.3955426 | Neuroendocrine |
| 75 | TSTD1 | 0.011480716 | 7.54707 | Neuroendocrine |
| 76 | TMEM14A | 0.011480716 | 4.5728464 | Neuroendocrine |
| 78 | PITX1 | 0.011568418 | 9.135858 | Neuroendocrine |
| 79 | IGFBP2 | 0.011741723 | 17.634024 | Neuroendocrine |
| 80 | FKBP8 | 0.012058909 | 5.244613 | Neuroendocrine |
| 81 | PNKD | 0.012058909 | 4.3293962 | Neuroendocrine |
| 82 | AHI1 | 0.012409175 | 3.9937127 | Neuroendocrine |
| 83 | EMC4 | 0.013323317 | 4.012028 | Neuroendocrine |
| 84 | SSR4 | 0.014091763 | 18.18389 | Neuroendocrine |
| 85 | KRT8 | 0.014500748 | 7.5062075 | Neuroendocrine |
| 86 | CADM1 | 0.014810979 | 4.6783056 | Neuroendocrine |

| | | | | |
|---|---|---|---|---|
| 87 | HAGH | 0.01522122 | 5.778144 | Neuroendocrine |
| 88 | HMGN3 | 0.01561599 | 6.644742 | Neuroendocrine |
| 89 | SFTPC | 0.015974158 | -194.53691 | Neuroendocrine |
| 90 | GPAA1 | 0.016316887 | 3.1806204 | Neuroendocrine |
| 91 | MIF | 0.016316887 | 19.227253 | Neuroendocrine |
| 92 | XIST | 0.016899582 | 7.3315887 | Neuroendocrine |
| 93 | LOC728392 | 0.01737004 | 8.827858 | Neuroendocrine |
| 94 | RABAC1 | 0.018334637 | 5.2733045 | Neuroendocrine |
| 95 | BAG1 | 0.018481191 | 5.278933 | Neuroendocrine |
| 96 | OCIAD2 | 0.021220282 | 5.5953374 | Neuroendocrine |
| 97 | DSP | 0.021403586 | 5.853502 | Neuroendocrine |
| 98 | SKP1 | 0.021405674 | 6.677549 | Neuroendocrine |
| 99 | PERP | 0.022281616 | 3.6627212 | Neuroendocrine |
| 0 | LST1 | 0 | 31.276834 | Nonclassical Monocyte |
| 12 | NAP1L1 | 0 | 9.214886 | Nonclassical Monocyte |
| 10 | C10orf54 | 0 | 6.614175 | Nonclassical Monocyte |
| 9 | FGL2 | 0 | 6.3425536 | Nonclassical Monocyte |
| 8 | IFITM2 | 0 | 16.931826 | Nonclassical Monocyte |
| 7 | CD48 | 0 | 6.745971 | Nonclassical Monocyte |
| 11 | TNFRSF1B | 0 | 5.472927 | Nonclassical Monocyte |
| 5 | LILRB2 | 0 | 8.986457 | Nonclassical Monocyte |
| 4 | FCN1 | 0 | 12.48771 | Nonclassical Monocyte |
| 3 | CORO1A | 0 | 9.225297 | Nonclassical Monocyte |
| 2 | FAM26F | 0 | 7.6476917 | Nonclassical Monocyte |
| 6 | CFP | 0 | 7.4696674 | Nonclassical Monocyte |
| 1 | COTL1 | 0 | 22.900227 | Nonclassical Monocyte |
| 13 | AIF1 | 9.82E-306 | 20.625446 | Nonclassical Monocyte |
| 14 | LILRA5 | 1.75E-298 | 6.5912876 | Nonclassical Monocyte |
| 15 | LINC01272 | 1.41E-297 | 7.9902577 | Nonclassical Monocyte |
| 16 | LIMD2 | 5.07E-295 | 5.53054 | Nonclassical Monocyte |
| 17 | LRRC25 | 7.60E-295 | 5.9351068 | Nonclassical Monocyte |
| 18 | FCGR3A | 4.13E-282 | 14.604664 | Nonclassical Monocyte |
| 19 | RGS2 | 5.71E-281 | 7.4545918 | Nonclassical Monocyte |
| 20 | RPS19 | 1.38E-254 | 45.826107 | Nonclassical Monocyte |
| 21 | MAFB | 1.55E-254 | 5.5916705 | Nonclassical Monocyte |
| 22 | LYN | 3.53E-238 | 4.3230777 | Nonclassical Monocyte |
| 23 | POU2F2 | 6.60E-233 | 5.5692825 | Nonclassical Monocyte |
| 24 | CTSS | 6.47E-230 | 14.670035 | Nonclassical Monocyte |
| 25 | TIMP1 | 2.28E-229 | 10.26797 | Nonclassical Monocyte |
| 26 | RNASET2 | 1.02E-228 | 5.11571 | Nonclassical Monocyte |
| 27 | NAAA | 3.28E-223 | 4.730754 | Nonclassical Monocyte |
| 28 | ABI3 | 9.29E-221 | 4.170163 | Nonclassical Monocyte |
| 29 | LILRA1 | 8.09E-220 | 7.705977 | Nonclassical Monocyte |
| 30 | FGR | 2.23E-219 | 4.1221685 | Nonclassical Monocyte |
| 31 | MPEG1 | 7.61E-218 | 4.2886157 | Nonclassical Monocyte |
| 32 | CSF1R | 8.79E-218 | 4.866049 | Nonclassical Monocyte |
| 33 | PTPRC | 2.09E-208 | 4.8759174 | Nonclassical Monocyte |
| 34 | STK17B | 6.06E-208 | 3.8297415 | Nonclassical Monocyte |
| 35 | SLC25A6 | 5.62E-202 | 7.4034195 | Nonclassical Monocyte |
| 36 | LYST | 9.14E-200 | 3.7567565 | Nonclassical Monocyte |
| 37 | C19orf38 | 1.75E-192 | 4.9849763 | Nonclassical Monocyte |
| 38 | CYTIP | 1.32E-188 | 4.2089443 | Nonclassical Monocyte |
| 39 | FCER1G | 1.08E-187 | 11.47885 | Nonclassical Monocyte |
| 40 | RAB24 | 2.77E-184 | 3.5193415 | Nonclassical Monocyte |
| 41 | SAT1 | 4.56E-184 | 22.409233 | Nonclassical Monocyte |
| 42 | EVI2B | 8.97E-184 | 3.7981539 | Nonclassical Monocyte |
| 43 | PSAP | 1.07E-182 | 10.433391 | Nonclassical Monocyte |
| 44 | FAM65B | 5.45E-182 | 3.8728545 | Nonclassical Monocyte |
| 45 | LSP1 | 1.17E-175 | 3.6836758 | Nonclassical Monocyte |
| 46 | AP1S2 | 2.71E-173 | 3.4905393 | Nonclassical Monocyte |
| 47 | PABPC1 | 3.05E-171 | 7.129283 | Nonclassical Monocyte |
| 48 | ICAM3 | 1.34E-167 | 3.5823517 | Nonclassical Monocyte |
| 49 | CFD | 1.40E-166 | 2.9387772 | Nonclassical Monocyte |
| 50 | CXCR4 | 1.62E-166 | 4.4852204 | Nonclassical Monocyte |

| | | | | |
|---|---|---|---|---|
| 51 | IFI30 | 4.47E-166 | 0.46674874 | Nonclassical Monocyte |
| 52 | CLEC7A | 6.37E-166 | 2.9127727 | Nonclassical Monocyte |
| 53 | SPI1 | 1.05E-164 | 2.8659706 | Nonclassical Monocyte |
| 54 | MS4A7 | 2.01E-162 | 3.2867644 | Nonclassical Monocyte |
| 55 | TYROBP | 1.32E-160 | 5.314 | Nonclassical Monocyte |
| 56 | ITGB2 | 5.94E-159 | 4.0284233 | Nonclassical Monocyte |
| 57 | SIGLEC10 | 4.33E-158 | 5.5968766 | Nonclassical Monocyte |
| 58 | NACA | 4.75E-158 | 9.6808405 | Nonclassical Monocyte |
| 59 | FYB | 1.15E-157 | 3.133022 | Nonclassical Monocyte |
| 60 | NEAT1 | 1.87E-156 | 9.569959 | Nonclassical Monocyte |
| 61 | ZFAND5 | 2.43E-155 | 3.5529144 | Nonclassical Monocyte |
| 62 | ACTB | 3.02E-155 | 40.043503 | Nonclassical Monocyte |
| 63 | PILRA | 8.19E-155 | 2.2618217 | Nonclassical Monocyte |
| 64 | STXBP2 | 1.23E-154 | 1.5955501 | Nonclassical Monocyte |
| 65 | SMAP2 | 2.28E-153 | 3.4483428 | Nonclassical Monocyte |
| 66 | PTPN6 | 3.28E-153 | 2.7936106 | Nonclassical Monocyte |
| 67 | TNFSF13B | 3.53E-153 | 3.0030756 | Nonclassical Monocyte |
| 68 | TYMP | 9.47E-151 | 3.5997753 | Nonclassical Monocyte |
| 69 | PRELID1 | 1.41E-150 | 3.752669 | Nonclassical Monocyte |
| 70 | CD37 | 2.69E-149 | 3.0103457 | Nonclassical Monocyte |
| 71 | APOBEC3A | 1.91E-148 | 6.3767843 | Nonclassical Monocyte |
| 72 | HCLS1 | 4.71E-147 | 2.6812823 | Nonclassical Monocyte |
| 73 | CD300A | 2.39E-146 | 3.7646306 | Nonclassical Monocyte |
| 74 | RHOG | 1.16E-145 | 2.7906525 | Nonclassical Monocyte |
| 75 | MTSS1 | 2.07E-145 | 3.979232 | Nonclassical Monocyte |
| 76 | S100A4 | 2.08E-144 | 7.4636984 | Nonclassical Monocyte |
| 77 | CST3 | 2.85E-144 | 8.149443 | Nonclassical Monocyte |
| 78 | GMFG | 7.01E-144 | 3.2735066 | Nonclassical Monocyte |
| 79 | IFITM3 | 2.12E-141 | 8.730392 | Nonclassical Monocyte |
| 80 | RGS18 | 4.11E-140 | 4.812952 | Nonclassical Monocyte |
| 81 | TKT | 1.87E-139 | 2.769743 | Nonclassical Monocyte |
| 82 | RHOC | 4.14E-138 | 3.9796586 | Nonclassical Monocyte |
| 83 | RPS9 | 1.30E-131 | 13.867625 | Nonclassical Monocyte |
| 84 | HCK | 3.41E-131 | 2.1794722 | Nonclassical Monocyte |
| 85 | MAPKAPK3 | 5.37E-131 | 2.963408 | Nonclassical Monocyte |
| 86 | PYCARD | 6.43E-131 | 1.3818523 | Nonclassical Monocyte |
| 87 | ADGRE2 | 1.44E-130 | 4.571729 | Nonclassical Monocyte |
| 88 | FTL | 6.27E-130 | -185.84064 | Nonclassical Monocyte |
| 89 | RGS19 | 1.02E-129 | 2.5051346 | Nonclassical Monocyte |
| 90 | RPL26 | 1.33E-129 | 15.521558 | Nonclassical Monocyte |
| 91 | UQCRB | 1.97E-129 | 5.045697 | Nonclassical Monocyte |
| 92 | CEBPB | 3.85E-129 | 5.1434402 | Nonclassical Monocyte |
| 93 | DRAP1 | 5.17E-128 | 2.5865102 | Nonclassical Monocyte |
| 94 | C1orf162 | 8.97E-128 | 1.6602525 | Nonclassical Monocyte |
| 95 | BCL2A1 | 1.11E-127 | 1.8227316 | Nonclassical Monocyte |
| 96 | LILRB1 | 1.53E-127 | 3.90085 | Nonclassical Monocyte |
| 97 | GPBAR1 | 3.29E-127 | 4.7608356 | Nonclassical Monocyte |
| 98 | ARHGDIB | 1.43E-126 | 4.3689733 | Nonclassical Monocyte |
| 99 | OAZ1 | 2.23E-126 | 2.9004138 | Nonclassical Monocyte |
| 0 | TIMP1 | 1.07E-105 | 84.37625 | OLR1+ Classical Monocyte |
| 1 | SOD2 | 2.21E-105 | 48.230003 | OLR1+ Classical Monocyte |
| 2 | FCN1 | 2.66E-100 | 11.965529 | OLR1+ Classical Monocyte |
| 3 | SERPINB9 | 5.12E-97 | 7.7798424 | OLR1+ Classical Monocyte |
| 4 | HIF1A | 4.36E-95 | 7.294676 | OLR1+ Classical Monocyte |
| 5 | VCAN | 1.31E-92 | 9.947903 | OLR1+ Classical Monocyte |
| 6 | CD300E | 7.72E-92 | 8.00127 | OLR1+ Classical Monocyte |
| 7 | ATP13A3 | 3.07E-91 | 6.1651783 | OLR1+ Classical Monocyte |
| 8 | IL1B | 6.24E-89 | 30.990505 | OLR1+ Classical Monocyte |
| 9 | NAMPT | 5.38E-88 | 15.382515 | OLR1+ Classical Monocyte |
| 10 | AQP9 | 2.97E-86 | 6.9703865 | OLR1+ Classical Monocyte |
| 11 | G0S2 | 3.50E-86 | 19.956139 | OLR1+ Classical Monocyte |
| 12 | MARCKS | 8.25E-86 | 9.192298 | OLR1+ Classical Monocyte |
| 13 | SRGN | 1.21E-85 | 65.663925 | OLR1+ Classical Monocyte |
| 14 | TNFRSF1B | 7.05E-84 | 7.6228085 | OLR1+ Classical Monocyte |

| | | | | |
|---|---|---|---|---|
| 15 | CD93 | 7.05E-84 | 5.2004347 | OLR1+ Classical Monocyte |
| 16 | TNFAIP3 | 2.93E-80 | 7.658799 | OLR1+ Classical Monocyte |
| 17 | C15orf48 | 3.54E-79 | 21.891575 | OLR1+ Classical Monocyte |
| 18 | PPIF | 3.79E-79 | 6.549402 | OLR1+ Classical Monocyte |
| 19 | S100A9 | 4.54E-77 | 73.72103 | OLR1+ Classical Monocyte |
| 20 | ACSL1 | 8.29E-76 | 6.202588 | OLR1+ Classical Monocyte |
| 21 | IER3 | 2.54E-75 | 7.72108 | OLR1+ Classical Monocyte |
| 22 | S100A8 | 2.81E-73 | 41.574745 | OLR1+ Classical Monocyte |
| 23 | TLR2 | 3.32E-72 | 5.0074863 | OLR1+ Classical Monocyte |
| 24 | BCL2A1 | 1.09E-71 | 9.091979 | OLR1+ Classical Monocyte |
| 25 | PFKFB3 | 1.01E-70 | 5.3198104 | OLR1+ Classical Monocyte |
| 26 | CD44 | 1.30E-69 | 11.639467 | OLR1+ Classical Monocyte |
| 27 | IFNGR2 | 1.79E-68 | 5.7626715 | OLR1+ Classical Monocyte |
| 28 | SAMSN1 | 2.20E-68 | 6.0153174 | OLR1+ Classical Monocyte |
| 29 | IL1RN | 2.08E-66 | 13.569891 | OLR1+ Classical Monocyte |
| 30 | LINC00152 | 1.17E-65 | 6.1453953 | OLR1+ Classical Monocyte |
| 31 | SLC25A37 | 4.18E-64 | 5.0260954 | OLR1+ Classical Monocyte |
| 32 | SLC43A2 | 4.60E-64 | 4.7689548 | OLR1+ Classical Monocyte |
| 33 | TNIP1 | 4.98E-63 | 4.851274 | OLR1+ Classical Monocyte |
| 34 | LILRB2 | 7.27E-63 | 5.40217 | OLR1+ Classical Monocyte |
| 35 | CD163 | 1.59E-61 | 6.4727964 | OLR1+ Classical Monocyte |
| 36 | NINJ1 | 9.30E-59 | 5.812452 | OLR1+ Classical Monocyte |
| 37 | PNRC1 | 9.70E-58 | 8.869752 | OLR1+ Classical Monocyte |
| 38 | S100A12 | 1.02E-56 | 7.5085716 | OLR1+ Classical Monocyte |
| 39 | NFKB1 | 1.09E-56 | 4.296232 | OLR1+ Classical Monocyte |
| 40 | WTAP | 3.30E-56 | 5.3710194 | OLR1+ Classical Monocyte |
| 41 | UPP1 | 3.23E-55 | 4.322301 | OLR1+ Classical Monocyte |
| 42 | GK | 6.30E-55 | 4.9307466 | OLR1+ Classical Monocyte |
| 43 | SAT1 | 7.42E-55 | 29.175945 | OLR1+ Classical Monocyte |
| 44 | MXD1 | 1.31E-53 | 5.222129 | OLR1+ Classical Monocyte |
| 45 | SPHK1 | 1.43E-53 | 5.1147857 | OLR1+ Classical Monocyte |
| 46 | CTSL | 2.34E-53 | 7.721184 | OLR1+ Classical Monocyte |
| 47 | LCP2 | 1.74E-49 | 4.220081 | OLR1+ Classical Monocyte |
| 48 | MIR4435-2HG | 2.30E-49 | 5.20935 | OLR1+ Classical Monocyte |
| 49 | CLEC4E | 2.57E-49 | 4.764018 | OLR1+ Classical Monocyte |
| 50 | VEGFA | 1.69E-48 | 4.8810525 | OLR1+ Classical Monocyte |
| 51 | DSE | 2.31E-48 | 3.631687 | OLR1+ Classical Monocyte |
| 52 | EHD1 | 2.26E-47 | 4.532378 | OLR1+ Classical Monocyte |
| 53 | TYMP | 4.38E-45 | 5.652867 | OLR1+ Classical Monocyte |
| 54 | APOBEC3A | 1.11E-44 | 7.6448264 | OLR1+ Classical Monocyte |
| 55 | NFKBIA | 1.50E-44 | 12.046293 | OLR1+ Classical Monocyte |
| 56 | LYZ | 3.46E-44 | 9.448557 | OLR1+ Classical Monocyte |
| 57 | FPR1 | 4.29E-44 | 4.1818395 | OLR1+ Classical Monocyte |
| 58 | CXCL8 | 7.49E-44 | 8.506725 | OLR1+ Classical Monocyte |
| 59 | LITAF | 9.08E-44 | 6.138665 | OLR1+ Classical Monocyte |
| 60 | CEBPB | 1.59E-43 | 8.993204 | OLR1+ Classical Monocyte |
| 61 | SLC2A3 | 2.77E-43 | 3.5087729 | OLR1+ Classical Monocyte |
| 62 | PLAUR | 3.27E-42 | 4.7477856 | OLR1+ Classical Monocyte |
| 63 | FTH1 | 2.48E-41 | 70.13752 | OLR1+ Classical Monocyte |
| 64 | BACH1 | 4.03E-41 | 3.624518 | OLR1+ Classical Monocyte |
| 65 | SLC7A5 | 4.27E-41 | 5.1269965 | OLR1+ Classical Monocyte |
| 66 | SLC39A8 | 9.59E-41 | 3.99655 | OLR1+ Classical Monocyte |
| 67 | FCER1G | 5.96E-40 | 6.943213 | OLR1+ Classical Monocyte |
| 68 | RAB31 | 5.47E-39 | 2.912832 | OLR1+ Classical Monocyte |
| 69 | ADAM19 | 4.85E-38 | 5.8252273 | OLR1+ Classical Monocyte |
| 70 | IVNS1ABP | 1.04E-37 | 3.2401137 | OLR1+ Classical Monocyte |
| 71 | EMP3 | 1.19E-36 | 4.729116 | OLR1+ Classical Monocyte |
| 72 | CCL20 | 4.70E-36 | -3.7133744 | OLR1+ Classical Monocyte |
| 73 | RGS2 | 7.30E-36 | 5.6378603 | OLR1+ Classical Monocyte |
| 74 | TNFSF8 | 1.82E-35 | 6.1416106 | OLR1+ Classical Monocyte |
| 75 | RNF19B | 2.46E-35 | 4.313494 | OLR1+ Classical Monocyte |
| 76 | CTSS | 2.84E-35 | 1.8799509 | OLR1+ Classical Monocyte |
| 77 | RIN2 | 7.07E-34 | 4.5062466 | OLR1+ Classical Monocyte |
| 78 | LCP1 | 7.69E-34 | 2.8001075 | OLR1+ Classical Monocyte |

| | | | | |
|---|---|---|---|---|
| 79 | LST1 | 1.07E-33 | 2.0419066 | OLR1+ Classical Monocyte |
| 80 | KYNU | 2.64E-33 | 2.4693997 | OLR1+ Classical Monocyte |
| 81 | ADGRE2 | 4.44E-33 | 4.303031 | OLR1+ Classical Monocyte |
| 82 | LINC01272 | 6.09E-33 | 2.6083095 | OLR1+ Classical Monocyte |
| 83 | TYROBP | 1.10E-32 | -3.5213215 | OLR1+ Classical Monocyte |
| 84 | THBS1 | 1.18E-32 | 8.207225 | OLR1+ Classical Monocyte |
| 85 | RPS26 | 1.18E-32 | 5.665729 | OLR1+ Classical Monocyte |
| 86 | AIF1 | 1.44E-32 | 1.1537079 | OLR1+ Classical Monocyte |
| 87 | JARID2 | 2.75E-32 | 4.513931 | OLR1+ Classical Monocyte |
| 88 | FCAR | 6.88E-32 | 5.4095774 | OLR1+ Classical Monocyte |
| 89 | CCL4L1 | 3.52E-31 | 10.815977 | OLR1+ Classical Monocyte |
| 90 | TPM4 | 5.58E-31 | 3.0745647 | OLR1+ Classical Monocyte |
| 91 | VASP | 6.18E-31 | 2.2175298 | OLR1+ Classical Monocyte |
| 92 | CD53 | 6.71E-31 | 2.267556 | OLR1+ Classical Monocyte |
| 93 | GNA15 | 9.22E-31 | 2.8659093 | OLR1+ Classical Monocyte |
| 94 | KDM6B | 3.04E-30 | 2.6898818 | OLR1+ Classical Monocyte |
| 95 | COTL1 | 3.11E-30 | 3.2051544 | OLR1+ Classical Monocyte |
| 96 | ATP5E | 3.67E-30 | 4.7839556 | OLR1+ Classical Monocyte |
| 97 | H3F3AP4 | 5.56E-30 | 2.5625885 | OLR1+ Classical Monocyte |
| 98 | ANPEP | 9.52E-30 | 2.596717 | OLR1+ Classical Monocyte |
| 99 | CSTA | 2.20E-29 | 1.2301171 | OLR1+ Classical Monocyte |
| 0 | COX4I2 | 0 | 12.255396 | Pericyte |
| 29 | TCF21 | 0 | 5.3775706 | Pericyte |
| 30 | HSPA2 | 0 | 5.635838 | Pericyte |
| 31 | KCNK3 | 0 | 8.4693775 | Pericyte |
| 32 | MEST | 0 | 6.6155224 | Pericyte |
| 33 | A2M | 0 | 3.1061015 | Pericyte |
| 34 | SPARCL1 | 0 | 3.3939767 | Pericyte |
| 35 | CADM1 | 0 | 4.295815 | Pericyte |
| 36 | SPARC | 0 | 3.7404313 | Pericyte |
| 37 | TGFB1I1 | 0 | 4.031652 | Pericyte |
| 38 | CEBPD | 0 | 9.439743 | Pericyte |
| 39 | SOD3 | 0 | 3.4535346 | Pericyte |
| 28 | RGN | 0 | 6.814736 | Pericyte |
| 40 | TACC1 | 0 | 3.854674 | Pericyte |
| 42 | PMP22 | 0 | 3.101492 | Pericyte |
| 43 | NOTCH3 | 0 | 5.608291 | Pericyte |
| 44 | ACTA2 | 0 | 3.6854553 | Pericyte |
| 45 | WFDC1 | 0 | 4.4644895 | Pericyte |
| 46 | COL1A2 | 0 | 3.3360624 | Pericyte |
| 47 | PCOLCE | 0 | 4.149474 | Pericyte |
| 48 | NDRG1 | 0 | 3.397621 | Pericyte |
| 50 | KCNK17 | 0 | 7.791259 | Pericyte |
| 51 | PAG1 | 0 | 3.3110278 | Pericyte |
| 52 | FAM105A | 0 | 3.7446575 | Pericyte |
| 53 | AGTR1 | 0 | 7.6304965 | Pericyte |
| 41 | HES4 | 0 | 3.5990388 | Pericyte |
| 27 | LAMC3 | 0 | 9.442597 | Pericyte |
| 49 | LAMB1 | 0 | 4.737743 | Pericyte |
| 25 | CHN1 | 0 | 6.5104513 | Pericyte |
| 1 | CALD1 | 0 | 14.024912 | Pericyte |
| 2 | BGN | 0 | 11.066929 | Pericyte |
| 3 | PDGFRB | 0 | 10.766837 | Pericyte |
| 4 | NDUFA4L2 | 0 | 10.844315 | Pericyte |
| 26 | MFGE8 | 0 | 4.1313396 | Pericyte |
| 5 | HIGD1B | 0 | 12.021802 | Pericyte |
| 6 | PTN | 0 | 10.702086 | Pericyte |
| 7 | EGFL6 | 0 | 9.014278 | Pericyte |
| 9 | IGFBP7 | 0 | 15.5876 | Pericyte |
| 10 | ITM2C | 0 | 5.869958 | Pericyte |
| 11 | TPM2 | 0 | 5.2643056 | Pericyte |
| 12 | PPP1R14A | 0 | 6.6993275 | Pericyte |
| 8 | MYL9 | 0 | 6.807132 | Pericyte |
| 14 | DCN | 0 | 3.3534353 | Pericyte |

| | | | | |
|---|---|---|---|---|
| 24 | GJA4 | 0 | 6.1960173 | Pericyte |
| 13 | LHFP | 0 | 5.982848 | Pericyte |
| 22 | COL6A2 | 0 | 4.330449 | Pericyte |
| 21 | SDC2 | 0 | 4.7830677 | Pericyte |
| 20 | TAGLN | 0 | 6.625836 | Pericyte |
| 23 | EFEMP1 | 0 | 4.6797667 | Pericyte |
| 18 | FAM162B | 0 | 9.58414 | Pericyte |
| 17 | GAS6 | 0 | 5.3457556 | Pericyte |
| 16 | GPX3 | 0 | 10.051005 | Pericyte |
| 15 | COL4A1 | 0 | 6.194925 | Pericyte |
| 19 | COL4A2 | 0 | 5.6634426 | Pericyte |
| 54 | F2R | 2.30E-300 | 4.020098 | Pericyte |
| 55 | SGCE | 2.53E-297 | 4.2348037 | Pericyte |
| 56 | ITGA1 | 1.26E-284 | 3.822866 | Pericyte |
| 57 | TPPP3 | 1.25E-273 | 0.36188135 | Pericyte |
| 58 | STOM | 2.50E-269 | 2.8196604 | Pericyte |
| 59 | CCDC102B | 1.49E-265 | 5.1142335 | Pericyte |
| 60 | TMEM204 | 1.41E-259 | 2.3590093 | Pericyte |
| 61 | CYGB | 4.67E-247 | 4.9047117 | Pericyte |
| 62 | TESC | 6.29E-246 | 4.1666446 | Pericyte |
| 63 | IGFBP2 | 4.10E-245 | 2.0349755 | Pericyte |
| 64 | NR2F2 | 8.87E-240 | 3.8303897 | Pericyte |
| 65 | ANXA6 | 3.72E-235 | 2.8262053 | Pericyte |
| 66 | CTGF | 2.02E-232 | 6.8036876 | Pericyte |
| 67 | MYO1B | 4.15E-227 | 3.3118744 | Pericyte |
| 68 | FXYD1 | 1.22E-219 | 3.3957937 | Pericyte |
| 69 | NID1 | 1.06E-216 | 5.370581 | Pericyte |
| 70 | DUSP4 | 1.27E-215 | 4.1280985 | Pericyte |
| 71 | PERP | 1.35E-215 | 1.3659239 | Pericyte |
| 72 | TBX5 | 4.83E-212 | 6.6098685 | Pericyte |
| 73 | RCN1 | 1.89E-211 | 2.7299416 | Pericyte |
| 74 | EHD2 | 2.48E-206 | 3.1485732 | Pericyte |
| 75 | DKK3 | 3.41E-206 | 3.2236137 | Pericyte |
| 76 | ID3 | 5.75E-200 | 4.4849052 | Pericyte |
| 77 | NDN | 1.67E-196 | 3.2702434 | Pericyte |
| 78 | SERPING1 | 8.13E-196 | -0.9691141 | Pericyte |
| 79 | ECM1 | 2.90E-193 | 4.495077 | Pericyte |
| 80 | APOE | 7.63E-190 | -8.367963 | Pericyte |
| 81 | LURAP1L | 2.48E-189 | 3.9143534 | Pericyte |
| 82 | RARRES2 | 1.62E-187 | 1.3768001 | Pericyte |
| 83 | UXS1 | 6.99E-187 | 2.3483772 | Pericyte |
| 84 | NEXN | 6.81E-186 | 3.728138 | Pericyte |
| 85 | RRAD | 1.37E-185 | 2.2426054 | Pericyte |
| 86 | RFTN1 | 2.07E-185 | 3.2194004 | Pericyte |
| 87 | PHLDA1 | 8.42E-183 | 2.2510204 | Pericyte |
| 88 | P2RY14 | 2.59E-178 | 5.035956 | Pericyte |
| 89 | F10 | 5.56E-176 | 4.552128 | Pericyte |
| 90 | FERMT2 | 4.85E-174 | 2.8787982 | Pericyte |
| 91 | COL6A1 | 1.73E-172 | 2.7645652 | Pericyte |
| 92 | EFEMP2 | 2.52E-172 | 2.9963193 | Pericyte |
| 93 | GUCY1B3 | 6.77E-170 | 5.183058 | Pericyte |
| 94 | SPRY1 | 6.56E-168 | 2.4322798 | Pericyte |
| 95 | IMPA2 | 6.05E-165 | 2.801944 | Pericyte |
| 96 | WLS | 1.39E-161 | 2.5413034 | Pericyte |
| 97 | PLAC9 | 7.33E-158 | 2.111845 | Pericyte |
| 98 | MXRA8 | 7.94E-158 | 3.7289255 | Pericyte |
| 99 | RGS5 | 1.79E-157 | 3.6995094 | Pericyte |
| 0 | MZB1 | 7.10E-118 | 79.88904 | Plasma |
| 1 | SSR4 | 9.84E-117 | 91.99019 | Plasma |
| 2 | IGLL5 | 1.22E-115 | inf | Plasma |
| 3 | JCHAIN | 1.63E-115 | inf | Plasma |
| 4 | FKBP11 | 1.03E-113 | 25.423082 | Plasma |
| 5 | HERPUD1 | 1.62E-109 | 51.794426 | Plasma |
| 6 | DERL3 | 2.34E-109 | 29.850138 | Plasma |

| | | | | |
|---|---|---|---|---|
| 7 | ITM2C | 2.66E-109 | 26.292519 | Plasma |
| 8 | CD79A | 9.18E-107 | 23.412066 | Plasma |
| 9 | SEC11C | 3.53E-103 | 20.469585 | Plasma |
| 10 | CD27 | 6.50E-101 | 18.996626 | Plasma |
| 11 | HSP90B1 | 1.37E-98 | 37.07416 | Plasma |
| 12 | PIM2 | 8.57E-90 | 12.850421 | Plasma |
| 13 | FKBP2 | 4.13E-86 | 12.304888 | Plasma |
| 14 | MANF | 1.52E-85 | 9.742978 | Plasma |
| 15 | DNAJB9 | 1.21E-83 | 9.030061 | Plasma |
| 16 | SPCS2 | 3.76E-83 | 12.297173 | Plasma |
| 17 | GNG7 | 5.22E-83 | 8.592624 | Plasma |
| 18 | PDIA6 | 1.68E-82 | 11.46163 | Plasma |
| 19 | SPCS1 | 2.57E-81 | 15.498479 | Plasma |
| 20 | CRELD2 | 3.42E-79 | 7.5458903 | Plasma |
| 21 | TNFRSF13B | 7.16E-78 | 12.156593 | Plasma |
| 22 | PRDX4 | 3.90E-77 | 12.506701 | Plasma |
| 23 | TNFRSF17 | 5.19E-76 | 13.655764 | Plasma |
| 24 | XBP1 | 6.90E-76 | 22.203516 | Plasma |
| 25 | TPD52 | 7.18E-76 | 5.667745 | Plasma |
| 26 | TSC22D3 | 8.77E-76 | 41.911568 | Plasma |
| 27 | POU2AF1 | 2.13E-74 | 9.715965 | Plasma |
| 28 | KRTCAP2 | 9.45E-74 | 10.69396 | Plasma |
| 29 | UBE2J1 | 3.11E-73 | 6.9229455 | Plasma |
| 30 | ISG20 | 8.01E-73 | 7.458499 | Plasma |
| 31 | SPCS3 | 1.03E-70 | 7.9640675 | Plasma |
| 32 | SDF2L1 | 2.27E-70 | 7.2099442 | Plasma |
| 33 | ICAM3 | 8.39E-70 | 5.8198047 | Plasma |
| 34 | IL2RG | 1.20E-69 | 7.6030145 | Plasma |
| 35 | LIME1 | 1.70E-69 | 5.9134936 | Plasma |
| 36 | VIMP | 1.98E-69 | 9.101904 | Plasma |
| 37 | RABAC1 | 1.36E-68 | 8.476075 | Plasma |
| 38 | SLAMF7 | 7.48E-68 | 7.019144 | Plasma |
| 39 | SSR3 | 8.94E-68 | 7.5202346 | Plasma |
| 40 | TXNDC15 | 4.13E-67 | 6.3106346 | Plasma |
| 41 | FAM46C | 7.77E-67 | 6.636148 | Plasma |
| 42 | TXNDC11 | 2.48E-66 | 6.489062 | Plasma |
| 43 | SELK | 3.35E-66 | 8.388248 | Plasma |
| 44 | MEI1 | 7.08E-66 | 7.338681 | Plasma |
| 45 | SUB1 | 1.57E-65 | 13.041314 | Plasma |
| 46 | TMEM258 | 2.85E-64 | 9.804365 | Plasma |
| 47 | SELM | 1.05E-63 | 7.3098693 | Plasma |
| 48 | SEC61B | 3.18E-63 | 9.570222 | Plasma |
| 49 | PPAPDC1B | 6.00E-63 | 5.6005387 | Plasma |
| 50 | PABPC4 | 1.86E-62 | 5.456978 | Plasma |
| 51 | RPL3 | 5.99E-62 | 46.769325 | Plasma |
| 52 | ANKRD36BP2 | 1.27E-60 | 10.968892 | Plasma |
| 53 | BIRC3 | 1.53E-60 | 10.229354 | Plasma |
| 54 | SDC1 | 3.91E-60 | 6.306828 | Plasma |
| 55 | TRAM1 | 1.89E-59 | 5.712494 | Plasma |
| 56 | RPS4X | 2.77E-58 | 47.355602 | Plasma |
| 57 | MYDGF | 1.00E-57 | 8.14349 | Plasma |
| 58 | LMAN2 | 1.52E-57 | 6.527189 | Plasma |
| 59 | CCDC167 | 1.92E-57 | 5.742358 | Plasma |
| 60 | DDOST | 5.64E-57 | 5.602856 | Plasma |
| 61 | HSH2D | 6.95E-57 | 5.073552 | Plasma |
| 62 | RPLP1 | 1.22E-56 | 81.37304 | Plasma |
| 63 | TMED9 | 7.53E-56 | 6.9885736 | Plasma |
| 64 | ERLEC1 | 7.70E-56 | 5.2486763 | Plasma |
| 65 | NUCB2 | 9.35E-56 | 5.923337 | Plasma |
| 66 | CUTA | 3.41E-55 | 5.4596043 | Plasma |
| 67 | ISCU | 3.78E-55 | 7.08474 | Plasma |
| 68 | PDIA4 | 6.92E-53 | 5.100588 | Plasma |
| 69 | SSR2 | 1.97E-52 | 6.0699754 | Plasma |
| 70 | PPIB | 1.15E-51 | 8.676759 | Plasma |

| | | | | |
|---|---|---|---|---|
| 71 | TMED10 | 3.40E-50 | 6.6232686 | Plasma |
| 72 | SEL1L3 | 4.25E-50 | 5.520777 | Plasma |
| 73 | HM13 | 4.72E-50 | 4.3728538 | Plasma |
| 74 | SLC25A6 | 1.29E-48 | 9.741176 | Plasma |
| 75 | TMEM59 | 1.59E-48 | 10.125816 | Plasma |
| 76 | RPL10 | 2.20E-48 | 81.84928 | Plasma |
| 77 | RGS1 | 4.43E-48 | 15.16377 | Plasma |
| 78 | EEF2 | 8.19E-48 | 11.688599 | Plasma |
| 79 | OST4 | 2.34E-47 | 7.298562 | Plasma |
| 80 | CYTIP | 3.27E-47 | 4.576633 | Plasma |
| 81 | RPLP0 | 4.14E-47 | 17.368258 | Plasma |
| 82 | SPAG4 | 4.97E-47 | 8.934663 | Plasma |
| 83 | HSPA5 | 1.44E-46 | 7.4862537 | Plasma |
| 84 | P4HB | 2.41E-46 | 4.704418 | Plasma |
| 85 | EEF1G | 3.81E-46 | 17.18819 | Plasma |
| 86 | SERP1 | 1.38E-45 | 6.073409 | Plasma |
| 87 | TMED2 | 2.34E-45 | 4.8866262 | Plasma |
| 88 | RPN2 | 7.45E-45 | 5.9399343 | Plasma |
| 89 | CHPF | 2.55E-42 | 4.956937 | Plasma |
| 90 | PLP2 | 2.64E-42 | 4.1070633 | Plasma |
| 91 | RPL7A | 6.12E-42 | 17.342646 | Plasma |
| 92 | ERP29 | 6.65E-42 | 5.020097 | Plasma |
| 93 | TP53INP1 | 6.65E-42 | 5.156942 | Plasma |
| 94 | ERGIC3 | 6.97E-42 | 4.560366 | Plasma |
| 95 | RPS14 | 8.43E-42 | 38.36256 | Plasma |
| 96 | CFLAR | 2.37E-41 | 5.02955 | Plasma |
| 97 | KDELR2 | 1.95E-40 | 4.270296 | Plasma |
| 98 | CYBA | 3.15E-40 | 16.34019 | Plasma |
| 99 | LMAN1 | 3.57E-40 | 3.8348298 | Plasma |
| 0 | IRF7 | 1.31E-80 | 16.850393 | Plasmacytoid Dendritic |
| 1 | GZMB | 1.90E-78 | 61.799934 | Plasmacytoid Dendritic |
| 2 | ITM2C | 7.05E-78 | 14.076507 | Plasmacytoid Dendritic |
| 3 | PPP1R14B | 1.66E-77 | 11.603047 | Plasmacytoid Dendritic |
| 4 | JCHAIN | 7.13E-77 | 14.804726 | Plasmacytoid Dendritic |
| 5 | LILRA4 | 7.35E-73 | 11.674637 | Plasmacytoid Dendritic |
| 6 | PLD4 | 2.94E-72 | 15.278667 | Plasmacytoid Dendritic |
| 7 | IRF8 | 1.18E-70 | 13.095418 | Plasmacytoid Dendritic |
| 8 | C12orf75 | 1.18E-70 | 9.7238 | Plasmacytoid Dendritic |
| 9 | MZB1 | 2.89E-70 | 7.9595714 | Plasmacytoid Dendritic |
| 10 | GPR183 | 9.39E-70 | 14.576397 | Plasmacytoid Dendritic |
| 11 | SERPINF1 | 6.66E-65 | 8.259203 | Plasmacytoid Dendritic |
| 12 | PLAC8 | 2.31E-63 | 15.364309 | Plasmacytoid Dendritic |
| 13 | SPIB | 4.21E-63 | 10.147239 | Plasmacytoid Dendritic |
| 14 | CXCR3 | 5.25E-63 | 11.462083 | Plasmacytoid Dendritic |
| 15 | TSPAN13 | 1.43E-62 | 8.10072 | Plasmacytoid Dendritic |
| 16 | CLIC3 | 2.71E-62 | 12.16828 | Plasmacytoid Dendritic |
| 17 | MAP1A | 6.83E-58 | 7.359739 | Plasmacytoid Dendritic |
| 18 | SEC61B | 7.19E-58 | 11.746684 | Plasmacytoid Dendritic |
| 19 | NAPSB | 1.60E-57 | 9.126286 | Plasmacytoid Dendritic |
| 20 | HERPUD1 | 2.15E-57 | 10.257376 | Plasmacytoid Dendritic |
| 21 | LIME1 | 4.62E-56 | 5.85146 | Plasmacytoid Dendritic |
| 22 | CXCR4 | 2.19E-55 | 12.009237 | Plasmacytoid Dendritic |
| 23 | PLP2 | 1.31E-54 | 7.0332966 | Plasmacytoid Dendritic |
| 24 | TCF4 | 1.32E-54 | 6.1899643 | Plasmacytoid Dendritic |
| 25 | CLN8 | 2.83E-54 | 7.643387 | Plasmacytoid Dendritic |
| 26 | SELL | 1.06E-53 | 6.4126368 | Plasmacytoid Dendritic |
| 27 | RNASE6 | 3.01E-53 | 6.333935 | Plasmacytoid Dendritic |
| 28 | SCT | 3.52E-52 | 12.76508 | Plasmacytoid Dendritic |
| 29 | DERL3 | 7.99E-52 | 7.541924 | Plasmacytoid Dendritic |
| 30 | RNASET2 | 5.41E-51 | 6.8321595 | Plasmacytoid Dendritic |
| 31 | IRF4 | 1.61E-49 | 9.210438 | Plasmacytoid Dendritic |
| 32 | UGCG | 9.76E-49 | 5.10557 | Plasmacytoid Dendritic |
| 33 | C9orf142 | 2.10E-47 | 5.62866 | Plasmacytoid Dendritic |
| 34 | CYB561A3 | 4.89E-47 | 6.0569835 | Plasmacytoid Dendritic |

| | | | | |
|---|---|---|---|---|
| 35 | LILRB4 | 1.69E-46 | 5.5802355 | Plasmacytoid Dendritic |
| 36 | IL3RA | 3.82E-46 | 5.1905475 | Plasmacytoid Dendritic |
| 37 | RPS3A | 3.90E-46 | 38.360863 | Plasmacytoid Dendritic |
| 38 | CCDC50 | 9.32E-46 | 6.0277953 | Plasmacytoid Dendritic |
| 39 | SMPD3 | 9.32E-46 | 8.8252325 | Plasmacytoid Dendritic |
| 40 | RPS8 | 9.55E-46 | 29.549099 | Plasmacytoid Dendritic |
| 41 | PTCRA | 1.79E-45 | 6.8748426 | Plasmacytoid Dendritic |
| 42 | MPEG1 | 3.94E-45 | 6.136932 | Plasmacytoid Dendritic |
| 43 | BCL11A | 5.23E-45 | 7.6354938 | Plasmacytoid Dendritic |
| 44 | IGFLR1 | 2.41E-44 | 4.8921685 | Plasmacytoid Dendritic |
| 45 | LRRC26 | 2.12E-43 | 8.04582 | Plasmacytoid Dendritic |
| 46 | RPS23 | 2.61E-43 | 30.519588 | Plasmacytoid Dendritic |
| 47 | UCP2 | 1.46E-42 | 5.9252467 | Plasmacytoid Dendritic |
| 48 | PTPRCAP | 2.48E-42 | 4.70465 | Plasmacytoid Dendritic |
| 49 | CLEC4C | 3.86E-41 | 12.886507 | Plasmacytoid Dendritic |
| 50 | DDIT4 | 3.69E-39 | 4.980178 | Plasmacytoid Dendritic |
| 51 | TRAF4 | 7.18E-38 | 5.1455283 | Plasmacytoid Dendritic |
| 52 | CORO1A | 1.77E-37 | 5.259588 | Plasmacytoid Dendritic |
| 53 | LTB | 7.32E-37 | 8.753316 | Plasmacytoid Dendritic |
| 54 | CNPY3 | 1.21E-36 | 3.6789944 | Plasmacytoid Dendritic |
| 55 | RPLP0 | 1.81E-35 | 12.902682 | Plasmacytoid Dendritic |
| 56 | CD4 | 2.11E-35 | 3.420838 | Plasmacytoid Dendritic |
| 57 | COMMD6 | 5.85E-35 | 5.383581 | Plasmacytoid Dendritic |
| 58 | RPL13 | 6.83E-35 | 35.410152 | Plasmacytoid Dendritic |
| 59 | RPL6 | 1.31E-34 | 19.060156 | Plasmacytoid Dendritic |
| 60 | RPS28 | 1.44E-34 | 26.696918 | Plasmacytoid Dendritic |
| 61 | RPL3 | 1.53E-34 | 23.248543 | Plasmacytoid Dendritic |
| 62 | GAPT | 2.17E-34 | 5.918047 | Plasmacytoid Dendritic |
| 63 | RPL13A | 4.26E-34 | 34.269253 | Plasmacytoid Dendritic |
| 64 | RPS2 | 6.76E-34 | 32.555973 | Plasmacytoid Dendritic |
| 65 | PTPRS | 3.06E-33 | 7.0749216 | Plasmacytoid Dendritic |
| 66 | PRKCB | 3.56E-33 | 4.4861655 | Plasmacytoid Dendritic |
| 67 | SPCS1 | 4.67E-33 | 4.8207564 | Plasmacytoid Dendritic |
| 68 | PTPRE | 6.89E-33 | 4.5464582 | Plasmacytoid Dendritic |
| 69 | RPS12 | 8.71E-33 | 32.43596 | Plasmacytoid Dendritic |
| 70 | P2RY14 | 1.81E-32 | 5.9675803 | Plasmacytoid Dendritic |
| 71 | RPS11 | 1.85E-32 | 14.2111025 | Plasmacytoid Dendritic |
| 72 | RPL23A | 2.61E-32 | 22.205273 | Plasmacytoid Dendritic |
| 73 | EEF1B2 | 3.34E-32 | 7.1927047 | Plasmacytoid Dendritic |
| 74 | IRF2BP2 | 4.94E-32 | 3.6037574 | Plasmacytoid Dendritic |
| 75 | EEF1G | 9.80E-32 | 11.977186 | Plasmacytoid Dendritic |
| 76 | SLC7A5P2 | 2.07E-31 | 5.4154687 | Plasmacytoid Dendritic |
| 77 | DNASE1L3 | 3.70E-31 | 4.8921556 | Plasmacytoid Dendritic |
| 78 | RPS18 | 1.21E-30 | 31.679577 | Plasmacytoid Dendritic |
| 79 | SMIM3 | 1.21E-30 | 4.34047 | Plasmacytoid Dendritic |
| 80 | RPS4X | 1.33E-30 | 21.563465 | Plasmacytoid Dendritic |
| 81 | RPL23 | 1.79E-30 | 10.391322 | Plasmacytoid Dendritic |
| 82 | RPS7 | 4.68E-30 | 13.569544 | Plasmacytoid Dendritic |
| 83 | TPM2 | 6.77E-30 | 3.1006172 | Plasmacytoid Dendritic |
| 84 | RPS6 | 8.26E-30 | 22.566916 | Plasmacytoid Dendritic |
| 85 | DCK | 1.11E-29 | 3.8179543 | Plasmacytoid Dendritic |
| 86 | RPL10A | 1.56E-29 | 13.489942 | Plasmacytoid Dendritic |
| 87 | FAM129C | 1.81E-29 | 8.466027 | Plasmacytoid Dendritic |
| 88 | PARK7 | 2.77E-29 | 3.9229603 | Plasmacytoid Dendritic |
| 89 | SLC15A4 | 6.22E-29 | 4.5847044 | Plasmacytoid Dendritic |
| 90 | RPS13 | 6.94E-29 | 19.133633 | Plasmacytoid Dendritic |
| 91 | RPL4 | 9.07E-29 | 8.579022 | Plasmacytoid Dendritic |
| 92 | MYBL2 | 9.75E-29 | 8.608915 | Plasmacytoid Dendritic |
| 93 | LAMP5 | 1.03E-28 | 8.885973 | Plasmacytoid Dendritic |
| 94 | ERP29 | 1.62E-28 | 3.5605886 | Plasmacytoid Dendritic |
| 95 | CBX6 | 1.90E-28 | 3.5542548 | Plasmacytoid Dendritic |
| 96 | SNHG5 | 2.03E-28 | 6.484496 | Plasmacytoid Dendritic |
| 97 | RPS27A | 2.29E-28 | 24.187225 | Plasmacytoid Dendritic |
| 98 | RPL37 | 2.29E-28 | 16.762392 | Plasmacytoid Dendritic |

| | | | | |
|---|---|---|---|---|
| 99 | SSR4 | 3.85E-28 | 4.3882165 | Plasmacytoid Dendritic |
| 0 | MAX | 3.38E-18 | 8.77804 | Platelet/Megakaryocyte |
| 1 | RGS10 | 1.31E-14 | 10.785911 | Platelet/Megakaryocyte |
| 2 | LIMS1 | 1.38E-14 | 8.36885 | Platelet/Megakaryocyte |
| 3 | RGS18 | 4.13E-14 | 18.333527 | Platelet/Megakaryocyte |
| 4 | CLU | 1.77E-13 | 12.300729 | Platelet/Megakaryocyte |
| 5 | CTTN | 1.77E-13 | 6.2218337 | Platelet/Megakaryocyte |
| 6 | PGRMC1 | 2.67E-13 | 5.219046 | Platelet/Megakaryocyte |
| 7 | ODC1 | 2.67E-13 | 6.422462 | Platelet/Megakaryocyte |
| 8 | GPX1 | 3.42E-13 | 27.792986 | Platelet/Megakaryocyte |
| 14 | PF4 | 4.56E-13 | 57.84415 | Platelet/Megakaryocyte |
| 13 | TREML1 | 4.56E-13 | 12.816275 | Platelet/Megakaryocyte |
| 12 | TUBB1 | 4.56E-13 | 22.257929 | Platelet/Megakaryocyte |
| 11 | GP9 | 4.56E-13 | 15.679065 | Platelet/Megakaryocyte |
| 10 | RUFY1 | 4.56E-13 | 7.2926917 | Platelet/Megakaryocyte |
| 9 | FERMT3 | 4.56E-13 | 6.390036 | Platelet/Megakaryocyte |
| 15 | PPBP | 6.07E-13 | 97.04555 | Platelet/Megakaryocyte |
| 16 | TMEM40 | 6.07E-13 | 10.59624 | Platelet/Megakaryocyte |
| 17 | TPM4 | 8.12E-13 | 9.028777 | Platelet/Megakaryocyte |
| 18 | NCOA4 | 1.14E-12 | 9.845521 | Platelet/Megakaryocyte |
| 19 | GRAP2 | 1.54E-12 | 9.729382 | Platelet/Megakaryocyte |
| 20 | NAP1L1 | 1.57E-12 | 8.827285 | Platelet/Megakaryocyte |
| 21 | PTPN18 | 3.25E-12 | 4.5711026 | Platelet/Megakaryocyte |
| 22 | TUBA4A | 3.99E-12 | 10.61509 | Platelet/Megakaryocyte |
| 23 | PTCRA | 5.19E-12 | 10.284495 | Platelet/Megakaryocyte |
| 24 | NT5C3A | 5.20E-12 | 6.9223113 | Platelet/Megakaryocyte |
| 25 | RAP1B | 5.27E-12 | 12.4147835 | Platelet/Megakaryocyte |
| 26 | NRGN | 5.58E-12 | 10.558661 | Platelet/Megakaryocyte |
| 27 | CTSA | 8.11E-12 | 6.476782 | Platelet/Megakaryocyte |
| 28 | MAP3K7CL | 1.26E-11 | 7.420743 | Platelet/Megakaryocyte |
| 29 | ACRBP | 2.55E-11 | 17.623415 | Platelet/Megakaryocyte |
| 30 | PRKAR2B | 3.24E-11 | 5.663494 | Platelet/Megakaryocyte |
| 31 | TLN1 | 4.17E-11 | 4.7738056 | Platelet/Megakaryocyte |
| 32 | KIF2A | 5.47E-11 | 5.916847 | Platelet/Megakaryocyte |
| 33 | CALM3 | 6.34E-11 | 5.8097463 | Platelet/Megakaryocyte |
| 34 | DAPP1 | 7.54E-11 | 4.998682 | Platelet/Megakaryocyte |
| 35 | MPP1 | 8.18E-11 | 6.0365877 | Platelet/Megakaryocyte |
| 36 | 02-Mar | 1.01E-10 | 5.3631043 | Platelet/Megakaryocyte |
| 37 | CA2 | 1.51E-10 | 5.555574 | Platelet/Megakaryocyte |
| 38 | CCL5 | 1.92E-10 | 11.186448 | Platelet/Megakaryocyte |
| 39 | PLA2G12A | 2.18E-10 | 4.999792 | Platelet/Megakaryocyte |
| 40 | DMTN | 3.48E-10 | 7.9203234 | Platelet/Megakaryocyte |
| 41 | GMPR | 4.22E-10 | 5.336977 | Platelet/Megakaryocyte |
| 42 | CMTM5 | 7.35E-10 | 14.422548 | Platelet/Megakaryocyte |
| 43 | MYL9 | 8.28E-10 | 3.7492437 | Platelet/Megakaryocyte |
| 44 | PTGS1 | 1.22E-09 | 5.8229094 | Platelet/Megakaryocyte |
| 45 | MMD | 1.29E-09 | 8.187517 | Platelet/Megakaryocyte |
| 46 | ETFA | 1.36E-09 | 3.6284437 | Platelet/Megakaryocyte |
| 47 | SNAP23 | 1.88E-09 | 4.0236254 | Platelet/Megakaryocyte |
| 48 | GNG11 | 2.08E-09 | 15.99963 | Platelet/Megakaryocyte |
| 49 | OST4 | 2.77E-09 | 7.856174 | Platelet/Megakaryocyte |
| 50 | GAS2L1 | 3.86E-09 | 3.9285562 | Platelet/Megakaryocyte |
| 51 | YWHAH | 3.94E-09 | 6.9307685 | Platelet/Megakaryocyte |
| 52 | SNCA | 4.21E-09 | 6.1415625 | Platelet/Megakaryocyte |
| 53 | CXCR2P1 | 4.24E-09 | 10.679381 | Platelet/Megakaryocyte |
| 54 | ITGA2B | 4.26E-09 | 11.2292 | Platelet/Megakaryocyte |
| 55 | C2orf88 | 5.54E-09 | 9.521566 | Platelet/Megakaryocyte |
| 56 | VIM-AS1 | 6.64E-09 | 4.236202 | Platelet/Megakaryocyte |
| 57 | F13A1 | 6.92E-09 | 8.271036 | Platelet/Megakaryocyte |
| 58 | R3HDM4 | 1.15E-08 | 3.7877886 | Platelet/Megakaryocyte |
| 59 | OAZ1 | 1.30E-08 | 21.991852 | Platelet/Megakaryocyte |
| 60 | MYL12A | 1.85E-08 | 12.191288 | Platelet/Megakaryocyte |
| 61 | CLEC1B | 2.14E-08 | 13.214213 | Platelet/Megakaryocyte |
| 62 | H3F3AP4 | 3.18E-08 | 3.9377725 | Platelet/Megakaryocyte |

| | | | | |
|---|---|---|---|---|
| 63 | SPARC | 3.60E-08 | 5.1605477 | Platelet/Megakaryocyte |
| 64 | RAB27B | 4.05E-08 | 5.3303523 | Platelet/Megakaryocyte |
| 65 | C9orf89 | 5.66E-08 | 4.397399 | Platelet/Megakaryocyte |
| 66 | NGFRAP1 | 1.31E-07 | 4.169204 | Platelet/Megakaryocyte |
| 67 | BIN2 | 1.43E-07 | 3.8095584 | Platelet/Megakaryocyte |
| 68 | LAT | 1.43E-07 | 5.1622 | Platelet/Megakaryocyte |
| 69 | GP1BA | 1.50E-07 | 8.3686905 | Platelet/Megakaryocyte |
| 70 | SDPR | 1.59E-07 | 14.332419 | Platelet/Megakaryocyte |
| 71 | FAM110A | 1.83E-07 | 4.383746 | Platelet/Megakaryocyte |
| 72 | EIF2AK1 | 2.88E-07 | 3.9633389 | Platelet/Megakaryocyte |
| 73 | SLC40A1 | 3.12E-07 | 4.7850666 | Platelet/Megakaryocyte |
| 74 | RNF11 | 3.52E-07 | 4.8742414 | Platelet/Megakaryocyte |
| 75 | TGFB1 | 4.14E-07 | 2.9347265 | Platelet/Megakaryocyte |
| 76 | TAGLN2 | 6.78E-07 | 10.361161 | Platelet/Megakaryocyte |
| 77 | HRAT92 | 7.03E-07 | 9.727798 | Platelet/Megakaryocyte |
| 78 | TSC22D1 | 1.03E-06 | 5.826267 | Platelet/Megakaryocyte |
| 79 | PIP4K2A | 1.30E-06 | 3.229347 | Platelet/Megakaryocyte |
| 80 | C19orf33 | 1.37E-06 | 4.6700587 | Platelet/Megakaryocyte |
| 81 | SH3BGRL2 | 1.45E-06 | 5.077546 | Platelet/Megakaryocyte |
| 82 | CCDC85B | 1.47E-06 | 2.7516296 | Platelet/Megakaryocyte |
| 83 | GSTO1 | 1.60E-06 | 7.4378624 | Platelet/Megakaryocyte |
| 84 | MIR4435-2HG | 1.68E-06 | 3.9517767 | Platelet/Megakaryocyte |
| 85 | AMD1 | 2.52E-06 | 2.8864038 | Platelet/Megakaryocyte |
| 86 | PDZK1IP1 | 3.21E-06 | 4.0430827 | Platelet/Megakaryocyte |
| 87 | TRIM58 | 4.22E-06 | 7.1316824 | Platelet/Megakaryocyte |
| 88 | EMC3 | 4.67E-06 | 2.498988 | Platelet/Megakaryocyte |
| 89 | TLK1 | 4.75E-06 | 4.2575064 | Platelet/Megakaryocyte |
| 90 | ACTN1 | 7.33E-06 | 3.5667973 | Platelet/Megakaryocyte |
| 91 | SMIM5 | 8.26E-06 | 5.7943015 | Platelet/Megakaryocyte |
| 92 | VDAC3 | 8.99E-06 | 2.5530715 | Platelet/Megakaryocyte |
| 93 | HIST1H2BK | 9.65E-06 | 2.8332837 | Platelet/Megakaryocyte |
| 94 | SH3BGRL3 | 9.93E-06 | 13.891773 | Platelet/Megakaryocyte |
| 95 | MFSD1 | 9.94E-06 | 2.5948434 | Platelet/Megakaryocyte |
| 96 | ENKUR | 1.01E-05 | 4.7441735 | Platelet/Megakaryocyte |
| 97 | RABGAP1L | 1.07E-05 | 3.3273966 | Platelet/Megakaryocyte |
| 98 | VCL | 1.11E-05 | 3.705714 | Platelet/Megakaryocyte |
| 99 | LYPLAL1 | 1.16E-05 | 2.6511025 | Platelet/Megakaryocyte |
| 0 | KRT19 | 1.03E-26 | 330.65448 | Proliferating Basal |
| 1 | TACSTD2 | 1.04E-26 | 65.25126 | Proliferating Basal |
| 2 | KRT15 | 1.04E-26 | 134.75554 | Proliferating Basal |
| 3 | PERP | 2.05E-26 | 51.253902 | Proliferating Basal |
| 4 | PTTG1 | 2.31E-26 | 19.88485 | Proliferating Basal |
| 5 | HMGB2 | 2.31E-26 | 36.30166 | Proliferating Basal |
| 6 | STMN1 | 2.31E-26 | 44.159557 | Proliferating Basal |
| 7 | HMGB1 | 3.23E-26 | 69.94465 | Proliferating Basal |
| 8 | FXYD3 | 6.74E-26 | 56.161488 | Proliferating Basal |
| 9 | S100A2 | 6.74E-26 | 101.29838 | Proliferating Basal |
| 10 | CKS1B | 7.13E-26 | 16.16849 | Proliferating Basal |
| 11 | TUBB | 9.19E-26 | 53.94266 | Proliferating Basal |
| 12 | RPLP0 | 1.65E-25 | 150.42578 | Proliferating Basal |
| 13 | IMPDH2 | 1.65E-25 | 15.570613 | Proliferating Basal |
| 14 | PLP2 | 1.81E-25 | 24.056969 | Proliferating Basal |
| 15 | PTMA | 2.22E-25 | 244.96228 | Proliferating Basal |
| 16 | H2AFZ | 2.25E-25 | 66.41789 | Proliferating Basal |
| 17 | HMGA1 | 2.25E-25 | 21.89178 | Proliferating Basal |
| 18 | RPS5 | 2.25E-25 | 156.21758 | Proliferating Basal |
| 19 | EEF1G | 2.66E-25 | 141.82863 | Proliferating Basal |
| 20 | KRT8 | 2.66E-25 | 78.936035 | Proliferating Basal |
| 21 | NGFRAP1 | 4.14E-25 | 16.738485 | Proliferating Basal |
| 22 | LRRC75A-AS1 | 5.04E-25 | 86.245 | Proliferating Basal |
| 24 | GSTP1 | 6.63E-25 | 296.37033 | Proliferating Basal |
| 23 | KRT17 | 6.63E-25 | 70.3829 | Proliferating Basal |
| 25 | HSPB1 | 7.26E-25 | 71.20648 | Proliferating Basal |
| 26 | HINT1 | 9.84E-25 | 55.534374 | Proliferating Basal |

| | | | | |
|---|---|---|---|---|
| 27 | HNRNPA1 | 1.19E-24 | 61.126904 | Proliferating Basal |
| 28 | CLDN4 | 1.19E-24 | 17.739641 | Proliferating Basal |
| 29 | HMGN2 | 1.37E-24 | 41.63491 | Proliferating Basal |
| 30 | SLC25A5 | 1.85E-24 | 48.90487 | Proliferating Basal |
| 31 | RPS2 | 1.95E-24 | 399.27643 | Proliferating Basal |
| 36 | KRT18 | 2.01E-24 | 56.078434 | Proliferating Basal |
| 34 | KRT5 | 2.01E-24 | 59.88299 | Proliferating Basal |
| 35 | RPS4X | 2.01E-24 | 318.01794 | Proliferating Basal |
| 32 | NUCKS1 | 2.01E-24 | 24.67309 | Proliferating Basal |
| 33 | RPL18 | 2.01E-24 | 145.30234 | Proliferating Basal |
| 37 | S100A14 | 2.41E-24 | 23.843365 | Proliferating Basal |
| 38 | GAPDH | 2.48E-24 | 191.76953 | Proliferating Basal |
| 39 | SLC25A6 | 2.86E-24 | 59.34939 | Proliferating Basal |
| 40 | RPSA | 2.87E-24 | 136.43289 | Proliferating Basal |
| 41 | MRPL51 | 3.14E-24 | 14.064724 | Proliferating Basal |
| 42 | SDC1 | 3.14E-24 | 11.124692 | Proliferating Basal |
| 43 | MIR205HG | 3.52E-24 | 46.476536 | Proliferating Basal |
| 45 | RPL10A | 3.73E-24 | 203.2048 | Proliferating Basal |
| 44 | RPL7 | 3.73E-24 | 221.35043 | Proliferating Basal |
| 46 | RANBP1 | 3.73E-24 | 19.184776 | Proliferating Basal |
| 47 | RPS3 | 4.11E-24 | 258.6173 | Proliferating Basal |
| 48 | SPINT2 | 6.05E-24 | 26.225782 | Proliferating Basal |
| 49 | NPM1 | 6.30E-24 | 62.271683 | Proliferating Basal |
| 50 | RAN | 8.70E-24 | 24.317646 | Proliferating Basal |
| 51 | RPL13A | 1.02E-23 | 344.95593 | Proliferating Basal |
| 52 | GNB2L1 | 1.03E-23 | 128.96953 | Proliferating Basal |
| 53 | ANP32B | 1.13E-23 | 16.145441 | Proliferating Basal |
| 54 | H2AFV | 1.13E-23 | 16.612616 | Proliferating Basal |
| 55 | AHCY | 1.39E-23 | 9.812408 | Proliferating Basal |
| 56 | GAS5 | 1.78E-23 | 42.01655 | Proliferating Basal |
| 57 | RPLP1 | 1.83E-23 | 554.7289 | Proliferating Basal |
| 58 | MZT2B | 1.92E-23 | 24.915178 | Proliferating Basal |
| 59 | CYC1 | 2.01E-23 | 16.429964 | Proliferating Basal |
| 60 | RPL4 | 2.01E-23 | 85.17019 | Proliferating Basal |
| 61 | GMNN | 2.02E-23 | 8.721064 | Proliferating Basal |
| 62 | SFN | 2.14E-23 | 19.435648 | Proliferating Basal |
| 63 | MGST1 | 2.20E-23 | 34.765625 | Proliferating Basal |
| 64 | HMGN3 | 2.23E-23 | 28.15359 | Proliferating Basal |
| 65 | C6orf48 | 2.25E-23 | 13.883796 | Proliferating Basal |
| 66 | KRTCAP3 | 2.25E-23 | 9.105744 | Proliferating Basal |
| 67 | UBE2C | 2.25E-23 | 20.081608 | Proliferating Basal |
| 68 | MDK | 2.25E-23 | 28.387253 | Proliferating Basal |
| 69 | RPL3 | 2.25E-23 | 277.46173 | Proliferating Basal |
| 70 | RPS19 | 2.51E-23 | 212.53276 | Proliferating Basal |
| 71 | RPS18 | 2.64E-23 | 402.26178 | Proliferating Basal |
| 72 | KIAA0101 | 3.11E-23 | 17.8865 | Proliferating Basal |
| 73 | PRDX2 | 3.35E-23 | 24.433025 | Proliferating Basal |
| 74 | S100A16 | 3.54E-23 | 18.092848 | Proliferating Basal |
| 75 | RPS3A | 4.45E-23 | 196.40962 | Proliferating Basal |
| 76 | EEF2 | 4.45E-23 | 48.265892 | Proliferating Basal |
| 77 | EEF1D | 4.53E-23 | 49.025555 | Proliferating Basal |
| 78 | MIF | 4.53E-23 | 44.83172 | Proliferating Basal |
| 79 | PKM | 4.64E-23 | 34.260975 | Proliferating Basal |
| 80 | ATP5B | 4.65E-23 | 34.00741 | Proliferating Basal |
| 81 | KLF5 | 4.66E-23 | 17.255453 | Proliferating Basal |
| 82 | TECR | 4.99E-23 | 16.306057 | Proliferating Basal |
| 83 | RPL29 | 4.99E-23 | 155.05605 | Proliferating Basal |
| 84 | ARPC1A | 5.05E-23 | 8.987088 | Proliferating Basal |
| 85 | FBL | 5.52E-23 | 11.819516 | Proliferating Basal |
| 86 | AQP3 | 6.57E-23 | 81.82496 | Proliferating Basal |
| 87 | RPL8 | 6.91E-23 | 215.31827 | Proliferating Basal |
| 88 | RPS16 | 8.93E-23 | 133.85216 | Proliferating Basal |
| 89 | ATXN10 | 8.97E-23 | 6.29534 | Proliferating Basal |
| 90 | RPS9 | 8.97E-23 | 184.96707 | Proliferating Basal |

| | | | | |
|---|---|---|---|---|
| 91 | RPL15 | 9.24E-23 | 221.22453 | Proliferating Basal |
| 92 | MTCH1 | 9.41E-23 | 9.258109 | Proliferating Basal |
| 93 | RPS7 | 1.06E-22 | 142.4356 | Proliferating Basal |
| 94 | HNRNPA2B1 | 1.09E-22 | 31.379416 | Proliferating Basal |
| 95 | SIVA1 | 1.38E-22 | 16.371899 | Proliferating Basal |
| 96 | PBK | 1.49E-22 | 10.989664 | Proliferating Basal |
| 97 | TPI1 | 1.54E-22 | 36.39064 | Proliferating Basal |
| 98 | GGH | 1.62E-22 | 9.422597 | Proliferating Basal |
| 99 | BTF3 | 1.84E-22 | 76.10535 | Proliferating Basal |
| 0 | STMN1 | 4.18E-140 | 77.1562 | Proliferating Macrophage |
| 1 | H2AFZ | 5.07E-137 | 130.91528 | Proliferating Macrophage |
| 2 | TUBB | 5.07E-137 | 87.55958 | Proliferating Macrophage |
| 3 | HMGN2 | 7.83E-136 | 74.09394 | Proliferating Macrophage |
| 4 | TUBA1B | 4.92E-134 | 223.73065 | Proliferating Macrophage |
| 5 | HMGB1 | 4.92E-134 | 70.05371 | Proliferating Macrophage |
| 6 | CKS1B | 2.79E-131 | 16.454605 | Proliferating Macrophage |
| 7 | CENPW | 1.85E-130 | 11.859951 | Proliferating Macrophage |
| 8 | HMGB2 | 1.82E-128 | 28.90239 | Proliferating Macrophage |
| 9 | DEK | 1.89E-126 | 18.53873 | Proliferating Macrophage |
| 10 | ANP32B | 7.23E-126 | 17.976072 | Proliferating Macrophage |
| 11 | H2AFV | 4.32E-125 | 18.205187 | Proliferating Macrophage |
| 12 | TK1 | 1.19E-123 | 19.912584 | Proliferating Macrophage |
| 13 | KIAA0101 | 3.08E-122 | 22.511864 | Proliferating Macrophage |
| 14 | RHEB | 7.92E-122 | 18.3675 | Proliferating Macrophage |
| 15 | RAN | 1.22E-119 | 24.424856 | Proliferating Macrophage |
| 16 | NUCKS1 | 4.52E-119 | 14.106658 | Proliferating Macrophage |
| 17 | ARL6IP1 | 3.12E-117 | 33.259354 | Proliferating Macrophage |
| 18 | SMC2 | 5.63E-115 | 8.312777 | Proliferating Macrophage |
| 19 | PTMA | 6.84E-115 | 86.86716 | Proliferating Macrophage |
| 20 | LSM4 | 2.13E-113 | 12.628017 | Proliferating Macrophage |
| 21 | HMGN1 | 4.28E-113 | 13.619325 | Proliferating Macrophage |
| 22 | AP2S1 | 6.57E-113 | 20.954142 | Proliferating Macrophage |
| 23 | GGH | 1.15E-112 | 12.096968 | Proliferating Macrophage |
| 24 | MZT2B | 2.32E-112 | 13.458903 | Proliferating Macrophage |
| 25 | VPS29 | 5.01E-112 | 15.985681 | Proliferating Macrophage |
| 26 | DTYMK | 5.65E-112 | 9.842801 | Proliferating Macrophage |
| 27 | PPIA | 1.88E-111 | 48.375053 | Proliferating Macrophage |
| 28 | NUSAP1 | 3.33E-111 | 15.843446 | Proliferating Macrophage |
| 29 | FAM111A | 4.04E-111 | 8.338366 | Proliferating Macrophage |
| 30 | MARCO | 9.16E-111 | 86.84648 | Proliferating Macrophage |
| 31 | ANAPC11 | 9.32E-111 | 10.541234 | Proliferating Macrophage |
| 32 | TMEM106C | 1.56E-110 | 11.242904 | Proliferating Macrophage |
| 33 | NENF | 5.44E-110 | 10.752351 | Proliferating Macrophage |
| 34 | MS4A4A | 1.79E-109 | 20.871408 | Proliferating Macrophage |
| 35 | RPA3 | 3.90E-109 | 9.426484 | Proliferating Macrophage |
| 36 | GAPDH | 1.47E-108 | 71.08811 | Proliferating Macrophage |
| 37 | TYMS | 1.55E-108 | 19.723703 | Proliferating Macrophage |
| 38 | GRN | 1.78E-108 | 113.312775 | Proliferating Macrophage |
| 39 | TUBA1C | 3.25E-108 | 16.510057 | Proliferating Macrophage |
| 40 | VIM | 3.52E-108 | 207.60614 | Proliferating Macrophage |
| 41 | DUT | 4.98E-108 | 17.715818 | Proliferating Macrophage |
| 42 | HN1 | 6.91E-108 | 18.826212 | Proliferating Macrophage |
| 43 | LSM3 | 8.30E-108 | 9.400742 | Proliferating Macrophage |
| 44 | YBX1 | 2.54E-107 | 62.42077 | Proliferating Macrophage |
| 45 | YWHAH | 2.89E-107 | 18.778263 | Proliferating Macrophage |
| 46 | TMEM160 | 7.81E-107 | 7.978177 | Proliferating Macrophage |
| 47 | MNDA | 1.14E-106 | 17.72707 | Proliferating Macrophage |
| 48 | HEXB | 1.38E-106 | 17.586975 | Proliferating Macrophage |
| 49 | TUBB6 | 2.15E-106 | 11.676661 | Proliferating Macrophage |
| 50 | SUMO2 | 3.43E-106 | 21.97408 | Proliferating Macrophage |
| 51 | TUBB4B | 3.43E-106 | 23.074741 | Proliferating Macrophage |
| 52 | C1QB | 6.00E-106 | 261.42532 | Proliferating Macrophage |
| 53 | TAGLN2 | 1.15E-105 | 55.43873 | Proliferating Macrophage |
| 54 | CKS2 | 1.65E-105 | 16.01655 | Proliferating Macrophage |

| | | | | |
|---|---|---|---|---|
| 55 | PLP2 | 2.77E-105 | 13.1216345 | Proliferating Macrophage |
| 56 | MAD2L1 | 3.38E-105 | 9.714752 | Proliferating Macrophage |
| 57 | SMC4 | 3.99E-105 | 9.356377 | Proliferating Macrophage |
| 58 | UBB | 7.61E-104 | 91.17841 | Proliferating Macrophage |
| 59 | RPS27L | 8.64E-104 | 20.835854 | Proliferating Macrophage |
| 60 | BLOC1S1 | 1.06E-103 | 13.640627 | Proliferating Macrophage |
| 61 | VSIG4 | 1.49E-103 | 37.615204 | Proliferating Macrophage |
| 62 | TKT | 1.86E-103 | 18.460194 | Proliferating Macrophage |
| 63 | LYZ | 1.87E-103 | 247.03297 | Proliferating Macrophage |
| 64 | RPS20 | 2.28E-103 | 68.37216 | Proliferating Macrophage |
| 65 | PSAP | 2.45E-103 | 72.658875 | Proliferating Macrophage |
| 66 | CALM2 | 2.60E-103 | 57.370026 | Proliferating Macrophage |
| 67 | RHOA | 7.59E-103 | 25.921743 | Proliferating Macrophage |
| 68 | CALM3 | 1.20E-102 | 10.772569 | Proliferating Macrophage |
| 69 | CDK1 | 1.46E-102 | 18.232265 | Proliferating Macrophage |
| 70 | FN1 | 1.74E-102 | 41.096752 | Proliferating Macrophage |
| 71 | NAP1L1 | 2.00E-102 | 11.603448 | Proliferating Macrophage |
| 72 | LSM5 | 2.41E-102 | 6.8872967 | Proliferating Macrophage |
| 73 | GSTO1 | 2.68E-102 | 35.042984 | Proliferating Macrophage |
| 74 | SNRPD1 | 3.76E-102 | 7.8042197 | Proliferating Macrophage |
| 75 | MRPL51 | 4.24E-102 | 8.794792 | Proliferating Macrophage |
| 76 | COX8A | 7.84E-102 | 17.43977 | Proliferating Macrophage |
| 77 | CTSC | 1.35E-101 | 46.658665 | Proliferating Macrophage |
| 78 | C1QC | 1.52E-101 | 105.00673 | Proliferating Macrophage |
| 79 | GNG5 | 1.59E-101 | 15.4335785 | Proliferating Macrophage |
| 80 | GPX4 | 1.64E-101 | 37.408653 | Proliferating Macrophage |
| 81 | ARPC3 | 3.30E-101 | 38.486893 | Proliferating Macrophage |
| 82 | HNRNPA2B1 | 3.30E-101 | 19.925058 | Proliferating Macrophage |
| 83 | PSMA4 | 3.79E-101 | 12.073158 | Proliferating Macrophage |
| 84 | SNRPB | 4.27E-101 | 10.857956 | Proliferating Macrophage |
| 85 | PFN1 | 4.63E-101 | 85.60131 | Proliferating Macrophage |
| 87 | NUDT1 | 4.70E-101 | 6.4491463 | Proliferating Macrophage |
| 86 | MS4A6A | 4.70E-101 | 14.277918 | Proliferating Macrophage |
| 88 | MDH1 | 7.18E-101 | 13.983376 | Proliferating Macrophage |
| 89 | MS4A7 | 7.94E-101 | 23.750622 | Proliferating Macrophage |
| 90 | FABP4 | 9.79E-101 | 185.38132 | Proliferating Macrophage |
| 91 | CSTA | 1.19E-100 | 20.269861 | Proliferating Macrophage |
| 92 | CRIP1 | 1.21E-100 | 184.56136 | Proliferating Macrophage |
| 93 | H2AFY | 1.29E-100 | 11.381867 | Proliferating Macrophage |
| 94 | RAC1 | 1.80E-100 | 27.714655 | Proliferating Macrophage |
| 95 | CHCHD2 | 1.87E-100 | 24.984892 | Proliferating Macrophage |
| 96 | UBE2C | 2.32E-100 | 24.645287 | Proliferating Macrophage |
| 97 | DBI | 2.49E-100 | 23.247137 | Proliferating Macrophage |
| 98 | C1QA | 2.88E-100 | 197.89964 | Proliferating Macrophage |
| 99 | CTSB | 2.97E-100 | 30.740114 | Proliferating Macrophage |
| 0 | HMGB2 | 6.93E-63 | 30.568062 | Proliferating NK/T |
| 1 | STMN1 | 1.03E-60 | 27.716675 | Proliferating NK/T |
| 2 | SMC4 | 1.14E-48 | 7.5507245 | Proliferating NK/T |
| 3 | GZMA | 2.02E-48 | 14.594669 | Proliferating NK/T |
| 4 | BIRC5 | 1.34E-46 | 9.906653 | Proliferating NK/T |
| 5 | H2AFV | 3.45E-46 | 7.9631653 | Proliferating NK/T |
| 6 | TUBB | 3.45E-46 | 28.21888 | Proliferating NK/T |
| 7 | CORO1A | 3.45E-46 | 12.022384 | Proliferating NK/T |
| 8 | DEK | 2.84E-45 | 7.289869 | Proliferating NK/T |
| 9 | CKS1B | 2.55E-44 | 6.5812664 | Proliferating NK/T |
| 10 | KIAA0101 | 3.62E-44 | 10.085524 | Proliferating NK/T |
| 11 | HMGB1 | 4.33E-44 | 21.237173 | Proliferating NK/T |
| 12 | MKI67 | 5.58E-44 | 11.716371 | Proliferating NK/T |
| 13 | HMGN2 | 7.19E-44 | 16.696817 | Proliferating NK/T |
| 14 | TYMS | 1.68E-43 | 9.007095 | Proliferating NK/T |
| 15 | PTMA | 1.88E-43 | 51.497578 | Proliferating NK/T |
| 16 | NUSAP1 | 2.58E-43 | 8.234677 | Proliferating NK/T |
| 17 | H2AFZ | 3.93E-43 | 15.882992 | Proliferating NK/T |
| 18 | PTPRCAP | 5.46E-43 | 8.07075 | Proliferating NK/T |

| | | | | |
|---|---|---|---|---|
| 19 | NKG7 | 5.20E-42 | 23.742624 | Proliferating NK/T |
| 20 | CALM3 | 9.85E-38 | 7.273368 | Proliferating NK/T |
| 21 | CXCR4 | 1.08E-37 | 11.785486 | Proliferating NK/T |
| 22 | HIST1H4C | 1.55E-37 | 12.480488 | Proliferating NK/T |
| 23 | RAC2 | 1.78E-36 | 6.823108 | Proliferating NK/T |
| 24 | C12orf75 | 2.13E-36 | 5.426178 | Proliferating NK/T |
| 25 | GZMB | 5.63E-36 | 13.015694 | Proliferating NK/T |
| 26 | TOP2A | 6.34E-36 | 8.2668915 | Proliferating NK/T |
| 27 | UBE2C | 7.99E-36 | 8.350592 | Proliferating NK/T |
| 28 | CENPF | 1.20E-35 | 8.898412 | Proliferating NK/T |
| 29 | CD247 | 1.67E-35 | 5.537334 | Proliferating NK/T |
| 30 | CD7 | 5.75E-34 | 6.8855233 | Proliferating NK/T |
| 31 | CTSW | 9.19E-34 | 7.249365 | Proliferating NK/T |
| 32 | ANP32B | 1.15E-33 | 6.983864 | Proliferating NK/T |
| 33 | DNAJC9 | 2.98E-33 | 5.1572227 | Proliferating NK/T |
| 34 | CST7 | 5.02E-33 | 6.527937 | Proliferating NK/T |
| 35 | CKS2 | 1.20E-32 | 5.8944287 | Proliferating NK/T |
| 36 | ANP32E | 5.14E-32 | 5.302 | Proliferating NK/T |
| 37 | PCNA | 1.03E-31 | 7.2789974 | Proliferating NK/T |
| 38 | CARHSP1 | 1.45E-31 | 3.8784559 | Proliferating NK/T |
| 39 | PRF1 | 2.91E-31 | 7.7512336 | Proliferating NK/T |
| 40 | IFI16 | 7.85E-31 | 4.2372856 | Proliferating NK/T |
| 41 | ZFP36L2 | 1.56E-30 | 11.094642 | Proliferating NK/T |
| 42 | ASPM | 5.36E-30 | 9.488414 | Proliferating NK/T |
| 43 | H2AFX | 8.06E-30 | 5.4977174 | Proliferating NK/T |
| 44 | NUCB2 | 1.19E-29 | 3.678448 | Proliferating NK/T |
| 45 | IDH2 | 1.88E-29 | 5.0994935 | Proliferating NK/T |
| 46 | PTTG1 | 2.94E-29 | 6.749871 | Proliferating NK/T |
| 47 | TUBA1B | 3.16E-29 | 26.90844 | Proliferating NK/T |
| 48 | CCL5 | 5.24E-29 | 12.881602 | Proliferating NK/T |
| 49 | SMC2 | 6.72E-29 | 4.5564866 | Proliferating NK/T |
| 50 | HMGN1 | 1.28E-28 | 5.857363 | Proliferating NK/T |
| 51 | MZT2B | 1.82E-28 | 5.548415 | Proliferating NK/T |
| 52 | USP1 | 2.04E-28 | 4.133096 | Proliferating NK/T |
| 53 | MCM7 | 3.10E-28 | 5.388861 | Proliferating NK/T |
| 54 | NUCKS1 | 3.10E-28 | 5.0203624 | Proliferating NK/T |
| 55 | ZWINT | 3.80E-27 | 5.7858796 | Proliferating NK/T |
| 56 | TK1 | 3.98E-27 | 6.106813 | Proliferating NK/T |
| 57 | DDX39A | 8.25E-27 | 4.316117 | Proliferating NK/T |
| 58 | CDKN2D | 1.09E-26 | 4.824534 | Proliferating NK/T |
| 59 | TMPO | 1.46E-26 | 4.2931004 | Proliferating NK/T |
| 60 | HNRNPA1 | 2.24E-26 | 12.9318285 | Proliferating NK/T |
| 61 | ASF1B | 3.54E-26 | 7.0741496 | Proliferating NK/T |
| 62 | GNLY | 4.24E-26 | 25.345327 | Proliferating NK/T |
| 63 | IFITM1 | 5.45E-26 | 9.773754 | Proliferating NK/T |
| 64 | RALY | 5.80E-26 | 3.3714857 | Proliferating NK/T |
| 65 | CDKN3 | 6.98E-26 | 6.6139174 | Proliferating NK/T |
| 66 | C9orf142 | 7.41E-26 | 3.6359293 | Proliferating NK/T |
| 67 | CENPM | 8.38E-26 | 5.1352797 | Proliferating NK/T |
| 68 | SYNE2 | 8.38E-26 | 3.662916 | Proliferating NK/T |
| 69 | CDK1 | 8.81E-26 | 5.3209896 | Proliferating NK/T |
| 70 | KLRB1 | 2.10E-25 | 5.498745 | Proliferating NK/T |
| 71 | GZMH | 2.37E-25 | 5.7739873 | Proliferating NK/T |
| 72 | DUT | 3.68E-25 | 6.042372 | Proliferating NK/T |
| 73 | IL2RG | 4.15E-25 | 3.9177086 | Proliferating NK/T |
| 74 | AURKB | 4.52E-25 | 7.931398 | Proliferating NK/T |
| 75 | BUB3 | 4.54E-25 | 3.6526632 | Proliferating NK/T |
| 76 | DDIT4 | 5.40E-25 | 5.616445 | Proliferating NK/T |
| 77 | RRM2 | 6.19E-25 | 7.6487436 | Proliferating NK/T |
| 78 | UBE2T | 6.55E-25 | 5.7426476 | Proliferating NK/T |
| 79 | TMEM106C | 1.91E-24 | 4.512339 | Proliferating NK/T |
| 80 | TPX2 | 3.96E-24 | 7.3618593 | Proliferating NK/T |
| 81 | GNG2 | 4.16E-24 | 4.6003146 | Proliferating NK/T |
| 82 | ARHGDIB | 9.04E-24 | 10.258334 | Proliferating NK/T |

| | | | | |
|---|---|---|---|---|
| 83 | MAD2L1 | 1.11E-23 | 5.9134364 | Proliferating NK/T |
| 84 | CDCA5 | 1.45E-23 | 7.7528973 | Proliferating NK/T |
| 85 | EZH2 | 2.56E-23 | 5.4940524 | Proliferating NK/T |
| 86 | CDT1 | 3.24E-23 | 7.0460277 | Proliferating NK/T |
| 87 | BIN2 | 3.26E-23 | 3.6947715 | Proliferating NK/T |
| 88 | KIF22 | 6.03E-23 | 4.058691 | Proliferating NK/T |
| 89 | SNRPB | 7.18E-23 | 4.696506 | Proliferating NK/T |
| 90 | MAD2L2 | 7.57E-23 | 3.4490082 | Proliferating NK/T |
| 91 | GAPDH | 1.26E-22 | 29.852297 | Proliferating NK/T |
| 92 | FGFBP2 | 1.98E-22 | 5.707076 | Proliferating NK/T |
| 93 | RPSA | 2.34E-22 | 15.681626 | Proliferating NK/T |
| 94 | ITGB2 | 2.40E-22 | 4.1148 | Proliferating NK/T |
| 95 | KLRD1 | 6.14E-22 | 5.586366 | Proliferating NK/T |
| 96 | PFN1 | 9.34E-22 | 26.08092 | Proliferating NK/T |
| 97 | SNHG6 | 1.07E-21 | 4.3300695 | Proliferating NK/T |
| 98 | SPC25 | 2.47E-21 | 6.970335 | Proliferating NK/T |
| 99 | GTSE1 | 2.59E-21 | 7.2160425 | Proliferating NK/T |
| 0 | MIR205HG | 9.46E-99 | 36.818836 | Proximal Basal |
| 1 | KRT19 | 7.49E-98 | 216.04959 | Proximal Basal |
| 2 | PERP | 3.71E-96 | 36.89718 | Proximal Basal |
| 3 | TACSTD2 | 4.62E-96 | 48.993176 | Proximal Basal |
| 4 | PRSS23 | 1.53E-95 | 29.855583 | Proximal Basal |
| 5 | S100A2 | 1.53E-95 | 43.28301 | Proximal Basal |
| 6 | AQP3 | 1.27E-93 | 60.26099 | Proximal Basal |
| 7 | KRT15 | 7.28E-92 | 82.580956 | Proximal Basal |
| 8 | F3 | 3.10E-91 | 19.845493 | Proximal Basal |
| 9 | FXYD3 | 1.31E-90 | 35.32843 | Proximal Basal |
| 10 | KLF5 | 1.46E-90 | 14.617402 | Proximal Basal |
| 11 | WFDC2 | 2.90E-90 | 63.365036 | Proximal Basal |
| 12 | KRT5 | 1.01E-89 | 28.637413 | Proximal Basal |
| 13 | CHST9 | 2.08E-89 | 11.682169 | Proximal Basal |
| 14 | SERPINF1 | 2.41E-89 | 17.91541 | Proximal Basal |
| 15 | RPS4X | 2.89E-89 | 204.16109 | Proximal Basal |
| 16 | AQP5 | 2.96E-89 | 26.889057 | Proximal Basal |
| 17 | RPL10A | 5.86E-89 | 139.96706 | Proximal Basal |
| 18 | GAS5 | 2.80E-88 | 33.37274 | Proximal Basal |
| 19 | AGR2 | 4.11E-88 | 35.60752 | Proximal Basal |
| 20 | RPLP0 | 1.27E-87 | 90.72042 | Proximal Basal |
| 21 | GSTP1 | 6.73E-87 | 125.03533 | Proximal Basal |
| 22 | KRT17 | 1.03E-86 | 47.710743 | Proximal Basal |
| 23 | RPL3 | 1.10E-86 | 190.89609 | Proximal Basal |
| 24 | CYP4B1 | 2.22E-86 | 14.149611 | Proximal Basal |
| 25 | LRRC75A-AS1 | 2.98E-86 | 53.30879 | Proximal Basal |
| 26 | KLK11 | 3.69E-86 | 13.794685 | Proximal Basal |
| 27 | KRT8 | 4.50E-86 | 32.207306 | Proximal Basal |
| 28 | EEF1G | 8.35E-86 | 86.885506 | Proximal Basal |
| 29 | RPLP1 | 7.38E-85 | 347.13885 | Proximal Basal |
| 30 | HMGB3 | 1.33E-84 | 11.317076 | Proximal Basal |
| 31 | RPS6 | 1.55E-84 | 170.79382 | Proximal Basal |
| 32 | RPS3 | 2.61E-84 | 147.06616 | Proximal Basal |
| 33 | RPS18 | 2.74E-84 | 248.70006 | Proximal Basal |
| 34 | RPSA | 4.96E-84 | 70.95492 | Proximal Basal |
| 35 | EEF2 | 6.79E-84 | 37.871967 | Proximal Basal |
| 36 | KRT18 | 6.91E-84 | 29.429054 | Proximal Basal |
| 37 | IGFBP3 | 6.94E-84 | 13.477646 | Proximal Basal |
| 38 | GNB2L1 | 7.78E-84 | 91.18498 | Proximal Basal |
| 39 | RPL18 | 8.44E-84 | 91.43879 | Proximal Basal |
| 40 | C19orf33 | 9.76E-84 | 15.652592 | Proximal Basal |
| 41 | RPL13A | 2.09E-83 | 226.70236 | Proximal Basal |
| 42 | SLPI | 2.09E-83 | 181.89226 | Proximal Basal |
| 43 | RPL7 | 2.71E-83 | 140.27448 | Proximal Basal |
| 44 | RPS2 | 3.07E-83 | 216.14772 | Proximal Basal |
| 45 | ELF3 | 4.68E-83 | 16.117414 | Proximal Basal |
| 46 | SPINT2 | 1.03E-82 | 16.998789 | Proximal Basal |

| | | | | |
|---|---|---|---|---|
| 47 | HS3ST1 | 1.42E-82 | 10.138452 | Proximal Basal |
| 48 | RPL13 | 1.70E-82 | 213.99316 | Proximal Basal |
| 49 | RPL7A | 2.11E-82 | 95.8568 | Proximal Basal |
| 50 | EPHX1 | 2.44E-82 | 13.766846 | Proximal Basal |
| 51 | CXCL17 | 2.99E-82 | 27.346233 | Proximal Basal |
| 52 | RPS7 | 3.57E-82 | 84.645424 | Proximal Basal |
| 53 | RPL8 | 4.65E-82 | 132.33292 | Proximal Basal |
| 54 | FMO2 | 8.50E-82 | 11.128609 | Proximal Basal |
| 55 | SERPINB3 | 3.84E-81 | 51.538975 | Proximal Basal |
| 56 | RPS5 | 3.91E-81 | 84.687515 | Proximal Basal |
| 57 | ADH1C | 6.05E-81 | 13.212114 | Proximal Basal |
| 58 | RPL15 | 6.64E-81 | 138.71808 | Proximal Basal |
| 59 | RPL5 | 7.75E-81 | 68.226776 | Proximal Basal |
| 60 | MDK | 9.15E-81 | 14.021516 | Proximal Basal |
| 61 | HMGN3 | 1.24E-80 | 18.228075 | Proximal Basal |
| 62 | RPL29 | 1.52E-80 | 97.01563 | Proximal Basal |
| 63 | PPAP2C | 5.97E-80 | 9.921581 | Proximal Basal |
| 64 | BTF3 | 1.35E-79 | 51.773438 | Proximal Basal |
| 65 | RPL12 | 2.07E-79 | 125.27509 | Proximal Basal |
| 66 | CLDN4 | 2.52E-79 | 11.363771 | Proximal Basal |
| 67 | SDC1 | 4.01E-79 | 8.278313 | Proximal Basal |
| 68 | RPS8 | 1.46E-78 | 104.05977 | Proximal Basal |
| 69 | LMO4 | 2.16E-78 | 14.057745 | Proximal Basal |
| 70 | RPL4 | 3.13E-78 | 55.52763 | Proximal Basal |
| 71 | EHF | 6.75E-78 | 8.745751 | Proximal Basal |
| 72 | NPM1 | 6.82E-78 | 32.546097 | Proximal Basal |
| 73 | RPL18A | 9.19E-78 | 125.53232 | Proximal Basal |
| 74 | CCND1 | 1.13E-77 | 12.105117 | Proximal Basal |
| 75 | RPL10 | 1.18E-77 | 248.2741 | Proximal Basal |
| 76 | ZFP36L1 | 1.33E-77 | 18.588457 | Proximal Basal |
| 77 | OAT | 1.45E-77 | 9.048789 | Proximal Basal |
| 78 | TSPAN1 | 2.32E-77 | 12.440402 | Proximal Basal |
| 79 | RPL6 | 2.94E-77 | 79.84043 | Proximal Basal |
| 80 | RPS9 | 3.61E-77 | 114.847946 | Proximal Basal |
| 81 | RPS14 | 1.15E-76 | 150.61333 | Proximal Basal |
| 82 | MGST1 | 1.35E-76 | 22.918097 | Proximal Basal |
| 83 | TSC22D1 | 1.41E-76 | 20.633629 | Proximal Basal |
| 84 | EEF1D | 1.75E-76 | 30.052832 | Proximal Basal |
| 85 | RPS3A | 2.75E-76 | 122.99467 | Proximal Basal |
| 86 | DDR1 | 4.07E-76 | 7.323189 | Proximal Basal |
| 87 | BAG1 | 9.89E-76 | 11.075407 | Proximal Basal |
| 88 | IGFBP2 | 2.35E-75 | 12.765765 | Proximal Basal |
| 89 | BCAM | 9.28E-75 | 17.69379 | Proximal Basal |
| 90 | SNHG5 | 1.12E-74 | 18.343037 | Proximal Basal |
| 91 | SYTL1 | 2.08E-74 | 8.451789 | Proximal Basal |
| 92 | IMPDH2 | 3.65E-74 | 9.38275 | Proximal Basal |
| 93 | RPS12 | 3.70E-74 | 142.15117 | Proximal Basal |
| 94 | SLC25A6 | 5.57E-74 | 36.585472 | Proximal Basal |
| 95 | SOX2 | 1.04E-73 | 8.902432 | Proximal Basal |
| 96 | EPCAM | 1.41E-73 | 8.663956 | Proximal Basal |
| 97 | HNRNPA1 | 2.17E-73 | 31.960785 | Proximal Basal |
| 98 | CLU | 2.39E-73 | 17.174454 | Proximal Basal |
| 99 | ALDH1A1 | 3.42E-73 | 13.457093 | Proximal Basal |
| 0 | MORN2 | 3.10E-54 | 38.46334 | Proximal Ciliated |
| 1 | SNTN | 3.10E-54 | 32.1922 | Proximal Ciliated |
| 2 | ROPN1L | 3.10E-54 | 20.064184 | Proximal Ciliated |
| 3 | C20orf85 | 3.10E-54 | 62.345394 | Proximal Ciliated |
| 4 | C9orf24 | 3.10E-54 | 57.78587 | Proximal Ciliated |
| 5 | PIFO | 3.10E-54 | 30.210974 | Proximal Ciliated |
| 6 | LRRC23 | 3.10E-54 | 16.175127 | Proximal Ciliated |
| 7 | CCDC17 | 3.10E-54 | 11.834471 | Proximal Ciliated |
| 8 | CD24 | 3.10E-54 | 33.25757 | Proximal Ciliated |
| 9 | SMIM22 | 3.10E-54 | 23.444654 | Proximal Ciliated |
| 11 | C1orf194 | 3.34E-54 | 27.481892 | Proximal Ciliated |

| | | | | |
|---|---|---|---|---|
| 10 | CFAP126 | 3.34E-54 | 18.140005 | Proximal Ciliated |
| 12 | FAM183A | 3.55E-54 | 29.655071 | Proximal Ciliated |
| 13 | CETN2 | 4.45E-54 | 29.314299 | Proximal Ciliated |
| 14 | DYNLT1 | 5.43E-54 | 38.96383 | Proximal Ciliated |
| 15 | EFHC1 | 5.49E-54 | 14.819499 | Proximal Ciliated |
| 16 | TSPAN1 | 6.20E-54 | 48.307087 | Proximal Ciliated |
| 17 | CAPS | 7.01E-54 | 72.026985 | Proximal Ciliated |
| 18 | CAPSL | 7.01E-54 | 33.745846 | Proximal Ciliated |
| 19 | DPCD | 7.01E-54 | 14.233261 | Proximal Ciliated |
| 20 | C11orf88 | 7.02E-54 | 17.966166 | Proximal Ciliated |
| 21 | ODF3B | 8.74E-54 | 24.561424 | Proximal Ciliated |
| 22 | MRPS31 | 9.43E-54 | 23.591337 | Proximal Ciliated |
| 23 | PRDX5 | 9.43E-54 | 80.01719 | Proximal Ciliated |
| 24 | TUBB4B | 9.99E-54 | 92.84289 | Proximal Ciliated |
| 25 | RSPH1 | 1.19E-53 | 40.62917 | Proximal Ciliated |
| 26 | LDLRAD1 | 1.21E-53 | 12.980035 | Proximal Ciliated |
| 27 | C9orf116 | 1.34E-53 | 27.63764 | Proximal Ciliated |
| 28 | FAM229B | 1.36E-53 | 20.797802 | Proximal Ciliated |
| 29 | ZMYND10 | 1.36E-53 | 22.165085 | Proximal Ciliated |
| 30 | FAM216B | 1.36E-53 | 13.646084 | Proximal Ciliated |
| 31 | SPA17 | 1.47E-53 | 15.528478 | Proximal Ciliated |
| 32 | FAM92B | 1.63E-53 | 15.914383 | Proximal Ciliated |
| 33 | DNALI1 | 1.63E-53 | 15.535385 | Proximal Ciliated |
| 34 | DYNLRB2 | 2.03E-53 | 18.913208 | Proximal Ciliated |
| 35 | LRRC46 | 2.12E-53 | 12.723304 | Proximal Ciliated |
| 38 | FXYD3 | 2.16E-53 | 39.487804 | Proximal Ciliated |
| 36 | WDR54 | 2.16E-53 | 18.84079 | Proximal Ciliated |
| 37 | CCDC170 | 2.16E-53 | 15.161766 | Proximal Ciliated |
| 39 | TPPP3 | 2.47E-53 | 61.818253 | Proximal Ciliated |
| 40 | POLR2I | 2.48E-53 | 16.520859 | Proximal Ciliated |
| 41 | C5orf49 | 3.74E-53 | 18.111097 | Proximal Ciliated |
| 42 | HSPB11 | 4.22E-53 | 15.735289 | Proximal Ciliated |
| 43 | ATPIF1 | 5.91E-53 | 36.1084 | Proximal Ciliated |
| 44 | TCTEX1D4 | 6.21E-53 | 15.192102 | Proximal Ciliated |
| 45 | CDS1 | 6.55E-53 | 9.161567 | Proximal Ciliated |
| 46 | DYNLL1 | 7.18E-53 | 80.351036 | Proximal Ciliated |
| 47 | TUBA1A | 7.37E-53 | 67.11649 | Proximal Ciliated |
| 48 | C11orf70 | 7.40E-53 | 14.074923 | Proximal Ciliated |
| 49 | WBSCR27 | 8.64E-53 | 12.102943 | Proximal Ciliated |
| 50 | TCTEX1D2 | 9.21E-53 | 21.843334 | Proximal Ciliated |
| 51 | CD164L2 | 1.16E-52 | 12.549773 | Proximal Ciliated |
| 52 | CDHR3 | 1.51E-52 | 15.184527 | Proximal Ciliated |
| 53 | C21orf59 | 1.59E-52 | 12.96765 | Proximal Ciliated |
| 54 | HSPH1 | 1.60E-52 | 12.203309 | Proximal Ciliated |
| 55 | FAM81B | 2.00E-52 | 11.557337 | Proximal Ciliated |
| 56 | CCDC78 | 2.26E-52 | 16.500814 | Proximal Ciliated |
| 57 | TSPAN19 | 2.72E-52 | 12.454651 | Proximal Ciliated |
| 58 | UFC1 | 3.06E-52 | 26.204708 | Proximal Ciliated |
| 59 | CRNDE | 3.12E-52 | 13.243965 | Proximal Ciliated |
| 61 | TMEM190 | 3.20E-52 | 50.65114 | Proximal Ciliated |
| 60 | GSTP1 | 3.20E-52 | 104.86186 | Proximal Ciliated |
| 62 | C1orf189 | 4.38E-52 | 14.249166 | Proximal Ciliated |
| 63 | KIF9 | 6.13E-52 | 13.785777 | Proximal Ciliated |
| 64 | AKAP14 | 6.28E-52 | 13.986428 | Proximal Ciliated |
| 65 | TMC5 | 7.47E-52 | 15.505571 | Proximal Ciliated |
| 66 | LRRIQ1 | 7.75E-52 | 14.490171 | Proximal Ciliated |
| 67 | C22orf15 | 1.02E-51 | 12.059924 | Proximal Ciliated |
| 68 | MORN5 | 1.07E-51 | 19.089418 | Proximal Ciliated |
| 69 | DYDC2 | 1.35E-51 | 11.221239 | Proximal Ciliated |
| 70 | NME5 | 1.46E-51 | 13.953581 | Proximal Ciliated |
| 71 | MS4A8 | 1.51E-51 | 13.789922 | Proximal Ciliated |
| 72 | C9orf135 | 1.76E-51 | 15.861865 | Proximal Ciliated |
| 73 | AGR3 | 2.09E-51 | 35.000507 | Proximal Ciliated |
| 74 | IFT57 | 2.39E-51 | 14.842768 | Proximal Ciliated |

| | | | | |
|---|---|---|---|---|
| 75 | IK | 2.43E-51 | 13.975696 | Proximal Ciliated |
| 76 | RSPH9 | 2.52E-51 | 15.038557 | Proximal Ciliated |
| 77 | HSBP1 | 2.91E-51 | 31.226067 | Proximal Ciliated |
| 78 | CCDC146 | 3.03E-51 | 16.395588 | Proximal Ciliated |
| 79 | CCDC42B | 3.66E-51 | 10.218449 | Proximal Ciliated |
| 80 | ENDOG | 5.02E-51 | 10.639967 | Proximal Ciliated |
| 81 | H2AFJ | 5.55E-51 | 15.451179 | Proximal Ciliated |
| 82 | SPATA18 | 6.15E-51 | 9.156608 | Proximal Ciliated |
| 83 | BASP1 | 6.46E-51 | 12.738556 | Proximal Ciliated |
| 84 | CIB1 | 1.57E-50 | 26.881931 | Proximal Ciliated |
| 85 | IFT22 | 1.67E-50 | 10.628421 | Proximal Ciliated |
| 86 | DNPH1 | 1.89E-50 | 11.514759 | Proximal Ciliated |
| 87 | CFAP53 | 3.00E-50 | 12.890606 | Proximal Ciliated |
| 88 | MNS1 | 3.65E-50 | 10.442508 | Proximal Ciliated |
| 89 | TMEM231 | 3.83E-50 | 9.439984 | Proximal Ciliated |
| 90 | C2orf40 | 4.07E-50 | 17.988848 | Proximal Ciliated |
| 91 | SPAG6 | 4.45E-50 | 11.720504 | Proximal Ciliated |
| 92 | HMGN3 | 4.45E-50 | 23.059772 | Proximal Ciliated |
| 93 | PSENEN | 5.56E-50 | 17.851988 | Proximal Ciliated |
| 94 | IFT27 | 6.93E-50 | 8.147499 | Proximal Ciliated |
| 95 | SLC44A4 | 1.13E-49 | 11.215113 | Proximal Ciliated |
| 96 | WDR34 | 1.93E-49 | 9.777329 | Proximal Ciliated |
| 97 | IFT43 | 2.01E-49 | 9.961898 | Proximal Ciliated |
| 98 | DNAH12 | 2.54E-49 | 12.866642 | Proximal Ciliated |
| 99 | DPY30 | 3.27E-49 | 13.282119 | Proximal Ciliated |
| 0 | PRR4 | 7.78E-14 | 908.466 | Serous |
| 1 | ZG16B | 7.78E-14 | 257.42258 | Serous |
| 2 | LTF | 7.78E-14 | 284.79315 | Serous |
| 3 | SLPI | 7.78E-14 | inf | Serous |
| 4 | PIP | 7.78E-14 | 508.5139 | Serous |
| 5 | SLC12A2 | 7.78E-14 | 15.6923 | Serous |
| 6 | AZGP1 | 7.78E-14 | 79.769684 | Serous |
| 7 | NDRG2 | 7.78E-14 | 27.078255 | Serous |
| 8 | LYZ | 7.78E-14 | inf | Serous |
| 9 | S100A1 | 7.78E-14 | 27.186083 | Serous |
| 12 | BPIFB1 | 8.41E-14 | 902.2882 | Serous |
| 11 | PIGR | 8.41E-14 | 97.91012 | Serous |
| 10 | DMBT1 | 8.41E-14 | 52.960682 | Serous |
| 13 | WFDC2 | 1.23E-13 | 250.66074 | Serous |
| 14 | CXCL17 | 4.53E-13 | 94.711945 | Serous |
| 15 | MARCKSL1 | 5.97E-13 | 9.798609 | Serous |
| 16 | TCN1 | 7.09E-13 | 34.639717 | Serous |
| 17 | LRRC26 | 7.52E-13 | 20.123638 | Serous |
| 18 | PART1 | 7.52E-13 | 12.478888 | Serous |
| 19 | GNAS | 8.29E-13 | 24.555946 | Serous |
| 20 | PPP1R1B | 8.94E-13 | 11.896581 | Serous |
| 21 | AQP5 | 1.38E-12 | 22.171389 | Serous |
| 22 | FXYD3 | 3.91E-12 | 20.02646 | Serous |
| 23 | EHF | 4.30E-12 | 9.059412 | Serous |
| 24 | CD24 | 7.05E-12 | 12.077041 | Serous |
| 25 | RNASE1 | 1.05E-11 | 44.25543 | Serous |
| 26 | PHLDA1 | 1.16E-11 | 9.434013 | Serous |
| 27 | SELM | 1.16E-11 | 14.015405 | Serous |
| 28 | SSR4 | 1.36E-11 | 42.908863 | Serous |
| 29 | SERPINA3 | 2.68E-11 | 20.32317 | Serous |
| 30 | NUCB2 | 7.92E-11 | 13.565014 | Serous |
| 31 | C6orf58 | 9.81E-11 | 513.8133 | Serous |
| 32 | SCGB3A1 | 1.18E-10 | inf | Serous |
| 33 | PRSS8 | 1.39E-10 | 7.4060645 | Serous |
| 34 | FAM3D | 1.55E-10 | 16.928005 | Serous |
| 35 | S100B | 1.65E-10 | 11.531933 | Serous |
| 37 | KIAA1324 | 3.29E-10 | 7.8124146 | Serous |
| 36 | CYP4X1 | 3.29E-10 | 8.139403 | Serous |
| 38 | XBP1 | 5.10E-10 | 32.889214 | Serous |

| | | | | |
|---|---|---|---|---|
| 39 | TPD52L1 | 5.28E-10 | 6.914424 | Serous |
| 40 | FOLR1 | 9.96E-10 | 15.970308 | Serous |
| 41 | SCGB3A2 | 9.96E-10 | 67.09675 | Serous |
| 42 | PRB3 | 9.97E-10 | 319.51346 | Serous |
| 43 | CCL28 | 1.79E-09 | 10.505794 | Serous |
| 44 | KRT19 | 2.74E-09 | 5.203251 | Serous |
| 45 | PPAP2C | 3.98E-09 | 5.4127946 | Serous |
| 46 | KRT18 | 3.98E-09 | 9.111273 | Serous |
| 47 | P4HB | 3.98E-09 | 9.203633 | Serous |
| 48 | KCNN4 | 3.98E-09 | 6.481852 | Serous |
| 49 | SFTPC | 4.13E-09 | -170.396 | Serous |
| 50 | FKBP2 | 5.90E-09 | 8.5895815 | Serous |
| 51 | SPINT2 | 9.11E-09 | 8.640325 | Serous |
| 52 | LPO | 9.35E-09 | 19.196758 | Serous |
| 53 | KRT10 | 9.87E-09 | 6.740754 | Serous |
| 54 | PHB | 1.61E-08 | 5.7573338 | Serous |
| 55 | FAM107B | 1.81E-08 | 6.5099897 | Serous |
| 56 | SERP1 | 2.02E-08 | 12.806824 | Serous |
| 57 | GATM | 2.20E-08 | 6.771651 | Serous |
| 58 | TMEM123 | 2.33E-08 | 5.803997 | Serous |
| 59 | OAT | 2.34E-08 | 4.320297 | Serous |
| 60 | LRRC75A-AS1 | 2.66E-08 | 14.784373 | Serous |
| 61 | TPT1 | 2.97E-08 | 109.1202 | Serous |
| 62 | PLTP | 3.06E-08 | 5.762111 | Serous |
| 63 | PABPC1 | 4.52E-08 | 14.34897 | Serous |
| 64 | NUPR1 | 4.52E-08 | 11.327218 | Serous |
| 65 | MZT2A | 5.44E-08 | 3.6494827 | Serous |
| 66 | PPP1R16A | 5.44E-08 | 4.624621 | Serous |
| 67 | CLU | 5.44E-08 | 32.407402 | Serous |
| 68 | MIA | 8.00E-08 | 11.263785 | Serous |
| 69 | TSPAN13 | 1.47E-07 | 7.6609607 | Serous |
| 70 | FKBP11 | 2.13E-07 | 7.031736 | Serous |
| 71 | WFDC21P | 2.37E-07 | 5.1867185 | Serous |
| 72 | PPDPF | 2.51E-07 | 21.712238 | Serous |
| 73 | DANCR | 2.51E-07 | 4.544731 | Serous |
| 74 | RPL12 | 2.62E-07 | 47.595886 | Serous |
| 75 | HERPUD1 | 2.62E-07 | 6.7430515 | Serous |
| 76 | MSMB | 2.86E-07 | 3.9553957 | Serous |
| 77 | RPL13 | 3.07E-07 | 87.23278 | Serous |
| 78 | NPDC1 | 3.39E-07 | 4.753155 | Serous |
| 79 | KRT8 | 3.97E-07 | 5.431861 | Serous |
| 80 | SFTPA1 | 4.26E-07 | -12.625215 | Serous |
| 81 | EPCAM | 4.38E-07 | 5.748682 | Serous |
| 82 | RPL13A | 4.94E-07 | 73.753586 | Serous |
| 83 | LCN2 | 5.13E-07 | 14.918333 | Serous |
| 84 | CDC42EP5 | 5.26E-07 | 5.1830745 | Serous |
| 85 | PHGDH | 5.66E-07 | 5.3264055 | Serous |
| 86 | RPS6 | 5.67E-07 | 52.908604 | Serous |
| 87 | SCGB1A1 | 6.28E-07 | -64.43106 | Serous |
| 88 | RPL36 | 6.46E-07 | 34.935337 | Serous |
| 89 | SMIM22 | 6.48E-07 | 4.3239255 | Serous |
| 90 | RPL10 | 7.18E-07 | 91.926735 | Serous |
| 91 | RPS12 | 7.61E-07 | 58.300625 | Serous |
| 92 | GMDS | 8.07E-07 | 6.2563453 | Serous |
| 93 | PERP | 8.63E-07 | 2.6451855 | Serous |
| 94 | SEC61B | 9.76E-07 | 9.880792 | Serous |
| 95 | RPL15 | 1.02E-06 | 49.774567 | Serous |
| 96 | RPL8 | 1.14E-06 | 44.178722 | Serous |
| 97 | CLDN10 | 1.15E-06 | 6.4183755 | Serous |
| 98 | INSR | 1.27E-06 | 3.8468432 | Serous |
| 99 | PAIP2B | 1.27E-06 | 6.5040326 | Serous |
| 0 | SFTPC | 0 | inf | Signaling Alveolar Epithelial Type 2 |
| 1 | SFTPA2 | 0 | 112.23028 | Signaling Alveolar Epithelial Type 2 |
| 2 | SFTPA1 | 0 | 104.543655 | Signaling Alveolar Epithelial Type 2 |

| | | | | |
|---|---|---|---|---|
| 3 | NAPSA | 0 | 27.143358 | Signaling Alveolar Epithelial Type 2 |
| 4 | SFTPD | 0 | 19.123281 | Signaling Alveolar Epithelial Type 2 |
| 5 | SFTPB | 0 | 63.368046 | Signaling Alveolar Epithelial Type 2 |
| 6 | SFTA2 | 0 | 10.366522 | Signaling Alveolar Epithelial Type 2 |
| 7 | SLC34A2 | 0 | 9.054954 | Signaling Alveolar Epithelial Type 2 |
| 8 | MUC1 | 0 | 8.586664 | Signaling Alveolar Epithelial Type 2 |
| 9 | SLPI | 5.60E-307 | 44.81461 | Signaling Alveolar Epithelial Type 2 |
| 10 | PGC | 3.54E-302 | 13.9768915 | Signaling Alveolar Epithelial Type 2 |
| 11 | AK1 | 1.41E-271 | 7.106877 | Signaling Alveolar Epithelial Type 2 |
| 12 | CXCL17 | 4.30E-265 | 6.519976 | Signaling Alveolar Epithelial Type 2 |
| 13 | MALL | 5.28E-236 | 4.829883 | Signaling Alveolar Epithelial Type 2 |
| 14 | SELENBP1 | 4.02E-233 | 4.7591295 | Signaling Alveolar Epithelial Type 2 |
| 15 | SFTA3 | 4.18E-227 | 4.6591783 | Signaling Alveolar Epithelial Type 2 |
| 16 | LAMP3 | 1.59E-226 | 4.302264 | Signaling Alveolar Epithelial Type 2 |
| 17 | PEBP4 | 1.23E-225 | 4.669559 | Signaling Alveolar Epithelial Type 2 |
| 18 | PIGR | 6.81E-225 | 4.076779 | Signaling Alveolar Epithelial Type 2 |
| 19 | LRRK2 | 1.69E-214 | 4.491867 | Signaling Alveolar Epithelial Type 2 |
| 20 | ABCA3 | 2.70E-213 | 4.7109323 | Signaling Alveolar Epithelial Type 2 |
| 21 | C16orf89 | 4.73E-212 | 4.221341 | Signaling Alveolar Epithelial Type 2 |
| 22 | HOPX | 1.30E-210 | 6.195725 | Signaling Alveolar Epithelial Type 2 |
| 23 | CYB5A | 1.38E-196 | 10.797457 | Signaling Alveolar Epithelial Type 2 |
| 24 | RNASE1 | 3.30E-185 | 8.9382515 | Signaling Alveolar Epithelial Type 2 |
| 25 | S100A14 | 5.42E-184 | 3.8341227 | Signaling Alveolar Epithelial Type 2 |
| 26 | NPC2 | 1.10E-183 | 18.925169 | Signaling Alveolar Epithelial Type 2 |
| 27 | C4BPA | 5.11E-181 | 3.7879913 | Signaling Alveolar Epithelial Type 2 |
| 28 | C3 | 2.44E-179 | 2.9591618 | Signaling Alveolar Epithelial Type 2 |
| 29 | AGR3 | 1.14E-173 | 3.162028 | Signaling Alveolar Epithelial Type 2 |
| 30 | CLDN18 | 8.55E-173 | 3.7686086 | Signaling Alveolar Epithelial Type 2 |
| 31 | SCGB3A2 | 1.92E-161 | 1.2057267 | Signaling Alveolar Epithelial Type 2 |
| 32 | LPCAT1 | 6.06E-154 | 3.7342896 | Signaling Alveolar Epithelial Type 2 |
| 33 | FOLR1 | 3.76E-141 | 2.6413023 | Signaling Alveolar Epithelial Type 2 |
| 34 | FXYD3 | 4.05E-140 | 1.6690223 | Signaling Alveolar Epithelial Type 2 |
| 35 | AGR2 | 2.81E-126 | 1.6434231 | Signaling Alveolar Epithelial Type 2 |
| 36 | WFDC2 | 9.65E-126 | -2.8889587 | Signaling Alveolar Epithelial Type 2 |
| 37 | TPPP3 | 1.61E-122 | 2.3855026 | Signaling Alveolar Epithelial Type 2 |
| 38 | PRDX5 | 3.50E-117 | 3.3835952 | Signaling Alveolar Epithelial Type 2 |
| 39 | MGST1 | 7.21E-105 | 1.9556646 | Signaling Alveolar Epithelial Type 2 |
| 40 | FGGY | 7.10E-103 | 3.151953 | Signaling Alveolar Epithelial Type 2 |
| 41 | ELF3 | 1.48E-101 | 1.3539866 | Signaling Alveolar Epithelial Type 2 |
| 42 | ADIRF | 1.22E-100 | 2.7426105 | Signaling Alveolar Epithelial Type 2 |
| 43 | C8orf4 | 1.06E-99 | 3.4289 | Signaling Alveolar Epithelial Type 2 |
| 44 | TSTD1 | 3.07E-99 | 2.1433558 | Signaling Alveolar Epithelial Type 2 |
| 45 | RGS16 | 3.24E-99 | 2.9466987 | Signaling Alveolar Epithelial Type 2 |
| 46 | FASN | 4.09E-99 | 3.643479 | Signaling Alveolar Epithelial Type 2 |
| 47 | DHCR24 | 4.32E-99 | 2.4509623 | Signaling Alveolar Epithelial Type 2 |
| 48 | PLA2G1B | 7.62E-98 | 3.7928324 | Signaling Alveolar Epithelial Type 2 |
| 49 | SEPP1 | 2.03E-97 | 1.6030669 | Signaling Alveolar Epithelial Type 2 |
| 50 | NGFRAP1 | 9.65E-94 | 2.3526697 | Signaling Alveolar Epithelial Type 2 |
| 51 | KRT8 | 3.05E-92 | 0.58534086 | Signaling Alveolar Epithelial Type 2 |
| 52 | TMEM125 | 6.31E-92 | 2.909309 | Signaling Alveolar Epithelial Type 2 |
| 53 | CHIAP2 | 4.87E-90 | 3.7595239 | Signaling Alveolar Epithelial Type 2 |
| 54 | XIST | 3.72E-88 | 1.8582503 | Signaling Alveolar Epithelial Type 2 |
| 55 | WIF1 | 1.14E-85 | 2.4268913 | Signaling Alveolar Epithelial Type 2 |
| 56 | C11orf96 | 1.27E-85 | 2.0926373 | Signaling Alveolar Epithelial Type 2 |
| 57 | NKX2-1 | 1.73E-84 | 2.735001 | Signaling Alveolar Epithelial Type 2 |
| 58 | SLC22A31 | 4.80E-84 | 3.1696784 | Signaling Alveolar Epithelial Type 2 |
| 59 | CACNA2D2 | 2.67E-83 | 3.5824175 | Signaling Alveolar Epithelial Type 2 |
| 60 | MID1IP1 | 1.01E-82 | 2.458459 | Signaling Alveolar Epithelial Type 2 |
| 61 | CLDN4 | 2.52E-82 | 1.3606087 | Signaling Alveolar Epithelial Type 2 |
| 62 | KRT18 | 3.13E-82 | 0.29694837 | Signaling Alveolar Epithelial Type 2 |
| 63 | CTSH | 1.12E-81 | 2.4799762 | Signaling Alveolar Epithelial Type 2 |
| 64 | GKN2 | 4.22E-81 | 3.4322343 | Signaling Alveolar Epithelial Type 2 |
| 65 | HSD17B6 | 1.96E-80 | 2.995894 | Signaling Alveolar Epithelial Type 2 |
| 66 | SLC39A8 | 3.35E-74 | 1.968086 | Signaling Alveolar Epithelial Type 2 |

| | | | | |
|---|---|---|---|---|
| 67 | PMM1 | 4.43E-73 | 2.683626 | Signaling Alveolar Epithelial Type 2 |
| 68 | FABP5 | 1.60E-72 | 1.5579767 | Signaling Alveolar Epithelial Type 2 |
| 69 | CEBPD | 1.37E-69 | 2.1322706 | Signaling Alveolar Epithelial Type 2 |
| 70 | CLIC3 | 6.32E-69 | 2.0003588 | Signaling Alveolar Epithelial Type 2 |
| 71 | TSC22D1 | 1.40E-68 | 2.2420704 | Signaling Alveolar Epithelial Type 2 |
| 72 | CRTAC1 | 1.43E-68 | 2.7657483 | Signaling Alveolar Epithelial Type 2 |
| 73 | ALPL | 2.49E-68 | 3.1242058 | Signaling Alveolar Epithelial Type 2 |
| 74 | CXCL2 | 6.24E-68 | 2.0493743 | Signaling Alveolar Epithelial Type 2 |
| 75 | DCXR | 1.05E-66 | 1.9911155 | Signaling Alveolar Epithelial Type 2 |
| 76 | STARD10 | 2.71E-65 | 2.3305783 | Signaling Alveolar Epithelial Type 2 |
| 77 | SCGB3A1 | 4.51E-65 | -67.00975 | Signaling Alveolar Epithelial Type 2 |
| 78 | NNMT | 1.70E-63 | 1.9527296 | Signaling Alveolar Epithelial Type 2 |
| 79 | SDC4 | 3.34E-62 | 0.8722406 | Signaling Alveolar Epithelial Type 2 |
| 80 | KRT19 | 8.51E-62 | -2.2136867 | Signaling Alveolar Epithelial Type 2 |
| 81 | STEAP4 | 3.39E-60 | 1.76896 | Signaling Alveolar Epithelial Type 2 |
| 82 | GSTP1 | 9.73E-59 | 1.0101087 | Signaling Alveolar Epithelial Type 2 |
| 83 | CRNDE | 2.91E-58 | 1.9111553 | Signaling Alveolar Epithelial Type 2 |
| 84 | DBI | 3.88E-58 | 2.6564121 | Signaling Alveolar Epithelial Type 2 |
| 85 | CA2 | 1.62E-57 | 1.6256227 | Signaling Alveolar Epithelial Type 2 |
| 86 | EPHX1 | 2.90E-57 | 1.5798213 | Signaling Alveolar Epithelial Type 2 |
| 87 | LGALSL | 3.69E-57 | 3.171669 | Signaling Alveolar Epithelial Type 2 |
| 88 | ETV5 | 5.27E-56 | 3.391778 | Signaling Alveolar Epithelial Type 2 |
| 89 | MRPL14 | 1.51E-55 | 1.4997156 | Signaling Alveolar Epithelial Type 2 |
| 90 | SDR16C5 | 4.10E-55 | 1.8663528 | Signaling Alveolar Epithelial Type 2 |
| 91 | NRGN | 1.39E-54 | 2.518319 | Signaling Alveolar Epithelial Type 2 |
| 92 | MFSD2A | 7.21E-54 | 2.0939577 | Signaling Alveolar Epithelial Type 2 |
| 93 | EPCAM | 1.51E-53 | 1.200414 | Signaling Alveolar Epithelial Type 2 |
| 94 | GAS5 | 1.77E-52 | 1.9406866 | Signaling Alveolar Epithelial Type 2 |
| 95 | F3 | 2.91E-52 | 1.2410415 | Signaling Alveolar Epithelial Type 2 |
| 96 | SECISBP2L | 5.47E-51 | 1.9268101 | Signaling Alveolar Epithelial Type 2 |
| 97 | CHI3L2 | 1.19E-50 | 2.1750212 | Signaling Alveolar Epithelial Type 2 |
| 98 | MAL2 | 3.32E-49 | 1.8466514 | Signaling Alveolar Epithelial Type 2 |
| 99 | C14orf1 | 1.63E-47 | 1.8913025 | Signaling Alveolar Epithelial Type 2 |
| 0 | TYMP | 4.01E-75 | 27.45624 | TREM2+ Dendritic |
| 1 | HLA-DQA1 | 1.82E-73 | 82.69439 | TREM2+ Dendritic |
| 2 | APOE | 5.38E-72 | 479.29114 | TREM2+ Dendritic |
| 3 | CTSB | 2.34E-70 | 51.97067 | TREM2+ Dendritic |
| 4 | HLA-DMB | 2.34E-70 | 28.007824 | TREM2+ Dendritic |
| 5 | LIPA | 1.81E-69 | 37.305676 | TREM2+ Dendritic |
| 6 | CTSZ | 3.17E-69 | 30.99467 | TREM2+ Dendritic |
| 7 | LILRB4 | 6.84E-69 | 17.115866 | TREM2+ Dendritic |
| 8 | TREM2 | 1.84E-68 | 15.052346 | TREM2+ Dendritic |
| 9 | HLA-DPB1 | 3.23E-68 | 120.37938 | TREM2+ Dendritic |
| 10 | CAPG | 5.88E-67 | 54.06115 | TREM2+ Dendritic |
| 11 | PSAP | 1.09E-64 | 108.62237 | TREM2+ Dendritic |
| 12 | HLA-DPA1 | 1.49E-64 | 121.618286 | TREM2+ Dendritic |
| 13 | LYZ | 1.63E-64 | 243.7055 | TREM2+ Dendritic |
| 14 | HLA-DMA | 2.43E-64 | 32.364277 | TREM2+ Dendritic |
| 15 | C15orf48 | 4.58E-64 | 26.781013 | TREM2+ Dendritic |
| 16 | HLA-DQB1 | 5.57E-64 | 48.425327 | TREM2+ Dendritic |
| 17 | CYBB | 9.20E-64 | 9.901893 | TREM2+ Dendritic |
| 18 | HLA-DRA | 1.14E-63 | 422.69373 | TREM2+ Dendritic |
| 19 | GPX1 | 1.41E-63 | 37.017097 | TREM2+ Dendritic |
| 20 | CD74 | 1.90E-63 | 478.81372 | TREM2+ Dendritic |
| 21 | CPM | 1.70E-62 | 11.127137 | TREM2+ Dendritic |
| 22 | CTSS | 1.58E-61 | 71.94013 | TREM2+ Dendritic |
| 23 | BRI3 | 6.82E-61 | 16.83466 | TREM2+ Dendritic |
| 24 | NEAT1 | 1.94E-60 | 29.764927 | TREM2+ Dendritic |
| 25 | CD68 | 1.31E-59 | 35.1392 | TREM2+ Dendritic |
| 26 | HLA-DRB1 | 2.95E-59 | 179.48196 | TREM2+ Dendritic |
| 27 | HLA-DRB5 | 5.70E-59 | 61.186123 | TREM2+ Dendritic |
| 28 | ACP5 | 1.88E-58 | 37.444233 | TREM2+ Dendritic |
| 29 | CD44 | 3.07E-58 | 15.765824 | TREM2+ Dendritic |
| 30 | MAN2B1 | 3.22E-58 | 7.084414 | TREM2+ Dendritic |

| | | | | |
|---|---|---|---|---|
| 31 | IFI30 | 3.33E-58 | 65.078026 | TREM2+ Dendritic |
| 32 | GRN | 6.34E-58 | 60.959515 | TREM2+ Dendritic |
| 33 | ATP6V1F | 6.34E-58 | 17.114063 | TREM2+ Dendritic |
| 34 | MS4A6A | 1.21E-57 | 11.598249 | TREM2+ Dendritic |
| 35 | ITGB2 | 1.59E-57 | 18.555925 | TREM2+ Dendritic |
| 36 | VIM | 2.29E-57 | 107.76992 | TREM2+ Dendritic |
| 37 | MGAT1 | 2.58E-57 | 6.457455 | TREM2+ Dendritic |
| 38 | MAFB | 2.59E-57 | 6.0701113 | TREM2+ Dendritic |
| 39 | TNFSF13B | 2.59E-57 | 7.939632 | TREM2+ Dendritic |
| 40 | SH3BGRL3 | 4.43E-57 | 46.229736 | TREM2+ Dendritic |
| 41 | FTH1 | 4.55E-57 | inf | TREM2+ Dendritic |
| 42 | DBI | 4.87E-57 | 24.738632 | TREM2+ Dendritic |
| 43 | HLA-DRB6 | 1.95E-56 | 64.5567 | TREM2+ Dendritic |
| 44 | CYBA | 5.13E-56 | 62.05595 | TREM2+ Dendritic |
| 45 | GABARAP | 5.56E-56 | 21.90446 | TREM2+ Dendritic |
| 46 | TMEM176B | 9.13E-56 | 20.226122 | TREM2+ Dendritic |
| 47 | GPNMB | 1.42E-55 | 36.947613 | TREM2+ Dendritic |
| 48 | GPX4 | 1.42E-55 | 25.258318 | TREM2+ Dendritic |
| 49 | CHCHD10 | 2.94E-55 | 8.802825 | TREM2+ Dendritic |
| 50 | PRDX1 | 3.23E-55 | 42.78193 | TREM2+ Dendritic |
| 51 | RGS10 | 5.25E-55 | 7.3999333 | TREM2+ Dendritic |
| 52 | APOC1 | 5.92E-55 | 176.84413 | TREM2+ Dendritic |
| 53 | ARPC1B | 6.79E-55 | 20.168005 | TREM2+ Dendritic |
| 54 | TXN | 2.19E-54 | 58.76875 | TREM2+ Dendritic |
| 55 | CTSD | 2.30E-54 | 84.97677 | TREM2+ Dendritic |
| 56 | CTSH | 4.77E-54 | 21.325594 | TREM2+ Dendritic |
| 57 | CSTB | 6.55E-54 | 58.8154 | TREM2+ Dendritic |
| 58 | FTL | 7.98E-54 | inf | TREM2+ Dendritic |
| 59 | LGALS1 | 9.59E-54 | 44.730957 | TREM2+ Dendritic |
| 60 | TMSB10 | 9.59E-54 | 127.11059 | TREM2+ Dendritic |
| 61 | H2AFY | 3.60E-53 | 6.6561747 | TREM2+ Dendritic |
| 62 | YBX1 | 7.78E-53 | 31.324045 | TREM2+ Dendritic |
| 63 | RNH1 | 1.20E-52 | 9.490065 | TREM2+ Dendritic |
| 64 | CCL18 | 1.76E-52 | 257.52032 | TREM2+ Dendritic |
| 65 | COX4I1 | 2.31E-52 | 20.239779 | TREM2+ Dendritic |
| 66 | LGMN | 4.19E-52 | 16.12854 | TREM2+ Dendritic |
| 67 | PILRA | 4.47E-52 | 7.1968365 | TREM2+ Dendritic |
| 68 | HLA-DQA2 | 6.36E-52 | 14.241804 | TREM2+ Dendritic |
| 69 | NPC2 | 8.34E-52 | 29.678177 | TREM2+ Dendritic |
| 70 | RNASE6 | 9.42E-52 | 5.6645875 | TREM2+ Dendritic |
| 71 | KLF6 | 9.62E-52 | 11.533436 | TREM2+ Dendritic |
| 72 | HEXB | 1.44E-51 | 13.798172 | TREM2+ Dendritic |
| 73 | CST3 | 1.64E-51 | 66.41771 | TREM2+ Dendritic |
| 74 | FYB | 2.08E-51 | 6.1698008 | TREM2+ Dendritic |
| 75 | TYROBP | 3.32E-51 | 61.085175 | TREM2+ Dendritic |
| 76 | AP2S1 | 3.57E-51 | 9.3140955 | TREM2+ Dendritic |
| 77 | AKR1A1 | 5.44E-51 | 7.742849 | TREM2+ Dendritic |
| 78 | PPT1 | 5.55E-51 | 8.068626 | TREM2+ Dendritic |
| 79 | LIMS1 | 6.04E-51 | 5.7223225 | TREM2+ Dendritic |
| 80 | RNF130 | 1.72E-50 | 7.747623 | TREM2+ Dendritic |
| 81 | ANXA2 | 2.36E-50 | 41.992035 | TREM2+ Dendritic |
| 82 | ZFAND5 | 2.78E-50 | 6.007375 | TREM2+ Dendritic |
| 83 | CD63 | 2.82E-50 | 72.95121 | TREM2+ Dendritic |
| 84 | ATOX1 | 2.85E-50 | 10.075736 | TREM2+ Dendritic |
| 85 | TMEM176A | 4.36E-50 | 9.35616 | TREM2+ Dendritic |
| 86 | SPI1 | 4.71E-50 | 12.428985 | TREM2+ Dendritic |
| 87 | SLAMF8 | 5.41E-50 | 6.6818156 | TREM2+ Dendritic |
| 88 | FUOM | 6.83E-50 | 4.8184657 | TREM2+ Dendritic |
| 89 | LY96 | 1.03E-49 | 6.115143 | TREM2+ Dendritic |
| 90 | SNX10 | 1.21E-49 | 9.712013 | TREM2+ Dendritic |
| 91 | AP1S2 | 1.79E-49 | 4.718705 | TREM2+ Dendritic |
| 92 | FPR3 | 1.92E-49 | 5.692519 | TREM2+ Dendritic |
| 93 | C1QC | 2.21E-49 | 53.146442 | TREM2+ Dendritic |
| 94 | ACTB | 3.20E-49 | 133.04834 | TREM2+ Dendritic |

| | | | | |
|---|---|---|---|---|
| 95 | PKM | 3.96E-49 | 16.32219 | TREM2+ Dendritic |
| 96 | TSPO | 4.56E-49 | 24.612005 | TREM2+ Dendritic |
| 97 | LAPTM5 | 4.61E-49 | 14.956222 | TREM2+ Dendritic |
| 98 | C1QB | 4.85E-49 | 120.028564 | TREM2+ Dendritic |
| 99 | SERF2 | 7.10E-49 | 38.043667 | TREM2+ Dendritic |
| 0 | ACTA2 | 2.16E-286 | 68.74844 | Vascular Smooth Muscle |
| 1 | TAGLN | 8.83E-285 | 66.28073 | Vascular Smooth Muscle |
| 2 | MYL9 | 1.17E-280 | 30.19646 | Vascular Smooth Muscle |
| 3 | TPM2 | 1.10E-276 | 22.262514 | Vascular Smooth Muscle |
| 4 | IGFBP7 | 5.56E-257 | 55.949917 | Vascular Smooth Muscle |
| 5 | SOD3 | 4.92E-250 | 12.7629595 | Vascular Smooth Muscle |
| 6 | CALD1 | 1.57E-244 | 16.577765 | Vascular Smooth Muscle |
| 7 | PPP1R14A | 1.74E-239 | 10.445327 | Vascular Smooth Muscle |
| 8 | BGN | 3.66E-230 | 12.606791 | Vascular Smooth Muscle |
| 9 | C2orf40 | 5.86E-207 | 15.999933 | Vascular Smooth Muscle |
| 10 | MFAP4 | 3.45E-206 | 11.382636 | Vascular Smooth Muscle |
| 11 | PLAC9 | 2.87E-202 | 7.195744 | Vascular Smooth Muscle |
| 12 | TPM1 | 1.73E-195 | 9.276076 | Vascular Smooth Muscle |
| 13 | TINAGL1 | 2.99E-185 | 6.967405 | Vascular Smooth Muscle |
| 14 | DCN | 2.65E-183 | 7.4142413 | Vascular Smooth Muscle |
| 15 | COL6A2 | 2.73E-182 | 7.729218 | Vascular Smooth Muscle |
| 16 | RARRES2 | 2.73E-182 | 5.915589 | Vascular Smooth Muscle |
| 17 | MGP | 7.27E-171 | 12.096676 | Vascular Smooth Muscle |
| 18 | PRKCDBP | 2.25E-167 | 5.582733 | Vascular Smooth Muscle |
| 19 | SDC2 | 2.09E-161 | 6.8750844 | Vascular Smooth Muscle |
| 20 | MFGE8 | 2.32E-159 | 5.4449525 | Vascular Smooth Muscle |
| 21 | EGFL6 | 2.19E-149 | 7.6507506 | Vascular Smooth Muscle |
| 22 | FXYD1 | 2.51E-149 | 6.229778 | Vascular Smooth Muscle |
| 23 | TGFB1I1 | 1.20E-147 | 5.450703 | Vascular Smooth Muscle |
| 24 | CRYAB | 1.01E-145 | 6.5898876 | Vascular Smooth Muscle |
| 25 | CAV1 | 1.29E-143 | 9.688255 | Vascular Smooth Muscle |
| 26 | AEBP1 | 6.43E-142 | 6.390928 | Vascular Smooth Muscle |
| 27 | SELM | 9.04E-140 | 5.438269 | Vascular Smooth Muscle |
| 28 | MYH11 | 1.16E-139 | 6.6263027 | Vascular Smooth Muscle |
| 29 | PRELP | 8.55E-139 | 5.151012 | Vascular Smooth Muscle |
| 30 | ID4 | 1.71E-136 | 7.5739455 | Vascular Smooth Muscle |
| 31 | SPARCL1 | 1.87E-135 | 5.6000257 | Vascular Smooth Muscle |
| 32 | COX4I2 | 5.29E-134 | 6.7571883 | Vascular Smooth Muscle |
| 33 | 07-Sep | 6.41E-130 | 6.224836 | Vascular Smooth Muscle |
| 34 | MYLK | 2.64E-129 | 4.937727 | Vascular Smooth Muscle |
| 35 | CTGF | 1.44E-128 | 14.944083 | Vascular Smooth Muscle |
| 36 | MAP1B | 4.14E-126 | 5.9171205 | Vascular Smooth Muscle |
| 37 | CNN1 | 3.77E-125 | 7.370187 | Vascular Smooth Muscle |
| 38 | ADIRF | 2.85E-121 | 7.3383937 | Vascular Smooth Muscle |
| 39 | NEXN | 4.84E-121 | 5.472949 | Vascular Smooth Muscle |
| 40 | DSTN | 6.61E-121 | 9.584266 | Vascular Smooth Muscle |
| 41 | TPPP3 | 1.11E-119 | 2.7237923 | Vascular Smooth Muscle |
| 42 | COL1A2 | 6.84E-119 | 6.344363 | Vascular Smooth Muscle |
| 43 | LMCD1 | 2.40E-118 | 5.3072343 | Vascular Smooth Muscle |
| 44 | NTN4 | 4.99E-118 | 5.692952 | Vascular Smooth Muscle |
| 45 | ACTG2 | 1.21E-117 | 8.203274 | Vascular Smooth Muscle |
| 46 | C1S | 2.76E-116 | 4.9461875 | Vascular Smooth Muscle |
| 47 | LMOD1 | 6.57E-111 | 6.1289167 | Vascular Smooth Muscle |
| 48 | ANXA6 | 1.09E-110 | 3.9665933 | Vascular Smooth Muscle |
| 49 | TCF21 | 3.11E-110 | 4.586201 | Vascular Smooth Muscle |
| 50 | ITM2C | 1.18E-108 | 3.8324554 | Vascular Smooth Muscle |
| 51 | COL6A1 | 4.10E-106 | 4.870023 | Vascular Smooth Muscle |
| 52 | ISYNA1 | 8.33E-103 | 5.049591 | Vascular Smooth Muscle |
| 53 | FILIP1L | 5.57E-102 | 4.6791935 | Vascular Smooth Muscle |
| 54 | IGFBP5 | 5.71E-101 | 7.8929553 | Vascular Smooth Muscle |
| 55 | AOC3 | 1.67E-100 | 3.9866407 | Vascular Smooth Muscle |
| 56 | CSRP1 | 1.50E-99 | 4.8330564 | Vascular Smooth Muscle |
| 57 | RRAD | 2.65E-99 | 4.1664634 | Vascular Smooth Muscle |
| 58 | PTRF | 8.83E-99 | 4.0215344 | Vascular Smooth Muscle |

| | | | | |
|---|---|---|---|---|
| 59 | NOTCH3 | 2.62E-98 | 5.3727503 | Vascular Smooth Muscle |
| 60 | LHFP | 4.11E-98 | 4.0623884 | Vascular Smooth Muscle |
| 61 | PDGFRB | 9.92E-98 | 4.4512486 | Vascular Smooth Muscle |
| 62 | GPX3 | 1.89E-97 | 7.78191 | Vascular Smooth Muscle |
| 63 | SPARC | 1.29E-96 | 4.501275 | Vascular Smooth Muscle |
| 64 | FRZB | 1.34E-96 | 6.354815 | Vascular Smooth Muscle |
| 65 | PDLIM7 | 2.55E-95 | 3.8960798 | Vascular Smooth Muscle |
| 66 | TM4SF1 | 3.43E-94 | 6.4212394 | Vascular Smooth Muscle |
| 67 | ADH1B | 9.72E-94 | 4.7393985 | Vascular Smooth Muscle |
| 68 | NR2F2 | 2.41E-91 | 4.5239644 | Vascular Smooth Muscle |
| 69 | MRGPRF | 7.76E-91 | 6.411738 | Vascular Smooth Muscle |
| 70 | HES4 | 8.72E-90 | 3.5648656 | Vascular Smooth Muscle |
| 71 | HSPA2 | 1.82E-89 | 5.6571016 | Vascular Smooth Muscle |
| 72 | GEM | 5.45E-89 | 4.9591575 | Vascular Smooth Muscle |
| 73 | PMP22 | 7.03E-88 | 3.1751287 | Vascular Smooth Muscle |
| 74 | PDLIM3 | 8.14E-86 | 4.279234 | Vascular Smooth Muscle |
| 75 | GUCY1A3 | 1.84E-85 | 5.2902203 | Vascular Smooth Muscle |
| 76 | WFDC1 | 1.11E-82 | 4.6723638 | Vascular Smooth Muscle |
| 77 | C1R | 1.60E-80 | 3.7790449 | Vascular Smooth Muscle |
| 78 | NDN | 2.91E-80 | 3.89722 | Vascular Smooth Muscle |
| 79 | CNN3 | 5.56E-80 | 3.2357981 | Vascular Smooth Muscle |
| 80 | PCOLCE | 4.65E-79 | 3.7543912 | Vascular Smooth Muscle |
| 81 | PLS3 | 7.67E-78 | 3.5900595 | Vascular Smooth Muscle |
| 82 | CRIP2 | 1.43E-77 | 2.6873734 | Vascular Smooth Muscle |
| 83 | THY1 | 1.46E-76 | 6.2508645 | Vascular Smooth Muscle |
| 84 | PRSS23 | 6.58E-76 | 3.5692985 | Vascular Smooth Muscle |
| 85 | COL1A1 | 4.21E-75 | 4.174053 | Vascular Smooth Muscle |
| 86 | ADAMTS1 | 1.62E-74 | 4.4650397 | Vascular Smooth Muscle |
| 87 | OLFML2B | 7.78E-74 | 6.241108 | Vascular Smooth Muscle |
| 88 | NDRG2 | 1.11E-73 | 3.1438053 | Vascular Smooth Muscle |
| 89 | ITGB1 | 3.58E-73 | 3.7090008 | Vascular Smooth Muscle |
| 90 | CRIM1 | 2.61E-72 | 3.786499 | Vascular Smooth Muscle |
| 91 | FOXS1 | 8.48E-72 | 6.740092 | Vascular Smooth Muscle |
| 92 | TIMP3 | 1.05E-71 | 3.4864936 | Vascular Smooth Muscle |
| 93 | LBH | 1.36E-71 | 3.1290228 | Vascular Smooth Muscle |
| 94 | CYR61 | 1.72E-71 | 5.749265 | Vascular Smooth Muscle |
| 95 | ACTN1 | 2.40E-71 | 2.9046328 | Vascular Smooth Muscle |
| 96 | CAV2 | 9.18E-71 | 2.815513 | Vascular Smooth Muscle |
| 97 | VASN | 3.25E-70 | 4.4181156 | Vascular Smooth Muscle |
| 98 | DKK3 | 3.98E-70 | 4.018252 | Vascular Smooth Muscle |
| 99 | GAS6 | 4.78E-70 | 2.7879944 | Vascular Smooth Muscle |
| 0 | ACKR1 | 0 | 35.58131 | Vein |
| 26 | CTNNAL1 | 0 | 4.399978 | Vein |
| 25 | RGS5 | 0 | 5.8789673 | Vein |
| 23 | FAM107A | 0 | 5.958948 | Vein |
| 22 | HYAL2 | 0 | 8.373361 | Vein |
| 21 | RNASE1 | 0 | 8.48297 | Vein |
| 20 | C7 | 0 | 6.52673 | Vein |
| 19 | IFITM3 | 0 | 15.313325 | Vein |
| 18 | CALCRL | 0 | 5.083821 | Vein |
| 17 | GPX3 | 0 | 9.469779 | Vein |
| 16 | TIMP3 | 0 | 14.551763 | Vein |
| 15 | EPAS1 | 0 | 10.588028 | Vein |
| 14 | SRPX | 0 | 5.8804736 | Vein |
| 24 | PCAT19 | 0 | 5.2696023 | Vein |
| 12 | PTPRB | 0 | 5.771452 | Vein |
| 1 | PRSS23 | 0 | 12.385643 | Vein |
| 2 | VWF | 0 | 14.267983 | Vein |
| 13 | CPE | 0 | 7.9207706 | Vein |
| 3 | CLU | 0 | 18.179504 | Vein |
| 5 | RAMP3 | 0 | 7.0048223 | Vein |
| 4 | IGFBP7 | 0 | 18.646185 | Vein |
| 7 | NNMT | 0 | 8.200702 | Vein |
| 8 | CLEC3B | 0 | 9.829462 | Vein |

| | | | | |
|---|---|---|---|---|
| 9 | SLCO2A1 | 0 | 6.6698017 | Vein |
| 10 | LIFR | 0 | 6.0733056 | Vein |
| 11 | PTGDS | 0 | 8.390159 | Vein |
| 6 | MGP | 0 | 18.328318 | Vein |
| 27 | RAMP2 | 2.71E-305 | 6.077506 | Vein |
| 28 | NPDC1 | 1.90E-299 | 4.053273 | Vein |
| 29 | TM4SF1 | 1.49E-295 | 13.628463 | Vein |
| 30 | IL33 | 4.16E-290 | 4.655494 | Vein |
| 31 | SDPR | 2.77E-286 | 7.4797416 | Vein |
| 32 | PLAT | 6.14E-286 | 6.4266615 | Vein |
| 33 | MT1X | 2.54E-274 | 36.075024 | Vein |
| 34 | MT1M | 2.55E-267 | 15.653169 | Vein |
| 35 | PLA1A | 4.55E-264 | 6.2270193 | Vein |
| 36 | CRIP2 | 2.16E-262 | 4.10004 | Vein |
| 37 | IFI27 | 1.78E-261 | 11.438392 | Vein |
| 38 | IL1R1 | 3.28E-257 | 4.64986 | Vein |
| 39 | CD93 | 2.39E-256 | 4.5012856 | Vein |
| 40 | PRCP | 4.67E-256 | 4.039346 | Vein |
| 41 | AQP1 | 5.30E-251 | 5.090811 | Vein |
| 42 | PTGIS | 3.54E-243 | 6.0616364 | Vein |
| 43 | TFPI | 4.24E-243 | 3.4927619 | Vein |
| 44 | CLEC14A | 8.15E-243 | 3.7538538 | Vein |
| 45 | MT2A | 1.23E-234 | 87.97682 | Vein |
| 46 | FCN3 | 1.69E-234 | 14.398841 | Vein |
| 47 | EGFL7 | 2.30E-230 | 4.6459594 | Vein |
| 48 | SOX7 | 1.35E-228 | 5.4108934 | Vein |
| 49 | SPARCL1 | 7.61E-226 | 5.591408 | Vein |
| 50 | IFITM1 | 1.17E-224 | 5.4523687 | Vein |
| 51 | CAV1 | 1.40E-224 | 5.2564344 | Vein |
| 52 | A2M | 3.31E-222 | 3.117967 | Vein |
| 53 | C10orf10 | 1.05E-220 | 8.019437 | Vein |
| 54 | GNG11 | 2.08E-214 | 3.2893667 | Vein |
| 55 | CD59 | 1.62E-213 | 8.51202 | Vein |
| 56 | SELP | 8.66E-212 | 6.017651 | Vein |
| 57 | PTRF | 2.85E-211 | 3.5091255 | Vein |
| 58 | CLDN5 | 1.89E-210 | 8.407784 | Vein |
| 59 | JAM2 | 5.07E-206 | 3.725047 | Vein |
| 60 | PECAM1 | 2.37E-203 | 4.5436916 | Vein |
| 61 | VCAM1 | 3.06E-197 | 8.373555 | Vein |
| 62 | ADIRF | 1.13E-196 | 5.443283 | Vein |
| 63 | TGM2 | 1.11E-195 | 3.3354619 | Vein |
| 64 | MT1E | 7.26E-194 | 12.524458 | Vein |
| 65 | SOCS3 | 2.29E-192 | 6.2566094 | Vein |
| 66 | CEBPD | 3.29E-192 | 7.79127 | Vein |
| 67 | CYP1B1 | 7.14E-184 | 6.3727427 | Vein |
| 68 | TSPAN7 | 6.15E-182 | 3.4143424 | Vein |
| 69 | ABI3BP | 1.25E-180 | 4.862981 | Vein |
| 70 | IGFBP4 | 5.42E-174 | 3.2014868 | Vein |
| 71 | MMRN1 | 1.14E-171 | 5.768254 | Vein |
| 72 | IL6ST | 1.46E-168 | 3.217956 | Vein |
| 73 | ENG | 1.27E-165 | 3.1463146 | Vein |
| 74 | ECSCR | 4.27E-164 | 2.8084855 | Vein |
| 75 | LYVE1 | 5.78E-164 | 4.764095 | Vein |
| 76 | IFITM2 | 4.16E-162 | 4.620211 | Vein |
| 77 | ADGRG6 | 2.89E-158 | 4.882856 | Vein |
| 78 | BMPR2 | 1.02E-157 | 3.1059384 | Vein |
| 79 | TINAGL1 | 3.14E-155 | 2.8034942 | Vein |
| 80 | STXBP6 | 2.24E-154 | 3.0069387 | Vein |
| 81 | ASRGL1 | 1.64E-151 | 3.2438278 | Vein |
| 82 | MT1A | 2.19E-149 | 8.840399 | Vein |
| 83 | NOSTRIN | 1.41E-145 | 3.0022848 | Vein |
| 84 | SPTBN1 | 1.22E-143 | 2.8614767 | Vein |
| 85 | EDN1 | 1.27E-139 | 5.862754 | Vein |
| 86 | LRRC32 | 4.51E-139 | 3.3548603 | Vein |

| | | | | |
|---|---|---|---|---|
| 87 | MMRN2 | 1.49E-138 | 3.3553476 | Vein |
| 88 | FLT1 | 1.72E-137 | 3.699339 | Vein |
| 89 | GSN | 3.51E-134 | 2.995926 | Vein |
| 90 | LTC4S | 6.81E-134 | 2.7232416 | Vein |
| 91 | TIMP1 | 1.28E-133 | 16.173962 | Vein |
| 92 | ADAMTS9 | 9.49E-130 | 5.790848 | Vein |
| 93 | CXCL2 | 3.28E-129 | 12.964409 | Vein |
| 94 | CRIM1 | 5.60E-128 | 3.516158 | Vein |
| 95 | HEG1 | 5.16E-125 | 3.5207748 | Vein |
| 96 | TEK | 1.39E-124 | 3.7709835 | Vein |
| 97 | THBD | 1.39E-124 | 2.8391263 | Vein |
| 98 | S1PR1 | 6.87E-123 | 2.980877 | Vein |
| 99 | C8orf4 | 7.71E-123 | 2.8722944 | Vein |
