## Supplementary Table 5 for "CellDART: Cell type inference by domain adaptation of single-cell and spatial transcriptomic data"

| **Table 5** | The cell types presenting highly different cell fraction in the lung tissue domain of interest compared to other domains. The corresponding cell types were ranked by the ratio of average min-max scaled cell fraction in the domain of interest (a) to the average min-max scaled cell fraction in the rest of the tissue domain (b). The result with Benjamini-Hochberg adjusted p-value below 0.05 were selected. |
|---|---|

**Lung tissue 1**

| Cell types | Tissue domain of interest | Adjusted p-values | Ratio (a/b) | Average scaled cell fraction in the tissue domain (a) | Average scaled cell fraction in the rest of tissue domains (b) |
|---|---|---|---|---|---|
| B | ImmuneCluster | 0.000915494 | 12.10914421 | 0.237038821 | 0.019575192 |
| Proximal Basal | Terminal_Bronchiole | 6.06E-05 | 9.543188095 | 0.384605825 | 0.04030161 |
| Proximal Basal | Bronchial_Epithelium | 4.92E-07 | 8.550522804 | 0.341571778 | 0.039947473 |
| Airway Smooth Muscle | Unknown_Stroma | 0.002532364 | 5.239579678 | 0.168814957 | 0.032219179 |
| Classical Monocyte | Bronchial_Epithelium | 4.92E-07 | 5.169167042 | 0.523485422 | 0.10127075 |
| Proximal Ciliated | Terminal_Bronchiole | 6.06E-05 | 4.734353542 | 0.50654912 | 0.106994361 |
| Proximal Ciliated | Bronchial_Epithelium | 4.83E-07 | 4.493933201 | 0.478300065 | 0.106432393 |
| OLR1+ Classical Monocyte | ImmuneCluster | 3.79E-05 | 4.302912712 | 0.244210988 | 0.056754809 |
| Myeloid Dendritic Type 1 | ImmuneCluster | 3.79E-05 | 4.185704708 | 0.219830424 | 0.052519336 |
| Vascular Smooth Muscle | Unknown_Stroma | 0.008196763 | 4.013421059 | 0.191762552 | 0.04778032 |
| EREG+ Dendritic | ImmuneCluster | 3.79E-05 | 3.995280504 | 0.324188381 | 0.081142835 |
| Plasma | Fibrous_Stroma | 2.02E-21 | 3.980188847 | 0.02565255 | 0.006445059 |
| Differentiating Basal | Terminal_Bronchiole | 0.000263469 | 3.800572872 | 0.197914779 | 0.052074987 |
| Differentiating Basal | Bronchial_Epithelium | 9.59E-06 | 3.789741755 | 0.196342841 | 0.051809028 |
| Plasmacytoid Dendritic | Fibrous_Stroma | 1.86E-22 | 3.567270279 | 0.027871385 | 0.007813085 |
| CD4+ Naive T | ImmuneCluster | 3.79E-05 | 3.416904688 | 0.337835848 | 0.098871894 |
| Nonclassical Monocyte | Terminal_Bronchiole | 0.000116853 | 3.370207071 | 0.278105795 | 0.082518905 |
| Classical Monocyte | Terminal_Bronchiole | 6.06E-05 | 3.329625607 | 0.34289664 | 0.102983542 |
| Club | Bronchial_Epithelium | 4.34E-06 | 3.284182549 | 0.292885631 | 0.089180678 |
| Intermediate Monocyte | ImmuneCluster | 0.000125658 | 3.257628918 | 0.13739866 | 0.042177506 |
| Nonclassical Monocyte | Bronchial_Epithelium | 2.88E-06 | 3.067627668 | 0.252543986 | 0.082325503 |
| Plasmacytoid Dendritic | ImmuneCluster | 0.000440992 | 3.033862352 | 0.056839075 | 0.018734889 |
| Mucous | Bronchial_Epithelium | 7.26E-05 | 2.978192806 | 0.097475499 | 0.032729749 |
| Platelet/Megakaryocyte | ImmuneCluster | 0.00037789 | 2.9362607 | 0.278955787 | 0.095003754 |
| Myeloid Dendritic Type 2 | ImmuneCluster | 0.000907403 | 2.854470968 | 0.201335654 | 0.07053344 |
| CD8+ Memory/Effector T | ImmuneCluster | 0.011625581 | 2.816564083 | 0.166363478 | 0.059066106 |
| Proximal Basal | Unknown_Stroma | 0.022358283 | 2.748407364 | 0.111717552 | 0.040648106 |
| Ciliated | Terminal_Bronchiole | 0.000199935 | 2.67619133 | 0.155464381 | 0.058091655 |
| Natural Killer | Bronchial_Epithelium | 6.88E-07 | 2.632586479 | 0.472270519 | 0.179394111 |
| Lymphatic | Fibrous_Stroma | 1.96E-09 | 2.458773851 | 0.018444041 | 0.007501316 |
| Macrophage | Bronchial_Epithelium | 1.47E-06 | 2.447047234 | 0.622727156 | 0.254481047 |
| Natural Killer | Terminal_Bronchiole | 0.000162986 | 2.444812298 | 0.440336466 | 0.180110544 |
| Fibromyocyte | Fibrous_Stroma | 8.36E-18 | 2.444451094 | 0.028441923 | 0.011635301 |
| Basophil/Mast 2 | ImmuneCluster | 0.000465886 | 2.411924839 | 0.142713845 | 0.059170105 |
| Mucous | Terminal_Bronchiole | 0.036301968 | 2.346082211 | 0.077313595 | 0.032954343 |
| Classical Monocyte | ImmuneCluster | 0.000129029 | 2.333250523 | 0.240481302 | 0.103067078 |
| Mesothelial | Fibrous_Stroma | 5.90E-23 | 2.332616329 | 0.092140213 | 0.039500799 |
| Ciliated | Bronchial_Epithelium | 0.00163826 | 2.293075323 | 0.133140981 | 0.058062192 |
| Club | Terminal_Bronchiole | 0.004699659 | 2.28663063 | 0.205812231 | 0.090006769 |
| Neuroendocrine | Fibrous_Stroma | 7.68E-28 | 2.25043273 | 0.107418023 | 0.047732163 |
| Fibromyocyte | Unknown_Stroma | 0.018891864 | 2.241523981 | 0.046064008 | 0.020550307 |
| IGSF21+ Dendritic | ImmuneCluster | 0.006859671 | 2.236110926 | 0.101469129 | 0.0453775 |
| Macrophage | Terminal_Bronchiole | 0.000192866 | 2.207215548 | 0.563891172 | 0.255476266 |
| Basophil/Mast 1 | ImmuneCluster | 0.000752654 | 2.183227301 | 0.142059118 | 0.065068401 |
| Differentiating Basal | Unknown_Stroma | 0.002532364 | 1.993701935 | 0.103284255 | 0.051805265 |
| Plasma | ImmuneCluster | 0.039606796 | 1.944273353 | 0.033188805 | 0.017070031 |
| Proximal Basal | ImmuneCluster | 0.001606612 | 1.941615224 | 0.080988541 | 0.041711941 |
| Myofibroblast | Fibrous_Stroma | 1.92E-21 | 1.926610947 | 0.036891442 | 0.019148361 |
| Nonclassical Monocyte | ImmuneCluster | 0.047305843 | 1.918529868 | 0.159008414 | 0.082880341 |
| Basal | Terminal_Bronchiole | 0.027642724 | 1.839063287 | 0.045975395 | 0.024999354 |
| CD4+ Naive T | Terminal_Bronchiole | 0.002368512 | 1.796958089 | 0.180454195 | 0.10042204 |
| Serous | Fibrous_Stroma | 1.55E-17 | 1.755307317 | 0.05860801 | 0.033389032 |
| TREM2+ Dendritic | ImmuneCluster | 0.002801543 | 1.694133401 | 0.279601187 | 0.165040836 |
| Intermediate Monocyte | Bronchial_Epithelium | 0.00163826 | 1.690521002 | 0.072272219 | 0.042751446 |
| Lipofibroblast | Fibrous_Stroma | 1.09E-10 | 1.64502573 | 0.071932182 | 0.043727085 |
| IGSF21+ Dendritic | Fibrous_Stroma | 9.43E-14 | 1.642819762 | 0.055367842 | 0.033702932 |
| EREG+ Dendritic | Fibrous_Stroma | 2.66E-16 | 1.609831333 | 0.099754892 | 0.06196605 |
| Adventitial Fibroblast | Fibrous_Stroma | 1.89E-07 | 1.584110141 | 0.118435293 | 0.074764557 |
| Basophil/Mast 2 | Fibrous_Stroma | 9.62E-11 | 1.572480083 | 0.071268372 | 0.045322273 |
| Airway Smooth Muscle | Fibrous_Stroma | 3.03E-07 | 1.567026615 | 0.041485339 | 0.026473921 |
| Natural Killer T | ImmuneCluster | 0.007048575 | 1.537902594 | 0.259808928 | 0.168937176 |
| Bronchial Vessel 1 | Fibrous_Stroma | 9.40E-09 | 1.520473361 | 0.061403345 | 0.04038436 |
| Vascular Smooth Muscle | Fibrous_Stroma | 9.36E-08 | 1.478051901 | 0.058978569 | 0.039902907 |
| Alveolar Epithelial Type 1 | Alveolar_Space_Capillary_Pneumocytes | 7.12E-10 | 1.475520492 | 0.056958985 | 0.038602639 |
| Capillary Aerocyte | Unknown | 0.01597972 | 1.432793736 | 0.131588459 | 0.091840476 |
| OLR1+ Classical Monocyte | Fibrous_Stroma | 3.92E-15 | 1.431779504 | 0.067203201 | 0.046936836 |
| B | Bronchial_Epithelium | 0.007397428 | 1.407913208 | 0.029977329 | 0.021292029 |
| Myeloid Dendritic Type 1 | Fibrous_Stroma | 1.56E-08 | 1.386788845 | 0.061423961 | 0.044292223 |
| Goblet | Fibrous_Stroma | 1.24E-07 | 1.381444931 | 0.090243116 | 0.065325163 |
| CD8+ Memory/Effector T | Bronchial_Epithelium | 0.007022879 | 1.379276991 | 0.082461528 | 0.059786055 |
| Proliferating Macrophage | Fibrous_Stroma | 1.96E-11 | 1.376450658 | 0.114434838 | 0.083137624 |
| Club | Unknown_Stroma | 0.018891864 | 1.362112761 | 0.122528031 | 0.089954399 |
| CD4+ Naive T | Bronchial_Epithelium | 0.045096124 | 1.345650196 | 0.135350987 | 0.100584082 |
| Proliferating NK/T | Fibrous_Stroma | 1.45E-10 | 1.318174243 | 0.050654214 | 0.038427554 |
| Alveolar Epithelial Type 2 | Alveolar_Space_Capillary_Pneumocytes | 1.45E-15 | 1.307768703 | 0.291957438 | 0.223248526 |
| Capillary Aerocyte | Alveolar_Space_Capillary_Pneumocytes | 7.28E-08 | 1.298945904 | 0.111941256 | 0.086178534 |
| Signaling Alveolar Epithelial Type 2 | Alveolar_Space_Capillary_Pneumocytes | 1.46E-09 | 1.290697575 | 0.138493538 | 0.10730131 |
| Platelet/Megakaryocyte | Fibrous_Stroma | 2.38E-07 | 1.275096297 | 0.106628247 | 0.083623685 |
| Capillary | Alveolar_Space_Capillary_Pneumocytes | 7.56E-15 | 1.252241969 | 0.236551508 | 0.188902393 |
| Basophil/Mast 1 | Fibrous_Stroma | 1.88E-05 | 1.250320673 | 0.072043456 | 0.057619981 |
| Capillary | Unknown | 0.009216363 | 1.230944037 | 0.247781634 | 0.20129399 |
| Capillary Intermediate 1 | Unknown | 0.01597972 | 1.225989461 | 0.221945658 | 0.181033909 |
| Natural Killer T | Fibrous_Stroma | 9.40E-09 | 1.223153591 | 0.18448849 | 0.150830194 |
| Capillary Intermediate 2 | Alveolar_Space_Capillary_Pneumocytes | 3.03E-07 | 1.223005772 | 0.08707498 | 0.071197525 |
| Mucous | Fibrous_Stroma | 1.25E-09 | 1.222027063 | 0.036059953 | 0.029508309 |
| Myeloid Dendritic Type 2 | Fibrous_Stroma | 0.001516973 | 1.212451577 | 0.077583484 | 0.063988931 |
| Mucous | Unknown_Stroma | 0.033663238 | 1.210781217 | 0.040006261 | 0.033041693 |
| Capillary Intermediate 1 | Alveolar_Space_Capillary_Pneumocytes | 1.46E-09 | 1.203413248 | 0.207456902 | 0.172390401 |
| CD8+ Naive T | Alveolar_Space_Capillary_Pneumocytes | 7.74E-09 | 1.189662099 | 0.160693243 | 0.13507469 |
| TREM2+ Dendritic | Fibrous_Stroma | 1.80E-06 | 1.177646399 | 0.17777583 | 0.150958583 |
| Ciliated | Alveolar_Space_Capillary_Pneumocytes | 3.88E-06 | 1.151467204 | 0.06424927 | 0.055797741 |
| Bronchial Vessel 2 | Fibrous_Stroma | 0.012090219 | 1.148283601 | 0.102019839 | 0.088845506 |
| Ionocyte | Fibrous_Stroma | 0.000224504 | 1.144553542 | 0.111251436 | 0.097200729 |
| B | Fibrous_Stroma | 3.91E-06 | 1.143601298 | 0.022600709 | 0.019762751 |
| Alveolar Fibroblast | Alveolar_Space_Capillary_Pneumocytes | 0.008077472 | 1.118638515 | 0.164742246 | 0.147270307 |
| Proliferating Basal | Fibrous_Stroma | 0.001156921 | 1.105084538 | 0.067887194 | 0.061431676 |
| Macrophage | Alveolar_Space_Capillary_Pneumocytes | 0.013573291 | 1.094022632 | 0.272747368 | 0.249306872 |
| Mucous | Unknown | 0.047828919 | 1.066132665 | 0.035206139 | 0.033022288 |
| Nonclassical Monocyte | Alveolar_Space_Capillary_Pneumocytes | 0.00092462 | 1.045039177 | 0.085987732 | 0.082281828 |
| CD4+ Memory/Effector T | Alveolar_Space_Capillary_Pneumocytes | 0.0278817 | 1.042314291 | 0.076126002 | 0.073035553 |
| Intermediate Monocyte | Fibrous_Stroma | 0.003243905 | 1.037217379 | 0.043644577 | 0.042078525 |
| Basal | Alveolar_Space_Capillary_Pneumocytes | 9.20E-06 | 0.988923669 | 0.024917658 | 0.025196746 |
| Classical Monocyte | Fibrous_Stroma | 0.020371789 | 0.920075834 | 0.100354418 | 0.109071895 |
| Classical Monocyte | Alveolar_Space_Capillary_Pneumocytes | 0.003166843 | 0.912844121 | 0.097919919 | 0.107269049 |
| Proximal Ciliated | Alveolar_Space_Capillary_Pneumocytes | 0.033429434 | 0.912255466 | 0.102398179 | 0.112247266 |
| Basal | Fibrous_Stroma | 7.43E-09 | 0.906883478 | 0.024019724 | 0.026486009 |
| Intermediate Monocyte | Alveolar_Space_Capillary_Pneumocytes | 0.000725102 | 0.894135118 | 0.039795615 | 0.044507384 |
| Natural Killer | Fibrous_Stroma | 0.003346598 | 0.885701537 | 0.171671823 | 0.193825811 |
| TREM2+ Dendritic | Alveolar_Space_Capillary_Pneumocytes | 0.002373343 | 0.884715617 | 0.152637914 | 0.172527656 |
| Natural Killer T | Alveolar_Space_Capillary_Pneumocytes | 0.000471076 | 0.874990761 | 0.154843375 | 0.176965728 |
| Macrophage | Fibrous_Stroma | 0.00011013 | 0.864781737 | 0.240483031 | 0.278085232 |
| Differentiating Basal | Alveolar_Space_Capillary_Pneumocytes | 0.036724171 | 0.863020539 | 0.047727883 | 0.055303298 |
| CD4+ Naive T | Fibrous_Stroma | 6.21E-05 | 0.861543953 | 0.094166249 | 0.109299414 |
| Capillary Intermediate 1 | Fibrous_Stroma | 1.14E-06 | 0.85649991 | 0.171311498 | 0.200013444 |
| Ciliated | Fibrous_Stroma | 3.49E-08 | 0.842957139 | 0.054143295 | 0.064230189 |
| Alveolar Epithelial Type 2 | Fibrous_Stroma | 2.06E-07 | 0.8389799 | 0.226732343 | 0.270247638 |
| Nonclassical Monocyte | Fibrous_Stroma | 4.81E-05 | 0.838475943 | 0.076977968 | 0.091807008 |
| Bronchial Vessel 2 | Alveolar_Space_Capillary_Pneumocytes | 0.002686868 | 0.837888062 | 0.085171998 | 0.101650804 |
| Proliferating Basal | Alveolar_Space_Capillary_Pneumocytes | 0.000309606 | 0.834346116 | 0.057403505 | 0.068800591 |
| CD8+ Naive T | Fibrous_Stroma | 2.65E-08 | 0.826854646 | 0.131405607 | 0.158922255 |
| Capillary Intermediate 2 | Fibrous_Stroma | 2.74E-08 | 0.82341373 | 0.06983728 | 0.084814325 |
| Basophil/Mast 1 | Alveolar_Space_Capillary_Pneumocytes | 0.002243645 | 0.821133494 | 0.057323337 | 0.06981001 |
| Myeloid Dendritic Type 2 | Alveolar_Space_Capillary_Pneumocytes | 0.019498111 | 0.818989813 | 0.062359102 | 0.076141484 |
| Differentiating Basal | Unknown | 0.029869373 | 0.807184219 | 0.043217894 | 0.053541549 |
| Intermediate Monocyte | Unknown | 0.026601753 | 0.802745283 | 0.034972325 | 0.043565907 |
| Signaling Alveolar Epithelial Type 2 | Fibrous_Stroma | 2.06E-07 | 0.802642763 | 0.1060967 | 0.132184207 |
| Capillary | Fibrous_Stroma | 2.84E-14 | 0.800724685 | 0.184404224 | 0.230296656 |
| CD8+ Memory/Effector T | Fibrous_Stroma | 0.01922045 | 0.792786837 | 0.053760499 | 0.067812048 |
| Alveolar Epithelial Type 1 | Fibrous_Stroma | 1.07E-05 | 0.787042737 | 0.039898515 | 0.050694216 |

**Lung tissue 2**

| Cell types | Tissue domain of interest | Adjusted p-values | Ratio (a/b) | Average scaled cell fraction in the tissue domain (a) | Average scaled cell fraction in the rest of tissue domains (b) |
|---|---|---|---|---|---|
| Ciliated | Terminal_Bronchiole | 0.009178934 | 12.97328091 | 0.247474819 | 0.019075731 |
| Proximal Ciliated | Terminal_Bronchiole | 0.009178934 | 10.59664631 | 0.27209121 | 0.025677105 |
| Club | Terminal_Bronchiole | 0.009178934 | 6.005847454 | 0.391915977 | 0.065255731 |
| Mucous | Terminal_Bronchiole | 0.013126002 | 5.991510391 | 0.243000746 | 0.040557511 |
| B | ImmuneCluster | 3.62E-47 | 5.324261665 | 0.054487679 | 0.010233847 |
| Goblet | Terminal_Bronchiole | 0.030231659 | 4.057803154 | 0.189264208 | 0.046642039 |
| Differentiating Basal | Terminal_Bronchiole | 0.009178934 | 3.468072891 | 0.054719485 | 0.015778067 |
| Plasma | ImmuneCluster | 2.98E-42 | 3.017703772 | 0.037893008 | 0.012556901 |
| Plasmacytoid Dendritic | ImmuneCluster | 9.47E-30 | 2.857616186 | 0.054689344 | 0.0191381 |
| OLR1+ Classical Monocyte | ImmuneCluster | 4.67E-48 | 2.695884228 | 0.048141878 | 0.017857546 |
| Signaling Alveolar Epithelial Type 2 | Alveolar_Space_Capillary_Pneumocytes | 1.74E-133 | 2.665920734 | 0.172733083 | 0.064793028 |
| CD4+ Memory/Effector T | ImmuneCluster | 2.66E-35 | 2.556840181 | 0.091772847 | 0.035893071 |
| Capillary Intermediate 2 | Alveolar_Space_Capillary_Pneumocytes | 1.32E-125 | 2.267900229 | 0.235634506 | 0.103899859 |
| EREG+ Dendritic | ImmuneCluster | 2.79E-35 | 2.172198057 | 0.064692236 | 0.029781925 |
| IGSF21+ Dendritic | ImmuneCluster | 1.24E-22 | 2.167957306 | 0.07187672 | 0.033154123 |
| Alveolar Epithelial Type 1 | Alveolar_Space_Capillary_Pneumocytes | 7.61E-68 | 2.074790239 | 0.094649225 | 0.045618698 |
| Adventitial Fibroblast | Fibrous_Stroma | 5.75E-62 | 2.068246365 | 0.063277923 | 0.030594965 |
| Proliferating NK/T | ImmuneCluster | 1.55E-29 | 2.019854784 | 0.078193665 | 0.038712516 |
| Myeloid Dendritic Type 1 | ImmuneCluster | 2.74E-19 | 1.900189757 | 0.097775139 | 0.05145546 |
| Proximal Basal | Terminal_Bronchiole | 0.030231659 | 1.888171315 | 0.021859046 | 0.011576834 |
| Lipofibroblast | ImmuneCluster | 1.18E-19 | 1.867130518 | 0.101832151 | 0.054539386 |
| Lipofibroblast | Fibrous_Stroma | 1.19E-64 | 1.818737626 | 0.078537077 | 0.043182191 |
| Alveolar Epithelial Type 2 | Alveolar_Space_Capillary_Pneumocytes | 5.51E-90 | 1.798917651 | 0.300969452 | 0.167305857 |
| Vein | Fibrous_Stroma | 1.33E-56 | 1.7955935 | 0.063868418 | 0.03556953 |
| Myeloid Dendritic Type 2 | ImmuneCluster | 1.69E-12 | 1.772127032 | 0.079728432 | 0.044990245 |
| Fibromyocyte | Fibrous_Stroma | 1.12E-59 | 1.737985253 | 0.039334647 | 0.022632325 |
| Mesothelial | Fibrous_Stroma | 8.37E-61 | 1.708923101 | 0.100923531 | 0.0590568 |
| Intermediate Monocyte | ImmuneCluster | 5.16E-19 | 1.705338478 | 0.118690439 | 0.069599345 |
| Lymphatic | Unknown | 0.002423852 | 1.638717175 | 0.031023063 | 0.018931311 |
| Ionocyte | ImmuneCluster | 1.39E-09 | 1.635771155 | 0.085115276 | 0.05203373 |
| Plasma | Unknown | 0.036173691 | 1.599439621 | 0.022136796 | 0.013840345 |
| Mucous | ImmuneCluster | 6.84E-09 | 1.588679314 | 0.061509807 | 0.038717572 |
| Artery | Fibrous_Stroma | 4.47E-51 | 1.581762791 | 0.230599836 | 0.145786613 |
| Natural Killer | Alveolar_Space_Capillary_Pneumocytes | 5.27E-66 | 1.573816419 | 0.228092909 | 0.144929811 |
| Bronchial Vessel 1 | ImmuneCluster | 2.30E-11 | 1.569506764 | 0.087599233 | 0.055813223 |
| Mesothelial | ImmuneCluster | 5.55E-15 | 1.56611383 | 0.115712121 | 0.073884875 |
| B | Unknown | 0.036173691 | 1.473710418 | 0.020072227 | 0.013620198 |
| CD8+ Memory/Effector T | Alveolar_Space_Capillary_Pneumocytes | 2.58E-41 | 1.446207881 | 0.118139282 | 0.081689008 |
| Natural Killer T | ImmuneCluster | 2.51E-16 | 1.440731406 | 0.198585242 | 0.137836412 |
| Myofibroblast | Fibrous_Stroma | 2.84E-33 | 1.439046621 | 0.04416804 | 0.030692572 |
| Nonclassical Monocyte | ImmuneCluster | 2.62E-05 | 1.43511343 | 0.093015276 | 0.064813882 |
| Capillary Aerocyte | Alveolar_Space_Capillary_Pneumocytes | 3.25E-39 | 1.426014066 | 0.213269606 | 0.149556458 |
| Platelet/Megakaryocyte | ImmuneCluster | 4.81E-09 | 1.418375611 | 0.119448692 | 0.084215134 |
| Neuroendocrine | ImmuneCluster | 4.19E-09 | 1.371965647 | 0.161510333 | 0.117721848 |
| Capillary | Alveolar_Space_Capillary_Pneumocytes | 2.15E-59 | 1.355276227 | 0.26881507 | 0.198347077 |
| Basophil/Mast 1 | Fibrous_Stroma | 2.81E-25 | 1.349327207 | 0.103055052 | 0.076375142 |
| Club | Alveolar_Space_Capillary_Pneumocytes | 3.47E-09 | 1.342730522 | 0.080210552 | 0.059736893 |
| Nonclassical Monocyte | Alveolar_Space_Capillary_Pneumocytes | 8.98E-16 | 1.297501922 | 0.08048898 | 0.062033802 |
| TREM2+ Dendritic | ImmuneCluster | 0.000578029 | 1.294934511 | 0.081031784 | 0.062575974 |
| Alveolar Fibroblast | Fibrous_Stroma | 1.22E-23 | 1.29471755 | 0.0797754 | 0.061616063 |
| Proliferating Macrophage | ImmuneCluster | 7.06E-07 | 1.289974928 | 0.067466207 | 0.052300401 |
| Classical Monocyte | ImmuneCluster | 6.87E-06 | 1.281867743 | 0.087154239 | 0.067990042 |
| Classical Monocyte | Fibrous_Stroma | 3.37E-22 | 1.275325656 | 0.079239786 | 0.062132981 |
| Artery | ImmuneCluster | 1.38E-06 | 1.25999856 | 0.226204097 | 0.179527268 |
| OLR1+ Classical Monocyte | Fibrous_Stroma | 1.40E-15 | 1.251359344 | 0.023380188 | 0.018683832 |
| Ciliated | Alveolar_Space_Capillary_Pneumocytes | 2.35E-49 | 1.248906255 | 0.022666808 | 0.018149327 |
| Adventitial Fibroblast | ImmuneCluster | 2.56E-10 | 1.24515748 | 0.055148631 | 0.044290487 |
| Proximal Ciliated | Alveolar_Space_Capillary_Pneumocytes | 1.29E-37 | 1.237912059 | 0.030203799 | 0.024398986 |
| EREG+ Dendritic | Fibrous_Stroma | 2.33E-12 | 1.204170704 | 0.036580846 | 0.030378455 |
| CD8+ Naive T | Alveolar_Space_Capillary_Pneumocytes | 6.75E-13 | 1.175918698 | 0.206773236 | 0.175839737 |
| Basophil/Mast 2 | Alveolar_Space_Capillary_Pneumocytes | 8.43E-06 | 1.168025732 | 0.114575423 | 0.098093234 |
| Capillary Intermediate 1 | Alveolar_Space_Capillary_Pneumocytes | 9.28E-14 | 1.16500175 | 0.364506096 | 0.312880307 |
| Neuroendocrine | Fibrous_Stroma | 1.17E-07 | 1.157700062 | 0.131834164 | 0.113875926 |
| Vein | ImmuneCluster | 0.000181454 | 1.147711992 | 0.054697338 | 0.047657721 |
| CD4+ Memory/Effector T | Fibrous_Stroma | 1.07E-14 | 1.144029021 | 0.04439548 | 0.038806252 |
| CD4+ Naive T | Alveolar_Space_Capillary_Pneumocytes | 7.47E-14 | 1.142274261 | 0.060078565 | 0.052595567 |
| IGSF21+ Dendritic | Fibrous_Stroma | 1.01E-11 | 1.136631012 | 0.039527901 | 0.034776371 |
| Lymphatic | Fibrous_Stroma | 7.74E-22 | 1.134971499 | 0.022094449 | 0.019466965 |
| Bronchial Vessel 1 | Fibrous_Stroma | 1.04E-08 | 1.13254559 | 0.062947117 | 0.055580206 |
| TREM2+ Dendritic | Alveolar_Space_Capillary_Pneumocytes | 3.78E-14 | 1.123720288 | 0.069752552 | 0.062072877 |
| Proliferating Macrophage | Alveolar_Space_Capillary_Pneumocytes | 0.000440111 | 1.121212125 | 0.058183234 | 0.051893152 |
| Pericyte | ImmuneCluster | 0.041773687 | 1.120534062 | 0.108661376 | 0.096972845 |
| Vascular Smooth Muscle | Alveolar_Space_Capillary_Pneumocytes | 9.00E-11 | 1.117769599 | 0.084061809 | 0.075204954 |
| Natural Killer T | Fibrous_Stroma | 4.64E-06 | 1.113829851 | 0.152284473 | 0.136721492 |
| Bronchial Vessel 2 | ImmuneCluster | 0.014225637 | 1.101474762 | 0.117340066 | 0.106529959 |
| Bronchial Vessel 2 | Fibrous_Stroma | 0.000130232 | 1.090930223 | 0.112736985 | 0.103340231 |
| Plasma | Fibrous_Stroma | 3.53E-10 | 1.090083241 | 0.015734984 | 0.014434662 |
| Proximal Basal | Fibrous_Stroma | 0.000714688 | 1.089095354 | 0.012140909 | 0.011147701 |
| Plasmacytoid Dendritic | Fibrous_Stroma | 7.27E-05 | 1.083386779 | 0.023592772 | 0.021776868 |
| Myeloid Dendritic Type 1 | Alveolar_Space_Capillary_Pneumocytes | 5.17E-08 | 1.038159728 | 0.057440851 | 0.05532949 |
| Proximal Basal | Alveolar_Space_Capillary_Pneumocytes | 1.58E-09 | 1.037702084 | 0.011898922 | 0.011466607 |
| Pericyte | Fibrous_Stroma | 0.002087224 | 1.027995944 | 0.09959802 | 0.096885622 |
| Intermediate Monocyte | Fibrous_Stroma | 0.036727153 | 1.02186811 | 0.075258493 | 0.073647954 |
| Airway Smooth Muscle | Alveolar_Space_Capillary_Pneumocytes | 1.06E-07 | 1.021635532 | 0.052264776 | 0.051157944 |
| Macrophage | Fibrous_Stroma | 0.036727153 | 1.017850637 | 0.23307696 | 0.228989348 |
| Differentiating Basal | Alveolar_Space_Capillary_Pneumocytes | 0.004889573 | 1.011337399 | 0.015976131 | 0.015797034 |
| Macrophage | Alveolar_Space_Capillary_Pneumocytes | 0.000316549 | 0.960335553 | 0.224341631 | 0.233607545 |
| Capillary Aerocyte | ImmuneCluster | 0.0063094 | 0.951062739 | 0.161140352 | 0.16943188 |
| Capillary Intermediate 1 | Fibrous_Stroma | 0.002421691 | 0.937227011 | 0.316694111 | 0.337905437 |
| CD8+ Naive T | Fibrous_Stroma | 0.013905681 | 0.93545264 | 0.178346261 | 0.19065237 |
| Proximal Basal | ImmuneCluster | 0.009149453 | 0.930566907 | 0.010864232 | 0.011674853 |
| Proximal Ciliated | ImmuneCluster | 0.040287733 | 0.929250896 | 0.024455512 | 0.026317447 |
| Natural Killer | ImmuneCluster | 0.000762252 | 0.90980792 | 0.155871019 | 0.171323001 |
| Capillary Intermediate 1 | ImmuneCluster | 0.001229174 | 0.909627855 | 0.301308364 | 0.331243545 |
| Myeloid Dendritic Type 2 | Fibrous_Stroma | 0.003512009 | 0.902937412 | 0.045675736 | 0.050585717 |
| Pericyte | Alveolar_Space_Capillary_Pneumocytes | 6.98E-07 | 0.898740768 | 0.090931125 | 0.101176143 |
| Proliferating Basal | Alveolar_Space_Capillary_Pneumocytes | 0.005416695 | 0.895641625 | 0.018219255 | 0.020342126 |
| Platelet/Megakaryocyte | Fibrous_Stroma | 7.50E-08 | 0.875562966 | 0.081303097 | 0.092858084 |
| CD8+ Naive T | ImmuneCluster | 1.06E-05 | 0.873715699 | 0.163734376 | 0.187400058 |
| Bronchial Vessel 2 | Alveolar_Space_Capillary_Pneumocytes | 7.30E-10 | 0.852542996 | 0.095952541 | 0.112548627 |
| Lymphatic | ImmuneCluster | 1.78E-05 | 0.849732399 | 0.017809786 | 0.020959288 |
| Capillary | Fibrous_Stroma | 1.11E-15 | 0.84800452 | 0.199790627 | 0.235600904 |
| Basophil/Mast 2 | ImmuneCluster | 0.003676709 | 0.846743405 | 0.088557042 | 0.104585454 |
| Proximal Ciliated | Fibrous_Stroma | 5.75E-20 | 0.839929998 | 0.02366226 | 0.028171705 |
| Ciliated | ImmuneCluster | 0.001518784 | 0.838534355 | 0.016613524 | 0.019812575 |
| TREM2+ Dendritic | Fibrous_Stroma | 1.31E-18 | 0.83445996 | 0.058052912 | 0.069569439 |
| Ciliated | Fibrous_Stroma | 1.95E-22 | 0.825464904 | 0.017474292 | 0.021169031 |
| Proliferating Macrophage | Fibrous_Stroma | 5.46E-13 | 0.803215265 | 0.047403935 | 0.059017722 |
| CD4+ Naive T | Fibrous_Stroma | 1.74E-12 | 0.786892235 | 0.047741555 | 0.06067102 |
| Capillary Aerocyte | Fibrous_Stroma | 1.02E-15 | 0.784095109 | 0.146494269 | 0.186832264 |
| Ionocyte | Alveolar_Space_Capillary_Pneumocytes | 0.000542982 | 0.778003991 | 0.046045519 | 0.059184168 |
| Basophil/Mast 1 | Alveolar_Space_Capillary_Pneumocytes | 2.04E-18 | 0.767581403 | 0.07294143 | 0.095027618 |
| Alveolar Fibroblast | Alveolar_Space_Capillary_Pneumocytes | 2.48E-20 | 0.760938227 | 0.057216786 | 0.075192422 |
| Myeloid Dendritic Type 1 | Fibrous_Stroma | 7.74E-21 | 0.743066728 | 0.047038034 | 0.063302569 |
| Intermediate Monocyte | Alveolar_Space_Capillary_Pneumocytes | 3.19E-13 | 0.736191571 | 0.059447918 | 0.080750607 |
| Mucous | Alveolar_Space_Capillary_Pneumocytes | 7.09E-10 | 0.727330208 | 0.032418724 | 0.044572223 |
| Club | ImmuneCluster | 1.89E-05 | 0.7247715 | 0.04904997 | 0.067676462 |
| CD8+ Memory/Effector T | Fibrous_Stroma | 2.93E-25 | 0.711524308 | 0.075748928 | 0.106460072 |
| Alveolar Epithelial Type 2 | Fibrous_Stroma | 7.10E-26 | 0.70376116 | 0.168415889 | 0.239308313 |
| Nonclassical Monocyte | Fibrous_Stroma | 2.76E-20 | 0.702212691 | 0.054806031 | 0.078047626 |
| Natural Killer T | Alveolar_Space_Capillary_Pneumocytes | 8.59E-32 | 0.696756244 | 0.110153258 | 0.158094391 |
| Capillary | ImmuneCluster | 2.10E-31 | 0.690341532 | 0.156191885 | 0.226253062 |
| Proliferating NK/T | Alveolar_Space_Capillary_Pneumocytes | 9.94E-19 | 0.689870358 | 0.032359812 | 0.046907093 |
| Natural Killer | Fibrous_Stroma | 1.77E-42 | 0.680719733 | 0.135063827 | 0.198413268 |
| Myofibroblast | Alveolar_Space_Capillary_Pneumocytes | 3.24E-27 | 0.671637058 | 0.027391277 | 0.040782854 |
| Alveolar Epithelial Type 1 | Fibrous_Stroma | 1.59E-20 | 0.661845982 | 0.04709246 | 0.071153201 |
| Classical Monocyte | Alveolar_Space_Capillary_Pneumocytes | 1.07E-34 | 0.659736633 | 0.051310848 | 0.077774748 |
| Lymphatic | Alveolar_Space_Capillary_Pneumocytes | 1.25E-58 | 0.64223659 | 0.014854621 | 0.023129515 |
| Neuroendocrine | Alveolar_Space_Capillary_Pneumocytes | 6.45E-31 | 0.621508479 | 0.085497715 | 0.137564838 |

| | | | | | |
|---|---|---|---|---|---|
| Capillary Aerocyte | Fibrous_Stroma | 4.17E-05 | 0.786444485 | 0.084560804 | 0.107522912 |
| Alveolar Fibroblast | Unknown | 0.012433206 | 0.785180092 | 0.122013599 | 0.155395687 |
| Proliferating NK/T | Alveolar_Space_Capillary_Pneumocytes | 2.40E-08 | 0.778850496 | 0.038032431 | 0.048831493 |
| Platelet/Megakaryocyte | Alveolar_Space_Capillary_Pneumocytes | 1.10E-05 | 0.767448127 | 0.080207169 | 0.104511522 |
| Proliferating Macrophage | Alveolar_Space_Capillary_Pneumocytes | 6.91E-09 | 0.734961867 | 0.0810607 | 0.11029239 |
| Adventitial Fibroblast | Alveolar_Space_Capillary_Pneumocytes | 0.010502712 | 0.717619896 | 0.078504257 | 0.109395318 |
| Classical Monocyte | Unknown | 0.00041666 | 0.713348389 | 0.075868316 | 0.106355205 |
| Myeloid Dendritic Type 1 | Alveolar_Space_Capillary_Pneumocytes | 1.19E-05 | 0.706434011 | 0.04214175 | 0.059654195 |
| Goblet | Alveolar_Space_Capillary_Pneumocytes | 1.91E-07 | 0.702223361 | 0.061725829 | 0.087900564 |
| Mucous | Alveolar_Space_Capillary_Pneumocytes | 8.28E-12 | 0.691150546 | 0.025527211 | 0.036934372 |
| Proximal Basal | Unknown | 0.047828919 | 0.684099019 | 0.029414611 | 0.042997591 |
| Myeloid Dendritic Type 2 | Unknown_Stroma | 0.04665154 | 0.67985487 | 0.048984677 | 0.072051667 |
| Lipofibroblast | Alveolar_Space_Capillary_Pneumocytes | 4.55E-05 | 0.671263516 | 0.044806603 | 0.066749647 |
| Basophil/Mast 2 | Alveolar_Space_Capillary_Pneumocytes | 3.03E-07 | 0.65928489 | 0.044447728 | 0.067418091 |
| IGSF21+ Dendritic | Alveolar_Space_Capillary_Pneumocytes | 2.29E-08 | 0.645901918 | 0.033513948 | 0.051887054 |
| OLR1+ Classical Monocyte | Alveolar_Space_Capillary_Pneumocytes | 5.34E-15 | 0.641661286 | 0.042404801 | 0.066085957 |
| Capillary | Bronchial_Epithelium | 0.034783709 | 0.637808263 | 0.130820453 | 0.205109373 |
| B | Alveolar_Space_Capillary_Pneumocytes | 1.44E-07 | 0.635689616 | 0.015423964 | 0.024263356 |
| Serous | Alveolar_Space_Capillary_Pneumocytes | 6.11E-11 | 0.634516418 | 0.034272857 | 0.054014135 |
| Bronchial Vessel 1 | Alveolar_Space_Capillary_Pneumocytes | 8.23E-10 | 0.62001878 | 0.036963273 | 0.059616376 |
| Alveolar Epithelial Type 2 | ImmuneCluster | 0.021607035 | 0.603604615 | 0.148895353 | 0.246676952 |
| EREG+ Dendritic | Alveolar_Space_Capillary_Pneumocytes | 6.44E-13 | 0.597728789 | 0.057277028 | 0.095824443 |
| Alveolar Fibroblast | Terminal_Bronchiole | 0.030932207 | 0.587802052 | 0.090135068 | 0.15334256 |
| Natural Killer T | Unknown_Stroma | 0.008053324 | 0.584484518 | 0.099984638 | 0.171064645 |
| Vascular Smooth Muscle | Alveolar_Space_Capillary_Pneumocytes | 7.27E-05 | 0.551363289 | 0.032726835 | 0.059356209 |
| Myofibroblast | Alveolar_Space_Capillary_Pneumocytes | 3.77E-14 | 0.534562647 | 0.018362572 | 0.034350645 |
| CD8+ Naive T | Bronchial_Epithelium | 0.001083491 | 0.506157815 | 0.072888702 | 0.144003913 |
| Airway Smooth Muscle | Alveolar_Space_Capillary_Pneumocytes | 7.31E-05 | 0.468952149 | 0.019824767 | 0.042274605 |
| Lymphatic | Alveolar_Space_Capillary_Pneumocytes | 7.54E-05 | 0.460070819 | 0.007626793 | 0.016577434 |
| Neuroendocrine | Alveolar_Space_Capillary_Pneumocytes | 3.85E-21 | 0.458090603 | 0.04525461 | 0.098789647 |
| Myeloid Dendritic Type 1 | Unknown_Stroma | 0.018891864 | 0.452566683 | 0.024650132 | 0.054467402 |
| Mesothelial | Alveolar_Space_Capillary_Pneumocytes | 5.34E-15 | 0.451639354 | 0.038020898 | 0.0841842 |
| CD8+ Naive T | Terminal_Bronchiole | 0.002114291 | 0.445816636 | 0.064158931 | 0.143913269 |
| Basophil/Mast 1 | Terminal_Bronchiole | 0.030696822 | 0.418659598 | 0.027585525 | 0.065890104 |
| Myeloid Dendritic Type 2 | Terminal_Bronchiole | 0.016690189 | 0.403979421 | 0.029012775 | 0.071817458 |
| Capillary | ImmuneCluster | 0.000101794 | 0.391048431 | 0.080406934 | 0.205618858 |
| Capillary Intermediate 1 | Terminal_Bronchiole | 0.002595966 | 0.380554646 | 0.070217803 | 0.184514374 |
| Fibromyocyte | Alveolar_Space_Capillary_Pneumocytes | 1.91E-12 | 0.373792291 | 0.009911559 | 0.02651622 |
| Myeloid Dendritic Type 2 | Bronchial_Epithelium | 0.003812498 | 0.357378393 | 0.025703287 | 0.071921773 |
| Vein | Terminal_Bronchiole | 0.026665077 | 0.348556966 | 0.021315213 | 0.061152741 |
| CD4+ Memory/Effector T | Terminal_Bronchiole | 0.000964377 | 0.332754582 | 0.024724562 | 0.074302696 |
| Platelet/Megakaryocyte | Bronchial_Epithelium | 0.01078554 | 0.328245282 | 0.031825718 | 0.096957125 |
| Adventitial Fibroblast | Terminal_Bronchiole | 0.013256818 | 0.299233884 | 0.029795354 | 0.09957213 |
| Capillary Intermediate 1 | ImmuneCluster | 3.79E-05 | 0.284519672 | 0.052642204 | 0.185021311 |
| Proliferating NK/T | Bronchial_Epithelium | 0.040982976 | 0.275229603 | 0.012523719 | 0.045502804 |
| Plasmacytoid Dendritic | Alveolar_Space_Capillary_Pneumocytes | 1.56E-16 | 0.268490076 | 0.006736784 | 0.025091369 |
| Alveolar Fibroblast | Bronchial_Epithelium | 6.53E-06 | 0.260622293 | 0.040086582 | 0.153811023 |
| Plasma | Alveolar_Space_Capillary_Pneumocytes | 9.64E-16 | 0.251646578 | 0.005744656 | 0.022828268 |
| Pericyte | ImmuneCluster | 0.000403258 | 0.245467499 | 0.031728573 | 0.129257739 |
| Vein | Bronchial_Epithelium | 0.00150159 | 0.242329404 | 0.014848349 | 0.061273407 |
| Capillary Intermediate 1 | Bronchial_Epithelium | 4.81E-06 | 0.242165685 | 0.044778489 | 0.184908479 |
| TREM2+ Dendritic | Terminal_Bronchiole | 0.000199935 | 0.227961212 | 0.037983749 | 0.166623741 |
| EREG+ Dendritic | Bronchial_Epithelium | 0.001235051 | 0.227840781 | 0.019041775 | 0.083574921 |
| Alveolar Epithelial Type 1 | ImmuneCluster | 0.008755252 | 0.216105163 | 0.00971094 | 0.044936176 |
| Bronchial Vessel 1 | Bronchial_Epithelium | 0.000113271 | 0.213972196 | 0.011220811 | 0.052440509 |
| OLR1+ Classical Monocyte | Terminal_Bronchiole | 0.003051104 | 0.189923301 | 0.011115242 | 0.058524899 |
| Adventitial Fibroblast | Bronchial_Epithelium | 4.67E-05 | 0.179541722 | 0.017915988 | 0.099787325 |
| Natural Killer T | Bronchial_Epithelium | 1.46E-06 | 0.178117365 | 0.03039562 | 0.170649394 |
| Proliferating Macrophage | Bronchial_Epithelium | 6.12E-05 | 0.16710268 | 0.016918795 | 0.101247899 |
| TREM2+ Dendritic | Bronchial_Epithelium | 6.88E-07 | 0.16513826 | 0.027568268 | 0.166940525 |
| Alveolar Epithelial Type 2 | Terminal_Bronchiole | 0.000116853 | 0.163654372 | 0.040408939 | 0.246916354 |
| Capillary Aerocyte | ImmuneCluster | 9.37E-05 | 0.154504329 | 0.014727678 | 0.095322102 |
| Natural Killer T | Terminal_Bronchiole | 6.06E-05 | 0.148321196 | 0.025275335 | 0.170409456 |
| Bronchial Vessel 1 | Terminal_Bronchiole | 0.000199935 | 0.143068567 | 0.007494106 | 0.052381221 |
| CD4+ Memory/Effector T | Bronchial_Epithelium | 7.97E-07 | 0.142176837 | 0.010591496 | 0.074495226 |
| EREG+ Dendritic | Terminal_Bronchiole | 0.000485188 | 0.136535391 | 0.011399512 | 0.083491258 |
| Myeloid Dendritic Type 1 | Terminal_Bronchiole | 0.000231749 | 0.13445127 | 0.007276769 | 0.054121979 |
| Capillary Aerocyte | Terminal_Bronchiole | 0.000485057 | 0.124535829 | 0.01184113 | 0.095082112 |
| Proliferating NK/T | Terminal_Bronchiole | 0.002114291 | 0.117076606 | 0.005324257 | 0.04547669 |
| Basophil/Mast 2 | Bronchial_Epithelium | 4.56E-06 | 0.109922692 | 0.006619913 | 0.060223348 |
| Alveolar Epithelial Type 2 | Bronchial_Epithelium | 6.88E-07 | 0.107677132 | 0.026639787 | 0.247404322 |
| Ionocyte | Terminal_Bronchiole | 0.000199935 | 0.1038922 | 0.010965307 | 0.105545051 |
| Basophil/Mast 2 | Terminal_Bronchiole | 9.22E-05 | 0.096344531 | 0.005792806 | 0.060125943 |
| Alveolar Epithelial Type 1 | Terminal_Bronchiole | 0.001004407 | 0.094211332 | 0.004225627 | 0.044852637 |
| IGSF21+ Dendritic | Bronchial_Epithelium | 5.08E-06 | 0.093848802 | 0.004328757 | 0.046124797 |
| Proliferating Macrophage | Terminal_Bronchiole | 6.06E-05 | 0.087006174 | 0.008798858 | 0.101129122 |
| Myeloid Dendritic Type 1 | Bronchial_Epithelium | 1.49E-06 | 0.084069088 | 0.004559054 | 0.054229844 |
| Serous | Bronchial_Epithelium | 2.44E-05 | 0.081816398 | 0.00391212 | 0.047815841 |
| Capillary Aerocyte | Bronchial_Epithelium | 3.01E-06 | 0.07963632 | 0.007586935 | 0.095269784 |
| Pericyte | Terminal_Bronchiole | 6.06E-05 | 0.071983598 | 0.009290425 | 0.129063085 |
| Serous | Terminal_Bronchiole | 0.000192866 | 0.065824442 | 0.00314223 | 0.047736529 |
| IGSF21+ Dendritic | Terminal_Bronchiole | 6.06E-05 | 0.063658826 | 0.002931658 | 0.04605265 |
| Neuroendocrine | Bronchial_Epithelium | 2.88E-06 | 0.06037816 | 0.004932197 | 0.081688426 |
| Pericyte | Bronchial_Epithelium | 4.92E-07 | 0.048897326 | 0.006322971 | 0.129311174 |
| Neuroendocrine | Terminal_Bronchiole | 7.45E-05 | 0.046238717 | 0.003770713 | 0.081548832 |
| Myofibroblast | Terminal_Bronchiole | 6.06E-05 | 0.042947102 | 0.001255154 | 0.029225577 |
| Lipofibroblast | Terminal_Bronchiole | 6.06E-05 | 0.041932005 | 0.002507986 | 0.05981078 |
| Alveolar Epithelial Type 1 | Bronchial_Epithelium | 1.32E-06 | 0.039631516 | 0.001781309 | 0.04494679 |
| Plasma | Terminal_Bronchiole | 0.000199935 | 0.03893584 | 0.000673016 | 0.017285267 |
| Plasma | Bronchial_Epithelium | 5.08E-06 | 0.033060256 | 0.000572521 | 0.017317509 |
| Airway Smooth Muscle | Terminal_Bronchiole | 6.36E-05 | 0.03187092 | 0.001117118 | 0.035051323 |
| Ionocyte | Bronchial_Epithelium | 4.92E-07 | 0.031362396 | 0.003317448 | 0.105777882 |
| Myofibroblast | Bronchial_Epithelium | 4.92E-07 | 0.029261265 | 0.000856812 | 0.029281456 |
| Lymphatic | Terminal_Bronchiole | 0.000116853 | 0.025976794 | 0.000355794 | 0.0136966 |
| Vascular Smooth Muscle | Terminal_Bronchiole | 6.06E-05 | 0.025369463 | 0.001289654 | 0.050834883 |
| Mesothelial | Terminal_Bronchiole | 6.06E-05 | 0.023924546 | 0.00165846 | 0.069320418 |
| Airway Smooth Muscle | Bronchial_Epithelium | 4.92E-07 | 0.019683009 | 0.000691242 | 0.035118718 |
| Mesothelial | Bronchial_Epithelium | 4.92E-07 | 0.018448394 | 0.001281269 | 0.069451518 |
| Fibromyocyte | Terminal_Bronchiole | 6.06E-05 | 0.012388092 | 0.000262039 | 0.021152474 |
| Lipofibroblast | Bronchial_Epithelium | 4.83E-07 | 0.011265309 | 0.000675156 | 0.05993234 |
| Lymphatic | Bronchial_Epithelium | 4.92E-07 | 0.010699533 | 0.000146834 | 0.013723385 |
| Vascular Smooth Muscle | Bronchial_Epithelium | 4.83E-07 | 0.010554298 | 0.000537575 | 0.050934192 |
| Plasmacytoid Dendritic | Bronchial_Epithelium | 6.88E-07 | 0.008304531 | 0.000159262 | 0.019177729 |
| Plasmacytoid Dendritic | Terminal_Bronchiole | 6.06E-05 | 0.007394232 | 0.000141539 | 0.019141776 |
| Fibromyocyte | Bronchial_Epithelium | 4.83E-07 | 0.006046976 | 0.000128154 | 0.021193072 |

| | | | | | |
|---|---|---|---|---|---|
| Bronchial Vessel 1 | Alveolar_Space_Capillary_Pneumocytes | 3.63E-34 | 0.6197474 | 0.041194651 | 0.066470064 |
| Capillary Intermediate 2 | Fibrous_Stroma | 1.29E-54 | 0.565979481 | 0.100883089 | 0.178245127 |
| Alveolar Epithelial Type 2 | ImmuneCluster | 7.92E-25 | 0.545116663 | 0.118240505 | 0.216908619 |
| Signaling Alveolar Epithelial Type 2 | Fibrous_Stroma | 1.12E-46 | 0.507512391 | 0.063363865 | 0.12485186 |
| EREG+ Dendritic | Alveolar_Space_Capillary_Pneumocytes | 9.41E-46 | 0.481320173 | 0.018903786 | 0.039274868 |
| Capillary Intermediate 2 | ImmuneCluster | 8.97E-20 | 0.476506054 | 0.07196077 | 0.151017532 |
| Artery | Alveolar_Space_Capillary_Pneumocytes | 2.25E-84 | 0.473446459 | 0.103449211 | 0.218502447 |
| Vascular Smooth Muscle | ImmuneCluster | 7.85E-34 | 0.441942662 | 0.036382902 | 0.082324937 |
| Fibromyocyte | Alveolar_Space_Capillary_Pneumocytes | 1.82E-86 | 0.426201433 | 0.015525145 | 0.036426779 |
| IGSF21+ Dendritic | Alveolar_Space_Capillary_Pneumocytes | 8.66E-48 | 0.423526585 | 0.018897491 | 0.044619374 |
| CD4+ Memory/Effector T | Alveolar_Space_Capillary_Pneumocytes | 8.66E-68 | 0.42347759 | 0.021151582 | 0.049947347 |
| Alveolar Epithelial Type 1 | ImmuneCluster | 3.18E-21 | 0.38148436 | 0.024473421 | 0.064153142 |
| Airway Smooth Muscle | ImmuneCluster | 2.60E-25 | 0.381264091 | 0.020888252 | 0.054786835 |
| Vein | Alveolar_Space_Capillary_Pneumocytes | 2.72E-96 | 0.345503062 | 0.020771287 | 0.060118966 |
| Signaling Alveolar Epithelial Type 2 | ImmuneCluster | 1.55E-29 | 0.325953931 | 0.033870593 | 0.103912205 |
| Plasmacytoid Dendritic | Alveolar_Space_Capillary_Pneumocytes | 1.38E-36 | 0.316972911 | 0.009002887 | 0.028402701 |
| OLR1+ Classical Monocyte | Alveolar_Space_Capillary_Pneumocytes | 2.54E-75 | 0.314726561 | 0.008236776 | 0.026171213 |
| Macrophage | Terminal_Bronchiole | 0.009178934 | 0.279588133 | 0.0646253 | 0.231144652 |
| Basophil/Mast 1 | Terminal_Bronchiole | 0.016329424 | 0.279240966 | 0.024722969 | 0.088536322 |
| Mesothelial | Alveolar_Space_Capillary_Pneumocytes | 1.99E-123 | 0.277385473 | 0.027590267 | 0.099465437 |
| Adventitial Fibroblast | Alveolar_Space_Capillary_Pneumocytes | 1.83E-102 | 0.275485784 | 0.015951283 | 0.057902381 |
| Bronchial Vessel 2 | Terminal_Bronchiole | 0.009178934 | 0.216452554 | 0.023320356 | 0.107738882 |
| B | Alveolar_Space_Capillary_Pneumocytes | 6.19E-36 | 0.207068384 | 0.003947295 | 0.019062761 |
| Alveolar Fibroblast | Terminal_Bronchiole | 0.016167229 | 0.197117254 | 0.013781863 | 0.069917083 |
| CD4+ Memory/Effector T | Terminal_Bronchiole | 0.010511866 | 0.171256259 | 0.0070888 | 0.041392941 |
| Lipofibroblast | Alveolar_Space_Capillary_Pneumocytes | 2.80E-133 | 0.159070373 | 0.012571443 | 0.0790307 |
| Plasma | Alveolar_Space_Capillary_Pneumocytes | 1.41E-72 | 0.136924922 | 0.002773169 | 0.020253209 |
| Bronchial Vessel 1 | Terminal_Bronchiole | 0.009178934 | 0.129754052 | 0.007655622 | 0.059001025 |
| B | Terminal_Bronchiole | 0.043471957 | 0.115089439 | 0.001676029 | 0.014562839 |
| Vein | Terminal_Bronchiole | 0.043471957 | 0.106321901 | 0.005148444 | 0.048423175 |
| Adventitial Fibroblast | Terminal_Bronchiole | 0.043471957 | 0.090352237 | 0.004104138 | 0.045423761 |
| Lymphatic | Terminal_Bronchiole | 0.009178934 | 0.055357095 | 0.001145301 | 0.020689329 |
